# Phylogenetic shifts in gene body methylation correlate with gene expression and reflect trait conservation

**DOI:** 10.1101/687186

**Authors:** Danelle K. Seymour, Brandon S. Gaut

## Abstract

A subset of genes in plant genomes are labeled with DNA methylation specifically at CG residues. These genes, known as gene-body methylated (gbM), have a number of associated characteristics. They tend to have longer sequences, to be enriched for intermediate expression levels, and to be associated with slower rates of molecular evolution. Most importantly, gbM genes tend to maintain their level of DNA methylation between species, suggesting that this trait is under evolutionary constraint. Given the degree of conservation in gbM, we still know surprisingly little about its function in plant genomes or whether gbM is itself a target of selection. To address these questions, we surveyed DNA methylation across eight grass (Poaceae) species that span a gradient of genome sizes. We first established that genome size correlates with genome-wide DNA methylation levels, but less so for genic levels. We then leveraged genomic data to identify a set of 2,982 putative orthologs among the eight species and examined shifts of methylation status for each ortholog in a phylogenetic context. A total of 55% of orthologs exhibited a shift in gbM, but these shifts occurred predominantly on terminal branches, indicating that shifts in gbM are rarely conveyed over time. Finally, we found that the degree of conservation of gbM across species is associated with increased gene length, reduced rates of molecular evolution, and increased gene expression level, but reduced gene expression variation across species. Overall, these observations suggest a basis for evolutionary pressure to maintain gbM status over evolutionary time.

## INTRODUCTION

DNA methylation is an epigenetic mark with known roles in chromatin packaging, genomic surveillance, and transcriptional regulation. The molecular machinery responsible for its initiation and maintenance are preserved in the genomes of both plants and animals, indicating a deep evolutionary origin (Suzuki and Bird 2008; Feng et al. 2010; Zemach et al. 2010; Huff and Zilberman 2014). Despite this conservation, DNA methylation has been independently lost on multiple occasions, including in the genomes of C*aenorhabditis elegans* and *Drosophila melanogaster (Yi 2017)*). Within plants, profiling of DNA methylation has revealed distinct methylation patterns between transposable elements (TEs) and genes (Feng et al. 2010; Zemach et al. 2010; Niederhuth et al. 2016; Takuno et al. 2016). TEs are typically enriched for DNA methylation in all three nucleotide contexts - i.e, CG, CHG (where H is A|T|C), and CHH (Gruenbaum et al. 1981) - and this enrichment is associated with chromatin compaction and reduced expression of the targeted TE (Slotkin and Martienssen 2007). In contrast to TEs, genic methylation tends to occur in the CG context within transcribed regions, and this gene-body methylation (gbM) is linked to active transcription (Zhang et al. 2006; Zilberman et al. 2007; Cokus et al. 2008; Lister et al. 2008; Lister et al. 2009). Unlike TE methylation, which turns over rapidly, relative levels of CG methylation can be highly conserved within specific genes, in some cases remaining stable over ~400 million years of land plant evolution (Takuno et al. 2016).

Most of the steps necessary to initiate TE methylation are well characterized (Law and Jacobsen 2010), but the molecular mechanism responsible for the initiation of gbM in plant genomes is not fully understood. It is known that the maintenance of CG methylation in genes is due to the action of *MET1* (*DNA METHYLTRANSFERASE 1*) (Finnegan and Dennis 1993; Finnegan et al. 1996; Ronemus et al. 1996), but *MET1* is not sufficient to initiate gbM (Stroud et al. 2013). Instead, a feed-forward regulatory loop may be necessary for initiation. The loop consists of the plant-specific methyltransferase *CMT3* (*CHROMOMETHYLTRANSFERASE 3*) and a histone methyltransferase *KYP* (*KRYPTONITE* or *SUVH4*) (reviewed in (Du et al. 2015)). Three observations are consistent with this feed-forward model: *i*) the deposition of H3K9me2 by *KYP* is recognized by *CMT3*, which then *de novo* methylates CHG trinucleotides (reviewed in (Du et al. 2015)), *ii*) loss of the histone demethylase IBM1 (INCREASED IN BONSAI METHYLATION 1) leads to the accumulation of both H3K9me2 and CHG methylation (Saze et al. 2008; Miura et al. 2009), and *iii*) two Brassicaceae species that lost *CMT3* also lack gbM (Bewick et al. 2016; Niederhuth et al. 2016). However, it is still unclear why epigenetic marks typically associated with heterochromatin (mCHG and H3K9me2) are enriched, albeit transiently, in the body of methylated genes, particularly because genic CHG methylation is associated with suppressed expression in *A. thaliana* (Slotkin and Martienssen 2007). One hypothesis is that CG dinucleotides are spuriously methylated by *CMT3* in this feed-forward regulatory loop (Bewick et al. 2016). However, recent work has shown that methylation of the first cytosine of CCG trinucleotides by CMT3 requires methylation of the second cytosine, suggesting that CG methylation is not spurious but required for the *CMT3/KYP* regulatory loop to function (Zabet et al. 2017).

Another mystery is why some, but not all, genes are methylated. It is nonetheless clear that gbM genes have distinct characteristics. They tend to be longer and contain more exons than unmethylated genes (Takuno and Gaut 2012; Takuno and Gaut 2013; Takuno et al. 2017); they are transcribed at moderate to high levels and across more tissues relative to unmethylated genes [e.g., (Zhang et al. 2006; Zilberman et al. 2007)]; they are enriched for housekeeping functions (Takuno and Gaut 2012; Takuno and Gaut 2013; Takuno et al. 2017); and they evolve more slowly than unmethylated genes, indicative of evolutionary constraint (Takuno and Gaut 2012; Takuno and Gaut 2013; Takuno et al. 2017). This last observation is surprising because DNA methylation catalyzes C to T substitutions (Bird 1980; Pfeifer 2006); hence the conservation of gbM implies that the benefit of maintaining CG methylation outweighs the costs associated with elevated mutation rates.

Despite these general characteristics, genic methylation also has some lineage-specific innovations. For example, gymnosperms tend to be enriched for CG *and* CHG methylation in gene bodies (Takuno et al. 2016). In *A. thaliana*, genic CHG methylation is associated with reduced expression (Slotkin and Martienssen 2007), but *Pinus taeda* genes with CHG methylation appear to exhibit classical gbM characteristics such as moderate levels of gene expression (Takuno et al. 2016). Some lineages also vary in the distribution of CG methylation across the gene body (Feng et al. 2010; Zemach et al. 2010; Niederhuth et al. 2016; Takuno et al. 2016). Angiosperms tend to have enriched CG methylation towards the 3’ end of genes (Feng et al. 2010; Zemach et al. 2010; Niederhuth et al. 2016; Takuno et al. 2016), but ferns and gymnosperms exhibit enriched CG methylation across the entire region (Takuno et al. 2016). Another source of variation among lineages is genome size (GS), because species with larger genomes tend to have higher overall levels of CHG in their genes (Takuno et al. 2016). This observation has, however, been chiefly driven by maize (*Zea mays* ssp. *mays*) as an outlier with the largest GS in some comparative methylomic analyses, which has correspondingly high levels of genic CHG methylation (Gent et al. 2013; Niederhuth et al. 2016; Takuno et al. 2016). Finally, we have already noted another lineage-specific feature, which is that at least two Brassicacae species completely lack gbM (Bewick et al. 2016; Niederhuth et al. 2016).

The absence of gbM in the two Brassicaceae species has been used to argue that gbM is dispensable and thus has no function (Bewick et al. 2016; Bewick et al. 2019). This argument is bolstered by molecular studies showing that the lack of gbM in *A. thaliana met1* mutants has only minor effects on gene expression (Zhang et al. 2006; Zilberman et al. 2007; Cokus et al. 2008; Lister et al. 2008; Bewick et al. 2019). However, the arguments against function are countered by at least four lines of evidence, including *i*) the evolutionary conservation of gbM, *ii*) the fact that loss of a functional trait does not *de facto* imply that a trait is non-functional (Zilberman 2017), *iii*) the possibility that the function of genic methylation has been replaced by a separate epigenetic regulatory marks in Brassicaceae species, as it has in *D. melanogaster (Sentmanat and Elgin 2012)*, and *iv*) observations that shifts in gbM status between lineages are weakly but consistently associated with shifts in gene expression. For example, Takuno *et. al*. (2017) studied gene expression on a small number of orthologs that did not have conserved gbM levels between *A. thaliana* and *A. lyrata (Takuno et al. 2017)*. These orthologs differed more widely in gene expression between lineages than did genes with conserved body-methylation, suggesting the possibility of an indirect link between gbM and gene expression. More recently, Muyle and Gaut (2019) re-investigated the effect of gbM loss in one of the Brassicaceae (*Eutrema selagineum*) that lacks gbM, and they found that the loss of gbM was associated with small but significant decrease in gene expression (Muyle and Gaut 2019).

Overall, the function of gbM in plant genomes remains elusive despite extensive investigation from both mechanistic and evolutionary perspectives. It is likely, however, that further methylome analyses will reveal additional lineage-specific phenomena that provide further insight into the evolution and function of methylation marks. It is also surprising to realize that gbM has not been studied in an explicit phylogenetic context. Such a context could prove valuable for inferring both the evolutionary forces that act on gbM, if there are any, and its potential relationship to gene expression. Imagine, for example, a multi-species phylogeny based on an orthologous gene, for which gbM status is known for all species. With this information, it is possible to determine whether a gene is body-methylated across all species or in only a subset of species. If gbM varies, one can infer whether gbM is lost or gained at an internal node of the phylogeny, such that the gbM shift is inherited by all descendants of that node. Such a shift is not only phylogenetically informative, but retention of the character state among descendant suggests (superficially) that the shift is evolutionarily constrained and perhaps beneficial. Furthermore, if gbM is associated with gene expression, the clade of species that inherited the gain or loss of gbM should exhibit differences in expression relative to the rest of the species on the phylogeny. By taking this phylogenetic approach, it may be possible to glean further insights into modes, patterns and function of gbM evolution.

Here we adopt this approach to study methylomes and gene expression across eight species from the grass family (Poaceae). The family originated ~130 million years ago (Mya) (Prasad et al. 2011) or perhaps more recently (~76 Mya) (Gaut 2002; Bouchenak-Khelladi et al. 2010; Christin et al. 2014) and is a model for genomic studies due to its economic importance. Here we focus on the grasses because they represent a system of ‘intermediate’ evolutionary distance and because previous surveys of gbM have either spanned large phylogenetic distances (i.e. land plants (Feng et al. 2010; Zemach et al. 2010; Niederhuth et al. 2016; Takuno et al. 2016) or angiosperms (Feng et al. 2010; Zemach et al. 2010; Niederhuth et al. 2016; Takuno et al. 2016) or closely related species, like *A. thaliana* and *A. lyrata* (Seymour et al. 2014; Takuno et al. 2017). Also, despite their intermediate evolutionary distances, grasses vary widely in GS, permitting the address of questions related to the covariance of GS and methylation.

By contrasting methylome and gene expression patterns across eight grass taxa, we address three sets of questions. The first explores the relationship between GS and methylation patterns. Our taxonomic sampling includes three diploid grasses with genomes larger than 2 Gb, which is comparable to that of maize (2.6 Gb). Given this sampling, do genome-wide methylation levels correlate with GS, and do those correlations hold in genes? If so, what might this imply about characteristics of gbM? The second set of questions focuses on methylation within genes. Do taxa vary in the degree to which genes are methylated? Is this variation largely in the CHG context or in the canonical CG context? Finally, we take advantage of the intermediate evolutionary distances among grasses to define a set of thousands of orthologous genes and explore gbM as a phylogenetic trait. Does the presence and absence of gbM vary across eight lineages for individual genes? If so, do shifts in gbM status tend to occur in internal or external phylogenetic branches? And do these shifts correlate with either the level or variance of gene expression among taxa? By taking this phylogenetic approach, our survey of the grasses provides insights into patterns of gbM evolution and a potential functional relationship to gene expression.

## RESULTS AND DISCUSSION

### Genome-wide patterns of DNA methylation correlate with genome size

We surveyed genome-wide levels of DNA methylation and gene expression from the leaves of eight members of the Poaceae (**Table 1 and Fig 1A**), generating new methylomic data for five of eight species and new expression data for three of eight species. The eight species span most of the evolutionary breadth of the grasses and include multiple samples from the two largest subfamilies (Panicoideae and Pooideae). The species also represent a range of GS, with a 15-fold difference between the largest (*Hordeum vulgare*, 5428 Mb) and smallest (*Brachypodium distachyon*, 355Mb) genomes. Our sample includes three diploid species (*Phyllostachys heterocycla, Hordeum vulgare, Triticum urartu*) with GS similar to or larger than *Zea mays* (**Table 1**).

**Table 1:** Species Sampling for Bisulfite and RNA sequencing

| Species | Genome Size (1C - pg) | Genome Size (1C - Mb)<sup>1</sup> | Subfamily |
| --- | --- | --- | --- |
| *Setaria italica* | 0.53 | 513 | Panicoideae |
| *Zea Mays* | 2.73 | 2665 | Panicoideae |
| *Sorghum bicolor* | 0.75 | 734 | Panicoideae |
| *Oryza sativa* | 0.5 | 489 | Ehrhartoideae |
| *Phyllostachys heterocycla* | 2.12 | 2075 | Bambusoideae |
| *Brachypodium distachyon* | 0.36 | 355 | Pooideae |
| *Triticum urartu* | 4.93 | 4817 | Pooideae |
| *Hordeum vulgare* | 5.55 | 5428 | Pooideae |

<sup>1</sup> Genome sizes are from the Kew C-value database (http://data.kew.org/cvalues/) except for *Phyllostachys heterocycla* (Peng et al. 2013)

**Figure 1.**
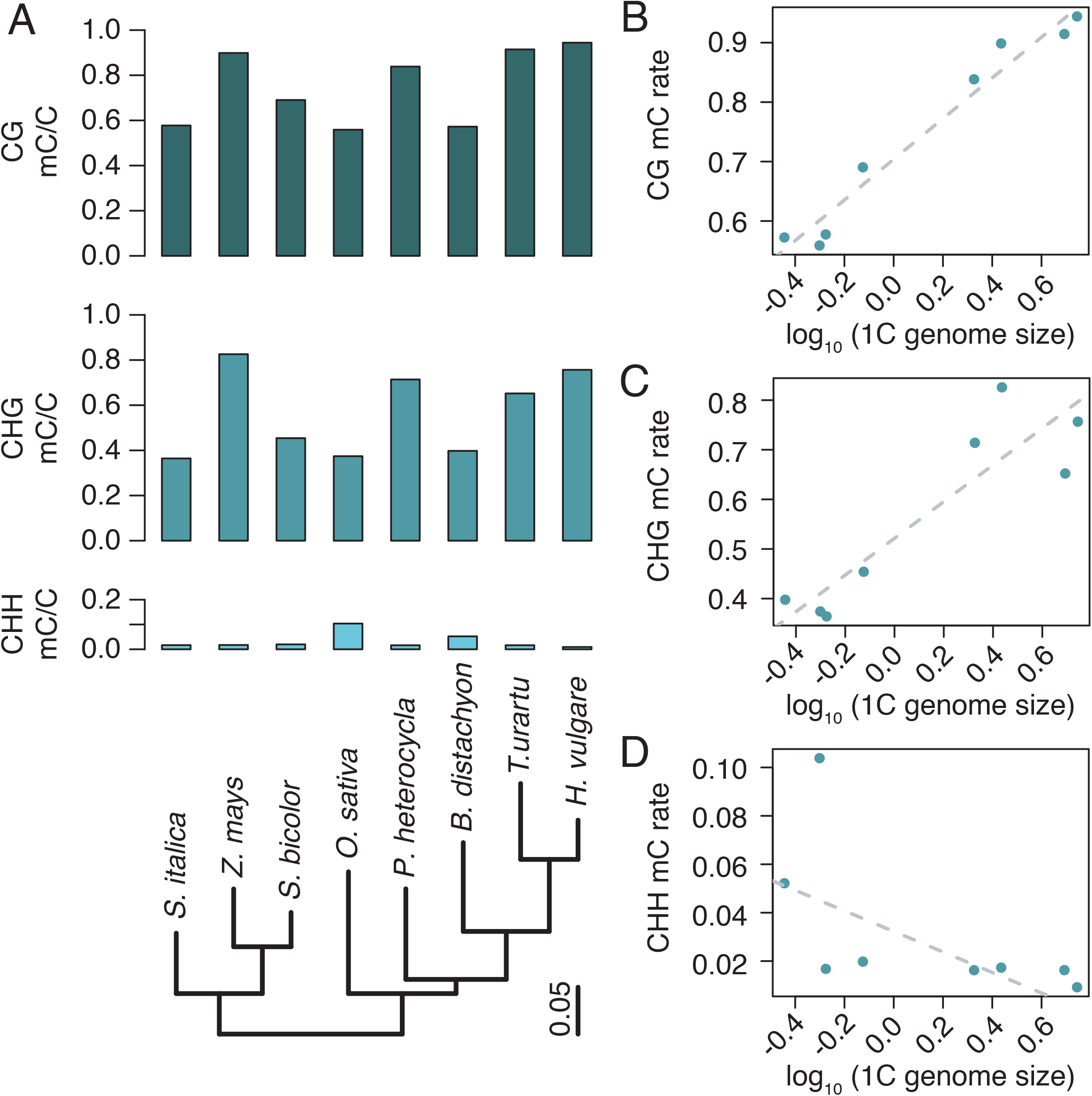
Genome-wide levels of DNA methylation and their relationship to genome size. A) Genome-wide levels of DNA methylation calculated as the proportion of methylated cytosine residues. The inferred relationship among the eight grass species is illustrated by the maximum likelihood phylogeny inferred from 2,982 single copy orthologous genes. B-D) Phylogenetic Generalized Least-Squares Regression (PGLS) of the proportion of genome-wide DNA methylation (A) on the 1C picogram genome size estimates of each species (Gregory et al. 2007; Peng et al. 2013). After accounting for the genetic relatedness between species, the proportion of CG (B) and CHG (C) methylation was significantly associated with genome size (CG b = 0.342, *P* = 8.1×10^−6^; CHG b = 0.370, *P* = 2.3×10^−3^). No significant relationship was found between the proportion of CHH (D) methylation and genome size (CHH b = −0.042, *P* = 7.0×10^−2^).

We first examined genome-wide DNA methylation patterns. Each species had detectable methylation in all three contexts – CG, CHG, and CHH. The median proportion of cytosine residues methylated in the CG and CHG contexts was 74.9% and 56.7% of sites across all species. In every case, CG methylation was higher than CHG methylation (by 7 to 26% more), and the proportion of methylated cytosines at these two contexts was highly correlated (spearman’s rho = 0.80952381; *P* = 0.02178). Each of the three species with larger genomes (*P. heterocycla, H. vulgare, T. urartu*) had a similar or slightly higher proportion of methylated CG sites than *Z. mays*, with 94.4% of CG residues methylated in *H. vulgare* (**Fig 1A**). Rates of CHG methylation were >65% in all of the four species with genome sizes greater than 2 Gb, but *Z. mays* continued to have the highest proportion of CHG methylation, at 82.6%. Similar to previous reports (Niederhuth et al. 2016), methylation levels at CHH residues were low, ranging from 0.9% to 10.3% among species. Surprisingly, the proportion of methylated CHH residues was negatively correlated with the other two contexts (CG: spearman’s *rho* = −0.8333; *P* = 0.01538; CHG spearman’s *rho* = −0.547619; *P* = 0.171). Altogether, the hierarchical relationship among contexts - i.e, the observation that the proportion of methylated cytosines is higher in CG than for either CHG or CHH - extends previous observations (Niederhuth et al. 2016; Roessler et al. 2016), but the mechanistic causes of this hierarchy remain obscure. One possibility is that the fidelity of the methylation maintenance machinery is higher than that of the initiation machinery. Even though mCHG is symmetric and can be maintained across cell divisions by *CMT3*, mCHG per-site levels are not 100% and, as a result, may require reinitiation after cell division. In fact, a subset of CHG sites are controlled by both *CMT3* and *DRM1/2* (Stroud et al. 2013).

We examined the relationship between genome size (1C pg) and the proportion of genome-wide DNA methylation using phylogenetic generalized least squares (PGLS) (Symonds and Blomberg 2014). To pursue this approach, we first inferred a maximum likelihood phylogeny based on the identification of 2,982 single-copy orthologs across the eight species (see **Methods**). The resulting phylogeny had 100% bootstrap support at each node and was topologically consistent with previous phylogenetic treatments of the grass family (Saarela et al. 2018) (**Fig 1A**). We used the branch lengths from the phylogeny in PGLS analyses, which identified significant positive relationships between GS and genome-wide DNA methylation levels at CG and CHG residues (**Fig 1B, 1C**; CG b = 0.342, *P* = 8.09×10^−6^; CHG b = 0.369, *P* = 2.3×10^−3^). In contrast, CHH methylation was negatively related to GS but not significantly so (*b* = −0.042, *P* = 7.0×10^−2^; **Figure 1D**).

In some respects, the positive correlations between GS and both CG and CHG methylation levels are not unexpected, for two reasons. The first is that significant relationship between CG and CHG methylation has been detected at a genome-wide level across a sample of 34 angiosperms (Niederhuth et al. 2016). However, this relationship only held for CHG after removing the species with the largest genome, *Z. mays (Niederhuth et al. 2016)*. Hence, our results solidify this previous finding for the grass family, based on a broader range of GS. The second reason is that larger genomes have higher transposable element (TE) content (Tenaillon et al. 2010) and hence have more DNA to methylate (Alonso et al. 2015). In this context, the higher average *level* (or proportion) of cytosine sites that are methylated in large genomes may reflect the fact that TEs are generally methylated at higher levels and that large genomes have a higher ratio of TEs relative to genes.

### Methylation patterns across all annotated genes

We generated genome-wide datasets in part to investigate patterns of methylation within genes. Moving toward that aim, we first measured absolute levels of genic methylation, including intron sequences, across all annotated genes in each of the genomes. Across our Poaceae sample, the average level of genic methylation was lower at both CG and CHG residues than the corresponding genome-wide levels, reflecting the well-established fact that genes tend to have lower methylation levels than TEs in these two contexts (Zhang et al. 2006; Zilberman et al. 2007; Cokus et al. 2008; Lister et al. 2008). This decrease was substantial; the level of CG methylation in genes was 23-47% lower than genomic levels, and there was a similar decrease for CHG sites (**Table S1**). Interestingly, for the four species with genomes larger than 2Mb, CHG methylation levels of genes exceeded 10%, reaching as high as 34.2% in *Z. mays* (**Table S1**). This contrasts with observations from Brassicaceae species and especially from *A. thaliana*, which has low levels of CHG methylation in genes (Seymour et al. 2014; Takuno et al. 2017). Levels of CG and CHG were both further reduced when excluding intron sequences (**Table S1**), indicating that the grass species surveyed here have considerable intronic methylation. *Z. mays*, in particular, exhibited a dramatic reduction (>20%) in CG and CHG mC when considering only exon sequences relative to entire genes (**Table S1**). In contrast, CHH methylation levels of genic regions were comparable to genome-wide levels, 1.1% versus 1.7% (**Table S1**). When introns were excluded, exons had a median CHH level of 0.2% (**Table S1**).

To query the relationship between GS and DNA methylation in genes, we repeated GS correlation analyses using genes. The correlations were reduced at genes relative to genome-wide measures (**Fig S1**); only CG remained significantly correlated with genome size (CG *b* = 0.204, *P* = 7.5×10^−4^; CHG *b* = 0.137, *P* = 1.03×10^−1^; CHH b=-0.01, *P* = 1.93×10^−1^). When the data were limited to exons only, no correlations remained (**Fig S1**). These analyses indicate that the effect of GS on the methylation level of genes is limited, suggesting that genic sequences are, for the most part, buffered from the consequences of neighboring TE methylation, including those TEs that reside in introns.

After examining these broad patterns, we turned to the methylation status of individual genes, applying a statistical approach to determine whether genes were methylated above or below the genic background level for each species (see **Methods**) (**Fig S2**). We found that a similar proportion (30.1%-39.6%) of genes were methylated significantly above the background CG level across species, and we categorized these genes as “body-methylated” (BM) (**Fig 2A**). Following convention (Takuno and Gaut 2012; Takuno et al. 2017), we also classified genes with significantly lower-than-background methylation levels as “under-methylated” (or UM) and genes that could not be assigned confidently as either BM or UM as “intermediately-methylated”, or IM. The distributions of absolute levels of DNA methylation at CG residues for each category and each context are provided in **Fig S3**.

**Figure 2.**
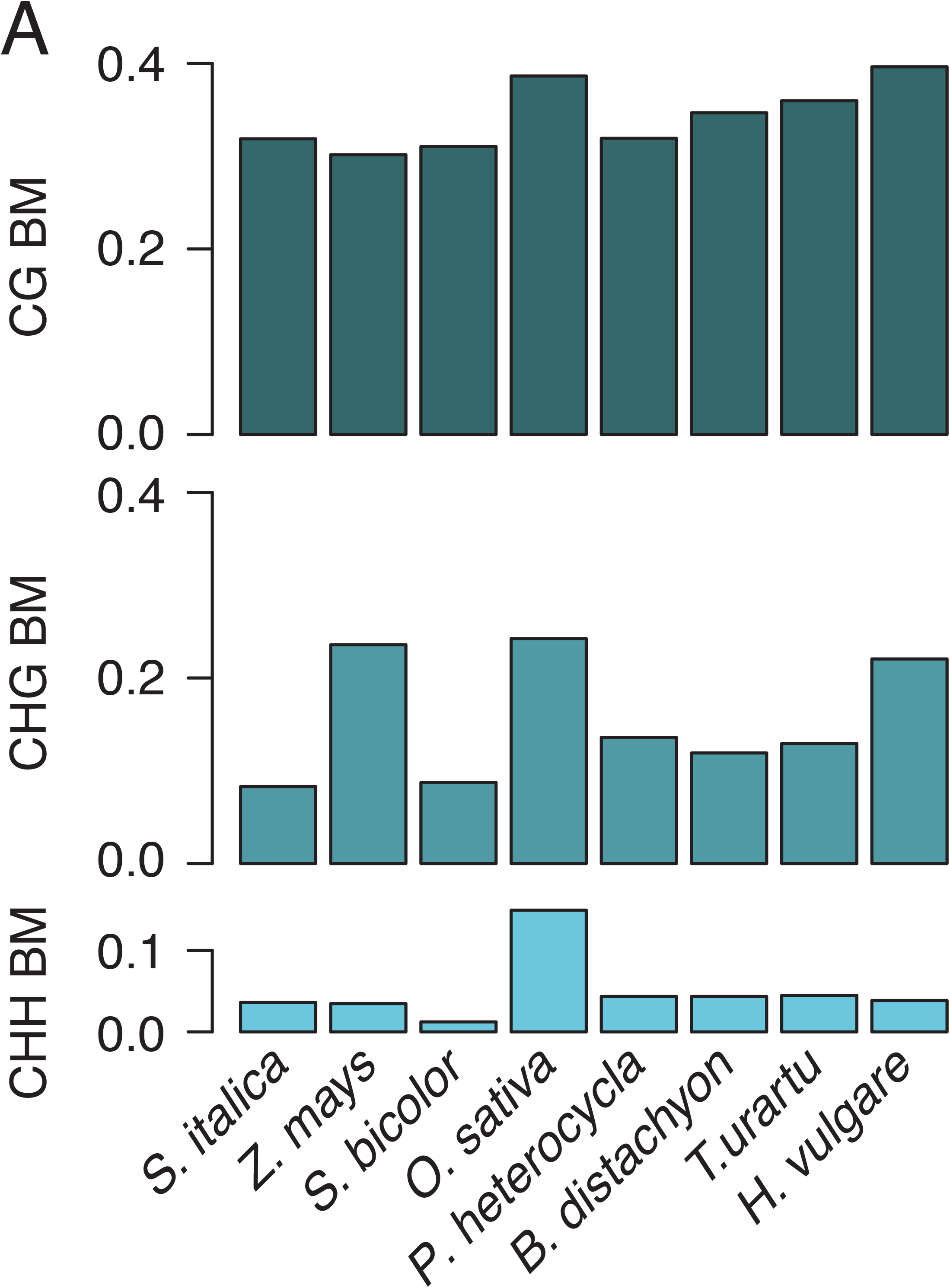
The proportion of all annotated genes within each species that are body methylate (BM). BM status of coding sequences (exon only) was inferred using a binomial test (see **Methods**) to identify coding sequences with methylation levels higher than the species’ background level in each in each context – CG, CHG, CHH.

Although gbM typically refers to methylation at CG residues located within a gene, and in this case within exons, we performed the same statistical test to identify genes (exon only) with hyper- and hypo-methylation at CHG or CHH sites. As expected, fewer genes contained CHG or CHH body methylation relative to CG methylation, and the proportion of BM genes for these two contexts was more variable across species than the CG context (CHG: 8.3%-24.2%; CHH: 1.2%-14.9%) (**Table S2**). Three species in particular were notable for having a high number of CHG BM genes - *Z. mays, O. sativa*, and *H. vulgare*. Given the small GS of rice (~500 Mb; **Table 1**), it is obvious that GS is not a good predictor of the percent of CHG BM genes. *Z. mays* was remarkable in its high CHG level and in one other respect, the degree of overlap between CG BM and CHG BM (**Table S2**). Only 23% of CG BM genes in *Z. mays* lacked CHG and CHH BM.

In most cases, methylation in CHG and CHH contexts has been associated with the suppression of gene expression. Accordingly, we measured gene expression in each taxon and explicitly contrasted expression level among BM genes in the CG, CHH and CHG contexts. Genes containing CHG and CHH BM were expressed at significantly lower levels than CG BM genes (**Fig S4**; one-sided Wilcoxon rank sum *P* < 0.01 for all comparisons except *T. urartu* CHH), suggesting that the methylation status of these genes is driven by the molecular mechanism responsible for suppressing TE activity. Unlike TE-linked methylation, which typically has a repressive effect on gene expression, CG only methylation in genes has been associated positively with gene expression levels (Zhang et al. 2006; Zilberman et al. 2007; Cokus et al. 2008; Lister et al. 2008; Takuno and Gaut 2012; Meng et al. 2016), and this positive correlation held for each of the surveyed species (**Fig S5**).

### Orthologous genes and methylation as an evolutionary trait

The first comparisons of gbM between species revealed that most orthologs had highly conserved methylation levels (Takuno and Gaut 2013; Seymour et al. 2014), because most (but not all) orthologs retain relative methylation levels between species. However, the exceptions to this rule - i.e, genes that have shifted gbM status from UM to BM or vice versa - have helped illustrate that shifts in gbM are weakly associated with shifts in gene expression (Takuno et al. 2017). To date, this approach has only been applied to pairs of closely related species, but here we examine shifts in status for orthologs among all eight grass species. To do so, we first identified a set of 2,982 putative orthologs that were filtered by reciprocal best-hits (see **Methods**). Given the set of putative orthologs, we contrasted their methylation statistics relative to the set of all annotated genes. One finding is that the ortholog set comprises a non-random representation of the complete gene set, because the orthologs were biased toward a higher proportion of CG-only methylation (**Table S2**). We suspect this is the case, because the ortholog set should be biased for genes that are highly conserved in both sequence and copy number and that also may be less likely to be pseudogenes or misannotated TEs. Consistent with previous studies, we find a high degree of conservation in the level of CG DNA methylation between species across the ortholog set (**Fig 3A**).

**Figure 3.**
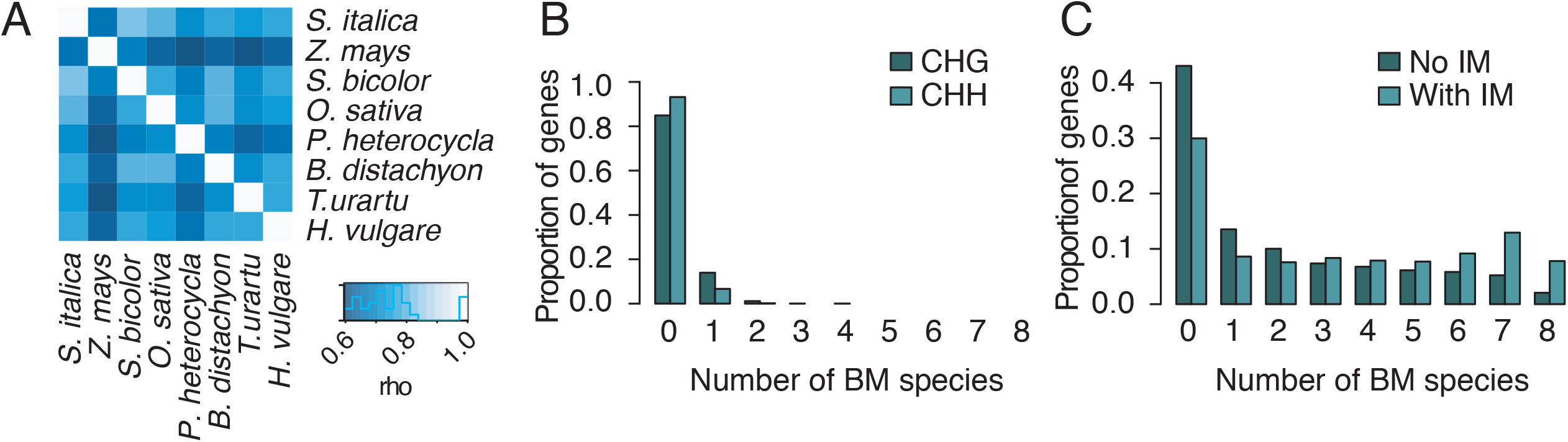
Methylation characteristics of orthologous genes. A) The correlation (Spearman’s rho) between absolute levels of gbM across orthologous genes, based on CG methylation. Orthologs with significant CHG/CHH methylation were excluded. B) The distribution of CHG or CHH BM across 2,982 orthologous genes. B) The distribution of CG BM across 2,336 orthologous genes. Orthologs with non-CG methylation (from B) were excluded. ‘No IM’ refers to cases where orthologs with intermediate methylation (IM) were considered to be under-methylated; ‘With IM’ plots the distribution when orthologs with IM were considered to be body-methylated.

#### CG status varies more often than CHG/CHH status

One advantage of the ortholog set is the ability to examine methylation as a phylogenetic trait. To do so, we examined methylation status for each of the eight species. Focusing first on CHG/CHH methylation, we found that 18.2% of orthologs (or 521 total) had CHG and/or CHH BM status. Only 103 of these 521 orthologs were BM for both CHG and CHH contexts. When CHG or CHH BM was detected, however, it was never conserved across all eight taxa. In fact, it was found in only one of the eight species for 479 of the 521 orthologs (92%) (**Fig 3B**), showing that shifts from UM to BM tend to be lineage-specific. Most species exhibited CHG/CHH BM gains in only a few orthologs (median = 32), but a few lineages were exceptional. For example, *P. heterocycla* and *T. urartu* gained BM at 199 and 215 orthologs, respectively. Given previous work (Meng et al. 2016) and our own finding that CHG is associated with suppression of expression (see above), it is reasonable to postulate that these observations represent potential pseudogenization events. Surprisingly, only 2 genes gained CHG or CHH BM in ≥ 2 taxa, and neither of these gains were phylogenetically informative. In other words, shifts in CHG/CHH BM status were rare and evolutionary labile over the timeframe defined by our data set.

We next examined CG BM status across the species, focusing on shifts in gbM status (i.e., BM to UM or vice versa) and the placement of these shifts on the phylogeny. To do so, we first excluded genes with any signal of CHH/CHG methylation, leaving 2,336 orthologs. Two patterns were evident in the distribution of CG BM status. The first was that a large proportion of orthologs maintained their BM status over evolutionary time (**Fig 3C**). When treating IM orthologs as UM, 45.2% of orthologs had a stable CG state across species, in that all eight species were either UM or BM. The majority (1,007 of 2,336 genes) were conserved UM and far fewer (48 of 2,336 genes) were conserved BM. When IM status was treated as BM, we found that a larger proportion of orthologs were methylated across all eight species (7.8% versus 2.1%) (**Fig S3**). For the remainder of the analyses presented here, IM genes were treated (conservatively) as UM. The second pattern, which was somewhat unexpected given previous observations of gbM conservation, was that nearly 55% of orthologs (1281) shifted their CG gbM status from BM to UM (or vice versa) across the phylogeny. Of these 1281 orthologs, 438 featured a shift in CG BM status within only a single species, with gains in CG BM occurring more frequently than losses (**Fig 3C**). The remainder included shifts in CG gbM status in more than one species (**Fig 3C**).

It has been demonstrated repeatedly that molecular characteristics differ between BM and UM genes, such as longer length and slower rates of molecular evolution for BM genes (Takuno and Gaut 2012; Takuno and Gaut 2013; Takuno et al. 2017) (**Fig S6**). We assessed whether these molecular signatures were associated with the degree of conservation in CG gbM status across our sample of grass species. Similar to previous work, we found that BM genes are associated with longer coding sequence lengths and slower rates of molecular evolution at both synonymous and non-synonymous sites (**Fig 4**). We also found that for each of these characteristics there is a significant linear relationship with the degree of CG gbM conservation (**Fig 4A,C,D**). For example, there was a positive relationship between gbM conservation and average gene length across the species (adjusted R^2^ = 0.2032, *P* = 1.91×10^−117^) (**Fig 4A**), and evolutionary rates at both synonymous and non-synonymous sites were negatively correlated with the degree of conservation in gbM (non-synonymous: adjusted R^2^ = 0.016, P = 9.31×10^−10^; synonymous: adjusted R^2^ = 0.049, P = 5.53×10^−27^) (**Fig 4C, 4D**). However, there does not appear to be a significant relationship between gbM conservation and degree of gene length variation, as measured by the coefficient of variation (adjusted R^2^ = 0.0004, P = 1.62×10^−1^) (**Fig 4B**).

**Figure 4.**
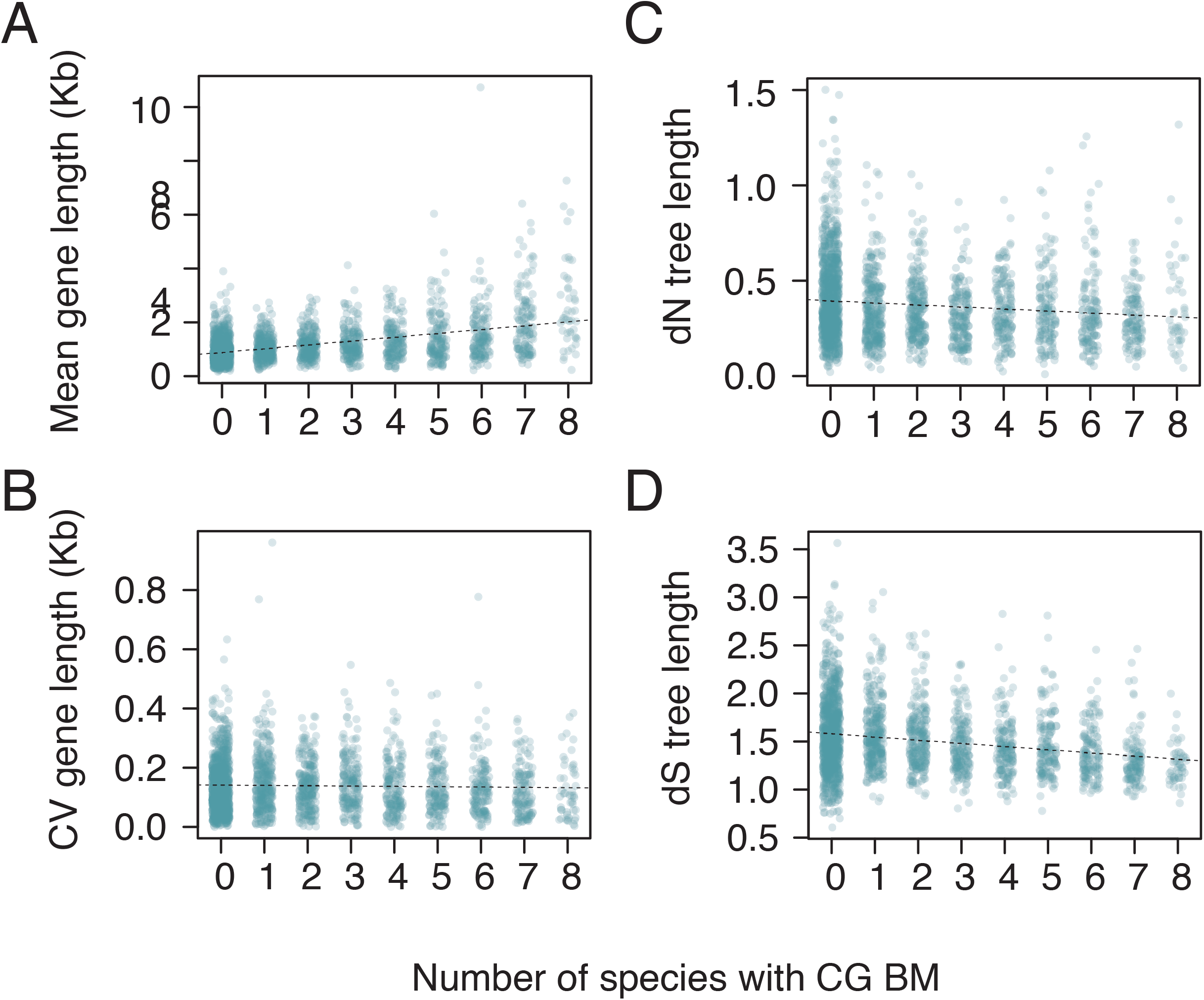
Ortholog characteristics and their relationships to conservation of CG gbM. A) Gene length (based only on the CDS) is positively correlated with CG gbM conservation (adjusted R^2^ = 0.2032, *P* = 1.91×10^−117^). B) The coefficient of variation (CV) for gene length is uncorrelated with gbM conservation (adjusted R^2^ = 0.0004, *P* = 1.62×10^−1^). B) The rate of nonsynonymous substitution (dN) is negatively related to gbM conservation (adjusted R^2^ = 0.01617, *P* = 9.31×10^−10^). Total tree length refers to the branch length of the distance tree built from pairwise dN estimates for each orthologous gene cluster. D) The rate of synonymous substitution (dS) is negatively related to gbM conservation (adjusted R^2^ = 0.4999, *P* = 5.53×10^−27^). Total tree length refers to the branch length of the distance tree built from pairwise dS estimates for each orthologous gene cluster. Genes with BM in the CHG or CHH context were excluded from these analyses.

If shifts in gbM are beneficial, then we might expect those changes to be maintained in subsequent lineages. Although a number of orthologs shift their gbM status in more than 2 species, we found only 16 orthologs for which shifts in gbM status corresponded to the species’ phylogeny (**Table S3**). Hence, gbM shifts, like shifts in CHH/CHG methylation status, rarely conveyed phylogenetic information. To more formally examine whether shifts in CG status were phylogenetically random, we used stochastic character state mapping to infer the location of BM shifts for the 1,281 orthologs that varied in gbM status across species (**Fig. 3C**) (Nielsen 2002; Huelsenbeck et al. 2003). The majority of gbM shifts occurred on terminal branches (**Fig 5A**). These terminal branches constituted 68.45% of the species tree (**Fig. 1A**), but we still found a significant enrichment of character state shifts on these branches (one-sided Wilcoxon rank sum test, *P* = 4.23×10^−136^). The dearth of shifts in gbM status on internal branches indicate that the dynamics of gbM are driven by lineage-specific changes and suggest that these shifts tend to be short-lived on evolutionary time-scales. Overall, the pattern of shifts on the phylogeny are consistent with an interpretation that changes in gbM status tend to be deleterious.

**Figure 5.**
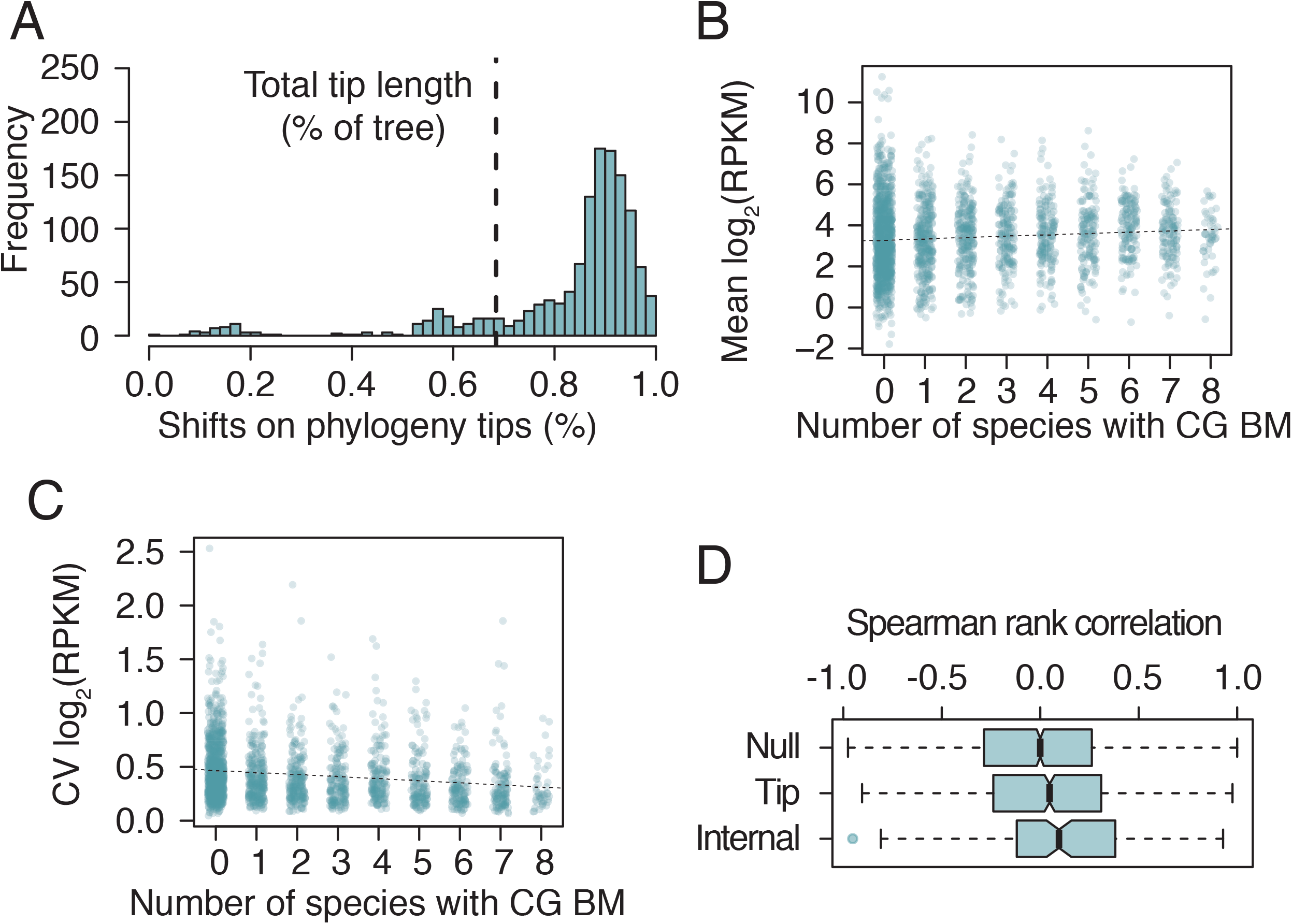
The evolutionary dynamics of gbM and impacts on gene expression. A) The location of gbM shifts on the species tree (Fig 1A) plotted as a percent of the total tree length. Stochastic character state mapping was used to predict the location of character state changes (i.e. from BM to UM or vice versa) (100 simulations per ortholog group). Plotted values per ortholog represent the average location of gbM shifts across 100 simulations. Dashed line indicates the total length of the tip branches as a percent of the total tree length (68.45%). The distribution has a mean significantly different from 68.45% (one-tailed Wilcoxon rank sum test, *P* = 4.23×10^−136^). B) The mean in gene expression (log_2_ RPKM) across species is positively correlated with the degree of CG gbM conservation (adjusted R^2^ = 0.007636; P = 1.53×10^−5^), and C) the coefficient in variation in gene expression (log_2_ RPKM) across species is negatively correlated with the degree of CG gbM conservation (adjusted R^2^ = 0.02924; P = 2.02×10^−15^). Genes with a log_2_(RPKM) < 1 were removed. Genes with BM in the CHG or CHH context were excluded in B) and C). D) Spearman’s rank correlation between gene body methylation rank and gene expression rank. The ranks for gene expression (log_2_(RPKM)) and quantitative levels of methylation were determined for each ortholog in each species. Orthologs were grouped by the location of gbM shifts on the phylogeny (A). Shifts on tip branches were those that occurred at >68.45% of the length of the phylogeny. Shifts on internal branches occurred <=68.45%. The null distribution of correlation between DNA methylation and expression was estimated by randomly sampling a vector of gene expression ranks for each ortholog.

#### CG shifts and gene expression

Thus far, the potential relationship between CG BM status and gene expression has been unclear, which in turn has cast doubts on the potential functionality of gbM. This potential relationship is important; if gbM status is associated with gene expression, then it provides either a direct or indirect phenotype, making gbM potentially visible to natural selection and providing a logical explanation for the long-term conservation of gbM status for some genes. Given that gbM has been observed to be correlated with gene expression (Muyle and Gaut 2019) and given the hypothesis that gbM has a homeostatic function (Zilberman 2017), our null expectations were straightforward. First, we expected a significant relationship between mean gene expression levels and the number of species with BM status. To test this expectation, we measured the mean gene expression level for each gene and categorized the gene as having BM from 0 up to 8 taxa. There was indeed a significant positive relationship (adjusted R^2^ = 0.007636; P = 1.53×10^−5^) (**Fig 5B**), albeit one that explains little of the variance.

The second -- and potentially more important -- prediction is that gbM plays a homeostatic role. If that is the case, the variance in gene expression among taxa should be lower for genes with conserved BM status across all eight species than for genes that have lost BM status in one, two and more species. Accordingly, we plotted the coefficient of variation (CV) in gene expression for individual orthologs against the number of species with BM status (**Fig 5C**). As predicted, CV scales negatively with the number of species, indicating that the reduction in variation is not only a function of the mean (adjusted R^2^ = 0.02924; P = 2.02×10^−15^). In other words, conserved BM genes are significantly more conserved in their level of gene expression across species than genes that vary in BM status.

We delved further into this phenomenon by assessing the data for orthologs that varied in gbM status across the phylogeny. First, we characterized two sets of orthologs – those with shifts of BM that occurred on the terminal branches (>68.45% in **Fig 5A**) and those with BM shifts that occurred on the internal branches (<68.45% in **Fig 5A**). The “tip branch” group contained 1103 orthologs and the second “internal branch” group contained 178 orthologs. We postulated that if gbM and gene expression are correlated, then methylation rank and gene expression rank should be related for both groups. This approach has the advantage that it does not rely only on BM status but on relative ranks of methylation and expression. For both of these ortholog sets, the two ranks were more correlated with gene expression than expected by chance (for terminal branches: Wilcoxson rank sum *P* = 0.006221; for internal branches: Wilcoxson rank sum *P* = 0.001388) (**Fig 5D**). Although significant, the median degree of correlation per ortholog was low, explaining only 4.7% (tip branches) to 9.5% (internal branches) of the relationship between DNA methylation and gene expression.

Taken together, these analyses provide evidence that changes in gbM status are related to gene expression. There are, of course, important caveats to this inference. For example, our method of defining status as BM, IM or UM is coarse, but for that reason we expect our approach should tend to miss, rather than over-estimate, associations. We also note that ranking statistics, which are largely independent of status designations, also find significant correlations. Second, the approach examines expression patterns across genes without taking into account phylogenetic relationships as does PGLS, for example. We note, however, that because shifts in CG BM status tend to be lineage specific, PGLS has no statistical basis (or power) in this case. Third, the approach identifies associations, but it does not allow us to discern causality - i.e. if gbM shifts lead to gene expression differences or vice versa or indeed whether the relationship is indirect. Previous work has shown that changes in gene expression may proceed alterations to DNA methylation (Secco et al. 2015), but this has not been investigated for gbM specifically. Finally, we note that correlations are significant but small, which is consistent with previous work that has found a relationship between expression and gbM (Takuno et al. 2017; Muyle and Gaut 2019). Thus, the effect on individual genes is likely small and may be too small to detect in an experimental framework. However, the power of natural selection comes from the fact that it can act on functional features that have small effects on fitness, such as codon usage bias (Duret and Mouchiroud 1999). We thus hypothesize that gbM is a trait imminently visible to natural selection under some conditions.

## CONCLUSIONS

The major debate about gbM, as defined by CG (and not TE-like) methylation, surrounds its function. Does it have a function, and if so, what is it? That gbM is maintained for millions of years across species indicates some evolutionary constraint, which indirectly implies function. Function has also been tested directly via potential effects on gene expression, and the outcomes have been mixed. On the one hand, several authors have looked for effects of gene expression in *A. thaliana met1* methylation mutants, and none have detected an association between loss of methylation and global shifts in gene expression at gbM genes (Zhang et al. 2006; Zilberman et al. 2007; Cokus et al. 2008; Lister et al. 2008; Bewick et al. 2019). Efforts to detect expression effects of de-methylation in *E. selagineum* have also yielded negative results (Bewick et al. 2016). On the other hand, increased levels of gbM correlate with increased expression within species (Zhang et al. 2006; Zilberman et al. 2007; Cokus et al. 2008; Lister et al. 2008; Takuno and Gaut 2012; Meng et al. 2016), gbM status was the strongest predictor of allele-specific expression in *Capsella grandiflora* (Steige et al. 2017), gbM genes have less transcriptional noise than non-methylation genes (Horvath et al. 2019), and, finally, evolutionary comparisons have suggested slight but detectable effects on gene expression when genes change in status from BM to UM or vice versa (Takuno et al. 2017; Muyle and Gaut 2019).

To date, evolutionary analyses have tested for associations between two species but not on larger samples. Here we have examined orthologs across multiple species in the context of the species phylogeny. We find, first, that shifts in CG gbM occur predominantly on the tips of the trees and are rarely phylogenetically informative. These observations corroborate previous work suggesting that gbM evolution is conservative - although, surprisingly, less conservative than CHG/CHH status in these taxa - and therefore also suggest that shifts in CG gbM status are predominantly deleterious. One weakness of this conclusion is the density of taxon sampling; with a denser tree, it becomes more likely to observe gbM shifts on internal branches that are passed on to descendants. We nonetheless predict that further study on denser trees will yield a similar bias toward change on the tips. Second, we show detectable associations between gene expression and the number of taxa that have an ortholog with gbM status. More specifically, orthologs that are BM have increased expression and lower coefficients of variation as a function of the number of species.

Based on this information, the evidence for an association between gbM and gene expression becomes more compelling, but it is also clear that the effect is very subtle - i.e., our linear fits explain only 0.7% for the mean and 2.9% for the CV of expression. Given the current range of evolutionary observations, one of the next frontiers is to evaluate whether there is evidence for selection on gbM within populations. There is indirect evidence that purifying selection acts on gbM genes in a population sample of *A. thaliana (Takuno et al. 2017)*, but no evidence yet for selection operating specifically at CG residues in *A. thaliana* genes (Vidalis et al. 2016). Another frontier is causality: is the association between gbM and gene expression causal? If not, what factors affect both quantities and are causal? If there is a causal relationship, in which direction does causality lie? Hence, even as the picture comes into tighter focus that gbM tends to be subject to evolutionary constraint and to associate weakly with patterns of gene expression, there are still many lingering questions about the functional significance gbM.

## MATERIALS AND METHODS

### Publicly Available Data

Our total datasets consist of whole-genome RNAseq and BSseq from each of eight species. Some of these data were generated previously and retrieved from public sources. For example, BSseq data was previously generated for three of the eight species. These were download from https://www.ncbi.nlm.nih.gov/sra for *B. distachyon* (SRR628921, SRR629088, SRR629207), *O. sativa* (SRR1035998, SRR1035999, SRR1036000) and *Z. mays* (SRR850328). The data were generated from seedlings from *Z. mays* and from leaves from *B. distachyon* and *O. sativa*. We generated BSseq from young leaves harvested from 6 week old plants for the remaining five species, following the methods outlined below. Note that methylation patterns do not vary substantially among plant tissues (Schmitz et al. 2013; Seymour et al. 2014; Widman et al. 2014; Roessler et al. 2016); hence the *Z. mays* seedling data was unlikely to contribute substantially to the differences among species.

RNAseq were publicly available for five of the eight species. We downloaded RNAseq data from https://www.ncbi.nlm.nih.gov/sra for *B. distachyon* leaves (SRR3170524, SRR3170525, SRR3170526), *O. sativa* leaves (SRR1005356, SRR1005354, SRR1005355), Z. *mays* leaves (SRR520998, SRR520999, SRR521003, SRR521006), *P. heterocycla* leaves (ERR105067, ERR105068) and *H. vulgare* leaves (SRR3290257, SRR3290258, SRR3290283, SRR3290284, SRR3290313, SRR3290314, SRR3290315). The data for *B. distachyon* and *O. sativa* had matched BSseq and RNAseq data from the same tissue. All of the RNAseq data had three replicates except those of *P. heterocycla* (2 replicates).

All of the RNAseq and BSseq data, whether generated for this study (see below) or downloaded from previous papers (see above), were mapped to reference genomes. The reference accessions were used for each of the eight Poaceae species (International Rice Genome Sequencing Project 2005; Paterson et al. 2009; Schnable et al. 2009; International Brachypodium Initiative 2010; Bennetzen et al. 2012; Ling et al. 2013; Peng et al. 2013; Mascher et al. 2017). The version of each reference genome can be found in **Table S4**.

### Bisulfite and RNA Sequencing

We generated bisulfite data for five of the species in this study - *P. heterocycla, H. vulgare, S. bicolor, T. urartu*, and *S. italic*a. For *P. heterocycla* a DNA sample from the reference accession was kindly provided by Dr. Zehui Jiang (Research Institute of Forestry, Chinese Academy of Forestry, Beijing, China). For the remaining four species, seeds for the reference accession were acquired and grown in soil under ambient light. Leaf tissue was harvested from 6-week old seedlings. Leaf tissue was harvested in triplicate to perform both bisulfite and RNA sequencing.

To generate BSseq data, DNA was isolated for *H. vulgare*, *T. urartu*, *S. italic*a, and *S. bicolor* with the Qiagen DNeasy Plant Mini kit. Bisulfite libraries were prepared as previously described (Urich et al. 2015) at the University of Georgia (R. Schmitz lab). Libraries were pooled and sequenced (150 bp paired-end) on the Illumina HiSeq2500. As a control for bisulfite conversion, lambda-DNA was spiked into each library preparation to measure the conversion rate of unmethylated cytosines (0.5% wt/wt). False methylation rates (FMR) for each library were estimated for each taxon (**Table S5**). In cases where lambda-DNA was not included in the library preparation (*B. distachyon* and *Z. mays*), FMRs were estimated using chloroplast DNA, which lacks DNA methylation.

RNAseq data were generated by isolating DNA using the Qiagen RNeasy Plant Mini kit, and TruSeq Illumina RNA libraries were prepared in triplicate for each species. RNAseq libraries were sequenced on the Illumina HiSeq3000 using single-end 100bp reads. The BSseq and RNAseq data generated for this study were deposited in NCBI Short Read Archive (https://www.ncbi.nlm.nih.gov/sra) under accession PRJNA340292.

### Processing of bisulfite sequencing reads and inferring methylated sites

BSseq reads were trimmed for quality and adapter sequences using trimmomatic (v0.35). Sequencing read length varied by data set. *B. distachyon, O. sativa*, and *Z. mays* were sequenced with 100 bp paired-end (PE) reads. As a result, the minimum read length threshold after quality trimming differed for *B. distachyon, O. sativa*, and *Z. mays* was 80 bp, compared to 100 bp for all remaining species. Bismark (v0.15.0), in conjunction with bowtie2 (v 2.2.7) was used to align trimmed reads to the reference genomes of each species. A seed of 20 bp with 0 mismatches was required for alignment (-N 0 -L 20) to the reference (**Table S4**). In total, the number of sequenced reads after trimming ranged from 34,010,995 to 476,202,473, and mapped coverages ranging from 6.7X to 81X was achieved for each species when all replicates were combined (**Table S6**).

The number of methylated and unmethylated reads per cytosine were calculated using the bismark_methylation_extractor (Bismark v0.15.0). Methylated cytosines were identified using a binomial test incorporating the estimated rates of bisulfite conversion errors (*P* < 0.05 after Benjamini-Yekutieli FDR correction) (Lister et al. 2008). Error rates, or false methylation rates (FMRs), were calculated as the fraction of methylated cytosines in either lambda-DNA or, when lambda-DNA was not present, the chloroplast genome (**Table S5**). A minimum coverage of 2 was required at each cytosine to determine methylation status.

### Estimation of gene body methylation (gbM)

Genome-wide levels of DNA methylation vary by species (**Fig 1A**) and as a result, we utilized a strategy to determine the gene body methylation (gbM) status of each gene relative to the species’ overall genic methylation rate independently for each context – CG, CHG, CHH. The genic background methylation rate for each context was calculated from cytosines residing in both introns and exons. Using a binomial test incorporating species-specific DNA methylation levels (Takuno and Gaut 2012), we determined whether a gene was body-methylated (BM), under-methylated (UM), or intermediately-methylated (IM). For this test, only cytosines residing in exons were considered. Following Takuno and Gaut (2012), the one-tailed *P* value for the departure of CG methylation levels from the average proportion of cytosine methylation was calculated as:

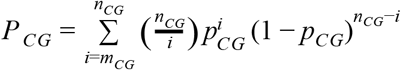

where *n_CG_* is the number of cytosine residues at CG sites in the gene of interest, *m_CG_* is the number of methylated cytosine residues at CG sites for the same gene, and *p_CG_* is the average proportion of methylated cytosine residues across all genes. When *P_CG_* < 0.05, a gene is more densely methylated than expected and termed body-methylated (BM). Under-methylated (UM) genes are those genes that are less densely methylated than expected (*P_CG_* > 0.95). Intermediately-methylated (IM genes) are those genes where 0.05 < *P_CG_* < 0.95. The distribution of the proportion of DNA methylation for each context relative to the average genic proportion of DNA methylation is shown in **Figure S2**.

### Processing of RNA sequencing reads

RNA sequencing reads were quality and adapter trimmed using trimmomatic (v0.35). A minimum read length threshold of 45 bp was required after trimming. Trimmed reads were aligned to the most recent coding sequence annotation available for each genome (**Table S4**). Alignments were performed using bwa (v0.7.12) allowing 2 mismatches (-n 2). Raw read counts were normalized (TMM) in edgeR (v3.20.9) for each species and reads-per-kilobase-mapped (RPKM) was estimated from the fitted values. See **Table S7** for RNAseq library and alignment statistics.

### Orthologous gene identification and substitution rate estimates

Orthologous genes with a single copy in each of the eight surveyed species were identified using orthomcl (v2.0.9) and blast (v.2.2.30) with the following options “-evalue 1e-5 -outfmt 6” used for blastp. A representative isoform for each protein sequence was obtained from the reference annotations (**Table S4**). For *Z. mays*, a representative protein sequence for each gene was chosen by selecting the longest isoform. A total of 2,982 single copy orthologous genes were identified among the eight species (**Table S8**).

To estimate substitution rates, protein sequences were aligned for each ortholog group using MUSCLE (v3.8). The command “tranalign” was used to extract aligned nucleotide sequences. Pairwise non-synonymous (dN) and synonymous (dS) substitution rates were calculated in R using the kaks() function in the R package seqinR (v3.4-5).

### Stochastic mapping

Stochastic character state mapping was performed using phytools (v.0.6.60) in the R statistical computing environment (v3.4.1). To construct a phylogenetic tree for character state mapping, protein sequences from each 1:1 ortholog were first aligned using MUSCLE (v3.8). Protein alignments were then converted to nucleotide alignments with the ‘tranalign’ from EMBOSS (v6.5.7). Concatenated nucleotide alignments of 1:1 orthologs were then used to infer a maximum likelihood phylogenetic tree using *ape* (v5.2) and phangorn (v2.4.0) using a GTR substitution model. Next, for each ortholog, character state transitions (from BM to UM or visa versa) were then simulated 100 times to estimate both the number of transitions and their location on the phylogeny.

### Phylogenetic generalized least-squares regression (PGLS)

The relationship between genome size (log_10_ 1C) and genome-wide levels of DNA methylation was queried after correcting for relatedness using PGLS regression. PGLS regression was performed using nlme (v3.1.131) in the R statistical computing environment (v3.4.1).

### Correlation between gbM and gene expression at orthologs

The degree of correlation between gene body methylation levels and gene expression was calculated to determine if orthologs with shifts in BM status, as determined by stochastic character state mapping, shifted gene expression levels in a coordinated fashion. First, ranks for gene expression and quantitative levels of methylation were determined for each ortholog in each species. This step was necessary to remove the species-specific differences in absolute gene expression and methylation levels. Next, the spearman rank correlation between expression rank and methylation rank was determined for each ortholog. Permutations were also performed to determine background levels of correlation between methylation levels of each ortholog and randomly sampled gene expression ranks.

## ACKNOWLEDGEMENTS

We thank Galen Martin, Aline Muyle, and Shohei Takuno for providing feedback on the manuscript and Rebecca Gaut for technical assistance and RNAseq library preparation. We also thank Robert Schmitz (U. Georgia) and his lab for BS-seq library preparation. This work was supported by the National Science Foundation (grant numbers 1542703 to B.S.G; 1609024 to D.K.S.). The BSseq and RNAseq data generated for this study were deposited in the NCBI Short Read Archive (https://www.ncbi.nlm.nih.gov/sra) under accession PRJNA340292.

