## Supplemental Figures and Tables for "Phylogenetic shifts in gene body methylation correlate with gene expression and reflect trait conservation"

### SUPPLEMENTAL FIGURE LEGENDS

**Figure S1. Relationship between genome size and methylation levels in genes.** Phylogenetic Generalized Least-Squares Regression (PGLS) of the proportion of DNA methylation in genic sequences (exons+introns) and CDS (exons only) on the 1C picogram genome size (Gregory et al. 2007; Peng et al. 2013) (Genic sequences: CG b = 0.204, $P = 7.5 \times 10^{-4}$; CHG b = 0.137, $P = 1.0 \times 10^{-1}$; CHH b = -0.01, $P = 1.9 \times 10^{-1}$. CDS: CG b = 0.082, $P = 7.2 \times 10^{-2}$; CHG b = 0.021, $P = 5.2 \times 10^{-1}$; CHH b = -0.006, $P = 1.9 \times 10^{-1}$).

**Figure S2. Distribution of DNA methylation in coding sequences.** The distribution of the proportion of methylated cytosines in each coding sequence (CDS) is shown for each species. The species-specific background level of genic methylation (exon+intron) is shown by the dashed line in each plot.

**Figure S3. Distribution of DNA methylation by gbM category.** The distribution of the proportion of methylated cytosines (CG, CHG, and CHH) in each coding sequence is shown for each gbM category – BM, IM, and UM – in each species.

**Figure S4. The relationship between gbM category and gene expression.** The levels of gene expression ($log_2$(RPKM)) for genes with either CG, CHG, or CHH BM are shown. CHG and CHH BM are associated with reduced gene expression levels in most surveyed species compared to the expression level of CG BM genes, with the exception of CHH BM genes in *T. urartu* (one-sided Wilcoxon rank sum test $P < 1.67 \times 10^{-4}$ for CHG vs CG and CHH vs CG comparisons; $P = 2.63 \times 10^{-1}$ for *T. urartu* CHH BM vs CG BM).

**Figure S5. The relationship between CG methylation and gene expression.** The quantitative level of DNA methylation (average of CG mC level of each cytosine covered with 2 or more reads) by gene expression quantile ($log_2$(RPKM)).

**Figure S6. Ortholog characteristics for BM and UM orthologs.** The difference between BM and UM orthologs for number of characteristics. In this case, UM refers to orthologs that lack CG methylation across all 8 species. BM refers to those orthologs with CG methylation in at least one species. Orthologs with CHG or CHH BM in any species are excluded.

### SUPPLEMENTARY TABLE LEGENDS

**Table S1. The level of DNA methylation at CG, CHG, and CHH residues.** Genome-wide, genic (exon+intron) and CDS (coding sequence, exon only) proportion of mCG, mCHG, and mCHH.

**Table S2. Summary of annotated genes and orthologs methylated in each context.** The total number of annotated genes (single isoform only) and orthologs for each species. The column "Total (no NA) refers to the number of genes (or orthologs) for which there was coverage and methylation information could therefore be obtained. For the orthologous gene set, all 8 species were required to have methylation information. The body-methylation status of each gene (or ortholog) was determined using a binomial test to identify genes (or orthologs) with methylation levels higher than the species' background level (see **Methods**).

**Table S3. Orthologs with phylogenetic signal.** Orthologs with BM gains/losses that directly follow the species phylogeny (**Fig 1A**) are listed. BM is indicated by 1 and UM is indicated by 0. Correspondence between ortholog IDs and gene IDs is found in Table S8.

**Table S4. Information on genomic resources.** Lists genome and annotation versions for each species.

**Table S5. Rate of false positive methylation errors for bisulfite sequencing libraries.** Error rates, or false methylation rates (FMRs), were calculated as the fraction of methylated cytosines in either lambda-DNA or, when lambda-DNA was not present, the chloroplast genome.

**Table S6. Bisulfite sequencing statistics.** SRA ID as well as the number of trimmed and mapped paired-end reads are listed for each library.

**Table S7. RNA sequencing statistics.** SRA ID as well as the number of trimmed and mapped paired-end reads are listed for each library.

**Table S8. List of orthologous gene groups.** Gene names for each species are listed for each single copy ortholog.

S1

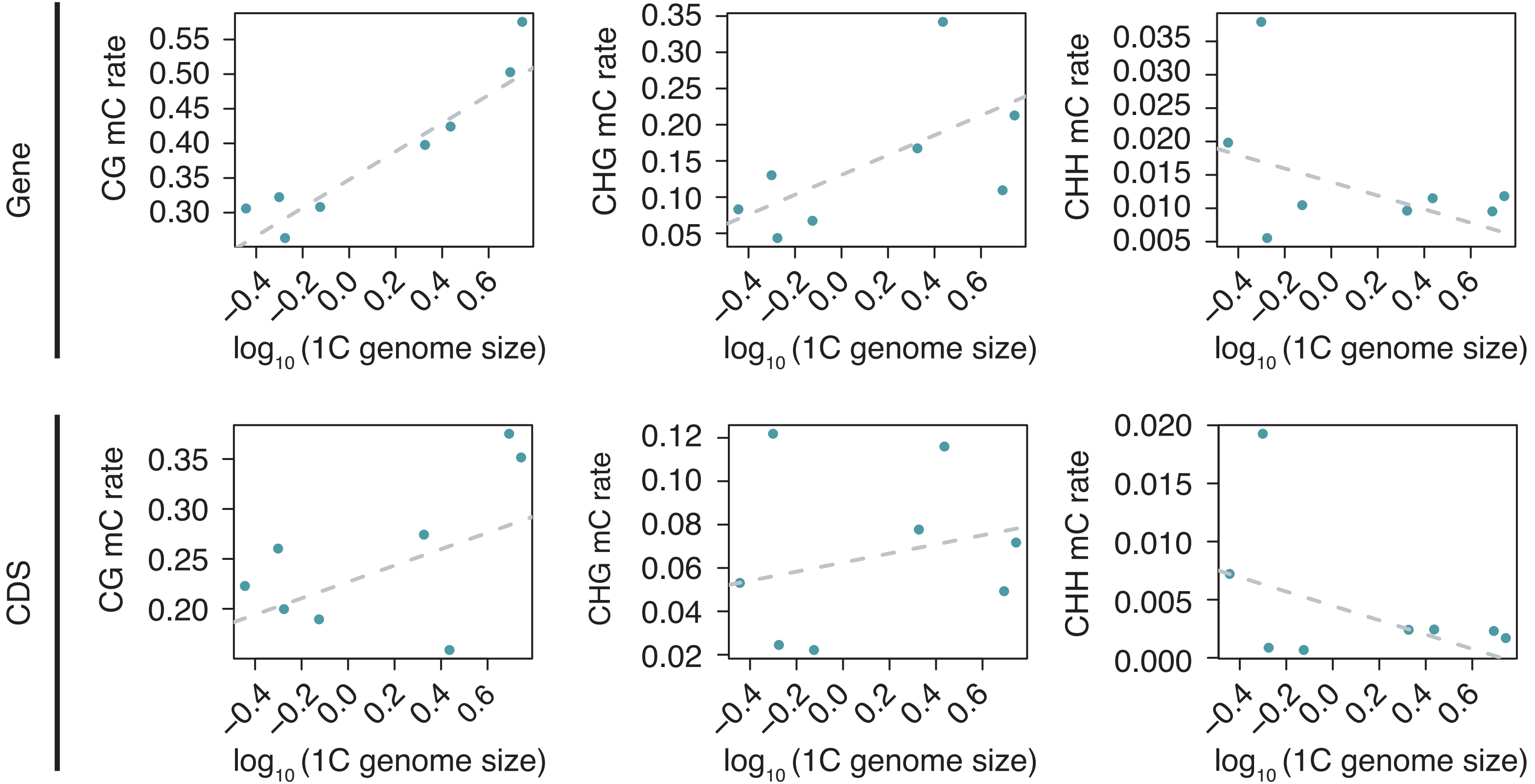
Gene
CDS
CG mC rate
CHG mC rate
CHH mC rate
0.55
0.50
0.45
0.40
0.35
0.30
0.25
0.20
0.15
0.10
0.05
0.035
0.030
0.025
0.020
0.015
0.010
0.005
0.12
0.08
0.06
0.04
0.02
0.000
-0.4
-0.2
0.0
0.2
0.4
0.6
log10 (1C genome size)

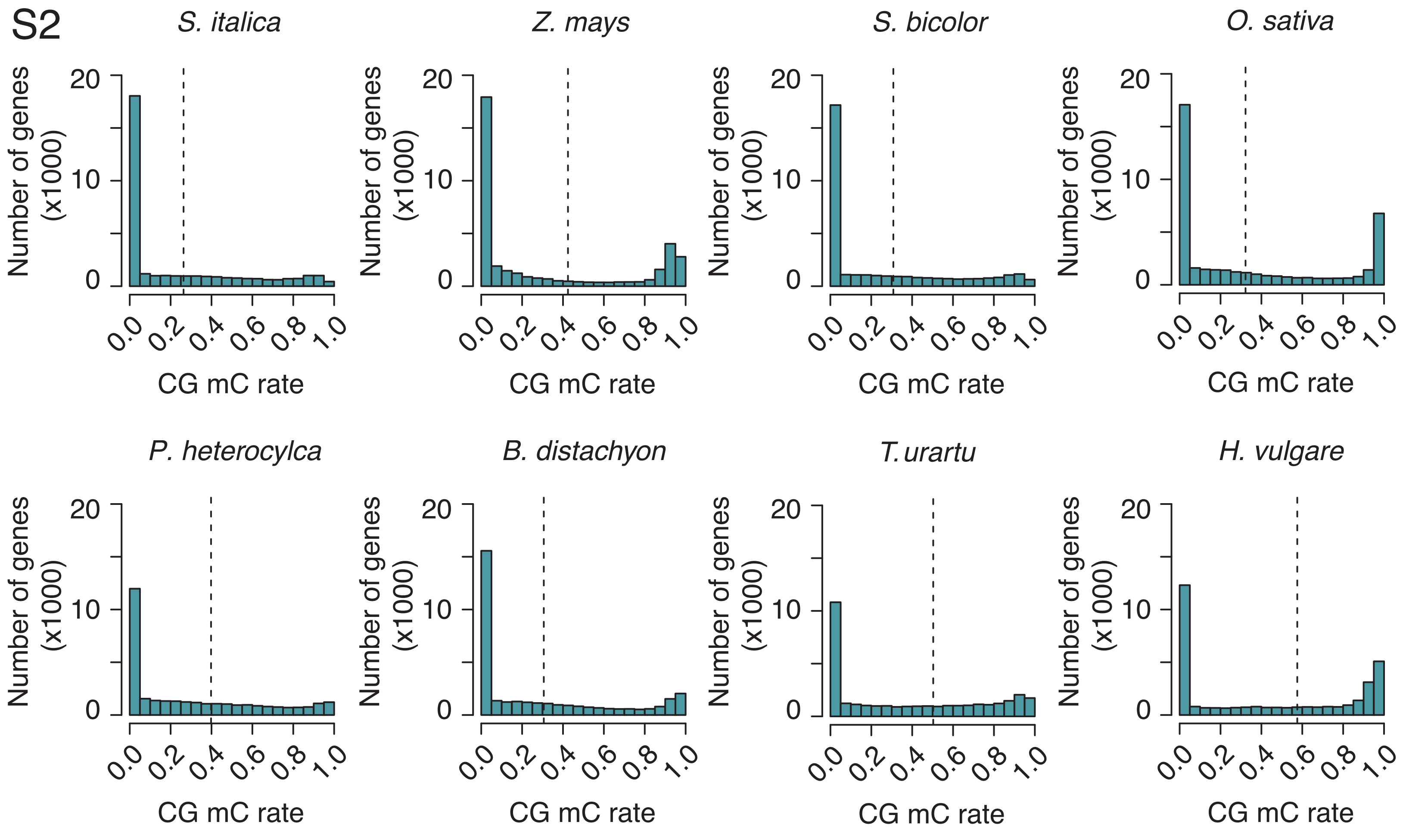
S2
S. italica
Z. mays
S. bicolor
O. sativa
P. heterocylca
B. distachyon
T. urartu
H. vulgare
Number of genes (x1000)
0
10
20
0.0
0.2
0.4
0.6
0.8
1.0
CG mC rate

S3

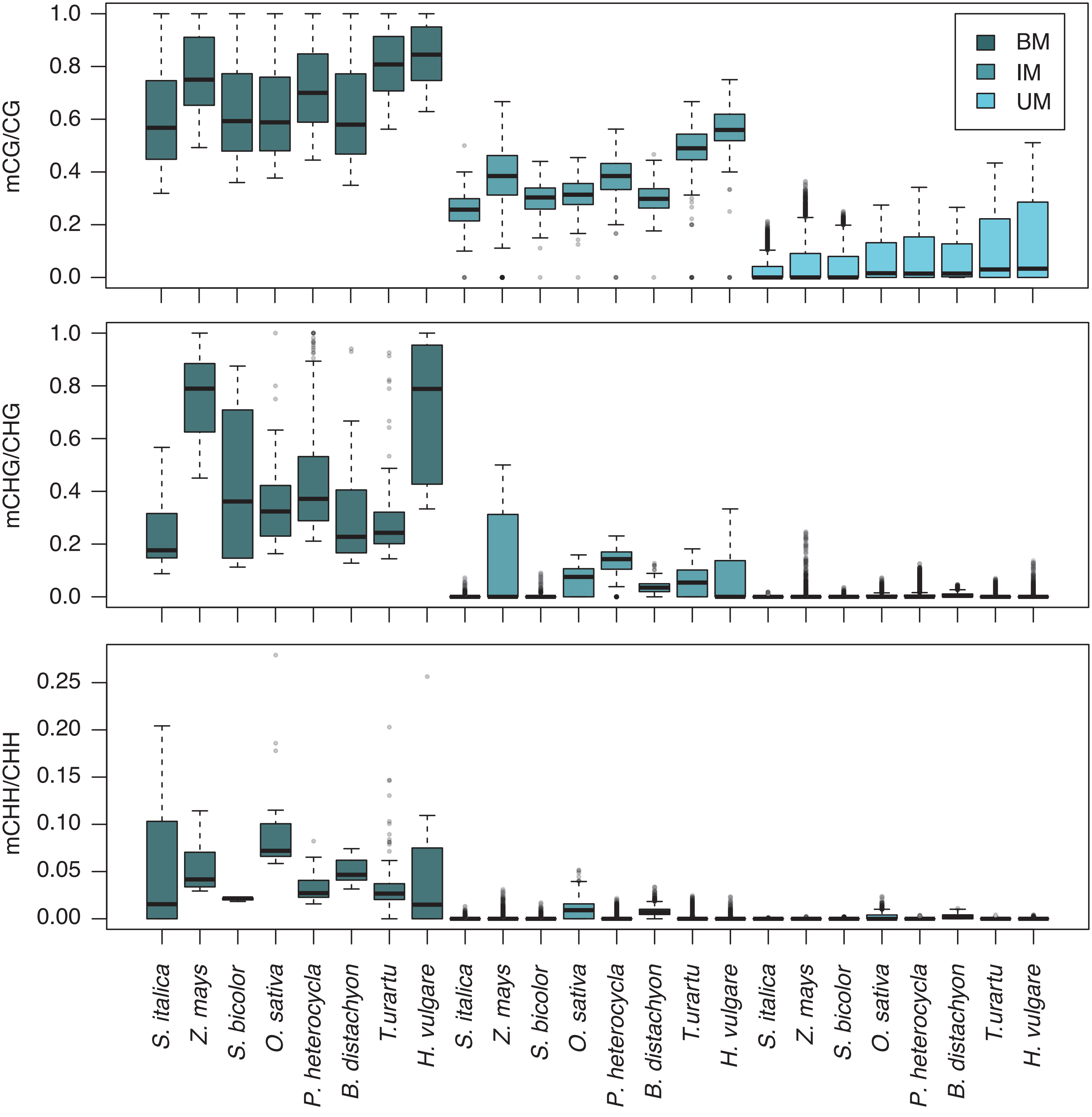

BM
IM
UM
mCG/CG
mCHG/CHG
mCHH/CHH
1.0
0.8
0.6
0.4
0.2
0.0
0.25
0.20
0.15
0.10
0.05
0.00
S. italica
Z. mays
S. bicolor
O. sativa
P. heterocycla
B. distachyon
T.urartu
H. vulgare

S4

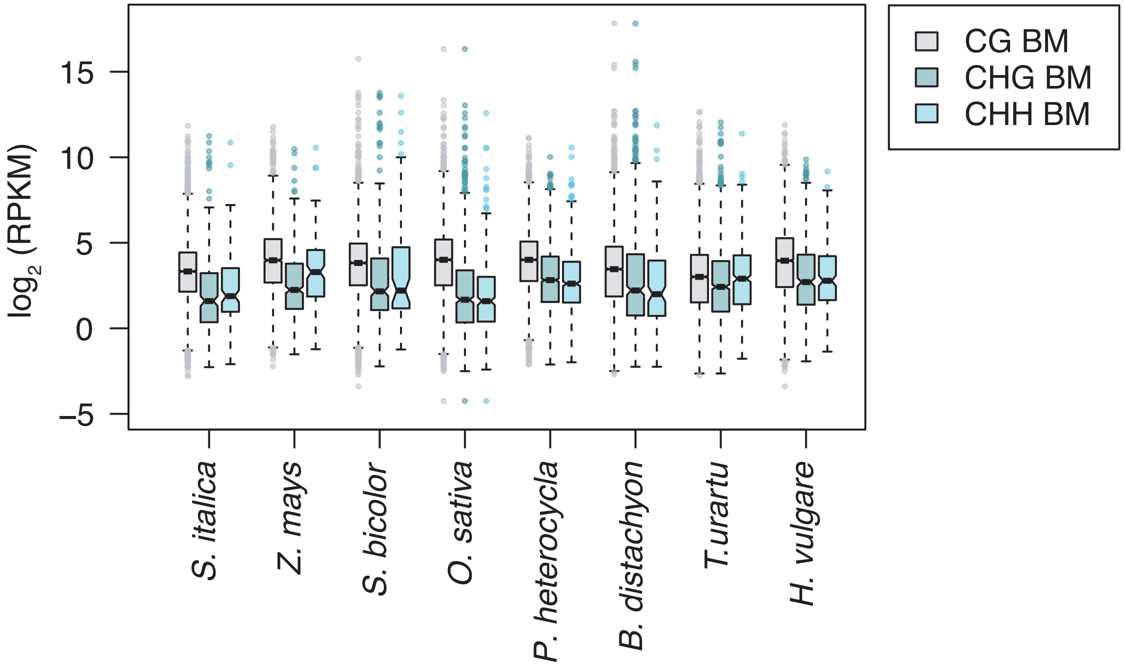

log2 (RPKM)
15
10
5
0
−5
CG BM
CHG BM
CHH BM
S. italica
Z. mays
S. bicolor
O. sativa
P. heterocycla
B. distachyon
T.urartu
H. vulgare

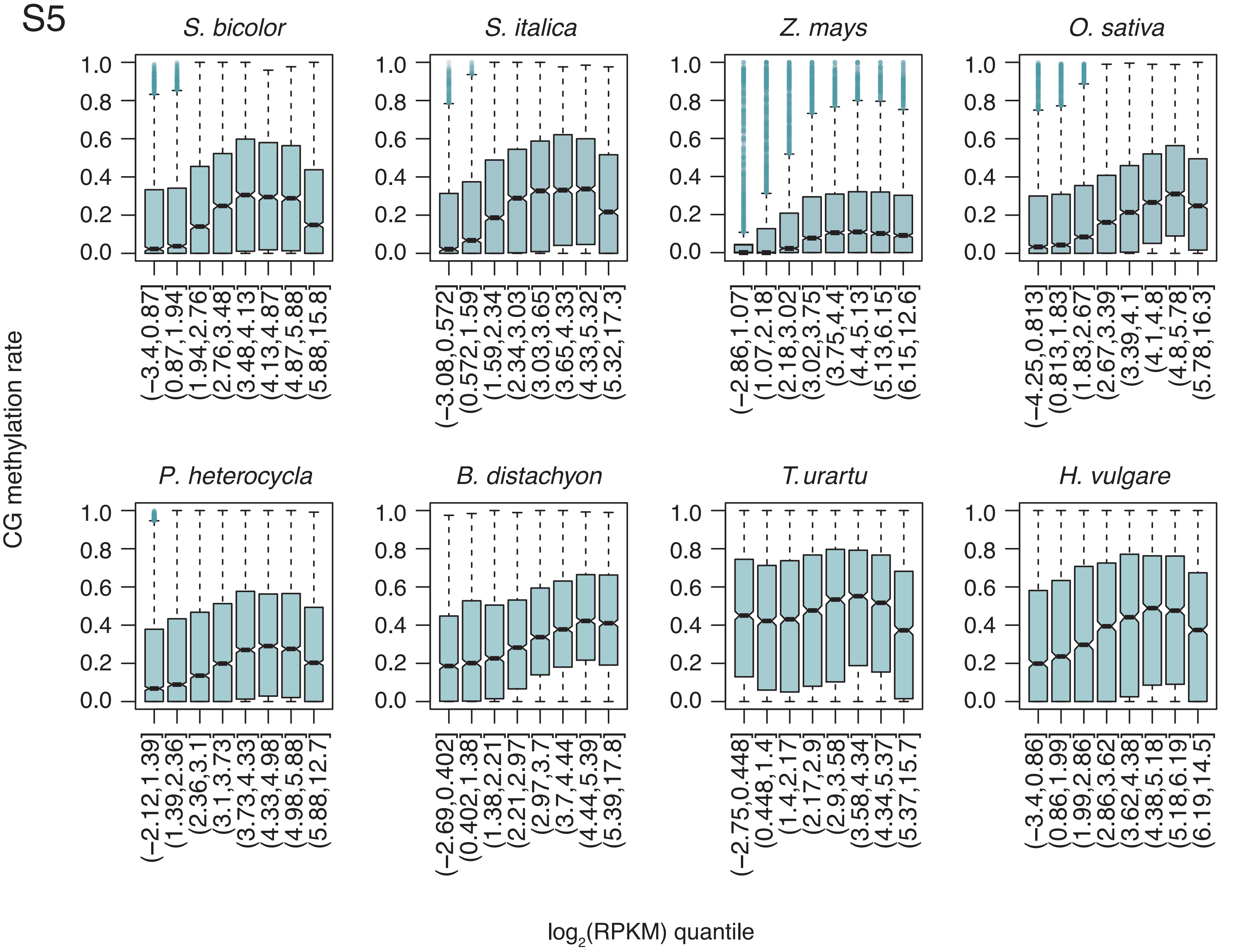
S5
S. bicolor
S. italica
Z. mays
O. sativa
P. heterocycla
B. distachyon
T. urartu
H. vulgare
CG methylation rate
log2(RPKM) quantile
0.0
0.2
0.4
0.6
0.8
1.0
(−3.4,0.87]
(0.87,1.94]
(1.94,2.76]
(2.76,3.48]
(3.48,4.13]
(4.13,4.87]
(4.87,5.88]
(5.88,15.8]
(−3.08,0.572]
(0.572,1.59]
(1.59,2.34]
(2.34,3.03]
(3.03,3.65]
(3.65,4.33]
(4.33,5.32]
(5.32,17.3]
(−2.86,1.07]
(1.07,2.18]
(2.18,3.02]
(3.02,3.75]
(3.75,4.4]
(4.4,5.13]
(5.13,6.15]
(6.15,12.6]
(−4.25,0.813]
(0.813,1.83]
(1.83,2.67]
(2.67,3.39]
(3.39,4.1]
(4.1,4.8]
(4.8,5.78]
(5.78,16.3]
(−2.12,1.39]
(1.39,2.36]
(2.36,3.1]
(3.1,3.73]
(3.73,4.33]
(4.33,4.98]
(4.98,5.88]
(5.88,12.7]
(−2.69,0.402]
(0.402,1.38]
(1.38,2.21]
(2.21,2.97]
(2.97,3.7]
(3.7,4.44]
(4.44,5.39]
(5.39,17.8]
(−2.75,0.448]
(0.448,1.4]
(1.4,2.17]
(2.17,2.9]
(2.9,3.58]
(3.58,4.34]
(4.34,5.37]
(5.37,15.7]
(−3.4,0.86]
(0.86,1.99]
(1.99,2.86]
(2.86,3.62]
(3.62,4.38]
(4.38,5.18]
(5.18,6.19]
(6.19,14.5]

S6

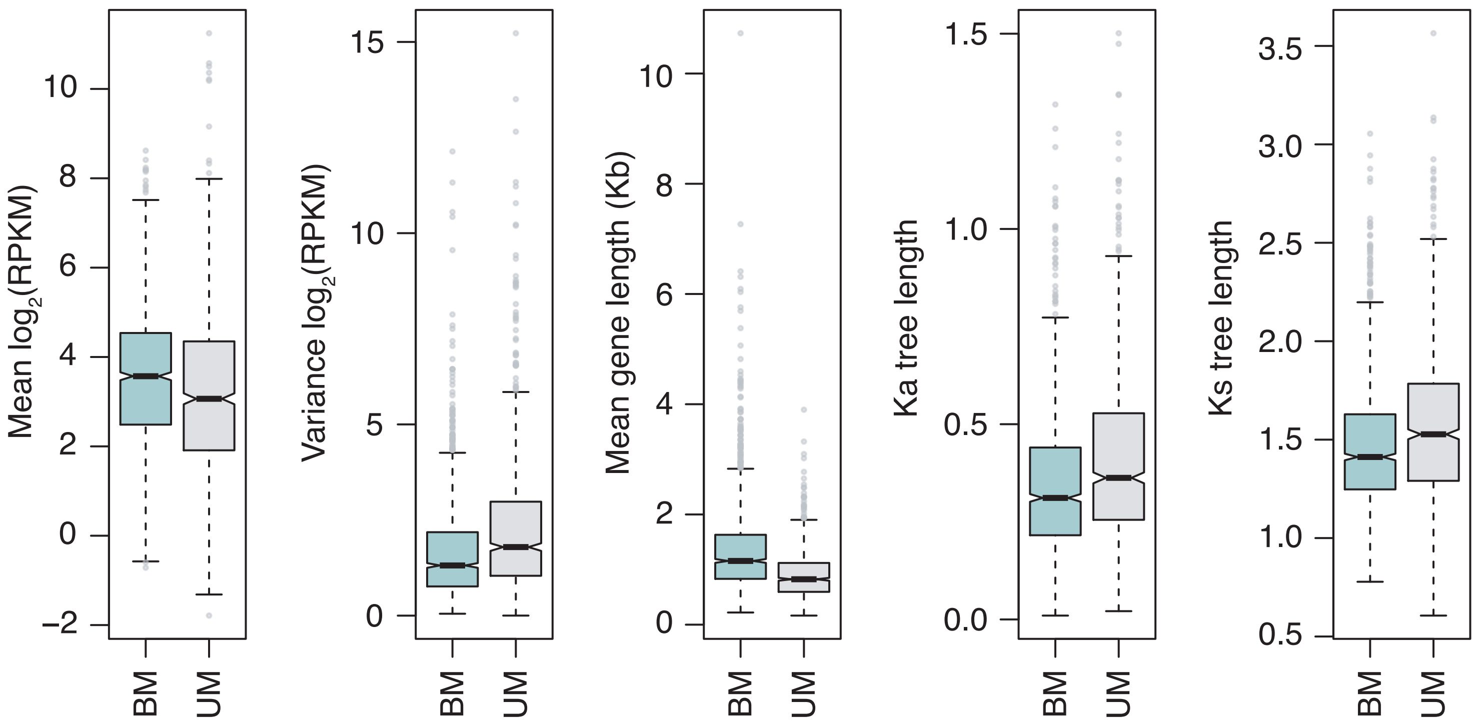

Mean log2(RPKM)
−2
0
2
4
6
8
10
BM
UM
Variance log2(RPKM)
0
5
10
15
BM
UM
Mean gene length (Kb)
0
2
4
6
8
10
BM
UM
Ka tree length
0.0
0.5
1.0
1.5
BM
UM
Ks tree length
0.5
1.0
1.5
2.0
2.5
3.0
3.5
BM
UM

**Table S1. The level of DNA methylation at CG, CHG, and CHH residues.**

| | Genome-wide | | | Genic (exon+intron) | | | CDS (exon only) | | |
|---|---|---|---|---|---|---|---|---|---|
| Species | CG | CHG | CHH | CG | CHG | CHH | CG | CHG | CHH |
| *Setaria italica* | 0.5774 | 0.3642 | 0.0168 | 0.2633 | 0.0434 | 0.0055 | 0.1997 | 0.0245 | 0.0009 |
| *Zea Mays* | 0.8987 | 0.8259 | 0.0173 | 0.4244 | 0.3420 | 0.0115 | 0.1588 | 0.1160 | 0.0024 |
| *Sorghum bicolor* | 0.6904 | 0.4540 | 0.0198 | 0.3082 | 0.0673 | 0.0105 | 0.1894 | 0.0222 | 0.0007 |
| *Oryza sativa* | 0.5589 | 0.3741 | 0.1038 | 0.3224 | 0.1302 | 0.0379 | 0.2603 | 0.1219 | 0.0193 |
| *Phyllostachys heterocycla* | 0.8384 | 0.7141 | 0.0162 | 0.3978 | 0.1673 | 0.0096 | 0.2742 | 0.0777 | 0.0024 |
| *Brachypodium distachyon* | 0.5723 | 0.3977 | 0.0522 | 0.3060 | 0.0831 | 0.0198 | 0.2228 | 0.0531 | 0.0072 |
| *Triticum urartu* | 0.9145 | 0.6521 | 0.0163 | 0.5028 | 0.1094 | 0.0095 | 0.3753 | 0.0493 | 0.0023 |
| *Hordeum vulgare* | 0.9441 | 0.7567 | 0.0092 | 0.5755 | 0.2127 | 0.0118 | 0.3516 | 0.0717 | 0.0017 |

**Table S2. Summary of annotated genes and orthologs methylated in each context.**

| | All Annotated Genes | | | | | | Orthologous Genes | | | | | |
|---|---|---|---|---|---|---|---|---|---|---|---|---|
| Species | Total | Total (no NA) | CG | CG only | CHG | CHH | Total | Total (no NA) | CG | CG only | CHG | CHH |
| *Setaria italica* | 34584 | 34041 | 10845 | 8394 | 2809 | 1186 | 2982 | 2860 | 1068 | 1060 | 11 | 9 |
| *Zea Mays* | 39502 | 37538 | 11298 | 2674 | 8756 | 1163 | 2982 | 2860 | 189 | 165 | 25 | 8 |
| *Sorghum bicolor* | 34211 | 33787 | 10481 | 7843 | 2937 | 393 | 2982 | 2860 | 868 | 862 | 10 | 3 |
| *Oryza sativa* | 42189 | 41530 | 16035 | 6511 | 10052 | 6201 | 2982 | 2860 | 666 | 651 | 22 | 19 |
| *Phyllostachys heterocycla* | 31987 | 31925 | 10191 | 6408 | 4335 | 1380 | 2982 | 2860 | 654 | 511 | 182 | 56 |
| *Brachypodium distachyon* | 34310 | 34176 | 11849 | 8242 | 4065 | 1476 | 2982 | 2860 | 859 | 838 | 25 | 12 |
| *Triticum urartu* | 33424 | 32692 | 11765 | 8445 | 4219 | 1421 | 2982 | 2860 | 1017 | 915 | 171 | 79 |
| *Hordeum vulgare* | 39734 | 33641 | 13324 | 6142 | 7345 | 1146 | 2982 | 2860 | 664 | 642 | 23 | 14 |

**Table S3. Orthologs with phylogenetic signal.**

| orthologID | *S. italica* | *Z. mays* | *S. bicolor* | *O. sativa* | *P. heterocycla* | *B. distachyon* | *T. urartu* | *H. vulgare* |
|---|---|---|---|---|---|---|---|---|
| orthologID_006242 | 0 | 0 | 0 | 0 | 0 | 1 | 1 | 1 |
| orthologID_006380 | 0 | 0 | 0 | 0 | 0 | 1 | 1 | 1 |
| orthologID_006589 | 0 | 0 | 0 | 0 | 0 | 1 | 1 | 1 |
| orthologID_006840 | 0 | 0 | 0 | 1 | 1 | 0 | 0 | 0 |
| orthologID_007698 | 0 | 0 | 0 | 1 | 1 | 1 | 1 | 1 |
| orthologID_007813 | 0 | 0 | 0 | 1 | 1 | 1 | 1 | 1 |
| orthologID_008143 | 0 | 0 | 0 | 0 | 0 | 1 | 1 | 1 |
| orthologID_008304 | 0 | 0 | 0 | 0 | 0 | 1 | 1 | 1 |
| orthologID_008387 | 0 | 0 | 0 | 0 | 0 | 1 | 1 | 1 |
| orthologID_008657 | 0 | 0 | 0 | 0 | 0 | 1 | 1 | 1 |
| orthologID_008677 | 0 | 0 | 0 | 1 | 1 | 1 | 1 | 1 |
| orthologID_008903 | 0 | 0 | 0 | 1 | 1 | 0 | 0 | 0 |
| orthologID_008959 | 0 | 0 | 0 | 0 | 0 | 1 | 1 | 1 |
| orthologID_009135 | 0 | 0 | 0 | 0 | 0 | 1 | 1 | 1 |
| orthologID_009140 | 1 | 1 | 1 | 0 | 0 | 0 | 0 | 0 |

**Table S4. Information on genomic resources.**

| Species | Genome | Annotation | Genome Resource |
|---|---|---|---|
| *Setaria italica* | Sitalica_312_v2.fa | Sitalica_312_v2.2.gene.gff3 | https://phytozome.jgi.doe.gov |
| *Zea Mays* | Zea_mays.AGPv4.dna.chromosome.all.fa | Zea_mays.AGPv4.cds.all.fa | http://plants.ensembl.org/index.html |
| *Sorghum bicolor* | Sbicolor_313_v3.0.fa | Sbicolor_313_v3.1.gene.gff3 | https://phytozome.jgi.doe.gov |
| *Oryza sativa* | Osativa_323_v7.0.fa | Osativa_323_v7.0.gene.gff3 | https://phytozome.jgi.doe.gov |
| *Phyllostachys heterocycla* | P_heterocycla_v1.0.Scaffolds.fa | P_heterocycla_v1.0.genemodel.gff | http://server.ncgr.ac.cn/bamboo/ |
| *Brachypodium distachyon* | Bdistachyon_314_v3.0.fa | Bdistachyon_314_v3.1.gene.gff3 | https://phytozome.jgi.doe.gov |
| *Triticum urartu* | Triticum_urartu.ASM34745v1.31.dna.genome.fa | Triticum_urartu.GCA_000347455.1.30.cds.all.fa | http://plants.ensembl.org/index.html |
| *Hordeum vulgare* | 150831_barley_pseudomolecules.fa | Hv_IBSC_PGSB_r1_HighConf.gtf | https://webblast.ipk-gatersleben.de |

**Table S5. Rate of false positive methylation errors for bisulfite sequencing libraries.**

| Sample | Species | Replicate | Method | CG FMR | CHG FMR | CHH FMR |
|---|---|---|---|---|---|---|
| P01 | *H. vulgare* | A | λ DNA | 0.001246019 | 0.001511980 | 0.001444878 |
| P02 | *H. vulgare* | B | λ DNA | 0.001197072 | 0.001442373 | 0.001368112 |
| P03 | *H. vulgare* | C | λ DNA | 0.001269014 | 0.001490355 | 0.001425880 |
| P04 | *S. italica* | A | λ DNA | 0.001216545 | 0.001488870 | 0.001368322 |
| P05 | *S. italica* | B | λ DNA | 0.001100060 | 0.001542065 | 0.001306192 |
| P06 | *S. italica* | C | λ DNA | 0.001622711 | 0.001741653 | 0.001697245 |
| P07 | *S. bicolor* | A | λ DNA | 0.001357007 | 0.001633894 | 0.001567219 |
| P08 | *S. bicolor* | B | λ DNA | 0.001241004 | 0.001539483 | 0.001460505 |
| P09 | *S. bicolor* | C | λ DNA | 0.001118252 | 0.001260037 | 0.001239415 |
| P10 | *P. heterocycla* | A | λ DNA | 0.001270735 | 0.001480494 | 0.001472576 |
| P11 | *P. heterocycla* | B | λ DNA | 0.001426921 | 0.001637409 | 0.001610093 |
| P12 | *P. heterocycla* | C | λ DNA | 0.001168522 | 0.001442413 | 0.001372062 |
| P13 | *T. urartu* | A | λ DNA | 0.001224012 | 0.001450066 | 0.001366496 |
| P14 | *T. urartu* | B | λ DNA | 0.001050812 | 0.001237936 | 0.001187400 |
| P15 | *T. urartu* | C | λ DNA | 0.001094182 | 0.001302801 | 0.001247631 |
| P18 | *O. sativa* | A | λ DNA | 0.005033481 | 0.005174992 | 0.005559796 |
| P19 | *O. sativa* | B | λ DNA | 0.004124629 | 0.004343137 | 0.004568394 |
| P20 | *O. sativa* | C | λ DNA | 0.004730641 | 0.004818811 | 0.005086037 |
| P22 | *B. distachyon* | A | Chloroplast DNA | 0.002710838 | 0.002878814 | 0.002627274 |
| P23 | *B. distachyon* | B | Chloroplast DNA | 0.003862086 | 0.003401056 | 0.002822095 |
| P24 | *B. distachyon* | C | Chloroplast DNA | 0.002962997 | 0.002422593 | 0.002922155 |
| P25 | *Z. mays* | A | Chloroplast DNA | 0.010563115 | 0.010949488 | 0.006790905 |

**Table S6. Bisulfite sequencing statistics.**

| Species | Replicate | Lane | SRA Run ID | SRA file ID | File name | Number of trimmed PE reads | Number of aligned PE reads |
|---|---|---|---|---|---|---|---|
| *H. vulgare* | A | 1 | SRR4099340 | Hvulgare_bisulfite_repA.1 | R259-L2-P01-CAACTA_trimmed_1P.fq.gz | 19225671 | 6045259 |
| *H. vulgare* | B | 1 | SRR4099341 | Hvulgare_bisulfite_repB.1 | R259-L2-P02-CACCGG_trimmed_1P.fq.gz | 16666869 | 5116811 |
| *H. vulgare* | C | 1 | SRR4099422 | Hvulgare_bisulfite_repC.1 | R259-L2-P03-CACGAT_trimmed_1P.fq.gz | 18627431 | 5803830 |
| *H. vulgare* | A | 2 | SRR4099433 | Hvulgare_bisulfite_repA.2 | R269-L1-P01-CAACTA_trimmed_1P.fq.gz | 16687836 | 11635578 |
| *H. vulgare* | B | 2 | SRR4099444 | Hvulgare_bisulfite_repB.2 | R269-L1-P02-CACCGG_trimmed_1P.fq.gz | 13037462 | 9078008 |
| *H. vulgare* | C | 2 | SRR4099475 | Hvulgare_bisulfite_repC.2 | R269-L1-P03-CACGAT_trimmed_1P.fq.gz | 17054734 | 11891245 |
| *H. vulgare* | A | 3 | SRR4099486 | Hvulgare_bisulfite_repA.3 | R269-L2-P01-CAACTA_trimmed_1P.fq.gz | 16355540 | 11395929 |
| *H. vulgare* | B | 3 | SRR4099497 | Hvulgare_bisulfite_repB.3 | R269-L2-P02-CACCGG_trimmed_1P.fq.gz | 12819865 | 8918607 |
| *H. vulgare* | C | 3 | SRR4099508 | Hvulgare_bisulfite_repC.3 | R269-L2-P03-CACGAT_trimmed_1P.fq.gz | 16728521 | 11646795 |
| *H. vulgare* | A | 4 | SRR4099519 | Hvulgare_bisulfite_repA.4 | R270-L1-P01-CAACTA_trimmed_1P.fq.gz | 17601572 | 12258100 |
| *H. vulgare* | B | 4 | SRR4099342 | Hvulgare_bisulfite_repB.4 | R270-L1-P02-CACCGG_trimmed_1P.fq.gz | 13278256 | 9230528 |
| *H. vulgare* | C | 4 | SRR4099353 | Hvulgare_bisulfite_repC.4 | R270-L1-P03-CACGAT_trimmed_1P.fq.gz | 17692400 | 12318399 |
| *H. vulgare* | A | 5 | SRR4099364 | Hvulgare_bisulfite_repA.5 | R270-L2-P01-CAACTA_trimmed_1P.fq.gz | 17674940 | 12311109 |
| *H. vulgare* | B | 5 | SRR4099395 | Hvulgare_bisulfite_repB.5 | R270-L2-P02-CACCGG_trimmed_1P.fq.gz | 13329683 | 9265241 |
| *H. vulgare* | C | 5 | SRR4099406 | Hvulgare_bisulfite_repC.5 | R270-L2-P03-CACGAT_trimmed_1P.fq.gz | 17753504 | 12364839 |
| *H. vulgare* | A | 6 | SRR4099417 | Hvulgare_bisulfite_repA.6 | R271-L1-P01-CAACTA_trimmed_1P.fq.gz | 16496756 | 11473294 |
| *H. vulgare* | B | 6 | SRR4099418 | Hvulgare_bisulfite_repB.6 | R271-L1-P02-CACCGG_trimmed_1P.fq.gz | 11858420 | 8233391 |
| *H. vulgare* | C | 6 | SRR4099419 | Hvulgare_bisulfite_repC.6 | R271-L1-P03-CACGAT_trimmed_1P.fq.gz | 16285671 | 11329084 |
| *H. vulgare* | A | 7 | SRR4099420 | Hvulgare_bisulfite_repA.7 | R271-L2-P01-CAACTA_trimmed_1P.fq.gz | 16691035 | 11619544 |
| *H. vulgare* | B | 7 | SRR4099421 | Hvulgare_bisulfite_repB.7 | R271-L2-P02-CACCGG_trimmed_1P.fq.gz | 11987345 | 8328752 |
| *H. vulgare* | C | 7 | SRR4099423 | Hvulgare_bisulfite_repC.7 | R271-L2-P03-CACGAT_trimmed_1P.fq.gz | 16475581 | 11476377 |
| *H. vulgare* | A | 8 | SRR4099424 | Hvulgare_bisulfite_repA.8 | R272-L1-P01-CAACTA_trimmed_1P.fq.gz | 17400640 | 12090992 |
| *H. vulgare* | B | 8 | SRR4099425 | Hvulgare_bisulfite_repB.8 | R272-L1-P02-CACCGG_trimmed_1P.fq.gz | 12300009 | 8527726 |
| *H. vulgare* | C | 8 | SRR4099426 | Hvulgare_bisulfite_repC.8 | R272-L1-P03-CACGAT_trimmed_1P.fq.gz | 17375742 | 12068214 |
| *H. vulgare* | A | 9 | SRR4099427 | Hvulgare_bisulfite_repA.9 | R272-L2-P01-CAACTA_trimmed_1P.fq.gz | 16862917 | 11700336 |
| *H. vulgare* | B | 9 | SRR4099428 | Hvulgare_bisulfite_repB.9 | R272-L2-P02-CACCGG_trimmed_1P.fq.gz | 11912806 | 8245150 |
| *H. vulgare* | C | 9 | SRR4099429 | Hvulgare_bisulfite_repC.9 | R272-L2-P03-CACGAT_trimmed_1P.fq.gz | 16857866 | 11684791 |
| *H. vulgare* | A | 10 | SRR4099430 | Hvulgare_bisulfite_repA.10 | R278-L1-P01-CAACTA_trimmed_1P.fq.gz | 17381631 | 11091644 |
| *H. vulgare* | B | 10 | SRR4099431 | Hvulgare_bisulfite_repB.10 | R278-L1-P02-CACCGG_trimmed_1P.fq.gz | 14900645 | 9469700 |
| *H. vulgare* | C | 10 | SRR4099432 | Hvulgare_bisulfite_repC.10 | R278-L1-P03-CACGAT_trimmed_1P.fq.gz | 16881125 | 10730006 |
| *P. heterocycla* | A | 1 | SRR4099434 | Pheterocylca_bisulfite_repA.1 | R259-L2-P10-CTAGCT_trimmed_1P.fq.gz | 5113204 | 1143675 |
| *P. heterocycla* | B | 1 | SRR4099435 | Pheterocylca_bisulfite_repB.1 | R259-L2-P11-CTATAC_trimmed_1P.fq.gz | 7742368 | 2752317 |
| *P. heterocycla* | C | 1 | SRR4099436 | Pheterocylca_bisulfite_repC.1 | R259-L2-P12-CTCAGA_trimmed_1P.fq.gz | 6447048 | 2387354 |
| *P. heterocycla* | A | 2 | SRR4099437 | Pheterocylca_bisulfite_repA.2 | R269-L1-P10-CTAGCT_trimmed_1P.fq.gz | 4784488 | 3571106 |
| *P. heterocycla* | B | 2 | SRR4099438 | Pheterocylca_bisulfite_repB.2 | R269-L1-P11-CTATAC_trimmed_1P.fq.gz | 6744146 | 5391514 |

| | | | | | | | |
|---|---|---|---|---|---|---|---|
| *P. heterocycla* | C | 2 | SRR4099439 | Pheterocylca_bisulfite_repC.2 | R269-L1-P12-CTCAGA_trimmed_1P.fq.gz | 5239480 | 4212078 |
| *P. heterocycla* | A | 3 | SRR4099440 | Pheterocylca_bisulfite_repA.3 | R269-L2-P10-CTAGCT_trimmed_1P.fq.gz | 4699620 | 3508054 |
| *P. heterocycla* | B | 3 | SRR4099441 | Pheterocylca_bisulfite_repB.3 | R269-L2-P11-CTATAC_trimmed_1P.fq.gz | 6609581 | 5283201 |
| *P. heterocycla* | C | 3 | SRR4099442 | Pheterocylca_bisulfite_repC.3 | R269-L2-P12-CTCAGA_trimmed_1P.fq.gz | 5127453 | 4125154 |
| *P. heterocycla* | A | 4 | SRR4099443 | Pheterocylca_bisulfite_repA.4 | R270-L1-P10-CTAGCT_trimmed_1P.fq.gz | 4793854 | 3574539 |
| *P. heterocycla* | B | 4 | SRR4099445 | Pheterocylca_bisulfite_repB.4 | R270-L1-P11-CTATAC_trimmed_1P.fq.gz | 6906711 | 5507520 |
| *P. heterocycla* | C | 4 | SRR4099446 | Pheterocylca_bisulfite_repC.4 | R270-L1-P12-CTCAGA_trimmed_1P.fq.gz | 5793103 | 4652179 |
| *P. heterocycla* | A | 5 | SRR4099447 | Pheterocylca_bisulfite_repA.5 | R270-L2-P10-CTAGCT_trimmed_1P.fq.gz | 4797059 | 3574752 |
| *P. heterocycla* | B | 5 | SRR4099448 | Pheterocylca_bisulfite_repB.5 | R270-L2-P11-CTATAC_trimmed_1P.fq.gz | 6931374 | 5524620 |
| *P. heterocycla* | C | 5 | SRR4099449 | Pheterocylca_bisulfite_repC.5 | R270-L2-P12-CTCAGA_trimmed_1P.fq.gz | 5820557 | 4673375 |
| *P. heterocycla* | A | 6 | SRR4099470 | Pheterocylca_bisulfite_repA.6 | R271-L1-P10-CTAGCT_trimmed_1P.fq.gz | 4782597 | 3556706 |
| *P. heterocycla* | B | 6 | SRR4099471 | Pheterocylca_bisulfite_repB.6 | R271-L1-P11-CTATAC_trimmed_1P.fq.gz | 6560903 | 5237167 |
| *P. heterocycla* | C | 6 | SRR4099472 | Pheterocylca_bisulfite_repC.6 | R271-L1-P12-CTCAGA_trimmed_1P.fq.gz | 5454481 | 4386086 |
| *P. heterocycla* | A | 7 | SRR4099473 | Pheterocylca_bisulfite_repA.7 | R271-L2-P10-CTAGCT_trimmed_1P.fq.gz | 4814560 | 3583491 |
| *P. heterocycla* | B | 7 | SRR4099474 | Pheterocylca_bisulfite_repB.7 | R271-L2-P11-CTATAC_trimmed_1P.fq.gz | 6622852 | 5286935 |
| *P. heterocycla* | C | 7 | SRR4099476 | Pheterocylca_bisulfite_repC.7 | R271-L2-P12-CTCAGA_trimmed_1P.fq.gz | 5531414 | 4446823 |
| *P. heterocycla* | A | 8 | SRR4099477 | Pheterocylca_bisulfite_repA.8 | R272-L1-P10-CTAGCT_trimmed_1P.fq.gz | 4768540 | 3543821 |
| *P. heterocycla* | B | 8 | SRR4099478 | Pheterocylca_bisulfite_repB.8 | R272-L1-P11-CTATAC_trimmed_1P.fq.gz | 6784261 | 5398976 |
| *P. heterocycla* | C | 8 | SRR4099479 | Pheterocylca_bisulfite_repC.8 | R272-L1-P12-CTCAGA_trimmed_1P.fq.gz | 5754516 | 4614788 |
| *P. heterocycla* | A | 9 | SRR4099480 | Pheterocylca_bisulfite_repA.9 | R272-L2-P10-CTAGCT_trimmed_1P.fq.gz | 4643415 | 3448322 |
| *P. heterocycla* | B | 9 | SRR4099481 | Pheterocylca_bisulfite_repB.9 | R272-L2-P11-CTATAC_trimmed_1P.fq.gz | 6584989 | 5241182 |
| *P. heterocycla* | C | 9 | SRR4099482 | Pheterocylca_bisulfite_repC.9 | R272-L2-P12-CTCAGA_trimmed_1P.fq.gz | 5574861 | 4466043 |
| *P. heterocycla* | A | 10 | SRR4099483 | Pheterocylca_bisulfite_repA.10 | R278-L1-P10-CTAGCT_trimmed_1P.fq.gz | 4657236 | 3213356 |
| *P. heterocycla* | B | 10 | SRR4099484 | Pheterocylca_bisulfite_repB.10 | R278-L1-P11-CTATAC_trimmed_1P.fq.gz | 6968912 | 5114606 |
| *P. heterocycla* | C | 10 | SRR4099485 | Pheterocylca_bisulfite_repC.10 | R278-L1-P12-CTCAGA_trimmed_1P.fq.gz | 5838731 | 4316189 |
| *S. bicolor* | A | 1 | SRR4099487 | Sbicolor_bisulfite_repA.1 | R259-L2-P07-CATTTT_trimmed_1P.fq.gz | 2673786 | 962047 |
| *S. bicolor* | B | 1 | SRR4099488 | Sbicolor_bisulfite_repB.1 | R259-L2-P08-CCAACA_trimmed_1P.fq.gz | 2280865 | 811971 |
| *S. bicolor* | C | 1 | SRR4099489 | Sbicolor_bisulfite_repC.1 | R259-L2-P09-CGGAAT_trimmed_1P.fq.gz | 2054599 | 712180 |
| *S. bicolor* | A | 2 | SRR4099490 | Sbicolor_bisulfite_repA.2 | R269-L1-P07-CATTTT_trimmed_1P.fq.gz | 2335360 | 1829634 |
| *S. bicolor* | B | 2 | SRR4099491 | Sbicolor_bisulfite_repB.2 | R269-L1-P08-CCAACA_trimmed_1P.fq.gz | 1939352 | 1489810 |
| *S. bicolor* | C | 2 | SRR4099492 | Sbicolor_bisulfite_repC.2 | R269-L1-P09-CGGAAT_trimmed_1P.fq.gz | 1610256 | 1238325 |
| *S. bicolor* | A | 3 | SRR4099493 | Sbicolor_bisulfite_repA.3 | R269-L2-P07-CATTTT_trimmed_1P.fq.gz | 2292601 | 1794918 |
| *S. bicolor* | B | 3 | SRR4099494 | Sbicolor_bisulfite_repB.3 | R269-L2-P08-CCAACA_trimmed_1P.fq.gz | 1901665 | 1460588 |
| *S. bicolor* | C | 3 | SRR4099495 | Sbicolor_bisulfite_repC.3 | R269-L2-P09-CGGAAT_trimmed_1P.fq.gz | 1575937 | 1211414 |
| *S. bicolor* | A | 4 | SRR4099496 | Sbicolor_bisulfite_repA.4 | R270-L1-P07-CATTTT_trimmed_1P.fq.gz | 2443135 | 1908650 |
| *S. bicolor* | B | 4 | SRR4099498 | Sbicolor_bisulfite_repB.4 | R270-L1-P08-CCAACA_trimmed_1P.fq.gz | 2053947 | 1571137 |
| *S. bicolor* | C | 4 | SRR4099499 | Sbicolor_bisulfite_repC.4 | R270-L1-P09-CGGAAT_trimmed_1P.fq.gz | 1667494 | 1279857 |
| *S. bicolor* | A | 5 | SRR4099500 | Sbicolor_bisulfite_repA.5 | R270-L2-P07-CATTTT_trimmed_1P.fq.gz | 2455578 | 1917904 |

| | | | | | | | |
|---|---|---|---|---|---|---|---|
| *S. bicolor* | B | 5 | SRR4099501 | Sbicolor_bisulfite_repB.5 | R270-L2-P08-CCAACA_trimmed_1P.fq.gz | 2057351 | 1576972 |
| *S. bicolor* | C | 5 | SRR4099502 | Sbicolor_bisulfite_repC.5 | R270-L2-P09-CGGAAT_trimmed_1P.fq.gz | 1674581 | 1284725 |
| *S. bicolor* | A | 6 | SRR4099503 | Sbicolor_bisulfite_repA.6 | R271-L1-P07-CATTTT_trimmed_1P.fq.gz | 2284978 | 1784298 |
| *S. bicolor* | B | 6 | SRR4099504 | Sbicolor_bisulfite_repB.6 | R271-L1-P08-CCAACA_trimmed_1P.fq.gz | 1924201 | 1475010 |
| *S. bicolor* | C | 6 | SRR4099505 | Sbicolor_bisulfite_repC.6 | R271-L1-P09-CGGAAT_trimmed_1P.fq.gz | 1575603 | 1209371 |
| *S. bicolor* | A | 7 | SRR4099506 | Sbicolor_bisulfite_repA.7 | R271-L2-P07-CATTTT_trimmed_1P.fq.gz | 2306786 | 1801231 |
| *S. bicolor* | B | 7 | SRR4099507 | Sbicolor_bisulfite_repB.7 | R271-L2-P08-CCAACA_trimmed_1P.fq.gz | 1944044 | 1492950 |
| *S. bicolor* | C | 7 | SRR4099509 | Sbicolor_bisulfite_repC.7 | R271-L2-P09-CGGAAT_trimmed_1P.fq.gz | 1588348 | 1219988 |
| *S. bicolor* | A | 8 | SRR4099510 | Sbicolor_bisulfite_repA.8 | R272-L1-P07-CATTTT_trimmed_1P.fq.gz | 2414804 | 1882311 |
| *S. bicolor* | B | 8 | SRR4099511 | Sbicolor_bisulfite_repB.8 | R272-L1-P08-CCAACA_trimmed_1P.fq.gz | 2023102 | 1546168 |
| *S. bicolor* | C | 8 | SRR4099512 | Sbicolor_bisulfite_repC.8 | R272-L1-P09-CGGAAT_trimmed_1P.fq.gz | 1634709 | 1250601 |
| *S. bicolor* | A | 9 | SRR4099513 | Sbicolor_bisulfite_repA.9 | R272-L2-P07-CATTTT_trimmed_1P.fq.gz | 2343421 | 1827072 |
| *S. bicolor* | B | 9 | SRR4099514 | Sbicolor_bisulfite_repB.9 | R272-L2-P08-CCAACA_trimmed_1P.fq.gz | 1970551 | 1504604 |
| *S. bicolor* | C | 9 | SRR4099515 | Sbicolor_bisulfite_repC.9 | R272-L2-P09-CGGAAT_trimmed_1P.fq.gz | 1592575 | 1218829 |
| *S. bicolor* | A | 10 | SRR4099516 | Sbicolor_bisulfite_repA.10 | R278-L1-P07-CATTTT_trimmed_1P.fq.gz | 2412194 | 1731108 |
| *S. bicolor* | B | 10 | SRR4099517 | Sbicolor_bisulfite_repB.10 | R278-L1-P08-CCAACA_trimmed_1P.fq.gz | 2069829 | 1457616 |
| *S. bicolor* | C | 10 | SRR4099518 | Sbicolor_bisulfite_repC.10 | R278-L1-P09-CGGAAT_trimmed_1P.fq.gz | 1849744 | 1279396 |
| *S. italica* | A | 1 | SRR4099520 | Sitalica_bisulfite_repA.1 | R259-L2-P04-CACTCA_trimmed_1P.fq.gz | 1473405 | 508257 |
| *S. italica* | B | 1 | SRR4099521 | Sitalica_bisulfite_repB.1 | R259-L2-P05-CAGGCG_trimmed_1P.fq.gz | 1346472 | 503868 |
| *S. italica* | C | 1 | SRR4099522 | Sitalica_bisulfite_repC.1 | R259-L2-P06-CATGGC_trimmed_1P.fq.gz | 1336939 | 502608 |
| *S. italica* | A | 2 | SRR4099523 | Sitalica_bisulfite_repA.2 | R269-L1-P04-CACTCA_trimmed_1P.fq.gz | 1229028 | 964117 |
| *S. italica* | B | 2 | SRR4099524 | Sitalica_bisulfite_repB.2 | R269-L1-P05-CAGGCG_trimmed_1P.fq.gz | 967335 | 803385 |
| *S. italica* | C | 2 | SRR4099525 | Sitalica_bisulfite_repC.2 | R269-L1-P06-CATGGC_trimmed_1P.fq.gz | 1177172 | 977449 |
| *S. italica* | A | 3 | SRR4099526 | Sitalica_bisulfite_repA.3 | R269-L2-P04-CACTCA_trimmed_1P.fq.gz | 1203668 | 944180 |
| *S. italica* | B | 3 | SRR4099547 | Sitalica_bisulfite_repB.3 | R269-L2-P05-CAGGCG_trimmed_1P.fq.gz | 949550 | 789357 |
| *S. italica* | C | 3 | SRR4099548 | Sitalica_bisulfite_repC.3 | R269-L2-P06-CATGGC_trimmed_1P.fq.gz | 1156499 | 960349 |
| *S. italica* | A | 4 | SRR4099549 | Sitalica_bisulfite_repA.4 | R270-L1-P04-CACTCA_trimmed_1P.fq.gz | 1304388 | 1022537 |
| *S. italica* | B | 4 | SRR4099343 | Sitalica_bisulfite_repB.4 | R270-L1-P05-CAGGCG_trimmed_1P.fq.gz | 970271 | 799749 |
| *S. italica* | C | 4 | SRR4099344 | Sitalica_bisulfite_repC.4 | R270-L1-P06-CATGGC_trimmed_1P.fq.gz | 1193464 | 986998 |
| *S. italica* | A | 5 | SRR4099345 | Sitalica_bisulfite_repA.5 | R270-L2-P04-CACTCA_trimmed_1P.fq.gz | 1309166 | 1024625 |
| *S. italica* | B | 5 | SRR4099346 | Sitalica_bisulfite_repB.5 | R270-L2-P05-CAGGCG_trimmed_1P.fq.gz | 976126 | 804296 |
| *S. italica* | C | 5 | SRR4099347 | Sitalica_bisulfite_repC.5 | R270-L2-P06-CATGGC_trimmed_1P.fq.gz | 1198041 | 989686 |
| *S. italica* | A | 6 | SRR4099348 | Sitalica_bisulfite_repA.6 | R271-L1-P04-CACTCA_trimmed_1P.fq.gz | 1239339 | 975510 |
| *S. italica* | B | 6 | SRR4099349 | Sitalica_bisulfite_repB.6 | R271-L1-P05-CAGGCG_trimmed_1P.fq.gz | 789600 | 654717 |
| *S. italica* | C | 6 | SRR4099350 | Sitalica_bisulfite_repC.6 | R271-L1-P06-CATGGC_trimmed_1P.fq.gz | 1000050 | 829784 |
| *S. italica* | A | 7 | SRR4099351 | Sitalica_bisulfite_repA.7 | R271-L2-P04-CACTCA_trimmed_1P.fq.gz | 1254076 | 985811 |
| *S. italica* | B | 7 | SRR4099352 | Sitalica_bisulfite_repB.7 | R271-L2-P05-CAGGCG_trimmed_1P.fq.gz | 797488 | 661084 |
| *S. italica* | C | 7 | SRR4099354 | Sitalica_bisulfite_repC.7 | R271-L2-P06-CATGGC_trimmed_1P.fq.gz | 1015739 | 842945 |

| | | | | | | | |
|---|---|---|---|---|---|---|---|
| *S. italica* | A | 8 | SRR4099355 | Sitalica_bisulfite_repA.8 | R272-L1-P04-CACTCA_trimmed_1P.fq.gz | 1290388 | 1008212 |
| *S. italica* | B | 8 | SRR4099356 | Sitalica_bisulfite_repB.8 | R272-L1-P05-CAGGCG_trimmed_1P.fq.gz | 853398 | 699946 |
| *S. italica* | C | 8 | SRR4099357 | Sitalica_bisulfite_repC.8 | R272-L1-P06-CATGGC_trimmed_1P.fq.gz | 1092555 | 901279 |
| *S. italica* | A | 9 | SRR4099358 | Sitalica_bisulfite_repA.9 | R272-L2-P04-CACTCA_trimmed_1P.fq.gz | 1254186 | 979589 |
| *S. italica* | B | 9 | SRR4099359 | Sitalica_bisulfite_repB.9 | R272-L2-P05-CAGGCG_trimmed_1P.fq.gz | 826087 | 678430 |
| *S. italica* | C | 9 | SRR4099360 | Sitalica_bisulfite_repC.9 | R272-L2-P06-CATGGC_trimmed_1P.fq.gz | 1058388 | 874703 |
| *S. italica* | A | 10 | SRR4099361 | Sitalica_bisulfite_repA.10 | R278-L1-P04-CACTCA_trimmed_1P.fq.gz | 1325725 | 944957 |
| *S. italica* | B | 10 | SRR4099362 | Sitalica_bisulfite_repB.10 | R278-L1-P05-CAGGCG_trimmed_1P.fq.gz | 1209851 | 906006 |
| *S. italica* | C | 10 | SRR4099363 | Sitalica_bisulfite_repC.10 | R278-L1-P06-CATGGC_trimmed_1P.fq.gz | 1212601 | 914187 |
| *T. urartu* | A | 1 | SRR4099365 | Turartu_bisulfite_repA.1 | R259-L2-P13-GCGCTA_trimmed_1P.fq.gz | 13310388 | 4159576 |
| *T. urartu* | B | 1 | SRR4099366 | Turartu_bisulfite_repB.1 | R259-L2-P14-TAATCG_trimmed_1P.fq.gz | 12920707 | 4057100 |
| *T. urartu* | C | 1 | SRR4099367 | Turartu_bisulfite_repC.1 | R259-L2-P15-TACAGC_trimmed_1P.fq.gz | 16009977 | 4976951 |
| *T. urartu* | A | 2 | SRR4099368 | Turartu_bisulfite_repA.2 | R269-L1-P13-GCGCTA_trimmed_1P.fq.gz | 10409319 | 7002505 |
| *T. urartu* | B | 2 | SRR4099369 | Turartu_bisulfite_repB.2 | R269-L1-P14-TAATCG_trimmed_1P.fq.gz | 9410113 | 6378560 |
| *T. urartu* | C | 2 | SRR4099379 | Turartu_bisulfite_repC.2 | R269-L1-P15-TACAGC_trimmed_1P.fq.gz | 13566467 | 9155105 |
| *T. urartu* | A | 3 | SRR4099391 | Turartu_bisulfite_repA.3 | R269-L2-P13-GCGCTA_trimmed_1P.fq.gz | 10244048 | 6886444 |
| *T. urartu* | B | 3 | SRR4099392 | Turartu_bisulfite_repB.3 | R269-L2-P14-TAATCG_trimmed_1P.fq.gz | 9239089 | 6259859 |
| *T. urartu* | C | 3 | SRR4099393 | Turartu_bisulfite_repC.3 | R269-L2-P15-TACAGC_trimmed_1P.fq.gz | 13314670 | 8979088 |
| *T. urartu* | A | 4 | SRR4099394 | Turartu_bisulfite_repA.4 | R270-L1-P13-GCGCTA_trimmed_1P.fq.gz | 10453609 | 7013244 |
| *T. urartu* | B | 4 | SRR4099396 | Turartu_bisulfite_repB.4 | R270-L1-P14-TAATCG_trimmed_1P.fq.gz | 9270920 | 6274338 |
| *T. urartu* | C | 4 | SRR4099397 | Turartu_bisulfite_repC.4 | R270-L1-P15-TACAGC_trimmed_1P.fq.gz | 13965748 | 9397187 |
| *T. urartu* | A | 5 | SRR4099398 | Turartu_bisulfite_repA.5 | R270-L2-P13-GCGCTA_trimmed_1P.fq.gz | 10493350 | 7026554 |
| *T. urartu* | B | 5 | SRR4099399 | Turartu_bisulfite_repB.5 | R270-L2-P14-TAATCG_trimmed_1P.fq.gz | 9295082 | 6274857 |
| *T. urartu* | C | 5 | SRR4099400 | Turartu_bisulfite_repC.5 | R270-L2-P15-TACAGC_trimmed_1P.fq.gz | 13979903 | 9388254 |
| *T. urartu* | A | 6 | SRR4099401 | Turartu_bisulfite_repA.6 | R271-L1-P13-GCGCTA_trimmed_1P.fq.gz | 9888929 | 6637852 |
| *T. urartu* | B | 6 | SRR4099402 | Turartu_bisulfite_repB.6 | R271-L1-P14-TAATCG_trimmed_1P.fq.gz | 8030669 | 5434694 |
| *T. urartu* | C | 6 | SRR4099403 | Turartu_bisulfite_repC.6 | R271-L1-P15-TACAGC_trimmed_1P.fq.gz | 12627033 | 8493153 |
| *T. urartu* | A | 7 | SRR4099404 | Turartu_bisulfite_repA.7 | R271-L2-P13-GCGCTA_trimmed_1P.fq.gz | 9964918 | 6684237 |
| *T. urartu* | B | 7 | SRR4099405 | Turartu_bisulfite_repB.7 | R271-L2-P14-TAATCG_trimmed_1P.fq.gz | 8111224 | 5485638 |
| *T. urartu* | C | 7 | SRR4099407 | Turartu_bisulfite_repC.7 | R271-L2-P15-TACAGC_trimmed_1P.fq.gz | 12770960 | 8588539 |
| *T. urartu* | A | 8 | SRR4099408 | Turartu_bisulfite_repA.8 | R272-L1-P13-GCGCTA_trimmed_1P.fq.gz | 10084654 | 6747602 |
| *T. urartu* | B | 8 | SRR4099409 | Turartu_bisulfite_repB.8 | R272-L1-P14-TAATCG_trimmed_1P.fq.gz | 8480738 | 5728189 |
| *T. urartu* | C | 8 | SRR4099410 | Turartu_bisulfite_repC.8 | R272-L1-P15-TACAGC_trimmed_1P.fq.gz | 13380291 | 8990380 |
| *T. urartu* | A | 9 | SRR4099411 | Turartu_bisulfite_repA.9 | R272-L2-P13-GCGCTA_trimmed_1P.fq.gz | 9803468 | 6548078 |
| *T. urartu* | B | 9 | SRR4099412 | Turartu_bisulfite_repB.9 | R272-L2-P14-TAATCG_trimmed_1P.fq.gz | 8199786 | 5528353 |
| *T. urartu* | C | 9 | SRR4099413 | Turartu_bisulfite_repC.9 | R272-L2-P15-TACAGC_trimmed_1P.fq.gz | 12969803 | 8697196 |
| *T. urartu* | A | 10 | SRR4099414 | Turartu_bisulfite_repA.10 | R278-L1-P13-GCGCTA_trimmed_1P.fq.gz | 11976303 | 7382851 |
| *T. urartu* | B | 10 | SRR4099415 | Turartu_bisulfite_repB.10 | R278-L1-P14-TAATCG_trimmed_1P.fq.gz | 11533811 | 7178592 |

| | | | | | | | |
|---|---|---|---|---|---|---|---|
| *T. urartu* | C | 10 | SRR4099416 | Turartu_bisulfite_repC.10 | R278-L1-P15-TACAGC_trimmed_1P.fq.gz | 14489217 | 8958357 |
| *B. distachyon* | A | 1 | SRR628921 | NA | Bdi-BSseq-P22-SRR628921_trimmed_1P.fq.gz | 59093673 | 50797778 |
| *B. distachyon* | B | 2 | SRR629088 | NA | Bdi-BSseq-P23-SRR629088_trimmed_1P.fq.gz | 53707397 | 44611305 |
| *B. distachyon* | C | 3 | SRR629207 | NA | Bdi-BSseq-P24-SRR629207_trimmed_1P.fq.gz | 55918529 | 48483304 |
| *O. sativa* | A | 1 | SRR1035998 | NA | Osa-BSseq-P18-SRR1035998_trimmed_1P.fq.gz | 53170824 | 42881350 |
| *O. sativa* | B | 2 | SRR1035999 | NA | Osa-BSseq-P19-SRR1035999_trimmed_1P.fq.gz | 75831729 | 61345637 |
| *O. sativa* | C | 3 | SRR1036000 | NA | Osa-BSseq-P20-SRR1036000_trimmed_1P.fq.gz | 53999088 | 43195962 |
| *Z. mays* | A | 1 | SRR850328 | NA | Zma-BSseq-P25-SRR850328_trimmed_1P.fq.gz | 140210381 | 89333801 |

**Table S7. RNA sequencing statistics.**

| Species | Replicate | Lane | SRA Run ID | SRA file ID | Filename | Number of trimmed PE reads | Number of aligned PE reads |
|---|---|---|---|---|---|---|---|
| *T. urartu* | A | NA | SRR4101189 | Turartu_RNAseq_repA | 10-T-urartuA_trimmed.fq.gz | 31141621 | 19817208 |
| *T. urartu* | B | NA | SRR4101190 | Turartu_RNAseq_repB | 11-T-urartuB_trimmed.fq.gz | 34818985 | 22105789 |
| *T. urartu* | C | NA | SRR4101193 | Turartu_RNAseq_repC | 12-T-urartuC_trimmed.fq.gz | 29032619 | 19121432 |
| *H. vulgare* | A | NA | SRR4101194 | Hvulgare_RNAseq_repA | 1-H-vulgareA_trimmed.fq.gz | 31909780 | 19529644 |
| *H. vulgare* | B | NA | SRR4101195 | Hvulgare_RNAseq_repB | 2-H-vulagareB_trimmed.fq.gz | 30598122 | 21442787 |
| *H. vulgare* | C | NA | SRR4101196 | Hvulgare_RNAseq_repC | 3-H-vulgareC_trimmed.fq.gz | 31158941 | 21040339 |
| *S. italica* | A | NA | SRR4101197 | Sitalica_RNAseq_repA | 4-FoxtailMilletA_trimmed.fq.gz | 33332438 | 18087577 |
| *S. italica* | B | NA | SRR4101198 | Sitalica_RNAseq_repB | 5-FoxtailMilletB_trimmed.fq.gz | 30088604 | 15907783 |
| *S. italica* | C | NA | SRR4101199 | Sitalica_RNAseq_repC | 6-FoxtailMilletC_trimmed.fq.gz | 32943951 | 17523123 |
| *S. bicolor* | A | NA | SRR4101200 | Sbicolor_RNAseq_repA | 7-S-bicolorA_trimmed.fq.gz | 32679198 | 10421991 |
| *S. bicolor* | B | NA | SRR4101191 | Sbicolor_RNAseq_repB | 8-S-bicolorB_trimmed.fq.gz | 35296889 | 11041613 |
| *S. bicolor* | C | NA | SRR4101192 | Sbicolor_RNAseq_repC | 9-S-bicolorC_trimmed.fq.gz | 38379088 | 12336047 |
| *O. sativa* | A | NA | SRR1005354 | NA | 13-Osativa_trimmed.fq.gz | 27886722 | 16233735 |
| *O. sativa* | B | NA | SRR1005355 | NA | 14-Osativa_trimmed.fq.gz | 30761608 | 18180057 |
| *O. sativa* | C | NA | SRR1005356 | NA | 15-Osativa_trimmed.fq.gz | 28332640 | 14913912 |
| *B. distachyon* | A | NA | SRR3170524 | NA | 16-Bdistachyon_trimmed.fq.gz | 24709474 | 8525378 |
| *B. distachyon* | B | NA | SRR3170525 | NA | 17-Bdistachyon_trimmed.fq.gz | 28812202 | 12453434 |
| *B. distachyon* | C | NA | SRR3170526 | NA | 18-Bdistachyon_trimmed.fq.gz | 22722774 | 8500020 |
| *Z. mays* | A | 1 | SRR520998 | NA | 19.1-Zmays_trimmed.fq.gz | 9915832 | 5878300 |
| *Z. mays* | A | 2 | SRR520999 | NA | 19.2-Zmays_trimmed.fq.gz | 9768383 | 5793666 |
| *Z. mays* | B | 1 | SRR521003 | NA | 20-Zmays_trimmed.fq.gz | 130384170 | 80929553 |
| *Z. mays* | C | 1 | SRR521006 | NA | 21-Zmays_trimmed.fq.gz | 112808856 | 72185826 |
| *H. vulgare* | D | 1 | SRR3290257 | NA | 22.1-Hvulgare_trimmed.fq.gz | 18650844 | 11124682 |
| *H. vulgare* | D | 2 | SRR3290258 | NA | 22.2-Hvulgare_trimmed.fq.gz | 18971790 | 11311129 |
| *H. vulgare* | E | 1 | SRR3290283 | NA | 23.1-Hvulgare_trimmed.fq.gz | 17009797 | 10428572 |
| *H. vulgare* | E | 2 | SRR3290284 | NA | 23.2-Hvulgare_trimmed.fq.gz | 16774537 | 10280760 |
| *H. vulgare* | F | 1 | SRR3290313 | NA | 24.1-Hvulgare_trimmed.fq.gz | 12340135 | 7250487 |
| *H. vulgare* | F | 2 | SRR3290314 | NA | 24.2-Hvulgare_trimmed.fq.gz | 12209087 | 7169406 |
| *H. vulgare* | F | 3 | SRR3290315 | NA | 24.3-Hvulgare_trimmed.fq.gz | 12063315 | 7073810 |
| *P. heterocycla* | A | NA | ERR105067 | NA | 25-Pheterocylca_trimmed.fq.gz | 20987458 | 3089804 |
| *P. heterocycla* | B | NA | ERR105068 | NA | 26-Pheterocylca_trimmed.fq.gz | 17997913 | 3088968 |

**Table S8. List of orthologous gene groups.**

| orthologID | *P. heterocylca* | *O. sativa* | *B. distachyon* | *H. vulgare* | *T. urartu* | *S. italica* | *S. bicolor* | *Z. mays* |
|---|---|---|---|---|---|---|---|---|
| orthologID_008488 | PH01000276G0310 | LOC_Os06g49880.1 | Bradi1g35030.1 | HORVU7Hr1G091180.1 | TRIUR3_18855 | Seita.4G277900.1 | Sobic.010G262200.1 | Zm00001d014765_T001 |
| orthologID_008958 | PH01001050G0230 | LOC_Os04g47020.2 | Bradi5g18060.6 | HORVU2Hr1G096850.3 | TRIUR3_18278 | Seita.7G195300.1 | Sobic.010G039500.1 | Zm00001d026148_T002 |
| orthologID_008513 | PH01000061G1050 | LOC_Os06g43810.1 | Bradi1g30710.2 | HORVU7Hr1G114700.1 | TRIUR3_13145 | Seita.4G199400.1 | Sobic.010G207250.1 | Zm00001d036737_T001 |
| orthologID_008656 | PH01000245G1130 | LOC_Os06g11640.1 | Bradi1g45607.2 | HORVU7Hr1G036690.2 | TRIUR3_14294 | Seita.4G090200.1 | Sobic.010G088300.1 | Zm00001d044866_T001 |
| orthologID_008387 | PH01002537G0050 | LOC_Os06g05400.1 | Bradi1g51010.1 | HORVU7Hr1G021280.2 | TRIUR3_03041 | Seita.4G034100.1 | Sobic.010G035500.1 | Zm00001d045372_T002 |
| orthologID_008367 | PH01000169G0740 | LOC_Os06g42850.1 | Bradi1g35460.2 | HORVU7Hr1G092300.5 | TRIUR3_27460 | Seita.4G207300.1 | Sobic.010G199900.1 | Zm00001d046652_T001 |
| orthologID_008673 | PH01000401G0680 | LOC_Os06g13570.1 | Bradi1g44577.1 | HORVU7Hr1G040220.3 | TRIUR3_19678 | Seita.4G107400.1 | Sobic.010G100516.1 | Zm00001d037268_T003 |
| orthologID_007724 | PH01000083G1350 | LOC_Os06g02120.1 | Bradi1g50250.1 | HORVU0Hr1G005920.1 | TRIUR3_10437 | Seita.4G009600.1 | Sobic.010G009466.1 | Zm00001d045550_T002 |
| orthologID_007559 | PH01000366G0740 | LOC_Os06g42810.1 | Bradi1g35470.1 | HORVU7Hr1G092320.1 | TRIUR3_19690 | Seita.4G207400.1 | Sobic.010G199800.1 | Zm00001d046654_T002 |
| orthologID_008442 | PH01001272G0440 | LOC_Os06g49770.1 | Bradi1g34930.1 | HORVU7Hr1G094980.1 | TRIUR3_12747 | Seita.4G276900.1 | Sobic.010G261200.2 | Zm00001d014758_T001 |
| orthologID_008337 | PH01000065G2100 | LOC_Os12g17080.1 | Bradi1g45290.1 | HORVU0Hr1G019780.1 | TRIUR3_25948 | Seita.4G096700.1 | Sobic.010G091500.1 | Zm00001d044843_T001 |
| orthologID_009592 | PH01000127G1010 | LOC_Os02g19990.1 | Bradi3g11320.1 | HORVU0Hr1G039530.2 | TRIUR3_12550 | Seita.1G136600.1 | Sobic.010G248800.1 | Zm00001d053272_T001 |
| orthologID_008316 | PH01003464G0160 | LOC_Os06g05550.1 | Bradi1g50970.1 | HORVU7Hr1G021580.1 | TRIUR3_03037 | Seita.4G036600.1 | Sobic.010G037600.1 | Zm00001d011661_T001 |
| orthologID_008378 | PH01002192G0260 | LOC_Os01g13080.1 | Bradi1g34200.1 | HORVU7Hr1G096990.2 | TRIUR3_24083 | Seita.4G241800.1 | Sobic.010G251600.1 | Zm00001d014679_T001 |
| orthologID_008605 | PH01001174G0210 | LOC_Os06g06820.1 | Bradi1g48440.1 | HORVU7Hr1G026370.1 | TRIUR3_28059 | Seita.4G061800.1 | Sobic.010G051300.1 | Zm00001d045216_T002 |
| orthologID_007967 | PH01000327G0710 | LOC_Os06g50120.1 | Bradi1g35200.2 | HORVU7Hr1G091500.1 | TRIUR3_22896 | Seita.4G280200.1 | Sobic.010G265900.1 | Zm00001d036406_T001 |
| orthologID_008189 | PH01000418G0400 | LOC_Os06g37570.1 | Bradi1g37430.1 | HORVU7Hr1G086640.4 | TRIUR3_09433 | Seita.4G233900.1 | Sobic.010G173900.2 | Zm00001d046358_T001 |
| orthologID_007840 | PH01000517G0660 | LOC_Os06g10520.1 | Bradi1g46310.2 | HORVU7Hr1G035030.2 | TRIUR3_19900 | Seita.4G085200.1 | Sobic.010G078100.1 | Zm00001d044985_T014 |
| orthologID_008596 | PH01001415G0170 | LOC_Os06g08170.1 | Bradi1g47600.1 | HORVU7Hr1G029420.3 | TRIUR3_14829 | Seita.4G052300.1 | Sobic.010G062800.1 | Zm00001d035820_T002 |
| orthologID_007748 | PH01000276G1230 | LOC_Os06g50360.1 | Bradi1g30180.1 | HORVU7Hr1G117540.4 | TRIUR3_00347 | Seita.4G282000.1 | Sobic.010G268100.1 | Zm00001d036396_T002 |
| orthologID_009321 | PH01000952G0490 | LOC_Os02g21630.1 | Bradi3g10920.2 | HORVU0Hr1G022240.6 | TRIUR3_06405 | Seita.2G205800.1 | Sobic.010G005600.3 | Zm00001d045574_T003 |
| orthologID_008689 | PH01002612G0250 | LOC_Os06g47640.1 | Bradi1g33520.1 | HORVU7Hr1G099560.1 | TRIUR3_13177 | Seita.4G251700.1 | Sobic.010G241300.1 | Zm00001d014559_T001 |
| orthologID_007681 | PH01004010G0130 | LOC_Os06g05190.1 | Bradi1g51220.1 | HORVU7Hr1G019990.4 | TRIUR3_16962 | Seita.4G032300.1 | Sobic.010G033600.1 | Zm00001d045381_T001 |
| orthologID_008259 | PH01000849G0330 | LOC_Os06g35960.1 | Bradi1g38140.2 | HORVU7Hr1G088920.3 | TRIUR3_12841 | Seita.4G180500.1 | Sobic.010G164400.1 | Zm00001d046299_T001 |
| orthologID_008163 | PH01000069G1630 | LOC_Os06g46950.1 | Bradi1g33050.1 | HORVU7Hr1G101790.1 | TRIUR3_10232 | Seita.4G255900.1 | Sobic.010G234000.1 | Zm00001d014491_T001 |
| orthologID_008107 | PH01000517G0200 | LOC_Os06g10610.1 | Bradi1g46237.2 | HORVU7Hr1G035270.4 | TRIUR3_25102 | Seita.4G084500.1 | Sobic.010G078800.1 | Zm00001d044972_T001 |
| orthologID_007934 | PH01000176G1140 | LOC_Os06g49150.1 | Bradi1g34550.4 | HORVU7Hr1G096150.11 | TRIUR3_27831 | Seita.4G271600.1 | Sobic.010G255500.1 | Zm00001d014717_T001 |
| orthologID_007527 | PH01001107G0490 | LOC_Os06g12690.1 | Bradi4g07420.1 | HORVU5Hr1G030970.2 | TRIUR3_30999 | Seita.4G101800.1 | Sobic.010G095700.1 | Zm00001d037246_T002 |
| orthologID_008496 | PH01002612G0080 | LOC_Os09g07020.1 | Bradi1g33340.1 | HORVU4Hr1G015340.2 | TRIUR3_18430 | Seita.4G253100.1 | Sobic.010G239900.1 | Zm00001d024865_T001 |
| orthologID_007647 | PH01000832G0670 | LOC_Os06g01230.1 | Bradi1g52480.2 | HORVU7Hr1G000910.1 | TRIUR3_04701 | Seita.4G000700.1 | Sobic.010G000100.1 | Zm00001d045617_T001 |
| orthologID_007765 | PH01000776G0090 | LOC_Os06g19480.1 | Bradi1g43200.1 | HORVU7Hr1G044860.3 | TRIUR3_25391 | Seita.4G129900.1 | Sobic.010G123700.1 | Zm00001d045812_T001 |
| orthologID_007591 | PH01000366G0250 | LOC_Os06g42560.4 | Bradi1g35600.1 | HORVU7Hr1G092800.3 | TRIUR3_11469 | Seita.4G209000.1 | Sobic.010G198000.1 | Zm00001d046676_T001 |
| orthologID_008376 | PH01000268G0530 | LOC_Os06g14470.1 | Bradi1g44147.1 | HORVU7Hr1G041520.2 | TRIUR3_22054 | Seita.4G113800.1 | Sobic.010G105600.1 | Zm00001d037352_T001 |
| orthologID_007561 | PH01002462G0030 | LOC_Os06g30750.1 | Bradi1g40150.1 | HORVU7Hr1G075700.2 | TRIUR3_31025 | Seita.4G162900.1 | Sobic.010G147600.1 | Zm00001d037453_T001 |
| orthologID_008025 | PH01000065G0330 | LOC_Os06g12630.1 | Bradi1g45010.1 | HORVU7Hr1G038770.2 | TRIUR3_30693 | Seita.4G101400.1 | Sobic.010G095200.1 | Zm00001d037240_T001 |
| orthologID_008089 | PH01000503G0190 | LOC_Os06g06280.1 | Bradi1g48880.2 | HORVU7Hr1G024690.2 | TRIUR3_20229 | Seita.4G067900.1 | Sobic.010G044800.2 | Zm00001d036240_T001 |
| orthologID_008409 | PH01004075G0160 | LOC_Os06g39140.1 | Bradi1g37100.1 | HORVU0Hr1G021640.3 | TRIUR3_27237 | Seita.4G228000.1 | Sobic.010G178200.1 | Zm00001d046472_T001 |
| orthologID_008327 | PH01000014G0510 | LOC_Os06g09620.1 | Bradi1g46616.2 | HORVU7Hr1G033760.1 | TRIUR3_18462 | Seita.4G040500.1 | Sobic.010G073300.2 | Zm00001d037107_T001 |
| orthologID_008716 | PH01000069G1120 | LOC_Os06g47220.1 | Bradi1g33230.1 | HORVU0Hr1G038990.4 | TRIUR3_23765 | Seita.4G258300.1 | Sobic.010G236900.1 | Zm00001d014518_T001 |
| orthologID_008143 | PH01000045G0150 | LOC_Os06g03682.1 | Bradi1g49590.1 | HORVU7Hr1G010490.25 | TRIUR3_03525 | Seita.4G016900.1 | Sobic.010G016900.1 | Zm00001d036008_T001 |
| orthologID_008221 | PH01001174G0220 | LOC_Os06g06810.1 | Bradi1g48447.1 | HORVU7Hr1G026040.3 | TRIUR3_06594 | Seita.4G061900.1 | Sobic.010G051200.1 | Zm00001d045217_T011 |
| orthologID_007867 | PH01000080G0350 | LOC_Os02g26890.1 | Bradi1g45780.3 | HORVU7Hr1G036170.8 | TRIUR3_03222 | Seita.4G076900.1 | Sobic.010G085900.1 | Zm00001d037190_T001 |
| orthologID_007789 | PH01000268G0820 | LOC_Os06g14240.1 | Bradi1g44230.1 | HORVU0Hr1G020420.1 | TRIUR3_16866 | Seita.4G111400.1 | Sobic.010G104300.1 | Zm00001d044728_T001 |
| orthologID_008561 | PH01000164G0170 | LOC_Os06g49450.1 | Bradi1g34717.3 | HORVU7Hr1G095760.1 | TRIUR3_02543 | Seita.4G274100.1 | Sobic.010G257500.1 | Zm00001d014728_T006 |
| orthologID_008193 | PH01000245G0100 | LOC_Os06g11380.1 | Bradi1g45833.1 | HORVU7Hr1G036090.2 | TRIUR3_08497 | Seita.4G077400.1 | Sobic.010G085100.1 | Zm00001d044903_T026 |
| orthologID_007807 | PH01004769G0030 | LOC_Os06g05110.1 | Bradi1g51140.1 | HORVU7Hr1G020590.16 | TRIUR3_00248 | Seita.4G031200.1 | Sobic.010G033000.1 | Zm00001d036135_T003 |
| orthologID_008237 | PH01000709G0240 | LOC_Os06g19730.1 | Bradi1g42980.1 | HORVU7Hr1G045410.14 | TRIUR3_16461 | Seita.4G132000.1 | Sobic.010G125700.1 | Zm00001d037401_T001 |
| orthologID_008372 | PH01000407G0920 | LOC_Os06g05090.1 | Bradi1g51130.1 | HORVU7Hr1G020620.2 | TRIUR3_00249 | Seita.4G031000.1 | Sobic.010G032700.1 | Zm00001d036131_T001 |

| | | | | | | | | |
|---|---|---|---|---|---|---|---|---|
| orthologID_007726 | PH01001814G0430 | LOC_Os06g43800.1 | Bradi1g30656.1 | HORVU7Hr1G114960.13 | TRIUR3_28784 | Seita.4G199800.1 | Sobic.010G207100.1 | Zm00001d036738_T001 |
| orthologID_007762 | PH01000080G1160 | LOC_Os06g11730.1 | Bradi1g45540.1 | HORVU7Hr1G036920.2 | TRIUR3_18546 | Seita.4G091000.1 | Sobic.010G088950.1 | Zm00001d037209_T001 |
| orthologID_008451 | PH01001933G0030 | LOC_Os06g44540.1 | Bradi1g31250.1 | HORVU0Hr1G022360.1 | TRIUR3_01327 | Seita.4G191700.1 | Sobic.010G214700.1 | Zm00001d036701_T001 |
| orthologID_009921 | PH01005300G0060 | LOC_Os10g42960.1 | Bradi3g21640.1 | HORVU1Hr1G051450.1 | TRIUR3_32881 | Seita.4G101500.1 | Sobic.010G095400.1 | Zm00001d037242_T001 |
| orthologID_008040 | PH01000673G0190 | LOC_Os06g07160.1 | Bradi1g48210.1 | HORVU7Hr1G027350.3 | TRIUR3_20796 | Seita.4G059100.1 | Sobic.010G054300.2 | Zm00001d045193_T002 |
| orthologID_008003 | PH01000069G1880 | LOC_Os06g46799.1 | Bradi1g32870.1 | HORVU7Hr1G102480.7 | TRIUR3_13001 | Seita.4G259700.1 | Sobic.010G232500.1 | Zm00001d014467_T002 |
| orthologID_008296 | PH01003084G0120 | LOC_Os06g01460.1 | Bradi1g52170.1 | HORVU7Hr1G000120.2 | TRIUR3_08849 | Seita.4G002800.1 | Sobic.010G002700.1 | Zm00001d035878_T003 |
| orthologID_007628 | PH01000439G0170 | LOC_Os06g04190.1 | Bradi1g50020.1 | HORVU7Hr1G012570.17 | TRIUR3_32406 | Seita.4G021400.1 | Sobic.010G021900.1 | Zm00001d036044_T001 |
| orthologID_007981 | PH01001291G0040 | LOC_Os06g10310.1 | Bradi1g46427.1 | HORVU7Hr1G034610.4 | TRIUR3_30418 | Seita.4G086400.1 | Sobic.010G077200.1 | Zm00001d000238_T001 |
| orthologID_008058 | PH01000069G1790 | LOC_Os06g46900.1 | Bradi1g32990.1 | HORVU7Hr1G101920.1 | TRIUR3_14263 | Seita.4G258900.1 | Sobic.010G233200.1 | Zm00001d014486_T001 |
| orthologID_008277 | PH01000069G1690 | LOC_Os06g46930.1 | Bradi1g33030.1 | HORVU7Hr1G101810.1 | TRIUR3_10234 | Seita.4G255700.1 | Sobic.010G233800.1 | Zm00001d014488_T001 |
| orthologID_008734 | PH01003036G0170 | LOC_Os06g21600.1 | Bradi1g42660.1 | HORVU7Hr1G046290.3 | TRIUR3_22829 | Seita.4G140000.1 | Sobic.010G132600.1 | Zm00001d045897_T003 |
| orthologID_008447 | PH01006257G0040 | LOC_Os06g44080.1 | Bradi1g30940.1 | HORVU7Hr1G113500.1 | TRIUR3_11368 | Seita.4G196800.1 | Sobic.010G210000.1 | Zm00001d036717_T001 |
| orthologID_007616 | PH01000948G0150 | LOC_Os06g43140.1 | Bradi1g30273.2 | HORVU7Hr1G116640.1 | TRIUR3_13503 | Seita.4G205400.1 | Sobic.010G201700.2 | Zm00001d046628_T001 |
| orthologID_008346 | PH01001262G0380 | LOC_Os06g13660.1 | Bradi1g44530.1 | HORVU7Hr1G040390.16 | TRIUR3_14369 | Seita.4G108000.1 | Sobic.010G100900.1 | Zm00001d044745_T006 |
| orthologID_007877 | PH01003132G0080 | LOC_Os06g35814.1 | Bradi1g38190.1 | HORVU7Hr1G089120.1 | TRIUR3_24954 | Seita.4G179100.1 | Sobic.010G163600.1 | Zm00001d046313_T001 |
| orthologID_008075 | PH01002537G0060 | LOC_Os06g05390.1 | Bradi1g51020.1 | HORVU7Hr1G096120.1 | TRIUR3_03042 | Seita.4G034000.1 | Sobic.010G035400.1 | Zm00001d045373_T001 |
| orthologID_008546 | PH01000517G0640 | LOC_Os06g10530.1 | Bradi1g46300.1 | HORVU7Hr1G035010.1 | TRIUR3_30099 | Seita.4G085100.1 | Sobic.010G078200.1 | Zm00001d044986_T001 |
| orthologID_007703 | PH01000075G0200 | LOC_Os06g39590.1 | Bradi1g36930.1 | HORVU7Hr1G084940.1 | TRIUR3_02488 | Seita.4G226200.1 | Sobic.010G180600.1 | Zm00001d036918_T001 |
| orthologID_008280 | PH01001523G0600 | LOC_Os06g13070.1 | Bradi1g44750.5 | HORVU7Hr1G039710.3 | TRIUR3_29041 | Seita.4G105300.1 | Sobic.010G098700.3 | Zm00001d037258_T002 |
| orthologID_006905 | PH01000460G0750 | LOC_Os09g06560.1 | Bradi2g18270.1 | HORVU5Hr1G060480.1 | TRIUR3_14637 | Seita.7G065800.1 | Sobic.010G147700.1 | Zm00001d019852_T004 |
| orthologID_007599 | PH01000440G0290 | LOC_Os06g05530.1 | Bradi1g50990.2 | HORVU7Hr1G021340.7 | TRIUR3_03039 | Seita.4G036400.1 | Sobic.010G037400.1 | Zm00001d045358_T001 |
| orthologID_008217 | PH01000045G0240 | LOC_Os06g03660.1 | Bradi1g49550.3 | HORVU7Hr1G010250.4 | TRIUR3_25990 | Seita.4G016000.1 | Sobic.010G016400.1 | Zm00001d036001_T006 |
| orthologID_008715 | PH01000440G0880 | LOC_Os06g05820.1 | Bradi1g54900.2 | HORVU7Hr1G022440.1 | TRIUR3_10157 | Seita.4G072400.1 | Sobic.010G041000.1 | Zm00001d036181_T001 |
| orthologID_007247 | PH01000385G0650 | LOC_Os04g28020.1 | Bradi4g44050.2 | HORVU2Hr1G067040.5 | TRIUR3_29897 | Seita.7G056400.1 | Sobic.010G222900.2 | Zm00001d046970_T002 |
| orthologID_008444 | PH01000119G1000 | LOC_Os06g08720.1 | Bradi1g47090.1 | HORVU7Hr1G031020.2 | TRIUR3_27270 | Seita.4G046300.1 | Sobic.010G067500.1 | Zm00001d045085_T001 |
| orthologID_007925 | PH01000178G0320 | LOC_Os06g41050.1 | Bradi1g36000.2 | HORVU3Hr1G023060.16 | TRIUR3_33395 | Seita.4G216500.1 | Sobic.010G191000.1 | Zm00001d046761_T002 |
| orthologID_008608 | PH01004270G0050 | LOC_Os06g48620.1 | Bradi1g34070.1 | HORVU7Hr1G097570.14 | TRIUR3_09542 | Seita.4G243000.1 | Sobic.010G250200.1 | Zm00001d036484_T001 |
| orthologID_007641 | PH01000503G0870 | LOC_Os06g06030.1 | Bradi1g49140.2 | HORVU5Hr1G110330.1 | TRIUR3_14979 | Seita.4G070600.1 | Sobic.010G042800.1 | Zm00001d045315_T003 |
| orthologID_007567 | PH01001672G0230 | LOC_Os09g15480.1 | Bradi1g29750.2 | HORVU7Hr1G121440.2 | TRIUR3_08626 | Seita.4G287200.1 | Sobic.010G275100.1 | Zm00001d014852_T001 |
| orthologID_007711 | PH01002645G0060 | LOC_Os06g04820.1 | Bradi1g51290.1 | HORVU7Hr1G018670.3 | TRIUR3_03177 | Seita.4G028900.1 | Sobic.010G029900.2 | Zm00001d036110_T002 |
| orthologID_008251 | PH01003751G0100 | LOC_Os06g41060.1 | Bradi1g35990.1 | HORVU7Hr1G094690.1 | TRIUR3_20483 | Seita.4G216400.1 | Sobic.010G191100.2 | Zm00001d046759_T001 |
| orthologID_008264 | PH01000083G1240 | LOC_Os06g02210.1 | Bradi1g51710.1 | HORVU7Hr1G011580.2 | TRIUR3_03688 | Seita.4G008400.1 | Sobic.010G008400.1 | Zm00001d045558_T003 |
| orthologID_008184 | PH01000268G0540 | LOC_Os06g14460.1 | Bradi1g44160.1 | HORVU7Hr1G041540.1 | TRIUR3_22055 | Seita.4G113500.1 | Sobic.010G105400.1 | Zm00001d045655_T013 |
| orthologID_008599 | PH01000216G0880 | LOC_Os06g06550.1 | Bradi1g48620.1 | HORVU7Hr1G025380.7 | TRIUR3_07810 | Seita.4G065600.1 | Sobic.010G047600.1 | Zm00001d036255_T001 |
| orthologID_008633 | PH01001381G0520 | LOC_Os06g36560.1 | Bradi1g37870.1 | HORVU7Hr1G088260.4 | TRIUR3_27642 | Seita.4G239800.1 | Sobic.010G168800.1 | Zm00001d046234_T001 |
| orthologID_008531 | PH01000178G1160 | LOC_Os06g41390.1 | Bradi1g35910.2 | HORVU7Hr1G094310.16 | TRIUR3_14465 | Seita.4G214700.1 | Sobic.010G193900.1 | Zm00001d036795_T002 |
| orthologID_008194 | PH01000401G0630 | LOC_Os06g13220.1 | Bradi1g44640.5 | HORVU7Hr1G040150.1 | TRIUR3_34441 | Seita.4G106700.1 | Sobic.010G099900.1 | Zm00001d037266_T003 |
| orthologID_008057 | PH01002313G0050 | LOC_Os06g27760.1 | Bradi1g41830.1 | HORVU7Hr1G082710.2 | TRIUR3_31396 | Seita.J011900.1 | Sobic.010G101000.1 | Zm00001d037273_T001 |
| orthologID_008639 | PH01002537G0110 | LOC_Os06g05350.1 | Bradi1g51040.1 | HORVU7Hr1G021180.2 | TRIUR3_12734 | Seita.4G033800.1 | Sobic.010G035200.1 | Zm00001d036148_T003 |
| orthologID_008477 | PH01002473G0050 | LOC_Os06g37450.1 | Bradi1g37480.1 | HORVU7Hr1G086810.1 | TRIUR3_30842 | Seita.4G234800.1 | Sobic.010G173400.1 | Zm00001d046354_T001 |
| orthologID_007573 | PH01000045G1340 | LOC_Os06g02555.1 | Bradi1g50590.1 | HORVU7Hr1G008690.19 | TRIUR3_01358 | Seita.4G012300.1 | Sobic.010G013200.1 | Zm00001d035964_T003 |
| orthologID_008437 | PH01000440G0850 | LOC_Os06g05800.1 | Bradi1g49340.2 | HORVU7Hr1G022410.1 | TRIUR3_25905 | Seita.4G072600.1 | Sobic.010G040800.1 | Zm00001d036196_T001 |
| orthologID_008502 | PH01001039G0450 | LOC_Os06g44060.1 | Bradi1g30930.1 | HORVU0Hr1G022470.8 | TRIUR3_11369 | Seita.4G196700.1 | Sobic.010G209900.1 | Zm00001d036718_T002 |
| orthologID_008199 | PH01008194G0030 | LOC_Os06g33930.1 | Bradi1g38720.1 | HORVU7Hr1G079510.5 | TRIUR3_14211 | Seita.4G169100.1 | Sobic.010G154900.1 | Zm00001d037028_T001 |
| orthologID_007821 | PH01000708G0760 | LOC_Os06g03940.1 | Bradi1g49940.2 | HORVU7Hr1G013190.10 | TRIUR3_21419 | Seita.4G019600.1 | Sobic.010G020300.1 | Zm00001d045483_T002 |
| orthologID_008703 | PH01001777G0110 | LOC_Os06g43990.1 | Bradi1g30850.4 | HORVU7Hr1G113890.3 | TRIUR3_06296 | Seita.4G197200.1 | Sobic.010G208900.1 | Zm00001d046803_T001 |
| orthologID_007665 | PH01004186G0090 | LOC_Os06g22670.1 | Bradi1g40407.4 | HORVU7Hr1G081540.1 | TRIUR3_11264 | Seita.4G161800.1 | Sobic.010G147000.1 | Zm00001d037461_T001 |
| orthologID_009030 | PH01004565G0080 | LOC_Os03g32090.1 | Bradi5g01406.1 | HORVU1Hr1G062240.2 | TRIUR3_21422 | Seita.4G122400.1 | Sobic.010G115400.2 | Zm00001d037296_T001 |
| orthologID_008647 | PH01000849G0460 | LOC_Os06g35970.1 | Bradi1g38120.1 | HORVU7Hr1G088890.2 | TRIUR3_20400 | Seita.4G180600.1 | Sobic.010G164500.1 | Zm00001d046297_T001 |

| | | | | | | | | |
|---|---|---|---|---|---|---|---|---|
| orthologID_008340 | PH01002727G0130 | LOC_Os06g44420.1 | Bradi1g31210.1 | HORVU1Hr1G003060.2 | TRIUR3_12088 | Seita.4G192500.1 | Sobic.010G213800.1 | Zm00001d046927_T001 |
| orthologID_007952 | PH01001297G0310 | LOC_Os06g41380.1 | Bradi1g35930.2 | HORVU7Hr1G094270.3 | TRIUR3_19485 | Seita.4G214900.1 | Sobic.010G193700.1 | Zm00001d036797_T002 |
| orthologID_008027 | PH01004659G0050 | LOC_Os06g43550.1 | Bradi1g30400.1 | HORVU0Hr1G022500.1 | TRIUR3_03383 | Seita.4G202900.1 | Sobic.010G204300.1 | Zm00001d046602_T002 |
| orthologID_008310 | PH01001240G0380 | LOC_Os06g07770.1 | Bradi1g47950.2 | HORVU7Hr1G028260.1 | TRIUR3_16412 | Seita.4G056300.1 | Sobic.010G058400.2 | Zm00001d037482_T001 |
| orthologID_008906 | PH01000057G0860 | LOC_Os04g56670.2 | Bradi5g24980.1 | HORVU2Hr1G110610.1 | TRIUR3_01832 | Seita.6G186900.1 | Sobic.010G100000.1 | Zm00001d037267_T005 |
| orthologID_008498 | PH01001272G0220 | LOC_Os06g49870.1 | Bradi1g35020.1 | HORVU0Hr1G021960.1 | TRIUR3_21013 | Seita.4G277800.1 | Sobic.010G262000.1 | Zm00001d014763_T001 |
| orthologID_008542 | PH01000484G0350 | LOC_Os06g12530.1 | Bradi1g45080.1 | HORVU7Hr1G038520.6 | TRIUR3_28228 | Seita.4G100200.1 | Sobic.010G094500.1 | Zm00001d037236_T001 |
| orthologID_008456 | PH01001523G0540 | LOC_Os06g13100.1 | Bradi1g44730.1 | HORVU7Hr1G039780.3 | TRIUR3_25605 | Seita.4G105500.1 | Sobic.010G099100.1 | Zm00001d037261_T001 |
| orthologID_008350 | PH01000014G0990 | LOC_Os06g09880.1 | Bradi1g46560.1 | HORVU7Hr1G033940.1 | TRIUR3_04301 | Seita.4G038900.1 | Sobic.010G074100.1 | Zm00001d045014_T003 |
| orthologID_007836 | PH01001814G0050 | LOC_Os06g43690.1 | Bradi1g30580.1 | HORVU7Hr1G115460.4 | TRIUR3_00213 | Seita.4G201000.1 | Sobic.010G205900.1 | Zm00001d046587_T003 |
| orthologID_007792 | PH01000818G0470 | LOC_Os06g05720.1 | Bradi1g50790.2 | HORVU7Hr1G021910.1 | TRIUR3_09799 | Seita.4G074400.1 | Sobic.010G039800.1 | Zm00001d036201_T001 |
| orthologID_007270 | PH01002126G0210 | LOC_Os12g13950.1 | Bradi4g39480.1 | HORVU7Hr1G102300.8 | TRIUR3_30551 | Seita.3G086800.1 | Sobic.010G103700.2 | Zm00001d023801_T004 |
| orthologID_007793 | PH01001019G0410 | LOC_Os06g48510.1 | Bradi1g33960.1 | HORVU7Hr1G097970.8 | TRIUR3_23657 | Seita.4G244500.1 | Sobic.010G248301.1 | Zm00001d036496_T001 |
| orthologID_008304 | PH01001120G0290 | LOC_Os06g19650.1 | Bradi1g43110.1 | HORVU7Hr1G045230.2 | TRIUR3_34811 | Seita.4G131400.1 | Sobic.010G125200.1 | Zm00001d045825_T001 |
| orthologID_008181 | PH01001867G0070 | LOC_Os06g23910.1 | Bradi1g41750.1 | HORVU7Hr1G082470.4 | TRIUR3_12837 | Seita.4G156300.1 | Sobic.010G096200.1 | Zm00001d044801_T001 |
| orthologID_008670 | PH01000065G0180 | LOC_Os06g12740.1 | Bradi1g44980.1 | HORVU7Hr1G038900.15 | TRIUR3_27811 | Seita.4G101900.1 | Sobic.010G095800.1 | Zm00001d044806_T001 |
| orthologID_009138 | PH01000246G0410 | LOC_Os06g02230.1 | Bradi3g18810.1 | HORVU7Hr1G004350.2 | TRIUR3_32580 | Seita.4G008200.1 | Sobic.010G008300.1 | Zm00001d035921_T001 |
| orthologID_008516 | PH01000169G1000 | LOC_Os06g43080.1 | Bradi1g35290.1 | HORVU7Hr1G091880.1 | TRIUR3_01431 | Seita.4G206000.1 | Sobic.010G201200.2 | Zm00001d046634_T001 |
| orthologID_008696 | PH01001187G0230 | LOC_Os09g10910.3 | Bradi1g41940.1 | HORVU7Hr1G083180.1 | TRIUR3_23096 | Seita.4G154400.1 | Sobic.010G144900.1 | Zm00001d037463_T001 |
| orthologID_008727 | PH01000407G0260 | LOC_Os06g04790.1 | Bradi1g51330.1 | HORVU7Hr1G018600.1 | TRIUR3_29121 | Seita.9G067000.1 | Sobic.010G029300.1 | Zm00001d036107_T001 |
| orthologID_008532 | PH01001309G0250 | LOC_Os06g46240.1 | Bradi1g32690.2 | HORVU7Hr1G106780.2 | TRIUR3_12485 | Seita.4G263600.1 | Sobic.010G227100.2 | Zm00001d036611_T001 |
| orthologID_007969 | PH01000494G0840 | LOC_Os06g34850.1 | Bradi1g39130.1 | HORVU7Hr1G080380.1 | TRIUR3_34436 | Seita.4G174700.1 | Sobic.010G160200.1 | Zm00001d046161_T001 |
| orthologID_008202 | PH01000352G0150 | LOC_Os06g04040.1 | Bradi1g50010.1 | HORVU3Hr1G081110.1 | TRIUR3_24753 | Seita.4G020900.1 | Sobic.010G021500.1 | Zm00001d045472_T002 |
| orthologID_008223 | PH01001174G0580 | LOC_Os06g06720.1 | Bradi1g48540.1 | HORVU7Hr1G025650.1 | TRIUR3_05125 | Seita.4G062900.1 | Sobic.010G050200.1 | Zm00001d045233_T004 |
| orthologID_008723 | PH01002999G0140 | LOC_Os06g22820.1 | Bradi1g39510.1 | HORVU7Hr1G080980.2 | TRIUR3_21705 | Seita.4G161200.1 | Sobic.010G146600.1 | Zm00001d037058_T001 |
| orthologID_008704 | PH01000450G0890 | LOC_Os06g45090.2 | Bradi1g31660.1 | HORVU7Hr1G109780.5 | TRIUR3_00935 | Seita.4G187900.1 | Sobic.010G218100.1 | Zm00001d046934_T003 |
| orthologID_008021 | PH01000045G2140 | LOC_Os06g03790.1 | Bradi1g49670.1 | HORVU7Hr1G011290.1 | TRIUR3_17076 | Seita.4G017900.1 | Sobic.010G018000.1 | Zm00001d045500_T001 |
| orthologID_008526 | PH01000327G0660 | LOC_Os06g50130.1 | Bradi1g35210.1 | HORVU7Hr1G091580.1 | TRIUR3_22050 | Seita.4G280300.1 | Sobic.010G266000.1 | Zm00001d014781_T001 |
| orthologID_007618 | PH01001291G0490 | LOC_Os06g10210.1 | Bradi1g46470.1 | HORVU7Hr1G034350.1 | TRIUR3_31008 | Seita.4G087400.1 | Sobic.010G076700.1 | Zm00001d000357_T002 |
| orthologID_007809 | PH01000407G0170 | LOC_Os06g04650.1 | Bradi1g51410.1 | HORVU7Hr1G018320.4 | TRIUR3_07484 | Seita.4G027500.1 | Sobic.010G028400.1 | Zm00001d045418_T001 |
| orthologID_008660 | PH01001530G0240 | LOC_Os06g37640.1 | Bradi1g37370.1 | HORVU7Hr1G086420.4 | TRIUR3_09792 | Seita.4G233300.1 | Sobic.010G174700.1 | Zm00001d046366_T001 |
| orthologID_008051 | PH01002336G0310 | LOC_Os06g07560.1 | Bradi1g48070.1 | HORVU7Hr1G027770.4 | TRIUR3_14314 | Seita.4G057300.1 | Sobic.010G057000.1 | Zm00001d037477_T001 |
| orthologID_008023 | PH01000352G0910 | LOC_Os06g04300.1 | Bradi1g51640.1 | HORVU7Hr1G014150.4 | TRIUR3_10579 | Seita.4G023500.1 | Sobic.010G024100.1 | Zm00001d016877_T002 |
| orthologID_008598 | PH01002597G0150 | LOC_Os06g03980.1 | Bradi1g49960.2 | HORVU7Hr1G012910.14 | TRIUR3_06615 | Seita.4G020000.1 | Sobic.010G020700.1 | Zm00001d045480_T003 |
| orthologID_008176 | PH01001342G0160 | LOC_Os06g14740.1 | Bradi1g44060.1 | HORVU7Hr1G041850.1 | TRIUR3_07773 | Seita.4G115500.1 | Sobic.010G106900.1 | Zm00001d037333_T002 |
| orthologID_008621 | PH01001631G0150 | LOC_Os06g06160.2 | Bradi1g49070.1 | HORVU7Hr1G023980.3 | TRIUR3_12918 | Seita.4G069300.1 | Sobic.010G043600.1 | Zm00001d045305_T001 |
| orthologID_008066 | PH01001174G0200 | LOC_Os06g06870.1 | Bradi1g48426.1 | HORVU7Hr1G026430.16 | TRIUR3_31488 | Seita.4G061700.1 | Sobic.010G051400.1 | Zm00001d045215_T001 |
| orthologID_006338 | PH01002127G0220 | LOC_Os01g10490.1 | Bradi2g06300.1 | HORVU3Hr1G026690.5 | TRIUR3_32452 | Seita.5G143000.1 | Sobic.010G070900.2 | Zm00001d037094_T001 |
| orthologID_007583 | PH01000075G0480 | LOC_Os06g39660.1 | Bradi1g36890.1 | HORVU7Hr1G084950.3 | TRIUR3_13861 | Seita.4G225700.1 | Sobic.010G181100.1 | Zm00001d046440_T001 |
| orthologID_008615 | PH01000724G0740 | LOC_Os06g50880.1 | Bradi1g29980.1 | HORVU7Hr1G118690.2 | TRIUR3_05241 | Seita.4G283400.1 | Sobic.010G269500.1 | Zm00001d036386_T001 |
| orthologID_008377 | PH01001494G0310 | LOC_Os06g18820.1 | Bradi1g43365.1 | HORVU7Hr1G043800.7 | TRIUR3_05195 | Seita.4G127400.1 | Sobic.010G122000.1 | Zm00001d037385_T002 |
| orthologID_008128 | PH01001182G0490 | LOC_Os06g39240.1 | Bradi1g37080.1 | HORVU7Hr1G085130.1 | TRIUR3_35007 | Seita.4G227700.1 | Sobic.010G178500.1 | Zm00001d046471_T001 |
| orthologID_008674 | PH01000178G0310 | LOC_Os06g41040.1 | Bradi1g36010.1 | HORVU7Hr1G094640.60 | TRIUR3_12435 | Seita.4G216600.1 | Sobic.010G190900.1 | Zm00001d046764_T001 |
| orthologID_006203 | PH01001082G0140 | LOC_Os04g17064.1 | Bradi2g13510.1 | HORVU5Hr1G113970.13 | TRIUR3_26341 | Seita.2G167800.1 | Sobic.010G081200.1 | Zm00001d044945_T001 |
| orthologID_008185 | PH01001077G0400 | LOC_Os06g09930.1 | Bradi1g46520.1 | HORVU7Hr1G034160.8 | TRIUR3_17411 | Seita.J001400.1 | Sobic.010G074900.1 | Zm00001d037110_T002 |
| orthologID_007824 | PH01000276G1300 | LOC_Os06g50370.1 | Bradi1g30190.1 | HORVU7Hr1G117640.2 | TRIUR3_02727 | Seita.4G282100.1 | Sobic.010G268200.1 | Zm00001d014796_T001 |
| orthologID_007997 | PH01000286G0480 | LOC_Os06g08670.1 | Bradi1g47140.1 | HORVU7Hr1G030810.1 | TRIUR3_07828 | Seita.4G046700.1 | Sobic.010G067100.1 | Zm00001d037079_T001 |
| orthologID_008726 | PH01000061G0920 | LOC_Os06g43780.1 | Bradi1g30640.1 | HORVU7Hr1G115200.4 | TRIUR3_23360 | Seita.4G200200.1 | Sobic.010G206800.1 | Zm00001d036739_T001 |
| orthologID_007922 | PH01001672G0220 | LOC_Os11g38930.1 | Bradi1g29740.2 | HORVU7Hr1G121430.7 | TRIUR3_08627 | Seita.4G287300.1 | Sobic.010G275150.1 | Zm00001d014853_T002 |
| orthologID_008172 | PH01000740G0150 | LOC_Os06g06900.1 | Bradi1g48400.1 | HORVU7Hr1G026560.2 | TRIUR3_21181 | Seita.4G061500.1 | Sobic.010G051800.1 | Zm00001d045212_T001 |

| | | | | | | | | |
|---|---|---|---|---|---|---|---|---|
| orthologID_008239 | PH01004129G0090 | LOC_Os06g36480.1 | Bradi1g37898.1 | HORVU7Hr1G088320.5 | TRIUR3_32198 | Seita.4G240200.1 | Sobic.010G168500.2 | Zm00001d046246_T001 |
| orthologID_007469 | PH01000574G0030 | LOC_Os12g18640.1 | Bradi4g39100.1 | HORVU3Hr1G088470.1 | TRIUR3_28517 | Seita.3G308800.1 | Sobic.010G249700.1 | Zm00001d036489_T001 |
| orthologID_008452 | PH01000439G0840 | LOC_Os06g04380.1 | Bradi1g51620.1 | HORVU7Hr1G015760.1 | TRIUR3_20250 | Seita.4G024000.1 | Sobic.010G024600.1 | Zm00001d045436_T001 |
| orthologID_008733 | PH01000069G0060 | LOC_Os06g47820.1 | Bradi1g33440.1 | HORVU7Hr1G099220.5 | TRIUR3_02921 | Seita.4G250000.1 | Sobic.010G243000.1 | Zm00001d014583_T001 |
| orthologID_007911 | PH01003143G0160 | LOC_Os06g38970.1 | Bradi1g37210.1 | HORVU7Hr1G085650.4 | TRIUR3_17011 | Seita.4G229500.1 | Sobic.010G177100.1 | Zm00001d046488_T001 |
| orthologID_007649 | PH01002141G0160 | LOC_Os06g45370.1 | Bradi1g31810.1 | HORVU7Hr1G109280.1 | TRIUR3_08419 | Seita.4G186100.1 | Sobic.010G219700.2 | Zm00001d046942_T001 |
| orthologID_006851 | PH01001733G0350 | LOC_Os01g32090.1 | Bradi2g13480.1 | HORVU5Hr1G079050.1 | TRIUR3_09168 | Seita.2G131400.1 | Sobic.010G081700.1 | Zm00001d044944_T001 |
| orthologID_008145 | PH01001158G0600 | LOC_Os06g09330.1 | Bradi1g46730.1 | HORVU7Hr1G032310.8 | TRIUR3_27038 | Seita.4G042200.1 | Sobic.010G071200.1 | Zm00001d045049_T001 |
| orthologID_008676 | PH01000286G0050 | LOC_Os06g09170.2 | Bradi1g46900.4 | HORVU7Hr1G031690.4 | TRIUR3_05470 | Seita.4G043700.1 | Sobic.010G069500.3 | Zm00001d045065_T001 |
| orthologID_007610 | PH01004008G0200 | LOC_Os06g12100.1 | Bradi1g45320.1 | HORVU7Hr1G037730.1 | TRIUR3_07705 | Seita.4G095900.1 | Sobic.010G091200.3 | Zm00001d006629_T001 |
| orthologID_008438 | PH01000246G0760 | LOC_Os06g02000.3 | Bradi1g51830.5 | HORVU3Hr1G026180.16 | TRIUR3_15688 | Seita.4G007200.1 | Sobic.010G007500.1 | Zm00001d035916_T001 |
| orthologID_008106 | PH01000724G0630 | LOC_Os06g50930.1 | Bradi1g30020.1 | HORVU0Hr1G022690.1 | TRIUR3_18768 | Seita.4G284100.1 | Sobic.010G270300.1 | Zm00001d014816_T001 |
| orthologID_008658 | PH01000269G0330 | LOC_Os07g27460.2 | Bradi1g29077.4 | HORVU1Hr1G002340.1 | TRIUR3_08602 | Seita.4G177700.1 | Sobic.010G162200.4 | Zm00001d046323_T001 |
| orthologID_007673 | PH01002125G0220 | LOC_Os06g51420.1 | Bradi1g29390.1 | HORVU7Hr1G119810.1 | TRIUR3_06239 | Seita.4G290500.1 | Sobic.010G278400.1 | Zm00001d014894_T001 |
| orthologID_010114 | PH01000176G0550 | LOC_Os06g48960.1 | Bradi1g34350.1 | HORVU7Hr1G096470.2 | TRIUR3_25751 | Seita.4G240500.1 | Sobic.010G253400.1 | Zm00001d014692_T001 |
| orthologID_007856 | PH01002059G0180 | LOC_Os06g44910.1 | Bradi1g31450.1 | HORVU7Hr1G110810.1 | TRIUR3_18444 | Seita.4G190300.1 | Sobic.010G216200.1 | Zm00001d046900_T003 |
| orthologID_008732 | PH01001156G0060 | LOC_Os06g36940.1 | Bradi1g37710.2 | HORVU7Hr1G087700.2 | TRIUR3_26020 | Seita.4G237800.1 | Sobic.010G170900.1 | Zm00001d036970_T001 |
| orthologID_008002 | PH01001087G0530 | LOC_Os06g21980.1 | Bradi1g42620.1 | HORVU7Hr1G046410.5 | TRIUR3_18240 | Seita.4G140800.1 | Sobic.010G133400.1 | Zm00001d045911_T002 |
| orthologID_008517 | PH01001174G0310 | LOC_Os06g06790.1 | Bradi1g48460.1 | HORVU7Hr1G026030.2 | TRIUR3_06595 | Seita.4G062000.1 | Sobic.010G051100.1 | Zm00001d045218_T003 |
| orthologID_008534 | PH01002297G0180 | LOC_Os06g39650.1 | Bradi1g36900.2 | HORVU7Hr1G084980.2 | TRIUR3_13860 | Seita.4G225800.1 | Sobic.010G181000.1 | Zm00001d036912_T001 |
| orthologID_007566 | PH01003426G0010 | LOC_Os06g45100.1 | Bradi1g31670.1 | HORVU7Hr1G109800.14 | TRIUR3_00934 | Seita.4G187800.1 | Sobic.010G218200.1 | Zm00001d046935_T001 |
| orthologID_009140 | PH01000026G2110 | LOC_Os02g56210.1 | Bradi3g54700.1 | HORVU6Hr1G088300.1 | TRIUR3_30918 | Seita.1G358200.1 | Sobic.010G239800.1 | Zm00001d036540_T002 |
| orthologID_007941 | PH01003168G0040 | LOC_Os06g17830.1 | Bradi1g43447.1 | HORVU0Hr1G039260.9 | TRIUR3_13944 | Seita.4G126100.1 | Sobic.010G121600.1 | Zm00001d045782_T001 |
| orthologID_007905 | PH01000722G0200 | LOC_Os06g10770.1 | Bradi1g46130.1 | HORVU7Hr1G035450.1 | TRIUR3_14805 | Seita.4G083100.1 | Sobic.010G080300.1 | Zm00001d037164_T001 |
| orthologID_008361 | PH01003413G0110 | LOC_Os06g45410.1 | Bradi1g31837.1 | HORVU7Hr1G109130.1 | TRIUR3_14321 | Seita.4G185800.1 | Sobic.010G220000.1 | Zm00001d046947_T001 |
| orthologID_008428 | PH01007388G0050 | LOC_Os06g45000.1 | Bradi1g31580.3 | HORVU7Hr1G110260.1 | TRIUR3_08353 | Seita.4G189400.1 | Sobic.010G217100.1 | Zm00001d046890_T001 |
| orthologID_008326 | PH01002017G0170 | LOC_Os06g04610.1 | Bradi1g51470.3 | HORVU7Hr1G017880.1 | TRIUR3_19379 | Seita.4G026900.1 | Sobic.010G028000.1 | Zm00001d045421_T001 |
| orthologID_008248 | PH01000475G0290 | LOC_Os06g20620.1 | Bradi1g42770.1 | HORVU7Hr1G045910.1 | TRIUR3_22228 | Seita.4G137000.1 | Sobic.010G130300.1 | Zm00001d045873_T002 |
| orthologID_007796 | PH01000834G0790 | LOC_Os06g42000.1 | Bradi1g35660.1 | HORVU7Hr1G093020.1 | TRIUR3_26997 | Seita.4G210300.1 | Sobic.010G197200.1 | Zm00001d046682_T001 |
| orthologID_008197 | PH01002626G0270 | LOC_Os06g46880.1 | Bradi1g32970.1 | HORVU7Hr1G102060.1 | TRIUR3_04265 | Seita.4G259400.1 | Sobic.010G232900.1 | Zm00001d036563_T001 |
| orthologID_008035 | PH01000246G0970 | LOC_Os06g01966.1 | Bradi1g51910.1 | HORVU7Hr1G002600.1 | TRIUR3_00683 | Seita.4G006500.1 | Sobic.010G006700.1 | Zm00001d045567_T001 |
| orthologID_010065 | PH01000494G0920 | LOC_Os06g35050.1 | Bradi1g39160.1 | HORVU7Hr1G080460.6 | TRIUR3_15197 | Seita.4G174900.1 | Sobic.010G160400.1 | Zm00001d046168_T001 |
| orthologID_008161 | PH01001102G0300 | LOC_Os06g09600.1 | Bradi1g46630.1 | HORVU7Hr1G033490.1 | TRIUR3_07874 | Seita.4G041000.1 | Sobic.010G073000.2 | Zm00001d045030_T001 |
| orthologID_008101 | PH01000673G0050 | LOC_Os06g07474.1 | Bradi1g48120.1 | HORVU7Hr1G027600.1 | TRIUR3_07849 | Seita.4G057900.1 | Sobic.010G055700.1 | Zm00001d045181_T001 |
| orthologID_007927 | PH01000014G2120 | LOC_Os06g10440.1 | Bradi1g46350.1 | HORVU7Hr1G034860.2 | TRIUR3_09952 | Seita.4G085800.1 | Sobic.010G077600.2 | Zm00001d037124_T001 |
| orthologID_007816 | PH01004275G0110 | LOC_Os06g39040.1 | Bradi1g37175.1 | HORVU7Hr1G085570.3 | TRIUR3_25935 | Seita.4G229200.1 | Sobic.010G177400.1 | Zm00001d046483_T001 |
| orthologID_008691 | PH01001588G0470 | LOC_Os06g41770.1 | Bradi1g35790.1 | HORVU7Hr1G093580.1 | TRIUR3_33804 | Seita.4G212900.1 | Sobic.010G194900.2 | Zm00001d046718_T001 |
| orthologID_008590 | PH01001272G0140 | LOC_Os06g49890.1 | Bradi1g35040.1 | HORVU7Hr1G091170.2 | TRIUR3_18854 | Seita.4G278100.1 | Sobic.010G262300.1 | Zm00001d036417_T004 |
| orthologID_008736 | PH01001478G0290 | LOC_Os06g38320.1 | Bradi1g37311.4 | HORVU7Hr1G086010.14 | TRIUR3_03680 | Seita.4G231400.1 | Sobic.010G176200.1 | Zm00001d046385_T001 |
| orthologID_008504 | PH01002032G0070 | LOC_Os06g34020.1 | Bradi1g38755.1 | HORVU7Hr1G079650.1 | TRIUR3_20131 | Seita.4G169700.1 | Sobic.010G155500.1 | Zm00001d046134_T001 |
| orthologID_008241 | PH01000887G0620 | LOC_Os06g10710.1 | Bradi1g46160.1 | HORVU7Hr1G035420.1 | TRIUR3_22237 | Seita.4G083500.1 | Sobic.010G079800.1 | Zm00001d037158_T001 |
| orthologID_008273 | PH01000820G0060 | LOC_Os06g16060.1 | Bradi1g43750.1 | HORVU7Hr1G042940.1 | TRIUR3_24140 | Seita.4G120300.1 | Sobic.010G113800.2 | Zm00001d037300_T001 |
| orthologID_008022 | PH01000080G0090 | LOC_Os06g11310.1 | Bradi1g45790.1 | HORVU7Hr1G036180.1 | TRIUR3_03221 | Seita.4G077000.1 | Sobic.010G085800.1 | Zm00001d044898_T001 |
| orthologID_008065 | PH01001751G0210 | LOC_Os06g44900.1 | Bradi1g31440.1 | HORVU7Hr1G110790.1 | TRIUR3_13027 | Seita.4G190400.1 | Sobic.010G216100.1 | Zm00001d036688_T001 |
| orthologID_008016 | PH01000286G0540 | LOC_Os06g08650.1 | Bradi1g47170.1 | HORVU7Hr1G030850.2 | TRIUR3_07830 | Seita.4G046900.1 | Sobic.010G066900.1 | Zm00001d045090_T001 |
| orthologID_007702 | PH01001356G0230 | LOC_Os06g34120.1 | Bradi1g38780.3 | HORVU7Hr1G079810.1 | TRIUR3_16783 | Seita.4G170500.1 | Sobic.010G156100.1 | Zm00001d046136_T001 |
| orthologID_008464 | PH01002645G0160 | LOC_Os06g04910.1 | Bradi1g51060.1 | HORVU7Hr1G021100.2 | TRIUR3_06620 | Seita.4G029400.1 | Sobic.010G030700.1 | Zm00001d045394_T001 |
| orthologID_008297 | PH01000776G0370 | LOC_Os06g19390.1 | Bradi1g43240.1 | HORVU7Hr1G044240.1 | TRIUR3_29531 | Seita.4G129400.1 | Sobic.010G123300.1 | Zm00001d045798_T001 |
| orthologID_008413 | PH01000618G0620 | LOC_Os06g37010.1 | Bradi1g37667.1 | HORVU7Hr1G087490.2 | TRIUR3_33178 | Seita.4G237500.1 | Sobic.010G171300.1 | Zm00001d036965_T001 |
| orthologID_008566 | PH01000418G0350 | LOC_Os06g37610.1 | Bradi1g37390.1 | HORVU7Hr1G086570.2 | TRIUR3_09795 | Seita.4G233600.1 | Sobic.010G174400.1 | Zm00001d046363_T001 |

| | | | | | | | | |
|---|---|---|---|---|---|---|---|---|
| orthologID_008416 | PH01003606G0270 | LOC_Os06g47900.1 | Bradi1g33657.1 | HORVU7Hr1G098610.3 | TRIUR3_21176 | Seita.4G248800.1 | Sobic.010G244000.1 | Zm00001d014595_T001 |
| orthologID_007757 | PH01000843G0230 | LOC_Os06g35160.1 | Bradi1g38340.1 | HORVU7Hr1G089510.1 | TRIUR3_27512 | Seita.4G175800.1 | Sobic.010G161000.1 | Zm00001d046175_T001 |
| orthologID_007845 | PH01001544G0320 | LOC_Os06g51480.1 | Bradi1g29240.1 | HORVU7Hr1G105880.1 | TRIUR3_10377 | Seita.4G292000.1 | Sobic.010G280100.1 | Zm00001d028780_T001 |
| orthologID_008595 | PH01000083G0100 | LOC_Os06g01470.1 | Bradi1g52180.1 | HORVU7Hr1G000130.4 | TRIUR3_08850 | Seita.4G002900.1 | Sobic.010G002800.2 | Zm00001d035881_T001 |
| orthologID_008434 | PH01005548G0040 | LOC_Os06g04530.1 | Bradi1g51530.1 | HORVU7Hr1G017670.6 | TRIUR3_23002 | Seita.4G026100.1 | Sobic.010G027200.1 | Zm00001d045430_T002 |
| orthologID_010057 | PH01001544G0040 | LOC_Os06g51520.1 | Bradi1g29321.1 | HORVU7Hr1G119620.4 | TRIUR3_00657 | Seita.4G290800.1 | Sobic.010G278700.1 | Zm00001d014896_T001 |
| orthologID_007569 | PH01000164G0280 | LOC_Os06g49300.1 | Bradi1g34647.1 | HORVU7Hr1G095920.1 | TRIUR3_19422 | Seita.4G272500.1 | Sobic.010G256400.1 | Zm00001d014721_T001 |
| orthologID_008329 | PH01000407G0050 | LOC_Os06g04580.1 | Bradi1g51500.1 | HORVU7Hr1G017740.3 | TRIUR3_16225 | Seita.4G026600.1 | Sobic.010G027600.1 | Zm00001d011122_T001 |
| orthologID_006136 | PH01000274G0630 | LOC_Os01g08850.1 | Bradi2g05357.1 | HORVU3Hr1G020560.7 | TRIUR3_10393 | Seita.5G131200.1 | Sobic.003G039500.1 | Zm00001d039568_T001 |
| orthologID_006903 | PH01000592G0620 | LOC_Os01g08320.1 | Bradi2g04910.1 | HORVU3Hr1G022540.2 | TRIUR3_08262 | Seita.5G126200.1 | Sobic.003G044900.1 | Zm00001d039624_T001 |
| orthologID_006643 | PH01003793G0200 | LOC_Os01g61810.2 | Bradi2g54200.2 | HORVU3Hr1G087390.1 | TRIUR3_13296 | Seita.5G371200.1 | Sobic.003G346500.1 | Zm00001d042968_T002 |
| orthologID_006256 | PH01000109G1380 | LOC_Os01g57450.1 | Bradi2g51890.1 | HORVU3Hr1G077000.2 | TRIUR3_19232 | Seita.5G342000.1 | Sobic.003G319000.1 | Zm00001d012610_T001 |
| orthologID_009963 | PH01000697G0140 | LOC_Os01g03100.1 | Bradi2g01580.1 | HORVU3Hr1G005920.1 | TRIUR3_08435 | Seita.5G082500.1 | Sobic.003G093500.1 | Zm00001d008503_T002 |
| orthologID_006897 | PH01000536G0060 | LOC_Os01g52830.1 | Bradi2g48790.1 | HORVU3Hr1G069140.1 | TRIUR3_26578 | Seita.5G304500.1 | Sobic.003G283700.2 | Zm00001d011755_T001 |
| orthologID_006276 | PH01001553G0280 | LOC_Os01g29230.1 | Bradi2g13251.1 | HORVU3Hr1G046600.1 | TRIUR3_26279 | Seita.5G176200.1 | Sobic.003G170700.5 | Zm00001d040864_T003 |
| orthologID_006547 | PH01002825G0080 | LOC_Os01g07260.1 | Bradi2g04117.1 | HORVU3Hr1G017570.2 | TRIUR3_14930 | Seita.5G117500.1 | Sobic.003G055600.1 | Zm00001d039718_T001 |
| orthologID_009961 | PH01000004G1720 | LOC_Os01g64262.1 | Bradi2g55740.1 | HORVU2Hr1G038320.1 | TRIUR3_22026 | Seita.5G390800.1 | Sobic.003G365100.1 | Zm00001d042762_T001 |
| orthologID_006340 | PH01000094G1290 | LOC_Os01g16430.1 | Bradi2g10146.1 | HORVU3Hr1G038190.1 | TRIUR3_22636 | Seita.5G046700.1 | Sobic.003G126800.1 | Zm00001d008268_T002 |
| orthologID_006157 | PH01001378G0310 | LOC_Os01g52160.1 | Bradi2g48360.4 | HORVU3Hr1G068200.1 | TRIUR3_20021 | Seita.5G299000.1 | Sobic.003G278700.1 | Zm00001d043615_T001 |
| orthologID_007001 | PH01000229G0180 | LOC_Os01g67960.1 | Bradi2g58140.1 | HORVU3Hr1G091850.2 | TRIUR3_10276 | Seita.5G420400.1 | Sobic.003G394500.1 | Zm00001d012118_T002 |
| orthologID_006514 | PH01000054G1710 | LOC_Os01g62060.1 | Bradi2g54400.1 | HORVU3Hr1G087010.1 | TRIUR3_18607 | Seita.5G374200.1 | Sobic.003G349100.1 | Zm00001d042940_T001 |
| orthologID_006414 | PH01002236G0060 | LOC_Os01g50440.1 | Bradi2g47520.2 | HORVU3Hr1G065650.1 | TRIUR3_24049 | Seita.5G284700.1 | Sobic.003G269100.1 | Zm00001d043737_T001 |
| orthologID_006647 | PH01000525G0810 | LOC_Os01g03490.1 | Bradi2g01800.1 | HORVU3Hr1G004420.2 | TRIUR3_22734 | Seita.5G085100.1 | Sobic.003G089300.1 | Zm00001d040040_T001 |
| orthologID_007388 | PH01005355G0010 | LOC_Os12g23200.1 | Bradi4g08020.1 | HORVU3Hr1G009210.1 | TRIUR3_07216 | Seita.3G304000.1 | Sobic.003G052500.1 | Zm00001d008706_T001 |
| orthologID_006979 | PH01004293G0020 | LOC_Os01g37130.1 | Bradi2g40950.3 | HORVU3Hr1G048950.9 | TRIUR3_22173 | Seita.5G195800.1 | Sobic.003G185900.1 | Zm00001d044525_T001 |
| orthologID_006649 | PH01000812G0040 | LOC_Os01g13100.1 | Bradi2g07801.1 | HORVU3Hr1G031590.25 | TRIUR3_26197 | Seita.5G007600.1 | Sobic.003G007300.3 | Zm00001d039294_T002 |
| orthologID_006274 | PH01000048G0690 | LOC_Os01g51250.1 | Bradi2g47880.2 | HORVU3Hr1G066850.1 | TRIUR3_25231 | Seita.5G290200.1 | Sobic.003G273900.1 | Zm00001d011697_T001 |
| orthologID_006392 | PH01002045G0100 | LOC_Os01g11040.1 | Bradi2g06610.1 | HORVU3Hr1G029210.3 | TRIUR3_31093 | Seita.5G146900.1 | Sobic.003G023400.1 | Zm00001d008906_T003 |
| orthologID_006850 | PH01000721G0350 | LOC_Os01g09750.2 | Bradi2g05816.2 | HORVU3Hr1G019340.3 | TRIUR3_24572 | Seita.5G136800.1 | Sobic.003G033100.1 | Zm00001d039477_T001 |
| orthologID_006584 | PH01000009G1940 | LOC_Os01g60490.1 | Bradi2g53486.1 | HORVU3Hr1G080860.1 | TRIUR3_27575 | Seita.5G361600.1 | Sobic.003G337500.1 | Zm00001d043066_T001 |
| orthologID_006405 | PH01000320G0960 | LOC_Os01g03820.1 | Bradi2g02010.3 | HORVU3Hr1G003820.2 | TRIUR3_16064 | Seita.5G087700.1 | Sobic.003G086900.1 | Zm00001d040008_T005 |
| orthologID_006269 | PH01000229G1200 | LOC_Os01g67590.1 | Bradi2g57910.1 | HORVU3Hr1G088910.2 | TRIUR3_35379 | Seita.5G417900.1 | Sobic.003G392100.1 | Zm00001d042475_T001 |
| orthologID_006511 | PH01000094G0540 | LOC_Os01g16530.1 | Bradi2g10278.1 | HORVU3Hr1G038610.1 | TRIUR3_19394 | Seita.5G044900.1 | Sobic.003G128400.1 | Zm00001d040322_T002 |
| orthologID_006888 | PH01000155G1180 | LOC_Os01g56000.1 | Bradi2g50890.3 | HORVU3Hr1G074330.1 | TRIUR3_22135 | Seita.5G329800.1 | Sobic.003G307000.2 | Zm00001d043347_T002 |
| orthologID_006437 | PH01000031G1740 | LOC_Os01g68020.1 | Bradi2g58152.1 | HORVU3Hr1G092020.7 | TRIUR3_25289 | Seita.5G421200.1 | Sobic.003G395400.1 | Zm00001d042443_T001 |
| orthologID_006689 | PH01003108G0210 | LOC_Os01g14640.1 | Bradi2g08737.1 | HORVU3Hr1G033950.1 | TRIUR3_15672 | Seita.5G062800.1 | Sobic.003G111600.1 | Zm00001d040246_T001 |
| orthologID_006843 | PH01001166G0230 | LOC_Os01g37460.1 | Bradi2g40970.1 | HORVU3Hr1G049390.3 | TRIUR3_16750 | Seita.5G195700.1 | Sobic.003G185700.1 | Zm00001d011123_T001 |
| orthologID_006580 | PH01000384G0160 | LOC_Os01g47070.1 | Bradi2g45610.1 | HORVU3Hr1G059480.1 | TRIUR3_23643 | Seita.5G251600.1 | Sobic.003G244600.1 | Zm00001d043988_T001 |
| orthologID_006590 | PH01000071G0570 | LOC_Os01g66320.1 | Bradi2g57230.1 | HORVU3Hr1G090690.1 | TRIUR3_13158 | Seita.5G406900.1 | Sobic.003G381300.2 | Zm00001d042589_T001 |
| orthologID_006135 | PH01000317G1120 | LOC_Os01g65600.1 | Bradi2g56710.1 | HORVU3Hr1G081660.23 | TRIUR3_17158 | Seita.5G400600.1 | Sobic.003G375500.1 | Zm00001d012233_T001 |
| orthologID_006945 | PH01000090G1290 | LOC_Os01g63354.1 | Bradi2g55170.2 | HORVU3Hr1G085290.1 | TRIUR3_21556 | Seita.5G384900.1 | Sobic.003G359300.1 | Zm00001d012349_T001 |
| orthologID_006238 | PH01000568G0330 | LOC_Os01g47050.1 | Bradi2g45600.1 | HORVU3Hr1G059460.1 | TRIUR3_33769 | Seita.5G251400.1 | Sobic.003G244400.1 | Zm00001d043989_T001 |
| orthologID_006170 | PH01000504G0460 | LOC_Os01g21610.1 | Bradi2g12037.1 | HORVU6Hr1G019930.1 | TRIUR3_17000 | Seita.5G153400.1 | Sobic.003G150500.1 | Zm00001d009156_T001 |
| orthologID_006934 | PH01000209G0280 | LOC_Os01g16540.1 | Bradi2g10270.1 | HORVU3Hr1G038580.1 | TRIUR3_35263 | Seita.5G045000.1 | Sobic.003G128300.1 | Zm00001d008256_T003 |
| orthologID_006965 | PH01000244G0580 | LOC_Os01g67180.1 | Bradi2g57608.1 | HORVU3Hr1G089580.5 | TRIUR3_12338 | Seita.5G413800.1 | Sobic.003G388251.1 | Zm00001d042512_T006 |
| orthologID_006423 | PH01002444G0150 | LOC_Os01g25600.1 | Bradi2g12780.1 | HORVU3Hr1G021910.3 | TRIUR3_32322 | Seita.5G164200.1 | Sobic.003G160800.1 | Zm00001d040804_T001 |
| orthologID_006331 | PH01002825G0030 | LOC_Os01g07330.1 | Bradi2g04151.2 | HORVU3Hr1G017490.32 | TRIUR3_18526 | Seita.5G117800.1 | Sobic.003G055200.2 | Zm00001d000382_T001 |
| orthologID_006991 | PH01001261G0170 | LOC_Os01g42070.1 | Bradi2g42970.1 | HORVU3Hr1G052930.1 | TRIUR3_18136 | Seita.5G218400.1 | Sobic.003G212800.1 | Zm00001d044420_T001 |
| orthologID_006919 | PH01002021G0130 | LOC_Os01g38500.1 | Bradi2g41360.1 | HORVU3Hr1G048630.1 | TRIUR3_33980 | Seita.5G204900.1 | Sobic.003G191400.1 | Zm00001d044230_T001 |
| orthologID_006384 | PH01000014G1830 | LOC_Os01g18320.1 | Bradi2g11107.1 | HORVU5Hr1G011100.1 | TRIUR3_14614 | Seita.5G033900.1 | Sobic.003G137100.1 | Zm00001d008203_T001 |

| | | | | | | | | |
|---|---|---|---|---|---|---|---|---|
| orthologID_006361 | PH01000316G1140 | LOC_Os01g68000.1 | Bradi2g58100.1 | HORVU3Hr1G091930.1 | TRIUR3_34361 | Seita.5G420900.1 | Sobic.003G395200.1 | Zm00001d042445_T001 |
| orthologID_006714 | PH01000907G0170 | LOC_Os01g21850.1 | Bradi2g12090.1 | HORVU3Hr1G044780.1 | TRIUR3_19847 | Seita.5G153900.1 | Sobic.003G150900.1 | Zm00001d009150_T002 |
| orthologID_006824 | PH01003812G0050 | LOC_Os01g37832.1 | Bradi2g41230.3 | HORVU3Hr1G048310.2 | TRIUR3_32614 | Seita.5G201500.1 | Sobic.003G189700.2 | Zm00001d044497_T002 |
| orthologID_006339 | PH01000018G1610 | LOC_Os01g55120.1 | Bradi2g50210.3 | HORVU3Hr1G072760.3 | TRIUR3_35368 | Seita.5G322700.1 | Sobic.003G300000.1 | Zm00001d043421_T001 |
| orthologID_006481 | PH01000229G1370 | LOC_Os01g67560.1 | Bradi2g57880.1 | HORVU3Hr1G088990.3 | TRIUR3_17955 | Seita.5G417700.1 | Sobic.003G391900.1 | Zm00001d042476_T002 |
| orthologID_006404 | PH01000277G0530 | LOC_Os01g54880.1 | Bradi2g50030.1 | HORVU3Hr1G072140.4 | TRIUR3_15508 | Seita.5G320200.1 | Sobic.003G297500.1 | Zm00001d012737_T001 |
| orthologID_006865 | PH01000066G1320 | LOC_Os01g65320.1 | Bradi2g56490.1 | HORVU3Hr1G082240.19 | TRIUR3_23429 | Seita.5G397700.1 | Sobic.003G372700.1 | Zm00001d042670_T027 |
| orthologID_006508 | PH01005955G0040 | LOC_Os01g11260.1 | Bradi2g06760.4 | HORVU3Hr1G028720.3 | TRIUR3_31596 | Seita.5G148100.1 | Sobic.003G022000.1 | Zm00001d039385_T001 |
| orthologID_006140 | PH01001213G0370 | LOC_Os01g69060.1 | Bradi2g58912.1 | HORVU3Hr1G093990.1 | TRIUR3_05973 | Seita.5G429500.1 | Sobic.003G403800.4 | Zm00001d042346_T001 |
| orthologID_006434 | PH01000336G0830 | LOC_Os01g55600.1 | Bradi2g50570.1 | HORVU3Hr1G073520.3 | TRIUR3_04583 | Seita.5G326800.1 | Sobic.003G304000.1 | Zm00001d043375_T001 |
| orthologID_006133 | PH01000099G0410 | LOC_Os01g55830.1 | Bradi2g50760.4 | HORVU3Hr1G074040.3 | TRIUR3_28363 | Seita.5G328300.1 | Sobic.003G305700.1 | Zm00001d012675_T001 |
| orthologID_006223 | PH01000627G0240 | LOC_Os01g53840.1 | Bradi2g49400.1 | HORVU3Hr1G070600.19 | TRIUR3_34616 | Seita.5G312500.1 | Sobic.003G291000.3 | Zm00001d043516_T001 |
| orthologID_006328 | PH01001028G0640 | LOC_Os01g39290.1 | Bradi2g41640.1 | HORVU3Hr1G049940.2 | TRIUR3_35042 | Seita.5G207900.1 | Sobic.003G195300.1 | Zm00001d011217_T001 |
| orthologID_007156 | PH01001126G0450 | LOC_Os12g13150.1 | Bradi4g33310.1 | HORVU6Hr1G080600.1 | TRIUR3_09235 | Seita.3G090800.1 | Sobic.003G233400.2 | Zm00001d011447_T001 |
| orthologID_006969 | PH01003318G0130 | LOC_Os01g74170.1 | Bradi2g62257.2 | HORVU3Hr1G115810.9 | TRIUR3_11542 | Seita.5G468900.1 | Sobic.003G444300.3 | Zm00001d041956_T002 |
| orthologID_006887 | PH01000441G0440 | LOC_Os01g07730.1 | Bradi2g04447.2 | HORVU3Hr1G024590.1 | TRIUR3_17628 | Seita.5G121600.1 | Sobic.003G050800.1 | Zm00001d039674_T001 |
| orthologID_006216 | PH01000015G2060 | LOC_Os01g05080.1 | Bradi2g02930.2 | HORVU3Hr1G011620.11 | TRIUR3_04710 | Seita.5G099500.1 | Sobic.003G074700.1 | Zm00001d008612_T002 |
| orthologID_007574 | PH01002470G0090 | LOC_Os06g26234.1 | Bradi1g41970.1 | HORVU7Hr1G082830.1 | TRIUR3_18316 | Seita.4G152500.1 | Sobic.003G213800.2 | Zm00001d011301_T001 |
| orthologID_006880 | PH01000002G2620 | LOC_Os01g70060.1 | Bradi2g59297.3 | HORVU3Hr1G095520.4 | TRIUR3_33286 | Seita.5G434900.1 | Sobic.003G409200.1 | Zm00001d011975_T002 |
| orthologID_006329 | PH01000098G0690 | LOC_Os01g06550.1 | Bradi2g00226.1 | HORVU3Hr1G025750.1 | TRIUR3_19081 | Seita.5G112600.1 | Sobic.003G062300.1 | Zm00001d008648_T003 |
| orthologID_006427 | PH01001932G0190 | LOC_Os01g12520.1 | Bradi2g07420.2 | HORVU3Hr1G030460.1 | TRIUR3_31041 | Seita.5G015100.1 | Sobic.003G012300.1 | Zm00001d008974_T001 |
| orthologID_006449 | PH01001208G0360 | LOC_Os01g44210.1 | Bradi2g44440.1 | HORVU3Hr1G056580.3 | TRIUR3_26998 | Seita.5G237100.1 | Sobic.003G230400.1 | Zm00001d044130_T001 |
| orthologID_006785 | PH01000214G0590 | LOC_Os01g53600.1 | Bradi2g49200.1 | HORVU3Hr1G070050.7 | TRIUR3_24874 | Seita.5G310300.1 | Sobic.003G289400.1 | Zm00001d043555_T002 |
| orthologID_006876 | PH01001797G0270 | LOC_Os01g15540.1 | Bradi2g09450.2 | HORVU3Hr1G036020.1 | TRIUR3_13671 | Seita.5G055100.1 | Sobic.003G119200.2 | Zm00001d040312_T001 |
| orthologID_006659 | PH01000058G1580 | LOC_Os01g72780.1 | Bradi2g61300.1 | HORVU3Hr1G110140.7 | TRIUR3_10775 | Seita.5G459200.1 | Sobic.003G432300.1 | Zm00001d042045_T001 |
| orthologID_006677 | PH01000002G1270 | LOC_Os01g69250.1 | Bradi2g59080.1 | HORVU7Hr1G075380.8 | TRIUR3_13562 | Seita.5G431500.1 | Sobic.003G405900.1 | Zm00001d042329_T001 |
| orthologID_007184 | PH01000250G0800 | LOC_Os09g09480.2 | Bradi4g26920.2 | HORVU5Hr1G052710.6 | TRIUR3_27429 | Seita.5G226800.1 | Sobic.003G220600.1 | Zm00001d044480_T001 |
| orthologID_009930 | PH01001669G0130 | LOC_Os01g47670.1 | Bradi2g46020.1 | HORVU3Hr1G062150.1 | TRIUR3_22118 | Seita.5G268900.1 | Sobic.003G255000.1 | Zm00001d011590_T001 |
| orthologID_006537 | PH01000244G0210 | LOC_Os01g67040.1 | Bradi2g57550.1 | HORVU3Hr1G089820.1 | TRIUR3_12967 | Seita.5G412700.1 | Sobic.003G386700.2 | Zm00001d012169_T001 |
| orthologID_006560 | PH01003429G0140 | LOC_Os01g10010.2 | Bradi2g05960.1 | HORVU3Hr1G018940.1 | TRIUR3_09694 | Seita.5G139000.1 | Sobic.003G030800.1 | Zm00001d039455_T002 |
| orthologID_006879 | PH01001169G0110 | LOC_Os01g07910.1 | Bradi2g04610.1 | HORVU3Hr1G024170.3 | TRIUR3_27879 | Seita.5G123100.1 | Sobic.003G049200.1 | Zm00001d039656_T003 |
| orthologID_006814 | PH01000402G0730 | LOC_Os01g56350.3 | Bradi2g51140.1 | HORVU3Hr1G074850.25 | TRIUR3_14919 | Seita.5G332400.1 | Sobic.003G309500.1 | Zm00001d043325_T001 |
| orthologID_006750 | PH01000333G0270 | LOC_Os01g53240.1 | Bradi2g49000.2 | HORVU3Hr1G069650.1 | TRIUR3_07593 | Seita.5G307500.1 | Sobic.003G287000.1 | Zm00001d043538_T001 |
| orthologID_006623 | PH01000633G0760 | LOC_Os01g40110.1 | Bradi2g41957.3 | HORVU3Hr1G050320.1 | TRIUR3_32500 | Seita.5G257500.1 | Sobic.003G198300.1 | Zm00001d011199_T002 |
| orthologID_006913 | PH01000032G0980 | LOC_Os03g02420.1 | Bradi2g52180.1 | HORVU4Hr1G015420.3 | TRIUR3_24423 | Seita.5G345000.1 | Sobic.003G321700.1 | Zm00001d043229_T001 |
| orthologID_006152 | PH01000783G0010 | LOC_Os01g51140.1 | Bradi2g47810.1 | HORVU3Hr1G066600.1 | TRIUR3_19402 | Seita.5G289400.1 | Sobic.003G273200.2 | Zm00001d043699_T001 |
| orthologID_006669 | PH01000066G0580 | LOC_Os01g65560.1 | Bradi2g56682.1 | HORVU3Hr1G081760.1 | TRIUR3_16173 | Seita.5G400300.1 | Sobic.003G375100.2 | Zm00001d042640_T001 |
| orthologID_006365 | PH01000111G0040 | LOC_Os01g65730.1 | Bradi2g56792.2 | HORVU3Hr1G091460.10 | TRIUR3_17186 | Seita.5G401600.1 | Sobic.003G376500.2 | Zm00001d012226_T001 |
| orthologID_006812 | PH01000015G1560 | LOC_Os01g04930.1 | Bradi2g02810.1 | HORVU3Hr1G010600.2 | TRIUR3_09932 | Seita.5G098300.1 | Sobic.003G076500.2 | Zm00001d008602_T001 |
| orthologID_006202 | PH01000015G2470 | LOC_Os01g05480.3 | Bradi2g03160.1 | HORVU3Hr1G013360.6 | TRIUR3_22283 | Seita.5G102400.1 | Sobic.003G072400.1 | Zm00001d039866_T001 |
| orthologID_006197 | PH01000015G2120 | LOC_Os01g05100.1 | Bradi2g02950.2 | HORVU3Hr1G011660.5 | TRIUR3_05801 | Seita.5G099800.1 | Sobic.003G074300.1 | Zm00001d039878_T003 |
| orthologID_010196 | PH01000252G0950 | LOC_Os01g54050.1 | Bradi2g49560.2 | HORVU3Hr1G070880.7 | TRIUR3_27991 | Seita.5G314500.1 | Sobic.003G292600.2 | Zm00001d043505_T001 |
| orthologID_008986 | PH01001656G0300 | LOC_Os04g02680.1 | Bradi5g01760.1 | HORVU2Hr1G001310.11 | TRIUR3_07334 | Seita.5G136000.1 | Sobic.003G034100.1 | Zm00001d039489_T002 |
| orthologID_006473 | PH01003561G0100 | LOC_Os01g63950.1 | Bradi2g55520.1 | HORVU3Hr1G084420.2 | TRIUR3_27707 | Seita.5G388800.1 | Sobic.003G363000.2 | Zm00001d042787_T001 |
| orthologID_006761 | PH01000443G0450 | LOC_Os01g12440.1 | Bradi2g07357.1 | HORVU3Hr1G030310.1 | TRIUR3_26876 | Seita.5G015800.1 | Sobic.003G012800.1 | Zm00001d039324_T001 |
| orthologID_006571 | PH01000090G1260 | LOC_Os01g63410.1 | Bradi2g55200.1 | HORVU3Hr1G085230.2 | TRIUR3_13642 | Seita.5G385100.1 | Sobic.003G359500.1 | Zm00001d012339_T001 |
| orthologID_006638 | PH01003546G0120 | LOC_Os03g47550.1 | Bradi2g02940.1 | HORVU3Hr1G011650.4 | TRIUR3_05800 | Seita.5G099700.1 | Sobic.003G074400.1 | Zm00001d039879_T001 |
| orthologID_006319 | PH01000076G0800 | LOC_Os01g52540.1 | Bradi2g48597.1 | HORVU3Hr1G068640.1 | TRIUR3_11727 | Seita.5G302100.1 | Sobic.003G281300.1 | Zm00001d011745_T001 |
| orthologID_006960 | PH01000775G0490 | LOC_Os01g56660.1 | Bradi2g51463.1 | HORVU3Hr1G075770.11 | TRIUR3_06422 | Seita.5G336500.1 | Sobic.003G312600.1 | Zm00001d043300_T001 |
| orthologID_006631 | PH01002707G0300 | LOC_Os01g36540.1 | Bradi2g40625.1 | HORVU3Hr1G084510.2 | TRIUR3_15047 | Seita.5G192500.1 | Sobic.003G183900.1 | Zm00001d044537_T001 |

| | | | | | | | | |
|---|---|---|---|---|---|---|---|---|
| orthologID_006817 | PH01000252G0840 | LOC_Os01g54020.1 | Bradi2g49520.1 | HORVU3Hr1G070780.4 | TRIUR3_09682 | Seita.5G314200.1 | Sobic.003G292300.1 | Zm00001d043508_T002 |
| orthologID_006148 | PH01000090G0900 | LOC_Os01g63500.1 | Bradi2g55260.1 | HORVU3Hr1G085080.3 | TRIUR3_30955 | Seita.5G385800.1 | Sobic.003G360100.1 | Zm00001d042822_T001 |
| orthologID_006218 | PH01001169G0070 | LOC_Os01g07880.1 | Bradi2g04590.1 | HORVU3Hr1G024220.2 | TRIUR3_07158 | Seita.5G122900.1 | Sobic.003G049400.1 | Zm00001d008734_T001 |
| orthologID_006382 | PH01000489G0400 | LOC_Os01g68770.1 | Bradi2g58677.2 | HORVU3Hr1G093350.2 | TRIUR3_04843 | Seita.5G427000.1 | Sobic.003G401200.1 | Zm00001d042367_T002 |
| orthologID_006775 | PH01000056G0160 | LOC_Os01g07250.1 | Bradi2g04110.1 | HORVU3Hr1G017580.1 | TRIUR3_29270 | Seita.5G117400.1 | Sobic.003G055700.1 | Zm00001d039719_T002 |
| orthologID_006773 | PH01000072G1610 | LOC_Os01g56990.1 | Bradi2g51670.1 | HORVU3Hr1G076310.2 | TRIUR3_03753 | Seita.5G338900.1 | Sobic.003G315332.1 | Zm00001d043280_T001 |
| orthologID_006952 | PH01001461G0500 | LOC_Os01g11990.1 | Bradi2g07122.1 | HORVU3Hr1G029340.3 | TRIUR3_16161 | Seita.5G020200.1 | Sobic.003G016200.1 | Zm00001d039352_T001 |
| orthologID_006378 | PH01000054G0470 | LOC_Os01g62440.1 | Bradi2g37760.1 | HORVU1Hr1G010820.13 | TRIUR3_25976 | Seita.5G377400.1 | Sobic.003G352500.1 | Zm00001d042908_T002 |
| orthologID_006491 | PH01000004G1570 | LOC_Os02g44132.2 | Bradi2g55807.1 | HORVU3Hr1G083750.1 | TRIUR3_28797 | Seita.5G391400.1 | Sobic.003G365700.1 | Zm00001d012284_T001 |
| orthologID_006277 | PH01000632G0190 | LOC_Os01g14830.1 | Bradi2g08840.2 | HORVU3Hr1G034240.1 | TRIUR3_20113 | Seita.5G061400.1 | Sobic.003G113400.1 | Zm00001d008360_T001 |
| orthologID_006575 | PH01001878G0010 | LOC_Os05g34880.1 | Bradi2g24967.1 | HORVU1Hr1G063720.6 | TRIUR3_11848 | Seita.3G236900.1 | Sobic.003G029500.1 | Zm00001d008865_T002 |
| orthologID_006193 | PH01000267G0590 | LOC_Os01g57440.1 | Bradi2g51880.2 | HORVU3Hr1G077010.3 | TRIUR3_19233 | Seita.5G341900.1 | Sobic.003G318900.1 | Zm00001d012611_T001 |
| orthologID_006465 | PH01000032G0550 | LOC_Os01g57730.1 | Bradi2g52077.1 | HORVU3Hr1G077580.4 | TRIUR3_32734 | Seita.5G344000.1 | Sobic.003G320800.1 | Zm00001d043238_T001 |
| orthologID_006917 | PH01000425G0820 | LOC_Os01g57930.1 | Bradi2g52187.1 | HORVU3Hr1G077710.2 | TRIUR3_31834 | Seita.5G345600.1 | Sobic.003G321900.2 | Zm00001d043228_T001 |
| orthologID_006507 | PH01000009G2590 | LOC_Os01g60830.1 | Bradi2g53620.1 | HORVU3Hr1G081280.1 | TRIUR3_18531 | Seita.5G363500.1 | Sobic.003G339200.1 | Zm00001d043045_T001 |
| orthologID_006578 | PH01000581G0380 | LOC_Os01g49740.1 | Bradi2g47130.1 | HORVU3Hr1G064340.3 | TRIUR3_14695 | Seita.5G279700.1 | Sobic.003G264700.1 | Zm00001d043786_T001 |
| orthologID_006564 | PH01002164G0300 | LOC_Os01g47650.1 | Bradi2g45987.1 | HORVU3Hr1G060780.1 | TRIUR3_15429 | Seita.5G262500.1 | Sobic.003G249600.1 | Zm00001d011532_T002 |
| orthologID_006297 | PH01000015G1390 | LOC_Os01g04870.1 | Bradi2g02750.1 | HORVU3Hr1G010370.2 | TRIUR3_09514 | Seita.5G097300.1 | Sobic.003G077600.1 | Zm00001d008597_T001 |
| orthologID_006767 | PH01000284G1160 | LOC_Os01g54440.1 | Bradi2g49774.1 | HORVU3Hr1G071410.1 | TRIUR3_24777 | Seita.5G317100.1 | Sobic.003G294900.2 | Zm00001d043454_T001 |
| orthologID_006679 | PH01000018G1270 | LOC_Os01g55094.2 | Bradi2g50180.2 | HORVU3Hr1G072640.15 | TRIUR3_14480 | Seita.5G322400.1 | Sobic.003G299500.4 | Zm00001d043425_T003 |
| orthologID_006563 | PH01000633G0470 | LOC_Os01g40050.2 | Bradi2g41910.2 | HORVU3Hr1G050400.4 | TRIUR3_14726 | Seita.6G138900.1 | Sobic.003G197800.1 | Zm00001d044297_T002 |
| orthologID_006836 | PH01005670G0050 | LOC_Os01g40630.1 | Bradi2g42190.1 | HORVU3Hr1G050770.10 | TRIUR3_21353 | Seita.5G208900.1 | Sobic.003G202200.1 | Zm00001d011246_T002 |
| orthologID_006422 | PH01002356G0150 | LOC_Os01g47400.1 | Bradi2g45790.1 | HORVU3Hr1G060060.8 | TRIUR3_25427 | Seita.5G260400.1 | Sobic.003G247000.1 | Zm00001d043965_T001 |
| orthologID_006861 | PH01000592G0720 | LOC_Os01g08270.1 | Bradi2g04860.7 | HORVU3Hr1G022770.2 | TRIUR3_35425 | Seita.5G125800.1 | Sobic.003G045600.1 | Zm00001d008744_T007 |
| orthologID_006325 | PH01001281G0100 | LOC_Os01g59080.1 | Bradi2g52720.1 | HORVU3Hr1G078830.2 | TRIUR3_09836 | Seita.1G000200.1 | Sobic.003G328100.1 | Zm00001d043168_T001 |
| orthologID_006139 | PH01000907G0160 | LOC_Os01g21860.1 | Bradi2g12100.1 | HORVU3Hr1G020810.1 | TRIUR3_10780 | Seita.5G154000.1 | Sobic.003G151000.1 | Zm00001d009147_T002 |
| orthologID_006159 | PH01006766G0010 | LOC_Os01g73230.1 | Bradi2g61570.1 | HORVU3Hr1G111910.7 | TRIUR3_25666 | Seita.5G463400.1 | Sobic.003G437500.1 | Zm00001d048147_T002 |
| orthologID_006552 | PH01000144G1500 | LOC_Os01g73500.1 | Bradi2g61767.1 | HORVU3Hr1G113020.2 | TRIUR3_02071 | Seita.9G386600.1 | Sobic.003G440300.1 | Zm00001d011799_T001 |
| orthologID_006488 | PH01001208G0220 | LOC_Os01g44360.1 | Bradi2g44510.3 | HORVU5Hr1G012770.3 | TRIUR3_24553 | Seita.5G238300.1 | Sobic.003G231600.1 | Zm00001d044119_T001 |
| orthologID_006540 | PH01004678G0090 | LOC_Os01g61440.1 | Bradi2g54000.1 | HORVU3Hr1G087770.2 | TRIUR3_06047 | Seita.5G368100.1 | Sobic.003G343900.1 | Zm00001d042996_T001 |
| orthologID_006475 | PH01002896G0090 | LOC_Os01g48700.1 | Bradi2g46527.2 | HORVU7Hr1G037090.2 | TRIUR3_22279 | Seita.5G270300.1 | Sobic.003G256300.2 | Zm00001d011597_T002 |
| orthologID_006352 | PH01001112G0470 | LOC_Os01g58390.1 | Bradi2g52356.2 | HORVU3Hr1G078090.1 | TRIUR3_17684 | Seita.5G347600.1 | Sobic.003G324100.1 | Zm00001d043207_T004 |
| orthologID_006317 | PH01000009G2140 | LOC_Os01g60660.1 | Bradi2g53530.1 | HORVU3Hr1G081060.4 | TRIUR3_17399 | Seita.5G362200.1 | Sobic.003G338100.1 | Zm00001d012503_T003 |
| orthologID_006745 | PH01000004G2200 | LOC_Os01g64030.2 | Bradi2g55580.3 | HORVU3Hr1G084380.1 | TRIUR3_12646 | Seita.5G389600.1 | Sobic.003G363700.1 | Zm00001d042774_T001 |
| orthologID_006620 | PH01005281G0050 | LOC_Os01g47270.1 | Bradi2g45710.1 | HORVU3Hr1G059720.1 | TRIUR3_30876 | Seita.5G253200.1 | Sobic.003G246200.1 | Zm00001d043974_T001 |
| orthologID_006441 | PH01000090G1040 | LOC_Os01g63470.1 | Bradi2g55240.1 | HORVU3Hr1G085130.1 | TRIUR3_15441 | Seita.5G385500.1 | Sobic.003G359900.1 | Zm00001d012336_T001 |
| orthologID_006522 | PH01003657G0070 | LOC_Os01g16152.1 | Bradi2g09910.1 | HORVU3Hr1G037720.1 | TRIUR3_20117 | Seita.5G049600.1 | Sobic.003G124300.2 | Zm00001d040341_T001 |
| orthologID_009988 | PH01001213G0390 | LOC_Os01g69040.1 | Bradi2g58870.3 | HORVU3Hr1G093850.2 | TRIUR3_27301 | Seita.5G429300.1 | Sobic.003G403400.1 | Zm00001d042350_T002 |
| orthologID_009732 | PH01000186G1170 | LOC_Os10g34310.1 | Bradi3g29181.1 | HORVU1Hr1G042440.2 | TRIUR3_33175 | Seita.9G216700.1 | Sobic.003G038200.1 | Zm00001d039548_T001 |
| orthologID_006199 | PH01001339G0550 | LOC_Os01g05830.1 | Bradi2g03370.1 | HORVU3Hr1G013860.2 | TRIUR3_15770 | Seita.5G104400.1 | Sobic.003G070700.1 | Zm00001d039851_T002 |
| orthologID_006872 | PH01000441G0690 | LOC_Os01g07540.1 | Bradi2g04317.14 | HORVU3Hr1G016360.19 | TRIUR3_22183 | Seita.5G120200.1 | Sobic.003G052200.2 | Zm00001d039683_T009 |
| orthologID_006947 | PH01000066G0890 | LOC_Os01g65450.1 | Bradi2g56587.2 | HORVU3Hr1G082010.1 | TRIUR3_10489 | Seita.5G399100.1 | Sobic.003G374000.1 | Zm00001d012244_T001 |
| orthologID_006164 | PH01000235G0630 | LOC_Os01g67120.1 | Bradi2g57590.1 | HORVU3Hr1G089650.2 | TRIUR3_12336 | Seita.5G413400.1 | Sobic.003G387800.1 | Zm00001d042526_T001 |
| orthologID_006435 | PH01000357G0520 | LOC_Os01g07800.2 | Bradi2g04530.1 | HORVU3Hr1G024390.3 | TRIUR3_16545 | Seita.5G122300.1 | Sobic.003G050000.1 | Zm00001d039667_T001 |
| orthologID_006520 | PH01000223G1150 | LOC_Os01g62910.1 | Bradi2g54927.3 | HORVU3Hr1G085770.2 | TRIUR3_07817 | Seita.5G381200.1 | Sobic.003G356100.1 | Zm00001d039873_T002 |
| orthologID_006314 | PH01001281G0230 | LOC_Os01g59110.1 | Bradi2g52750.1 | HORVU3Hr1G078860.2 | TRIUR3_13826 | Seita.5G352600.1 | Sobic.003G328400.1 | Zm00001d043165_T002 |
| orthologID_006143 | PH01000966G0470 | LOC_Os01g10850.1 | Bradi2g06497.1 | HORVU3Hr1G026010.5 | TRIUR3_18632 | Seita.5G145500.1 | Sobic.003G024700.1 | Zm00001d008898_T001 |
| orthologID_007343 | PH01001205G0060 | LOC_Os11g13850.2 | Bradi4g21270.1 | HORVU4Hr1G015330.1 | TRIUR3_12677 | Seita.8G081500.1 | Sobic.003G409700.2 | Zm00001d042289_T001 |
| orthologID_006685 | PH01001317G0490 | LOC_Os01g15490.1 | Bradi2g09380.1 | HORVU3Hr1G035900.1 | TRIUR3_22519 | Seita.5G055900.1 | Sobic.003G118200.1 | Zm00001d008320_T004 |
| orthologID_006477 | PH01000667G0350 | LOC_Os01g13740.1 | Bradi2g08287.1 | HORVU3Hr1G032440.2 | TRIUR3_07027 | Seita.5G003100.1 | Sobic.003G002600.1 | Zm00001d039260_T002 |

| | | | | | | | | |
|---|---|---|---|---|---|---|---|---|
| orthologID_006231 | PH01002312G0230 | LOC_Os01g72570.1 | Bradi2g61190.4 | HORVU3Hr1G109300.13 | TRIUR3_26464 | Seita.5G457300.1 | Sobic.003G430900.1 | Zm00001d042055_T001 |
| orthologID_006363 | PH01000783G0510 | LOC_Os01g50930.1 | Bradi2g47720.4 | HORVU3Hr1G066380.1 | TRIUR3_28652 | Seita.5G288000.1 | Sobic.003G271900.3 | Zm00001d043707_T002 |
| orthologID_006707 | PH01000031G1690 | LOC_Os01g68010.1 | Bradi2g58110.1 | HORVU3Hr1G091910.5 | TRIUR3_19303 | Seita.5G421000.1 | Sobic.003G395300.1 | Zm00001d012126_T004 |
| orthologID_006804 | PH01001385G0420 | LOC_Os01g16080.1 | Bradi2g09800.1 | HORVU3Hr1G037180.3 | TRIUR3_22005 | Seita.5G051700.1 | Sobic.003G122400.1 | Zm00001d040348_T002 |
| orthologID_006326 | PH01000260G0890 | LOC_Os01g01770.1 | Bradi2g00677.1 | HORVU4Hr1G083140.1 | TRIUR3_08176 | Seita.5G072200.1 | Sobic.003G102800.1 | Zm00001d040158_T001 |
| orthologID_006166 | PH01001208G0460 | LOC_Os01g44090.1 | Bradi2g44390.1 | HORVU3Hr1G056800.4 | TRIUR3_14153 | Seita.5G236400.1 | Sobic.003G229800.1 | Zm00001d011431_T001 |
| orthologID_006430 | PH01000781G0130 | LOC_Os01g73270.1 | Bradi2g61649.1 | HORVU3Hr1G112480.2 | TRIUR3_07689 | Seita.5G463900.1 | Sobic.003G438100.1 | Zm00001d011812_T001 |
| orthologID_007008 | PH01004235G0060 | LOC_Os01g69910.1 | Bradi2g59137.1 | HORVU3Hr1G094860.1 | TRIUR3_22499 | Seita.5G433100.1 | Sobic.003G407200.1 | Zm00001d042313_T004 |
| orthologID_006920 | PH01000155G1150 | LOC_Os01g55974.1 | Bradi2g50870.1 | HORVU3Hr1G074280.2 | TRIUR3_10183 | Seita.9G511200.1 | Sobic.003G306900.2 | Zm00001d012671_T001 |
| orthologID_006151 | PH01003988G0090 | LOC_Os01g37842.1 | Bradi2g41240.1 | HORVU3Hr1G048340.1 | TRIUR3_25491 | Seita.5G201600.1 | Sobic.003G189800.1 | Zm00001d011167_T001 |
| orthologID_006298 | PH01001028G0620 | LOC_Os01g39270.1 | Bradi2g41630.1 | HORVU3Hr1G049900.2 | TRIUR3_14252 | Seita.5G207800.1 | Sobic.003G195150.1 | Zm00001d044267_T004 |
| orthologID_006857 | PH01000783G0230 | LOC_Os01g51120.1 | Bradi2g47800.1 | HORVU3Hr1G066580.1 | TRIUR3_19403 | Seita.5G289300.1 | Sobic.003G273100.1 | Zm00001d011688_T001 |
| orthologID_006771 | PH01002733G0270 | LOC_Os01g21120.1 | Bradi2g11890.1 | HORVU5Hr1G080790.1 | TRIUR3_34295 | Seita.5G022000.1 | Sobic.003G148600.1 | Zm00001d040651_T001 |
| orthologID_006596 | PH01004414G0170 | LOC_Os01g63220.2 | Bradi2g55077.2 | HORVU3Hr1G085480.4 | TRIUR3_04898 | Seita.5G383200.1 | Sobic.003G358000.4 | Zm00001d042847_T001 |
| orthologID_006406 | PH01000004G3220 | LOC_Os01g63780.1 | Bradi2g55351.1 | HORVU3Hr1G084740.1 | TRIUR3_23836 | Seita.5G387300.1 | Sobic.003G361400.1 | Zm00001d012314_T005 |
| orthologID_006258 | PH01001004G0040 | LOC_Os01g40200.1 | Bradi2g41973.5 | HORVU3Hr1G011330.8 | TRIUR3_10824 | Seita.5G257300.1 | Sobic.003G198500.2 | Zm00001d011197_T007 |
| orthologID_008740 | PH01004186G0030 | LOC_Os06g22700.1 | Bradi41430s00200.1 | HORVU7Hr1G081490.3 | TRIUR3_33005 | Seita.4G161500.1 | Sobic.003G361500.1 | Zm00001d042807_T001 |
| orthologID_006683 | PH01000094G1490 | LOC_Os01g16330.1 | Bradi2g10080.1 | HORVU3Hr1G038170.2 | TRIUR3_21909 | Seita.5G047600.1 | Sobic.003G126300.1 | Zm00001d040411_T001 |
| orthologID_006172 | PH01001737G0190 | LOC_Os01g61200.1 | Bradi2g53830.1 | HORVU3Hr1G088120.1 | TRIUR3_02407 | Seita.5G366300.1 | Sobic.003G341900.1 | Zm00001d012476_T001 |
| orthologID_006451 | PH01000209G0010 | LOC_Os01g16310.1 | Bradi2g10060.1 | HORVU3Hr1G038060.10 | TRIUR3_33100 | Seita.5G048300.1 | Sobic.003G126100.1 | Zm00001d008273_T001 |
| orthologID_006639 | PH01000214G0650 | LOC_Os01g53630.1 | Bradi2g49226.1 | HORVU3Hr1G070080.5 | TRIUR3_19687 | Seita.5G310600.1 | Sobic.003G289700.1 | Zm00001d043560_T006 |
| orthologID_009964 | PH01000141G0120 | LOC_Os01g66120.1 | Bradi2g57100.1 | HORVU3Hr1G090920.1 | TRIUR3_14109 | Seita.5G405100.1 | Sobic.003G379700.1 | Zm00001d042609_T001 |
| orthologID_006484 | PH01000775G0640 | LOC_Os01g56670.1 | Bradi2g51467.1 | HORVU3Hr1G075850.2 | TRIUR3_18747 | Seita.5G336600.1 | Sobic.003G312700.1 | Zm00001d012643_T007 |
| orthologID_006968 | PH01000209G0210 | LOC_Os01g16650.1 | Bradi2g10220.1 | HORVU3Hr1G038430.3 | TRIUR3_21860 | Seita.5G046000.1 | Sobic.003G127600.1 | Zm00001d008260_T001 |
| orthologID_009676 | PH01000738G0100 | LOC_Os08g21840.1 | Bradi3g20440.1 | HORVU4Hr1G078100.1 | TRIUR3_10050 | Seita.5G029300.1 | Sobic.003G141700.1 | Zm00001d040588_T004 |
| orthologID_006316 | PH01000872G0090 | LOC_Os01g47450.1 | Bradi2g45830.1 | HORVU3Hr1G060150.6 | TRIUR3_24690 | Seita.5G260800.1 | Sobic.003G247400.2 | Zm00001d011518_T001 |
| orthologID_006402 | PH01000578G0420 | LOC_Os01g19800.1 | Bradi2g11577.1 | HORVU1Hr1G052850.6 | TRIUR3_32903 | Seita.5G026700.1 | Sobic.003G144600.1 | Zm00001d040614_T002 |
| orthologID_006729 | PH01000260G1140 | LOC_Os01g01970.2 | Bradi2g00870.3 | HORVU0Hr1G005960.2 | TRIUR3_25255 | Seita.5G073500.1 | Sobic.003G101300.2 | Zm00001d040124_T001 |
| orthologID_006165 | PH01000357G1080 | LOC_Os01g07480.2 | Bradi2g04270.7 | HORVU3Hr1G016690.1 | TRIUR3_10339 | Seita.5G119400.1 | Sobic.003G052900.1 | Zm00001d039694_T001 |
| orthologID_006334 | PH01000032G0100 | LOC_Os01g57580.1 | Bradi2g51990.1 | HORVU3Hr1G077260.2 | TRIUR3_07341 | Seita.5G342800.1 | Sobic.003G319800.1 | Zm00001d043248_T001 |
| orthologID_009606 | PH01002367G0070 | LOC_Os01g72200.1 | Bradi3g55370.2 | HORVU3Hr1G107330.1 | TRIUR3_02660 | Seita.5G453700.1 | Sobic.003G426800.1 | Zm00001d011873_T001 |
| orthologID_006155 | PH01004678G0100 | LOC_Os01g61430.1 | Bradi2g53990.1 | HORVU3Hr1G087780.10 | TRIUR3_30599 | Seita.5G368000.1 | Sobic.003G343700.1 | Zm00001d042997_T001 |
| orthologID_006600 | PH01000037G0560 | LOC_Os01g11350.1 | Bradi2g06790.2 | HORVU3Hr1G040920.4 | TRIUR3_20118 | Seita.5G148300.1 | Sobic.003G021800.2 | Zm00001d039383_T001 |
| orthologID_006781 | PH01000425G0730 | LOC_Os01g57880.1 | Bradi2g52155.1 | HORVU3Hr1G077680.1 | TRIUR3_16942 | Seita.5G344700.1 | Sobic.003G321400.1 | Zm00001d012602_T001 |
| orthologID_006463 | PH01005670G0060 | LOC_Os01g40620.1 | Bradi2g42180.2 | HORVU3Hr1G050760.1 | TRIUR3_21352 | Seita.5G208800.1 | Sobic.003G202100.1 | Zm00001d011245_T001 |
| orthologID_006757 | PH01001234G0470 | LOC_Os01g42800.1 | Bradi2g43437.2 | HORVU3Hr1G053910.4 | TRIUR3_11740 | Seita.5G225000.1 | Sobic.003G219200.1 | Zm00001d044468_T002 |
| orthologID_006604 | PH01000018G2550 | LOC_Os01g55540.1 | Bradi2g50500.1 | HORVU3Hr1G073220.3 | TRIUR3_04934 | Seita.5G326000.1 | Sobic.003G303300.1 | Zm00001d043382_T001 |
| orthologID_006432 | PH01000990G0300 | LOC_Os01g52390.1 | Bradi2g48500.1 | HORVU3Hr1G068500.2 | TRIUR3_33029 | Seita.5G301000.1 | Sobic.003G279800.1 | Zm00001d043632_T001 |
| orthologID_006859 | PH01002329G0250 | LOC_Os01g40550.1 | Bradi2g42130.1 | HORVU3Hr1G051000.1 | TRIUR3_14496 | Seita.5G254800.1 | Sobic.003G201400.2 | Zm00001d011241_T001 |
| orthologID_006358 | PH01001186G0590 | LOC_Os01g06460.1 | Bradi2g00310.1 | HORVU3Hr1G025590.1 | TRIUR3_09567 | Seita.5G111800.1 | Sobic.003G063200.1 | Zm00001d039776_T001 |
| orthologID_006910 | PH01000701G0600 | LOC_Os01g52960.1 | Bradi2g48860.1 | HORVU3Hr1G069240.1 | TRIUR3_18360 | Seita.5G305400.1 | Sobic.003G284400.1 | Zm00001d043572_T001 |
| orthologID_006640 | PH01001488G0350 | LOC_Os01g09252.1 | Bradi2g05567.2 | HORVU1Hr1G020470.15 | TRIUR3_16797 | Seita.5G133600.1 | Sobic.003G036800.1 | Zm00001d039521_T002 |
| orthologID_006627 | PH01001880G0260 | LOC_Os01g46430.1 | Bradi2g45370.1 | HORVU3Hr1G058900.1 | TRIUR3_19247 | Seita.5G247200.1 | Sobic.003G240500.1 | Zm00001d044030_T001 |
| orthologID_006954 | PH01000393G0240 | LOC_Os01g71256.1 | Bradi2g60357.1 | HORVU3Hr1G104070.4 | TRIUR3_32455 | Seita.5G447200.1 | Sobic.003G420600.2 | Zm00001d011895_T001 |
| orthologID_006972 | PH01000906G0290 | LOC_Os01g09120.1 | Bradi2g05520.1 | HORVU3Hr1G020100.2 | TRIUR3_34216 | Seita.5G132900.1 | Sobic.003G037300.1 | Zm00001d039530_T001 |
| orthologID_006373 | PH01000632G0810 | LOC_Os01g15110.1 | Bradi2g09120.1 | HORVU3Hr1G035380.1 | TRIUR3_33439 | Seita.5G058000.1 | Sobic.003G116600.1 | Zm00001d040286_T003 |
| orthologID_006121 | PH01000622G0660 | LOC_Os01g74010.1 | Bradi2g62090.1 | HORVU3Hr1G114370.2 | TRIUR3_22116 | Seita.5G468000.1 | Sobic.003G443500.1 | Zm00001d041961_T001 |
| orthologID_006998 | PH01000235G0100 | LOC_Os01g66690.1 | Bradi2g57415.1 | HORVU3Hr1G090150.2 | TRIUR3_25865 | Seita.5G409300.1 | Sobic.003G384100.1 | Zm00001d042558_T002 |
| orthologID_006608 | PH01000260G0780 | LOC_Os01g01740.1 | Bradi2g00670.1 | HORVU3Hr1G000350.4 | TRIUR3_02741 | Seita.5G072100.1 | Sobic.003G103000.1 | Zm00001d040160_T001 |
| orthologID_006205 | PH01004745G0050 | LOC_Os01g08820.1 | Bradi2g05320.1 | HORVU3Hr1G020740.2 | TRIUR3_16855 | Seita.5G130600.1 | Sobic.003G040100.1 | Zm00001d039579_T001 |

| | | | | | | | | |
|---|---|---|---|---|---|---|---|---|
| orthologID_006377 | PH01000002G2740 | LOC_Os01g70140.1 | Bradi2g59350.1 | HORVU3Hr1G095890.1 | TRIUR3_08122 | Seita.5G436600.1 | Sobic.003G410200.1 | Zm00001d011966_T003 |
| orthologID_006591 | PH01000309G1170 | LOC_Os01g64690.1 | Bradi2g56060.2 | HORVU3Hr1G083300.1 | TRIUR3_07508 | Seita.5G393500.1 | Sobic.003G367900.1 | Zm00001d042725_T001 |
| orthologID_006946 | PH01000214G0400 | LOC_Os01g53500.1 | Bradi2g49152.1 | HORVU3Hr1G069960.1 | TRIUR3_21646 | Seita.5G309700.1 | Sobic.003G288401.1 | Zm00001d024313_T001 |
| orthologID_006293 | PH01000155G0840 | LOC_Os01g55860.1 | Bradi2g50770.4 | HORVU3Hr1G074120.2 | TRIUR3_10090 | Seita.5G328500.1 | Sobic.003G305900.1 | Zm00001d043358_T001 |
| orthologID_006443 | PH01000358G0890 | LOC_Os01g09300.1 | Bradi2g05600.1 | HORVU3Hr1G019820.11 | TRIUR3_27418 | Seita.5G134100.1 | Sobic.003G036300.1 | Zm00001d008820_T002 |
| orthologID_009524 | PH01004316G0080 | LOC_Os10g11340.1 | Bradi3g22960.3 | HORVU6Hr1G013180.3 | TRIUR3_09325 | Seita.9G221400.1 | Sobic.003G376800.1 | Zm00001d042626_T001 |
| orthologID_006420 | PH01000814G0820 | LOC_Os01g53040.1 | Bradi2g48907.1 | HORVU3Hr1G069320.3 | TRIUR3_31683 | Seita.5G306000.1 | Sobic.003G285500.1 | Zm00001d043569_T001 |
| orthologID_006625 | PH01000263G0260 | LOC_Os01g55270.1 | Bradi2g50300.1 | HORVU3Hr1G072900.7 | TRIUR3_09946 | Seita.5G323600.1 | Sobic.003G301000.1 | Zm00001d012710_T001 |
| orthologID_006978 | PH01000587G0940 | LOC_Os01g65400.2 | Bradi2g56537.1 | HORVU3Hr1G082130.10 | TRIUR3_20694 | Seita.5G398500.1 | Sobic.003G373300.1 | Zm00001d042663_T001 |
| orthologID_006318 | PH01004403G0090 | LOC_Os01g61720.8 | Bradi2g54170.8 | HORVU3Hr1G087500.2 | TRIUR3_05501 | Seita.5G371000.1 | Sobic.003G346000.5 | Zm00001d042973_T003 |
| orthologID_006889 | PH01004823G0140 | LOC_Os01g45900.1 | Bradi2g45130.1 | HORVU3Hr1G058150.3 | TRIUR3_19556 | Seita.5G243500.1 | Sobic.003G237300.1 | Zm00001d044061_T001 |
| orthologID_006878 | PH01001571G0100 | LOC_Os01g09990.1 | Bradi2g05950.2 | HORVU3Hr1G018980.6 | TRIUR3_35423 | Seita.5G138900.1 | Sobic.003G031100.1 | Zm00001d008853_T001 |
| orthologID_006250 | PH01000076G0050 | LOC_Os01g52750.1 | Bradi2g48740.1 | HORVU3Hr1G069030.7 | TRIUR3_08947 | Seita.5G303900.1 | Sobic.003G283100.1 | Zm00001d043581_T001 |
| orthologID_006390 | PH01000133G1110 | LOC_Os01g14820.1 | Bradi2g08832.1 | HORVU3Hr1G034170.6 | TRIUR3_19513 | Seita.5G061500.1 | Sobic.003G113300.1 | Zm00001d040259_T001 |
| orthologID_006759 | PH01002011G0130 | LOC_Os01g51754.1 | Bradi2g48150.1 | HORVU3Hr1G067620.1 | TRIUR3_11156 | Seita.5G295100.1 | Sobic.003G276400.1 | Zm00001d043662_T001 |
| orthologID_006483 | PH01002648G0250 | LOC_Os07g16950.4 | Bradi2g14350.1 | HORVU0Hr1G022580.4 | TRIUR3_33591 | Seita.1G148400.1 | Sobic.003G407600.1 | Zm00001d012003_T001 |
| orthologID_006147 | PH01000002G4080 | LOC_Os01g70710.1 | Bradi2g59810.1 | HORVU3Hr1G097770.1 | TRIUR3_32515 | Seita.5G441700.1 | Sobic.003G415900.1 | Zm00001d042225_T002 |
| orthologID_006663 | PH01000099G1290 | LOC_Os01g56180.1 | Bradi2g51010.1 | HORVU3Hr1G074680.1 | TRIUR3_20746 | Seita.5G331400.1 | Sobic.003G308400.1 | Zm00001d012662_T001 |
| orthologID_006171 | PH01000072G0300 | LOC_Os01g56400.1 | Bradi2g51200.1 | HORVU3Hr1G075100.1 | TRIUR3_20947 | Seita.5G332900.1 | Sobic.003G310100.1 | Zm00001d012645_T001 |
| orthologID_006516 | PH01000263G0390 | LOC_Os01g55310.1 | Bradi2g50350.1 | HORVU3Hr1G072950.3 | TRIUR3_32246 | Seita.5G324200.1 | Sobic.003G301500.1 | Zm00001d012707_T003 |
| orthologID_006959 | PH01001231G0470 | LOC_Os01g11570.1 | Bradi2g06900.1 | HORVU3Hr1G028210.3 | TRIUR3_26412 | Seita.5G149300.1 | Sobic.003G018800.1 | Zm00001d039370_T002 |
| orthologID_006550 | PH01000263G0270 | LOC_Os01g55280.1 | Bradi2g50317.1 | HORVU3Hr1G072880.3 | TRIUR3_09945 | Seita.5G323800.1 | Sobic.003G301200.1 | Zm00001d043406_T001 |
| orthologID_006704 | PH01000525G0570 | LOC_Os01g03650.1 | Bradi2g01870.1 | HORVU3Hr1G004330.1 | TRIUR3_30071 | Seita.5G089600.1 | Sobic.003G085000.1 | Zm00001d008549_T002 |
| orthologID_006572 | PH01000277G0630 | LOC_Os01g54910.1 | Bradi2g50050.3 | HORVU3Hr1G072230.3 | TRIUR3_24687 | Seita.5G320600.1 | Sobic.003G297800.1 | Zm00001d043494_T005 |
| orthologID_006398 | PH01000071G1820 | LOC_Os01g66720.1 | Bradi2g57440.2 | HORVU3Hr1G090050.25 | TRIUR3_06202 | Seita.5G409600.1 | Sobic.003G384300.1 | Zm00001d042555_T001 |
| orthologID_006637 | PH01000592G0380 | LOC_Os01g08420.1 | Bradi2g05017.1 | HORVU3Hr1G022000.3 | TRIUR3_23256 | Seita.5G127300.1 | Sobic.003G043200.1 | Zm00001d039612_T001 |
| orthologID_006301 | PH01000317G0080 | LOC_Os01g66030.1 | Bradi2g57000.1 | HORVU3Hr1G091000.8 | TRIUR3_15782 | Seita.5G404600.1 | Sobic.003G379000.1 | Zm00001d042618_T001 |
| orthologID_006898 | PH01000812G0610 | LOC_Os01g12840.1 | Bradi2g07660.1 | HORVU3Hr1G031220.4 | TRIUR3_28161 | Seita.5G009000.1 | Sobic.003G008800.2 | Zm00001d008993_T001 |
| orthologID_006236 | PH01000336G0700 | LOC_Os01g55560.1 | Bradi2g50520.2 | HORVU3Hr1G073240.1 | TRIUR3_35335 | Seita.5G326200.1 | Sobic.003G303600.1 | Zm00001d012690_T002 |
| orthologID_007539 | PH01000235G0760 | LOC_Os08g06474.1 | Bradi4g41070.1 | HORVU6Hr1G091140.1 | TRIUR3_01944 | Seita.5G414400.1 | Sobic.003G388700.1 | Zm00001d042510_T002 |
| orthologID_006229 | PH01000018G2540 | LOC_Os01g55520.1 | Bradi2g50490.2 | HORVU3Hr1G073200.7 | TRIUR3_30594 | Seita.5G325900.1 | Sobic.003G303200.1 | Zm00001d043383_T001 |
| orthologID_006161 | PH01000443G0530 | LOC_Os01g12460.1 | Bradi2g07370.1 | HORVU3Hr1G030360.1 | TRIUR3_17524 | Seita.5G015600.1 | Sobic.003G012600.2 | Zm00001d008970_T001 |
| orthologID_006808 | PH01000166G0280 | LOC_Os01g36080.1 | Bradi2g40550.2 | HORVU3Hr1G047160.17 | TRIUR3_22959 | Seita.5G191000.1 | Sobic.003G182600.1 | Zm00001d009364_T001 |
| orthologID_006470 | PH01000309G0680 | LOC_Os01g64440.1 | Bradi2g55886.1 | HORVU3Hr1G083650.1 | TRIUR3_27866 | Seita.5G391800.1 | Sobic.003G366100.2 | Zm00001d042751_T001 |
| orthologID_006765 | PH01000281G0780 | LOC_Os01g04720.1 | Bradi2g02700.1 | HORVU3Hr1G010070.2 | TRIUR3_27817 | Seita.5G096500.1 | Sobic.003G078300.1 | Zm00001d039911_T002 |
| orthologID_006209 | PH01000058G1810 | LOC_Os01g72880.1 | Bradi2g61370.2 | HORVU3Hr1G110480.5 | TRIUR3_24234 | Seita.5G460100.1 | Sobic.003G433300.1 | Zm00001d011829_T001 |
| orthologID_006347 | PH01000260G1250 | LOC_Os01g02050.1 | Bradi2g00910.1 | HORVU3Hr1G000940.3 | TRIUR3_03888 | Seita.5G074200.1 | Sobic.003G100600.1 | Zm00001d040129_T001 |
| orthologID_006273 | PH01002733G0260 | LOC_Os01g21160.1 | Bradi2g11900.1 | HORVU2Hr1G012440.2 | TRIUR3_22686 | Seita.5G021800.1 | Sobic.003G148800.1 | Zm00001d040670_T004 |
| orthologID_006178 | PH01000207G0760 | LOC_Os01g22550.1 | Bradi2g12320.1 | HORVU1Hr1G013730.2 | TRIUR3_04434 | Seita.5G156400.1 | Sobic.003G153000.1 | Zm00001d040724_T001 |
| orthologID_006684 | PH01001186G0530 | LOC_Os01g06500.1 | Bradi2g00232.1 | HORVU3Hr1G025670.4 | TRIUR3_18044 | Seita.5G112200.1 | Sobic.003G062500.1 | Zm00001d008646_T001 |
| orthologID_006158 | PH01000121G0050 | LOC_Os01g43220.1 | Bradi2g43770.1 | HORVU3Hr1G055200.3 | TRIUR3_10841 | Seita.5G229700.1 | Sobic.003G223300.1 | Zm00001d011370_T001 |
| orthologID_006711 | PH01000622G0600 | LOC_Os01g73990.1 | Bradi2g62010.1 | HORVU3Hr1G114360.3 | TRIUR3_04685 | Seita.5G467800.1 | Sobic.003G443300.1 | Zm00001d041963_T001 |
| orthologID_006892 | PH01001108G0520 | LOC_Os01g47690.1 | Bradi2g46040.1 | HORVU3Hr1G060920.3 | TRIUR3_22119 | Seita.5G262900.1 | Sobic.003G249900.1 | Zm00001d043941_T001 |
| orthologID_007009 | PH01000031G0320 | LOC_Os01g67520.1 | Bradi2g57837.2 | HORVU3Hr1G089030.13 | TRIUR3_21856 | Seita.5G416800.1 | Sobic.003G391500.1 | Zm00001d042478_T003 |
| orthologID_006242 | PH01000207G1290 | LOC_Os01g22660.1 | Bradi2g12340.1 | HORVU1Hr1G010670.3 | TRIUR3_30685 | Seita.5G157000.1 | Sobic.003G153500.1 | Zm00001d040731_T004 |
| orthologID_008747 | PH01000012G0930 | LOC_Os02g20280.1 | Bradi5g22027.1 | HORVU2Hr1G103800.3 | TRIUR3_16033 | Seita.5G330900.1 | Sobic.003G308000.1 | Zm00001d043337_T001 |
| orthologID_006267 | PH01002649G0270 | LOC_Os01g09280.1 | Bradi2g05590.1 | HORVU3Hr1G019840.4 | TRIUR3_12855 | Seita.5G133800.1 | Sobic.003G036600.1 | Zm00001d008198_T001 |
| orthologID_006456 | PH01001169G0230 | LOC_Os01g07960.3 | Bradi2g04645.1 | HORVU3Hr1G023960.1 | TRIUR3_19634 | Seita.5G123600.1 | Sobic.003G048500.2 | Zm00001d039653_T005 |
| orthologID_006517 | PH01000002G3460 | LOC_Os01g70430.1 | Bradi2g59570.5 | HORVU3Hr1G096690.8 | TRIUR3_00309 | Seita.5G439100.1 | Sobic.003G413100.1 | Zm00001d042250_T001 |
| orthologID_006498 | PH01001385G0140 | LOC_Os01g15780.1 | Bradi2g09620.1 | HORVU3Hr1G036630.6 | TRIUR3_13738 | Seita.5G053500.1 | Sobic.003G120700.2 | Zm00001d040371_T002 |

| | | | | | | | | |
|---|---|---|---|---|---|---|---|---|
| orthologID_006688 | PH01000525G0560 | LOC_Os01g03660.2 | Bradi2g01890.1 | HORVU3Hr1G004320.3 | TRIUR3_20384 | Seita.5G089500.1 | Sobic.003G085100.1 | Zm00001d039987_T003 |
| orthologID_006628 | PH01000423G0190 | LOC_Os01g55820.1 | Bradi2g50750.1 | HORVU3Hr1G074010.3 | TRIUR3_16147 | Seita.5G328200.1 | Sobic.003G305601.1 | Zm00001d043361_T001 |
| orthologID_006243 | PH01000032G2590 | LOC_Os01g58680.1 | Bradi2g52540.3 | HORVU3Hr1G078470.2 | TRIUR3_16333 | Seita.5G350300.1 | Sobic.003G326500.1 | Zm00001d012571_T002 |
| orthologID_006294 | PH01000277G1080 | LOC_Os01g55050.1 | Bradi2g50145.1 | HORVU3Hr1G072520.1 | TRIUR3_23019 | Seita.5G321900.1 | Sobic.003G299000.1 | Zm00001d012728_T001 |
| orthologID_006691 | PH01000778G0770 | LOC_Os01g48420.1 | Bradi2g46380.2 | HORVU3Hr1G062040.1 | TRIUR3_13961 | Seita.5G268100.1 | Sobic.003G254300.1 | Zm00001d011581_T001 |
| orthologID_006215 | PH01000260G1020 | LOC_Os01g01840.1 | Bradi2g00720.1 | HORVU3Hr1G000280.1 | TRIUR3_27194 | Seita.5G072700.1 | Sobic.003G102300.1 | Zm00001d040152_T001 |
| orthologID_006255 | PH01003318G0230 | LOC_Os01g74190.1 | Bradi2g62280.1 | HORVU3Hr1G115840.1 | TRIUR3_23810 | Seita.5G469100.1 | Sobic.003G444500.1 | Zm00001d011780_T002 |
| orthologID_006299 | PH01000697G0480 | LOC_Os01g03500.1 | Bradi2g01810.1 | HORVU3Hr1G004440.11 | TRIUR3_22735 | Seita.5G085200.1 | Sobic.003G089200.3 | Zm00001d008516_T001 |
| orthologID_006583 | PH01002011G0020 | LOC_Os01g51790.1 | Bradi2g48180.1 | HORVU3Hr1G067650.1 | TRIUR3_22087 | Seita.5G295500.1 | Sobic.003G276800.1 | Zm00001d043656_T001 |
| orthologID_006141 | PH01000018G0640 | LOC_Os01g54784.1 | Bradi2g49980.1 | HORVU3Hr1G071910.8 | TRIUR3_34462 | Seita.5G319700.1 | Sobic.003G296900.1 | Zm00001d043483_T001 |
| orthologID_006748 | PH01000746G0590 | LOC_Os01g54670.1 | Bradi2g49930.1 | HORVU3Hr1G071800.7 | TRIUR3_30088 | Seita.5G319300.1 | Sobic.003G296600.1 | Zm00001d043479_T001 |
| orthologID_007632 | PH01000737G0320 | LOC_Os01g35990.1 | Bradi1g07710.1 | HORVU7Hr1G010600.19 | TRIUR3_06099 | Seita.5G188200.1 | Sobic.003G178900.1 | Zm00001d044578_T001 |
| orthologID_006681 | PH01000425G0420 | LOC_Os01g57710.1 | Bradi2g52060.3 | HORVU3Hr1G077530.9 | TRIUR3_18111 | Seita.5G343800.1 | Sobic.003G320600.1 | Zm00001d043240_T002 |
| orthologID_006924 | PH01001615G0050 | LOC_Os01g37770.1 | Bradi2g41140.3 | HORVU3Hr1G048890.1 | TRIUR3_23714 | Seita.5G198800.1 | Sobic.003G188500.1 | Zm00001d011159_T001 |
| orthologID_006534 | PH01001578G0280 | LOC_Os01g18280.1 | Bradi2g11090.1 | HORVU3Hr1G030490.1 | TRIUR3_34421 | Seita.5G034100.1 | Sobic.003G136800.1 | Zm00001d040535_T007 |
| orthologID_006412 | PH01001186G0560 | LOC_Os01g06470.1 | Bradi2g00320.1 | HORVU3Hr1G025640.8 | TRIUR3_09566 | Seita.5G111900.1 | Sobic.003G063000.1 | Zm00001d008640_T002 |
| orthologID_006200 | PH01000277G0730 | LOC_Os01g54940.1 | Bradi2g50080.1 | HORVU3Hr1G072320.2 | TRIUR3_09364 | Seita.5G320900.1 | Sobic.003G298200.1 | Zm00001d043432_T001 |
| orthologID_008648 | PH01000465G0960 | LOC_Os03g18620.1 | Bradi1g65120.1 | HORVU4Hr1G055160.2 | TRIUR3_15623 | Seita.9G552500.1 | Sobic.003G388150.1 | Zm00001d042519_T002 |
| orthologID_006624 | PH01000182G1310 | LOC_Os01g46720.1 | Bradi2g45450.1 | HORVU3Hr1G059130.8 | TRIUR3_25852 | Seita.5G248500.1 | Sobic.003G241900.1 | Zm00001d044017_T001 |
| orthologID_006532 | PH01000032G2690 | LOC_Os01g58730.1 | Bradi2g52566.1 | HORVU3Hr1G078520.3 | TRIUR3_34271 | Seita.5G350500.1 | Sobic.003G326700.1 | Zm00001d012568_T001 |
| orthologID_006955 | PH01000018G1200 | LOC_Os01g55070.1 | Bradi2g50160.1 | HORVU3Hr1G072560.2 | TRIUR3_14482 | Seita.5G322100.1 | Sobic.003G299200.1 | Zm00001d043426_T001 |
| orthologID_006942 | PH01000094G0860 | LOC_Os01g16640.1 | Bradi2g10230.1 | HORVU3Hr1G038460.1 | TRIUR3_18268 | Seita.5G153100.1 | Sobic.003G127700.1 | Zm00001d008259_T001 |
| orthologID_006360 | PH01000002G1040 | LOC_Os01g69140.1 | Bradi2g58987.1 | HORVU3Hr1G094250.8 | TRIUR3_04729 | Seita.5G430300.1 | Sobic.003G404600.1 | Zm00001d042337_T001 |
| orthologID_006419 | PH01000357G0190 | LOC_Os01g07890.1 | Bradi2g04597.1 | HORVU3Hr1G024200.1 | TRIUR3_07159 | Seita.5G123000.1 | Sobic.003G049300.1 | Zm00001d039657_T001 |
| orthologID_006830 | PH01004111G0150 | LOC_Os01g36630.1 | Bradi2g40677.2 | HORVU3Hr1G049210.1 | TRIUR3_03449 | Seita.5G193900.1 | Sobic.003G184400.1 | Zm00001d009396_T002 |
| orthologID_006948 | PH01000121G0040 | LOC_Os01g43170.1 | Bradi2g43760.1 | HORVU3Hr1G055000.1 | TRIUR3_10840 | Seita.5G229600.1 | Sobic.003G223200.1 | Zm00001d044195_T018 |
| orthologID_008914 | PH01007367G0020 | LOC_Os09g09320.1 | Bradi5g02510.1 | HORVU2Hr1G007080.13 | TRIUR3_11930 | Seita.5G063900.1 | Sobic.003G110700.2 | Zm00001d040242_T001 |
| orthologID_006459 | PH01002160G0050 | LOC_Os01g70230.1 | Bradi2g59440.2 | HORVU3Hr1G096240.1 | TRIUR3_02326 | Seita.5G437700.1 | Sobic.003G411500.2 | Zm00001d042271_T001 |
| orthologID_008698 | PH01000040G1390 | LOC_Os03g61640.1 | Bradi1g02950.1 | HORVU2Hr1G109460.2 | TRIUR3_05675 | Seita.3G205100.1 | Sobic.003G037900.1 | Zm00001d039541_T002 |
| orthologID_006342 | PH01001932G0030 | LOC_Os01g12480.1 | Bradi2g07400.1 | HORVU3Hr1G030330.3 | TRIUR3_17522 | Seita.5G015300.1 | Sobic.003G012500.1 | Zm00001d052074_T001 |
| orthologID_006603 | PH01001004G0410 | LOC_Os01g40420.1 | Bradi2g42080.1 | HORVU2Hr1G032570.2 | TRIUR3_34291 | Seita.5G255500.1 | Sobic.003G200600.1 | Zm00001d044315_T002 |
| orthologID_006724 | PH01000252G0880 | LOC_Os01g54040.2 | Bradi2g49550.6 | HORVU3Hr1G070860.1 | TRIUR3_09680 | Seita.5G314400.1 | Sobic.003G292500.1 | Zm00001d043506_T003 |
| orthologID_006187 | PH01001210G0050 | LOC_Os01g51860.1 | Bradi2g48230.1 | HORVU3Hr1G067770.1 | TRIUR3_08389 | Seita.5G296000.1 | Sobic.003G277400.1 | Zm00001d011720_T002 |
| orthologID_006721 | PH01003928G0040 | LOC_Os01g07100.1 | Bradi2g03980.1 | HORVU3Hr1G017970.1 | TRIUR3_03987 | Seita.5G114800.1 | Sobic.003G058500.1 | Zm00001d039740_T001 |
| orthologID_006304 | PH01000109G0730 | LOC_Os01g56940.1 | Bradi2g51637.1 | HORVU3Hr1G076270.15 | TRIUR3_20621 | Seita.5G338600.1 | Sobic.003G314900.1 | Zm00001d043287_T002 |
| orthologID_006875 | PH01004326G0140 | LOC_Os01g72490.1 | Bradi2g61120.1 | HORVU3Hr1G108950.2 | TRIUR3_31630 | Seita.5G456500.1 | Sobic.003G429800.1 | Zm00001d011843_T001 |
| orthologID_006555 | PH01005426G0050 | LOC_Os01g19490.1 | Bradi2g11510.2 | HORVU2Hr1G120330.3 | TRIUR3_01095 | Seita.5G027300.1 | Sobic.003G144000.1 | Zm00001d009122_T001 |
| orthologID_006476 | PH01000291G0340 | LOC_Os01g14690.1 | Bradi2g08560.1 | HORVU3Hr1G033520.1 | TRIUR3_28185 | Seita.5G001400.1 | Sobic.003G001100.1 | Zm00001d039252_T001 |
| orthologID_006755 | PH01001261G0410 | LOC_Os01g41990.1 | Bradi2g42920.1 | HORVU3Hr1G052770.1 | TRIUR3_31124 | Seita.5G217800.1 | Sobic.003G212000.1 | Zm00001d044411_T001 |
| orthologID_006510 | PH01000203G0700 | LOC_Os11g13420.1 | Bradi2g06000.1 | HORVU3Hr1G018840.6 | TRIUR3_13250 | Seita.5G139500.1 | Sobic.003G030400.1 | Zm00001d039451_T005 |
| orthologID_006619 | PH01000121G0740 | LOC_Os01g43420.1 | Bradi2g43930.1 | HORVU3Hr1G055780.2 | TRIUR3_24588 | Seita.5G231600.1 | Sobic.003G225300.1 | Zm00001d044184_T001 |
| orthologID_006168 | PH01001378G0110 | LOC_Os01g52100.1 | Bradi2g48330.1 | HORVU3Hr1G068120.7 | TRIUR3_35140 | Seita.5G298300.1 | Sobic.003G278400.1 | Zm00001d043612_T001 |
| orthologID_006610 | PH01000425G0570 | LOC_Os01g57770.1 | Bradi2g52110.1 | HORVU3Hr1G077610.1 | TRIUR3_27782 | Seita.5G344200.1 | Sobic.003G321000.1 | Zm00001d043235_T001 |
| orthologID_008556 | PH01000363G1010 | LOC_Os07g38100.1 | Bradi1g24260.1 | HORVU2Hr1G058860.1 | TRIUR3_18518 | Seita.2G357600.1 | Sobic.003G350600.1 | Zm00001d042923_T003 |
| orthologID_006655 | PH01002896G0100 | LOC_Os01g48690.1 | Bradi2g46520.1 | HORVU3Hr1G062490.1 | TRIUR3_22278 | Seita.5G270200.1 | Sobic.003G256200.1 | Zm00001d011596_T001 |
| orthologID_006248 | PH01001108G0340 | LOC_Os01g47620.1 | Bradi2g45961.1 | HORVU3Hr1G060640.5 | TRIUR3_20530 | Seita.5G262200.1 | Sobic.003G249300.1 | Zm00001d043945_T001 |
| orthologID_006930 | PH01001170G0180 | LOC_Os01g22510.1 | Bradi2g12300.1 | HORVU3Hr1G062700.1 | TRIUR3_29844 | Seita.5G156200.1 | Sobic.003G152800.1 | Zm00001d040720_T001 |
| orthologID_006633 | PH01002772G0130 | LOC_Os05g28300.1 | Bradi2g28900.2 | HORVU1Hr1G060010.4 | TRIUR3_19453 | Seita.3G278300.1 | Sobic.003G422500.1 | Zm00001d042136_T005 |
| orthologID_006823 | PH01001722G0280 | LOC_Os01g08680.2 | Bradi2g05218.5 | HORVU3Hr1G021310.1 | TRIUR3_34375 | Seita.5G129300.1 | Sobic.003G041100.1 | Zm00001d039590_T002 |
| orthologID_006858 | PH01001236G0280 | LOC_Os01g70950.1 | Bradi2g60097.1 | HORVU3Hr1G098660.3 | TRIUR3_01253 | Seita.5G444200.1 | Sobic.003G418200.1 | Zm00001d042191_T002 |

| | | | | | | | | |
|---|---|---|---|---|---|---|---|---|
| orthologID_006632 | PH01002355G0120 | LOC_Os01g72100.1 | Bradi2g60660.1 | HORVU3Hr1G107090.2 | TRIUR3_31196 | Seita.5G451500.1 | Sobic.003G425500.1 | Zm00001d042107_T001 |
| orthologID_006794 | PH01000214G0600 | LOC_Os01g53610.1 | Bradi2g49207.1 | HORVU3Hr1G070040.3 | TRIUR3_24875 | Seita.5G310400.1 | Sobic.003G289500.1 | Zm00001d043556_T001 |
| orthologID_006661 | PH01000320G0990 | LOC_Os01g03740.1 | Bradi2g01967.1 | HORVU3Hr1G003860.1 | TRIUR3_13240 | Seita.5G087300.1 | Sobic.003G087300.1 | Zm00001d040014_T002 |
| orthologID_006554 | PH01000031G0250 | LOC_Os01g67480.1 | Bradi2g57800.4 | HORVU3Hr1G089090.2 | TRIUR3_33816 | Seita.5G416200.1 | Sobic.003G391100.1 | Zm00001d042482_T002 |
| orthologID_006838 | PH01000240G0480 | LOC_Os01g48810.1 | Bradi2g46610.1 | HORVU3Hr1G062710.1 | TRIUR3_27555 | Seita.5G271300.1 | Sobic.003G257600.1 | Zm00001d011605_T001 |
| orthologID_006145 | PH01000502G0910 | LOC_Os01g48140.1 | Bradi2g46220.1 | HORVU3Hr1G061620.1 | TRIUR3_20371 | Seita.5G265700.1 | Sobic.003G252000.1 | Zm00001d011559_T001 |
| orthologID_006134 | PH01000398G0430 | LOC_Os01g56150.1 | Bradi2g51000.2 | HORVU3Hr1G074620.2 | TRIUR3_16432 | Seita.5G331300.1 | Sobic.003G308300.1 | Zm00001d012667_T009 |
| orthologID_006848 | PH01002623G0080 | LOC_Os01g01720.1 | Bradi2g00660.1 | HORVU3Hr1G005640.6 | TRIUR3_16081 | Seita.5G072000.1 | Sobic.003G103200.1 | Zm00001d040161_T002 |
| orthologID_006676 | PH01004403G0020 | LOC_Os01g61670.1 | Bradi2g54127.1 | HORVU3Hr1G087450.3 | TRIUR3_08708 | Seita.5G370000.1 | Sobic.003G345500.1 | Zm00001d042976_T001 |
| orthologID_009948 | PH01001615G0350 | LOC_Os01g37630.1 | Bradi2g41070.1 | HORVU2Hr1G105690.1 | TRIUR3_07262 | Seita.5G194600.1 | Sobic.003G184900.1 | Zm00001d009398_T003 |
| orthologID_006949 | PH01000704G0200 | LOC_Os01g49250.1 | Bradi2g46830.2 | HORVU3Hr1G063690.10 | TRIUR3_09273 | Seita.5G274900.1 | Sobic.003G260900.1 | Zm00001d043827_T001 |
| orthologID_006221 | PH01000812G0370 | LOC_Os01g12930.1 | Bradi2g07750.2 | HORVU3Hr1G052520.1 | TRIUR3_10633 | Seita.5G008400.1 | Sobic.003G008100.1 | Zm00001d039302_T001 |
| orthologID_006806 | PH01002203G0260 | LOC_Os01g16290.1 | Bradi2g10040.1 | HORVU3Hr1G038050.2 | TRIUR3_25046 | Seita.5G048400.1 | Sobic.003G125900.1 | Zm00001d040416_T004 |
| orthologID_006568 | PH01000155G0520 | LOC_Os01g55740.1 | Bradi2g50680.3 | HORVU3Hr1G073790.2 | TRIUR3_11555 | Seita.5G327500.1 | Sobic.003G304800.1 | Zm00001d012681_T003 |
| orthologID_006996 | PH01000270G1150 | LOC_Os01g04200.1 | Bradi2g02250.1 | HORVU3Hr1G007230.1 | TRIUR3_03392 | Seita.5G091000.1 | Sobic.003G083900.1 | Zm00001d039971_T002 |
| orthologID_006621 | PH01000746G0310 | LOC_Os01g54530.1 | Bradi2g49840.2 | HORVU3Hr1G071540.4 | TRIUR3_30509 | Seita.5G318100.1 | Sobic.003G295700.1 | Zm00001d043465_T002 |
| orthologID_006208 | PH01000009G0870 | LOC_Os01g60020.1 | Bradi2g53260.1 | HORVU3Hr1G080100.7 | TRIUR3_04332 | Seita.5G358900.1 | Sobic.003G334600.1 | Zm00001d012527_T001 |
| orthologID_006156 | PH01000270G0210 | LOC_Os01g43250.1 | Bradi2g43780.1 | HORVU3Hr1G055410.2 | TRIUR3_10843 | Seita.5G229800.1 | Sobic.003G223500.1 | Zm00001d044193_T002 |
| orthologID_006933 | PH01001892G0060 | LOC_Os01g10350.1 | Bradi2g06190.1 | HORVU3Hr1G026980.2 | TRIUR3_11638 | Seita.5G141500.1 | Sobic.003G028100.1 | Zm00001d008871_T001 |
| orthologID_006288 | PH01000454G0450 | LOC_Os01g27210.1 | Bradi2g13020.1 | HORVU5Hr1G118800.4 | TRIUR3_34711 | Seita.5G168800.1 | Sobic.003G164800.1 | Zm00001d043344_T001 |
| orthologID_006670 | PH01000568G0340 | LOC_Os01g47040.1 | Bradi2g45590.1 | HORVU3Hr1G059440.4 | TRIUR3_30315 | Seita.9G508500.1 | Sobic.003G244300.1 | Zm00001d043990_T001 |
| orthologID_006597 | PH01000002G0240 | LOC_Os01g68890.1 | Bradi2g58790.1 | HORVU3Hr1G093570.4 | TRIUR3_29728 | Seita.5G428200.1 | Sobic.003G402300.1 | Zm00001d012047_T001 |
| orthologID_006285 | PH01000781G0250 | LOC_Os01g73234.1 | Bradi2g61580.1 | HORVU5Hr1G116290.1 | TRIUR3_23193 | Seita.5G463500.1 | Sobic.003G437600.2 | Zm00001d042017_T002 |
| orthologID_006307 | PH01000333G1120 | LOC_Os01g43060.1 | Bradi2g43670.1 | HORVU3Hr1G054630.1 | TRIUR3_28201 | Seita.5G228100.1 | Sobic.003G221600.1 | Zm00001d044219_T004 |
| orthologID_006220 | PH01000817G0610 | LOC_Os01g49280.1 | Bradi2g46835.3 | HORVU3Hr1G063680.11 | TRIUR3_09274 | Seita.5G275200.1 | Sobic.003G261000.3 | Zm00001d011616_T001 |
| orthologID_007012 | PH01008877G0020 | LOC_Os01g36600.1 | Bradi2g40660.1 | HORVU3Hr1G049360.1 | TRIUR3_16748 | Seita.5G193200.1 | Sobic.003G184200.1 | Zm00001d009387_T001 |
| orthologID_006753 | PH01000881G0350 | LOC_Os01g36870.1 | Bradi2g40770.1 | HORVU7Hr1G022270.11 | TRIUR3_31220 | Seita.5G197800.1 | Sobic.003G187200.1 | Zm00001d044511_T001 |
| orthologID_008632 | PH01003380G0040 | LOC_Os07g09720.1 | Bradi1g54610.1 | HORVU7Hr1G039880.3 | TRIUR3_20006 | Seita.5G152200.1 | Sobic.003G017400.1 | Zm00001d039361_T001 |
| orthologID_006144 | PH01000004G2230 | LOC_Os01g64010.2 | Bradi2g55560.1 | HORVU3Hr1G084350.4 | TRIUR3_12644 | Seita.5G389400.1 | Sobic.003G363500.1 | Zm00001d042778_T002 |
| orthologID_006797 | PH01000592G0690 | LOC_Os01g08280.1 | Bradi2g04870.1 | HORVU3Hr1G022600.6 | TRIUR3_08265 | Seita.5G125900.1 | Sobic.003G045200.1 | Zm00001d008748_T001 |
| orthologID_006311 | PH01000417G0840 | LOC_Os01g01380.1 | Bradi2g03497.2 | HORVU3Hr1G014240.28 | TRIUR3_06044 | Seita.5G068700.1 | Sobic.003G106200.2 | Zm00001d040193_T005 |
| orthologID_006558 | PH01003102G0110 | LOC_Os01g13930.1 | Bradi2g08370.1 | HORVU3Hr1G032760.4 | TRIUR3_09135 | Seita.5G002600.1 | Sobic.003G001500.1 | Zm00001d039256_T001 |
| orthologID_006993 | PH01000004G1750 | LOC_Os01g64250.1 | Bradi2g55720.1 | HORVU3Hr1G084020.3 | TRIUR3_22778 | Seita.5G390600.1 | Sobic.003G364900.1 | Zm00001d042764_T001 |
| orthologID_006210 | PH01002623G0240 | LOC_Os01g01600.1 | Bradi2g03427.1 | HORVU3Hr1G014060.13 | TRIUR3_23381 | Seita.5G069700.1 | Sobic.003G105200.1 | Zm00001d040186_T001 |
| orthologID_006662 | PH01000058G0460 | LOC_Os01g72350.1 | Bradi2g61017.2 | HORVU3Hr1G108620.3 | TRIUR3_22215 | Seita.5G455500.1 | Sobic.003G428600.1 | Zm00001d042064_T010 |
| orthologID_006254 | PH01000972G0320 | LOC_Os01g45370.1 | Bradi2g44900.1 | HORVU3Hr1G057020.4 | TRIUR3_18998 | Seita.5G240500.1 | Sobic.003G234900.1 | Zm00001d011457_T001 |
| orthologID_006700 | PH01000489G0090 | LOC_Os01g68930.1 | Bradi2g58820.1 | HORVU3Hr1G093590.18 | TRIUR3_13825 | Seita.5G428400.1 | Sobic.003G402500.1 | Zm00001d042357_T002 |
| orthologID_006431 | PH01000231G0020 | LOC_Os01g42280.1 | Bradi2g43100.3 | HORVU3Hr1G053290.1 | TRIUR3_34360 | Seita.5G220400.1 | Sobic.003G214800.1 | Zm00001d011311_T001 |
| orthologID_006211 | PH01000993G0650 | LOC_Os01g61630.1 | Bradi2g54110.1 | HORVU3Hr1G087540.1 | TRIUR3_02742 | Seita.5G369700.1 | Sobic.003G345300.1 | Zm00001d042978_T001 |
| orthologID_006664 | PH01002854G0220 | LOC_Os01g13210.1 | Bradi2g07940.1 | HORVU3Hr1G031850.2 | TRIUR3_26766 | Seita.5G005900.1 | Sobic.003G005800.1 | Zm00001d039283_T004 |
| orthologID_006943 | PH01000684G0180 | LOC_Os01g40690.1 | Bradi2g42270.1 | HORVU3Hr1G051220.3 | TRIUR3_28509 | Seita.5G209500.1 | Sobic.003G202800.1 | Zm00001d044332_T001 |
| orthologID_006795 | PH01000076G0220 | LOC_Os01g52720.1 | Bradi2g48720.1 | HORVU3Hr1G069000.1 | TRIUR3_08945 | Seita.5G303600.1 | Sobic.003G051200.1 | Zm00001d011750_T004 |
| orthologID_006798 | PH01000500G0770 | LOC_Os01g50320.1 | Bradi2g47450.1 | HORVU3Hr1G065510.9 | TRIUR3_08166 | Seita.5G284100.1 | Sobic.003G268300.1 | Zm00001d011652_T001 |
| orthologID_006658 | PH01000207G0700 | LOC_Os01g22500.1 | Bradi2g12290.1 | HORVU7Hr1G027240.3 | TRIUR3_21936 | Seita.5G156100.1 | Sobic.003G152700.1 | Zm00001d040717_T002 |
| orthologID_006355 | PH01000240G1100 | LOC_Os01g48570.1 | Bradi2g46490.1 | HORVU3Hr1G062350.3 | TRIUR3_28670 | Seita.5G269400.1 | Sobic.003G255600.1 | Zm00001d043872_T001 |
| orthologID_006780 | PH01000144G1410 | LOC_Os01g73560.1 | Bradi2g61810.1 | HORVU3Hr1G113080.1 | TRIUR3_02066 | Seita.5G465400.1 | Sobic.003G440700.1 | Zm00001d041994_T002 |
| orthologID_006973 | PH01000632G0690 | LOC_Os01g15039.1 | Bradi2g09090.1 | HORVU3Hr1G035210.3 | TRIUR3_28927 | Seita.5G058300.1 | Sobic.003G116100.1 | Zm00001d008334_T001 |
| orthologID_006523 | PH01001739G0230 | LOC_Os01g71990.1 | Bradi2g60730.1 | HORVU3Hr1G106720.2 | TRIUR3_30591 | Seita.5G450800.1 | Sobic.003G424300.1 | Zm00001d042117_T002 |
| orthologID_006472 | PH01000155G0470 | LOC_Os01g55720.1 | Bradi2g50650.1 | HORVU3Hr1G073770.2 | TRIUR3_11552 | Seita.5G327300.1 | Sobic.003G304500.1 | Zm00001d043371_T001 |
| orthologID_006446 | PH01000284G0930 | LOC_Os01g54340.1 | Bradi2g49690.1 | HORVU3Hr1G071220.1 | TRIUR3_21627 | Seita.5G316100.1 | Sobic.003G293900.1 | Zm00001d043444_T001 |

| | | | | | | | | |
|---|---|---|---|---|---|---|---|---|
| orthologID_006828 | PH01000098G1600 | LOC_Os01g07420.1 | Bradi2g04230.1 | HORVU3Hr1G017100.1 | TRIUR3_25286 | Seita.5G118800.1 | Sobic.003G053500.1 | Zm00001d008693_T002 |
| orthologID_006911 | PH01000235G0500 | LOC_Os01g66970.1 | Bradi2g57520.3 | HORVU3Hr1G089990.6 | TRIUR3_20055 | Seita.5G412000.1 | Sobic.003G386200.1 | Zm00001d042534_T002 |
| orthologID_006730 | PH01000284G0970 | LOC_Os01g54380.1 | Bradi2g49720.1 | HORVU3Hr1G071290.3 | TRIUR3_17595 | Seita.5G316500.1 | Sobic.003G294200.2 | Zm00001d043446_T001 |
| orthologID_006877 | PH01001385G0330 | LOC_Os01g16000.1 | Bradi2g09760.1 | HORVU3Hr1G037050.3 | TRIUR3_30793 | Seita.5G052200.1 | Sobic.003G121800.1 | Zm00001d040360_T001 |
| orthologID_006167 | PH01001731G0490 | LOC_Os01g51810.1 | Bradi2g48200.2 | HORVU3Hr1G067720.1 | TRIUR3_22086 | Seita.5G295700.1 | Sobic.003G277100.1 | Zm00001d042067_T001 |
| orthologID_006253 | PH01001461G0110 | LOC_Os01g11880.1 | Bradi2g07040.1 | HORVU1Hr1G001090.1 | TRIUR3_19547 | Seita.5G151900.1 | Sobic.003G017700.2 | Zm00001d039363_T001 |
| orthologID_006252 | PH01000778G0710 | LOC_Os01g48380.1 | Bradi2g46360.1 | HORVU3Hr1G061950.1 | TRIUR3_29300 | Seita.5G267700.1 | Sobic.003G254000.1 | Zm00001d011579_T001 |
| orthologID_006411 | PH01000240G0240 | LOC_Os01g48930.1 | Bradi2g46660.1 | HORVU3Hr1G062980.1 | TRIUR3_19603 | Seita.5G272200.1 | Sobic.003G258400.1 | Zm00001d043850_T001 |
| orthologID_006305 | PH01000622G0770 | LOC_Os01g74152.1 | Bradi2g62150.2 | HORVU3Hr1G115580.5 | TRIUR3_02366 | Seita.5G468600.1 | Sobic.003G444200.1 | Zm00001d011782_T002 |
| orthologID_007176 | PH01000641G0480 | LOC_Os10g08940.1 | Bradi4g05550.1 | HORVU5Hr1G024500.6 | TRIUR3_04874 | Seita.5G060000.1 | Sobic.003G114400.1 | Zm00001d040268_T002 |
| orthologID_006425 | PH01000155G0900 | LOC_Os01g55870.1 | Bradi2g50800.1 | HORVU3Hr1G074220.2 | TRIUR3_10088 | Seita.5G328800.1 | Sobic.003G306100.1 | Zm00001d012674_T010 |
| orthologID_006772 | PH01000972G0260 | LOC_Os01g45450.1 | Bradi2g44936.1 | HORVU3Hr1G057540.3 | TRIUR3_10202 | Seita.5G240900.1 | Sobic.003G235300.1 | Zm00001d011459_T001 |
| orthologID_006388 | PH01001418G0370 | LOC_Os01g69970.1 | Bradi2g59226.1 | HORVU3Hr1G095360.26 | TRIUR3_03779 | Seita.5G433800.1 | Sobic.003G408300.1 | Zm00001d042306_T004 |
| orthologID_006512 | PH01000155G0370 | LOC_Os01g55700.1 | Bradi2g50630.3 | HORVU3Hr1G073750.4 | TRIUR3_26430 | Seita.5G327200.1 | Sobic.003G304400.1 | Zm00001d012685_T004 |
| orthologID_006565 | PH01005022G0030 | LOC_Os01g13040.1 | Bradi2g07780.2 | HORVU3Hr1G031540.2 | TRIUR3_25658 | Seita.5G007900.1 | Sobic.003G007600.1 | Zm00001d039297_T001 |
| orthologID_006653 | PH01000309G0360 | LOC_Os01g64310.1 | Bradi2g55780.1 | HORVU3Hr1G083820.9 | TRIUR3_34272 | Seita.5G391200.1 | Sobic.003G365500.1 | Zm00001d042758_T001 |
| orthologID_006237 | PH01000032G1630 | LOC_Os01g58380.1 | Bradi2g52350.1 | HORVU3Hr1G078100.1 | TRIUR3_17685 | Seita.5G347500.1 | Sobic.003G324000.1 | Zm00001d012590_T001 |
| orthologID_006737 | PH01000689G0520 | LOC_Os01g32350.1 | Bradi2g13680.1 | HORVU3Hr1G044530.1 | TRIUR3_29433 | Seita.5G177800.1 | Sobic.003G170200.1 | Zm00001d009339_T001 |
| orthologID_006854 | PH01000762G0530 | LOC_Os01g01080.1 | Bradi2g03740.1 | HORVU3Hr1G015150.11 | TRIUR3_09166 | Seita.5G066000.1 | Sobic.003G109100.1 | Zm00001d008389_T003 |
| orthologID_006502 | PH01000004G3000 | LOC_Os01g63840.1 | Bradi2g55417.1 | HORVU3Hr1G084600.10 | TRIUR3_16473 | Seita.5G387800.1 | Sobic.003G362000.1 | Zm00001d042802_T001 |
| orthologID_006652 | PH01008070G0030 | LOC_Os07g09330.1 | Bradi2g09540.1 | HORVU3Hr1G036430.4 | TRIUR3_22111 | Seita.5G054400.1 | Sobic.003G119900.1 | Zm00001d008310_T001 |
| orthologID_006776 | PH01000525G0040 | LOC_Os01g03730.1 | Bradi2g01980.1 | HORVU3Hr1G003850.2 | TRIUR3_10610 | Seita.5G087400.1 | Sobic.003G087200.1 | Zm00001d008529_T001 |
| orthologID_006774 | PH01004720G0080 | LOC_Os01g36220.1 | Bradi2g40582.1 | HORVU3Hr1G047180.1 | TRIUR3_24759 | Seita.5G191600.1 | Sobic.003G183000.1 | Zm00001d044546_T001 |
| orthologID_006226 | PH01000346G0740 | LOC_Os01g42480.1 | Bradi2g43200.1 | HORVU3Hr1G053460.3 | TRIUR3_20926 | Seita.5G222700.1 | Sobic.003G217100.1 | Zm00001d011325_T001 |
| orthologID_006932 | PH01002045G0300 | LOC_Os01g11130.1 | Bradi2g06660.2 | HORVU3Hr1G029060.18 | TRIUR3_13431 | Seita.5G147300.1 | Sobic.003G022900.1 | Zm00001d039400_T002 |
| orthologID_006656 | PH01000504G0650 | LOC_Os01g21940.1 | Bradi2g12110.1 | HORVU0Hr1G006640.1 | TRIUR3_14051 | Seita.5G154200.1 | Sobic.003G151200.1 | Zm00001d040689_T001 |
| orthologID_006601 | PH01003150G0250 | LOC_Os01g10250.1 | Bradi2g06120.1 | HORVU3Hr1G027160.1 | TRIUR3_29756 | Seita.5G140900.1 | Sobic.003G028900.1 | Zm00001d039416_T001 |
| orthologID_006468 | PH01002809G0060 | LOC_Os01g70580.1 | Bradi2g59740.2 | HORVU3Hr1G097210.1 | TRIUR3_02891 | Seita.5G440600.1 | Sobic.003G414600.1 | Zm00001d042241_T001 |
| orthologID_006882 | PH01003107G0180 | LOC_Os01g10260.1 | Bradi2g06150.1 | HORVU3Hr1G027080.2 | TRIUR3_31796 | Seita.5G141000.1 | Sobic.003G028800.1 | Zm00001d039420_T001 |
| orthologID_006474 | PH01000048G0920 | LOC_Os01g51320.1 | Bradi2g47960.1 | HORVU3Hr1G066920.1 | TRIUR3_28466 | Seita.5G291400.1 | Sobic.003G274600.1 | Zm00001d043681_T001 |
| orthologID_006961 | PH01000048G1000 | LOC_Os01g51360.1 | Bradi2g47970.1 | HORVU3Hr1G066960.1 | TRIUR3_32253 | Seita.5G291500.1 | Sobic.003G274700.1 | Zm00001d043680_T001 |
| orthologID_006841 | PH01000293G0600 | LOC_Os01g64870.1 | Bradi2g56190.1 | HORVU3Hr1G082900.1 | TRIUR3_32632 | Seita.5G394700.1 | Sobic.003G369400.2 | Zm00001d042712_T001 |
| orthologID_006722 | PH01000214G1070 | LOC_Os01g53720.2 | Bradi2g49310.3 | HORVU3Hr1G070370.1 | TRIUR3_08074 | Seita.5G311500.1 | Sobic.003G290200.1 | Zm00001d043526_T004 |
| orthologID_006581 | PH01000090G1270 | LOC_Os01g63400.1 | Bradi2g55190.2 | HORVU3Hr1G085220.1 | TRIUR3_13641 | Seita.5G385000.1 | Sobic.003G359400.1 | Zm00001d012340_T001 |
| orthologID_006150 | PH01001236G0210 | LOC_Os01g70930.1 | Bradi2g60070.1 | HORVU3Hr1G098580.2 | TRIUR3_03078 | Seita.5G444000.1 | Sobic.003G418000.1 | Zm00001d042192_T001 |
| orthologID_008155 | PH01002305G0030 | LOC_Os07g23244.1 | Bradi1g60830.1 | HORVU7Hr1G057640.1 | TRIUR3_35120 | Seita.9G380300.1 | Sobic.003G022800.1 | Zm00001d039395_T001 |
| orthologID_006732 | PH01000622G0360 | LOC_Os01g73750.1 | Bradi2g61920.1 | HORVU3Hr1G113720.11 | TRIUR3_08581 | Seita.5G466600.1 | Sobic.003G441900.1 | Zm00001d041983_T006 |
| orthologID_006594 | PH01002356G0280 | LOC_Os01g47340.1 | Bradi2g45740.1 | HORVU3Hr1G059830.8 | TRIUR3_18614 | Seita.5G253800.1 | Sobic.003G246700.1 | Zm00001d043971_T001 |
| orthologID_006950 | PH01001378G0360 | LOC_Os01g52180.1 | Bradi2g48380.1 | HORVU3Hr1G068280.1 | TRIUR3_24326 | Seita.5G299200.1 | Sobic.003G278800.2 | Zm00001d043616_T001 |
| orthologID_006909 | PH01001753G0040 | LOC_Os01g48130.2 | Bradi2g46197.2 | HORVU3Hr1G061560.1 | TRIUR3_27393 | Seita.5G265500.1 | Sobic.003G251800.2 | Zm00001d043921_T001 |
| orthologID_006515 | PH01000031G0410 | LOC_Os01g67570.1 | Bradi2g57887.1 | HORVU3Hr1G088970.1 | TRIUR3_17954 | Seita.5G417800.1 | Sobic.003G392000.1 | Zm00001d012141_T001 |
| orthologID_007859 | PH01000357G0030 | LOC_Os01g34620.2 | Bradi1g08020.1 | HORVU3Hr1G023970.2 | TRIUR3_19635 | Seita.5G123500.1 | Sobic.003G048600.1 | Zm00001d008736_T001 |
| orthologID_006626 | PH01000235G1050 | LOC_Os01g67390.1 | Bradi2g57740.2 | HORVU3Hr1G089170.8 | TRIUR3_12122 | Seita.5G415700.1 | Sobic.003G390500.1 | Zm00001d042493_T001 |
| orthologID_006675 | PH01000260G1030 | LOC_Os01g01870.1 | Bradi2g00740.1 | HORVU3Hr1G000150.2 | TRIUR3_14872 | Seita.5G073000.1 | Sobic.003G102200.1 | Zm00001d040148_T001 |
| orthologID_006727 | PH01000209G0270 | LOC_Os01g16550.1 | Bradi2g10260.1 | HORVU3Hr1G038540.1 | TRIUR3_27234 | Seita.5G045100.1 | Sobic.003G128200.1 | Zm00001d008257_T001 |
| orthologID_006407 | PH01002127G0260 | LOC_Os01g10504.1 | Bradi2g06330.1 | HORVU3Hr1G026650.1 | TRIUR3_34584 | Seita.5G143100.1 | Sobic.003G027000.1 | Zm00001d039434_T001 |
| orthologID_007003 | PH01002846G0040 | LOC_Os01g39630.1 | Bradi2g41710.15 | HORVU0Hr1G003380.14 | TRIUR3_34347 | Seita.5G259300.1 | Sobic.003G023200.1 | Zm00001d044278_T001 |
| orthologID_006259 | PH01000489G0600 | LOC_Os01g68710.2 | Bradi2g58640.1 | HORVU3Hr1G093290.1 | TRIUR3_20703 | Seita.5G426600.1 | Sobic.003G400800.1 | Zm00001d042372_T002 |
| orthologID_006779 | PH01000015G2160 | LOC_Os01g05130.1 | Bradi2g02990.1 | HORVU3Hr1G012280.7 | TRIUR3_29287 | Seita.5G100100.1 | Sobic.003G074000.1 | Zm00001d039874_T001 |
| orthologID_006204 | PH01001474G0360 | LOC_Os01g54390.1 | Bradi2g49727.2 | HORVU3Hr1G071300.1 | TRIUR3_17594 | Seita.5G316600.1 | Sobic.003G294300.1 | Zm00001d043449_T001 |

| | | | | | | | | |
|---|---|---|---|---|---|---|---|---|
| orthologID_006728 | PH01000182G0790 | LOC_Os01g46800.1 | Bradi2g45480.1 | HORVU3Hr1G059220.1 | TRIUR3_29889 | Seita.5G249900.1 | Sobic.003G242800.2 | Zm00001d011496_T002 |
| orthologID_006906 | PH01000333G1160 | LOC_Os01g43070.1 | Bradi2g43680.1 | HORVU7Hr1G113210.5 | TRIUR3_01566 | Seita.5G228200.1 | Sobic.003G221700.1 | Zm00001d044217_T001 |
| orthologID_006696 | PH01001108G0350 | LOC_Os01g47630.1 | Bradi2g45970.5 | HORVU3Hr1G060730.2 | TRIUR3_26099 | Seita.5G262300.1 | Sobic.003G249400.1 | Zm00001d043944_T001 |
| orthologID_006504 | PH01000803G0670 | LOC_Os01g38610.1 | Bradi2g41390.1 | HORVU3Hr1G048770.1 | TRIUR3_28638 | Seita.5G204500.1 | Sobic.003G191800.1 | Zm00001d011187_T001 |
| orthologID_006802 | PH01000155G1290 | LOC_Os01g56030.1 | Bradi2g50920.1 | HORVU3Hr1G074300.1 | TRIUR3_24034 | Seita.5G330300.1 | Sobic.003G307400.1 | Zm00001d043346_T001 |
| orthologID_006754 | PH01000051G0980 | LOC_Os01g45530.1 | Bradi2g44970.1 | HORVU3Hr1G057550.1 | TRIUR3_34231 | Seita.5G241400.1 | Sobic.003G235700.1 | Zm00001d044086_T001 |
| orthologID_006196 | PH01001461G0510 | LOC_Os01g12000.1 | Bradi2g07130.2 | HORVU3Hr1G029400.1 | TRIUR3_16162 | Seita.5G020100.1 | Sobic.003G016100.1 | Zm00001d008946_T001 |
| orthologID_006995 | PH01001641G0310 | LOC_Os01g51010.1 | Bradi2g47747.1 | HORVU3Hr1G066420.1 | TRIUR3_35035 | Seita.5G288500.1 | Sobic.003G272400.1 | Zm00001d011684_T002 |
| orthologID_006811 | PH01004950G0030 | LOC_Os01g47190.1 | Bradi2g45670.1 | HORVU3Hr1G059610.1 | TRIUR3_13076 | Seita.5G253600.1 | Sobic.003G245800.1 | Zm00001d011510_T001 |
| orthologID_006582 | PH01000260G0380 | LOC_Os01g01680.1 | Bradi2g00600.1 | HORVU3Hr1G001540.1 | TRIUR3_02085 | Seita.5G071300.1 | Sobic.003G104200.1 | Zm00001d008408_T001 |
| orthologID_007492 | PH01000372G0480 | LOC_Os09g09790.1 | Bradi4g26940.1 | HORVU5Hr1G052770.1 | TRIUR3_12407 | Seita.1G308100.1 | Sobic.003G125400.1 | Zm00001d008281_T001 |
| orthologID_006573 | PH01000632G0390 | LOC_Os01g14960.1 | Bradi2g08970.1 | HORVU3Hr1G034760.5 | TRIUR3_09620 | Seita.5G059200.1 | Sobic.003G115200.1 | Zm00001d040275_T001 |
| orthologID_006321 | PH01002566G0180 | LOC_Os01g39860.1 | Bradi2g41840.1 | HORVU7Hr1G116830.1 | TRIUR3_35049 | Seita.5G258500.1 | Sobic.003G197200.1 | Zm00001d011208_T001 |
| orthologID_006268 | PH01000346G0930 | LOC_Os01g42630.1 | Bradi2g43310.1 | HORVU3Hr1G053540.26 | TRIUR3_34080 | Seita.5G223800.1 | Sobic.003G218300.1 | Zm00001d011329_T004 |
| orthologID_006366 | PH01000295G0730 | LOC_Os01g58750.1 | Bradi2g52580.1 | HORVU3Hr1G078540.1 | TRIUR3_32989 | Seita.5G350700.1 | Sobic.003G326900.1 | Zm00001d043185_T001 |
| orthologID_006440 | PH01000228G0970 | LOC_Os01g32080.1 | Bradi2g06680.1 | HORVU3Hr1G097230.3 | TRIUR3_12490 | Seita.5G011400.1 | Sobic.003G047100.1 | Zm00001d008739_T001 |
| orthologID_009665 | PH01002609G0030 | LOC_Os08g23410.1 | Bradi3g20660.1 | HORVU5Hr1G021340.1 | TRIUR3_30178 | Seita.6G112800.1 | Sobic.003G439000.1 | Zm00001d042008_T001 |
| orthologID_006496 | PH01002142G0270 | LOC_Os01g08710.1 | Bradi2g05234.2 | HORVU3Hr1G021120.2 | TRIUR3_32723 | Seita.5G129700.1 | Sobic.003G040800.1 | Zm00001d039584_T001 |
| orthologID_006884 | PH01003720G0070 | LOC_Os01g50120.2 | Bradi2g47380.3 | HORVU3Hr1G065350.1 | TRIUR3_26509 | Seita.5G283300.1 | Sobic.003G267500.1 | Zm00001d043764_T005 |
| orthologID_006296 | PH01000284G0450 | LOC_Os01g54090.1 | Bradi2g49580.2 | HORVU3Hr1G070910.3 | TRIUR3_12759 | Seita.5G314700.1 | Sobic.003G292800.1 | Zm00001d043503_T001 |
| orthologID_006699 | PH01001932G0270 | LOC_Os01g12560.1 | Bradi2g07470.1 | HORVU3Hr1G030620.2 | TRIUR3_12973 | Seita.5G014400.1 | Sobic.003G011800.1 | Zm00001d008976_T001 |
| orthologID_006731 | PH01002011G0380 | LOC_Os01g51690.1 | Bradi2g48090.1 | HORVU3Hr1G067510.3 | TRIUR3_33662 | Seita.5G294500.1 | Sobic.003G276000.1 | Zm00001d043663_T001 |
| orthologID_006479 | PH01000267G1220 | LOC_Os01g57240.1 | Bradi2g51790.1 | HORVU3Hr1G076790.3 | TRIUR3_18055 | Seita.5G340400.1 | Sobic.003G317100.1 | Zm00001d043270_T001 |
| orthologID_006490 | PH01000076G1510 | LOC_Os01g52400.1 | Bradi2g48510.3 | HORVU3Hr1G068520.10 | TRIUR3_25770 | Seita.5G301100.1 | Sobic.003G279900.5 | Zm00001d011738_T002 |
| orthologID_006194 | PH01000004G1260 | LOC_Os01g64520.1 | Bradi2g55950.2 | HORVU3Hr1G083560.1 | TRIUR3_32670 | Seita.5G392500.1 | Sobic.003G366400.1 | Zm00001d042746_T003 |
| orthologID_006371 | PH01000346G0710 | LOC_Os01g42470.2 | Bradi2g43190.1 | HORVU3Hr1G053490.2 | TRIUR3_20927 | Seita.5G222100.1 | Sobic.003G216700.2 | Zm00001d044446_T001 |
| orthologID_007091 | PH01002722G0190 | LOC_Os12g08220.1 | Bradi4g40960.1 | HORVU5Hr1G037920.1 | TRIUR3_18154 | Seita.7G281200.1 | Sobic.003G345900.1 | Zm00001d042974_T001 |
| orthologID_006173 | PH01002565G0140 | LOC_Os01g34060.1 | Bradi2g13860.1 | HORVU0Hr1G018970.2 | TRIUR3_30912 | Seita.5G183700.1 | Sobic.003G132300.1 | Zm00001d040464_T001 |
| orthologID_006890 | PH01000568G0020 | LOC_Os01g47330.1 | Bradi2g45730.1 | HORVU3Hr1G059810.2 | TRIUR3_18615 | Seita.5G253700.1 | Sobic.003G246400.1 | Zm00001d043972_T001 |
| orthologID_006819 | PH01005955G0080 | LOC_Os01g11240.1 | Bradi2g06740.1 | HORVU3Hr1G028780.1 | TRIUR3_21435 | Seita.5G147900.1 | Sobic.003G022200.1 | Zm00001d008910_T001 |
| orthologID_006217 | PH01001932G0350 | LOC_Os01g12580.1 | Bradi2g07480.1 | HORVU3Hr1G030650.1 | TRIUR3_26240 | Seita.5G014300.1 | Sobic.003G011400.1 | Zm00001d009382_T001 |
| orthologID_006421 | PH01000190G0040 | LOC_Os01g12540.1 | Bradi2g07437.1 | HORVU3Hr1G030640.21 | TRIUR3_12971 | Seita.5G014700.1 | Sobic.003G012000.1 | Zm00001d039319_T008 |
| orthologID_006807 | PH01000309G1260 | LOC_Os01g64720.1 | Bradi2g56090.3 | HORVU3Hr1G083260.1 | TRIUR3_15888 | Seita.5G393700.1 | Sobic.003G368200.1 | Zm00001d042722_T001 |
| orthologID_006359 | PH01003108G0030 | LOC_Os01g14510.1 | Bradi2g08660.1 | HORVU3Hr1G033790.2 | TRIUR3_23117 | Seita.5G000900.1 | Sobic.003G000500.1 | Zm00001d039243_T001 |
| orthologID_006958 | PH01000317G0560 | LOC_Os01g65890.1 | Bradi2g56900.1 | HORVU3Hr1G091240.3 | TRIUR3_26380 | Seita.5G403400.1 | Sobic.003G377800.1 | Zm00001d000223_T001 |
| orthologID_008667 | PH01000238G0630 | LOC_Os01g23630.1 | Bradi1g31307.2 | HORVU1Hr1G024140.2 | TRIUR3_18086 | Seita.5G159000.1 | Sobic.003G155700.1 | Zm00001d040757_T001 |
| orthologID_006912 | PH01000112G0590 | LOC_Os01g60270.1 | Bradi2g53390.3 | HORVU3Hr1G080660.4 | TRIUR3_23636 | Seita.5G360500.1 | Sobic.003G336600.1 | Zm00001d043076_T001 |
| orthologID_007006 | PH01000284G1000 | LOC_Os01g54400.1 | Bradi2g49740.1 | HORVU3Hr1G071320.1 | TRIUR3_12192 | Seita.5G316700.1 | Sobic.003G294400.1 | Zm00001d043450_T001 |
| orthologID_006769 | PH01000425G0390 | LOC_Os01g57690.1 | Bradi2g52050.1 | HORVU3Hr1G077560.2 | TRIUR3_18110 | Seita.5G343700.1 | Sobic.003G320500.1 | Zm00001d043242_T001 |
| orthologID_006486 | PH01000697G0130 | LOC_Os01g03090.1 | Bradi2g01570.1 | HORVU3Hr1G005950.1 | TRIUR3_08434 | Seita.5G082400.1 | Sobic.003G093600.1 | Zm00001d008502_T001 |
| orthologID_006916 | PH01000293G0890 | LOC_Os01g64760.1 | Bradi2g56110.1 | HORVU3Hr1G083180.2 | TRIUR3_32185 | Seita.5G394000.1 | Sobic.003G368500.1 | Zm00001d042719_T001 |
| orthologID_006672 | PH01002706G0040 | LOC_Os01g36940.1 | Bradi2g40830.1 | HORVU3Hr1G047440.2 | TRIUR3_27944 | Seita.5G196900.1 | Sobic.003G186600.1 | Zm00001d011145_T001 |
| orthologID_007563 | PH01000622G0380 | LOC_Os01g73760.1 | Bradi1g70200.1 | HORVU2Hr1G062320.8 | TRIUR3_10896 | Seita.5G466700.1 | Sobic.003G442000.1 | Zm00001d041982_T001 |
| orthologID_006976 | PH01003207G0120 | LOC_Os01g57050.1 | Bradi2g51696.1 | HORVU3Hr1G076550.1 | TRIUR3_34457 | Seita.5G339300.1 | Sobic.003G316000.1 | Zm00001d043276_T001 |
| orthologID_006525 | PH01000090G1490 | LOC_Os01g63260.1 | Bradi2g55110.1 | HORVU3Hr1G085400.1 | TRIUR3_34530 | Seita.5G383700.1 | Sobic.003G358400.2 | Zm00001d042843_T001 |
| orthologID_006181 | PH01006336G0020 | LOC_Os01g39780.1 | Bradi2g41770.1 | HORVU3Hr1G050210.2 | TRIUR3_21318 | Seita.5G259000.1 | Sobic.003G196600.1 | Zm00001d011204_T001 |
| orthologID_006345 | PH01003393G0080 | LOC_Os01g42190.1 | Bradi2g43040.1 | HORVU3Hr1G053060.4 | TRIUR3_25341 | Seita.5G219300.1 | Sobic.003G214000.1 | Zm00001d044426_T001 |
| orthologID_006417 | PH01000527G0500 | LOC_Os01g50750.1 | Bradi2g47600.1 | HORVU3Hr1G065900.2 | TRIUR3_26831 | Seita.5G286300.1 | Sobic.003G270400.3 | Zm00001d043728_T001 |
| orthologID_006577 | PH01001548G0260 | LOC_Os01g49010.1 | Bradi2g46720.1 | HORVU3Hr1G063320.34 | TRIUR3_28541 | Seita.5G273300.1 | Sobic.003G259600.1 | Zm00001d043840_T001 |
| orthologID_006241 | PH01002236G0120 | LOC_Os01g50460.1 | Bradi2g47540.2 | HORVU3Hr1G065770.8 | TRIUR3_25029 | Seita.5G284900.1 | Sobic.003G269300.1 | Zm00001d043735_T001 |

| | | | | | | | | |
|---|---|---|---|---|---|---|---|---|
| orthologID_006499 | PH01005377G0030 | LOC_Os01g46600.1 | Bradi2g45410.1 | HORVU3Hr1G059030.1 | TRIUR3_33623 | Seita.5G248000.1 | Sobic.003G241400.1 | Zm00001d044022_T001 |
| orthologID_006668 | PH01000316G0160 | LOC_Os01g68390.1 | Bradi2g58410.3 | HORVU3Hr1G092770.7 | TRIUR3_23315 | Seita.5G424100.1 | Sobic.003G398500.2 | Zm00001d042393_T002 |
| orthologID_006701 | PH01000111G0080 | LOC_Os01g65740.1 | Bradi2g56800.2 | HORVU3Hr1G091440.2 | TRIUR3_31642 | Seita.5G401700.1 | Sobic.003G376600.2 | Zm00001d042631_T001 |
| orthologID_006460 | PH01001641G0340 | LOC_Os01g50960.1 | Bradi2g47740.1 | HORVU3Hr1G066410.2 | TRIUR3_30405 | Seita.5G288400.1 | Sobic.003G272300.3 | Zm00001d011680_T002 |
| orthologID_006381 | PH01000402G1070 | LOC_Os01g56235.1 | Bradi2g51060.3 | HORVU3Hr1G074720.6 | TRIUR3_13484 | Seita.5G331900.1 | Sobic.003G308900.1 | Zm00001d043333_T001 |
| orthologID_009641 | PH01005781G0020 | LOC_Os08g40230.1 | Bradi3g11230.1 | HORVU3Hr1G073590.1 | TRIUR3_33076 | Seita.1G147800.1 | Sobic.003G396800.2 | Zm00001d042427_T001 |
| orthologID_006966 | PH01000633G0150 | LOC_Os01g39790.1 | Bradi2g41780.1 | HORVU3Hr1G050240.2 | TRIUR3_29702 | Seita.5G258800.1 | Sobic.003G196900.1 | Zm00001d044288_T001 |
| orthologID_006234 | PH01000058G1900 | LOC_Os01g72950.1 | Bradi2g61410.1 | HORVU3Hr1G110660.1 | TRIUR3_24045 | Seita.5G460600.1 | Sobic.003G433900.1 | Zm00001d011826_T001 |
| orthologID_009723 | PH01000074G1160 | LOC_Os08g31810.3 | Bradi3g35650.1 | HORVU6Hr1G002240.4 | TRIUR3_16202 | Seita.1G092300.1 | Sobic.003G424800.1 | Zm00001d050300_T001 |
| orthologID_006694 | PH01000066G1200 | LOC_Os01g65370.1 | Bradi2g56520.1 | HORVU3Hr1G082210.1 | TRIUR3_22555 | Seita.5G398000.1 | Sobic.003G373000.1 | Zm00001d042665_T001 |
| orthologID_006574 | PH01000781G0740 | LOC_Os01g72800.1 | Bradi2g61320.1 | HORVU3Hr1G110160.4 | TRIUR3_10773 | Seita.5G459400.1 | Sobic.003G432500.1 | Zm00001d042044_T001 |
| orthologID_006126 | PH01001418G0380 | LOC_Os01g69960.1 | Bradi2g59220.1 | HORVU3Hr1G095340.1 | TRIUR3_17715 | Seita.5G433700.1 | Sobic.003G408200.1 | Zm00001d042307_T001 |
| orthologID_006796 | PH01002354G0080 | LOC_Os01g12090.1 | Bradi2g07170.1 | HORVU3Hr1G029510.1 | TRIUR3_18443 | Seita.5G019300.1 | Sobic.003G015400.1 | Zm00001d039348_T001 |
| orthologID_006374 | PH01001090G0370 | LOC_Os01g02870.1 | Bradi2g01340.1 | HORVU3Hr1G002710.4 | TRIUR3_11376 | Seita.5G079300.1 | Sobic.003G096100.1 | Zm00001d040089_T001 |
| orthologID_006190 | PH01000762G0520 | LOC_Os01g01070.1 | Bradi2g03750.1 | HORVU3Hr1G015180.14 | TRIUR3_09164 | Seita.5G065100.1 | Sobic.003G109500.1 | Zm00001d008388_T002 |
| orthologID_006837 | PH01000817G0410 | LOC_Os01g49350.1 | Bradi2g46900.1 | HORVU3Hr1G063860.2 | TRIUR3_03192 | Seita.5G275900.1 | Sobic.003G261400.4 | Zm00001d043813_T002 |
| orthologID_007541 | PH01002240G0140 | LOC_Os12g27070.1 | Bradi4g07780.1 | HORVU7Hr1G043340.7 | TRIUR3_14600 | Seita.3G315200.1 | Sobic.003G212500.1 | Zm00001d044417_T001 |
| orthologID_006736 | PH01000317G0690 | LOC_Os01g65840.1 | Bradi2g56860.1 | HORVU3Hr1G091310.2 | TRIUR3_19777 | Seita.5G402800.1 | Sobic.003G377200.1 | Zm00001d000220_T001 |
| orthologID_006988 | PH01001004G0320 | LOC_Os01g40310.1 | Bradi2g42040.1 | HORVU3Hr1G050800.1 | TRIUR3_32760 | Seita.5G256300.1 | Sobic.003G199800.5 | Zm00001d044310_T001 |
| orthologID_006493 | PH01003126G0140 | LOC_Os01g05680.1 | Bradi2g03310.1 | HORVU3Hr1G013720.1 | TRIUR3_23789 | Seita.5G103600.1 | Sobic.003G071500.2 | Zm00001d039856_T002 |
| orthologID_006356 | PH01001024G0200 | LOC_Os01g72340.1 | Bradi2g61010.1 | HORVU3Hr1G108610.11 | TRIUR3_22214 | Seita.5G455400.1 | Sobic.003G428400.3 | Zm00001d042066_T016 |
| orthologID_006678 | PH01000139G0430 | LOC_Os01g62970.1 | Bradi2g54960.1 | HORVU3Hr1G085690.2 | TRIUR3_20661 | Seita.5G381700.1 | Sobic.003G356400.2 | Zm00001d042863_T001 |
| orthologID_006846 | PH01000317G0210 | LOC_Os01g66000.1 | Bradi2g56960.1 | HORVU3Hr1G091140.2 | TRIUR3_05116 | Seita.5G404400.1 | Sobic.003G378600.1 | Zm00001d042620_T001 |
| orthologID_006245 | PH01000817G0470 | LOC_Os01g49330.1 | Bradi2g46860.3 | HORVU3Hr1G063710.13 | TRIUR3_03195 | Seita.5G275600.1 | Sobic.003G261200.1 | Zm00001d043815_T005 |
| orthologID_006455 | PH01000009G2290 | LOC_Os01g60700.1 | Bradi2g53550.1 | HORVU3Hr1G081100.1 | TRIUR3_24752 | Seita.5G362700.1 | Sobic.003G338400.1 | Zm00001d043056_T002 |
| orthologID_006899 | PH01000207G0010 | LOC_Os01g21990.1 | Bradi2g12140.1 | HORVU0Hr1G006680.2 | TRIUR3_32125 | Seita.5G154500.1 | Sobic.003G151500.1 | Zm00001d040696_T001 |
| orthologID_006735 | PH01001108G0360 | LOC_Os01g47635.1 | Bradi2g45980.1 | HORVU3Hr1G060720.3 | TRIUR3_26098 | Seita.5G262400.1 | Sobic.003G249500.1 | Zm00001d043943_T001 |
| orthologID_006323 | PH01000098G0880 | LOC_Os01g06640.1 | Bradi2g03830.4 | HORVU3Hr1G018680.1 | TRIUR3_07421 | Seita.5G113600.1 | Sobic.003G061400.1 | Zm00001d039764_T002 |
| orthologID_006766 | PH01001339G0170 | LOC_Os01g06000.1 | Bradi2g00520.1 | HORVU3Hr1G104500.10 | TRIUR3_22823 | Seita.5G107500.1 | Sobic.003G067201.1 | Zm00001d039808_T001 |
| orthologID_006849 | PH01001150G0090 | LOC_Os05g37930.1 | Bradi2g23580.1 | HORVU1Hr1G073010.2 | TRIUR3_28048 | Seita.3G221600.1 | Sobic.009G159100.1 | Zm00001d038346_T002 |
| orthologID_006646 | PH01002872G0020 | LOC_Os05g43860.1 | Bradi2g19935.1 | HORVU1Hr1G076890.1 | TRIUR3_14725 | Seita.3G180800.1 | Sobic.009G196001.1 | Zm00001d038692_T001 |
| orthologID_006869 | PH01000680G0470 | LOC_Os05g02650.1 | Bradi2g37530.1 | HORVU1Hr1G007580.4 | TRIUR3_17287 | Seita.3G041300.1 | Sobic.009G020500.1 | Zm00001d035574_T001 |
| orthologID_006644 | PH01000391G0160 | LOC_Os05g51450.1 | Bradi2g14460.1 | HORVU1Hr1G094480.1 | TRIUR3_32709 | Seita.3G122600.1 | Sobic.009G255300.1 | Zm00001d039167_T001 |
| orthologID_006275 | PH01089747G0010 | LOC_Os05g36990.1 | Bradi2g23996.1 | HORVU1Hr1G074030.1 | TRIUR3_12577 | Seita.3G225700.1 | Sobic.009G154000.1 | Zm00001d038284_T001 |
| orthologID_006901 | PH01000445G0600 | LOC_Os05g04370.1 | Bradi2g38650.1 | HORVU1Hr1G016630.3 | TRIUR3_22581 | Seita.3G052300.1 | Sobic.009G032500.1 | Zm00001d010927_T001 |
| orthologID_006592 | PH01001316G0020 | LOC_Os05g33810.1 | Bradi2g25580.1 | HORVU1Hr1G060770.10 | TRIUR3_20168 | Seita.3G246000.1 | Sobic.009G135000.1 | Zm00001d010309_T006 |
| orthologID_007759 | PH01000750G0610 | LOC_Os03g52475.1 | Bradi1g09520.1 | HORVU4Hr1G010970.2 | TRIUR3_23855 | Seita.9G102700.1 | Sobic.009G013800.1 | Zm00001d035658_T001 |
| orthologID_006740 | PH01002000G0220 | LOC_Os05g35540.1 | Bradi2g24370.2 | HORVU1Hr1G074420.2 | TRIUR3_07667 | Seita.3G232000.1 | Sobic.009G148500.1 | Zm00001d038337_T001 |
| orthologID_006519 | PH01000298G0890 | LOC_Os05g48840.1 | Bradi2g16640.2 | HORVU1Hr1G087380.1 | TRIUR3_27181 | Seita.3G147300.1 | Sobic.009G231500.1 | Zm00001d009557_T001 |
| orthologID_006224 | PH01001740G0230 | LOC_Os02g57540.1 | Bradi2g18150.1 | HORVU1Hr1G079370.1 | TRIUR3_05237 | Seita.3G162800.1 | Sobic.009G215200.2 | Zm00001d009417_T006 |
| orthologID_006576 | PH01000428G0730 | LOC_Os05g03070.1 | Bradi2g37850.1 | HORVU1Hr1G011890.1 | TRIUR3_21414 | Seita.3G045000.1 | Sobic.009G025100.1 | Zm00001d010985_T001 |
| orthologID_006387 | PH01000502G0320 | LOC_Os05g45920.1 | Bradi2g18650.1 | HORVU1Hr1G080410.11 | TRIUR3_08665 | Seita.3G165500.1 | Sobic.009G212000.1 | Zm00001d038835_T002 |
| orthologID_009501 | PH01001445G0220 | LOC_Os05g27340.1 | Bradi3g37450.2 | HORVU2Hr1G068150.5 | TRIUR3_14175 | Seita.3G282800.1 | Sobic.009G099500.1 | Zm00001d038038_T001 |
| orthologID_006616 | PH01000423G0800 | LOC_Os05g43610.1 | Bradi2g20260.1 | HORVU1Hr1G077590.4 | TRIUR3_20214 | Seita.3G183600.1 | Sobic.009G192700.1 | Zm00001d038667_T001 |
| orthologID_006312 | PH01000348G0920 | LOC_Os05g38170.1 | Bradi2g23490.1 | HORVU1Hr1G072750.2 | TRIUR3_22547 | Seita.3G220200.1 | Sobic.009G160500.1 | Zm00001d038362_T002 |
| orthologID_006132 | PH01002959G0010 | LOC_Os05g48360.2 | Bradi2g17010.1 | HORVU1Hr1G085480.1 | TRIUR3_20323 | Seita.3G150700.1 | Sobic.009G227800.1 | Zm00001d038971_T001 |
| orthologID_006227 | PH01001556G0450 | LOC_Os05g05560.1 | Bradi2g36020.1 | HORVU1Hr1G020340.1 | TRIUR3_30639 | Seita.3G039100.1 | Sobic.009G042900.1 | Zm00001d010872_T001 |
| orthologID_006940 | PH01000271G0660 | LOC_Os05g45910.1 | Bradi2g18660.1 | HORVU1Hr1G080440.1 | TRIUR3_08666 | Seita.3G165700.1 | Sobic.009G211900.1 | Zm00001d000345_T001 |
| orthologID_006309 | PH01001057G0380 | LOC_Os05g02990.2 | Bradi2g37770.1 | HORVU1Hr1G010940.3 | TRIUR3_16970 | Seita.3G044200.1 | Sobic.009G024200.1 | Zm00001d035519_T001 |
| orthologID_006497 | PH01001331G0210 | LOC_Os05g04180.1 | Bradi2g38497.1 | HORVU1Hr1G016240.1 | TRIUR3_15613 | Seita.3G051600.1 | Sobic.009G031500.1 | Zm00001d010944_T001 |

| | | | | | | | | |
|---|---|---|---|---|---|---|---|---|
| orthologID_006266 | PH01003128G0230 | LOC_Os05g38330.1 | Bradi2g23430.1 | HORVU1Hr1G072640.2 | TRIUR3_08456 | Seita.3G218700.1 | Sobic.009G161900.1 | Zm00001d010519_T002 |
| orthologID_006962 | PH01000206G1300 | LOC_Os05g39250.1 | Bradi2g22660.1 | HORVU1Hr1G070690.1 | TRIUR3_32762 | Seita.3G209800.1 | Sobic.009G168800.1 | Zm00001d010586_T001 |
| orthologID_007011 | PH01001131G0060 | LOC_Os05g08920.1 | Bradi2g33630.1 | HORVU1Hr1G027490.3 | TRIUR3_32654 | Seita.3G107400.1 | Sobic.009G068000.2 | Zm00001d009943_T001 |
| orthologID_006526 | PH01000011G2340 | LOC_Os05g33500.1 | Bradi2g25830.1 | HORVU1Hr1G061210.7 | TRIUR3_11739 | Seita.3G248600.1 | Sobic.009G131600.1 | Zm00001d010340_T001 |
| orthologID_006783 | PH01000431G0220 | LOC_Os05g07210.1 | Bradi2g34560.1 | HORVU1Hr1G024390.1 | TRIUR3_25241 | Seita.7G273400.1 | Sobic.009G057700.1 | Zm00001d010838_T001 |
| orthologID_006394 | PH01000237G0030 | LOC_Os05g43810.1 | Bradi2g19997.3 | HORVU1Hr1G076940.1 | TRIUR3_03396 | Seita.3G181400.1 | Sobic.009G195500.2 | Zm00001d038687_T001 |
| orthologID_006528 | PH01008552G0020 | LOC_Os05g38250.1 | Bradi2g23450.1 | HORVU1Hr1G072710.2 | TRIUR3_08459 | Seita.3G219300.1 | Sobic.009G161200.1 | Zm00001d010053_T001 |
| orthologID_006494 | PH01000296G0640 | LOC_Os05g27870.1 | Bradi2g29750.1 | HORVU1Hr1G051570.3 | TRIUR3_12220 | Seita.3G281100.1 | Sobic.009G101200.1 | Zm00001d038006_T001 |
| orthologID_006926 | PH01000974G0220 | LOC_Os05g38680.1 | Bradi2g23060.1 | HORVU1Hr1G071910.1 | TRIUR3_13829 | Seita.3G215000.1 | Sobic.009G165200.1 | Zm00001d038387_T001 |
| orthologID_006657 | PH01001655G0240 | LOC_Os05g38850.1 | Bradi2g22907.1 | HORVU1Hr1G071560.4 | TRIUR3_25923 | Seita.3G213500.1 | Sobic.009G166400.1 | Zm00001d010567_T005 |
| orthologID_006357 | PH01000022G2160 | LOC_Os05g38710.1 | Bradi2g23040.5 | HORVU1Hr1G071890.16 | TRIUR3_29916 | Seita.3G214700.1 | Sobic.009G165400.1 | Zm00001d038389_T008 |
| orthologID_006713 | PH01000271G0420 | LOC_Os05g46020.1 | Bradi2g18530.1 | HORVU1Hr1G080300.1 | TRIUR3_01979 | Seita.3G164900.1 | Sobic.009G212800.1 | Zm00001d038843_T001 |
| orthologID_006611 | PH01000530G0360 | LOC_Os05g51754.1 | Bradi2g14190.1 | HORVU1Hr1G095210.4 | TRIUR3_27901 | Seita.3G118600.1 | Sobic.009G259700.1 | Zm00001d009776_T001 |
| orthologID_009504 | PH01000530G0690 | LOC_Os05g51640.1 | Bradi3g37880.2 | HORVU1Hr1G064630.6 | TRIUR3_29407 | Seita.3G119900.1 | Sobic.009G258400.1 | Zm00001d039154_T001 |
| orthologID_006674 | PH01000445G0250 | LOC_Os05g04584.1 | Bradi2g36910.1 | HORVU1Hr1G017410.1 | TRIUR3_27396 | Seita.3G054100.1 | Sobic.009G034300.1 | Zm00001d024763_T001 |
| orthologID_006408 | PH01002959G0020 | LOC_Os05g48340.1 | Bradi2g17000.1 | HORVU1Hr1G085470.1 | TRIUR3_20322 | Seita.3G151000.1 | Sobic.009G227700.1 | Zm00001d038970_T004 |
| orthologID_006744 | PH01000391G0080 | LOC_Os05g51240.1 | Bradi2g14500.1 | HORVU1Hr1G094180.2 | TRIUR3_17235 | Seita.3G123300.1 | Sobic.009G254700.1 | Zm00001d039175_T001 |
| orthologID_006350 | PH01000165G1550 | LOC_Os05g46340.1 | Bradi2g18237.1 | HORVU1Hr1G079730.1 | TRIUR3_34287 | Seita.3G163400.1 | Sobic.009G214300.2 | Zm00001d038860_T001 |
| orthologID_006386 | PH01109121G0010 | LOC_Os05g10630.1 | Bradi2g33300.4 | HORVU1Hr1G028330.1 | TRIUR3_22527 | Seita.3G111100.1 | Sobic.009G071700.1 | Zm00001d037675_T001 |
| orthologID_009985 | PH01000005G0920 | LOC_Os05g43050.1 | Bradi2g20530.1 | HORVU1Hr1G075230.1 | TRIUR3_12934 | Seita.3G186500.1 | Sobic.009G190100.1 | Zm00001d038643_T001 |
| orthologID_009954 | PH01000297G0550 | LOC_Os05g46760.1 | Bradi2g17820.1 | HORVU1Hr1G078710.3 | TRIUR3_33794 | Seita.3G160700.1 | Sobic.009G217600.1 | Zm00001d038884_T001 |
| orthologID_006175 | PH01000659G0110 | LOC_Os05g40040.1 | Bradi2g22250.1 | HORVU1Hr1G069650.1 | TRIUR3_20904 | Seita.3G204300.1 | Sobic.009G174000.1 | Zm00001d010615_T001 |
| orthologID_006341 | PH01001592G0370 | LOC_Os05g05720.1 | Bradi2g35800.1 | HORVU1Hr1G021230.11 | TRIUR3_21762 | Seita.3G037500.1 | Sobic.009G045000.1 | Zm00001d024858_T004 |
| orthologID_006810 | PH01000022G0190 | LOC_Os05g39560.1 | Bradi2g22520.1 | HORVU1Hr1G070460.1 | TRIUR3_26930 | Seita.3G208000.1 | Sobic.009G170500.1 | Zm00001d038437_T001 |
| orthologID_006207 | PH01152620G0010 | LOC_Os05g35970.1 | Bradi2g24230.1 | HORVU1Hr1G074850.1 | TRIUR3_22125 | Seita.3G229400.1 | Sobic.009G150700.3 | Zm00001d010183_T001 |
| orthologID_006495 | PH01001794G0430 | LOC_Os05g11570.1 | Bradi2g32880.1 | HORVU6Hr1G022070.1 | TRIUR3_29241 | Seita.3G073400.1 | Sobic.009G076200.1 | Zm00001d010008_T001 |
| orthologID_006986 | PH01000871G0250 | LOC_Os05g41610.1 | Bradi2g21142.1 | HORVU1Hr1G067460.6 | TRIUR3_31119 | Seita.3G192900.1 | Sobic.009G183400.1 | Zm00001d038577_T001 |
| orthologID_006984 | PH01000523G0060 | LOC_Os05g43230.1 | Bradi2g20460.1 | HORVU1Hr1G075530.1 | TRIUR3_18397 | Seita.3G185600.1 | Sobic.009G190600.1 | Zm00001d038650_T001 |
| orthologID_006228 | PH01000949G0530 | LOC_Os05g01950.1 | Bradi2g39200.1 | HORVU1Hr1G004400.6 | TRIUR3_00731 | Seita.3G066700.1 | Sobic.009G010700.1 | Zm00001d011073_T001 |
| orthologID_006992 | PH01000659G0320 | LOC_Os05g39950.1 | Bradi2g22310.2 | HORVU1Hr1G069900.1 | TRIUR3_29052 | Seita.3G205300.1 | Sobic.009G173400.1 | Zm00001d038473_T001 |
| orthologID_006902 | PH01000003G2480 | LOC_Os05g15630.1 | Bradi2g31700.1 | HORVU7Hr1G027410.1 | TRIUR3_26513 | Seita.3G084500.1 | Sobic.009G088900.1 | Zm00001d010038_T001 |
| orthologID_006424 | PH01000212G0990 | LOC_Os05g04830.1 | Bradi2g36720.2 | HORVU1Hr1G018010.2 | TRIUR3_24239 | Seita.3G056000.1 | Sobic.009G036800.1 | Zm00001d010899_T001 |
| orthologID_006397 | PH01000766G0340 | LOC_Os05g33880.1 | Bradi2g25507.3 | HORVU1Hr1G063320.1 | TRIUR3_02297 | Seita.3G244700.1 | Sobic.009G135900.1 | Zm00001d038173_T003 |
| orthologID_006599 | PH01000448G0100 | LOC_Os05g51180.1 | Bradi2g14540.1 | HORVU1Hr1G094100.1 | TRIUR3_18406 | Seita.3G124100.1 | Sobic.009G253400.1 | Zm00001d009740_T004 |
| orthologID_006536 | PH01003212G0030 | LOC_Os05g27000.1 | Bradi2g31080.1 | HORVU3Hr1G085990.1 | TRIUR3_09470 | Seita.3G284100.1 | Sobic.009G098000.1 | Zm00001d010396_T001 |
| orthologID_006332 | PH01000021G2400 | LOC_Os05g34040.1 | Bradi2g25410.1 | HORVU1Hr1G063510.1 | TRIUR3_02783 | Seita.3G243000.1 | Sobic.009G137000.1 | Zm00001d038189_T010 |
| orthologID_006548 | PH01001379G0290 | LOC_Os05g28500.1 | Bradi2g29350.1 | HORVU1Hr1G060170.1 | TRIUR3_17248 | Seita.3G279200.1 | Sobic.009G103000.2 | Zm00001d038022_T001 |
| orthologID_006793 | PH01001837G0380 | LOC_Os05g42190.1 | Bradi2g20710.3 | HORVU1Hr1G066240.1 | TRIUR3_23484 | Seita.3G188900.1 | Sobic.009G188000.1 | Zm00001d010698_T002 |
| orthologID_006939 | PH01000423G0070 | LOC_Os05g43760.1 | Bradi2g20060.4 | HORVU1Hr1G077080.2 | TRIUR3_31674 | Seita.3G181900.1 | Sobic.009G195000.2 | Zm00001d038683_T001 |
| orthologID_006535 | PH01000445G0570 | LOC_Os05g04380.1 | Bradi2g38660.1 | HORVU1Hr1G016660.1 | TRIUR3_32743 | Seita.3G052400.1 | Sobic.009G032600.1 | Zm00001d024734_T001 |
| orthologID_007205 | PH01000344G0050 | LOC_Os11g19460.1 | Bradi4g20190.2 | HORVU4Hr1G012400.12 | TRIUR3_24982 | Seita.8G107900.1 | Sobic.009G094700.1 | Zm00001d049153_T007 |
| orthologID_006518 | PH01000242G1370 | LOC_Os05g49650.1 | Bradi2g15870.1 | HORVU1Hr1G090200.2 | TRIUR3_28336 | Seita.3G139300.1 | Sobic.009G238300.1 | Zm00001d039076_T001 |
| orthologID_006393 | PH01000391G0850 | LOC_Os04g29920.1 | Bradi2g14138.1 | HORVU1Hr1G095310.7 | TRIUR3_34595 | Seita.3G117700.1 | Sobic.009G260000.1 | Zm00001d039213_T001 |
| orthologID_008201 | PH01001655G0300 | LOC_Os10g21000.1 | Bradi1g55820.3 | HORVU1Hr1G071470.1 | TRIUR3_11191 | Seita.3G213200.1 | Sobic.009G166600.1 | Zm00001d038400_T001 |
| orthologID_006372 | PH01000429G0070 | LOC_Os05g45350.1 | Bradi2g19010.1 | HORVU1Hr1G081410.7 | TRIUR3_23776 | Seita.3G169400.1 | Sobic.009G207600.1 | Zm00001d038751_T001 |
| orthologID_006987 | PH01000046G0970 | LOC_Os05g49220.1 | Bradi2g16140.1 | HORVU1Hr1G089430.5 | TRIUR3_06408 | Seita.3G143300.1 | Sobic.009G235100.1 | Zm00001d009596_T001 |
| orthologID_006742 | PH01000237G1180 | LOC_Os05g44170.1 | Bradi2g19680.5 | HORVU7Hr1G035190.2 | TRIUR3_34923 | Seita.3G177400.1 | Sobic.009G199800.1 | Zm00001d038724_T001 |
| orthologID_006871 | PH01001110G0400 | LOC_Os05g07280.1 | Bradi2g34507.1 | HORVU1Hr1G024540.1 | TRIUR3_33954 | Seita.7G274400.1 | Sobic.009G058800.1 | Zm00001d009820_T001 |
| orthologID_006813 | PH01000271G1010 | LOC_Os05g45810.1 | Bradi2g18740.2 | HORVU1Hr1G080820.6 | TRIUR3_19549 | Seita.3G166700.1 | Sobic.009G210300.1 | Zm00001d038730_T001 |
| orthologID_006665 | PH01000024G1060 | LOC_Os05g48160.1 | Bradi2g17200.3 | HORVU2Hr1G042420.1 | TRIUR3_01768 | Seita.3G153500.1 | Sobic.009G225600.1 | Zm00001d038946_T001 |

| | | | | | | | | |
|---|---|---|---|---|---|---|---|---|
| orthologID_006786 | PH01002221G0280 | LOC_Os05g49920.1 | Bradi2g15700.1 | HORVU1Hr1G090780.5 | TRIUR3_24525 | Seita.3G137300.1 | Sobic.009G241200.3 | Zm00001d039096_T001 |
| orthologID_006286 | PH01000772G0320 | LOC_Os05g31254.1 | Bradi2g27097.1 | HORVU1Hr1G057600.5 | TRIUR3_25487 | Seita.3G263200.1 | Sobic.009G119700.1 | Zm00001d038054_T002 |
| orthologID_009978 | PH01000823G0100 | LOC_Os05g04070.1 | Bradi2g38432.1 | HORVU1Hr1G015940.2 | TRIUR3_05963 | Seita.3G050900.1 | Sobic.009G030800.1 | Zm00001d024718_T004 |
| orthologID_006346 | PH01001057G0130 | LOC_Os05g02730.1 | Bradi2g37650.1 | HORVU1Hr1G010580.1 | TRIUR3_16981 | Seita.3G043800.1 | Sobic.009G023900.1 | Zm00001d035535_T001 |
| orthologID_006818 | PH01001512G0140 | LOC_Os05g06050.1 | Bradi2g35570.1 | HORVU1Hr1G021890.1 | TRIUR3_19448 | Seita.3G036100.1 | Sobic.009G046900.1 | Zm00001d010851_T001 |
| orthologID_006500 | PH01001489G0170 | LOC_Os05g41810.1 | Bradi2g21017.3 | HORVU1Hr1G067000.1 | TRIUR3_26377 | Seita.3G191400.1 | Sobic.009G184900.1 | Zm00001d038589_T005 |
| orthologID_006697 | PH01000022G1030 | LOC_Os05g39060.1 | Bradi2g22780.1 | HORVU1Hr1G071200.1 | TRIUR3_11860 | Seita.3G212000.1 | Sobic.009G167600.1 | Zm00001d038408_T001 |
| orthologID_006937 | PH01000273G0650 | LOC_Os05g11810.1 | Bradi2g32577.1 | HORVU1Hr1G029520.3 | TRIUR3_29606 | Seita.3G074700.1 | Sobic.009G077500.1 | Zm00001d037724_T002 |
| orthologID_009709 | PH01001556G0330 | LOC_Os05g05450.1 | Bradi3g49040.6 | HORVU1Hr1G019650.6 | TRIUR3_09282 | Seita.3G040000.1 | Sobic.009G042100.1 | Zm00001d024828_T002 |
| orthologID_006182 | PH01002938G0200 | LOC_Os05g05810.1 | Bradi2g35707.4 | HORVU1Hr1G021560.26 | TRIUR3_26479 | Seita.3G037100.1 | Sobic.009G045400.1 | Zm00001d010866_T004 |
| orthologID_006129 | PH01003376G0110 | LOC_Os05g33360.1 | Bradi2g25870.1 | HORVU1Hr1G061500.2 | TRIUR3_20458 | Seita.3G249300.1 | Sobic.009G131100.1 | Zm00001d038147_T001 |
| orthologID_006821 | PH01000261G1160 | LOC_Os05g40990.1 | Bradi2g21860.1 | HORVU1Hr1G068870.1 | TRIUR3_11643 | Seita.3G198900.1 | Sobic.009G178200.1 | Zm00001d038529_T003 |
| orthologID_006825 | PH01002183G0280 | LOC_Os05g44320.1 | Bradi2g19560.1 | HORVU1Hr1G086380.1 | TRIUR3_12607 | Seita.3G175900.1 | Sobic.009G201200.1 | Zm00001d038807_T005 |
| orthologID_006364 | PH01003432G0030 | LOC_Os05g32270.1 | Bradi2g26987.1 | HORVU1Hr1G056500.15 | TRIUR3_07433 | Seita.3G257700.1 | Sobic.009G124200.1 | Zm00001d038087_T002 |
| orthologID_006682 | PH01003596G0230 | LOC_Os05g51860.1 | Bradi2g14090.3 | HORVU1Hr1G095110.2 | TRIUR3_32880 | Seita.3G117300.1 | Sobic.009G260900.1 | Zm00001d039224_T001 |
| orthologID_006800 | PH01002139G0400 | LOC_Os05g02410.1 | Bradi2g37410.1 | HORVU1Hr1G006880.6 | TRIUR3_04726 | Seita.3G062500.1 | Sobic.009G016500.1 | Zm00001d011049_T001 |
| orthologID_006281 | PH01001950G0080 | LOC_Os05g09640.1 | Bradi2g33340.1 | HORVU1Hr1G028270.2 | TRIUR3_34922 | Seita.3G110000.1 | Sobic.009G070500.1 | Zm00001d037682_T001 |
| orthologID_007506 | PH01003741G0040 | LOC_Os11g25860.1 | Bradi4g08410.1 | HORVU5Hr1G050470.1 | TRIUR3_33897 | Seita.8G112000.1 | Sobic.009G246800.1 | Zm00001d009694_T001 |
| orthologID_007370 | PH01003188G0230 | LOC_Os05g33300.1 | Bradi4g07620.1 | HORVU5Hr1G030330.4 | TRIUR3_27409 | Seita.3G249500.1 | Sobic.009G130800.1 | Zm00001d009928_T001 |
| orthologID_010205 | PH01000595G0780 | LOC_Os05g05030.1 | Bradi2g36620.1 | HORVU1Hr1G018360.1 | TRIUR3_11356 | Seita.3G057200.1 | Sobic.009G038100.1 | Zm00001d024799_T001 |
| orthologID_006401 | PH01000011G1350 | LOC_Os05g33020.1 | Bradi2g26310.11 | HORVU1Hr1G062340.4 | TRIUR3_17103 | Seita.3G251700.1 | Sobic.009G129100.1 | Zm00001d038128_T004 |
| orthologID_006464 | PH01007234G0010 | LOC_Os05g48410.1 | Bradi2g16970.6 | HORVU1Hr1G085550.2 | TRIUR3_26342 | Seita.3G150300.1 | Sobic.009G228300.1 | Zm00001d038973_T002 |
| orthologID_006896 | PH01000258G0460 | LOC_Os05g05480.1 | Bradi2g36130.1 | HORVU1Hr1G020000.3 | TRIUR3_02440 | Seita.3G039500.1 | Sobic.009G042200.1 | Zm00001d024830_T001 |
| orthologID_006636 | PH01001131G0080 | LOC_Os05g08930.1 | Bradi2g33620.1 | HORVU1Hr1G027550.1 | TRIUR3_10581 | Seita.3G107800.1 | Sobic.009G068300.1 | Zm00001d037616_T001 |
| orthologID_006458 | PH01001556G0280 | LOC_Os05g05320.1 | Bradi2g36217.3 | HORVU1Hr1G019600.8 | TRIUR3_18906 | Seita.3G060100.1 | Sobic.009G041800.2 | Zm00001d010880_T001 |
| orthologID_006125 | PH01000348G0940 | LOC_Os05g38180.1 | Bradi2g23480.1 | HORVU1Hr1G072650.3 | TRIUR3_32692 | Seita.3G220100.1 | Sobic.009G160600.1 | Zm00001d038361_T001 |
| orthologID_006893 | PH01004198G0130 | LOC_Os05g34720.1 | Bradi2g25060.1 | HORVU1Hr1G064040.3 | TRIUR3_34854 | Seita.3G237700.1 | Sobic.009G141400.1 | Zm00001d010248_T001 |
| orthologID_006452 | PH01001980G0030 | LOC_Os05g28200.1 | Bradi2g28460.1 | HORVU1Hr1G059890.2 | TRIUR3_08904 | Seita.5G460000.1 | Sobic.009G104700.2 | Zm00001d037984_T004 |
| orthologID_006230 | PH01000410G0260 | LOC_Os05g51470.1 | Bradi2g14450.2 | HORVU1Hr1G094490.1 | TRIUR3_31565 | Seita.3G122400.1 | Sobic.009G255500.1 | Zm00001d039166_T002 |
| orthologID_006693 | PH01003208G0110 | LOC_Os05g03174.2 | Bradi2g37970.1 | HORVU1Hr1G013190.2 | TRIUR3_31415 | Seita.3G046000.1 | Sobic.009G026101.1 | Zm00001d010976_T001 |
| orthologID_006867 | PH01001592G0500 | LOC_Os05g05690.1 | Bradi2g35840.1 | HORVU1Hr1G020420.2 | TRIUR3_27200 | Seita.3G037800.1 | Sobic.009G044700.1 | Zm00001d024854_T001 |
| orthologID_006306 | PH01000203G1070 | LOC_Os05g07632.1 | Bradi2g34490.1 | HORVU1Hr1G024580.5 | TRIUR3_31456 | Seita.3G099200.1 | Sobic.009G059300.1 | Zm00001d037512_T001 |
| orthologID_007005 | PH01003433G0130 | LOC_Os05g44390.1 | Bradi2g19490.1 | HORVU1Hr1G076010.4 | TRIUR3_23963 | Seita.3G175300.1 | Sobic.009G201900.1 | Zm00001d010784_T005 |
| orthologID_006720 | PH01000271G1020 | LOC_Os05g45770.1 | Bradi2g18750.1 | HORVU1Hr1G080840.2 | TRIUR3_19548 | Seita.3G166800.1 | Sobic.009G210200.3 | Zm00001d038732_T001 |
| orthologID_006690 | PH01000342G1020 | LOC_Os05g12140.1 | Bradi2g32277.2 | HORVU3Hr1G084260.1 | TRIUR3_28310 | Seita.3G076000.1 | Sobic.009G079500.1 | Zm00001d010447_T001 |
| orthologID_006320 | PH01000254G0360 | LOC_Os05g49810.1 | Bradi2g15780.1 | HORVU1Hr1G090510.1 | TRIUR3_08107 | Seita.3G138200.1 | Sobic.009G240000.3 | Zm00001d039085_T001 |
| orthologID_008071 | PH01001720G0230 | LOC_Os07g37030.1 | Bradi1g24980.1 | HORVU2Hr1G041610.4 | TRIUR3_06189 | Seita.1G016500.1 | Sobic.009G143100.1 | Zm00001d016134_T001 |
| orthologID_006839 | PH01001601G0390 | LOC_Os05g50720.1 | Bradi2g15420.1 | HORVU1Hr1G092540.1 | TRIUR3_14962 | Seita.3G129300.1 | Sobic.009G248000.1 | Zm00001d039148_T001 |
| orthologID_006847 | PH01000429G0240 | LOC_Os05g45300.1 | Bradi2g19050.1 | HORVU1Hr1G081470.4 | TRIUR3_08630 | Seita.3G170100.1 | Sobic.009G207100.3 | Zm00001d038755_T001 |
| orthologID_006928 | PH01000022G0600 | LOC_Os05g39400.1 | Bradi2g22620.1 | HORVU1Hr1G070600.1 | TRIUR3_05904 | Seita.3G209200.1 | Sobic.009G169300.1 | Zm00001d010591_T001 |
| orthologID_009581 | PH01000796G0390 | LOC_Os02g21240.1 | Bradi3g10990.1 | HORVU1Hr1G004970.3 | TRIUR3_13725 | Seita.1G142600.1 | Sobic.009G034400.1 | Zm00001d010914_T001 |
| orthologID_006169 | PH01008407G0020 | LOC_Os05g31160.1 | Bradi2g27130.1 | HORVU1Hr1G057640.8 | TRIUR3_15861 | Seita.3G263600.1 | Sobic.009G119300.1 | Zm00001d038051_T001 |
| orthologID_006509 | PH01001465G0380 | LOC_Os05g08820.1 | Bradi2g33690.1 | HORVU1Hr1G026800.1 | TRIUR3_29520 | Seita.3G105900.1 | Sobic.009G066600.1 | Zm00001d037714_T001 |
| orthologID_006873 | PH01000794G0690 | LOC_Os05g34760.1 | Bradi2g25040.6 | HORVU1Hr1G063930.5 | TRIUR3_20040 | Seita.3G237500.1 | Sobic.009G141700.1 | Zm00001d038217_T002 |
| orthologID_007956 | PH01004067G0120 | LOC_Os05g27780.1 | Bradi1g11710.1 | HORVU3Hr1G083470.3 | TRIUR3_04413 | Seita.3G281800.1 | Sobic.009G100800.1 | Zm00001d038003_T001 |
| orthologID_006219 | PH01000391G0740 | LOC_Os05g51690.1 | Bradi2g14220.1 | HORVU1Hr1G095410.3 | TRIUR3_20859 | Seita.3G119200.1 | Sobic.009G259100.1 | Zm00001d009773_T001 |
| orthologID_006809 | PH01000165G1190 | LOC_Os05g46395.1 | Bradi2g18170.1 | HORVU1Hr1G079410.3 | TRIUR3_10930 | Seita.3G163000.1 | Sobic.009G214900.1 | Zm00001d038864_T001 |
| orthologID_007351 | PH01001222G0410 | LOC_Os11g06810.1 | Bradi4g24237.1 | HORVU4Hr1G019910.13 | TRIUR3_21010 | Seita.8G047300.1 | Sobic.009G249700.1 | Zm00001d039196_T001 |
| orthologID_006747 | PH01000949G0500 | LOC_Os05g01970.5 | Bradi2g39190.1 | HORVU1Hr1G004430.4 | TRIUR3_00730 | Seita.3G066500.1 | Sobic.009G010900.1 | Zm00001d011069_T004 |
| orthologID_006589 | PH01000622G0450 | LOC_Os05g01710.1 | Bradi2g39377.1 | HORVU1Hr1G003460.1 | TRIUR3_11607 | Seita.3G069800.1 | Sobic.009G004600.1 | Zm00001d011092_T001 |

| | | | | | | | | |
|---|---|---|---|---|---|---|---|---|
| orthologID_006482 | PH01000423G0270 | LOC_Os05g43670.1 | Bradi2g20087.1 | HORVU1Hr1G077190.1 | TRIUR3_05939 | Seita.3G182200.1 | Sobic.009G194700.1 | Zm00001d010727_T001 |
| orthologID_006137 | PH01002234G0150 | LOC_Os05g30710.1 | Bradi2g27440.2 | HORVU1Hr1G058280.1 | TRIUR3_15376 | Seita.3G267700.1 | Sobic.009G116400.4 | Zm00001d037891_T001 |
| orthologID_007883 | PH01000078G1830 | LOC_Os05g01120.1 | Bradi1g09917.1 | HORVU1Hr1G051300.10 | TRIUR3_03766 | Seita.3G116300.1 | Sobic.009G000700.1 | Zm00001d011117_T001 |
| orthologID_006556 | PH01000268G0100 | LOC_Os05g30030.1 | Bradi2g27840.1 | HORVU1Hr1G058770.1 | TRIUR3_24379 | Seita.3G271200.1 | Sobic.009G112900.1 | Zm00001d009910_T001 |
| orthologID_006977 | PH01001318G0310 | LOC_Os05g37330.1 | Bradi2g23830.1 | HORVU1Hr1G073640.1 | TRIUR3_28523 | Seita.3G224100.1 | Sobic.009G155600.1 | Zm00001d010128_T003 |
| orthologID_006941 | PH01000823G0530 | LOC_Os05g03830.1 | Bradi2g38350.1 | HORVU1Hr1G015560.2 | TRIUR3_25790 | Seita.3G050100.1 | Sobic.009G029700.1 | Zm00001d035443_T001 |
| orthologID_006687 | PH01002801G0250 | LOC_Os05g03460.1 | Bradi2g38090.1 | HORVU1Hr1G013840.1 | TRIUR3_04430 | Seita.3G047700.1 | Sobic.009G026900.1 | Zm00001d010971_T003 |
| orthologID_006654 | PH01002011G0250 | LOC_Os05g45954.1 | Bradi2g48130.2 | HORVU1Hr1G080350.3 | TRIUR3_01982 | Seita.3G165200.1 | Sobic.009G212500.1 | Zm00001d011134_T001 |
| orthologID_006770 | PH01001274G0440 | LOC_Os05g01300.1 | Bradi2g39900.1 | HORVU1Hr1G000940.3 | TRIUR3_08334 | Seita.3G114500.1 | Sobic.009G001900.1 | Zm00001d011103_T001 |
| orthologID_006894 | PH01004770G0010 | LOC_Os05g34810.1 | Bradi2g25010.1 | HORVU1Hr1G063880.1 | TRIUR3_21088 | Seita.3G237200.1 | Sobic.009G142100.1 | Zm00001d038220_T001 |
| orthologID_006506 | PH01001064G0420 | LOC_Os05g02400.1 | Bradi2g37420.1 | HORVU1Hr1G006890.1 | TRIUR3_04727 | Seita.3G062600.1 | Sobic.009G016400.1 | Zm00001d035608_T001 |
| orthologID_006513 | PH01007103G0020 | LOC_Os05g42010.1 | Bradi2g20810.3 | HORVU1Hr1G066460.1 | TRIUR3_19123 | Seita.3G190200.1 | Sobic.009G186700.2 | Zm00001d038600_T001 |
| orthologID_006480 | PH01000005G2890 | LOC_Os05g50930.1 | Bradi2g14790.3 | HORVU1Hr1G093180.1 | TRIUR3_33378 | Seita.3G126900.1 | Sobic.009G250400.1 | Zm00001d039194_T001 |
| orthologID_006790 | PH01000297G0990 | LOC_Os05g47690.1 | Bradi2g17560.1 | HORVU1Hr1G083110.6 | TRIUR3_03746 | Seita.3G157600.1 | Sobic.009G221800.1 | Zm00001d038910_T001 |
| orthologID_006444 | PH01002536G0030 | LOC_Os05g37950.1 | Bradi2g23570.1 | HORVU1Hr1G072990.17 | TRIUR3_28549 | Seita.3G221400.1 | Sobic.009G159200.1 | Zm00001d038348_T001 |
| orthologID_006262 | PH01000871G0240 | LOC_Os05g41620.1 | Bradi2g21136.1 | HORVU1Hr1G067370.3 | TRIUR3_23885 | Seita.3G192800.1 | Sobic.009G183500.1 | Zm00001d038578_T001 |
| orthologID_006629 | PH01001183G0400 | LOC_Os05g31970.1 | Bradi2g26817.1 | HORVU1Hr1G056820.1 | TRIUR3_14419 | Seita.3G260000.1 | Sobic.009G122400.1 | Zm00001d009975_T001 |
| orthologID_006840 | PH01001122G0420 | LOC_Os05g06760.1 | Bradi2g34760.1 | HORVU1Hr1G023670.1 | TRIUR3_33101 | Seita.7G270600.1 | Sobic.009G054700.1 | Zm00001d037553_T002 |
| orthologID_008007 | PH01000495G0780 | LOC_Os05g09370.1 | Bradi1g36960.1 | HORVU1Hr1G043150.7 | TRIUR3_25676 | Seita.3G271900.1 | Sobic.009G112300.1 | Zm00001d009903_T001 |
| orthologID_006615 | PH01000523G0300 | LOC_Os05g43380.1 | Bradi2g20347.1 | HORVU1Hr1G078000.1 | TRIUR3_10153 | Seita.3G184600.1 | Sobic.009G191600.1 | Zm00001d038660_T003 |
| orthologID_007323 | PH01001514G0250 | LOC_Os12g15460.1 | Bradi4g39360.1 | HORVU2Hr1G120430.2 | TRIUR3_00806 | Seita.3G085000.1 | Sobic.009G211100.1 | Zm00001d038825_T001 |
| orthologID_006545 | PH01000445G0850 | LOC_Os05g04160.1 | Bradi2g38481.1 | HORVU1Hr1G016190.1 | TRIUR3_15611 | Seita.3G051300.1 | Sobic.009G031300.1 | Zm00001d010942_T001 |
| orthologID_006695 | PH01000502G0600 | LOC_Os05g46200.1 | Bradi2g18440.4 | HORVU1Hr1G080090.9 | TRIUR3_26159 | Seita.3G164600.1 | Sobic.009G213100.1 | Zm00001d038847_T001 |
| orthologID_007074 | PH01001953G0400 | LOC_Os05g44160.1 | Bradi4g28905.1 | HORVU5Hr1G057770.1 | TRIUR3_09650 | Seita.3G177500.1 | Sobic.009G199700.1 | Zm00001d038719_T001 |
| orthologID_006667 | PH01074589G0010 | LOC_Os05g29020.1 | Bradi2g28000.1 | HORVU1Hr1G059170.2 | TRIUR3_18079 | Seita.3G273300.1 | Sobic.009G109600.3 | Zm00001d009890_T001 |
| orthologID_006907 | PH01001648G0280 | LOC_Os05g12380.1 | Bradi2g32220.1 | HORVU1Hr1G029950.1 | TRIUR3_31384 | Seita.3G077200.1 | Sobic.009G080600.1 | Zm00001d010443_T001 |
| orthologID_006284 | PH01009819G0010 | LOC_Os05g28860.1 | Bradi2g28110.1 | HORVU1Hr1G059450.3 | TRIUR3_07376 | Seita.3G275200.1 | Sobic.009G107300.1 | Zm00001d010021_T004 |
| orthologID_006864 | PH01000297G0870 | LOC_Os05g47660.1 | Bradi2g17600.1 | HORVU1Hr1G082900.6 | TRIUR3_05290 | Seita.3G158200.1 | Sobic.009G221500.1 | Zm00001d038908_T004 |
| orthologID_006450 | PH01000949G0420 | LOC_Os05g02010.1 | Bradi2g39150.1 | HORVU3Hr1G024040.1 | TRIUR3_00721 | Seita.3G065900.1 | Sobic.009G011600.2 | Zm00001d011067_T001 |
| orthologID_006822 | PH01000046G1280 | LOC_Os05g11980.1 | Bradi2g15960.1 | HORVU1Hr1G090050.1 | TRIUR3_18342 | Seita.3G140900.1 | Sobic.009G237400.2 | Zm00001d039062_T006 |
| orthologID_006648 | PH01002872G0090 | LOC_Os05g43850.1 | Bradi2g19960.1 | HORVU1Hr1G076900.2 | TRIUR3_14724 | Seita.3G180900.1 | Sobic.009G195900.1 | Zm00001d038691_T001 |
| orthologID_009566 | PH01002121G0220 | LOC_Os05g41880.1 | Bradi3g14127.1 | HORVU1Hr1G066830.36 | TRIUR3_00283 | Seita.3G191000.1 | Sobic.009G185700.1 | Zm00001d000329_T002 |
| orthologID_006177 | PH01004198G0120 | LOC_Os05g34700.1 | Bradi2g25070.1 | HORVU1Hr1G064020.1 | TRIUR3_27514 | Seita.3G237800.1 | Sobic.009G141300.1 | Zm00001d038211_T001 |
| orthologID_006971 | PH01000595G0160 | LOC_Os05g04700.1 | Bradi2g36770.1 | HORVU1Hr1G017700.2 | TRIUR3_27809 | Seita.3G055400.1 | Sobic.009G036100.1 | Zm00001d024778_T001 |
| orthologID_006469 | PH01001110G0420 | LOC_Os05g07640.1 | Bradi2g34480.1 | HORVU1Hr1G024670.1 | TRIUR3_12622 | Seita.3G099300.1 | Sobic.009G059400.1 | Zm00001d037509_T001 |
| orthologID_006994 | PH01007103G0040 | LOC_Os05g42000.1 | Bradi2g20820.3 | HORVU1Hr1G066450.2 | TRIUR3_19122 | Seita.3G190300.1 | Sobic.009G186600.1 | Zm00001d038599_T001 |
| orthologID_006418 | PH01002734G0140 | LOC_Os05g33710.1 | Bradi2g25610.3 | HORVU1Hr1G060850.2 | TRIUR3_28002 | Seita.3G246500.1 | Sobic.009G134500.1 | Zm00001d010305_T004 |
| orthologID_006885 | PH01000046G1000 | LOC_Os05g49250.1 | Bradi2g16110.1 | HORVU1Hr1G089550.5 | TRIUR3_01740 | Seita.3G143000.1 | Sobic.009G235300.1 | Zm00001d039050_T001 |
| orthologID_006287 | PH01000823G0670 | LOC_Os05g03750.1 | Bradi2g38237.1 | HORVU1Hr1G015000.2 | TRIUR3_32247 | Seita.3G049400.1 | Sobic.009G028400.1 | Zm00001d035456_T001 |
| orthologID_006925 | PH01002277G0170 | LOC_Os05g34210.1 | Bradi2g25350.1 | HORVU1Hr1G064830.4 | TRIUR3_33136 | Seita.3G241800.1 | Sobic.009G137800.1 | Zm00001d010271_T003 |
| orthologID_006429 | PH01000758G0390 | LOC_Os05g42280.1 | Bradi2g20600.1 | HORVU1Hr1G065950.7 | TRIUR3_09016 | Seita.3G187900.1 | Sobic.009G189000.1 | Zm00001d010702_T001 |
| orthologID_006335 | PH01000046G0020 | LOC_Os05g48800.2 | Bradi2g16670.1 | HORVU1Hr1G087100.4 | TRIUR3_13495 | Seita.3G147600.1 | Sobic.009G231300.1 | Zm00001d038999_T001 |
| orthologID_009958 | PH01000046G0590 | LOC_Os05g49070.1 | Bradi2g16396.1 | HORVU7Hr1G120950.2 | TRIUR3_19460 | Seita.3G144800.1 | Sobic.009G233900.1 | Zm00001d009582_T001 |
| orthologID_006233 | PH01000502G0390 | LOC_Os12g40440.1 | Bradi2g18550.2 | HORVU1Hr1G080320.2 | TRIUR3_01981 | Seita.3G165100.1 | Sobic.009G212600.1 | Zm00001d038841_T007 |
| orthologID_008417 | PH01000246G0580 | LOC_Os06g02028.1 | Bradi1g51790.1 | HORVU7Hr1G003100.2 | TRIUR3_06880 | Seita.4G007800.1 | Sobic.009G030400.1 | Zm00001d024715_T001 |
| orthologID_006561 | PH01000918G0720 | LOC_Os05g01750.1 | Bradi2g39361.1 | HORVU1Hr1G003530.3 | TRIUR3_11605 | Seita.3G068100.1 | Sobic.009G007800.1 | Zm00001d035698_T001 |
| orthologID_006461 | PH01000905G0580 | LOC_Os05g07860.1 | Bradi2g34310.1 | HORVU1Hr1G025340.1 | TRIUR3_30750 | Seita.3G101900.1 | Sobic.009G062400.1 | Zm00001d009800_T001 |
| orthologID_006953 | PH01000891G0430 | LOC_Os02g49190.1 | Bradi2g36760.2 | HORVU1Hr1G017710.2 | TRIUR3_24457 | Seita.3G055500.1 | Sobic.009G036200.1 | Zm00001d010910_T002 |
| orthologID_006918 | PH01001489G0180 | LOC_Os05g41800.1 | Bradi2g21030.1 | HORVU1Hr1G067020.22 | TRIUR3_26376 | Seita.3G191500.1 | Sobic.009G184800.1 | Zm00001d010679_T001 |
| orthologID_006686 | PH01001516G0010 | LOC_Os05g03540.1 | Bradi2g38140.1 | HORVU1Hr1G014080.2 | TRIUR3_16669 | Seita.3G048000.1 | Sobic.009G027100.2 | Zm00001d035473_T001 |

| | | | | | | | | |
|---|---|---|---|---|---|---|---|---|
| orthologID_006692 | PH01000298G0170 | LOC_Os05g49100.1 | Bradi2g16357.2 | HORVU1Hr1G088300.1 | TRIUR3_33327 | Seita.3G145000.1 | Sobic.009G234100.1 | Zm00001d039032_T001 |
| orthologID_008594 | PH01003078G0220 | LOC_Os06g01180.1 | Bradi1g52420.1 | HORVU7Hr1G000870.6 | TRIUR3_22231 | Seita.3G152000.1 | Sobic.009G226700.1 | Zm00001d038960_T001 |
| orthologID_006845 | PH01000046G1830 | LOC_Os05g49750.1 | Bradi2g15820.2 | HORVU1Hr1G090410.1 | TRIUR3_11544 | Seita.3G138900.1 | Sobic.009G238800.1 | Zm00001d009627_T001 |
| orthologID_006176 | PH01001592G0130 | LOC_Os05g05820.1 | Bradi2g35700.1 | HORVU1Hr1G021620.2 | TRIUR3_16001 | Seita.3G037000.1 | Sobic.009G045600.1 | Zm00001d035783_T001 |
| orthologID_006553 | PH01000685G0710 | LOC_Os05g06420.1 | Bradi2g35030.1 | HORVU1Hr1G022800.1 | TRIUR3_17796 | Seita.7G268500.1 | Sobic.009G051500.1 | Zm00001d009787_T001 |
| orthologID_006471 | PH01003163G0210 | LOC_Os05g25070.1 | Bradi2g31230.2 | HORVU1Hr1G030630.6 | TRIUR3_24382 | Seita.3G289800.1 | Sobic.009G095100.2 | Zm00001d010455_T013 |
| orthologID_006453 | PH01000005G2880 | LOC_Os05g50950.1 | Bradi2g14750.1 | HORVU1Hr1G093220.1 | TRIUR3_11139 | Seita.3G126600.1 | Sobic.009G250700.1 | Zm00001d039193_T001 |
| orthologID_006799 | PH01000011G1530 | LOC_Os05g33080.1 | Bradi2g26063.1 | HORVU1Hr1G062130.3 | TRIUR3_27493 | Seita.3G251200.1 | Sobic.009G129700.1 | Zm00001d038135_T001 |
| orthologID_006162 | PH01000445G0230 | LOC_Os05g04620.1 | Bradi2g36890.1 | HORVU1Hr1G017480.1 | TRIUR3_25925 | Seita.3G054300.1 | Sobic.009G034600.1 | Zm00001d024765_T001 |
| orthologID_006938 | PH01002254G0140 | LOC_Os05g26954.1 | Bradi2g31070.1 | HORVU1Hr1G079240.6 | TRIUR3_03527 | Seita.3G284400.1 | Sobic.009G097200.1 | Zm00001d010395_T001 |
| orthologID_006295 | PH01000878G0460 | LOC_Os05g11010.1 | Bradi2g33100.1 | HORVU1Hr1G029180.2 | TRIUR3_26644 | Seita.3G113000.1 | Sobic.009G073300.1 | Zm00001d037751_T001 |
| orthologID_006935 | PH01001141G0550 | LOC_Os05g42180.1 | Bradi2g20720.3 | HORVU1Hr1G066250.1 | TRIUR3_23565 | Seita.3G189000.1 | Sobic.009G187900.1 | Zm00001d038612_T001 |
| orthologID_006923 | PH01000949G0440 | LOC_Os05g01994.1 | Bradi2g39170.2 | HORVU1Hr1G004490.1 | TRIUR3_00723 | Seita.3G066100.1 | Sobic.009G011400.1 | Zm00001d035669_T002 |
| orthologID_006881 | PH01000980G0280 | LOC_Os05g32590.1 | Bradi2g26520.3 | HORVU1Hr1G063010.17 | TRIUR3_33022 | Seita.3G254500.1 | Sobic.009G126500.1 | Zm00001d038108_T001 |
| orthologID_006180 | PH01002060G0210 | LOC_Os05g28950.1 | Bradi2g28087.1 | HORVU1Hr1G059360.1 | TRIUR3_35030 | Seita.3G274700.1 | Sobic.009G108400.1 | Zm00001d037967_T002 |
| orthologID_006524 | PH01006486G0060 | LOC_Os05g43040.1 | Bradi2g20540.1 | HORVU1Hr1G075210.1 | TRIUR3_12933 | Seita.3G186600.1 | Sobic.009G190000.1 | Zm00001d038642_T001 |
| orthologID_007007 | PH01000298G0530 | LOC_Os05g48950.2 | Bradi2g16530.2 | HORVU1Hr1G087620.1 | TRIUR3_03633 | Seita.3G146300.1 | Sobic.009G232500.1 | Zm00001d039013_T001 |
| orthologID_006153 | PH01001947G0400 | LOC_Os05g43360.1 | Bradi2g20370.3 | HORVU1Hr1G077910.2 | TRIUR3_10155 | Seita.3G184800.1 | Sobic.009G191400.1 | Zm00001d038658_T001 |
| orthologID_006951 | PH01001217G0300 | LOC_Os05g32220.1 | Bradi2g26940.1 | HORVU1Hr1G056510.2 | TRIUR3_07435 | Seita.3G258100.1 | Sobic.009G123900.1 | Zm00001d038084_T001 |
| orthologID_006379 | PH01000523G0440 | LOC_Os05g43470.1 | Bradi2g20290.1 | HORVU1Hr1G077770.9 | TRIUR3_01109 | Seita.3G184000.1 | Sobic.009G192200.1 | Zm00001d000311_T003 |
| orthologID_006855 | PH01000046G0990 | LOC_Os05g49240.1 | Bradi2g16120.1 | HORVU1Hr1G089490.2 | TRIUR3_26254 | Seita.3G143100.1 | Sobic.009G235500.1 | Zm00001d009599_T001 |
| orthologID_006706 | PH01000258G1050 | LOC_Os05g05130.1 | Bradi2g36490.7 | HORVU1Hr1G018700.1 | TRIUR3_12594 | Seita.3G057600.1 | Sobic.009G038600.1 | Zm00001d024803_T001 |
| orthologID_006831 | PH01001725G0060 | LOC_Os05g11250.1 | Bradi2g32950.2 | HORVU1Hr1G029280.1 | TRIUR3_28220 | Seita.3G072600.1 | Sobic.009G075100.1 | Zm00001d010007_T001 |
| orthologID_006587 | PH01003188G0110 | LOC_Os05g33220.1 | Bradi2g25960.1 | HORVU1Hr1G061980.2 | TRIUR3_17420 | Seita.3G250100.1 | Sobic.009G130300.1 | Zm00001d009932_T001 |
| orthologID_006278 | PH01001110G0360 | LOC_Os05g07270.1 | Bradi2g34520.1 | HORVU1Hr1G024490.1 | TRIUR3_18328 | Seita.7G273900.1 | Sobic.009G058600.2 | Zm00001d009823_T001 |
| orthologID_006302 | PH01003109G0220 | LOC_Os01g01130.1 | Bradi2g03710.1 | HORVU3Hr1G015060.4 | TRIUR3_16757 | Seita.5G065600.1 | Sobic.009G033900.1 | Zm00001d010915_T001 |
| orthologID_006410 | PH01001601G0380 | LOC_Os05g50710.1 | Bradi2g15410.1 | HORVU1Hr1G092670.3 | TRIUR3_08299 | Seita.3G130700.1 | Sobic.009G247900.1 | Zm00001d009700_T001 |
| orthologID_006622 | PH01000298G0270 | LOC_Os05g49060.1 | Bradi2g16402.1 | HORVU1Hr1G088270.4 | TRIUR3_35134 | Seita.3G144600.1 | Sobic.009G233800.1 | Zm00001d039029_T001 |
| orthologID_006280 | PH01001734G0370 | LOC_Os05g09410.1 | Bradi2g33460.1 | HORVU1Hr1G027980.33 | TRIUR3_22886 | Seita.3G109200.1 | Sobic.009G069500.2 | Zm00001d037694_T001 |
| orthologID_006521 | PH01000042G2270 | LOC_Os05g05220.1 | Bradi2g36460.1 | HORVU1Hr1G018750.3 | TRIUR3_00872 | Seita.3G058800.1 | Sobic.009G040500.1 | Zm00001d024815_T005 |
| orthologID_006214 | PH01000423G0790 | LOC_Os05g43582.1 | Bradi2g20250.1 | HORVU1Hr1G078110.8 | TRIUR3_01020 | Seita.3G183500.1 | Sobic.009G192800.1 | Zm00001d038668_T001 |
| orthologID_008628 | PH01001782G0440 | LOC_Os05g04120.1 | Bradi1g67620.2 | HORVU4Hr1G062070.14 | TRIUR3_22503 | Seita.3G051000.1 | Sobic.009G031000.1 | Zm00001d010957_T001 |
| orthologID_006883 | PH01000157G1310 | LOC_Os05g47880.1 | Bradi2g17360.1 | HORVU1Hr1G083700.1 | TRIUR3_33292 | Seita.3G155600.1 | Sobic.009G223400.1 | Zm00001d038924_T001 |
| orthologID_006466 | PH01001141G0400 | LOC_Os05g42100.1 | Bradi2g20790.1 | HORVU1Hr1G066510.7 | TRIUR3_09097 | Seita.3G189700.1 | Sobic.009G187100.1 | Zm00001d038606_T001 |
| orthologID_006789 | PH01000766G0300 | LOC_Os05g33890.1 | Bradi2g25500.1 | HORVU1Hr1G063330.5 | TRIUR3_20200 | Seita.3G244600.1 | Sobic.009G136000.1 | Zm00001d038174_T002 |
| orthologID_006900 | PH01004121G0170 | LOC_Os05g02640.1 | Bradi2g37517.1 | HORVU1Hr1G007640.1 | TRIUR3_32155 | Seita.3G041100.1 | Sobic.009G020100.1 | Zm00001d035579_T007 |
| orthologID_006270 | PH01000523G0430 | LOC_Os05g43460.1 | Bradi2g20302.1 | HORVU1Hr1G077760.1 | TRIUR3_01108 | Seita.3G184100.1 | Sobic.009G192100.1 | Zm00001d000310_T001 |
| orthologID_006123 | PH01000206G1340 | LOC_Os05g39230.1 | Bradi2g22680.2 | HORVU1Hr1G070740.5 | TRIUR3_23522 | Seita.3G210000.1 | Sobic.009G168600.2 | Zm00001d010585_T002 |
| orthologID_006426 | PH01000577G0020 | LOC_Os05g13370.1 | Bradi2g31930.1 | HORVU1Hr1G030040.2 | TRIUR3_29335 | Seita.3G079400.1 | Sobic.009G083000.1 | Zm00001d010424_T002 |
| orthologID_006816 | PH01000556G0040 | LOC_Os05g41030.1 | Bradi2g21830.1 | HORVU1Hr1G068830.1 | TRIUR3_11641 | Seita.3G198600.1 | Sobic.009G178500.1 | Zm00001d038532_T001 |
| orthologID_006598 | PH01000414G0300 | LOC_Os05g32440.1 | Bradi2g26620.1 | HORVU1Hr1G063150.1 | TRIUR3_10525 | Seita.3G255700.1 | Sobic.009G125800.1 | Zm00001d010465_T001 |
| orthologID_006244 | PH01000528G0920 | LOC_Os05g40280.1 | Bradi2g22130.3 | HORVU1Hr1G069420.1 | TRIUR3_13028 | Seita.3G202300.1 | Sobic.009G175800.1 | Zm00001d038504_T001 |
| orthologID_006279 | PH01001664G0370 | LOC_Os05g35140.1 | Bradi2g24850.1 | HORVU1Hr1G065100.2 | TRIUR3_19593 | Seita.3G235900.1 | Sobic.009G143500.1 | Zm00001d038226_T001 |
| orthologID_006462 | PH01001143G0650 | LOC_Os05g30130.1 | Bradi2g27830.3 | HORVU1Hr1G058700.1 | TRIUR3_24067 | Seita.3G271000.1 | Sobic.009G113000.1 | Zm00001d009912_T001 |
| orthologID_006915 | PH01000078G1720 | LOC_Os05g01250.1 | Bradi2g39980.1 | HORVU0Hr1G000030.13 | TRIUR3_21486 | Seita.3G115200.1 | Sobic.009G001400.1 | Zm00001d035762_T001 |
| orthologID_006543 | PH01002248G0180 | LOC_Os05g33550.1 | Bradi2g25777.1 | HORVU1Hr1G061190.1 | TRIUR3_30954 | Seita.3G248100.1 | Sobic.009G132800.3 | Zm00001d010322_T001 |
| orthologID_006541 | PH01000297G0090 | LOC_Os05g46490.2 | Bradi2g18080.3 | HORVU1Hr1G079200.1 | TRIUR3_03528 | Seita.3G162300.1 | Sobic.009G216000.1 | Zm00001d038871_T001 |
| orthologID_006272 | PH01001277G0220 | LOC_Os05g06510.1 | Bradi2g34900.1 | HORVU1Hr1G023320.3 | TRIUR3_10625 | Seita.7G269200.1 | Sobic.009G052700.1 | Zm00001d011124_T002 |
| orthologID_006380 | PH01000242G0910 | LOC_Os05g49420.1 | Bradi2g15940.1 | HORVU1Hr1G090030.11 | TRIUR3_18341 | Seita.3G140700.1 | Sobic.009G237600.1 | Zm00001d039065_T003 |
| orthologID_006963 | PH01002139G0100 | LOC_Os05g02200.1 | Bradi2g37150.1 | HORVU5Hr1G102900.1 | TRIUR3_31305 | Seita.3G063900.1 | Sobic.009G014700.1 | Zm00001d035646_T001 |

| | | | | | | | | |
|---|---|---|---|---|---|---|---|---|
| orthologID_006842 | PH01003614G0140 | LOC_Os05g10840.2 | Bradi2g33150.1 | HORVU1Hr1G028900.11 | TRIUR3_31101 | Seita.3G112600.1 | Sobic.009G072900.2 | Zm00001d037759_T002 |
| orthologID_006673 | PH01000974G0600 | LOC_Os01g62350.1 | Bradi2g23260.3 | HORVU1Hr1G072270.1 | TRIUR3_18142 | Seita.3G217000.1 | Sobic.009G163800.1 | Zm00001d010528_T003 |
| orthologID_006146 | PH01002248G0150 | LOC_Os05g33520.1 | Bradi2g25800.1 | HORVU1Hr1G061230.3 | TRIUR3_11736 | Seita.3G248300.1 | Sobic.009G132000.2 | Zm00001d038161_T001 |
| orthologID_006128 | PH01000348G0830 | LOC_Os05g38140.1 | Bradi2g23520.1 | HORVU1Hr1G072810.2 | TRIUR3_14889 | Seita.3G220600.1 | Sobic.009G160000.1 | Zm00001d038357_T001 |
| orthologID_006327 | PH01000271G1030 | LOC_Os05g45760.1 | Bradi2g18760.1 | HORVU1Hr1G080900.3 | TRIUR3_11937 | Seita.3G166900.1 | Sobic.009G210100.1 | Zm00001d038733_T001 |
| orthologID_006349 | PH01003060G0240 | LOC_Os05g38040.1 | Bradi2g23540.2 | HORVU1Hr1G072890.7 | TRIUR3_22405 | Seita.3G221000.1 | Sobic.009G159700.1 | Zm00001d038352_T001 |
| orthologID_006982 | PH01000423G1050 | LOC_Os05g43440.1 | Bradi2g20320.2 | HORVU1Hr1G077860.2 | TRIUR3_01103 | Seita.3G184300.1 | Sobic.009G191800.4 | Zm00001d000241_T002 |
| orthologID_006956 | PH01000298G1040 | LOC_Os05g48750.1 | Bradi2g16710.2 | HORVU1Hr1G086980.13 | TRIUR3_15578 | Seita.3G147800.1 | Sobic.009G231100.1 | Zm00001d009554_T001 |
| orthologID_009846 | PH01000530G0880 | LOC_Os05g51620.1 | Bradi3g46247.2 | HORVU3Hr1G030250.1 | TRIUR3_28115 | Seita.3G120400.1 | Sobic.009G257900.1 | Zm00001d009765_T008 |
| orthologID_006712 | PH01002864G0120 | LOC_Os05g31490.1 | Bradi2g27060.1 | HORVU1Hr1G057490.4 | TRIUR3_12926 | Seita.3G261700.1 | Sobic.009G120500.1 | Zm00001d038059_T001 |
| orthologID_006868 | PH01000523G0190 | LOC_Os05g43310.1 | Bradi2g20380.1 | HORVU1Hr1G078140.4 | TRIUR3_01017 | Seita.3G184900.1 | Sobic.009G191300.1 | Zm00001d010715_T001 |
| orthologID_006292 | PH01000206G1100 | LOC_Os05g39410.1 | Bradi2g22610.1 | HORVU1Hr1G070580.5 | TRIUR3_05903 | Seita.3G209100.1 | Sobic.009G169400.1 | Zm00001d038432_T006 |
| orthologID_007882 | PH01002189G0040 | LOC_Os07g05580.1 | Bradi1g53740.1 | HORVU2Hr1G053290.6 | TRIUR3_27985 | Seita.1G205300.1 | Sobic.009G242400.1 | Zm00001d039106_T003 |
| orthologID_007245 | PH01000005G2030 | LOC_Os11g34360.1 | Bradi4g16790.1 | HORVU4Hr1G038270.2 | TRIUR3_33295 | Seita.8G145700.1 | Sobic.005G144201.1 | Zm00001d007441_T001 |
| orthologID_007099 | PH01001519G0070 | LOC_Os11g47690.1 | Bradi4g09380.2 | HORVU4Hr1G028140.1 | TRIUR3_18867 | Seita.8G248200.1 | Sobic.005G226600.1 | Zm00001d004769_T001 |
| orthologID_007309 | PH01001759G0240 | LOC_Os11g05970.1 | Bradi4g24990.1 | HORVU4Hr1G020790.1 | TRIUR3_12755 | Seita.8G040200.1 | Sobic.005G044600.1 | Zm00001d052784_T001 |
| orthologID_007395 | PH01002104G0150 | LOC_Os11g10130.1 | Bradi4g22730.1 | HORVU4Hr1G017040.2 | TRIUR3_35269 | Seita.8G101000.1 | Sobic.005G104800.1 | Zm00001d049079_T001 |
| orthologID_010003 | PH01000005G1450 | LOC_Os11g35320.1 | Bradi4g16327.1 | HORVU4Hr1G007670.3 | TRIUR3_12640 | Seita.8G152400.1 | Sobic.005G155300.1 | Zm00001d007479_T001 |
| orthologID_007481 | PH01002512G0320 | LOC_Os11g06040.1 | Bradi4g25047.2 | HORVU4Hr1G020680.1 | TRIUR3_29271 | Seita.8G041000.1 | Sobic.005G045300.2 | Zm00001d053155_T002 |
| orthologID_009114 | PH01004915G0060 | LOC_Os11g32320.1 | Bradi3g36577.1 | HORVU3Hr1G107320.1 | TRIUR3_02659 | Seita.8G138600.1 | Sobic.005G133700.1 | Zm00001d052682_T001 |
| orthologID_007426 | PH01001659G0340 | LOC_Os11g39020.1 | Bradi4g14020.1 | HORVU0Hr1G008550.1 | TRIUR3_27993 | Seita.8G181700.1 | Sobic.005G179300.1 | Zm00001d007597_T001 |
| orthologID_009263 | PH01000373G0820 | LOC_Os10g06610.1 | Bradi3g20952.1 | HORVU7Hr1G031100.1 | TRIUR3_26754 | Seita.9G285600.1 | Sobic.005G010900.2 | Zm00001d030340_T037 |
| orthologID_007157 | PH01006272G0040 | LOC_Os11g05760.1 | Bradi4g25160.1 | HORVU4Hr1G021620.2 | TRIUR3_27569 | Seita.8G037000.1 | Sobic.005G041500.1 | Zm00001d052592_T001 |
| orthologID_007535 | PH01000005G0960 | LOC_Os11g36430.3 | Bradi4g16000.2 | HORVU4Hr1G034400.7 | TRIUR3_28356 | Seita.8G158300.1 | Sobic.005G160400.1 | Zm00001d052930_T003 |
| orthologID_007042 | PH01002767G0160 | LOC_Os11g32470.1 | Bradi4g17287.4 | HORVU5Hr1G007320.2 | TRIUR3_12224 | Seita.8G138700.1 | Sobic.005G133800.1 | Zm00001d007388_T002 |
| orthologID_009858 | PH01001230G0370 | LOC_Os11g29370.1 | Bradi3g57750.4 | HORVU5Hr1G026730.1 | TRIUR3_32573 | Seita.8G121300.1 | Sobic.005G110541.1 | Zm00001d049191_T001 |
| orthologID_007059 | PH01001659G0170 | LOC_Os11g39360.2 | Bradi4g14010.3 | HORVU7Hr1G028150.4 | TRIUR3_16737 | Seita.8G185900.1 | Sobic.005G182300.2 | Zm00001d048880_T001 |
| orthologID_007295 | PH01000631G0590 | LOC_Os11g32480.1 | Bradi4g17280.1 | HORVU5Hr1G106840.1 | TRIUR3_02167 | Seita.8G138900.1 | Sobic.005G133900.1 | Zm00001d052678_T001 |
| orthologID_007267 | PH01000315G0430 | LOC_Os11g11050.1 | Bradi4g22150.1 | HORVU4Hr1G016410.2 | TRIUR3_18098 | Seita.8G094900.1 | Sobic.005G100800.1 | Zm00001d049042_T001 |
| orthologID_007490 | PH01000474G0310 | LOC_Os12g05000.1 | Bradi4g42940.1 | HORVU5Hr1G040870.2 | TRIUR3_29581 | Seita.8G028100.1 | Sobic.005G033200.1 | Zm00001d052488_T001 |
| orthologID_007478 | PH01004116G0020 | LOC_Os11g10100.1 | Bradi4g22760.1 | HORVU4Hr1G017100.25 | TRIUR3_33594 | Seita.8G101200.1 | Sobic.005G105000.1 | Zm00001d049080_T001 |
| orthologID_007381 | PH01000051G1240 | LOC_Os11g29630.1 | Bradi4g19020.1 | HORVU3Hr1G026350.1 | TRIUR3_26125 | Seita.8G122300.1 | Sobic.005G116300.1 | Zm00001d049178_T001 |
| orthologID_007095 | PH01002736G0110 | LOC_Os11g40080.1 | Bradi4g13710.1 | HORVU4Hr1G035360.12 | TRIUR3_19196 | Seita.8G193100.1 | Sobic.005G187300.1 | Zm00001d048835_T004 |
| orthologID_007427 | PH01000599G0370 | LOC_Os11g14070.1 | Bradi4g21025.1 | HORVU4Hr1G015040.1 | TRIUR3_24391 | Seita.8G078800.1 | Sobic.005G085800.1 | Zm00001d049011_T001 |
| orthologID_007409 | PH01002777G0100 | LOC_Os11g05614.1 | Bradi4g42180.1 | HORVU5Hr1G040700.6 | TRIUR3_33275 | Seita.8G036100.1 | Sobic.005G041000.1 | Zm00001d052564_T001 |
| orthologID_008499 | PH01000294G0080 | LOC_Os03g25380.1 | Bradi1g61520.3 | HORVU6Hr1G005380.1 | TRIUR3_30319 | Seita.9G392300.1 | Sobic.005G017100.1 | Zm00001d052377_T001 |
| orthologID_007164 | PH01002964G0120 | LOC_Os11g25780.1 | Bradi4g19634.1 | HORVU6Hr1G055230.2 | TRIUR3_21026 | Seita.8G111900.1 | Sobic.005G111200.1 | Zm00001d049157_T001 |
| orthologID_007313 | PH01000367G0850 | LOC_Os11g38260.1 | Bradi4g14630.1 | HORVU0Hr1G039090.1 | TRIUR3_03928 | Seita.8G173800.1 | Sobic.005G172400.1 | Zm00001d007581_T001 |
| orthologID_007114 | PH01004058G0120 | LOC_Os08g28180.1 | Bradi4g18590.3 | HORVU3Hr1G089060.1 | TRIUR3_17217 | Seita.4G100700.1 | Sobic.005G122700.1 | Zm00001d018219_T001 |
| orthologID_007111 | PH01003533G0090 | LOC_Os11g07970.1 | Bradi4g23700.1 | HORVU4Hr1G018020.2 | TRIUR3_30228 | Seita.8G059600.1 | Sobic.005G063700.3 | Zm00001d004739_T001 |
| orthologID_007259 | PH01003363G0150 | LOC_Os12g05550.1 | Bradi4g42490.1 | HORVU5Hr1G042220.1 | TRIUR3_32937 | Seita.8G033700.1 | Sobic.005G037900.1 | Zm00001d052533_T001 |
| orthologID_007435 | PH01001205G0030 | LOC_Os11g13840.1 | Bradi4g21265.1 | HORVU4Hr1G015350.1 | TRIUR3_32331 | Seita.8G081600.1 | Sobic.005G087600.2 | Zm00001d048991_T001 |
| orthologID_007373 | PH01002368G0210 | LOC_Os11g29910.1 | Bradi4g18980.1 | HORVU4Hr1G061950.1 | TRIUR3_15882 | Seita.8G123400.1 | Sobic.005G118300.1 | Zm00001d049170_T001 |
| orthologID_007525 | PH01001088G0550 | LOC_Os11g40210.1 | Bradi4g13773.1 | HORVU2Hr1G084090.2 | TRIUR3_29892 | Seita.8G195600.1 | Sobic.005G188400.1 | Zm00001d048829_T001 |
| orthologID_007532 | PH01001994G0120 | LOC_Os11g14190.1 | Bradi4g20940.1 | HORVU4Hr1G014900.1 | TRIUR3_08813 | Seita.8G077300.1 | Sobic.005G084600.1 | Zm00001d007280_T001 |
| orthologID_007021 | PH01004818G0100 | LOC_Os11g45710.7 | Bradi4g09920.1 | HORVU4Hr1G028910.6 | TRIUR3_28967 | Seita.8G237500.1 | Sobic.005G217000.5 | Zm00001d007716_T002 |
| orthologID_007445 | PH01000078G0200 | LOC_Os11g41140.1 | Bradi4g12920.1 | HORVU4Hr1G030950.7 | TRIUR3_29537 | Seita.8G202000.1 | Sobic.005G194700.2 | Zm00001d048796_T003 |
| orthologID_006595 | PH01000649G0080 | LOC_Os10g14180.8 | Bradi2g38870.1 | HORVU1Hr1G004920.4 | TRIUR3_06163 | Seita.1G154800.1 | Sobic.005G072400.1 | Zm00001d053144_T001 |
| orthologID_007336 | PH01001994G0160 | LOC_Os11g14300.1 | Bradi4g20900.1 | HORVU4Hr1G014800.1 | TRIUR3_29667 | Seita.8G077100.1 | Sobic.005G084400.1 | Zm00001d007279_T001 |
| orthologID_007357 | PH01000315G0410 | LOC_Os11g11020.1 | Bradi4g22160.1 | HORVU4Hr1G016440.1 | TRIUR3_26753 | Seita.8G095000.1 | Sobic.005G100900.1 | Zm00001d049043_T001 |

| | | | | | | | | |
|---|---|---|---|---|---|---|---|---|
| orthologID_007497 | PH01000465G0570 | LOC_Os11g43610.1 | Bradi4g11137.1 | HORVU6Hr1G080830.6 | TRIUR3_25645 | Seita.8G220200.1 | Sobic.005G204400.3 | Zm00001d007682_T002 |
| orthologID_007252 | PH01004098G0040 | LOC_Os11g38030.1 | Bradi4g14840.2 | HORVU4Hr1G029570.2 | TRIUR3_33146 | Seita.8G170600.1 | Sobic.005G169900.1 | Zm00001d007573_T001 |
| orthologID_007364 | PH01001484G0220 | LOC_Os04g14450.1 | Bradi4g15307.3 | HORVU3Hr1G019900.13 | TRIUR3_16796 | Seita.8G163400.1 | Sobic.005G020000.7 | Zm00001d052396_T001 |
| orthologID_007051 | PH01001230G0270 | LOC_Os11g29360.1 | Bradi4g19150.1 | HORVU1Hr1G087370.1 | TRIUR3_14245 | Seita.8G121200.1 | Sobic.005G110550.1 | Zm00001d049196_T001 |
| orthologID_007503 | PH01000005G2210 | LOC_Os11g34200.1 | Bradi4g16840.1 | HORVU4Hr1G034360.6 | TRIUR3_34926 | Seita.8G144800.1 | Sobic.005G143400.1 | Zm00001d052621_T002 |
| orthologID_007224 | PH01002392G0070 | LOC_Os11g09350.1 | Bradi4g23107.1 | HORVU4Hr1G043990.3 | TRIUR3_21358 | Seita.8G070400.1 | Sobic.005G076600.1 | Zm00001d049107_T001 |
| orthologID_007150 | PH01004081G0040 | LOC_Os11g32880.1 | Bradi4g17160.1 | HORVU4Hr1G040140.10 | TRIUR3_17794 | Seita.8G140800.1 | Sobic.005G138600.1 | Zm00001d007417_T001 |
| orthologID_007356 | PH01000637G0310 | LOC_Os11g36450.1 | Bradi4g16020.3 | HORVU7Hr1G100800.2 | TRIUR3_19243 | Seita.8G158500.1 | Sobic.005G160600.1 | Zm00001d052933_T001 |
| orthologID_009597 | PH01003340G0240 | LOC_Os02g17870.1 | Bradi3g39560.2 | HORVU3Hr1G096160.3 | TRIUR3_14354 | Seita.1G125300.1 | Sobic.005G033300.1 | Zm00001d052489_T001 |
| orthologID_007438 | PH01004446G0090 | LOC_Os11g13980.1 | Bradi4g21070.2 | HORVU4Hr1G015170.1 | TRIUR3_23889 | Seita.8G079900.1 | Sobic.005G086400.1 | Zm00001d007271_T001 |
| orthologID_007485 | PH01005812G0020 | LOC_Os11g47840.1 | Bradi4g09210.1 | HORVU4Hr1G027850.1 | TRIUR3_26991 | Seita.8G249900.1 | Sobic.005G229166.1 | Zm00001d048606_T001 |
| orthologID_007032 | PH01005178G0030 | LOC_Os11g37670.1 | Bradi4g14970.1 | HORVU7Hr1G104830.4 | TRIUR3_35151 | Seita.8G168600.1 | Sobic.005G168400.1 | Zm00001d052981_T001 |
| orthologID_007422 | PH01000466G0540 | LOC_Os11g43510.1 | Bradi4g11330.1 | HORVU4Hr1G033860.1 | TRIUR3_24906 | Seita.8G219600.1 | Sobic.005G203800.2 | Zm00001d048721_T001 |
| orthologID_007199 | PH01000368G0980 | LOC_Os11g07450.1 | Bradi4g24480.1 | HORVU4Hr1G019360.6 | TRIUR3_32207 | Seita.8G053400.1 | Sobic.005G058900.1 | Zm00001d053018_T004 |
| orthologID_007056 | PH01003323G0170 | LOC_Os11g29850.1 | Bradi4g19047.7 | HORVU3Hr1G089880.8 | TRIUR3_30766 | Seita.8G123000.1 | Sobic.005G117300.1 | Zm00001d007328_T003 |
| orthologID_007554 | PH01000005G1820 | LOC_Os11g34870.1 | Bradi4g16510.1 | HORVU4Hr1G044460.4 | TRIUR3_30104 | Seita.8G148600.1 | Sobic.005G149700.1 | Zm00001d053102_T001 |
| orthologID_007502 | PH01001822G0070 | LOC_Os11g34140.1 | Bradi4g16870.2 | HORVU5Hr1G063470.1 | TRIUR3_18155 | Seita.8G144500.1 | Sobic.005G143100.1 | Zm00001d052618_T001 |
| orthologID_007261 | PH01003993G0100 | LOC_Os11g31940.1 | Bradi4g18187.1 | HORVU3Hr1G024970.12 | TRIUR3_08342 | Seita.8G133300.1 | Sobic.005G127200.1 | Zm00001d007351_T001 |
| orthologID_007257 | PH01000130G0370 | LOC_Os12g02520.1 | Bradi4g44336.1 | HORVU5Hr1G046370.1 | TRIUR3_18973 | Seita.7G302800.1 | Sobic.005G014600.1 | Zm00001d005052_T002 |
| orthologID_007302 | PH01001348G0440 | LOC_Os11g47820.1 | Bradi4g09230.1 | HORVU4Hr1G027740.1 | TRIUR3_20849 | Seita.8G249700.1 | Sobic.005G228900.1 | Zm00001d004768_T001 |
| orthologID_007183 | PH01000205G0750 | LOC_Os11g20360.2 | Bradi4g20000.1 | HORVU4Hr1G012480.3 | TRIUR3_22893 | Seita.8G109000.1 | Sobic.005G092700.1 | Zm00001d048958_T003 |
| orthologID_007353 | PH01000957G0440 | LOC_Os11g05260.1 | Bradi4g39570.4 | HORVU4Hr1G025330.3 | TRIUR3_07409 | Seita.7G320900.1 | Sobic.005G036100.1 | Zm00001d052519_T004 |
| orthologID_007410 | PH01002934G0030 | LOC_Os11g37070.1 | Bradi4g15360.1 | HORVU4Hr1G034110.2 | TRIUR3_14763 | Seita.8G161600.1 | Sobic.005G164500.1 | Zm00001d052951_T001 |
| orthologID_007484 | PH01000631G0540 | LOC_Os11g32500.1 | Bradi4g17260.1 | HORVU4Hr1G043650.2 | TRIUR3_19235 | Seita.8G139100.1 | Sobic.005G134100.1 | Zm00001d007394_T001 |
| orthologID_008494 | PH01000864G0660 | LOC_Os11g29230.1 | Bradi1g19790.1 | HORVU3Hr1G005560.1 | TRIUR3_02228 | Seita.9G202900.1 | Sobic.005G110400.1 | Zm00001d049203_T001 |
| orthologID_007082 | PH01002178G0160 | LOC_Os11g43860.1 | Bradi4g11030.1 | HORVU4Hr1G033020.4 | TRIUR3_22032 | Seita.8G222300.1 | Sobic.005G206500.1 | Zm00001d048692_T003 |
| orthologID_007310 | PH01000315G0720 | LOC_Os11g11100.1 | Bradi4g22130.3 | HORVU4Hr1G016340.4 | TRIUR3_26059 | Seita.8G094700.1 | Sobic.005G100600.2 | Zm00001d049135_T001 |
| orthologID_007461 | PH01003847G0060 | LOC_Os11g31880.1 | Bradi4g17940.1 | HORVU4Hr1G037160.3 | TRIUR3_22379 | Seita.8G133900.1 | Sobic.005G129201.1 | Zm00001d049270_T002 |
| orthologID_007516 | PH01001088G0220 | LOC_Os11g39540.1 | Bradi4g13970.3 | HORVU4Hr1G031620.1 | TRIUR3_34887 | Seita.8G187000.1 | Sobic.005G183200.1 | Zm00001d048868_T002 |
| orthologID_007399 | PH01000005G1170 | LOC_Os11g36060.1 | Bradi4g16160.1 | HORVU2Hr1G019570.1 | TRIUR3_29372 | Seita.8G156800.1 | Sobic.005G157100.1 | Zm00001d052921_T002 |
| orthologID_006448 | PH01000232G0420 | LOC_Os01g74610.1 | Bradi2g62760.1 | HORVU5Hr1G123700.7 | TRIUR3_08679 | Seita.8G025600.1 | Sobic.005G028600.1 | Zm00001d004979_T003 |
| orthologID_007244 | PH01001798G0230 | LOC_Os11g48040.1 | Bradi4g09060.3 | HORVU4Hr1G027150.1 | TRIUR3_19305 | Seita.8G252200.1 | Sobic.005G231900.2 | Zm00001d048583_T001 |
| orthologID_007131 | PH01005277G0080 | LOC_Os11g44930.1 | Bradi4g10130.1 | HORVU4Hr1G032070.2 | TRIUR3_30164 | Seita.8G229600.1 | Sobic.005G210700.1 | Zm00001d048670_T001 |
| orthologID_009917 | PH01001701G0130 | LOC_Os05g05600.3 | Bradi3g10770.4 | HORVU6Hr1G050440.2 | TRIUR3_30205 | Seita.8G238500.1 | Sobic.005G206300.2 | Zm00001d048693_T001 |
| orthologID_007038 | PH01000902G0420 | LOC_Os11g42450.1 | Bradi4g11651.1 | HORVU4Hr1G031780.1 | TRIUR3_29651 | Seita.8G213600.1 | Sobic.005G199700.1 | Zm00001d048740_T001 |
| orthologID_007119 | PH01002517G0100 | LOC_Os11g38780.1 | Bradi4g14270.1 | HORVU1Hr1G086900.1 | TRIUR3_24401 | Seita.8G176000.1 | Sobic.005G175800.1 | Zm00001d048913_T001 |
| orthologID_007028 | PH01150188G0010 | LOC_Os11g35870.1 | Bradi4g16230.1 | HORVU4Hr1G036670.3 | TRIUR3_26469 | Seita.8G154900.1 | Sobic.005G158300.1 | Zm00001d052911_T002 |
| orthologID_007444 | PH01002557G0050 | LOC_Os11g40610.5 | Bradi4g13227.2 | HORVU4Hr1G030640.9 | TRIUR3_33924 | Seita.8G200900.1 | Sobic.005G194200.1 | Zm00001d048801_T001 |
| orthologID_007262 | PH01000005G1390 | LOC_Os11g35390.1 | Bradi4g16290.1 | HORVU5Hr1G123880.1 | TRIUR3_10415 | Seita.8G153200.1 | Sobic.005G155900.1 | Zm00001d053124_T001 |
| orthologID_007320 | PH01001551G0040 | LOC_Os11g16310.1 | Bradi4g20710.3 | HORVU4Hr1G014590.30 | TRIUR3_18766 | Seita.8G101400.1 | Sobic.005G105300.1 | Zm00001d049081_T001 |
| orthologID_007486 | PH01000315G0180 | LOC_Os11g10990.1 | Bradi4g22220.1 | HORVU4Hr1G016510.3 | TRIUR3_03738 | Seita.8G095800.1 | Sobic.005G101300.1 | Zm00001d049048_T001 |
| orthologID_007178 | PH01000375G0520 | LOC_Os06g27650.1 | Bradi4g19628.7 | HORVU6Hr1G026840.4 | TRIUR3_27289 | Seita.9G135200.1 | Sobic.005G113766.1 | Zm00001d004644_T003 |
| orthologID_007124 | PH01000005G1420 | LOC_Os11g35340.1 | Bradi4g16310.3 | HORVU2Hr1G031830.5 | TRIUR3_18705 | Seita.8G152900.1 | Sobic.005G155700.1 | Zm00001d053122_T001 |
| orthologID_007329 | PH01000957G0570 | LOC_Os11g05300.1 | Bradi4g42620.1 | HORVU0Hr1G011460.2 | TRIUR3_17500 | Seita.7G321400.1 | Sobic.005G036400.1 | Zm00001d004917_T001 |
| orthologID_007419 | PH01003146G0190 | LOC_Os11g47944.1 | Bradi4g09130.1 | HORVU4Hr1G027300.4 | TRIUR3_32353 | Seita.8G251000.1 | Sobic.005G231300.1 | Zm00001d048595_T001 |
| orthologID_007098 | PH01000634G0070 | LOC_Os11g16320.1 | Bradi4g20700.2 | HORVU4Hr1G014540.1 | TRIUR3_15416 | Seita.8G101500.1 | Sobic.005G105500.1 | Zm00001d049082_T001 |
| orthologID_007465 | PH01002050G0200 | LOC_Os12g07650.1 | Bradi4g45030.2 | HORVU5Hr1G048120.8 | TRIUR3_17081 | Seita.2G350100.1 | Sobic.005G002900.1 | Zm00001d044606_T001 |
| orthologID_007646 | PH01001918G0160 | LOC_Os03g25470.1 | Bradi1g24180.1 | HORVU2Hr1G039050.1 | TRIUR3_24790 | Seita.3G293100.1 | Sobic.005G059300.1 | Zm00001d053025_T001 |
| orthologID_009775 | PH01002370G0100 | LOC_Os10g28120.1 | Bradi3g26840.1 | HORVU1Hr1G072250.1 | TRIUR3_18141 | Seita.8G077000.1 | Sobic.005G084300.1 | Zm00001d049028_T001 |
| orthologID_007264 | PH01002332G0080 | LOC_Os11g32030.1 | Bradi4g17890.1 | HORVU1Hr1G005730.1 | TRIUR3_04471 | Seita.8G134500.1 | Sobic.005G129700.1 | Zm00001d049277_T001 |

| | | | | | | | | |
|---|---|---|---|---|---|---|---|---|
| orthologID_007480 | PH01002847G0260 | LOC_Os11g12800.1 | Bradi4g21756.1 | HORVU4Hr1G015590.7 | TRIUR3_35147 | Seita.8G083900.1 | Sobic.005G090300.1 | Zm00001d048970_T003 |
| orthologID_007170 | PH01002178G0130 | LOC_Os11g43890.1 | Bradi4g10930.1 | HORVU4Hr1G033170.1 | TRIUR3_08590 | Seita.8G222600.1 | Sobic.005G206700.1 | Zm00001d048691_T001 |
| orthologID_007265 | PH01000524G0220 | LOC_Os11g01890.1 | Bradi4g44660.1 | HORVU5Hr1G047060.7 | TRIUR3_26072 | Seita.8G016200.1 | Sobic.005G009300.1 | Zm00001d052329_T002 |
| orthologID_007547 | PH01005130G0050 | LOC_Os11g42000.1 | Bradi4g12039.1 | HORVU3Hr1G017190.1 | TRIUR3_00682 | Seita.8G209500.1 | Sobic.005G198100.1 | Zm00001d007645_T002 |
| orthologID_007493 | PH01004097G0110 | LOC_Os11g35360.1 | Bradi4g16300.1 | HORVU2Hr1G017920.2 | TRIUR3_14569 | Seita.8G153100.1 | Sobic.005G155800.1 | Zm00001d053123_T001 |
| orthologID_007064 | PH01001787G0070 | LOC_Os11g11980.1 | Bradi4g21880.2 | HORVU4Hr1G015820.1 | TRIUR3_35445 | Seita.8G087900.1 | Sobic.005G093400.1 | Zm00001d052985_T002 |
| orthologID_007242 | PH01007697G0010 | LOC_Os11g36070.3 | Bradi4g16140.1 | HORVU4Hr1G005740.21 | TRIUR3_07443 | Seita.8G156900.1 | Sobic.005G157000.2 | Zm00001d052922_T003 |
| orthologID_007272 | PH01000802G0090 | LOC_Os11g11500.1 | Bradi4g22020.2 | HORVU4Hr1G016050.3 | TRIUR3_33810 | Seita.8G091200.1 | Sobic.005G098000.1 | Zm00001d049148_T001 |
| orthologID_007201 | PH01003612G0020 | LOC_Os11g15570.1 | Bradi4g20750.1 | HORVU4Hr1G014670.1 | TRIUR3_27212 | Seita.8G074800.1 | Sobic.005G080200.2 | Zm00001d049127_T001 |
| orthologID_007198 | PH01001100G0140 | LOC_Os11g29840.1 | Bradi4g19040.2 | HORVU6Hr1G093860.1 | TRIUR3_25633 | Seita.8G122900.1 | Sobic.005G117100.1 | Zm00001d049174_T001 |
| orthologID_007279 | PH01000367G0600 | LOC_Os11g38010.1 | Bradi4g14870.3 | HORVU3Hr1G001680.6 | TRIUR3_05320 | Seita.8G170400.1 | Sobic.005G169700.1 | Zm00001d048945_T001 |
| orthologID_007448 | PH01003233G0100 | LOC_Os11g29700.1 | Bradi4g19220.1 | HORVU3Hr1G005490.1 | TRIUR3_24496 | Seita.8G122000.1 | Sobic.005G116100.1 | Zm00001d007320_T002 |
| orthologID_007240 | PH01002557G0230 | LOC_Os11g40500.1 | Bradi4g13337.1 | HORVU6Hr1G003910.3 | TRIUR3_01297 | Seita.7G228300.1 | Sobic.005G191600.1 | Zm00001d048825_T001 |
| orthologID_007377 | PH01001100G0090 | LOC_Os11g29810.1 | Bradi4g19000.2 | HORVU4Hr1G085150.1 | TRIUR3_10887 | Seita.8G122600.1 | Sobic.005G116900.1 | Zm00001d049176_T001 |
| orthologID_007299 | PH01000053G2340 | LOC_Os11g03060.1 | Bradi4g44120.7 | HORVU5Hr1G045960.3 | TRIUR3_15954 | Seita.8G009200.1 | Sobic.005G017300.2 | Zm00001d052379_T004 |
| orthologID_007328 | PH01000375G0650 | LOC_Os11g22960.1 | Bradi4g19620.1 | HORVU4Hr1G080590.1 | TRIUR3_10520 | Seita.5G329100.1 | Sobic.005G125300.1 | Zm00001d049256_T001 |
| orthologID_007231 | PH01000136G1390 | LOC_Os11g41910.1 | Bradi4g12180.2 | HORVU3Hr1G013770.2 | TRIUR3_29921 | Seita.8G208000.1 | Sobic.005G197300.1 | Zm00001d007639_T003 |
| orthologID_007145 | PH01003533G0050 | LOC_Os11g14170.1 | Bradi4g20960.2 | HORVU4Hr1G014960.21 | TRIUR3_33350 | Seita.8G077600.1 | Sobic.005G084900.4 | Zm00001d049018_T001 |
| orthologID_009605 | PH01002718G0260 | LOC_Os10g20600.1 | Bradi3g25380.1 | HORVU1Hr1G030480.3 | TRIUR3_23994 | Seita.9G274700.1 | Sobic.K004100.1 | Zm00001d030421_T001 |
| orthologID_007274 | PH01000151G1120 | LOC_Os11g16580.1 | Bradi4g20480.4 | HORVU4Hr1G014110.14 | TRIUR3_07934 | Seita.8G104300.1 | Sobic.008G038500.3 | Zm00001d023430_T001 |
| orthologID_007540 | PH01003053G0050 | LOC_Os12g42300.1 | Bradi4g01430.1 | HORVU5Hr1G007980.1 | TRIUR3_18471 | Seita.3G390600.1 | Sobic.008G173800.2 | Zm00001d041308_T001 |
| orthologID_007133 | PH01000491G0110 | LOC_Os12g08820.2 | Bradi4g40720.5 | HORVU2Hr1G086020.2 | TRIUR3_09571 | Seita.7G278300.1 | Sobic.008G064900.1 | Zm00001d041823_T003 |
| orthologID_007255 | PH01000245G0690 | LOC_Os11g16550.1 | Bradi4g20500.1 | HORVU4Hr1G014080.4 | TRIUR3_19301 | Seita.8G104100.1 | Sobic.008G038300.1 | Zm00001d023427_T001 |
| orthologID_007148 | PH01000674G0560 | LOC_Os12g34880.1 | Bradi4g05890.2 | HORVU5Hr1G025950.6 | TRIUR3_08393 | Seita.3G346100.1 | Sobic.008G116300.1 | Zm00001d023888_T001 |
| orthologID_009996 | PH01000006G2010 | LOC_Os12g32650.1 | Bradi4g06434.1 | HORVU5Hr1G028110.2 | TRIUR3_35322 | Seita.3G352700.1 | Sobic.008G110600.3 | Zm00001d035297_T001 |
| orthologID_009660 | PH01001140G0400 | LOC_Os02g20160.1 | Bradi3g11437.1 | HORVU5Hr1G111970.18 | TRIUR3_21652 | Seita.4G000500.1 | Sobic.008G150200.1 | Zm00001d041380_T001 |
| orthologID_007471 | PH01000006G2800 | LOC_Os12g33080.1 | Bradi4g06300.2 | HORVU5Hr1G027720.1 | TRIUR3_18353 | Seita.3G350700.1 | Sobic.008G112300.1 | Zm00001d023929_T001 |
| orthologID_007414 | PH01000172G0030 | LOC_Os12g10720.1 | Bradi4g40182.1 | HORVU7Hr1G107350.1 | TRIUR3_31805 | Seita.3G097200.1 | Sobic.008G074300.2 | Zm00001d023968_T005 |
| orthologID_007536 | PH01000925G0250 | LOC_Os12g06980.1 | Bradi4g41696.1 | HORVU5Hr1G035800.7 | TRIUR3_34701 | Seita.7G291300.1 | Sobic.008G049701.1 | Zm00001d041889_T048 |
| orthologID_007304 | PH01004953G0070 | LOC_Os12g26880.1 | Bradi4g07800.1 | HORVU6Hr1G071700.2 | TRIUR3_17654 | Seita.3G314400.1 | Sobic.008G020100.2 | Zm00001d052950_T001 |
| orthologID_006985 | PH01000248G0850 | LOC_Os07g06660.1 | Bradi2g33500.1 | HORVU1Hr1G027840.2 | TRIUR3_31278 | Seita.3G108700.1 | Sobic.008G188600.1 | Zm00001d031186_T003 |
| orthologID_006719 | PH01002124G0020 | LOC_Os12g06410.1 | Bradi2g10920.2 | HORVU6Hr1G034250.1 | TRIUR3_34523 | Seita.7G295400.1 | Sobic.008G044300.1 | Zm00001d023480_T001 |
| orthologID_007390 | PH01000417G0640 | LOC_Os12g43580.1 | Bradi4g00967.1 | HORVU5Hr1G002340.2 | TRIUR3_25679 | Seita.3G401200.1 | Sobic.008G185600.2 | Zm00001d041258_T001 |
| orthologID_007462 | PH01000084G1870 | LOC_Os12g37370.1 | Bradi4g04820.1 | HORVU5Hr1G023090.12 | TRIUR3_10000 | Seita.3G335700.1 | Sobic.008G128900.1 | Zm00001d041609_T001 |
| orthologID_007191 | PH01001605G0150 | LOC_Os12g29760.1 | Bradi4g07350.1 | HORVU5Hr1G013840.1 | TRIUR3_04456 | Seita.3G322800.1 | Sobic.008G097000.1 | Zm00001d023652_T002 |
| orthologID_007288 | PH01000053G0820 | LOC_Os06g50110.1 | Bradi4g43780.1 | HORVU0Hr1G010750.4 | TRIUR3_15217 | Seita.7G309300.1 | Sobic.008G018500.1 | Zm00001d023270_T001 |
| orthologID_007477 | PH01002156G0090 | LOC_Os12g12170.1 | Bradi4g40100.1 | HORVU7Hr1G100040.1 | TRIUR3_23903 | Seita.3G095800.1 | Sobic.008G077500.1 | Zm00001d023978_T001 |
| orthologID_007052 | PH01000463G1090 | LOC_Os12g32330.1 | Bradi4g06650.1 | HORVU0Hr1G011860.3 | TRIUR3_22853 | Seita.3G354800.1 | Sobic.008G108300.1 | Zm00001d035318_T001 |
| orthologID_007475 | PH01002722G0170 | LOC_Os12g08200.1 | Bradi4g40980.1 | HORVU5Hr1G037890.1 | TRIUR3_18152 | Seita.7G281400.1 | Sobic.008G061000.1 | Zm00001d041792_T001 |
| orthologID_007142 | PH01000910G0680 | LOC_Os12g38640.1 | Bradi4g03820.1 | HORVU5Hr1G020500.1 | TRIUR3_16301 | Seita.3G324600.1 | Sobic.008G139900.1 | Zm00001d030762_T001 |
| orthologID_007046 | PH01000602G0110 | LOC_Os12g44370.1 | Bradi4g00300.1 | HORVU2Hr1G126550.1 | TRIUR3_13941 | Seita.3G408900.1 | Sobic.008G193200.1 | Zm00001d041191_T001 |
| orthologID_007366 | PH01000925G0430 | LOC_Os12g07060.1 | Bradi4g41597.1 | HORVU5Hr1G036400.20 | TRIUR3_08599 | Seita.7G289900.1 | Sobic.008G051200.1 | Zm00001d023537_T002 |
| orthologID_007367 | PH01000185G1010 | LOC_Os12g39080.1 | Bradi4g03610.1 | HORVU5Hr1G019110.1 | TRIUR3_29862 | Seita.3G365700.1 | Sobic.008G142600.1 | Zm00001d030832_T001 |
| orthologID_007236 | PH01002107G0270 | LOC_Os11g03920.1 | Bradi4g43660.1 | HORVU5Hr1G044970.6 | TRIUR3_16346 | Seita.4G031800.1 | Sobic.008G016700.1 | Zm00001d015730_T001 |
| orthologID_007368 | PH01000025G1400 | LOC_Os12g40830.1 | Bradi4g02640.2 | HORVU5Hr1G014350.2 | TRIUR3_27325 | Seita.3G379000.1 | Sobic.008G156300.1 | Zm00001d041407_T002 |
| orthologID_007301 | PH01003243G0030 | LOC_Os03g25260.1 | Bradi4g44820.1 | HORVU5Hr1G047490.2 | TRIUR3_09182 | Seita.9G526000.1 | Sobic.008G007300.1 | Zm00001d052325_T001 |
| orthologID_007053 | PH01000183G0200 | LOC_Os12g42210.1 | Bradi4g01490.1 | HORVU5Hr1G008160.4 | TRIUR3_06414 | Seita.3G391500.1 | Sobic.008G172700.1 | Zm00001d000230_T002 |
| orthologID_007080 | PH01000245G0660 | LOC_Os11g16560.1 | Bradi4g20490.1 | HORVU4Hr1G014070.2 | TRIUR3_19300 | Seita.8G104200.1 | Sobic.008G038400.1 | Zm00001d023426_T005 |
| orthologID_007549 | PH01000134G0880 | LOC_Os12g44160.2 | Bradi4g00497.2 | HORVU5Hr1G000770.28 | TRIUR3_25362 | Seita.3G406800.1 | Sobic.008G190600.1 | Zm00001d031216_T002 |
| orthologID_007166 | PH01001390G0400 | LOC_Os12g06080.1 | Bradi4g42167.1 | HORVU5Hr1G040620.2 | TRIUR3_16157 | Seita.7G298100.1 | Sobic.008G041100.1 | Zm00001d023446_T001 |

| | | | | | | | | |
|---|---|---|---|---|---|---|---|---|
| orthologID_007514 | PH01000183G1300 | LOC_Os12g41700.1 | Bradi4g02040.1 | HORVU5Hr1G011690.2 | TRIUR3_03600 | Seita.3G386400.1 | Sobic.008G165000.1 | Zm00001d041474_T001 |
| orthologID_007425 | PH01002447G0330 | LOC_Os12g37610.1 | Bradi4g04560.2 | HORVU5Hr1G022510.1 | TRIUR3_17721 | Seita.3G334300.1 | Sobic.008G130900.1 | Zm00001d041595_T001 |
| orthologID_007402 | PH01000116G0770 | LOC_Os12g29400.1 | Bradi4g07520.1 | HORVU5Hr1G115100.2 | TRIUR3_11344 | Seita.3G320700.1 | Sobic.008G095000.1 | Zm00001d023664_T001 |
| orthologID_010016 | PH01000202G1270 | LOC_Os05g19910.1 | Bradi4g06067.1 | HORVU3Hr1G066740.3 | TRIUR3_04135 | Seita.3G292000.1 | Sobic.008G068300.1 | Zm00001d035246_T001 |
| orthologID_007496 | PH01000006G3570 | LOC_Os12g34500.1 | Bradi4g06017.1 | HORVU4Hr1G031530.1 | TRIUR3_23548 | Seita.3G348500.1 | Sobic.008G113800.1 | Zm00001d032945_T001 |
| orthologID_006586 | PH01000762G0700 | LOC_Os01g01302.1 | Bradi2g03680.1 | HORVU3Hr1G014780.6 | TRIUR3_05441 | Seita.3G330300.1 | Sobic.008G134800.1 | Zm00001d030688_T005 |
| orthologID_007077 | PH01000185G0030 | LOC_Os12g39640.1 | Bradi4g03160.1 | HORVU5Hr1G018220.11 | TRIUR3_17056 | Seita.3G370000.1 | Sobic.008G147101.1 | Zm00001d030891_T001 |
| orthologID_007246 | PH01000006G2340 | LOC_Os12g32950.1 | Bradi4g06402.1 | HORVU5Hr1G028000.1 | TRIUR3_14983 | Seita.3G351800.1 | Sobic.008G111400.1 | Zm00001d035288_T002 |
| orthologID_007326 | PH01000910G0290 | LOC_Os12g38800.1 | Bradi4g03736.1 | HORVU5Hr1G020200.13 | TRIUR3_18382 | Seita.3G363900.1 | Sobic.008G140700.1 | Zm00001d030774_T004 |
| orthologID_007558 | PH01007343G0030 | LOC_Os12g12730.1 | Bradi4g39900.1 | HORVU7Hr1G008910.2 | TRIUR3_31105 | Seita.3G093800.1 | Sobic.008G079200.2 | Zm00001d023992_T001 |
| orthologID_007129 | PH01000185G0920 | LOC_Os12g39110.1 | Bradi4g03530.2 | HORVU5Hr1G019040.1 | TRIUR3_27570 | Seita.3G367000.1 | Sobic.008G143100.1 | Zm00001d030843_T001 |
| orthologID_007189 | PH01000162G0420 | LOC_Os12g42400.2 | Bradi4g01380.3 | HORVU5Hr1G007890.11 | TRIUR3_28596 | Seita.3G390000.1 | Sobic.008G174650.1 | Zm00001d031092_T001 |
| orthologID_007332 | PH01000910G0320 | LOC_Os12g38790.1 | Bradi4g03742.2 | HORVU5Hr1G020250.2 | TRIUR3_08619 | Seita.3G363800.1 | Sobic.008G140600.1 | Zm00001d041523_T001 |
| orthologID_007115 | PH01000674G0710 | LOC_Os06g07200.1 | Bradi4g05940.1 | HORVU5Hr1G026120.1 | TRIUR3_12376 | Seita.3G347200.1 | Sobic.008G115500.1 | Zm00001d041716_T002 |
| orthologID_007312 | PH01000169G0440 | LOC_Os12g42220.1 | Bradi4g01480.1 | HORVU5Hr1G008180.2 | TRIUR3_06412 | Seita.3G391400.1 | Sobic.008G172800.3 | Zm00001d000229_T001 |
| orthologID_007213 | PH01000351G0700 | LOC_Os12g07950.1 | Bradi4g41127.2 | HORVU5Hr1G037540.3 | TRIUR3_26696 | Seita.7G283400.1 | Sobic.008G058100.5 | Zm00001d023585_T001 |
| orthologID_007520 | PH01000006G1570 | LOC_Os12g32620.1 | Bradi4g06510.1 | HORVU5Hr1G028180.1 | TRIUR3_20387 | Seita.3G353200.1 | Sobic.008G110200.1 | Zm00001d035304_T001 |
| orthologID_007429 | PH01000574G0140 | LOC_Os12g18630.1 | Bradi4g39110.1 | HORVU5Hr1G030040.1 | TRIUR3_22949 | Seita.3G307400.1 | Sobic.008G090900.1 | Zm00001d023791_T002 |
| orthologID_009517 | PH01000600G0220 | LOC_Os12g40450.1 | Bradi3g06440.3 | HORVU5Hr1G016750.6 | TRIUR3_07457 | Seita.3G374500.1 | Sobic.008G152000.2 | Zm00001d030944_T001 |
| orthologID_008573 | PH01004864G0030 | LOC_Os08g28830.1 | Bradi1g21907.4 | HORVU6Hr1G010120.6 | TRIUR3_03243 | Seita.6G130100.1 | Sobic.008G179000.1 | Zm00001d031130_T001 |
| orthologID_007215 | PH01000809G0120 | LOC_Os12g17530.1 | Bradi4g39190.1 | HORVU2Hr1G014550.5 | TRIUR3_20625 | Seita.3G306400.1 | Sobic.008G093600.2 | Zm00001d023630_T005 |
| orthologID_007523 | PH01000574G0360 | LOC_Os12g18120.1 | Bradi4g39137.3 | HORVU5Hr1G033710.3 | TRIUR3_24688 | Seita.3G307100.1 | Sobic.008G092300.3 | Zm00001d023636_T001 |
| orthologID_007019 | PH01004313G0100 | LOC_Os12g34870.2 | Bradi4g05917.2 | HORVU5Hr1G026160.2 | TRIUR3_12378 | Seita.3G346400.1 | Sobic.008G115900.1 | Zm00001d023895_T002 |
| orthologID_007524 | PH01000862G0270 | LOC_Os12g25720.1 | Bradi4g07862.2 | HORVU5Hr1G029910.8 | TRIUR3_30030 | Seita.3G300300.1 | Sobic.008G092800.1 | Zm00001d041083_T005 |
| orthologID_007061 | PH01000510G0060 | LOC_Os12g35560.1 | Bradi4g05580.1 | HORVU5Hr1G025310.8 | TRIUR3_04877 | Seita.3G343400.1 | Sobic.008G118400.1 | Zm00001d041692_T001 |
| orthologID_007174 | PH01000302G0960 | LOC_Os12g37720.1 | Bradi4g04650.1 | HORVU5Hr1G022740.1 | TRIUR3_16215 | Seita.3G333500.1 | Sobic.008G131800.1 | Zm00001d041582_T005 |
| orthologID_007544 | PH01002063G0250 | LOC_Os12g34340.1 | Bradi4g06130.1 | HORVU5Hr1G027150.1 | TRIUR3_25805 | Seita.3G349700.1 | Sobic.008G113200.2 | Zm00001d023915_T001 |
| orthologID_007470 | PH01006576G0030 | LOC_Os12g07290.1 | Bradi4g41520.4 | HORVU5Hr1G036120.2 | TRIUR3_29249 | Seita.7G287400.1 | Sobic.008G053600.1 | Zm00001d023557_T001 |
| orthologID_007436 | PH01001660G0380 | LOC_Os12g06850.1 | Bradi4g41760.1 | HORVU5Hr1G035760.3 | TRIUR3_20981 | Seita.7G292500.1 | Sobic.008G048300.1 | Zm00001d023514_T003 |
| orthologID_007483 | PH01002374G0040 | LOC_Os12g35620.1 | Bradi4g05517.1 | HORVU5Hr1G024440.1 | TRIUR3_18529 | Seita.3G343000.1 | Sobic.008G118800.1 | Zm00001d041691_T002 |
| orthologID_007226 | PH01000674G0270 | LOC_Os12g35270.1 | Bradi4g05720.1 | HORVU5Hr1G025810.1 | TRIUR3_20688 | Seita.3G345000.1 | Sobic.008G117100.1 | Zm00001d041696_T001 |
| orthologID_007203 | PH01087490G0010 | LOC_Os12g08770.1 | Bradi4g40780.1 | HORVU2Hr1G019820.1 | TRIUR3_13284 | Seita.7G279400.1 | Sobic.008G063500.1 | Zm00001d041819_T001 |
| orthologID_007537 | PH01003703G0160 | LOC_Os05g18950.1 | Bradi4g21120.4 | HORVU4Hr1G044120.2 | TRIUR3_21898 | Seita.3G392900.1 | Sobic.008G170600.1 | Zm00001d041326_T001 |
| orthologID_007451 | PH01001693G0200 | LOC_Os12g09290.1 | Bradi4g40560.1 | HORVU5Hr1G029010.1 | TRIUR3_25709 | Seita.7G276800.1 | Sobic.008G068100.1 | Zm00001d041833_T003 |
| orthologID_007063 | PH01000021G2170 | LOC_Os12g23180.1 | Bradi4g08030.1 | HORVU4Hr1G052270.4 | TRIUR3_10465 | Seita.3G304400.1 | Sobic.008G066100.1 | Zm00001d035212_T001 |
| orthologID_008246 | PH01001638G0260 | LOC_Os07g01200.1 | Bradi1g59740.1 | HORVU2Hr1G064900.3 | TRIUR3_25072 | Seita.3G375600.1 | Sobic.008G152500.5 | Zm00001d030951_T002 |
| orthologID_007344 | PH01000866G0030 | LOC_Os12g31380.1 | Bradi4g07040.1 | HORVU5Hr1G038720.1 | TRIUR3_13639 | Seita.3G359800.1 | Sobic.008G101600.1 | Zm00001d041758_T001 |
| orthologID_007037 | PH01000110G0250 | LOC_Os12g41590.1 | Bradi4g02160.1 | HORVU5Hr1G011960.2 | TRIUR3_12942 | Seita.3G383900.1 | Sobic.008G162300.2 | Zm00001d031026_T001 |
| orthologID_007076 | PH01001070G0650 | LOC_Os12g02340.1 | Bradi4g44400.1 | HORVU5Hr1G046520.1 | TRIUR3_14525 | Seita.8G013500.1 | Sobic.008G030600.1 | Zm00001d044685_T001 |
| orthologID_007067 | PH01003590G0040 | LOC_Os12g12740.1 | Bradi4g39887.1 | HORVU2Hr1G016030.2 | TRIUR3_15927 | Seita.3G093700.1 | Sobic.008G080200.1 | Zm00001d041175_T002 |
| orthologID_007433 | PH01000491G0120 | LOC_Os12g08830.1 | Bradi4g40710.1 | HORVU2Hr1G086040.4 | TRIUR3_09570 | Seita.7G278200.1 | Sobic.008G065000.1 | Zm00001d041824_T001 |
| orthologID_009617 | PH01002010G0170 | LOC_Os02g58590.1 | Bradi3g60820.1 | HORVU3Hr1G106650.3 | TRIUR3_02381 | Seita.4G270000.1 | Sobic.008G042600.1 | Zm00001d025232_T002 |
| orthologID_007378 | PH01001390G0320 | LOC_Os12g06100.2 | Bradi4g42110.1 | HORVU5Hr1G040400.6 | TRIUR3_32645 | Seita.7G297600.1 | Sobic.008G041700.1 | Zm00001d023452_T001 |
| orthologID_007508 | PH01000463G0650 | LOC_Os12g32190.1 | Bradi4g06700.1 | HORVU5Hr1G028460.1 | TRIUR3_23675 | Seita.3G355700.1 | Sobic.008G107000.1 | Zm00001d035236_T001 |
| orthologID_009645 | PH01002456G0150 | LOC_Os03g47470.1 | Bradi3g38300.1 | HORVU5Hr1G059030.1 | TRIUR3_04912 | Seita.3G312800.1 | Sobic.008G011200.1 | Zm00001d023240_T001 |
| orthologID_007494 | PH01001336G0010 | LOC_Os12g08260.2 | Bradi4g40870.2 | HORVU5Hr1G034820.4 | TRIUR3_13101 | Seita.7G280800.1 | Sobic.008G062200.1 | Zm00001d023694_T001 |
| orthologID_007531 | PH01002056G0230 | LOC_Os12g32760.1 | Bradi4g06420.2 | HORVU5Hr1G028060.1 | TRIUR3_31810 | Seita.3G352300.1 | Sobic.008G111100.1 | Zm00001d023941_T001 |
| orthologID_007321 | PH01001092G0040 | LOC_Os12g06520.1 | Bradi4g41890.1 | HORVU5Hr1G039870.5 | TRIUR3_31416 | Seita.7G294400.1 | Sobic.008G046600.1 | Zm00001d041920_T001 |
| orthologID_007202 | PH01001815G0110 | LOC_Os12g25180.1 | Bradi4g07896.1 | HORVU7Hr1G086720.3 | TRIUR3_28591 | Seita.3G300500.1 | Sobic.008G130700.1 | Zm00001d023693_T001 |
| orthologID_007519 | PH01000417G1130 | LOC_Os12g42970.1 | Bradi4g01140.1 | HORVU5Hr1G005990.5 | TRIUR3_20013 | Seita.3G397100.1 | Sobic.008G179800.1 | Zm00001d031135_T001 |

| | | | | | | | | |
|---|---|---|---|---|---|---|---|---|
| orthologID_007016 | PH01001154G0580 | LOC_Os12g40880.1 | Bradi4g02607.2 | HORVU5Hr1G014310.2 | TRIUR3_13986 | Seita.3G379300.1 | Sobic.008G156700.1 | Zm00001d041415_T001 |
| orthologID_007065 | PH01000183G0950 | LOC_Os12g41830.1 | Bradi4g01900.8 | HORVU5Hr1G011020.9 | TRIUR3_15808 | Seita.3G395100.1 | Sobic.008G168000.1 | Zm00001d031060_T001 |
| orthologID_007428 | PH01000134G0230 | LOC_Os12g43930.1 | Bradi4g00760.10 | HORVU5Hr1G001400.5 | TRIUR3_02060 | Seita.3G404600.1 | Sobic.008G188700.3 | Zm00001d031184_T001 |
| orthologID_007518 | PH01000105G0720 | LOC_Os12g41390.1 | Bradi4g02350.1 | HORVU5Hr1G012780.2 | TRIUR3_24552 | Seita.3G382700.1 | Sobic.008G161700.1 | Zm00001d041455_T002 |
| orthologID_007393 | PH01006110G0040 | LOC_Os12g13320.1 | Bradi4g01920.1 | HORVU1Hr1G062900.2 | TRIUR3_07046 | Seita.3G090300.1 | Sobic.008G083400.1 | Zm00001d024048_T004 |
| orthologID_007041 | PH01000302G0760 | LOC_Os12g37630.1 | Bradi4g04567.2 | HORVU5Hr1G022520.4 | TRIUR3_17720 | Seita.3G334200.1 | Sobic.008G131000.1 | Zm00001d030639_T001 |
| orthologID_007044 | PH01000866G0070 | LOC_Os12g31460.1 | Bradi4g06980.1 | HORVU5Hr1G039210.1 | TRIUR3_30292 | Seita.3G359500.1 | Sobic.008G102000.1 | Zm00001d023757_T001 |
| orthologID_007482 | PH01000957G0020 | LOC_Os11g05100.1 | Bradi4g42817.2 | HORVU5Hr1G041100.7 | TRIUR3_28513 | Seita.7G322600.1 | Sobic.008G033500.1 | Zm00001d044696_T002 |
| orthologID_007300 | PH01002122G0230 | LOC_Os12g38400.2 | Bradi4g03970.1 | HORVU5Hr1G021230.1 | TRIUR3_13997 | Seita.3G325900.1 | Sobic.008G137500.1 | Zm00001d030737_T001 |
| orthologID_007241 | PH01000302G0660 | LOC_Os12g37590.1 | Bradi4g04540.1 | HORVU5Hr1G022470.1 | TRIUR3_24548 | Seita.3G334500.1 | Sobic.008G130400.1 | Zm00001d041597_T001 |
| orthologID_007358 | PH01001065G0050 | LOC_Os12g39840.1 | Bradi4g03114.1 | HORVU5Hr1G018150.3 | TRIUR3_24454 | Seita.3G370400.1 | Sobic.008G147700.1 | Zm00001d041346_T001 |
| orthologID_007495 | PH01005541G0050 | LOC_Os12g07050.1 | Bradi4g41610.1 | HORVU5Hr1G036490.1 | TRIUR3_35038 | Seita.7G290000.1 | Sobic.008G051100.1 | Zm00001d041884_T001 |
| orthologID_007479 | PH01000110G1100 | LOC_Os12g41810.1 | Bradi4g01930.1 | HORVU5Hr1G011090.3 | TRIUR3_14612 | Seita.3G395400.1 | Sobic.008G167700.1 | Zm00001d041484_T001 |
| orthologID_007162 | PH01000105G1660 | LOC_Os12g40920.1 | Bradi4g02570.1 | HORVU5Hr1G014170.10 | TRIUR3_15470 | Seita.3G379700.1 | Sobic.008G157100.2 | Zm00001d030995_T002 |
| orthologID_007101 | PH01000510G0080 | LOC_Os12g35580.1 | Bradi4g05560.1 | HORVU5Hr1G024590.1 | TRIUR3_04875 | Seita.3G343200.1 | Sobic.008G118600.1 | Zm00001d023863_T001 |
| orthologID_007529 | PH01000006G2680 | LOC_Os12g33070.1 | Bradi4g06317.1 | HORVU5Hr1G027750.1 | TRIUR3_33254 | Seita.3G350800.1 | Sobic.008G112200.1 | Zm00001d023932_T001 |
| orthologID_007392 | PH01000025G1090 | LOC_Os12g40500.1 | Bradi4g02747.3 | HORVU5Hr1G016340.1 | TRIUR3_16239 | Seita.3G375000.1 | Sobic.008G152700.1 | Zm00001d030954_T001 |
| orthologID_007454 | PH01000084G1890 | LOC_Os12g37380.1 | Bradi4g04810.2 | HORVU5Hr1G023070.1 | TRIUR3_10001 | Seita.3G335800.1 | Sobic.008G128800.4 | Zm00001d041610_T001 |
| orthologID_007904 | PH01000551G0450 | LOC_Os12g06510.1 | Bradi1g28160.1 | HORVU5Hr1G040030.5 | TRIUR3_34679 | Seita.7G294700.1 | Sobic.008G046500.1 | Zm00001d023504_T001 |
| orthologID_006225 | PH01001816G0040 | LOC_Os05g22920.1 | Bradi2g05445.3 | HORVU0Hr1G014270.22 | TRIUR3_15083 | Seita.2G011900.1 | Sobic.008G032900.1 | Zm00001d023373_T004 |
| orthologID_008062 | PH01000567G0170 | LOC_Os08g10480.1 | Bradi1g41790.1 | HORVU7Hr1G082540.2 | TRIUR3_18185 | Seita.8G110600.1 | Sobic.008G179500.1 | Zm00001d031132_T002 |
| orthologID_008076 | PH01000475G0490 | LOC_Os06g20320.1 | Bradi1g42820.1 | HORVU7Hr1G045680.1 | TRIUR3_34610 | Seita.4G136300.1 | Sobic.008G021700.1 | Zm00001d023292_T001 |
| orthologID_007123 | PH01001450G0370 | LOC_Os12g37100.1 | Bradi4g04840.4 | HORVU5Hr1G023140.9 | TRIUR3_19530 | Seita.1G257400.1 | Sobic.008G127550.1 | Zm00001d000137_T003 |
| orthologID_007322 | PH01000866G0050 | LOC_Os12g31450.1 | Bradi4g06987.2 | HORVU5Hr1G035440.4 | TRIUR3_33358 | Seita.3G359600.1 | Sobic.008G101900.1 | Zm00001d023760_T001 |
| orthologID_008404 | PH01002689G0060 | LOC_Os05g19150.1 | Bradi1g42880.1 | HORVU7Hr1G045640.2 | TRIUR3_24031 | Seita.4G133900.1 | Sobic.008G097100.1 | Zm00001d023651_T001 |
| orthologID_007058 | PH01000006G0610 | LOC_Os12g32180.1 | Bradi4g06710.1 | HORVU5Hr1G028480.1 | TRIUR3_23674 | Seita.3G355900.1 | Sobic.008G106900.1 | Zm00001d035235_T001 |
| orthologID_007447 | PH01004133G0040 | LOC_Os12g13110.1 | Bradi4g39710.1 | HORVU7Hr1G021750.1 | TRIUR3_05386 | Seita.3G091200.1 | Sobic.008G082800.1 | Zm00001d037069_T001 |
| orthologID_010020 | PH01002056G0260 | LOC_Os12g32640.1 | Bradi4g06485.3 | HORVU5Hr1G028140.9 | TRIUR3_20412 | Seita.3G353000.1 | Sobic.008G110400.1 | Zm00001d035300_T001 |
| orthologID_007023 | PH01002353G0300 | LOC_Os12g43560.1 | Bradi4g00980.1 | HORVU5Hr1G003010.3 | TRIUR3_25318 | Seita.3G401100.1 | Sobic.008G185500.2 | Zm00001d031178_T002 |
| orthologID_007314 | PH01000183G0850 | LOC_Os12g41880.2 | Bradi4g01820.7 | HORVU5Hr1G010600.4 | TRIUR3_30486 | Seita.3G394900.1 | Sobic.008G168300.1 | Zm00001d041491_T001 |
| orthologID_007171 | PH01000510G0680 | LOC_Os12g36170.1 | Bradi4g05317.1 | HORVU5Hr1G113260.8 | TRIUR3_34675 | Seita.3G340500.1 | Sobic.008G123400.1 | Zm00001d023839_T001 |
| orthologID_009117 | PH01000012G2610 | LOC_Os10g37180.1 | Bradi3g30500.1 | HORVU6Hr1G076880.1 | TRIUR3_12946 | Seita.7G280500.1 | Sobic.008G062500.1 | Zm00001d023699_T001 |
| orthologID_009577 | PH01005219G0020 | LOC_Os05g22930.1 | Bradi3g04310.1 | HORVU7Hr1G084900.14 | TRIUR3_02486 | Seita.3G075200.1 | Sobic.008G033000.1 | Zm00001d023374_T001 |
| orthologID_007442 | PH01000302G0810 | LOC_Os12g37650.1 | Bradi4g04590.1 | HORVU5Hr1G022550.1 | TRIUR3_28272 | Seita.3G334000.1 | Sobic.008G131200.1 | Zm00001d041578_T001 |
| orthologID_009848 | PH01000473G0370 | LOC_Os02g42300.1 | Bradi3g49580.1 | HORVU6Hr1G059530.2 | TRIUR3_13373 | Seita.1G245400.1 | Sobic.004G224600.1 | Zm00001d017331_T002 |
| orthologID_009642 | PH01004222G0040 | LOC_Os02g10860.1 | Bradi3g07540.1 | HORVU6Hr1G037760.2 | TRIUR3_20602 | Seita.1G043800.1 | Sobic.004G085600.1 | Zm00001d015743_T002 |
| orthologID_009266 | PH01000026G0210 | LOC_Os02g54980.1 | Bradi3g53860.1 | HORVU6Hr1G084510.1 | TRIUR3_10619 | Seita.1G348100.1 | Sobic.004G327200.1 | Zm00001d018358_T001 |
| orthologID_009092 | PH01002669G0080 | LOC_Os02g04450.1 | Bradi3g03260.1 | HORVU6Hr1G018770.3 | TRIUR3_32638 | Seita.1G094000.1 | Sobic.004G034400.1 | Zm00001d054004_T001 |
| orthologID_009749 | PH01000019G2100 | LOC_Os02g02510.1 | Bradi3g01560.1 | HORVU0Hr1G040320.1 | TRIUR3_21893 | Seita.1G118800.1 | Sobic.004G012800.1 | Zm00001d054033_T001 |
| orthologID_009724 | PH01000210G0150 | LOC_Os02g53770.1 | Bradi3g57220.1 | HORVU6Hr1G081370.3 | TRIUR3_34367 | Seita.1G339300.1 | Sobic.004G317700.1 | Zm00001d018274_T003 |
| orthologID_007546 | PH01006035G0050 | LOC_Os02g31070.1 | Bradi4g12940.1 | HORVU4Hr1G031120.1 | TRIUR3_32307 | Seita.1G172400.1 | Sobic.004G157600.1 | Zm00001d016587_T001 |
| orthologID_008634 | PH01001268G0200 | LOC_Os03g04580.1 | Bradi1g75800.1 | HORVU4Hr1G083800.1 | TRIUR3_04848 | Seita.9G546300.1 | Sobic.004G016700.1 | Zm00001d015031_T001 |
| orthologID_009364 | PH01000473G1010 | LOC_Os02g42560.1 | Bradi3g49780.1 | HORVU6Hr1G060080.1 | TRIUR3_16105 | Seita.1G248000.1 | Sobic.004G227000.1 | Zm00001d051235_T001 |
| orthologID_009863 | PH01000005G3860 | LOC_Os09g02130.1 | Bradi3g03180.1 | HORVU6Hr1G018530.2 | TRIUR3_25074 | Seita.4G250100.1 | Sobic.004G028100.1 | Zm00001d053977_T001 |
| orthologID_009766 | PH01000227G0940 | LOC_Os02g06370.1 | Bradi3g04450.1 | HORVU6Hr1G025290.3 | TRIUR3_19131 | Seita.1G081000.1 | Sobic.004G047500.1 | Zm00001d053855_T001 |
| orthologID_006188 | PH01005384G0040 | LOC_Os05g27100.1 | Bradi2g30370.1 | HORVU1Hr1G064750.1 | TRIUR3_12682 | Seita.3G283700.1 | Sobic.004G317900.1 | Zm00001d038047_T001 |
| orthologID_009814 | PH01000765G0270 | LOC_Os02g08240.1 | Bradi3g05890.2 | HORVU6Hr1G029780.1 | TRIUR3_28599 | Seita.1G064300.1 | Sobic.004G064200.1 | Zm00001d053735_T002 |
| orthologID_009914 | PH01002859G0070 | LOC_Os02g54520.1 | Bradi3g60220.2 | HORVU6Hr1G083250.1 | TRIUR3_13802 | Seita.1G344700.1 | Sobic.004G322800.1 | Zm00001d052069_T001 |
| orthologID_009390 | PH01000585G0800 | LOC_Os02g03840.2 | Bradi3g02910.2 | HORVU6Hr1G016750.9 | TRIUR3_11299 | Seita.1G097800.1 | Sobic.004G030200.1 | Zm00001d015138_T004 |
| orthologID_009222 | PH01000965G0480 | LOC_Os02g09750.1 | Bradi3g06597.1 | HORVU6Hr1G035000.1 | TRIUR3_26327 | Seita.1G055200.1 | Sobic.004G074000.1 | Zm00001d015651_T001 |

| | | | | | | | | |
|---|---|---|---|---|---|---|---|---|
| orthologID_009845 | PH01003050G0110 | LOC_Os07g40920.1 | Bradi3g45970.1 | HORVU6Hr1G064120.2 | TRIUR3_04546 | Seita.1G195900.1 | Sobic.004G179500.1 | Zm00001d016853_T001 |
| orthologID_009618 | PH01000359G0810 | LOC_Os02g34490.1 | Bradi3g45402.2 | HORVU2Hr1G016070.6 | TRIUR3_25463 | Seita.1G189000.1 | Sobic.004G172400.1 | Zm00001d050714_T001 |
| orthologID_009257 | PH01149302G0010 | LOC_Os02g46190.1 | Bradi3g52016.1 | HORVU6Hr1G066610.2 | TRIUR3_29605 | Seita.1G276400.1 | Sobic.004G278200.1 | Zm00001d051492_T001 |
| orthologID_009588 | PH01000095G0630 | LOC_Os02g07250.1 | Bradi3g05210.1 | HORVU6Hr1G033690.10 | TRIUR3_08892 | Seita.1G073600.1 | Sobic.004G055000.1 | Zm00001d015374_T001 |
| orthologID_009671 | PH01000765G0460 | LOC_Os02g08300.1 | Bradi3g05920.3 | HORVU6Hr1G029760.3 | TRIUR3_24946 | Seita.1G063800.1 | Sobic.004G064600.1 | Zm00001d053738_T001 |
| orthologID_009402 | PH01001232G0500 | LOC_Os02g16550.1 | Bradi3g09820.1 | HORVU6Hr1G048780.2 | TRIUR3_33746 | Seita.1G009600.1 | Sobic.004G117800.1 | Zm00001d053371_T001 |
| orthologID_009812 | PH01002043G0120 | LOC_Os02g35890.1 | Bradi3g46410.1 | HORVU2Hr1G102120.1 | TRIUR3_05508 | Seita.1G202900.1 | Sobic.004G186100.1 | Zm00001d016892_T001 |
| orthologID_008142 | PH01003499G0050 | LOC_Os01g49360.1 | Bradi1g26360.1 | HORVU6Hr1G029240.4 | TRIUR3_12722 | Seita.1G127500.1 | Sobic.004G128800.1 | Zm00001d015927_T001 |
| orthologID_009349 | PH01000773G0240 | LOC_Os02g02570.1 | Bradi3g01650.1 | HORVU2Hr1G049500.2 | TRIUR3_22368 | Seita.1G112300.1 | Sobic.004G013600.1 | Zm00001d015009_T002 |
| orthologID_009338 | PH01000860G0270 | LOC_Os02g44570.2 | Bradi3g51060.1 | HORVU6Hr1G064010.6 | TRIUR3_06650 | Seita.1G264000.1 | Sobic.004G289400.1 | Zm00001d051396_T004 |
| orthologID_009612 | PH01107796G0010 | LOC_Os02g50100.1 | Bradi3g56840.1 | HORVU6Hr1G073650.2 | TRIUR3_10865 | Seita.1G310500.1 | Sobic.004G247800.1 | Zm00001d051796_T003 |
| orthologID_009383 | PH01002573G0120 | LOC_Os02g28820.1 | Bradi3g11150.1 | HORVU5Hr1G063600.1 | TRIUR3_00476 | Seita.1G156900.1 | Sobic.004G033900.1 | Zm00001d015198_T001 |
| orthologID_009852 | PH01000203G0860 | LOC_Os02g55120.1 | Bradi3g53970.2 | HORVU6Hr1G084690.1 | TRIUR3_18061 | Seita.1G349300.1 | Sobic.004G328400.2 | Zm00001d018366_T001 |
| orthologID_009902 | PH01000123G0160 | LOC_Os02g07740.1 | Bradi3g05437.2 | HORVU6Hr1G031520.6 | TRIUR3_12676 | Seita.1G069700.1 | Sobic.004G058400.1 | Zm00001d015400_T004 |
| orthologID_009694 | PH01000026G2150 | LOC_Os02g56260.1 | Bradi3g54730.2 | HORVU6Hr1G088380.2 | TRIUR3_19566 | Seita.1G358500.1 | Sobic.004G337700.1 | Zm00001d018422_T001 |
| orthologID_009836 | PH01000585G0760 | LOC_Os02g03860.1 | Bradi3g02890.1 | HORVU0Hr1G018510.1 | TRIUR3_14987 | Seita.1G098000.1 | Sobic.004G030000.1 | Zm00001d015134_T001 |
| orthologID_009594 | PH01002035G0220 | LOC_Os02g47000.1 | Bradi3g52380.1 | HORVU0Hr1G032120.1 | TRIUR3_18133 | Seita.1G283500.1 | Sobic.004G272300.1 | Zm00001d017708_T001 |
| orthologID_009137 | PH01001760G0290 | LOC_Os02g46120.1 | Bradi3g51990.2 | HORVU6Hr1G066510.9 | TRIUR3_07944 | Seita.1G275900.1 | Sobic.004G278900.2 | Zm00001d051500_T001 |
| orthologID_009297 | PH01000447G0740 | LOC_Os02g44930.1 | Bradi3g51140.1 | HORVU6Hr1G064720.1 | TRIUR3_34670 | Seita.1G267900.1 | Sobic.004G285900.3 | Zm00001d017555_T001 |
| orthologID_009262 | PH01000227G0530 | LOC_Os02g06660.1 | Bradi3g04770.1 | HORVU6Hr1G026330.5 | TRIUR3_20397 | Seita.1G078600.1 | Sobic.004G050600.1 | Zm00001d015334_T001 |
| orthologID_009282 | PH01000089G0730 | LOC_Os02g39400.1 | Bradi3g48140.1 | HORVU6Hr1G055250.1 | TRIUR3_20600 | Seita.1G224700.1 | Sobic.004G208000.1 | Zm00001d051015_T001 |
| orthologID_009120 | PH01001580G0330 | LOC_Os02g57180.1 | Bradi3g55340.1 | HORVU6Hr1G090720.1 | TRIUR3_14360 | Seita.1G365400.1 | Sobic.004G344800.1 | Zm00001d018479_T001 |
| orthologID_009316 | PH01000343G0240 | LOC_Os02g43760.1 | Bradi3g50470.4 | HORVU6Hr1G062360.2 | TRIUR3_19019 | Seita.1G257100.1 | Sobic.004G297300.1 | Zm00001d051337_T003 |
| orthologID_009335 | PH01003125G0200 | LOC_Os02g18870.1 | Bradi3g51720.1 | HORVU1Hr1G064460.2 | TRIUR3_25020 | Seita.1G130400.1 | Sobic.004G130500.1 | Zm00001d015960_T001 |
| orthologID_009096 | PH01000311G0400 | LOC_Os02g53270.1 | Bradi3g57560.1 | HORVU6Hr1G079360.5 | TRIUR3_31397 | Seita.1G335900.1 | Sobic.004G314000.1 | Zm00001d018226_T001 |
| orthologID_009276 | PH01003968G0140 | LOC_Os02g45620.1 | Bradi3g51707.1 | HORVU6Hr1G065900.1 | TRIUR3_27873 | Seita.1G272200.1 | Sobic.004G282200.2 | Zm00001d017606_T001 |
| orthologID_009228 | PH01000796G0220 | LOC_Os02g21110.1 | Bradi3g11000.1 | HORVU6Hr1G024710.1 | TRIUR3_20919 | Seita.1G142500.1 | Sobic.004G119000.1 | Zm00001d015698_T001 |
| orthologID_009200 | PH01000124G0280 | LOC_Os02g10770.2 | Bradi3g07457.1 | HORVU6Hr1G038030.5 | TRIUR3_22375 | Seita.1G044600.1 | Sobic.004G084700.2 | Zm00001d015758_T001 |
| orthologID_009851 | PH01000582G0620 | LOC_Os02g48640.1 | Bradi3g53440.1 | HORVU6Hr1G070610.13 | TRIUR3_15225 | Seita.1G297100.1 | Sobic.004G259900.1 | Zm00001d017849_T001 |
| orthologID_009372 | PH01002105G0330 | LOC_Os02g43180.1 | Bradi3g50172.1 | HORVU6Hr1G061200.1 | TRIUR3_18604 | Seita.1G253200.1 | Sobic.004G300900.1 | Zm00001d017417_T001 |
| orthologID_009135 | PH01000114G1290 | LOC_Os02g05820.1 | Bradi3g04100.3 | HORVU6Hr1G022570.2 | TRIUR3_18693 | Seita.1G086800.1 | Sobic.004G042800.1 | Zm00001d053876_T011 |
| orthologID_009225 | PH01000773G0320 | LOC_Os02g02500.1 | Bradi3g01570.1 | HORVU0Hr1G040310.2 | TRIUR3_21892 | Seita.1G118700.1 | Sobic.004G012700.2 | Zm00001d054034_T001 |
| orthologID_009248 | PH01001226G0550 | LOC_Os02g46030.1 | Bradi3g51960.1 | HORVU6Hr1G066450.5 | TRIUR3_14822 | Seita.1G275400.1 | Sobic.004G279300.1 | Zm00001d017642_T001 |
| orthologID_009544 | PH01000355G0200 | LOC_Os02g51700.1 | Bradi3g58930.1 | HORVU6Hr1G074930.1 | TRIUR3_14599 | Seita.1G323800.1 | Sobic.004G232800.1 | Zm00001d052225_T001 |
| orthologID_009911 | PH01000672G0550 | LOC_Os02g46440.1 | Bradi3g52090.1 | HORVU6Hr1G066780.1 | TRIUR3_32746 | Seita.1G278300.1 | Sobic.004G276500.2 | Zm00001d017659_T001 |
| orthologID_009576 | PH01003547G0120 | LOC_Os02g17640.1 | Bradi3g10240.1 | HORVU7Hr1G025910.1 | TRIUR3_19726 | Seita.1G123700.1 | Sobic.004G125700.1 | Zm00001d015986_T002 |
| orthologID_009711 | PH01002448G0210 | LOC_Os02g33130.1 | Bradi3g45090.1 | HORVU6Hr1G070590.1 | TRIUR3_35385 | Seita.1G182800.1 | Sobic.004G167800.3 | Zm00001d016740_T001 |
| orthologID_009744 | PH01000706G0540 | LOC_Os02g50780.1 | Bradi3g59640.1 | HORVU6Hr1G076440.9 | TRIUR3_22041 | Seita.1G316100.1 | Sobic.004G242500.1 | Zm00001d017997_T010 |
| orthologID_009494 | PH01000483G0080 | LOC_Os02g30840.1 | Bradi3g44230.1 | HORVU5Hr1G106170.2 | TRIUR3_27238 | Seita.1G171200.1 | Sobic.004G156800.1 | Zm00001d016595_T014 |
| orthologID_009408 | PH01000120G1380 | LOC_Os02g10310.1 | Bradi3g07130.1 | HORVU6Hr1G033350.1 | TRIUR3_12626 | Seita.1G049000.1 | Sobic.004G080100.1 | Zm00001d053663_T011 |
| orthologID_009133 | PH01000210G0210 | LOC_Os02g53830.1 | Bradi3g57180.1 | HORVU6Hr1G081600.2 | TRIUR3_21960 | Seita.1G339800.1 | Sobic.004G318400.1 | Zm00001d018280_T001 |
| orthologID_009832 | PH01000062G1200 | LOC_Os02g48480.1 | Bradi3g53327.1 | HORVU6Hr1G070350.2 | TRIUR3_06526 | Seita.1G296000.1 | Sobic.004G261100.1 | Zm00001d051635_T001 |
| orthologID_009151 | PH01002513G0060 | LOC_Os02g14910.1 | Bradi3g09340.1 | HORVU5Hr1G045690.1 | TRIUR3_29751 | Seita.1G019600.1 | Sobic.004G107900.1 | Zm00001d016154_T001 |
| orthologID_009130 | PH01000036G2040 | LOC_Os02g56950.1 | Bradi3g55130.1 | HORVU6Hr1G090150.10 | TRIUR3_02290 | Seita.1G363000.1 | Sobic.004G342300.1 | Zm00001d051930_T002 |
| orthologID_009512 | PH01000197G0470 | LOC_Os02g47620.1 | Bradi3g52760.3 | HORVU6Hr1G068910.1 | TRIUR3_08032 | Seita.1G289300.1 | Sobic.004G267700.2 | Zm00001d051567_T001 |
| orthologID_009446 | PH01003043G0040 | LOC_Os02g58470.1 | Bradi3g60680.1 | HORVU6Hr1G094910.1 | TRIUR3_10347 | Seita.1G378500.1 | Sobic.004G357500.1 | Zm00001d018552_T002 |
| orthologID_007916 | PH01000069G1200 | LOC_Os02g06480.1 | Bradi1g33190.1 | HORVU7Hr1G101260.2 | TRIUR3_21641 | Seita.1G080200.1 | Sobic.004G048700.1 | Zm00001d015313_T001 |
| orthologID_009360 | PH01000127G1100 | LOC_Os02g20270.1 | Bradi3g11414.1 | HORVU5Hr1G111940.1 | TRIUR3_35060 | Seita.1G137400.1 | Sobic.004G137500.1 | Zm00001d053275_T001 |
| orthologID_009734 | PH01004130G0110 | LOC_Os02g40110.1 | Bradi3g48740.1 | HORVU6Hr1G057170.4 | TRIUR3_18089 | Seita.1G232600.1 | Sobic.004G214700.1 | Zm00001d017210_T001 |
| orthologID_009627 | PH01003540G0140 | LOC_Os02g17250.1 | Bradi3g10090.1 | HORVU6Hr1G051240.4 | TRIUR3_23635 | Seita.1G120900.1 | Sobic.004G123000.1 | Zm00001d053306_T001 |

| | | | | | | | | |
|---|---|---|---|---|---|---|---|---|
| orthologID_009693 | PH01000501G0350 | LOC_Os02g36870.1 | Bradi3g46890.1 | HORVU3Hr1G095850.1 | TRIUR3_08126 | Seita.1G208800.1 | Sobic.004G191500.1 | Zm00001d016949_T001 |
| orthologID_007568 | PH01002808G0140 | LOC_Os06g24730.1 | Bradi1g35780.1 | HORVU7Hr1G083370.9 | TRIUR3_23162 | Seita.7G261400.1 | Sobic.004G034600.1 | Zm00001d015205_T001 |
| orthologID_009244 | PH01001651G0320 | LOC_Os02g21520.1 | Bradi3g10937.2 | HORVU1Hr1G000430.4 | TRIUR3_28264 | Seita.1G143800.1 | Sobic.004G109000.1 | Zm00001d016144_T002 |
| orthologID_009100 | PH01000345G0560 | LOC_Os02g40700.1 | Bradi3g48940.1 | HORVU0Hr1G016180.2 | TRIUR3_28556 | Seita.1G237200.1 | Sobic.004G217700.1 | Zm00001d017248_T001 |
| orthologID_009093 | PH01000642G0470 | LOC_Os02g58460.1 | Bradi3g60667.1 | HORVU7Hr1G116590.1 | TRIUR3_01855 | Seita.1G378400.1 | Sobic.004G357400.1 | Zm00001d018545_T001 |
| orthologID_007679 | PH01001544G0140 | LOC_Os11g43960.1 | Bradi1g29260.1 | HORVU3Hr1G004820.22 | TRIUR3_00631 | Seita.1G045300.1 | Sobic.004G102800.1 | Zm00001d016230_T001 |
| orthologID_006834 | PH01001032G0100 | LOC_Os12g15590.2 | Bradi2g13367.1 | HORVU3Hr1G000130.1 | TRIUR3_25740 | Seita.3G273800.1 | Sobic.004G254800.3 | Zm00001d040941_T004 |
| orthologID_009805 | PH01001907G0340 | LOC_Os02g12760.1 | Bradi3g08210.1 | HORVU6Hr1G039660.1 | TRIUR3_35067 | Seita.1G034000.1 | Sobic.004G093800.1 | Zm00001d016363_T002 |
| orthologID_009729 | PH01000092G1040 | LOC_Os02g50560.1 | Bradi3g59790.4 | HORVU6Hr1G076860.2 | TRIUR3_12945 | Seita.1G314400.1 | Sobic.004G244400.3 | Zm00001d017982_T001 |
| orthologID_009125 | PH01002064G0120 | LOC_Os02g38680.1 | Bradi3g47867.1 | HORVU6Hr1G054070.1 | TRIUR3_34735 | Seita.1G221000.1 | Sobic.004G204200.1 | Zm00001d017114_T004 |
| orthologID_009491 | PH01000019G2280 | LOC_Os02g02710.1 | Bradi3g01750.1 | HORVU6Hr1G011160.8 | TRIUR3_07800 | Seita.1G111400.1 | Sobic.004G015800.1 | Zm00001d054011_T002 |
| orthologID_009681 | PH01001308G0210 | LOC_Os02g52940.1 | Bradi3g57820.2 | HORVU6Hr1G080160.1 | TRIUR3_11817 | Seita.1G333200.1 | Sobic.004G311100.1 | Zm00001d018197_T001 |
| orthologID_009339 | PH01000569G0700 | LOC_Os02g49950.1 | Bradi3g56960.1 | HORVU6Hr1G073280.1 | TRIUR3_26210 | Seita.1G309100.1 | Sobic.004G249100.1 | Zm00001d000381_T001 |
| orthologID_006702 | PH01000078G1430 | LOC_Os05g01500.1 | Bradi2g39750.1 | HORVU1Hr1G001890.20 | TRIUR3_07869 | Seita.3G070300.1 | Sobic.004G057100.1 | Zm00001d015389_T002 |
| orthologID_009108 | PH01002624G0160 | LOC_Os02g45870.1 | Bradi3g51850.1 | HORVU6Hr1G066210.12 | TRIUR3_16849 | Seita.1G274400.1 | Sobic.004G280200.1 | Zm00001d017620_T005 |
| orthologID_009813 | PH01001599G0330 | LOC_Os02g02880.1 | Bradi3g01970.2 | HORVU6Hr1G012530.3 | TRIUR3_05931 | Seita.1G109400.1 | Sobic.004G018300.1 | Zm00001d053909_T002 |
| orthologID_009827 | PH01002448G0220 | LOC_Os02g33240.1 | Bradi3g45050.1 | HORVU3Hr1G081740.1 | TRIUR3_12274 | Seita.1G182300.1 | Sobic.004G167300.1 | Zm00001d016745_T001 |
| orthologID_009418 | PH01002704G0080 | LOC_Os02g16909.2 | Bradi3g10020.1 | HORVU7Hr1G011400.2 | TRIUR3_12902 | Seita.1G007800.1 | Sobic.004G122500.1 | Zm00001d016016_T001 |
| orthologID_009563 | PH01002704G0220 | LOC_Os02g17190.1 | Bradi3g10067.3 | HORVU6Hr1G044640.8 | TRIUR3_33519 | Seita.1G007400.1 | Sobic.004G123400.2 | Zm00001d016000_T004 |
| orthologID_010098 | PH01005292G0070 | LOC_Os02g53000.1 | Bradi3g57756.2 | HORVU6Hr1G080030.2 | TRIUR3_26587 | Seita.1G333700.1 | Sobic.004G311600.1 | Zm00001d018201_T001 |
| orthologID_009435 | PH01000354G0430 | LOC_Os02g33020.1 | Bradi3g44930.2 | HORVU7Hr1G041080.5 | TRIUR3_22012 | Seita.1G180900.1 | Sobic.004G165900.1 | Zm00001d016707_T001 |
| orthologID_009085 | PH01001580G0170 | LOC_Os02g57250.1 | Bradi3g55410.1 | HORVU6Hr1G091260.4 | TRIUR3_01945 | Seita.1G366100.1 | Sobic.004G345600.1 | Zm00001d051911_T002 |
| orthologID_009704 | PH01001538G0330 | LOC_Os02g35690.1 | Bradi3g46210.1 | HORVU6Hr1G050140.2 | TRIUR3_32867 | Seita.1G201000.1 | Sobic.004G184400.1 | Zm00001d016876_T001 |
| orthologID_009628 | PH01003357G0110 | LOC_Os02g53500.1 | Bradi3g57410.1 | HORVU6Hr1G078830.4 | TRIUR3_21037 | Seita.1G337200.1 | Sobic.004G315500.1 | Zm00001d018243_T001 |
| orthologID_009807 | PH01000324G0500 | LOC_Os02g13360.1 | Bradi3g08610.2 | HORVU2Hr1G109960.26 | TRIUR3_13837 | Seita.1G030300.1 | Sobic.004G098000.2 | Zm00001d016286_T014 |
| orthologID_009591 | PH01002594G0130 | LOC_Os02g13300.1 | Bradi3g08557.2 | HORVU6Hr1G042080.3 | TRIUR3_26685 | Seita.1G030900.1 | Sobic.004G097400.1 | Zm00001d016299_T001 |
| orthologID_009756 | PH01000019G1730 | LOC_Os02g02350.1 | Bradi3g01360.4 | HORVU6Hr1G007030.3 | TRIUR3_11493 | Seita.1G116800.1 | Sobic.004G010900.3 | Zm00001d014986_T009 |
| orthologID_009737 | PH01000114G0040 | LOC_Os02g04800.1 | Bradi3g03420.2 | HORVU6Hr1G020320.5 | TRIUR3_19225 | Seita.1G091000.1 | Sobic.004G037700.1 | Zm00001d015242_T002 |
| orthologID_009718 | PH01000251G0570 | LOC_Os02g35620.1 | Bradi3g46140.1 | HORVU6Hr1G009280.4 | TRIUR3_05280 | Seita.1G199400.1 | Sobic.004G183300.1 | Zm00001d016865_T001 |
| orthologID_010142 | PH01000210G0740 | LOC_Os02g54050.1 | Bradi3g60018.1 | HORVU6Hr1G081980.1 | TRIUR3_18975 | Seita.1G341100.1 | Sobic.004G319700.1 | Zm00001d052102_T001 |
| orthologID_009554 | PH01000140G1390 | LOC_Os02g57510.2 | Bradi3g55710.2 | HORVU6Hr1G091720.6 | TRIUR3_02303 | Seita.1G369900.1 | Sobic.004G349100.1 | Zm00001d018507_T002 |
| orthologID_009509 | PH01001246G0580 | LOC_Os02g49420.1 | Bradi3g56390.1 | HORVU6Hr1G072160.1 | TRIUR3_32711 | Seita.1G303000.1 | Sobic.004G254300.1 | Zm00001d051699_T001 |
| orthologID_009549 | PH01001973G0150 | LOC_Os02g53320.1 | Bradi3g57525.1 | HORVU6Hr1G079150.1 | TRIUR3_20184 | Seita.1G336200.1 | Sobic.004G314400.1 | Zm00001d018229_T001 |
| orthologID_009731 | PH01000835G0430 | LOC_Os02g36490.2 | Bradi3g46650.2 | HORVU2Hr1G107660.8 | TRIUR3_06137 | Seita.1G205200.1 | Sobic.004G188800.2 | Zm00001d016922_T004 |
| orthologID_009486 | PH01000019G1400 | LOC_Os02g02000.1 | Bradi3g01110.1 | HORVU6Hr1G005250.1 | TRIUR3_04662 | Seita.1G007200.1 | Sobic.004G007500.1 | Zm00001d054067_T001 |
| orthologID_009505 | PH01000197G0580 | LOC_Os02g47570.1 | Bradi3g52697.3 | HORVU6Hr1G068790.4 | TRIUR3_30575 | Seita.1G288700.1 | Sobic.004G268300.1 | Zm00001d017767_T001 |
| orthologID_009286 | PH01001539G0160 | LOC_Os02g10020.1 | Bradi3g06840.1 | HORVU6Hr1G034110.4 | TRIUR3_06420 | Seita.1G052600.1 | Sobic.004G077100.1 | Zm00001d015863_T001 |
| orthologID_009084 | PH01000021G1440 | LOC_Os02g57370.1 | Bradi3g55510.1 | HORVU6Hr1G091290.1 | TRIUR3_35221 | Seita.1G368000.1 | Sobic.004G347100.1 | Zm00001d018494_T002 |
| orthologID_009743 | PH01001936G0180 | LOC_Os02g12570.1 | Bradi3g08080.2 | HORVU6Hr1G038740.5 | TRIUR3_09742 | Seita.1G032400.1 | Sobic.004G092400.1 | Zm00001d053591_T002 |
| orthologID_009202 | PH01000355G0510 | LOC_Os02g51810.1 | Bradi3g58760.1 | HORVU6Hr1G078210.1 | TRIUR3_23396 | Seita.1G324900.1 | Sobic.004G231600.1 | Zm00001d018098_T005 |
| orthologID_009507 | PH01000585G0580 | LOC_Os02g03690.1 | Bradi3g02800.1 | HORVU6Hr1G015670.15 | TRIUR3_33568 | Seita.1G099800.1 | Sobic.004G027800.2 | Zm00001d053975_T003 |
| orthologID_009431 | PH01000092G0230 | LOC_Os02g50210.1 | Bradi3g60000.1 | HORVU6Hr1G073910.1 | TRIUR3_09875 | Seita.1G311200.1 | Sobic.004G247200.1 | Zm00001d017956_T001 |
| orthologID_009768 | PH01003506G0010 | LOC_Os02g54200.1 | Bradi3g60150.1 | HORVU6Hr1G082970.1 | TRIUR3_08930 | Seita.1G343400.1 | Sobic.004G321700.2 | Zm00001d052081_T003 |
| orthologID_009332 | PH01000608G0850 | LOC_Os02g08110.1 | Bradi3g05755.1 | HORVU6Hr1G030310.1 | TRIUR3_04884 | Seita.1G065700.1 | Sobic.004G063000.1 | Zm00001d015463_T001 |
| orthologID_009624 | PH01000062G0620 | LOC_Os02g48130.1 | Bradi3g53130.1 | HORVU0Hr1G016930.9 | TRIUR3_14937 | Seita.1G293700.1 | Sobic.004G263200.1 | Zm00001d017812_T002 |
| orthologID_009897 | PH01000197G0520 | LOC_Os02g47600.2 | Bradi3g52740.2 | HORVU6Hr1G068760.5 | TRIUR3_20171 | Seita.1G289000.1 | Sobic.004G268000.1 | Zm00001d017770_T001 |
| orthologID_009210 | PH01029494G0010 | LOC_Os02g47840.1 | Bradi3g52900.3 | HORVU6Hr1G069260.1 | TRIUR3_20108 | Seita.1G291000.1 | Sobic.004G266000.1 | Zm00001d051584_T001 |
| orthologID_009819 | PH01000217G0880 | LOC_Os02g17500.3 | Bradi3g10210.3 | HORVU7Hr1G115370.2 | TRIUR3_00206 | Seita.1G123000.1 | Sobic.004G124500.1 | Zm00001d053334_T001 |
| orthologID_009434 | PH01001738G0130 | LOC_Os02g42890.1 | Bradi3g50010.2 | HORVU6Hr1G060720.1 | TRIUR3_02969 | Seita.1G251100.1 | Sobic.004G303200.1 | Zm00001d051272_T001 |
| orthologID_009670 | PH01000019G1410 | LOC_Os02g02020.1 | Bradi3g01116.1 | HORVU6Hr1G005240.3 | TRIUR3_04661 | Seita.1G120100.1 | Sobic.004G007700.1 | Zm00001d014966_T001 |

| | | | | | | | | |
|---|---|---|---|---|---|---|---|---|
| orthologID_009104 | PH01001295G0130 | LOC_Os02g47190.1 | Bradi3g52500.2 | HORVU6Hr1G068100.2 | TRIUR3_08190 | Seita.1G286400.1 | Sobic.004G270600.1 | Zm00001d017726_T001 |
| orthologID_009738 | PH01003122G0060 | LOC_Os02g48000.1 | Bradi3g53040.1 | HORVU6Hr1G069570.7 | TRIUR3_26184 | Seita.1G292600.1 | Sobic.004G264400.1 | Zm00001d051599_T004 |
| orthologID_009872 | PH01004969G0020 | LOC_Os02g49470.1 | Bradi3g56350.1 | HORVU6Hr1G072310.1 | TRIUR3_26425 | Seita.1G303800.1 | Sobic.004G253400.1 | Zm00001d017907_T001 |
| orthologID_009668 | PH01003425G0180 | LOC_Os02g38950.1 | Bradi3g47936.1 | HORVU6Hr1G054540.1 | TRIUR3_31575 | Seita.1G221900.1 | Sobic.004G205200.1 | Zm00001d017124_T001 |
| orthologID_009871 | PH01002569G0310 | LOC_Os08g23320.1 | Bradi3g60400.6 | HORVU6Hr1G094110.3 | TRIUR3_07224 | Seita.1G376500.1 | Sobic.004G355300.2 | Zm00001d051848_T002 |
| orthologID_009809 | PH01000218G0360 | LOC_Os02g43600.1 | Bradi3g50390.1 | HORVU6Hr1G062040.3 | TRIUR3_13325 | Seita.1G256000.1 | Sobic.004G298100.1 | Zm00001d017449_T001 |
| orthologID_009503 | PH01001610G0530 | LOC_Os02g44220.1 | Bradi3g50800.1 | HORVU6Hr1G063300.2 | TRIUR3_28283 | Seita.1G261900.1 | Sobic.004G293700.1 | Zm00001d017501_T002 |
| orthologID_009534 | PH01001007G0110 | LOC_Os02g47450.1 | Bradi3g52650.2 | HORVU0Hr1G023150.6 | TRIUR3_28357 | Seita.1G288000.1 | Sobic.004G268900.1 | Zm00001d017761_T002 |
| orthologID_009658 | PH01000197G1290 | LOC_Os02g47310.1 | Bradi3g52560.1 | HORVU6Hr1G068410.1 | TRIUR3_32581 | Seita.1G287200.1 | Sobic.004G269800.1 | Zm00001d017746_T001 |
| orthologID_009250 | PH01000043G1650 | LOC_Os02g08540.1 | Bradi3g06160.1 | HORVU6Hr1G028660.1 | TRIUR3_29863 | Seita.1G060700.1 | Sobic.004G067000.1 | Zm00001d015546_T001 |
| orthologID_009115 | PH01001562G0090 | LOC_Os02g49480.1 | Bradi3g56340.2 | HORVU6Hr1G072300.14 | TRIUR3_26424 | Seita.1G303900.1 | Sobic.004G253300.1 | Zm00001d017908_T001 |
| orthologID_009545 | PH01003284G0180 | LOC_Os02g40900.1 | Bradi3g49130.1 | HORVU6Hr1G058450.9 | TRIUR3_13561 | Seita.1G239000.1 | Sobic.004G219400.1 | Zm00001d051139_T002 |
| orthologID_009829 | PH01001515G0120 | LOC_Os02g26440.1 | Bradi3g43090.1 | HORVU2Hr1G057210.11 | TRIUR3_28073 | Seita.1G154600.1 | Sobic.004G138800.1 | Zm00001d016458_T002 |
| orthologID_007055 | PH01000356G0530 | LOC_Os02g57060.1 | Bradi4g37555.1 | HORVU5Hr1G088200.8 | TRIUR3_08102 | Seita.1G364200.1 | Sobic.004G343500.1 | Zm00001d018470_T001 |
| orthologID_009161 | PH01000466G0910 | LOC_Os02g57380.1 | Bradi3g55530.1 | HORVU6Hr1G091330.7 | TRIUR3_08414 | Seita.1G368100.1 | Sobic.004G347300.2 | Zm00001d018495_T001 |
| orthologID_007033 | PH01004038G0020 | LOC_Os12g16210.1 | Bradi4g39300.1 | HORVU7Hr1G085240.2 | TRIUR3_07976 | Seita.3G297400.1 | Sobic.004G257600.1 | Zm00001d017871_T002 |
| orthologID_009119 | PH01007485G0020 | LOC_Os02g19804.1 | Bradi3g09570.1 | HORVU4Hr1G062570.1 | TRIUR3_18095 | Seita.1G014000.1 | Sobic.004G113700.1 | Zm00001d053397_T001 |
| orthologID_009260 | PH01001872G0180 | LOC_Os02g44740.1 | Bradi3g51270.1 | HORVU6Hr1G064440.2 | TRIUR3_17588 | Seita.1G266300.1 | Sobic.004G287500.2 | Zm00001d017537_T001 |
| orthologID_009830 | PH01001295G0530 | LOC_Os02g47120.1 | Bradi3g52437.2 | HORVU6Hr1G067890.26 | TRIUR3_08834 | Seita.1G284500.1 | Sobic.004G271600.1 | Zm00001d051545_T003 |
| orthologID_009447 | PH01002010G0240 | LOC_Os02g58650.1 | Bradi3g60850.1 | HORVU6Hr1G095250.2 | TRIUR3_04195 | Seita.1G379900.1 | Sobic.004G359100.2 | Zm00001d051823_T001 |
| orthologID_009587 | PH01000135G1520 | LOC_Os02g51030.1 | Bradi3g59480.1 | HORVU0Hr1G017140.11 | TRIUR3_26777 | Seita.1G317900.1 | Sobic.004G238900.1 | Zm00001d052266_T001 |
| orthologID_009908 | PH01004036G0180 | LOC_Os02g10900.1 | Bradi3g07580.4 | HORVU6Hr1G037680.1 | TRIUR3_23370 | Seita.1G043600.1 | Sobic.004G085900.2 | Zm00001d053623_T001 |
| orthologID_009141 | PH01000092G1670 | LOC_Os02g50830.1 | Bradi3g59590.2 | HORVU6Hr1G076310.2 | TRIUR3_24873 | Seita.1G316500.1 | Sobic.004G242100.1 | Zm00001d052276_T005 |
| orthologID_009548 | PH01000062G1170 | LOC_Os02g48450.1 | Bradi3g53300.2 | HORVU6Hr1G070320.1 | TRIUR3_06523 | Seita.1G295700.1 | Sobic.004G261400.1 | Zm00001d017834_T001 |
| orthologID_009715 | PH01000049G1170 | LOC_Os02g51500.1 | Bradi3g59100.1 | HORVU6Hr1G075180.1 | TRIUR3_17073 | Seita.1G322200.1 | Sobic.004G234900.1 | Zm00001d018072_T001 |
| orthologID_009784 | PH01000642G0510 | LOC_Os02g58510.1 | Bradi3g60720.1 | HORVU6Hr1G094950.1 | TRIUR3_03615 | Seita.1G378900.1 | Sobic.004G357900.1 | Zm00001d018555_T001 |
| orthologID_009147 | PH01000062G1350 | LOC_Os02g48590.1 | Bradi3g53390.1 | HORVU6Hr1G070500.2 | TRIUR3_21079 | Seita.1G296500.1 | Sobic.004G260400.1 | Zm00001d017845_T001 |
| orthologID_009540 | PH01000642G0120 | LOC_Os02g58300.1 | Bradi3g60510.1 | HORVU6Hr1G094400.1 | TRIUR3_07307 | Seita.1G377100.1 | Sobic.004G356300.1 | Zm00001d018534_T001 |
| orthologID_009614 | PH01000021G1210 | LOC_Os02g57460.1 | Bradi3g55650.3 | HORVU6Hr1G091610.1 | TRIUR3_17529 | Seita.1G369300.1 | Sobic.004G348500.1 | Zm00001d051894_T001 |
| orthologID_009359 | PH01000123G0190 | LOC_Os02g07720.2 | Bradi3g05420.1 | HORVU6Hr1G031590.2 | TRIUR3_26638 | Seita.1G069900.1 | Sobic.004G058200.1 | Zm00001d015399_T002 |
| orthologID_009515 | PH01000952G0380 | LOC_Os02g21650.1 | Bradi3g10900.1 | HORVU7Hr1G091920.1 | TRIUR3_12830 | Seita.1G144500.1 | Sobic.004G117300.1 | Zm00001d016060_T001 |
| orthologID_009686 | PH01000437G0390 | LOC_Os02g52670.1 | Bradi3g58015.1 | HORVU6Hr1G078520.2 | TRIUR3_03778 | Seita.1G330800.1 | Sobic.004G308900.1 | Zm00001d018158_T001 |
| orthologID_009423 | PH01002102G0240 | LOC_Os03g11990.3 | Bradi3g49671.1 | HORVU6Hr1G059920.2 | TRIUR3_04312 | Seita.1G246300.1 | Sobic.004G225500.1 | Zm00001d017347_T001 |
| orthologID_009596 | PH01000095G0770 | LOC_Os02g07420.1 | Bradi3g05247.1 | HORVU0Hr1G018370.1 | TRIUR3_17946 | Seita.1G072600.1 | Sobic.004G055800.1 | Zm00001d015379_T001 |
| orthologID_009788 | PH01001392G0200 | LOC_Os02g13640.1 | Bradi3g08770.1 | HORVU6Hr1G041650.2 | TRIUR3_24291 | Seita.1G027400.1 | Sobic.004G100200.1 | Zm00001d053555_T001 |
| orthologID_009722 | PH01002092G0430 | LOC_Os02g50050.1 | Bradi3g56861.1 | HORVU6Hr1G073560.1 | TRIUR3_19657 | Seita.1G310300.1 | Sobic.004G248000.1 | Zm00001d051793_T001 |
| orthologID_009707 | PH01000311G0070 | LOC_Os02g53400.1 | Bradi3g57470.1 | HORVU6Hr1G078960.8 | TRIUR3_11825 | Seita.1G336800.1 | Sobic.004G315100.1 | Zm00001d018238_T001 |
| orthologID_009533 | PH01000019G0150 | LOC_Os02g32860.1 | Bradi3g44840.1 | HORVU1Hr1G077820.10 | TRIUR3_01106 | Seita.1G180000.1 | Sobic.004G164800.1 | Zm00001d016694_T003 |
| orthologID_009405 | PH01002624G0180 | LOC_Os02g45880.1 | Bradi3g51860.1 | HORVU6Hr1G066250.1 | TRIUR3_16848 | Seita.1G274600.1 | Sobic.004G280100.1 | Zm00001d017621_T001 |
| orthologID_009287 | PH01003332G0200 | LOC_Os02g02970.1 | Bradi3g02160.3 | HORVU6Hr1G012060.3 | TRIUR3_05464 | Seita.1G108100.1 | Sobic.004G019700.1 | Zm00001d015059_T001 |
| orthologID_009249 | PH01000197G0440 | LOC_Os02g47650.1 | Bradi3g52780.1 | HORVU6Hr1G068990.2 | TRIUR3_17847 | Seita.1G289500.1 | Sobic.004G267500.1 | Zm00001d017777_T003 |
| orthologID_009853 | PH01004961G0020 | LOC_Os02g18480.1 | Bradi3g10550.1 | HORVU3Hr1G039760.1 | TRIUR3_23015 | Seita.1G128800.1 | Sobic.004G129100.1 | Zm00001d053348_T001 |
| orthologID_009369 | PH01000569G0270 | LOC_Os02g49700.1 | Bradi3g56180.1 | HORVU6Hr1G072810.6 | TRIUR3_23073 | Seita.1G306100.1 | Sobic.004G251300.1 | Zm00001d017920_T001 |
| orthologID_009462 | PH01002053G0300 | LOC_Os02g48116.1 | Bradi3g53107.3 | HORVU0Hr1G016880.9 | TRIUR3_19470 | Seita.1G293500.1 | Sobic.004G263400.2 | Zm00001d017810_T010 |
| orthologID_009556 | PH01000437G0570 | LOC_Os02g52540.1 | Bradi3g58086.1 | HORVU6Hr1G078670.1 | TRIUR3_05364 | Seita.1G329900.1 | Sobic.004G308000.1 | Zm00001d052172_T001 |
| orthologID_009421 | PH01000437G0280 | LOC_Os02g52744.1 | Bradi3g57980.1 | HORVU0Hr1G017250.4 | TRIUR3_06446 | Seita.1G331400.1 | Sobic.004G309400.1 | Zm00001d052163_T001 |
| orthologID_009155 | PH01001247G0490 | LOC_Os02g11940.1 | Bradi3g07997.4 | HORVU6Hr1G038500.17 | TRIUR3_34024 | Seita.1G039600.1 | Sobic.004G089000.1 | Zm00001d016397_T008 |
| orthologID_009488 | PH01005517G0020 | LOC_Os02g22070.1 | Bradi3g10760.2 | HORVU7Hr1G045830.2 | TRIUR3_11518 | Seita.1G129500.1 | Sobic.004G066300.1 | Zm00001d015519_T001 |
| orthologID_009817 | PH01001160G0560 | LOC_Os02g35750.1 | Bradi3g46270.3 | HORVU1Hr1G012440.1 | TRIUR3_20537 | Seita.1G201500.1 | Sobic.004G184900.1 | Zm00001d016878_T001 |
| orthologID_009353 | PH01000049G1410 | LOC_Os02g51570.1 | Bradi3g59057.1 | HORVU6Hr1G075080.4 | TRIUR3_05164 | Seita.1G322800.1 | Sobic.004G234200.3 | Zm00001d018077_T002 |

| | | | | | | | | |
|---|---|---|---|---|---|---|---|---|
| orthologID_009168 | PH01006137G0060 | LOC_Os02g33180.1 | Bradi3g45030.1 | HORVU2Hr1G097800.8 | TRIUR3_34558 | Seita.1G182100.1 | Sobic.004G167100.1 | Zm00001d050655_T001 |
| orthologID_009613 | PH01000457G0590 | LOC_Os02g03050.2 | Bradi3g02232.1 | HORVU6Hr1G012160.1 | TRIUR3_13179 | Seita.1G106900.1 | Sobic.004G020800.2 | Zm00001d053927_T021 |
| orthologID_009543 | PH01002577G0020 | LOC_Os02g15250.1 | Bradi3g09480.2 | HORVU1Hr1G082820.2 | TRIUR3_08536 | Seita.1G015800.1 | Sobic.004G111500.1 | Zm00001d016128_T001 |
| orthologID_009242 | PH01001907G0160 | LOC_Os02g12790.1 | Bradi3g08240.3 | HORVU6Hr1G039720.5 | TRIUR3_13697 | Seita.1G034200.1 | Sobic.004G094100.1 | Zm00001d016361_T001 |
| orthologID_009314 | PH01000608G0130 | LOC_Os02g07850.1 | Bradi3g05583.1 | HORVU0Hr1G015650.2 | TRIUR3_19434 | Seita.1G068500.1 | Sobic.004G060000.1 | Zm00001d053766_T001 |
| orthologID_009490 | PH01002010G0330 | LOC_Os02g58790.1 | Bradi3g60920.1 | HORVU6Hr1G095380.12 | TRIUR3_27990 | Seita.1G380300.1 | Sobic.004G359600.1 | Zm00001d018575_T001 |
| orthologID_009716 | PH01001226G0110 | LOC_Os02g46140.1 | Bradi3g52000.4 | HORVU6Hr1G066560.1 | TRIUR3_07943 | Seita.1G276000.1 | Sobic.004G278600.1 | Zm00001d051499_T001 |
| orthologID_009611 | PH01004821G0050 | LOC_Os02g44810.1 | Bradi3g51240.1 | HORVU6Hr1G064480.1 | TRIUR3_08588 | Seita.1G266700.1 | Sobic.004G287100.1 | Zm00001d017544_T002 |
| orthologID_007474 | PH01000105G0750 | LOC_Os12g41380.1 | Bradi4g02360.4 | HORVU5Hr1G012790.18 | TRIUR3_28699 | Seita.3G382600.1 | Sobic.004G020900.1 | Zm00001d053928_T001 |
| orthologID_009302 | PH01000743G0080 | LOC_Os02g40460.1 | Bradi3g48860.1 | HORVU6Hr1G057570.8 | TRIUR3_09078 | Seita.1G235700.1 | Sobic.004G215900.1 | Zm00001d017216_T001 |
| orthologID_009513 | PH01000140G0100 | LOC_Os02g58280.1 | Bradi3g60477.2 | HORVU6Hr1G094240.3 | TRIUR3_03452 | Seita.1G377000.1 | Sobic.004G356200.1 | Zm00001d018533_T003 |
| orthologID_009607 | PH01000089G0450 | LOC_Os02g39290.1 | Bradi3g48250.3 | HORVU6Hr1G055570.18 | TRIUR3_16236 | Seita.1G225900.1 | Sobic.004G209000.1 | Zm00001d017157_T001 |
| orthologID_009669 | PH01000123G0470 | LOC_Os02g07670.1 | Bradi3g05390.3 | HORVU6Hr1G031610.4 | TRIUR3_34150 | Seita.1G070700.1 | Sobic.004G057800.1 | Zm00001d015395_T001 |
| orthologID_009247 | PH01000021G0790 | LOC_Os02g57630.1 | Bradi3g55860.1 | HORVU6Hr1G092480.4 | TRIUR3_05876 | Seita.1G371300.1 | Sobic.004G350100.3 | Zm00001d018512_T001 |
| orthologID_009180 | PH01000642G0070 | LOC_Os02g58250.1 | Bradi3g60460.1 | HORVU0Hr1G017800.3 | TRIUR3_06764 | Seita.1G376800.1 | Sobic.004G356000.1 | Zm00001d018532_T001 |
| orthologID_009095 | PH01000773G0480 | LOC_Os02g02390.1 | Bradi3g01450.1 | HORVU6Hr1G008590.4 | TRIUR3_06495 | Seita.1G117300.1 | Sobic.004G011300.1 | Zm00001d014991_T004 |
| orthologID_009267 | PH01001018G0370 | LOC_Os02g29464.1 | Bradi3g43737.7 | HORVU5Hr1G020580.2 | TRIUR3_18330 | Seita.1G165000.1 | Sobic.004G151000.1 | Zm00001d050612_T001 |
| orthologID_009404 | PH01000355G0380 | LOC_Os02g51770.1 | Bradi3g58800.3 | HORVU6Hr1G074360.11 | TRIUR3_17313 | Seita.1G324500.1 | Sobic.004G232000.1 | Zm00001d052219_T002 |
| orthologID_009528 | PH01000514G0680 | LOC_Os02g12600.1 | Bradi3g08100.1 | HORVU6Hr1G038870.11 | TRIUR3_34639 | Seita.1G032600.1 | Sobic.004G092600.1 | Zm00001d016374_T002 |
| orthologID_009132 | PH01000343G0340 | LOC_Os02g43810.1 | Bradi3g50497.1 | HORVU6Hr1G062410.3 | TRIUR3_33734 | Seita.1G257500.1 | Sobic.004G297000.4 | Zm00001d051339_T001 |
| orthologID_009786 | PH01000049G0720 | LOC_Os02g51450.2 | Bradi3g59167.14 | HORVU6Hr1G075340.1 | TRIUR3_31159 | Seita.1G321400.1 | Sobic.004G235600.1 | Zm00001d018066_T001 |
| orthologID_009243 | PH01000026G1290 | LOC_Os02g55630.1 | Bradi3g54350.1 | HORVU6Hr1G086970.1 | TRIUR3_15918 | Seita.1G353900.1 | Sobic.004G333400.1 | Zm00001d018401_T001 |
| orthologID_009273 | PH01000173G0200 | LOC_Os02g52260.1 | Bradi3g58290.1 | HORVU6Hr1G077440.3 | TRIUR3_12498 | Seita.1G327800.1 | Sobic.004G305800.1 | Zm00001d018130_T001 |
| orthologID_008711 | PH01003036G0080 | LOC_Os06g21590.1 | Bradi1g42670.1 | HORVU7Hr1G046320.1 | TRIUR3_22828 | Seita.4G139900.1 | Sobic.004G056900.1 | Zm00001d015385_T001 |
| orthologID_009245 | PH01001872G0190 | LOC_Os02g44730.1 | Bradi3g51280.1 | HORVU6Hr1G064430.3 | TRIUR3_17589 | Seita.1G266200.1 | Sobic.004G287600.1 | Zm00001d017536_T001 |
| orthologID_009691 | PH01000280G1020 | LOC_Os02g03450.1 | Bradi3g02630.1 | HORVU6Hr1G014800.4 | TRIUR3_01145 | Seita.1G101800.1 | Sobic.004G025300.1 | Zm00001d015102_T001 |
| orthologID_009761 | PH01003006G0150 | LOC_Os02g30910.1 | Bradi3g44260.1 | HORVU7Hr1G030160.4 | TRIUR3_28114 | Seita.1G171800.1 | Sobic.004G157100.1 | Zm00001d050577_T001 |
| orthologID_009416 | PH01000791G0820 | LOC_Os02g10760.1 | Bradi3g07450.1 | HORVU6Hr1G038120.7 | TRIUR3_22374 | Seita.1G044700.1 | Sobic.004G084600.1 | Zm00001d015759_T001 |
| orthologID_009328 | PH01001462G0110 | LOC_Os02g39600.1 | Bradi3g48410.1 | HORVU6Hr1G055870.1 | TRIUR3_34044 | Seita.1G227800.1 | Sobic.004G210500.1 | Zm00001d017171_T001 |
| orthologID_009539 | PH01000731G0530 | LOC_Os02g37970.1 | Bradi3g47530.1 | HORVU6Hr1G053000.2 | TRIUR3_30202 | Seita.1G215900.1 | Sobic.004G199600.1 | Zm00001d017065_T001 |
| orthologID_009687 | PH01000145G1320 | LOC_Os02g03950.3 | Bradi3g02970.17 | HORVU6Hr1G017680.3 | TRIUR3_20670 | Seita.1G097100.1 | Sobic.004G030900.1 | Zm00001d015152_T006 |
| orthologID_009170 | PH01003332G0190 | LOC_Os02g02960.4 | Bradi3g02150.1 | HORVU6Hr1G012050.4 | TRIUR3_05463 | Seita.1G108200.1 | Sobic.004G019600.1 | Zm00001d015058_T001 |
| orthologID_009099 | PH01084788G0010 | LOC_Os02g34540.1 | Bradi3g45535.1 | HORVU6Hr1G059720.1 | TRIUR3_23443 | Seita.1G189400.1 | Sobic.004G172600.1 | Zm00001d016767_T001 |
| orthologID_009368 | PH01000345G0740 | LOC_Os02g40530.1 | Bradi3g48886.2 | HORVU6Hr1G057670.3 | TRIUR3_31527 | Seita.1G236400.1 | Sobic.004G216900.1 | Zm00001d017243_T001 |
| orthologID_009482 | PH01002043G0130 | LOC_Os02g35900.1 | Bradi3g46420.1 | HORVU2Hr1G124800.1 | TRIUR3_09417 | Seita.1G203000.1 | Sobic.004G186200.1 | Zm00001d016893_T001 |
| orthologID_009160 | PH01003604G0060 | LOC_Os02g49570.1 | Bradi3g56280.1 | HORVU6Hr1G072540.8 | TRIUR3_34994 | Seita.1G304700.1 | Sobic.004G252500.1 | Zm00001d051740_T005 |
| orthologID_009518 | PH01000280G0520 | LOC_Os04g35305.1 | Bradi3g02795.2 | HORVU6Hr1G015690.7 | TRIUR3_21743 | Seita.1G099900.1 | Sobic.004G027700.2 | Zm00001d015124_T001 |
| orthologID_009818 | PH01000093G1530 | LOC_Os02g06560.1 | Bradi3g04585.1 | HORVU6Hr1G025830.1 | TRIUR3_10130 | Seita.1G079500.1 | Sobic.004G049400.1 | Zm00001d053843_T001 |
| orthologID_009903 | PH01000203G0690 | LOC_Os02g55000.1 | Bradi3g53880.1 | HORVU6Hr1G084530.2 | TRIUR3_21941 | Seita.1G348300.1 | Sobic.004G327400.1 | Zm00001d018360_T001 |
| orthologID_010177 | PH01003715G0090 | LOC_Os02g09180.1 | Bradi3g06250.1 | HORVU6Hr1G028290.2 | TRIUR3_13974 | Seita.1G060300.1 | Sobic.004G068500.1 | Zm00001d015571_T002 |
| orthologID_009271 | PH01000019G0750 | LOC_Os02g01440.1 | Bradi3g00690.1 | HORVU6Hr1G002550.3 | TRIUR3_01271 | Seita.1G003800.1 | Sobic.004G003900.1 | Zm00001d054079_T002 |
| orthologID_009281 | PH01000628G0100 | LOC_Os02g16030.1 | Bradi3g09780.1 | HORVU1Hr1G037110.2 | TRIUR3_27352 | Seita.1G011800.1 | Sobic.004G116800.1 | Zm00001d053352_T001 |
| orthologID_009526 | PH01002823G0300 | LOC_Os02g49780.1 | Bradi3g57060.1 | HORVU6Hr1G072940.1 | TRIUR3_21159 | Seita.1G307600.1 | Sobic.004G250500.1 | Zm00001d017927_T001 |
| orthologID_007774 | PH01001221G0570 | LOC_Os04g38430.2 | Bradi1g21860.1 | HORVU6Hr1G000600.1 | TRIUR3_18370 | Seita.1G003300.1 | Sobic.004G003100.1 | Zm00001d054081_T001 |
| orthologID_009212 | PH01002064G0190 | LOC_Os02g38470.1 | Bradi3g47847.1 | HORVU6Hr1G054050.7 | TRIUR3_34439 | Seita.1G220700.1 | Sobic.004G204000.1 | Zm00001d017112_T001 |
| orthologID_009778 | PH01000140G1400 | LOC_Os02g57500.1 | Bradi3g55690.1 | HORVU0Hr1G017680.1 | TRIUR3_01476 | Seita.1G369700.1 | Sobic.004G348900.1 | Zm00001d018504_T001 |
| orthologID_009630 | PH01000135G0570 | LOC_Os02g51300.1 | Bradi3g59300.1 | HORVU6Hr1G075640.21 | TRIUR3_24654 | Seita.1G319700.1 | Sobic.004G237100.1 | Zm00001d018050_T003 |
| orthologID_009574 | PH01000628G0560 | LOC_Os02g15900.1 | Bradi3g09730.2 | HORVU6Hr1G041020.1 | TRIUR3_20852 | Seita.1G012400.1 | Sobic.004G116100.1 | Zm00001d016072_T001 |
| orthologID_009798 | PH01001874G0300 | LOC_Os02g39820.1 | Bradi3g48510.1 | HORVU6Hr1G056180.4 | TRIUR3_12616 | Seita.1G229700.1 | Sobic.004G212100.1 | Zm00001d017185_T001 |
| orthologID_009511 | PH01000218G0100 | LOC_Os02g43560.1 | Bradi3g50360.1 | HORVU6Hr1G061940.2 | TRIUR3_27182 | Seita.1G255500.1 | Sobic.004G298400.1 | Zm00001d017444_T001 |

| | | | | | | | | |
|---|---|---|---|---|---|---|---|---|
| orthologID_006344 | PH01002799G0090 | LOC_Os06g27970.1 | Bradi2g03820.1 | HORVU7Hr1G048110.2 | TRIUR3_25122 | Seita.1G202000.1 | Sobic.004G033100.1 | Zm00001d015194_T001 |
| orthologID_009896 | PH01000049G1110 | LOC_Os02g51470.3 | Bradi3g59130.1 | HORVU6Hr1G075280.1 | TRIUR3_16628 | Seita.1G321900.1 | Sobic.004G235200.1 | Zm00001d052242_T001 |
| orthologID_008381 | PH01003342G0140 | LOC_Os02g54624.1 | Bradi1g70460.1 | HORVU6Hr1G083430.1 | TRIUR3_16496 | Seita.1G345800.1 | Sobic.004G323700.3 | Zm00001d018328_T003 |
| orthologID_009514 | PH01000356G0520 | LOC_Os02g57050.1 | Bradi3g55230.1 | HORVU6Hr1G090270.2 | TRIUR3_31383 | Seita.1G364100.1 | Sobic.004G343400.1 | Zm00001d018469_T001 |
| orthologID_009221 | PH01000569G0190 | LOC_Os02g49620.1 | Bradi3g56230.1 | HORVU6Hr1G072640.1 | TRIUR3_03913 | Seita.1G305200.1 | Sobic.004G252000.1 | Zm00001d017915_T001 |
| orthologID_009562 | PH01001563G0300 | LOC_Os02g58139.2 | Bradi3g60300.4 | HORVU6Hr1G087430.7 | TRIUR3_04820 | Seita.1G375500.1 | Sobic.004G354000.1 | Zm00001d018528_T001 |
| orthologID_009701 | PH01000049G2030 | LOC_Os02g51760.1 | Bradi3g58820.1 | HORVU6Hr1G074410.1 | TRIUR3_17314 | Seita.1G324400.1 | Sobic.004G232100.1 | Zm00001d018093_T001 |
| orthologID_009816 | PH01004036G0040 | LOC_Os02g10920.1 | Bradi3g07610.7 | HORVU6Hr1G037610.3 | TRIUR3_27781 | Seita.1G043500.1 | Sobic.004G086000.1 | Zm00001d053622_T002 |
| orthologID_009355 | PH01002225G0300 | LOC_Os02g17330.1 | Bradi3g10127.1 | HORVU6Hr1G035160.2 | TRIUR3_11895 | Seita.1G121100.1 | Sobic.004G122800.1 | Zm00001d016006_T001 |
| orthologID_009191 | PH01000345G0470 | LOC_Os02g40750.1 | Bradi3g48960.1 | HORVU0Hr1G015970.1 | TRIUR3_24347 | Seita.1G237400.1 | Sobic.004G217900.2 | Zm00001d017250_T001 |
| orthologID_009649 | PH01000140G0540 | LOC_Os02g58020.1 | Bradi3g56780.1 | HORVU6Hr1G085890.2 | TRIUR3_22569 | Seita.1G374500.1 | Sobic.004G353000.1 | Zm00001d018522_T001 |
| orthologID_009326 | PH01000437G0190 | LOC_Os02g52780.1 | Bradi3g57960.4 | HORVU6Hr1G080670.1 | TRIUR3_27679 | Seita.1G331600.1 | Sobic.004G309600.1 | Zm00001d018178_T001 |
| orthologID_009211 | PH01000569G0180 | LOC_Os02g49610.1 | Bradi3g56240.1 | HORVU6Hr1G072650.3 | TRIUR3_03914 | Seita.1G305100.1 | Sobic.004G252100.1 | Zm00001d017914_T001 |
| orthologID_009458 | PH01003475G0070 | LOC_Os02g42520.2 | Bradi3g49740.1 | HORVU6Hr1G060010.19 | TRIUR3_13311 | Seita.1G247500.1 | Sobic.004G226500.1 | Zm00001d051227_T001 |
| orthologID_009309 | PH01000026G0730 | LOC_Os08g19370.1 | Bradi3g54150.1 | HORVU6Hr1G085830.1 | TRIUR3_20302 | Seita.1G351700.1 | Sobic.004G331200.2 | Zm00001d052027_T001 |
| orthologID_009844 | PH01000642G0360 | LOC_Os02g58410.1 | Bradi3g60610.3 | HORVU6Hr1G094660.5 | TRIUR3_06758 | Seita.1G378100.1 | Sobic.004G357100.1 | Zm00001d051839_T004 |
| orthologID_009181 | PH01000019G2350 | LOC_Os02g02740.3 | Bradi3g01817.1 | HORVU6Hr1G013000.2 | TRIUR3_10082 | Seita.1G110900.1 | Sobic.004G016300.2 | Zm00001d054009_T001 |
| orthologID_009113 | PH01000359G0710 | LOC_Os02g33860.1 | Bradi3g45337.1 | HORVU0Hr1G027010.1 | TRIUR3_18871 | Seita.1G187700.1 | Sobic.004G171200.1 | Zm00001d050708_T002 |
| orthologID_009432 | PH01000359G0660 | LOC_Os02g33780.1 | Bradi3g45310.1 | HORVU5Hr1G061040.3 | TRIUR3_29528 | Seita.1G187000.1 | Sobic.004G170800.2 | Zm00001d016756_T001 |
| orthologID_009295 | PH01001618G0390 | LOC_Os02g56600.1 | Bradi3g54817.1 | HORVU0Hr1G017490.7 | TRIUR3_32754 | Seita.1G359600.1 | Sobic.004G338800.1 | Zm00001d018435_T001 |
| orthologID_009319 | PH01000215G1010 | LOC_Os02g48600.1 | Bradi3g53400.1 | HORVU6Hr1G070510.2 | TRIUR3_21078 | Seita.1G296600.1 | Sobic.004G260300.1 | Zm00001d051643_T001 |
| orthologID_009365 | PH01000234G0310 | LOC_Os02g36850.1 | Bradi3g46880.1 | HORVU6Hr1G051650.2 | TRIUR3_29408 | Seita.1G208500.1 | Sobic.004G191200.1 | Zm00001d016943_T001 |
| orthologID_009733 | PH01001164G0340 | LOC_Os02g39890.1 | Bradi3g48560.1 | HORVU6Hr1G056260.5 | TRIUR3_13118 | Seita.1G230600.1 | Sobic.004G212700.1 | Zm00001d017193_T002 |
| orthologID_009238 | PH01002186G0270 | LOC_Os02g37000.1 | Bradi3g46980.1 | HORVU6Hr1G047300.2 | TRIUR3_27459 | Seita.1G210300.1 | Sobic.004G192800.1 | Zm00001d016979_T001 |
| orthologID_009826 | PH01000019G2400 | LOC_Os02g02770.1 | Bradi3g01770.1 | HORVU6Hr1G063510.3 | TRIUR3_31699 | Seita.1G110600.1 | Sobic.004G016500.1 | Zm00001d015030_T001 |
| orthologID_009391 | PH01000922G0160 | LOC_Os02g43340.2 | Bradi3g50230.1 | HORVU6Hr1G061380.5 | TRIUR3_22394 | Seita.1G253800.1 | Sobic.004G300100.1 | Zm00001d051320_T001 |
| orthologID_009894 | PH01000437G0850 | LOC_Os02g52380.1 | Bradi3g58180.1 | HORVU6Hr1G077230.2 | TRIUR3_26139 | Seita.1G328800.1 | Sobic.004G307000.1 | Zm00001d052179_T001 |
| orthologID_009657 | PH01000922G0620 | LOC_Os02g43550.1 | Bradi3g50350.1 | HORVU6Hr1G061870.1 | TRIUR3_19932 | Seita.1G255400.1 | Sobic.004G298500.2 | Zm00001d017441_T001 |
| orthologID_006782 | PH01002356G0260 | LOC_Os01g47350.1 | Bradi2g45757.1 | HORVU3Hr1G059840.4 | TRIUR3_26938 | Seita.5G253900.1 | Sobic.004G044200.1 | Zm00001d043970_T001 |
| orthologID_009824 | PH01001963G0320 | LOC_Os02g42620.1 | Bradi3g49836.1 | HORVU6Hr1G060250.7 | TRIUR3_03647 | Seita.1G248700.1 | Sobic.004G227700.1 | Zm00001d017374_T002 |
| orthologID_009201 | PH01000732G0570 | LOC_Os02g55430.1 | Bradi3g54210.2 | HORVU6Hr1G085880.3 | TRIUR3_07978 | Seita.1G352300.1 | Sobic.004G331800.1 | Zm00001d018388_T001 |
| orthologID_009561 | PH01002249G0120 | LOC_Os02g30610.2 | Bradi3g44100.1 | HORVU2Hr1G084420.1 | TRIUR3_03698 | Seita.1G169400.1 | Sobic.004G155400.1 | Zm00001d050563_T001 |
| orthologID_009162 | PH01000036G1460 | LOC_Os02g56570.1 | Bradi3g54840.1 | HORVU6Hr1G088820.5 | TRIUR3_03827 | Seita.1G359700.1 | Sobic.004G339000.1 | Zm00001d018438_T003 |
| orthologID_007106 | PH01000590G0070 | LOC_Os09g04490.1 | Bradi4g08540.1 | HORVU0Hr1G012140.1 | TRIUR3_21469 | Seita.4G289600.1 | Sobic.004G326100.1 | Zm00001d052058_T001 |
| orthologID_009712 | PH01000203G0800 | LOC_Os02g55080.1 | Bradi3g53940.1 | HORVU0Hr1G040430.3 | TRIUR3_15960 | Seita.1G348900.1 | Sobic.004G328000.3 | Zm00001d018364_T001 |
| orthologID_009828 | PH01002051G0150 | LOC_Os02g34700.1 | Bradi3g45620.1 | HORVU5Hr1G079090.7 | TRIUR3_09170 | Seita.1G191400.1 | Sobic.004G174800.2 | Zm00001d016798_T001 |
| orthologID_009710 | PH01001580G0090 | LOC_Os02g57280.1 | Bradi3g55440.1 | HORVU0Hr1G017570.2 | TRIUR3_23031 | Seita.1G366500.1 | Sobic.004G345900.1 | Zm00001d051908_T001 |
| orthologID_009467 | PH01000459G0200 | LOC_Os02g39630.2 | Bradi3g48430.1 | HORVU6Hr1G055910.2 | TRIUR3_23456 | Seita.1G228000.1 | Sobic.004G210800.2 | Zm00001d017174_T004 |
| orthologID_009437 | PH01000021G0050 | LOC_Os02g57990.1 | Bradi3g56110.1 | HORVU6Hr1G093310.1 | TRIUR3_01245 | Seita.1G374100.1 | Sobic.004G352700.1 | Zm00001d018517_T001 |
| orthologID_009833 | PH01000156G0910 | LOC_Os02g27740.1 | Bradi3g43470.1 | HORVU1Hr1G058580.1 | TRIUR3_31132 | Seita.1G160400.1 | Sobic.004G145600.1 | Zm00001d016490_T001 |
| orthologID_009492 | PH01000026G0230 | LOC_Os02g55030.1 | Bradi3g53887.1 | HORVU6Hr1G084590.3 | TRIUR3_09386 | Seita.1G348400.1 | Sobic.004G327500.1 | Zm00001d052050_T001 |
| orthologID_009091 | PH01000347G0200 | LOC_Os02g03280.2 | Bradi3g02520.2 | HORVU6Hr1G014450.1 | TRIUR3_21706 | Seita.1G103700.1 | Sobic.004G024200.1 | Zm00001d053952_T001 |
| orthologID_009246 | PH01000026G0180 | LOC_Os02g54910.1 | Bradi3g53850.1 | HORVU6Hr1G084430.3 | TRIUR3_17432 | Seita.1G348000.1 | Sobic.004G327100.1 | Zm00001d018355_T005 |
| orthologID_009632 | PH01002416G0270 | LOC_Os02g10090.1 | Bradi3g06970.1 | HORVU6Hr1G033750.1 | TRIUR3_21044 | Seita.1G051100.1 | Sobic.004G078300.1 | Zm00001d053680_T001 |
| orthologID_009112 | PH01002197G0130 | LOC_Os02g26860.1 | Bradi3g43180.1 | HORVU1Hr1G079270.2 | TRIUR3_11092 | Seita.1G157800.1 | Sobic.004G141400.1 | Zm00001d016472_T001 |
| orthologID_009750 | PH01000140G0530 | LOC_Os02g58030.1 | Bradi3g56757.1 | HORVU6Hr1G093360.11 | TRIUR3_25609 | Seita.1G374600.1 | Sobic.004G353400.1 | Zm00001d018525_T004 |
| orthologID_009294 | PH01000731G0510 | LOC_Os02g37940.1 | Bradi3g47520.1 | HORVU6Hr1G052970.2 | TRIUR3_19339 | Seita.1G215800.1 | Sobic.004G199500.1 | Zm00001d050942_T001 |
| orthologID_009424 | PH01000135G0910 | LOC_Os02g51160.1 | Bradi3g59337.1 | HORVU6Hr1G075730.11 | TRIUR3_15714 | Seita.1G319300.1 | Sobic.004G237500.1 | Zm00001d018042_T002 |
| orthologID_009797 | PH01000496G0570 | LOC_Os02g43330.1 | Bradi3g50220.1 | HORVU6Hr1G061390.1 | TRIUR3_22395 | Seita.1G253700.1 | Sobic.004G300300.1 | Zm00001d017422_T001 |
| orthologID_009197 | PH01000026G1530 | LOC_Os02g55950.1 | Bradi3g54430.1 | HORVU6Hr1G087150.8 | TRIUR3_19871 | Seita.1G355100.1 | Sobic.004G334300.1 | Zm00001d052009_T002 |

| | | | | | | | | |
|---|---|---|---|---|---|---|---|---|
| orthologID_009156 | PH01001846G0070 | LOC_Os02g13290.1 | Bradi3g08530.1 | HORVU6Hr1G040770.4 | TRIUR3_05914 | Seita.1G031100.1 | Sobic.004G097300.1 | Zm00001d016301_T001 |
| orthologID_009925 | PH01000019G0770 | LOC_Os02g01480.1 | Bradi3g00804.3 | HORVU2Hr1G021980.24 | TRIUR3_01800 | Seita.1G004000.1 | Sobic.004G004100.1 | Zm00001d054077_T003 |
| orthologID_009378 | PH01003576G0140 | LOC_Os02g43830.1 | Bradi3g50520.2 | HORVU6Hr1G062440.1 | TRIUR3_33648 | Seita.1G257700.1 | Sobic.004G296800.1 | Zm00001d017467_T001 |
| orthologID_009476 | PH01000036G0350 | LOC_Os02g55980.1 | Bradi3g54470.1 | HORVU6Hr1G087340.1 | TRIUR3_11885 | Seita.1G355400.1 | Sobic.004G334800.1 | Zm00001d052007_T001 |
| orthologID_009037 | PH01000023G0490 | LOC_Os04g46610.2 | Bradi5g17740.1 | HORVU2Hr1G097670.1 | TRIUR3_27591 | Seita.7G190200.1 | Sobic.004G326500.1 | Zm00001d018348_T001 |
| orthologID_006438 | PH01001803G0260 | LOC_Os05g22670.1 | Bradi2g31100.1 | HORVU1Hr1G034730.1 | TRIUR3_34207 | Seita.3G293700.1 | Sobic.004G252900.1 | Zm00001d009878_T001 |
| orthologID_009283 | PH01002092G0390 | LOC_Os02g50010.1 | Bradi3g56890.2 | HORVU6Hr1G073590.1 | TRIUR3_16773 | Seita.1G309900.1 | Sobic.004G248400.3 | Zm00001d051790_T001 |
| orthologID_008044 | PH01002224G0190 | LOC_Os12g20150.1 | Bradi1g50530.1 | HORVU7Hr1G008070.6 | TRIUR3_30998 | Seita.1G107700.1 | Sobic.004G120100.1 | Zm00001d023792_T005 |
| orthologID_009880 | PH01000145G0510 | LOC_Os02g04500.2 | Bradi3g03340.1 | HORVU6Hr1G019230.14 | TRIUR3_24221 | Seita.1G093300.1 | Sobic.004G035000.1 | Zm00001d015210_T001 |
| orthologID_009152 | PH01001614G0230 | LOC_Os02g37070.1 | Bradi3g47050.1 | HORVU7Hr1G115360.1 | TRIUR3_00204 | Seita.1G210900.1 | Sobic.004G193500.1 | Zm00001d050908_T001 |
| orthologID_009401 | PH01000672G0750 | LOC_Os02g46360.1 | Bradi3g52060.1 | HORVU6Hr1G066720.1 | TRIUR3_31564 | Seita.1G277800.1 | Sobic.004G277100.1 | Zm00001d051503_T001 |
| orthologID_009387 | PH01002102G0180 | LOC_Os02g42374.1 | Bradi3g49650.1 | HORVU6Hr1G059860.1 | TRIUR3_35244 | Seita.1G246100.1 | Sobic.004G225300.1 | Zm00001d017351_T001 |
| orthologID_009622 | PH01001148G0360 | LOC_Os02g46700.1 | Bradi3g52240.1 | HORVU6Hr1G067300.15 | TRIUR3_04159 | Seita.1G280800.1 | Sobic.004G274400.1 | Zm00001d017691_T001 |
| orthologID_009787 | PH01000140G1000 | LOC_Os02g57690.2 | Bradi3g56040.1 | HORVU6Hr1G093000.13 | TRIUR3_35388 | Seita.1G372000.1 | Sobic.004G350700.1 | Zm00001d051876_T002 |
| orthologID_009139 | PH01000021G1280 | LOC_Os02g57430.1 | Bradi3g55590.2 | HORVU6Hr1G091500.1 | TRIUR3_16284 | Seita.1G368800.1 | Sobic.004G348100.2 | Zm00001d018501_T001 |
| orthologID_009565 | PH01000145G1450 | LOC_Os02g03930.1 | Bradi3g02950.1 | HORVU6Hr1G017080.17 | TRIUR3_13851 | Seita.1G097300.1 | Sobic.004G030600.2 | Zm00001d053986_T001 |
| orthologID_009730 | PH01000065G1920 | LOC_Os02g51280.1 | Bradi3g59320.2 | HORVU6Hr1G075650.1 | TRIUR3_29014 | Seita.1G319500.1 | Sobic.004G237300.1 | Zm00001d018045_T001 |
| orthologID_009183 | PH01000093G0950 | LOC_Os02g06880.1 | Bradi3g04890.1 | HORVU0Hr1G015630.5 | TRIUR3_35165 | Seita.1G077500.1 | Sobic.004G051600.1 | Zm00001d053802_T001 |
| orthologID_009745 | PH01000062G1480 | LOC_Os02g48650.1 | Bradi3g53450.1 | HORVU6Hr1G070650.1 | TRIUR3_18350 | Seita.1G297200.1 | Sobic.004G259800.1 | Zm00001d051663_T001 |
| orthologID_009350 | PH01000482G0510 | LOC_Os02g06500.1 | Bradi3g04550.1 | HORVU6Hr1G025670.1 | TRIUR3_09738 | Seita.1G080100.1 | Sobic.004G048800.1 | Zm00001d015314_T003 |
| orthologID_009105 | PH01001371G0430 | LOC_Os02g32780.1 | Bradi3g44810.1 | HORVU6Hr1G066600.3 | TRIUR3_31762 | Seita.1G179400.1 | Sobic.004G164100.1 | Zm00001d016690_T001 |
| orthologID_009124 | PH01000998G0010 | LOC_Os02g45390.1 | Bradi3g51600.1 | HORVU0Hr1G016510.1 | TRIUR3_35420 | Seita.1G270700.1 | Sobic.004G283400.1 | Zm00001d051448_T001 |
| orthologID_009465 | PH01000973G0370 | LOC_Os02g13100.1 | Bradi3g08390.1 | HORVU6Hr1G039940.1 | TRIUR3_34991 | Seita.1G032000.1 | Sobic.004G096300.1 | Zm00001d016332_T001 |
| orthologID_009407 | PH01001018G0090 | LOC_Os02g29550.1 | Bradi3g43822.2 | HORVU6Hr1G042680.5 | TRIUR3_27883 | Seita.1G165600.1 | Sobic.004G151500.3 | Zm00001d016535_T002 |
| orthologID_009882 | PH01000049G1120 | LOC_Os02g51480.1 | Bradi3g59120.1 | HORVU6Hr1G075250.1 | TRIUR3_19073 | Seita.1G322000.1 | Sobic.004G235100.1 | Zm00001d052240_T001 |
| orthologID_006585 | PH01000284G0690 | LOC_Os02g57150.1 | Bradi2g49650.1 | HORVU5Hr1G007960.1 | TRIUR3_18472 | Seita.9G575400.1 | Sobic.004G344500.1 | Zm00001d018478_T001 |
| orthologID_009420 | PH01000313G0880 | LOC_Os01g32790.1 | Bradi3g28570.1 | HORVU5Hr1G119260.1 | TRIUR3_00694 | Seita.1G380800.1 | Sobic.004G010700.1 | Zm00001d017529_T001 |
| orthologID_009103 | PH01000173G0590 | LOC_Os02g52550.1 | Bradi3g58080.1 | HORVU6Hr1G078660.4 | TRIUR3_05363 | Seita.1G330000.1 | Sobic.004G308100.1 | Zm00001d052171_T001 |
| orthologID_009380 | PH01000501G0660 | LOC_Os02g36960.1 | Bradi3g46940.1 | HORVU6Hr1G052250.2 | TRIUR3_34770 | Seita.1G209800.1 | Sobic.004G192200.1 | Zm00001d016972_T001 |
| orthologID_007384 | PH01000600G0350 | LOC_Os02g49830.1 | Bradi4g07750.1 | HORVU6Hr1G073000.1 | TRIUR3_23012 | Seita.1G308000.1 | Sobic.004G250100.1 | Zm00001d017931_T001 |
| orthologID_009363 | PH01000215G1180 | LOC_Os02g48660.1 | Bradi3g53460.1 | HORVU6Hr1G070670.1 | TRIUR3_18352 | Seita.1G297300.1 | Sobic.004G259700.1 | Zm00001d017850_T001 |
| orthologID_009110 | PH01000917G0260 | LOC_Os02g07910.2 | Bradi3g05620.1 | HORVU6Hr1G030940.1 | TRIUR3_04147 | Seita.1G067800.1 | Sobic.004G060500.1 | Zm00001d015431_T001 |
| orthologID_009101 | PH01000965G0170 | LOC_Os02g09590.1 | Bradi3g06530.1 | HORVU6Hr1G035260.2 | TRIUR3_28847 | Seita.1G056700.1 | Sobic.004G072000.1 | Zm00001d053713_T001 |
| orthologID_007777 | PH01002755G0250 | LOC_Os12g18960.1 | Bradi1g29097.3 | HORVU3Hr1G080200.1 | TRIUR3_25037 | Seita.1G116600.1 | Sobic.004G003300.2 | Zm00001d014933_T001 |
| orthologID_009036 | PH01000224G0100 | LOC_Os04g51960.1 | Bradi5g21090.1 | HORVU2Hr1G100290.1 | TRIUR3_25783 | Seita.1G026200.1 | Sobic.004G102000.1 | Zm00001d053524_T002 |
| orthologID_009626 | PH01000569G0260 | LOC_Os02g49680.1 | Bradi3g56190.1 | HORVU6Hr1G072740.1 | TRIUR3_28250 | Seita.1G305900.1 | Sobic.004G251500.1 | Zm00001d051746_T001 |
| orthologID_009310 | PH01003202G0160 | LOC_Os02g42960.1 | Bradi3g50060.1 | HORVU6Hr1G060810.2 | TRIUR3_20706 | Seita.1G251800.1 | Sobic.004G302500.1 | Zm00001d017395_T002 |
| orthologID_009840 | PH01002151G0040 | LOC_Os02g46640.1 | Bradi3g52190.1 | HORVU6Hr1G067130.3 | TRIUR3_19755 | Seita.1G280000.1 | Sobic.004G275000.1 | Zm00001d017685_T005 |
| orthologID_009382 | PH01000145G1020 | LOC_Os02g04170.1 | Bradi3g03080.1 | HORVU6Hr1G017860.16 | TRIUR3_31522 | Seita.1G095500.1 | Sobic.004G032400.1 | Zm00001d015183_T002 |
| orthologID_009398 | PH01001529G0060 | LOC_Os11g18830.1 | Bradi3g54880.1 | HORVU6Hr1G088840.1 | TRIUR3_03823 | Seita.8G106500.1 | Sobic.004G227100.1 | Zm00001d017363_T001 |
| orthologID_009703 | PH01000473G0620 | LOC_Os02g42412.1 | Bradi3g49687.2 | HORVU6Hr1G059950.1 | TRIUR3_32774 | Seita.1G246500.1 | Sobic.004G225700.1 | Zm00001d051219_T001 |
| orthologID_009760 | PH01000043G0300 | LOC_Os02g08310.1 | Bradi3g05930.1 | HORVU6Hr1G029710.9 | TRIUR3_24947 | Seita.1G063700.1 | Sobic.004G064700.1 | Zm00001d015485_T001 |
| orthologID_009862 | PH01000043G1970 | LOC_Os02g09380.1 | Bradi3g06280.1 | HORVU6Hr1G035820.1 | TRIUR3_17888 | Seita.1G058600.1 | Sobic.004G069500.1 | Zm00001d015586_T002 |
| orthologID_009700 | PH01002988G0200 | LOC_Os02g09340.1 | Bradi3g06260.1 | HORVU6Hr1G036020.1 | TRIUR3_10761 | Seita.1G058900.1 | Sobic.004G069000.1 | Zm00001d053726_T001 |
| orthologID_009708 | PH01000642G0080 | LOC_Os02g58270.1 | Bradi3g60470.1 | HORVU1Hr1G053070.1 | TRIUR3_28013 | Seita.1G376900.1 | Sobic.004G356100.1 | Zm00001d051842_T003 |
| orthologID_009808 | PH01000893G0520 | LOC_Os02g40810.1 | Bradi3g49030.1 | HORVU6Hr1G058100.1 | TRIUR3_30480 | Seita.1G237900.1 | Sobic.004G218300.1 | Zm00001d017252_T002 |
| orthologID_009919 | PH01000135G1100 | LOC_Os02g51110.1 | Bradi3g59390.1 | HORVU6Hr1G075850.2 | TRIUR3_22736 | Seita.1G318800.1 | Sobic.004G238100.1 | Zm00001d018037_T001 |
| orthologID_009475 | PH01003222G0150 | LOC_Os02g31030.2 | Bradi3g44290.1 | HORVU3Hr1G079900.5 | TRIUR3_27322 | Seita.1G171900.1 | Sobic.004G157300.1 | Zm00001d016588_T002 |
| orthologID_009876 | PH01000514G0160 | LOC_Os02g12360.1 | Bradi3g08040.1 | HORVU6Hr1G038590.4 | TRIUR3_27519 | Seita.1G037800.1 | Sobic.004G091100.1 | Zm00001d016382_T004 |
| orthologID_009679 | PH01000457G0620 | LOC_Os02g03070.1 | Bradi3g02320.1 | HORVU6Hr1G013760.2 | TRIUR3_18034 | Seita.1G106700.1 | Sobic.004G021100.1 | Zm00001d053931_T003 |

| | | | | | | | | |
|---|---|---|---|---|---|---|---|---|
| orthologID_009886 | PH01006188G0020 | LOC_Os02g46610.1 | Bradi3g52170.1 | HORVU6Hr1G067060.1 | TRIUR3_30366 | Seita.1G279800.1 | Sobic.004G275200.1 | Zm00001d017682_T001 |
| orthologID_009177 | PH01000649G0680 | LOC_Os10g15390.1 | Bradi3g23060.1 | HORVU4Hr1G075580.8 | TRIUR3_03829 | Seita.9G278800.1 | Sobic.004G093500.2 | Zm00001d016371_T001 |
| orthologID_009857 | PH01000584G0320 | LOC_Os02g13970.2 | Bradi3g09010.2 | HORVU2Hr1G022680.11 | TRIUR3_24322 | Seita.1G024100.1 | Sobic.004G103300.1 | Zm00001d053545_T003 |
| orthologID_009253 | PH01000347G1380 | LOC_Os02g02990.1 | Bradi3g02180.1 | HORVU6Hr1G012010.1 | TRIUR3_08208 | Seita.1G107600.1 | Sobic.004G020000.1 | Zm00001d015061_T001 |
| orthologID_009656 | PH01000124G0720 | LOC_Os02g10820.1 | Bradi3g07530.2 | HORVU6Hr1G037860.2 | TRIUR3_32595 | Seita.1G044000.1 | Sobic.004G085300.1 | Zm00001d053624_T001 |
| orthologID_009868 | PH01000089G0980 | LOC_Os02g39140.1 | Bradi3g48070.3 | HORVU6Hr1G054910.7 | TRIUR3_16394 | Seita.1G223700.1 | Sobic.004G207200.1 | Zm00001d017138_T001 |
| orthologID_007315 | PH01000356G0540 | LOC_Os02g57070.1 | Bradi4g37550.1 | HORVU5Hr1G088210.8 | TRIUR3_30029 | Seita.1G364300.1 | Sobic.004G343600.1 | Zm00001d018471_T002 |
| orthologID_009609 | PH01000026G1980 | LOC_Os02g56170.1 | Bradi3g54660.1 | HORVU6Hr1G087990.2 | TRIUR3_03504 | Seita.1G357600.1 | Sobic.004G337000.2 | Zm00001d051988_T001 |
| orthologID_010164 | PH01001308G0400 | LOC_Os02g52910.1 | Bradi3g57850.1 | HORVU7Hr1G060240.1 | TRIUR3_07479 | Seita.1G332900.1 | Sobic.004G310800.1 | Zm00001d018193_T001 |
| orthologID_009909 | PH01053188G0010 | LOC_Os02g54100.1 | Bradi3g60070.2 | HORVU6Hr1G082310.4 | TRIUR3_29343 | Seita.1G341600.1 | Sobic.004G320300.1 | Zm00001d018292_T001 |
| orthologID_009331 | PH01000691G0480 | LOC_Os02g30460.1 | Bradi3g44060.1 | HORVU6Hr1G044700.5 | TRIUR3_31029 | Seita.1G168800.1 | Sobic.004G155100.1 | Zm00001d016566_T001 |
| orthologID_009856 | PH01003332G0210 | LOC_Os02g02980.1 | Bradi3g02170.5 | HORVU6Hr1G012070.9 | TRIUR3_05465 | Seita.1G108000.1 | Sobic.004G019800.2 | Zm00001d015060_T001 |
| orthologID_009838 | PH01007027G0030 | LOC_Os05g19954.1 | Bradi3g27090.2 | HORVU1Hr1G031370.1 | TRIUR3_34434 | Seita.1G156600.1 | Sobic.004G193200.5 | Zm00001d016988_T001 |
| orthologID_009516 | PH01000943G0250 | LOC_Os02g52140.1 | Bradi3g58600.1 | HORVU6Hr1G077700.12 | TRIUR3_01192 | Seita.1G326500.1 | Sobic.004G229000.1 | Zm00001d052198_T002 |
| orthologID_009638 | PH01000197G0410 | LOC_Os02g47660.1 | Bradi3g52790.2 | HORVU6Hr1G068980.2 | TRIUR3_17846 | Seita.1G289600.1 | Sobic.004G267400.1 | Zm00001d017778_T001 |
| orthologID_009413 | PH01000447G0170 | LOC_Os02g44860.1 | Bradi3g51207.1 | HORVU6Hr1G064610.1 | TRIUR3_21895 | Seita.1G267100.1 | Sobic.004G286700.1 | Zm00001d051419_T002 |
| orthologID_009371 | PH01000021G1200 | LOC_Os02g57470.1 | Bradi3g55660.1 | HORVU6Hr1G091620.4 | TRIUR3_17528 | Seita.1G369400.1 | Sobic.004G348600.1 | Zm00001d051893_T001 |
| orthologID_009619 | PH01002103G0390 | LOC_Os02g01150.1 | Bradi3g00330.1 | HORVU6Hr1G000650.1 | TRIUR3_20270 | Seita.1G001300.1 | Sobic.004G001300.1 | Zm00001d014919_T002 |
| orthologID_009773 | PH01003778G0160 | LOC_Os02g50700.1 | Bradi3g59700.1 | HORVU6Hr1G076530.1 | TRIUR3_24830 | Seita.1G315300.1 | Sobic.004G243300.1 | Zm00001d017990_T001 |
| orthologID_009842 | PH01000145G0580 | LOC_Os02g04460.1 | Bradi3g03270.1 | HORVU6Hr1G018830.4 | TRIUR3_08790 | Seita.1G093900.1 | Sobic.004G034500.1 | Zm00001d015204_T001 |
| orthologID_009127 | PH01000888G0360 | LOC_Os08g05750.1 | Bradi3g16967.2 | HORVU7Hr1G071780.9 | TRIUR3_21073 | Seita.6G059600.1 | Sobic.004G255300.1 | Zm00001d017894_T001 |
| orthologID_009457 | PH01000026G2100 | LOC_Os02g56200.1 | Bradi3g54690.1 | HORVU6Hr1G088200.8 | TRIUR3_03499 | Seita.1G358100.1 | Sobic.004G337400.1 | Zm00001d018420_T008 |
| orthologID_009226 | PH01000095G0580 | LOC_Os02g02840.1 | Bradi3g01930.1 | HORVU6Hr1G012690.3 | TRIUR3_32502 | Seita.1G109800.1 | Sobic.004G017900.1 | Zm00001d015036_T007 |
| orthologID_009623 | PH01001618G0340 | LOC_Os02g56565.1 | Bradi3g54847.1 | HORVU6Hr1G088800.8 | TRIUR3_03826 | Seita.1G359800.1 | Sobic.004G339100.1 | Zm00001d018439_T003 |
| orthologID_007161 | PH01000005G2160 | LOC_Os11g34310.1 | Bradi4g16820.1 | HORVU7Hr1G091900.1 | TRIUR3_01429 | Seita.8G145000.1 | Sobic.004G072100.1 | Zm00001d007585_T006 |
| orthologID_009179 | PH01001462G0120 | LOC_Os02g39580.1 | Bradi3g48390.1 | HORVU6Hr1G055830.1 | TRIUR3_25723 | Seita.1G227700.1 | Sobic.004G210101.1 | Zm00001d017169_T001 |
| orthologID_009089 | PH01000943G0760 | LOC_Os02g52290.1 | Bradi3g58255.1 | HORVU6Hr1G077340.1 | TRIUR3_12181 | Seita.1G328100.1 | Sobic.004G306200.1 | Zm00001d018134_T001 |
| orthologID_009584 | PH01000519G0150 | LOC_Os02g51100.1 | Bradi3g59400.1 | HORVU0Hr1G017120.2 | TRIUR3_22737 | Seita.1G318700.1 | Sobic.004G238200.1 | Zm00001d018036_T002 |
| orthologID_009166 | PH01000145G0330 | LOC_Os02g04550.1 | Bradi3g03391.1 | HORVU6Hr1G019350.2 | TRIUR3_32969 | Seita.1G092800.1 | Sobic.004G035900.2 | Zm00001d015215_T001 |
| orthologID_009673 | PH01000197G1080 | LOC_Os02g47440.1 | Bradi3g52640.1 | HORVU6Hr1G068640.1 | TRIUR3_19543 | Seita.1G287900.1 | Sobic.004G269000.1 | Zm00001d017756_T001 |
| orthologID_009298 | PH01000089G0560 | LOC_Os02g39350.1 | Bradi3g48210.1 | HORVU6Hr1G055460.11 | TRIUR3_16792 | Seita.1G225100.1 | Sobic.004G208500.1 | Zm00001d017150_T001 |
| orthologID_009442 | PH01002244G0320 | LOC_Os02g37150.1 | Bradi3g47157.1 | HORVU6Hr1G051740.2 | TRIUR3_33755 | Seita.1G211600.1 | Sobic.004G194900.1 | Zm00001d017000_T001 |
| orthologID_009373 | PH01000019G1100 | LOC_Os02g01720.1 | Bradi3g00960.1 | HORVU6Hr1G004260.1 | TRIUR3_25851 | Seita.1G005300.1 | Sobic.004G005200.1 | Zm00001d014951_T002 |
| orthologID_009255 | PH01000215G0810 | LOC_Os02g48470.1 | Bradi3g53320.1 | HORVU6Hr1G070290.1 | TRIUR3_06525 | Seita.1G295900.1 | Sobic.004G261200.1 | Zm00001d017837_T001 |
| orthologID_009384 | PH01000922G0500 | LOC_Os02g43500.1 | Bradi3g50330.1 | HORVU6Hr1G061800.1 | TRIUR3_32622 | Seita.1G255000.1 | Sobic.004G298900.1 | Zm00001d017435_T001 |
| orthologID_009417 | PH01000006G3200 | LOC_Os02g28040.1 | Bradi3g43530.1 | HORVU6Hr1G045800.2 | TRIUR3_22501 | Seita.1G161500.1 | Sobic.004G146600.1 | Zm00001d053247_T005 |
| orthologID_009904 | PH01006176G0030 | LOC_Os02g09530.1 | Bradi3g06500.1 | HORVU6Hr1G035300.19 | TRIUR3_23606 | Seita.1G056900.1 | Sobic.004G071700.1 | Zm00001d015627_T002 |
| orthologID_009654 | PH01002101G0240 | LOC_Os02g53620.1 | Bradi3g57320.1 | HORVU6Hr1G081080.12 | TRIUR3_28928 | Seita.1G338100.1 | Sobic.004G316500.2 | Zm00001d018255_T001 |
| orthologID_009269 | PH01000123G0400 | LOC_Os02g07680.1 | Bradi3g05400.1 | HORVU6Hr1G031650.5 | TRIUR3_19610 | Seita.1G070400.1 | Sobic.004G057900.2 | Zm00001d053782_T006 |
| orthologID_007360 | PH01000604G0250 | LOC_Os04g33360.1 | Bradi4g01693.1 | HORVU3Hr1G117870.1 | TRIUR3_06980 | Seita.4G015700.1 | Sobic.004G329100.2 | Zm00001d035994_T001 |
| orthologID_008518 | PH01001935G0170 | LOC_Os03g12720.1 | Bradi1g69090.1 | HORVU4Hr1G065610.1 | TRIUR3_33987 | Seita.9G480600.1 | Sobic.004G067400.1 | Zm00001d015553_T001 |
| orthologID_009763 | PH01000356G0410 | LOC_Os02g57010.1 | Bradi3g55190.1 | HORVU6Hr1G090190.2 | TRIUR3_03557 | Seita.1G363700.1 | Sobic.004G343000.1 | Zm00001d018466_T002 |
| orthologID_009746 | PH01001025G0190 | LOC_Os02g42230.1 | Bradi3g49530.1 | HORVU6Hr1G059440.7 | TRIUR3_09467 | Seita.1G244500.1 | Sobic.004G224100.1 | Zm00001d017324_T001 |
| orthologID_009900 | PH01000114G0340 | LOC_Os02g04970.1 | Bradi3g03637.1 | HORVU6Hr1G020420.1 | TRIUR3_25313 | Seita.1G090600.1 | Sobic.004G038100.1 | Zm00001d015245_T002 |
| orthologID_008309 | PH01001321G0350 | LOC_Os06g30020.1 | Bradi1g42420.1 | HORVU4Hr1G000670.2 | TRIUR3_21505 | Seita.1G197100.1 | Sobic.004G180900.3 | Zm00001d050793_T001 |
| orthologID_009289 | PH01000120G1260 | LOC_Os02g10380.1 | Bradi3g07180.3 | HORVU6Hr1G033200.2 | TRIUR3_18290 | Seita.1G048400.1 | Sobic.004G080500.2 | Zm00001d053660_T001 |
| orthologID_009875 | PH01000307G0680 | LOC_Os02g20870.1 | Bradi3g11450.2 | HORVU6Hr1G054340.1 | TRIUR3_24807 | Seita.1G141400.1 | Sobic.004G067500.3 | Zm00001d053751_T001 |
| orthologID_009241 | PH01000197G1460 | LOC_Os02g47280.1 | Bradi3g52547.1 | HORVU6Hr1G068370.1 | TRIUR3_22138 | Seita.1G287100.1 | Sobic.004G269900.2 | Zm00001d017742_T001 |
| orthologID_009111 | PH01215153G0010 | LOC_Os02g44690.1 | Bradi3g51340.1 | HORVU6Hr1G064230.1 | TRIUR3_26462 | Seita.1G265700.1 | Sobic.004G288100.1 | Zm00001d051405_T001 |
| orthologID_009336 | PH01001610G0110 | LOC_Os02g44330.1 | Bradi3g50910.1 | HORVU6Hr1G063590.2 | TRIUR3_23240 | Seita.1G262700.1 | Sobic.004G291400.2 | Zm00001d017512_T003 |

| | | | | | | | | |
|---|---|---|---|---|---|---|---|---|
| orthologID_009292 | PH01000943G0260 | LOC_Os02g52150.1 | Bradi3g58590.2 | HORVU6Hr1G077710.2 | TRIUR3_01193 | Seita.1G326600.1 | Sobic.004G228900.2 | Zm00001d052194_T001 |
| orthologID_009208 | PH01002051G0260 | LOC_Os02g34870.1 | Bradi3g45750.1 | HORVU0Hr1G021540.1 | TRIUR3_16781 | Seita.1G192500.1 | Sobic.004G176100.1 | Zm00001d047151_T001 |
| orthologID_009425 | PH01001447G0440 | LOC_Os02g31140.1 | Bradi3g44330.1 | HORVU0Hr1G016140.1 | TRIUR3_27140 | Seita.1G172900.1 | Sobic.004G158000.1 | Zm00001d016599_T002 |
| orthologID_009762 | PH01001148G0260 | LOC_Os02g46750.1 | Bradi3g52250.1 | HORVU6Hr1G067390.2 | TRIUR3_33521 | Seita.1G281200.1 | Sobic.004G274100.3 | Zm00001d051518_T001 |
| orthologID_009803 | PH01000173G0210 | LOC_Os02g52270.1 | Bradi3g58280.1 | HORVU6Hr1G077430.1 | TRIUR3_12499 | Seita.1G327900.1 | Sobic.004G305900.1 | Zm00001d018131_T004 |
| orthologID_009280 | PH01003357G0140 | LOC_Os02g53430.1 | Bradi3g57425.3 | HORVU6Hr1G078890.13 | TRIUR3_08499 | Seita.1G337000.1 | Sobic.004G315300.1 | Zm00001d018241_T001 |
| orthologID_009898 | PH01001885G0240 | LOC_Os02g35770.1 | Bradi3g46320.2 | HORVU1Hr1G082910.2 | TRIUR3_05289 | Seita.1G201700.1 | Sobic.004G185200.1 | Zm00001d050834_T001 |
| orthologID_009725 | PH01000399G0070 | LOC_Os08g28970.1 | Bradi3g35120.4 | HORVU7Hr1G044280.2 | TRIUR3_02904 | Seita.6G129400.1 | Sobic.007G115700.1 | Zm00001d032339_T001 |
| orthologID_009603 | PH01000975G0190 | LOC_Os08g14200.1 | Bradi3g18690.1 | HORVU7Hr1G106530.2 | TRIUR3_03898 | Seita.6G014200.1 | Sobic.007G086300.1 | Zm00001d050201_T001 |
| orthologID_009598 | PH01000624G0460 | LOC_Os08g07880.1 | Bradi3g15607.2 | HORVU7Hr1G116040.8 | TRIUR3_03381 | Seita.6G045000.1 | Sobic.007G060700.1 | Zm00001d024421_T001 |
| orthologID_009535 | PH01000406G0320 | LOC_Os08g35710.1 | Bradi3g37640.1 | HORVU7Hr1G073190.2 | TRIUR3_30141 | Seita.6G173500.1 | Sobic.007G150800.1 | Zm00001d050106_T001 |
| orthologID_009590 | PH01000289G0290 | LOC_Os10g24800.1 | Bradi3g18300.1 | HORVU5Hr1G011050.1 | TRIUR3_14613 | Seita.6G021800.1 | Sobic.007G079500.1 | Zm00001d024344_T001 |
| orthologID_009866 | PH01001896G0320 | LOC_Os08g02390.1 | Bradi3g13790.1 | HORVU7Hr1G075300.4 | TRIUR3_31309 | Seita.6G085200.1 | Sobic.007G018900.1 | Zm00001d035092_T002 |
| orthologID_009469 | PH01001561G0010 | LOC_Os08g37490.1 | Bradi3g38640.1 | HORVU7Hr1G061920.2 | TRIUR3_22458 | Seita.6G188100.1 | Sobic.007G226100.1 | Zm00001d052796_T003 |
| orthologID_009343 | PH01001430G0210 | LOC_Os08g43480.1 | Bradi3g42410.1 | HORVU7Hr1G057310.5 | TRIUR3_15763 | Seita.6G238600.1 | Sobic.007G178100.1 | Zm00001d053212_T002 |
| orthologID_009088 | PH01001310G0040 | LOC_Os08g43230.1 | Bradi3g42050.3 | HORVU7Hr1G056790.4 | TRIUR3_13244 | Seita.6G235200.1 | Sobic.007G181400.3 | Zm00001d031729_T001 |
| orthologID_006246 | PH01000109G1450 | LOC_Os01g57470.1 | Bradi2g51910.2 | HORVU3Hr1G077070.2 | TRIUR3_11689 | Seita.5G342200.1 | Sobic.007G212700.1 | Zm00001d031419_T001 |
| orthologID_009864 | PH01001711G0360 | LOC_Os08g38060.1 | Bradi3g38870.1 | HORVU7Hr1G050950.1 | TRIUR3_29225 | Seita.6G190900.1 | Sobic.007G224300.1 | Zm00001d031273_T001 |
| orthologID_009794 | PH01002403G0150 | LOC_Os08g16570.1 | Bradi3g19310.1 | HORVU7Hr1G122750.2 | TRIUR3_31010 | Seita.6G099300.1 | Sobic.007G095500.1 | Zm00001d050173_T001 |
| orthologID_009702 | PH01001492G0240 | LOC_Os08g34700.1 | Bradi3g37230.2 | HORVU4Hr1G013760.1 | TRIUR3_28536 | Seita.6G167300.1 | Sobic.007G144900.1 | Zm00001d032115_T001 |
| orthologID_009377 | PH01001388G0420 | LOC_Os08g37140.1 | Bradi3g38010.1 | HORVU6Hr1G020310.3 | TRIUR3_25150 | Seita.6G183400.1 | Sobic.007G158300.1 | Zm00001d031809_T001 |
| orthologID_009131 | PH01001046G0260 | LOC_Os08g04110.1 | Bradi3g14406.2 | HORVU7Hr1G073550.1 | TRIUR3_03006 | Seita.6G074100.1 | Sobic.007G030700.1 | Zm00001d024708_T001 |
| orthologID_009374 | PH01002756G0180 | LOC_Os08g16660.1 | Bradi3g19360.1 | HORVU5Hr1G093290.1 | TRIUR3_17787 | Seita.6G101200.1 | Sobic.007G097100.1 | Zm00001d050178_T001 |
| orthologID_009261 | PH01003703G0070 | LOC_Os08g01770.1 | Bradi3g13360.1 | HORVU7Hr1G077170.3 | TRIUR3_13378 | Seita.6G009100.1 | Sobic.007G009200.1 | Zm00001d035034_T002 |
| orthologID_009165 | PH01001052G0270 | LOC_Os08g29100.1 | Bradi3g35080.1 | HORVU7Hr1G078300.3 | TRIUR3_28140 | Seita.6G129200.1 | Sobic.007G115900.2 | Zm00001d049329_T001 |
| orthologID_007079 | PH01004825G0030 | LOC_Os09g25550.1 | Bradi4g30570.3 | HORVU5Hr1G064130.1 | TRIUR3_11904 | Seita.7G074800.1 | Sobic.007G087100.1 | Zm00001d024409_T001 |
| orthologID_009860 | PH01000888G0440 | LOC_Os08g05790.1 | Bradi3g16950.2 | HORVU7Hr1G071740.7 | TRIUR3_15288 | Seita.6G059400.1 | Sobic.007G043200.1 | Zm00001d024596_T002 |
| orthologID_009625 | PH01000383G0720 | LOC_Os08g05890.1 | Bradi3g16680.1 | HORVU0Hr1G021390.2 | TRIUR3_23350 | Seita.6G058400.1 | Sobic.007G044400.1 | Zm00001d024588_T001 |
| orthologID_009106 | PH01001137G0180 | LOC_Os08g37280.1 | Bradi3g38350.3 | HORVU6Hr1G016610.2 | TRIUR3_14988 | Seita.6G184500.1 | Sobic.007G159200.1 | Zm00001d031817_T003 |
| orthologID_009589 | PH01003483G0210 | LOC_Os08g34150.1 | Bradi3g36970.1 | HORVU6Hr1G079030.4 | TRIUR3_29324 | Seita.6G162500.1 | Sobic.007G140200.2 | Zm00001d032162_T001 |
| orthologID_009174 | PH01000264G0330 | LOC_Os08g04120.1 | Bradi3g14412.1 | HORVU7Hr1G073520.1 | TRIUR3_03005 | Seita.6G074000.1 | Sobic.007G030800.1 | Zm00001d049615_T001 |
| orthologID_009885 | PH01000264G0060 | LOC_Os08g04310.1 | Bradi3g14570.1 | HORVU7Hr1G073390.1 | TRIUR3_20963 | Seita.6G070700.1 | Sobic.007G033300.1 | Zm00001d024691_T001 |
| orthologID_009272 | PH01002747G0210 | LOC_Os08g01130.1 | Bradi3g12930.1 | HORVU7Hr1G078710.3 | TRIUR3_28091 | Seita.6G001900.1 | Sobic.007G000900.2 | Zm00001d034968_T001 |
| orthologID_009441 | PH01000289G1080 | LOC_Os08g10450.1 | Bradi3g17700.1 | HORVU6Hr1G083770.2 | TRIUR3_08022 | Seita.6G029500.1 | Sobic.007G073700.1 | Zm00001d024294_T001 |
| orthologID_009406 | PH01005310G0030 | LOC_Os08g33120.1 | Bradi3g36320.1 | HORVU2Hr1G011700.2 | TRIUR3_10802 | Seita.6G153800.1 | Sobic.007G132400.1 | Zm00001d050357_T006 |
| orthologID_009783 | PH01000303G0090 | LOC_Os08g07430.1 | Bradi3g15890.1 | HORVU7Hr1G033450.4 | TRIUR3_33620 | Seita.6G047600.1 | Sobic.007G056600.1 | Zm00001d024467_T004 |
| orthologID_009674 | PH01004106G0180 | LOC_Os08g39270.1 | Bradi3g39740.1 | HORVU7Hr1G049860.1 | TRIUR3_27852 | Seita.6G201700.1 | Sobic.007G214700.1 | Zm00001d052864_T001 |
| orthologID_009595 | PH01002426G0150 | LOC_Os08g40120.1 | Bradi3g40147.1 | HORVU7Hr1G051120.5 | TRIUR3_20145 | Seita.6G207400.1 | Sobic.007G208200.1 | Zm00001d031475_T001 |
| orthologID_009579 | PH01001063G0380 | LOC_Os08g06190.1 | Bradi3g16460.1 | HORVU1Hr1G091500.1 | TRIUR3_18460 | Seita.6G055000.1 | Sobic.007G048600.1 | Zm00001d024541_T001 |
| orthologID_009839 | PH01001266G0420 | LOC_Os08g32850.1 | Bradi3g36140.1 | HORVU7Hr1G021890.4 | TRIUR3_02994 | Seita.6G151000.1 | Sobic.007G130600.1 | Zm00001d050336_T003 |
| orthologID_008725 | PH01000404G0390 | LOC_Os03g42650.2 | Bradi1g22070.1 | HORVU3Hr1G075600.2 | TRIUR3_14194 | Seita.5G180800.1 | Sobic.007G069101.1 | Zm00001d024243_T001 |
| orthologID_009362 | PH01004290G0030 | LOC_Os08g29530.2 | Bradi3g34900.2 | HORVU6Hr1G035180.4 | TRIUR3_11892 | Seita.6G126600.1 | Sobic.007G118000.1 | Zm00001d032326_T002 |
| orthologID_009116 | PH01078407G0010 | LOC_Os08g41840.1 | Bradi3g41180.1 | HORVU7Hr1G054070.1 | TRIUR3_24120 | Seita.6G222400.1 | Sobic.007G194200.1 | Zm00001d031618_T001 |
| orthologID_009268 | PH01000323G0640 | LOC_Os08g38410.1 | Bradi3g39090.1 | HORVU7Hr1G060930.1 | TRIUR3_27703 | Seita.6G193500.1 | Sobic.007G221300.1 | Zm00001d052820_T001 |
| orthologID_009274 | PH01003169G0020 | LOC_Os08g43360.1 | Bradi3g42150.2 | HORVU7Hr1G056910.1 | TRIUR3_29274 | Seita.6G236300.1 | Sobic.007G180000.1 | Zm00001d053197_T006 |
| orthologID_009474 | PH01167993G0010 | LOC_Os08g06240.1 | Bradi3g16420.1 | HORVU7Hr1G070730.1 | TRIUR3_31492 | Seita.6G054000.1 | Sobic.007G049900.3 | Zm00001d024538_T001 |
| orthologID_009765 | PH01001902G0320 | LOC_Os08g44250.1 | Bradi3g44590.1 | HORVU7Hr1G058490.1 | TRIUR3_21130 | Seita.6G246100.1 | Sobic.007G172300.1 | Zm00001d031975_T001 |
| orthologID_009887 | PH01001790G0140 | LOC_Os08g17080.1 | Bradi3g19530.1 | HORVU1Hr1G059490.1 | TRIUR3_07372 | Seita.6G101700.1 | Sobic.007G100500.1 | Zm00001d049986_T001 |
| orthologID_007214 | PH01003201G0040 | LOC_Os12g18760.2 | Bradi4g39067.1 | HORVU7Hr1G081420.5 | TRIUR3_11266 | Seita.3G309300.1 | Sobic.007G090430.1 | Zm00001d041108_T019 |
| orthologID_009460 | PH01000289G1030 | LOC_Os08g10470.1 | Bradi3g17710.1 | HORVU7Hr1G086320.1 | TRIUR3_04246 | Seita.6G027900.1 | Sobic.007G073900.1 | Zm00001d024298_T001 |

| | | | | | | | | |
|---|---|---|---|---|---|---|---|---|
| orthologID_009735 | PH01000303G0040 | LOC_Os08g07280.1 | Bradi3g15930.4 | HORVU1Hr1G008530.1 | TRIUR3_10748 | Seita.6G048100.1 | Sobic.007G056100.4 | Zm00001d024462_T003 |
| orthologID_009899 | PH01007618G0010 | LOC_Os08g03600.1 | Bradi3g14277.1 | HORVU7Hr1G074050.2 | TRIUR3_25295 | Seita.6G075900.1 | Sobic.007G028600.2 | Zm00001d049624_T001 |
| orthologID_009661 | PH01000915G0030 | LOC_Os08g41090.1 | Bradi3g40330.1 | HORVU7Hr1G051570.2 | TRIUR3_01974 | Seita.6G210000.1 | Sobic.007G205600.1 | Zm00001d031494_T001 |
| orthologID_009357 | PH01001659G0150 | LOC_Os08g39090.1 | Bradi3g39520.1 | HORVU7Hr1G049250.1 | TRIUR3_24536 | Seita.6G199500.1 | Sobic.007G216900.1 | Zm00001d031328_T001 |
| orthologID_009094 | PH01000915G0220 | LOC_Os08g40270.1 | Bradi3g40250.2 | HORVU7Hr1G051420.2 | TRIUR3_24930 | Seita.6G208700.1 | Sobic.007G206800.1 | Zm00001d052906_T001 |
| orthologID_008707 | PH01001843G0290 | LOC_Os07g12320.1 | Bradi1g53790.1 | HORVU7Hr1G117550.1 | TRIUR3_00348 | Seita.7G244600.1 | Sobic.007G137350.1 | Zm00001d019200_T001 |
| orthologID_009344 | PH01003366G0130 | LOC_Os07g09800.1 | Bradi3g21280.2 | HORVU1Hr1G022660.1 | TRIUR3_08967 | Seita.9G294600.1 | Sobic.007G223500.1 | Zm00001d014349_T001 |
| orthologID_009527 | PH01000264G1180 | LOC_Os08g03430.2 | Bradi3g14221.1 | HORVU7Hr1G074520.1 | TRIUR3_14364 | Seita.6G077400.1 | Sobic.007G026700.1 | Zm00001d035184_T001 |
| orthologID_009229 | PH01000399G0510 | LOC_Os08g29160.1 | Bradi3g34990.2 | HORVU1Hr1G093880.4 | TRIUR3_07762 | Seita.6G128700.1 | Sobic.007G116600.1 | Zm00001d032333_T004 |
| orthologID_009146 | PH01000573G0720 | LOC_Os07g25420.1 | Bradi3g17960.1 | HORVU4Hr1G002140.1 | TRIUR3_27145 | Seita.6G025200.1 | Sobic.007G076300.1 | Zm00001d024319_T001 |
| orthologID_009728 | PH01000161G0070 | LOC_Os08g15060.1 | Bradi3g19020.1 | HORVU7Hr1G112910.1 | TRIUR3_12608 | Seita.6G096700.1 | Sobic.007G091900.1 | Zm00001d024199_T002 |
| orthologID_009799 | PH01001780G0460 | LOC_Os08g01880.1 | Bradi3g13440.3 | HORVU7Hr1G076840.1 | TRIUR3_29539 | Seita.6G010600.1 | Sobic.007G011000.1 | Zm00001d000424_T001 |
| orthologID_009772 | PH01002800G0270 | LOC_Os08g29520.1 | Bradi3g34910.1 | HORVU7Hr1G065590.7 | TRIUR3_28163 | Seita.6G126700.1 | Sobic.007G117800.1 | Zm00001d032327_T001 |
| orthologID_008528 | PH01000774G0080 | LOC_Os06g27980.1 | Bradi1g41890.1 | HORVU0Hr1G020310.2 | TRIUR3_13635 | Seita.4G150500.1 | Sobic.007G007300.2 | Zm00001d035018_T001 |
| orthologID_009126 | PH01001388G0160 | LOC_Os08g37070.1 | Bradi3g38033.1 | HORVU3Hr1G031970.1 | TRIUR3_26909 | Seita.6G183100.1 | Sobic.007G158000.1 | Zm00001d031806_T001 |
| orthologID_009337 | PH01000003G0680 | LOC_Os08g02520.1 | Bradi3g13892.1 | HORVU7Hr1G075080.1 | TRIUR3_34394 | Seita.6G083700.1 | Sobic.007G020800.1 | Zm00001d049659_T001 |
| orthologID_006189 | PH01001032G0160 | LOC_Os01g29409.1 | Bradi2g13350.1 | HORVU3Hr1G030270.2 | TRIUR3_19798 | Seita.J001500.1 | Sobic.007G058001.1 | Zm00001d040920_T002 |
| orthologID_009219 | PH01001742G0260 | LOC_Os08g06430.1 | Bradi3g16290.1 | HORVU7Hr1G070150.2 | TRIUR3_12553 | Seita.6G052100.1 | Sobic.007G051000.1 | Zm00001d049507_T001 |
| orthologID_009426 | PH01000799G0530 | LOC_Os10g25000.1 | Bradi3g23267.1 | HORVU7Hr1G002890.4 | TRIUR3_22640 | Seita.9G249600.1 | Sobic.007G076400.1 | Zm00001d014272_T002 |
| orthologID_009879 | PH01005515G0090 | LOC_Os08g33640.1 | Bradi3g36640.1 | HORVU5Hr1G115070.4 | TRIUR3_11345 | Seita.6G157500.1 | Sobic.007G136200.1 | Zm00001d032209_T001 |
| orthologID_009522 | PH01002506G0270 | LOC_Os08g41500.1 | Bradi3g40980.1 | HORVU7Hr1G053440.5 | TRIUR3_26293 | Seita.6G220000.1 | Sobic.007G196300.1 | Zm00001d049805_T001 |
| orthologID_009560 | PH01000104G1120 | LOC_Os08g37520.1 | Bradi3g38620.1 | HORVU3Hr1G067580.1 | TRIUR3_24641 | Seita.6G187900.1 | Sobic.007G226400.1 | Zm00001d052794_T001 |
| orthologID_009411 | PH01000937G0140 | LOC_Os08g36420.3 | Bradi3g37910.1 | HORVU3Hr1G022500.4 | TRIUR3_31920 | Seita.6G178000.1 | Sobic.007G153500.1 | Zm00001d031764_T002 |
| orthologID_009653 | PH01000150G1040 | LOC_Os08g39430.1 | Bradi3g39830.1 | HORVU7Hr1G050340.1 | TRIUR3_10908 | Seita.6G203500.1 | Sobic.007G212400.2 | Zm00001d031421_T001 |
| orthologID_009077 | PH01002532G0110 | LOC_Os08g31140.1 | Bradi3g35250.3 | HORVU4Hr1G017070.1 | TRIUR3_33177 | Seita.6G139900.1 | Sobic.007G123200.1 | Zm00001d032291_T001 |
| orthologID_009727 | PH01001289G0260 | LOC_Os08g34130.1 | Bradi3g36980.1 | HORVU2Hr1G113990.2 | TRIUR3_08090 | Seita.6G162200.1 | Sobic.007G140100.1 | Zm00001d032163_T001 |
| orthologID_009867 | PH01157007G0010 | LOC_Os08g44840.1 | Bradi3g12497.1 | HORVU1Hr1G075900.1 | TRIUR3_18568 | Seita.6G252400.1 | Sobic.007G165800.1 | Zm00001d031893_T001 |
| orthologID_009149 | PH01000008G1510 | LOC_Os08g33810.1 | Bradi3g36750.1 | HORVU2Hr1G112480.4 | TRIUR3_07357 | Seita.6G158700.1 | Sobic.007G137000.4 | Zm00001d012891_T001 |
| orthologID_009811 | PH01000003G1680 | LOC_Os08g02210.2 | Bradi3g13670.1 | HORVU7Hr1G076030.9 | TRIUR3_07382 | Seita.6G088800.1 | Sobic.007G016600.1 | Zm00001d035081_T001 |
| orthologID_009905 | PH01000462G0610 | LOC_Os08g04790.1 | Bradi3g14860.1 | HORVU7Hr1G072980.1 | TRIUR3_23761 | Seita.6G065800.1 | Sobic.007G036800.1 | Zm00001d024646_T001 |
| orthologID_009190 | PH01002403G0100 | LOC_Os08g44520.1 | Bradi3g42920.1 | HORVU7Hr1G059320.3 | TRIUR3_23441 | Seita.6G249000.1 | Sobic.007G168200.1 | Zm00001d031937_T001 |
| orthologID_009567 | PH01000873G0580 | LOC_Os08g14660.1 | Bradi3g18940.1 | HORVU3Hr1G099430.2 | TRIUR3_16437 | Seita.6G093300.1 | Sobic.007G089800.1 | Zm00001d024420_T001 |
| orthologID_009265 | PH01000383G0260 | LOC_Os08g06130.1 | Bradi3g16490.1 | HORVU3Hr1G021980.1 | TRIUR3_13394 | Seita.6G055500.1 | Sobic.007G047600.3 | Zm00001d024544_T002 |
| orthologID_009785 | PH01003333G0140 | LOC_Os08g01710.1 | Bradi3g13307.2 | HORVU0Hr1G040190.2 | TRIUR3_28551 | Seita.6G008400.1 | Sobic.007G008600.1 | Zm00001d035028_T001 |
| orthologID_009804 | PH01003871G0100 | LOC_Os08g01600.1 | Bradi3g13260.1 | HORVU7Hr1G077560.2 | TRIUR3_22947 | Seita.6G007100.1 | Sobic.007G006700.1 | Zm00001d034990_T001 |
| orthologID_009776 | PH01000852G0270 | LOC_Os08g39130.1 | Bradi3g39610.1 | HORVU7Hr1G049380.2 | TRIUR3_07639 | Seita.6G199900.1 | Sobic.007G216401.1 | Zm00001d052854_T001 |
| orthologID_009370 | PH01001830G0080 | LOC_Os08g36440.1 | Bradi3g37920.1 | HORVU7Hr1G045630.2 | TRIUR3_21112 | Seita.6G178100.1 | Sobic.007G153600.1 | Zm00001d031773_T003 |
| orthologID_009748 | PH01000161G0830 | LOC_Os08g42950.1 | Bradi3g41860.1 | HORVU7Hr1G056410.2 | TRIUR3_06529 | Seita.6G231400.1 | Sobic.007G184200.3 | Zm00001d031705_T001 |
| orthologID_008933 | PH01000751G0410 | LOC_Os07g04160.2 | Bradi5g02990.3 | HORVU6Hr1G030430.1 | TRIUR3_14867 | Seita.7G037600.1 | Sobic.007G143700.1 | Zm00001d032127_T002 |
| orthologID_008363 | PH01000083G0260 | LOC_Os05g45270.1 | Bradi1g52250.1 | HORVU7Hr1G001290.1 | TRIUR3_05054 | Seita.6G084200.1 | Sobic.007G220900.2 | Zm00001d052865_T001 |
| orthologID_009564 | PH01001401G0380 | LOC_Os08g02340.1 | Bradi3g13750.2 | HORVU7Hr1G075360.2 | TRIUR3_11987 | Seita.6G085600.1 | Sobic.007G018300.1 | Zm00001d035087_T002 |
| orthologID_006542 | PH01001046G0420 | LOC_Os09g04810.1 | Bradi2g08410.2 | HORVU3Hr1G032920.1 | TRIUR3_21162 | Seita.6G075300.1 | Sobic.007G029200.1 | Zm00001d035199_T001 |
| orthologID_009821 | PH01000289G0250 | LOC_Os08g12850.1 | Bradi3g18320.1 | HORVU6Hr1G015490.1 | TRIUR3_25867 | Seita.6G021600.1 | Sobic.007G079700.1 | Zm00001d024347_T001 |
| orthologID_007643 | PH01000325G0710 | LOC_Os07g38300.1 | Bradi1g24110.3 | HORVU2Hr1G038570.2 | TRIUR3_32329 | Seita.6G013200.1 | Sobic.007G013400.2 | Zm00001d049694_T004 |
| orthologID_009570 | PH01002663G0030 | LOC_Os08g31340.1 | Bradi3g35490.1 | HORVU1Hr1G072500.1 | TRIUR3_00913 | Seita.6G140600.1 | Sobic.007G123400.1 | Zm00001d050277_T001 |
| orthologID_009345 | PH01000853G0250 | LOC_Os03g38000.1 | Bradi3g16150.1 | HORVU5Hr1G034770.1 | TRIUR3_22767 | Seita.6G050300.1 | Sobic.007G053300.1 | Zm00001d024511_T001 |
| orthologID_009726 | PH01001550G0390 | LOC_Os08g36140.1 | Bradi3g37770.1 | HORVU2Hr1G011960.1 | TRIUR3_01609 | Seita.6G176000.1 | Sobic.007G151900.1 | Zm00001d031738_T001 |
| orthologID_009530 | PH01007976G0010 | LOC_Os08g02060.1 | Bradi3g13560.1 | HORVU7Hr1G076320.1 | TRIUR3_13652 | Seita.6G013300.1 | Sobic.007G013500.1 | Zm00001d035052_T001 |
| orthologID_009171 | PH01000462G0550 | LOC_Os08g04810.1 | Bradi3g14840.1 | HORVU7Hr1G072940.3 | TRIUR3_30278 | Seita.6G065600.1 | Sobic.007G037000.1 | Zm00001d024645_T001 |
| orthologID_009721 | PH01000877G0280 | LOC_Os08g01220.1 | Bradi3g13027.1 | HORVU7Hr1G078320.4 | TRIUR3_34190 | Seita.6G003000.1 | Sobic.007G001900.1 | Zm00001d034977_T001 |

| | | | | | | | | |
|---|---|---|---|---|---|---|---|---|
| orthologID_009194 | PH01000853G0140 | LOC_Os08g06530.1 | Bradi3g16187.1 | HORVU7Hr1G067110.2 | TRIUR3_33886 | Seita.6G050900.1 | Sobic.007G052400.2 | Zm00001d024519_T005 |
| orthologID_009218 | PH01002106G0130 | LOC_Os08g44000.1 | Bradi3g42580.1 | HORVU7Hr1G058030.3 | TRIUR3_18085 | Seita.6G247600.1 | Sobic.007G173800.1 | Zm00001d031997_T001 |
| orthologID_009585 | PH01000729G0350 | LOC_Os08g33900.1 | Bradi3g37080.1 | HORVU7Hr1G031990.2 | TRIUR3_30489 | Seita.6G159800.1 | Sobic.007G138300.1 | Zm00001d032183_T002 |
| orthologID_009837 | PH01154077G0010 | LOC_Os08g33082.1 | Bradi3g36277.1 | HORVU7Hr1G043410.2 | TRIUR3_22290 | Seita.6G153000.1 | Sobic.007G132200.1 | Zm00001d032245_T003 |
| orthologID_006642 | PH01002711G0020 | LOC_Os01g74180.1 | Bradi2g62270.1 | HORVU3Hr1G115880.4 | TRIUR3_01874 | Seita.5G469000.1 | Sobic.007G019300.1 | Zm00001d035093_T002 |
| orthologID_009087 | PH01002275G0130 | LOC_Os08g34070.1 | Bradi3g37000.1 | HORVU7Hr1G061180.4 | TRIUR3_28355 | Seita.6G161900.1 | Sobic.007G139700.1 | Zm00001d050440_T004 |
| orthologID_009324 | PH01001419G0180 | LOC_Os08g35160.2 | Bradi3g37480.1 | HORVU1Hr1G071180.6 | TRIUR3_20680 | Seita.6G170300.1 | Sobic.007G148000.3 | Zm00001d032084_T001 |
| orthologID_009270 | PH01001652G0290 | LOC_Os08g44770.1 | Bradi3g43070.1 | HORVU7Hr1G060130.2 | TRIUR3_28595 | Seita.6G251600.1 | Sobic.007G166600.1 | Zm00001d031908_T001 |
| orthologID_009356 | PH01000394G0800 | LOC_Os08g20610.1 | Bradi3g20110.3 | HORVU7Hr1G062240.7 | TRIUR3_35292 | Seita.6G180600.1 | Sobic.007G155500.2 | Zm00001d031782_T001 |
| orthologID_009477 | PH01000462G0070 | LOC_Os08g05650.1 | Bradi3g17050.2 | HORVU7Hr1G071940.2 | TRIUR3_20514 | Seita.6G060600.1 | Sobic.007G041500.1 | Zm00001d049572_T004 |
| orthologID_008353 | PH01002199G0150 | LOC_Os02g11030.1 | Bradi1g36880.2 | HORVU4Hr1G043660.1 | TRIUR3_10745 | Seita.2G151400.1 | Sobic.007G012900.1 | Zm00001d049703_T001 |
| orthologID_009389 | PH01000117G0310 | LOC_Os08g42470.1 | Bradi3g41510.1 | HORVU7Hr1G055180.1 | TRIUR3_31628 | Seita.6G226500.1 | Sobic.007G189400.1 | Zm00001d031665_T001 |
| orthologID_009795 | PH01000104G0480 | LOC_Os08g38020.1 | Bradi3g38840.1 | HORVU7Hr1G037230.2 | TRIUR3_16978 | Seita.6G190600.1 | Sobic.007G224400.1 | Zm00001d031272_T001 |
| orthologID_009740 | PH01001046G0340 | LOC_Os08g03630.1 | Bradi3g14330.1 | HORVU7Hr1G073860.2 | TRIUR3_17800 | Seita.6G075200.1 | Sobic.007G029300.1 | Zm00001d035200_T001 |
| orthologID_009203 | PH01002324G0320 | LOC_Os04g25400.2 | Bradi3g12690.1 | HORVU0Hr1G040200.12 | TRIUR3_14417 | Seita.6G144300.1 | Sobic.007G162600.1 | Zm00001d031860_T002 |
| orthologID_009224 | PH01000399G0290 | LOC_Os08g29110.1 | Bradi3g35050.1 | HORVU4Hr1G012560.14 | TRIUR3_11458 | Seita.6G129100.1 | Sobic.007G116100.1 | Zm00001d032337_T001 |
| orthologID_009546 | PH01005522G0100 | LOC_Os08g01100.1 | Bradi3g12920.1 | HORVU7Hr1G078760.2 | TRIUR3_35367 | Seita.6G001400.1 | Sobic.007G000500.1 | Zm00001d052749_T001 |
| orthologID_009629 | PH01001470G0430 | LOC_Os08g41420.1 | Bradi3g40920.1 | HORVU7Hr1G053230.2 | TRIUR3_17615 | Seita.6G219600.1 | Sobic.007G196900.1 | Zm00001d031573_T001 |
| orthologID_009558 | PH01002058G0170 | LOC_Os08g14580.1 | Bradi3g18890.1 | HORVU3Hr1G109780.2 | TRIUR3_03377 | Seita.6G091500.1 | Sobic.007G088200.2 | Zm00001d024413_T001 |
| orthologID_009800 | PH01000003G0200 | LOC_Os08g02840.1 | Bradi3g14070.1 | HORVU7Hr1G074810.4 | TRIUR3_26253 | Seita.6G081100.1 | Sobic.007G024200.1 | Zm00001d049647_T001 |
| orthologID_008041 | PH01157543G0010 | LOC_Os07g24090.1 | Bradi1g16160.2 | HORVU1Hr1G067320.1 | TRIUR3_04793 | Seita.6G130600.1 | Sobic.007G114700.3 | Zm00001d050234_T001 |
| orthologID_009358 | PH01001954G0010 | LOC_Os08g35060.1 | Bradi3g37410.2 | HORVU2Hr1G116300.5 | TRIUR3_12011 | Seita.6G169400.1 | Sobic.007G147000.1 | Zm00001d050090_T001 |
| orthologID_009573 | PH01002747G0230 | LOC_Os10g11110.1 | Bradi3g12940.1 | HORVU7Hr1G078750.7 | TRIUR3_16175 | Seita.6G001800.1 | Sobic.007G000800.1 | Zm00001d052747_T001 |
| orthologID_009199 | PH01002097G0170 | LOC_Os08g34340.1 | Bradi3g36830.3 | HORVU7Hr1G062120.2 | TRIUR3_22989 | Seita.6G164600.1 | Sobic.007G142600.1 | Zm00001d032145_T001 |
| orthologID_009752 | PH01000594G0430 | LOC_Os08g39880.1 | Bradi3g40020.1 | HORVU0Hr1G020800.3 | TRIUR3_23873 | Seita.6G205400.1 | Sobic.007G210300.3 | Zm00001d031450_T001 |
| orthologID_009497 | PH01000259G1010 | LOC_Os08g40860.1 | Bradi3g40480.1 | HORVU7Hr1G052060.4 | TRIUR3_25998 | Seita.6G212500.1 | Sobic.007G203000.1 | Zm00001d049760_T001 |
| orthologID_009254 | PH01000299G0320 | LOC_Os08g44320.1 | Bradi3g42730.1 | HORVU7Hr1G058760.3 | TRIUR3_23596 | Seita.6G244600.1 | Sobic.007G171400.1 | Zm00001d031963_T001 |
| orthologID_009747 | PH01000150G0830 | LOC_Os08g39560.1 | Bradi3g39900.1 | HORVU5Hr1G107000.6 | TRIUR3_01878 | Seita.6G204400.1 | Sobic.007G211300.4 | Zm00001d031436_T001 |
| orthologID_009308 | PH01000427G0640 | LOC_Os08g43370.1 | Bradi3g42160.1 | HORVU7Hr1G056980.1 | TRIUR3_30322 | Seita.6G236700.1 | Sobic.007G179900.2 | Zm00001d032045_T001 |
| orthologID_009461 | PH01000623G0200 | LOC_Os08g40940.1 | Bradi3g40382.1 | HORVU7Hr1G051650.1 | TRIUR3_18233 | Seita.6G211100.1 | Sobic.007G204700.1 | Zm00001d031510_T001 |
| orthologID_009388 | PH01003169G0140 | LOC_Os08g43290.1 | Bradi3g42070.1 | HORVU7Hr1G056650.1 | TRIUR3_16362 | Seita.6G235600.1 | Sobic.007G180700.1 | Zm00001d052736_T001 |
| orthologID_009185 | PH01000008G1080 | LOC_Os08g33700.1 | Bradi3g37120.2 | HORVU4Hr1G009150.1 | TRIUR3_14403 | Seita.6G159500.1 | Sobic.007G137800.3 | Zm00001d032185_T001 |
| orthologID_009354 | PH01001490G0340 | LOC_Os08g43560.1 | Bradi3g42340.1 | HORVU7Hr1G057430.1 | TRIUR3_11069 | Seita.6G239800.1 | Sobic.007G177000.1 | Zm00001d053223_T001 |
| orthologID_006154 | PH01000018G2640 | LOC_Os12g08140.1 | Bradi2g50550.5 | HORVU3Hr1G073540.14 | TRIUR3_04581 | Seita.5G326600.1 | Sobic.007G006800.3 | Zm00001d034989_T001 |
| orthologID_009157 | PH01001305G0100 | LOC_Os08g30480.1 | Bradi3g35470.1 | HORVU4Hr1G019140.5 | TRIUR3_23875 | Seita.6G134100.1 | Sobic.007G119600.1 | Zm00001d050256_T001 |
| orthologID_009575 | PH01000299G0550 | LOC_Os08g44430.2 | Bradi3g42850.1 | HORVU7Hr1G059240.4 | TRIUR3_04369 | Seita.6G243100.1 | Sobic.007G168600.4 | Zm00001d031942_T002 |
| orthologID_009234 | PH01000435G0280 | LOC_Os08g38780.1 | Bradi3g39280.1 | HORVU7Hr1G048760.2 | TRIUR3_12611 | Seita.6G196700.1 | Sobic.007G219500.1 | Zm00001d031310_T001 |
| orthologID_009717 | PH01001168G0440 | LOC_Os08g02190.1 | Bradi3g13640.1 | HORVU7Hr1G076060.1 | TRIUR3_07384 | Seita.6G088400.1 | Sobic.007G016400.1 | Zm00001d049685_T001 |
| orthologID_009485 | PH01007031G0020 | LOC_Os08g43190.1 | Bradi3g42030.1 | HORVU7Hr1G056770.2 | TRIUR3_13245 | Seita.6G235000.1 | Sobic.007G181600.1 | Zm00001d031727_T004 |
| orthologID_009906 | PH01002954G0010 | LOC_Os08g44620.1 | Bradi3g43000.1 | HORVU7Hr1G059900.2 | TRIUR3_34039 | Seita.6G250600.1 | Sobic.007G167300.1 | Zm00001d031925_T001 |
| orthologID_009780 | PH01001093G0240 | LOC_Os08g15650.1 | Bradi3g19130.1 | HORVU0Hr1G039280.1 | TRIUR3_33262 | Seita.6G094700.1 | Sobic.007G090300.1 | Zm00001d021250_T001 |
| orthologID_009439 | PH01001046G0270 | LOC_Os02g21510.1 | Bradi3g14400.1 | HORVU7Hr1G073570.4 | TRIUR3_03007 | Seita.6G074200.1 | Sobic.007G030600.1 | Zm00001d024709_T002 |
| orthologID_006778 | PH01001383G0140 | LOC_Os08g24390.1 | Bradi2g31560.1 | HORVU7Hr1G078020.23 | TRIUR3_25179 | Seita.6G114500.1 | Sobic.007G103400.1 | Zm00001d030581_T001 |
| orthologID_009610 | PH01000462G0080 | LOC_Os08g05640.2 | Bradi3g17060.3 | HORVU7Hr1G072050.4 | TRIUR3_35122 | Seita.6G060700.1 | Sobic.007G041400.1 | Zm00001d024605_T001 |
| orthologID_009317 | PH01003460G0030 | LOC_Os08g13320.1 | Bradi3g18527.3 | HORVU5Hr1G105810.1 | TRIUR3_14745 | Seita.6G018000.1 | Sobic.007G084100.1 | Zm00001d024357_T001 |
| orthologID_009635 | PH01000594G0730 | LOC_Os08g39980.1 | Bradi3g40060.1 | HORVU7Hr1G050770.2 | TRIUR3_28812 | Seita.6G206100.1 | Sobic.007G209500.1 | Zm00001d031454_T001 |
| orthologID_009288 | PH01004035G0040 | LOC_Os08g17320.1 | Bradi3g19560.1 | HORVU1Hr1G088440.2 | TRIUR3_23269 | Seita.6G102300.1 | Sobic.007G111200.1 | Zm00001d024134_T001 |
| orthologID_008872 | PH01002155G0190 | LOC_Os06g25439.1 | Bradi5g22240.1 | HORVU7Hr1G083360.1 | TRIUR3_04356 | Seita.9G195600.1 | Sobic.007G157300.1 | Zm00001d031799_T001 |
| orthologID_009278 | PH01000289G0890 | LOC_Os08g10604.1 | Bradi3g17880.1 | HORVU4Hr1G073610.10 | TRIUR3_21416 | Seita.6G026700.1 | Sobic.007G075600.1 | Zm00001d024308_T001 |
| orthologID_009895 | PH01000299G0560 | LOC_Os08g44440.1 | Bradi3g42860.1 | HORVU7Hr1G059270.1 | TRIUR3_30556 | Seita.6G248600.1 | Sobic.007G168500.1 | Zm00001d031941_T001 |

| | | | | | | | | |
|---|---|---|---|---|---|---|---|---|
| orthologID_009236 | PH01000383G0420 | LOC_Os08g06090.1 | Bradi3g16540.1 | HORVU7Hr1G071020.5 | TRIUR3_21410 | Seita.6G056400.1 | Sobic.007G046400.1 | Zm00001d049545_T001 |
| orthologID_009841 | PH01003692G0010 | LOC_Os08g36170.1 | Bradi3g37800.1 | HORVU7Hr1G043120.1 | TRIUR3_16991 | Seita.6G176300.1 | Sobic.007G152101.1 | Zm00001d031741_T001 |
| orthologID_009572 | PH01000150G1310 | LOC_Os08g39310.1 | Bradi3g39760.1 | HORVU7Hr1G049900.8 | TRIUR3_23070 | Seita.6G202500.1 | Sobic.007G213600.1 | Zm00001d052873_T001 |
| orthologID_009080 | PH01003170G0110 | LOC_Os08g17400.1 | Bradi3g19640.1 | HORVU7Hr1G066670.4 | TRIUR3_27317 | Seita.6G103400.1 | Sobic.007G111600.1 | Zm00001d050023_T001 |
| orthologID_009453 | PH01000289G0280 | LOC_Os08g12840.1 | Bradi3g18306.1 | HORVU2Hr1G089160.4 | TRIUR3_24581 | Seita.6G021700.1 | Sobic.007G079600.1 | Zm00001d049351_T002 |
| orthologID_009757 | PH01000612G0270 | LOC_Os08g34750.1 | Bradi3g37270.1 | HORVU7Hr1G060790.8 | TRIUR3_21384 | Seita.6G167600.1 | Sobic.007G145200.1 | Zm00001d032111_T003 |
| orthologID_009843 | PH01005189G0110 | LOC_Os08g04440.1 | Bradi3g14640.1 | HORVU7Hr1G073220.1 | TRIUR3_27646 | Seita.6G068900.1 | Sobic.007G034400.1 | Zm00001d024675_T001 |
| orthologID_009870 | PH01000670G0070 | LOC_Os08g32120.1 | Bradi3g35880.1 | HORVU2Hr1G113300.3 | TRIUR3_01780 | Seita.6G145300.1 | Sobic.007G126800.1 | Zm00001d032318_T001 |
| orthologID_009791 | PH01003078G0280 | LOC_Os01g25880.1 | Bradi3g06370.1 | HORVU3Hr1G001670.1 | TRIUR3_05321 | Seita.5G164500.1 | Sobic.007G140300.1 | Zm00001d032160_T001 |
| orthologID_009559 | PH01003910G0050 | LOC_Os08g02490.1 | Bradi3g13871.1 | HORVU7Hr1G075040.9 | TRIUR3_20490 | Seita.6G084100.1 | Sobic.007G020300.1 | Zm00001d049661_T001 |
| orthologID_009376 | PH01001770G0220 | LOC_Os08g10080.1 | Bradi3g17287.1 | HORVU7Hr1G072670.3 | TRIUR3_29250 | Seita.6G032700.1 | Sobic.007G071000.1 | Zm00001d024268_T001 |
| orthologID_009580 | PH01000303G0730 | LOC_Os08g08020.1 | Bradi3g15580.1 | HORVU3Hr1G112440.37 | TRIUR3_34390 | Seita.6G043800.1 | Sobic.007G061600.2 | Zm00001d024170_T001 |
| orthologID_009644 | PH01003138G0030 | LOC_Os08g10020.1 | Bradi3g17640.1 | HORVU6Hr1G085170.1 | TRIUR3_28442 | Seita.6G032200.1 | Sobic.007G071500.1 | Zm00001d049387_T001 |
| orthologID_009536 | PH01000104G1110 | LOC_Os08g37500.1 | Bradi3g38630.1 | HORVU3Hr1G067590.8 | TRIUR3_24640 | Seita.6G188000.1 | Sobic.007G226300.1 | Zm00001d052795_T001 |
| orthologID_009810 | PH01004673G0010 | LOC_Os08g01840.1 | Bradi3g13410.1 | HORVU7Hr1G076900.3 | TRIUR3_24898 | Seita.1G024700.1 | Sobic.007G009700.1 | Zm00001d049712_T006 |
| orthologID_009205 | PH01002506G0280 | LOC_Os08g41480.1 | Bradi3g40960.1 | HORVU7Hr1G053390.5 | TRIUR3_34064 | Seita.6G219900.1 | Sobic.007G196500.1 | Zm00001d040370_T001 |
| orthologID_009873 | PH01001044G0240 | LOC_Os08g34270.2 | Bradi3g36900.1 | HORVU7Hr1G089040.1 | TRIUR3_11317 | Seita.6G163300.1 | Sobic.007G141100.1 | Zm00001d032153_T002 |
| orthologID_009541 | PH01003871G0070 | LOC_Os12g18880.1 | Bradi3g13270.1 | HORVU7Hr1G077420.1 | TRIUR3_25682 | Seita.6G007600.1 | Sobic.007G007900.1 | Zm00001d049723_T001 |
| orthologID_006527 | PH01000344G0430 | LOC_Os09g20010.1 | Bradi2g01490.3 | HORVU1Hr1G000310.1 | TRIUR3_08141 | Seita.3G116100.1 | Sobic.007G203700.1 | Zm00001d031520_T001 |
| orthologID_009237 | PH01000161G0230 | LOC_Os08g15204.1 | Bradi3g19070.1 | HORVU3Hr1G089950.1 | TRIUR3_25066 | Seita.6G095500.1 | Sobic.007G091000.1 | Zm00001d050225_T001 |
| orthologID_009599 | PH01000623G0490 | LOC_Os08g40890.1 | Bradi3g40450.1 | HORVU7Hr1G051960.1 | TRIUR3_26421 | Seita.6G212100.1 | Sobic.007G203400.1 | Zm00001d031523_T001 |
| orthologID_009233 | PH01001046G0190 | LOC_Os08g04150.1 | Bradi3g14440.1 | HORVU7Hr1G073500.2 | TRIUR3_03002 | Seita.6G073700.1 | Sobic.007G031100.1 | Zm00001d049612_T001 |
| orthologID_009615 | PH01000123G0940 | LOC_Os02g07450.2 | Bradi3g05270.1 | HORVU6Hr1G032200.3 | TRIUR3_16626 | Seita.1G071900.1 | Sobic.007G063200.2 | Zm00001d024230_T001 |
| orthologID_008333 | PH01000946G0240 | LOC_Os11g16540.1 | Bradi1g08940.1 | HORVU5Hr1G094360.1 | TRIUR3_11829 | Seita.6G027800.1 | Sobic.007G074600.1 | Zm00001d049379_T013 |
| orthologID_009427 | PH01000003G0870 | LOC_Os08g02380.1 | Bradi3g13780.4 | HORVU7Hr1G075310.29 | TRIUR3_27576 | Seita.6G085300.1 | Sobic.007G018700.1 | Zm00001d035091_T001 |
| orthologID_009770 | PH01000729G0220 | LOC_Os08g33710.1 | Bradi3g37130.4 | HORVU2Hr1G062880.8 | TRIUR3_20252 | Seita.6G159400.1 | Sobic.007G137700.1 | Zm00001d032186_T001 |
| orthologID_008653 | PH01000649G0720 | LOC_Os10g13940.1 | Bradi3g23150.1 | HORVU1Hr1G068960.4 | TRIUR3_06851 | Seita.9G204700.1 | Sobic.007G020600.1 | Zm00001d035115_T004 |
| orthologID_007097 | PH01000205G0530 | LOC_Os11g20689.1 | Bradi4g19975.1 | HORVU4Hr1G012650.8 | TRIUR3_15774 | Seita.8G109500.1 | Sobic.007G191600.1 | Zm00001d031643_T001 |
| orthologID_009913 | PH01001457G0150 | LOC_Os08g27824.1 | Bradi3g21700.1 | HORVU7Hr1G089720.2 | TRIUR3_15121 | Seita.6G125900.1 | Sobic.007G106700.1 | Zm00001d049313_T004 |
| orthologID_009550 | PH01004430G0090 | LOC_Os08g05280.1 | Bradi3g17550.1 | HORVU7Hr1G072750.10 | TRIUR3_13137 | Seita.6G064500.1 | Sobic.007G037900.1 | Zm00001d024640_T003 |
| orthologID_009489 | PH01000117G0680 | LOC_Os08g42350.1 | Bradi3g41410.1 | HORVU7Hr1G054710.2 | TRIUR3_18312 | Seita.6G225300.1 | Sobic.007G191200.1 | Zm00001d031647_T001 |
| orthologID_009454 | PH01001052G0090 | LOC_Os08g29170.1 | Bradi3g34980.1 | HORVU6Hr1G025730.2 | TRIUR3_29213 | Seita.6G128600.1 | Sobic.007G116700.1 | Zm00001d032332_T001 |
| orthologID_009315 | PH01000796G0480 | LOC_Os02g21280.1 | Bradi3g10980.1 | HORVU3Hr1G032570.1 | TRIUR3_26793 | Seita.1G143000.1 | Sobic.007G096700.1 | Zm00001d050179_T001 |
| orthologID_009320 | PH01001044G0070 | LOC_Os08g34290.1 | Bradi3g36880.1 | HORVU0Hr1G020780.4 | TRIUR3_31153 | Seita.6G163700.1 | Sobic.007G141500.1 | Zm00001d050412_T002 |
| orthologID_009884 | PH01000323G0970 | LOC_Os08g38570.1 | Bradi3g39160.1 | HORVU1Hr1G020620.7 | TRIUR3_17159 | Seita.6G194600.1 | Sobic.007G220300.1 | Zm00001d031291_T004 |
| orthologID_009306 | PH01000117G0420 | LOC_Os08g42430.1 | Bradi3g41495.1 | HORVU0Hr1G021040.1 | TRIUR3_25258 | Seita.6G226100.1 | Sobic.007G189900.1 | Zm00001d049865_T001 |
| orthologID_009136 | PH01000259G1320 | LOC_Os08g40930.1 | Bradi3g40410.1 | HORVU7Hr1G051710.9 | TRIUR3_13775 | Seita.6G211200.1 | Sobic.007G204600.1 | Zm00001d049753_T001 |
| orthologID_009464 | PH01005189G0020 | LOC_Os08g04390.2 | Bradi3g14590.1 | HORVU7Hr1G073310.4 | TRIUR3_32784 | Seita.6G069500.1 | Sobic.007G033800.1 | Zm00001d024679_T002 |
| orthologID_009198 | PH01002426G0020 | LOC_Os08g40200.1 | Bradi3g40230.1 | HORVU7Hr1G051280.1 | TRIUR3_35003 | Seita.6G208400.1 | Sobic.007G207400.4 | Zm00001d031486_T001 |
| orthologID_009521 | PH01005828G0050 | LOC_Os08g33140.1 | Bradi3g36357.1 | HORVU2Hr1G101900.5 | TRIUR3_19256 | Seita.6G154000.1 | Sobic.007G132500.1 | Zm00001d050359_T001 |
| orthologID_009396 | PH01000161G0100 | LOC_Os08g15080.1 | Bradi3g19040.1 | HORVU7Hr1G067800.6 | TRIUR3_32911 | Seita.6G096400.1 | Sobic.007G091600.1 | Zm00001d049346_T001 |
| orthologID_009782 | PH01003661G0090 | LOC_Os10g09930.1 | Bradi3g22697.2 | HORVU7Hr1G047120.2 | TRIUR3_33272 | Seita.9G116600.1 | Sobic.007G052800.1 | Zm00001d049502_T001 |
| orthologID_009823 | PH01000150G0690 | LOC_Os08g39630.1 | Bradi3g39927.1 | HORVU7Hr1G050530.1 | TRIUR3_26625 | Seita.6G204700.1 | Sobic.007G211000.3 | Zm00001d006578_T003 |
| orthologID_010144 | PH01003787G0060 | LOC_Os08g41000.1 | Bradi3g40357.1 | HORVU7Hr1G051640.1 | TRIUR3_06791 | Seita.6G210800.1 | Sobic.007G204900.2 | Zm00001d049748_T001 |
| orthologID_009082 | PH01001444G0340 | LOC_Os08g34220.1 | Bradi3g36920.1 | HORVU2Hr1G035220.1 | TRIUR3_12044 | Seita.6G163000.1 | Sobic.007G140800.1 | Zm00001d032156_T003 |
| orthologID_009918 | PH01000788G0760 | LOC_Os08g44940.1 | Bradi3g12550.1 | HORVU4Hr1G000880.4 | TRIUR3_07751 | Seita.6G253100.1 | Sobic.007G165100.1 | Zm00001d031882_T001 |
| orthologID_009924 | PH01000411G0760 | LOC_Os08g39320.1 | Bradi3g39771.1 | HORVU7Hr1G049890.6 | TRIUR3_23071 | Seita.6G202600.1 | Sobic.007G213500.1 | Zm00001d052874_T001 |
| orthologID_010183 | PH01002106G0340 | LOC_Os08g44180.1 | Bradi3g42627.3 | HORVU7Hr1G058180.9 | TRIUR3_25060 | Seita.2G400100.1 | Sobic.007G172900.1 | Zm00001d050498_T001 |
| orthologID_009422 | PH01000915G0720 | LOC_Os06g27790.1 | Bradi3g40131.1 | HORVU7Hr1G051030.9 | TRIUR3_25473 | Seita.6G207200.1 | Sobic.007G208400.1 | Zm00001d031471_T002 |
| orthologID_009393 | PH01000462G0040 | LOC_Os08g05670.1 | Bradi3g17027.4 | HORVU7Hr1G071900.42 | TRIUR3_23893 | Seita.6G060400.1 | Sobic.007G042400.1 | Zm00001d024601_T003 |

| | | | | | | | | |
|---|---|---|---|---|---|---|---|---|
| orthologID_009301 | PH01000853G0770 | LOC_Os08g07010.1 | Bradi3g16130.1 | HORVU7Hr1G014870.1 | TRIUR3_07051 | Seita.6G049400.1 | Sobic.007G054500.1 | Zm00001d024497_T001 |
| orthologID_007768 | PH01000750G0210 | LOC_Os03g52750.2 | Bradi1g09370.3 | HORVU4Hr1G011340.7 | TRIUR3_13657 | Seita.9G100500.1 | Sobic.001G098100.1 | Zm00001d033928_T001 |
| orthologID_008188 | PH01002084G0060 | LOC_Os03g63140.1 | Bradi1g01730.3 | HORVU5Hr1G121530.2 | TRIUR3_00718 | Seita.9G012000.1 | Sobic.001G013000.1 | Zm00001d012852_T003 |
| orthologID_008476 | PH01000430G0330 | LOC_Os03g20470.2 | Bradi1g63960.2 | HORVU4Hr1G052100.1 | TRIUR3_12287 | Seita.9G422100.1 | Sobic.001G389400.1 | Zm00001d028955_T001 |
| orthologID_009346 | PH01000059G1780 | LOC_Os10g39460.2 | Bradi3g32220.1 | HORVU1Hr1G052060.4 | TRIUR3_13821 | Seita.9G341000.1 | Sobic.001G312700.1 | Zm00001d029764_T001 |
| orthologID_008592 | PH01000376G0960 | LOC_Os03g56700.1 | Bradi1g06840.1 | HORVU5Hr1G100800.1 | TRIUR3_01924 | Seita.9G069100.1 | Sobic.001G070300.1 | Zm00001d013144_T001 |
| orthologID_007263 | PH01001088G0510 | LOC_Os11g40200.1 | Bradi4g13770.1 | HORVU5Hr1G082670.2 | TRIUR3_08308 | Seita.8G194700.1 | Sobic.001G204700.1 | Zm00001d048831_T001 |
| orthologID_008478 | PH01001259G0200 | LOC_Os03g01130.1 | Bradi1g78570.1 | HORVU0Hr1G009550.3 | TRIUR3_25311 | Seita.9G579500.1 | Sobic.001G543800.1 | Zm00001d027266_T001 |
| orthologID_009480 | PH01002259G0210 | LOC_Os10g31970.1 | Bradi3g27970.2 | HORVU1Hr1G038370.1 | TRIUR3_22775 | Seita.9G232900.1 | Sobic.001G230000.1 | Zm00001d032801_T021 |
| orthologID_009214 | PH01000012G0560 | LOC_Os10g35920.1 | Bradi3g30187.1 | HORVU1Hr1G045640.4 | TRIUR3_30802 | Seita.9G201100.1 | Sobic.001G202266.1 | Zm00001d032530_T001 |
| orthologID_008162 | PH01000181G1390 | LOC_Os03g19520.2 | Bradi1g64420.1 | HORVU4Hr1G053650.1 | TRIUR3_26142 | Seita.9G428700.1 | Sobic.001G395200.1 | Zm00001d028896_T006 |
| orthologID_007675 | PH01254916G0010 | LOC_Os03g58710.1 | Bradi1g05116.3 | HORVU5Hr1G057950.4 | TRIUR3_12859 | Seita.9G051100.1 | Sobic.001G052600.1 | Zm00001d034493_T001 |
| orthologID_008423 | PH01000147G1250 | LOC_Os03g16870.1 | Bradi1g66580.1 | HORVU4Hr1G059340.23 | TRIUR3_21477 | Seita.3G205200.1 | Sobic.001G420000.1 | Zm00001d028631_T001 |
| orthologID_007169 | PH01001970G0280 | LOC_Os11g16430.1 | Bradi4g20557.4 | HORVU4Hr1G013960.2 | TRIUR3_25821 | Seita.6G237100.1 | Sobic.001G017700.1 | Zm00001d034797_T002 |
| orthologID_007709 | PH01000589G0780 | LOC_Os03g56540.3 | Bradi1g06980.2 | HORVU5Hr1G099410.4 | TRIUR3_25308 | Seita.9G069800.1 | Sobic.001G071100.1 | Zm00001d034270_T001 |
| orthologID_007750 | PH01002028G0290 | LOC_Os03g49630.1 | Bradi1g10960.1 | HORVU4Hr1G006670.2 | TRIUR3_02351 | Seita.9G125100.1 | Sobic.001G122900.1 | Zm00001d033654_T004 |
| orthologID_008709 | PH01000081G1610 | LOC_Os03g06090.1 | Bradi1g74680.1 | HORVU4Hr1G072520.2 | TRIUR3_10039 | Seita.9G535200.1 | Sobic.001G499300.1 | Zm00001d027721_T001 |
| orthologID_007805 | PH01003184G0200 | LOC_Os03g60920.1 | Bradi1g03440.1 | HORVU5Hr1G114620.2 | TRIUR3_12155 | Seita.9G029900.1 | Sobic.001G031200.1 | Zm00001d012932_T001 |
| orthologID_007848 | PH01000614G0780 | LOC_Os03g50960.1 | Bradi1g10580.1 | HORVU4Hr1G008420.1 | TRIUR3_16048 | Seita.9G114800.1 | Sobic.001G112200.1 | Zm00001d033793_T001 |
| orthologID_007689 | PH01000365G0800 | LOC_Os03g62780.1 | Bradi1g01880.1 | HORVU5Hr1G120650.5 | TRIUR3_19934 | Seita.9G014000.1 | Sobic.001G015200.1 | Zm00001d034828_T002 |
| orthologID_009451 | PH01000560G1080 | LOC_Os10g32960.1 | Bradi3g28320.1 | HORVU1Hr1G039230.2 | TRIUR3_10871 | Seita.9G227600.1 | Sobic.001G224600.1 | Zm00001d032773_T003 |
| orthologID_009323 | PH01000741G0210 | LOC_Os10g34740.1 | Bradi3g29470.1 | HORVU1Hr1G044440.3 | TRIUR3_20807 | Seita.9G212700.1 | Sobic.001G211900.2 | Zm00001d032633_T004 |
| orthologID_007896 | PH01000081G1510 | LOC_Os03g06120.3 | Bradi1g74650.2 | HORVU4Hr1G072630.1 | TRIUR3_02105 | Seita.9G535100.1 | Sobic.001G499200.1 | Zm00001d027722_T003 |
| orthologID_007253 | PH01001299G0500 | LOC_Os11g26030.1 | Bradi4g19590.1 | HORVU5Hr1G002310.3 | TRIUR3_25678 | Seita.8G113000.1 | Sobic.001G246500.1 | Zm00001d032944_T001 |
| orthologID_008586 | PH01001584G0330 | LOC_Os03g47940.1 | Bradi1g12350.1 | HORVU4Hr1G004100.1 | TRIUR3_15388 | Seita.9G138600.1 | Sobic.001G135600.1 | Zm00001d033522_T001 |
| orthologID_008729 | PH01002899G0100 | LOC_Os03g55720.2 | Bradi1g07530.3 | HORVU5Hr1G097870.1 | TRIUR3_15043 | Seita.9G078500.1 | Sobic.001G078000.1 | Zm00001d034179_T001 |
| orthologID_009922 | PH01000059G1540 | LOC_Os10g39230.1 | Bradi3g32160.1 | HORVU1Hr1G051970.2 | TRIUR3_30958 | Seita.9G341700.1 | Sobic.001G313400.1 | Zm00001d029754_T004 |
| orthologID_009705 | PH01000012G1890 | LOC_Os10g36680.1 | Bradi3g30680.3 | HORVU1Hr1G047410.5 | TRIUR3_13173 | Seita.9G194800.1 | Sobic.001G197100.1 | Zm00001d032478_T001 |
| orthologID_008226 | PH01002084G0030 | LOC_Os03g63270.1 | Bradi1g01750.1 | HORVU5Hr1G121570.9 | TRIUR3_17308 | Seita.9G011800.1 | Sobic.001G012800.1 | Zm00001d012850_T001 |
| orthologID_009264 | PH01000103G0240 | LOC_Os10g28040.1 | Bradi3g26800.1 | HORVU1Hr1G034070.10 | TRIUR3_27913 | Seita.9G263000.1 | Sobic.001G248100.1 | Zm00001d014175_T004 |
| orthologID_008085 | PH01001799G0020 | LOC_Os03g38640.1 | Bradi1g15460.1 | HORVU2Hr1G079600.1 | TRIUR3_33879 | Seita.9G179400.1 | Sobic.001G180300.1 | Zm00001d013794_T001 |
| orthologID_007833 | PH01000138G0660 | LOC_Os03g16570.2 | Bradi1g66761.1 | HORVU4Hr1G060040.2 | TRIUR3_21710 | Seita.9G461900.1 | Sobic.001G422700.1 | Zm00001d028593_T002 |
| orthologID_008095 | PH01000542G0840 | LOC_Os03g21490.2 | Bradi1g63380.1 | HORVU4Hr1G050780.2 | TRIUR3_21788 | Seita.9G414900.1 | Sobic.001G382800.1 | Zm00001d029039_T001 |
| orthologID_008064 | PH01002430G0030 | LOC_Os03g20860.1 | Bradi1g63730.1 | HORVU4Hr1G051620.3 | TRIUR3_22595 | Seita.9G419500.1 | Sobic.001G387200.1 | Zm00001d028981_T002 |
| orthologID_008481 | PH01000033G1960 | LOC_Os03g17590.1 | Bradi1g65890.1 | HORVU4Hr1G057360.2 | TRIUR3_19534 | Seita.9G445000.1 | Sobic.001G411200.1 | Zm00001d047760_T001 |
| orthologID_008038 | PH01000113G1150 | LOC_Os03g07790.1 | Bradi1g73040.1 | HORVU4Hr1G055690.6 | TRIUR3_04866 | Seita.9G521900.1 | Sobic.001G487200.1 | Zm00001d027868_T004 |
| orthologID_007657 | PH01001798G0220 | LOC_Os03g61030.1 | Bradi1g03400.1 | HORVU5Hr1G114890.4 | TRIUR3_11998 | Seita.9G028800.1 | Sobic.001G030200.1 | Zm00001d034699_T001 |
| orthologID_008589 | PH01001624G0040 | LOC_Os03g21830.1 | Bradi1g63147.2 | HORVU4Hr1G050130.1 | TRIUR3_22625 | Seita.9G412500.1 | Sobic.001G380500.1 | Zm00001d029065_T001 |
| orthologID_007696 | PH01001995G0090 | LOC_Os03g40170.1 | Bradi1g15090.3 | HORVU3Hr1G080040.2 | TRIUR3_23419 | Seita.9G174000.1 | Sobic.001G173600.1 | Zm00001d033170_T001 |
| orthologID_008355 | PH01001584G0350 | LOC_Os03g47970.1 | Bradi1g12330.5 | HORVU4Hr1G004080.9 | TRIUR3_15386 | Seita.9G138400.1 | Sobic.001G135400.1 | Zm00001d033523_T001 |
| orthologID_008681 | PH01002172G0060 | LOC_Os03g53590.2 | Bradi1g08780.1 | HORVU5Hr1G094660.1 | TRIUR3_16983 | Seita.9G094500.1 | Sobic.001G091400.1 | Zm00001d034000_T005 |
| orthologID_008394 | PH01000537G0430 | LOC_Os03g15340.1 | Bradi1g67580.1 | HORVU4Hr1G062010.1 | TRIUR3_25890 | Seita.9G463100.1 | Sobic.001G431100.1 | Zm00001d028486_T001 |
| orthologID_007572 | PH01003290G0010 | LOC_Os03g64310.1 | Bradi1g00563.2 | HORVU5Hr1G125660.1 | TRIUR3_14954 | Seita.9G001900.1 | Sobic.001G001800.2 | Zm00001d012799_T007 |
| orthologID_009275 | PH01004021G0080 | LOC_Os10g33270.1 | Bradi3g28470.3 | HORVU1Hr1G039870.1 | TRIUR3_05199 | Seita.9G226000.1 | Sobic.001G222600.1 | Zm00001d032736_T001 |
| orthologID_009327 | PH01002340G0030 | LOC_Os10g42540.1 | Bradi3g34360.1 | HORVU1Hr1G050740.2 | TRIUR3_14766 | Seita.9G309200.1 | Sobic.001G286200.1 | Zm00001d030080_T001 |
| orthologID_008559 | PH01000017G0730 | LOC_Os03g02720.1 | Bradi1g77280.2 | HORVU4Hr1G085740.6 | TRIUR3_14181 | Seita.9G564800.1 | Sobic.001G530000.1 | Zm00001d027384_T001 |
| orthologID_007612 | PH01000468G0630 | LOC_Os03g20680.1 | Bradi1g63816.1 | HORVU4Hr1G051780.1 | TRIUR3_33086 | Seita.9G420500.1 | Sobic.001G388100.1 | Zm00001d047567_T001 |
| orthologID_008690 | PH01002698G0260 | LOC_Os03g48080.1 | Bradi1g12230.1 | HORVU4Hr1G004240.1 | TRIUR3_13982 | Seita.9G137100.1 | Sobic.001G134400.1 | Zm00001d033537_T002 |
| orthologID_009178 | PH01000186G0420 | LOC_Os10g34670.1 | Bradi3g29410.1 | HORVU1Hr1G043250.1 | TRIUR3_24065 | Seita.9G213600.1 | Sobic.001G212800.1 | Zm00001d032649_T001 |
| orthologID_009677 | PH01000841G0440 | LOC_Os01g10290.1 | Bradi3g33350.1 | HORVU1Hr1G056110.1 | TRIUR3_32121 | Seita.9G323500.1 | Sobic.001G298300.1 | Zm00001d000178_T001 |

| | | | | | | | | |
|---|---|---|---|---|---|---|---|---|
| orthologID_008485 | PH01004629G0100 | LOC_Os03g61980.1 | Bradi1g02575.1 | HORVU5Hr1G118020.4 | TRIUR3_01600 | Seita.9G021400.1 | Sobic.001G022700.1 | Zm00001d034769_T001 |
| orthologID_008480 | PH01001627G0410 | LOC_Os03g64370.1 | Bradi1g00491.2 | HORVU5Hr1G124580.1 | TRIUR3_04895 | Seita.9G001600.1 | Sobic.001G001200.1 | Zm00001d034933_T001 |
| orthologID_007791 | PH01002110G0080 | LOC_Os05g15890.1 | Bradi1g14910.4 | HORVU2Hr1G037590.3 | TRIUR3_23149 | Seita.9G188400.1 | Sobic.001G191500.1 | Zm00001d013828_T002 |
| orthologID_008224 | PH01003371G0130 | LOC_Os03g58780.1 | Bradi1g05070.1 | HORVU5Hr1G107360.1 | TRIUR3_01315 | Seita.9G050100.1 | Sobic.001G051700.1 | Zm00001d034498_T007 |
| orthologID_007988 | PH01001253G0290 | LOC_Os03g52400.1 | Bradi1g09577.1 | HORVU4Hr1G010700.2 | TRIUR3_23700 | Seita.9G103300.1 | Sobic.001G100400.2 | Zm00001d013332_T001 |
| orthologID_008708 | PH01001293G0060 | LOC_Os03g46260.1 | Bradi1g12960.5 | HORVU4Hr1G002700.3 | TRIUR3_26044 | Seita.9G147300.1 | Sobic.001G144200.1 | Zm00001d013595_T001 |
| orthologID_008570 | PH01001015G0380 | LOC_Os05g44080.1 | Bradi1g62770.7 | HORVU4Hr1G047790.1 | TRIUR3_17153 | Seita.9G408500.1 | Sobic.001G375600.3 | Zm00001d047497_T003 |
| orthologID_007280 | PH01001214G0460 | LOC_Os04g52850.1 | Bradi4g02970.1 | HORVU5Hr1G017680.1 | TRIUR3_20894 | Seita.9G045200.1 | Sobic.001G047100.1 | Zm00001d013040_T008 |
| orthologID_008054 | PH01004012G0050 | LOC_Os03g01016.2 | Bradi1g78690.1 | HORVU4Hr1G090320.3 | TRIUR3_03031 | Seita.9G580200.1 | Sobic.001G544500.1 | Zm00001d027257_T008 |
| orthologID_008368 | PH01000753G0130 | LOC_Os03g48480.1 | Bradi1g12080.1 | HORVU4Hr1G004990.1 | TRIUR3_20208 | Seita.9G133200.1 | Sobic.001G131400.2 | Zm00001d013522_T001 |
| orthologID_008512 | PH01000243G0810 | LOC_Os03g03810.1 | Bradi1g76420.2 | HORVU4Hr1G082400.2 | TRIUR3_15796 | Seita.9G553900.1 | Sobic.001G518700.1 | Zm00001d027500_T001 |
| orthologID_007705 | PH01000687G0860 | LOC_Os03g14570.1 | Bradi1g68030.1 | HORVU4Hr1G062970.3 | TRIUR3_07111 | Seita.9G468600.1 | Sobic.001G436000.1 | Zm00001d047913_T001 |
| orthologID_008644 | PH01001042G0160 | LOC_Os03g55200.1 | Bradi1g07950.1 | HORVU5Hr1G096870.1 | TRIUR3_18181 | Seita.9G084200.1 | Sobic.001G082800.1 | Zm00001d013238_T001 |
| orthologID_008627 | PH01000395G0480 | LOC_Os03g15480.1 | Bradi1g67450.2 | HORVU4Hr1G061700.1 | TRIUR3_10372 | Seita.9G454400.1 | Sobic.001G429800.1 | Zm00001d028509_T001 |
| orthologID_009342 | PH01000230G0450 | LOC_Os10g05220.2 | Bradi3g17770.4 | HORVU5Hr1G105040.5 | TRIUR3_05763 | Seita.9G289100.1 | Sobic.001G268300.1 | Zm00001d030299_T001 |
| orthologID_007901 | PH01000278G0240 | LOC_Os03g40130.1 | Bradi1g15100.1 | HORVU5Hr1G061520.3 | TRIUR3_03873 | Seita.9G174100.1 | Sobic.001G173700.1 | Zm00001d033169_T001 |
| orthologID_008620 | PH01006466G0040 | LOC_Os03g18180.1 | Bradi1g65490.2 | HORVU4Hr1G056140.3 | TRIUR3_28516 | Seita.9G440200.1 | Sobic.001G405900.1 | Zm00001d047727_T002 |
| orthologID_008473 | PH01000154G1300 | LOC_Os03g16050.1 | Bradi1g67020.2 | HORVU4Hr1G060630.1 | TRIUR3_12456 | Seita.9G458800.1 | Sobic.001G425400.1 | Zm00001d028562_T001 |
| orthologID_007652 | PH01002188G0200 | LOC_Os03g45750.1 | Bradi1g13166.1 | HORVU4Hr1G002480.2 | TRIUR3_11841 | Seita.9G149300.1 | Sobic.001G147000.1 | Zm00001d013625_T001 |
| orthologID_009330 | PH01001683G0040 | LOC_Os10g42240.1 | Bradi3g34040.1 | HORVU0Hr1G000280.1 | TRIUR3_22509 | Seita.9G312300.1 | Sobic.001G289500.1 | Zm00001d030048_T001 |
| orthologID_007947 | PH01000010G1590 | LOC_Os03g04980.1 | Bradi1g75507.1 | HORVU4Hr1G078800.2 | TRIUR3_10173 | Seita.9G543200.1 | Sobic.001G507400.1 | Zm00001d048385_T001 |
| orthologID_009347 | PH01000012G1530 | LOC_Os10g36470.2 | Bradi3g30810.1 | HORVU1Hr1G047680.2 | TRIUR3_14953 | Seita.9G197100.1 | Sobic.001G199100.4 | Zm00001d013892_T010 |
| orthologID_007974 | PH01001926G0020 | LOC_Os07g02340.1 | Bradi1g59020.1 | HORVU2Hr1G063350.1 | TRIUR3_26862 | Seita.9G356800.1 | Sobic.001G329300.1 | Zm00001d029595_T005 |
| orthologID_007571 | PH01000308G0440 | LOC_Os03g14860.1 | Bradi1g67880.2 | HORVU4Hr1G062580.2 | TRIUR3_18096 | Seita.9G467000.1 | Sobic.001G434500.1 | Zm00001d047904_T001 |
| orthologID_009258 | PH01001857G0090 | LOC_Os10g42066.1 | Bradi3g34420.2 | HORVU1Hr1G050590.2 | TRIUR3_19362 | Seita.9G309900.1 | Sobic.001G286800.2 | Zm00001d030068_T001 |
| orthologID_007751 | PH01000953G0630 | LOC_Os03g42810.1 | Bradi1g14370.1 | HORVU0Hr1G007680.12 | TRIUR3_00034 | Seita.9G164900.1 | Sobic.001G162700.1 | Zm00001d033300_T001 |
| orthologID_007697 | PH01000606G0800 | LOC_Os03g53910.1 | Bradi1g08510.1 | HORVU5Hr1G095150.2 | TRIUR3_05187 | Seita.9G091000.1 | Sobic.001G088500.1 | Zm00001d034028_T003 |
| orthologID_007944 | PH01000339G0870 | LOC_Os03g13860.1 | Bradi1g68250.1 | HORVU4Hr1G063900.2 | TRIUR3_26308 | Seita.9G472900.1 | Sobic.001G440000.1 | Zm00001d047945_T002 |
| orthologID_009083 | PH01000362G0410 | LOC_Os10g37690.1 | Bradi3g31430.1 | HORVU1Hr1G048480.10 | TRIUR3_09719 | Seita.9G350300.1 | Sobic.001G322300.1 | Zm00001d029657_T001 |
| orthologID_007936 | PH01005158G0020 | LOC_Os03g52980.1 | Bradi1g09200.2 | HORVU5Hr1G094010.1 | TRIUR3_02691 | Seita.9G098500.1 | Sobic.001G096000.1 | Zm00001d033954_T001 |
| orthologID_008338 | PH01000040G0370 | LOC_Os03g62370.1 | Bradi1g02270.1 | HORVU5Hr1G119180.1 | TRIUR3_19354 | Seita.9G018500.1 | Sobic.001G019300.1 | Zm00001d012881_T001 |
| orthologID_008448 | PH01002172G0330 | LOC_Os03g53490.1 | Bradi1g08840.3 | HORVU5Hr1G094560.1 | TRIUR3_15503 | Seita.9G095200.1 | Sobic.001G092300.1 | Zm00001d033992_T009 |
| orthologID_008597 | PH01000371G0670 | LOC_Os03g63690.1 | Bradi1g01070.1 | HORVU5Hr1G123490.3 | TRIUR3_26623 | Seita.9G006500.1 | Sobic.001G007900.1 | Zm00001d012830_T002 |
| orthologID_009869 | PH01000562G0860 | LOC_Os10g27280.1 | Bradi3g26630.2 | HORVU1Hr1G021650.6 | TRIUR3_05853 | Seita.9G261300.1 | Sobic.001G250100.1 | Zm00001d032972_T001 |
| orthologID_007595 | PH01003422G0200 | LOC_Os03g56830.1 | Bradi1g06780.1 | HORVU5Hr1G100880.10 | TRIUR3_01932 | Seita.9G068400.1 | Sobic.001G069600.2 | Zm00001d034317_T001 |
| orthologID_009923 | PH01000335G0710 | LOC_Os10g33730.1 | Bradi3g28780.3 | HORVU1Hr1G041110.3 | TRIUR3_32067 | Seita.9G223000.1 | Sobic.001G219700.1 | Zm00001d014035_T001 |
| orthologID_007939 | PH01000010G1610 | LOC_Os03g04990.1 | Bradi1g75500.1 | HORVU4Hr1G078780.5 | TRIUR3_10172 | Seita.9G543100.1 | Sobic.001G507300.1 | Zm00001d027630_T002 |
| orthologID_007722 | PH01000804G0510 | LOC_Os03g61200.1 | Bradi1g03160.1 | HORVU5Hr1G115510.3 | TRIUR3_31534 | Seita.9G027400.1 | Sobic.001G028500.1 | Zm00001d034718_T001 |
| orthologID_007818 | PH01003165G0160 | LOC_Os03g30092.1 | Bradi1g59990.1 | HORVU3Hr1G013600.2 | TRIUR3_31448 | Seita.9G366500.1 | Sobic.001G337600.1 | Zm00001d039070_T001 |
| orthologID_007606 | PH01000395G0770 | LOC_Os03g15590.1 | Bradi1g67380.1 | HORVU4Hr1G061500.4 | TRIUR3_22975 | Seita.9G455100.1 | Sobic.001G429100.1 | Zm00001d028515_T001 |
| orthologID_008430 | PH01000430G0800 | LOC_Os03g20730.1 | Bradi1g63810.1 | HORVU4Hr1G051730.1 | TRIUR3_20136 | Seita.9G420300.1 | Sobic.001G387900.1 | Zm00001d028968_T001 |
| orthologID_007296 | PH01000287G0170 | LOC_Os03g01830.1 | Bradi4g13830.1 | HORVU4Hr1G090840.3 | TRIUR3_11647 | Seita.9G573100.1 | Sobic.001G537700.1 | Zm00001d048531_T001 |
| orthologID_008398 | PH01000122G0930 | LOC_Os03g59760.1 | Bradi1g04180.4 | HORVU5Hr1G111530.1 | TRIUR3_30605 | Seita.9G040300.1 | Sobic.001G040500.1 | Zm00001d013005_T004 |
| orthologID_009381 | PH01000321G0850 | LOC_Os10g38020.1 | Bradi3g31610.1 | HORVU1Hr1G048900.2 | TRIUR3_28628 | Seita.9G348000.1 | Sobic.001G320500.2 | Zm00001d029680_T001 |
| orthologID_008506 | PH01000370G0930 | LOC_Os03g03650.1 | Bradi1g76590.1 | HORVU4Hr1G082870.1 | TRIUR3_04853 | Seita.9G556100.1 | Sobic.001G521100.1 | Zm00001d027479_T002 |
| orthologID_008186 | PH01000081G2070 | LOC_Os03g05820.1 | Bradi1g74827.3 | HORVU4Hr1G072270.57 | TRIUR3_13257 | Seita.9G536800.1 | Sobic.001G500400.3 | Zm00001d027715_T001 |
| orthologID_007989 | PH01000287G1090 | LOC_Os03g01222.1 | Bradi1g78480.1 | HORVU4Hr1G088440.10 | TRIUR3_10282 | Seita.9G577800.1 | Sobic.001G542800.1 | Zm00001d048550_T001 |
| orthologID_007881 | PH01000101G0940 | LOC_Os03g12010.1 | Bradi1g69655.1 | HORVU4Hr1G067320.8 | TRIUR3_06357 | Seita.9G487700.1 | Sobic.001G453500.1 | Zm00001d028237_T002 |
| orthologID_008285 | PH01000532G0960 | LOC_Os03g14590.1 | Bradi1g68010.1 | HORVU4Hr1G062940.1 | TRIUR3_07109 | Seita.9G468400.1 | Sobic.001G435800.1 | Zm00001d028428_T001 |
| orthologID_007907 | PH01002421G0280 | LOC_Os03g38930.1 | Bradi1g15410.1 | HORVU4Hr1G037400.2 | TRIUR3_26709 | Seita.9G178800.1 | Sobic.001G179200.2 | Zm00001d033094_T001 |

| | | | | | | | | |
|---|---|---|---|---|---|---|---|---|
| orthologID_008156 | PH01000047G0210 | LOC_Os03g22870.1 | Bradi1g62340.1 | HORVU4Hr1G048800.3 | TRIUR3_11682 | Seita.9G403100.1 | Sobic.001G371200.1 | Zm00001d047474_T003 |
| orthologID_007764 | PH01000930G0290 | LOC_Os03g41530.1 | Bradi1g14690.1 | HORVU4Hr1G013610.2 | TRIUR3_06489 | Seita.9G169000.1 | Sobic.001G167800.1 | Zm00001d033337_T002 |
| orthologID_008322 | PH01000707G0940 | LOC_Os03g63554.1 | Bradi1g01330.1 | HORVU5Hr1G123220.1 | TRIUR3_07634 | Seita.9G007300.1 | Sobic.001G008700.1 | Zm00001d012832_T003 |
| orthologID_009303 | PH01002251G0100 | LOC_Os10g08840.1 | Bradi3g22600.3 | HORVU1Hr1G031310.10 | TRIUR3_07629 | Seita.9G280900.1 | Sobic.001G263200.1 | Zm00001d030385_T002 |
| orthologID_007720 | PH01000930G0020 | LOC_Os03g41600.1 | Bradi1g14677.7 | HORVU4Hr1G013630.8 | TRIUR3_06488 | Seita.9G168800.1 | Sobic.001G167600.1 | Zm00001d033335_T001 |
| orthologID_009696 | PH01000103G1690 | LOC_Os10g26660.1 | Bradi3g24960.1 | HORVU1Hr1G006950.3 | TRIUR3_04151 | Seita.9G259500.1 | Sobic.001G252500.1 | Zm00001d014196_T001 |
| orthologID_007761 | PH01000113G1310 | LOC_Os03g07850.1 | Bradi1g72967.1 | HORVU4Hr1G075740.2 | TRIUR3_33341 | Seita.9G521300.1 | Sobic.001G486500.3 | Zm00001d027872_T001 |
| orthologID_007866 | PH01001135G0640 | LOC_Os03g51700.1 | Bradi1g10040.1 | HORVU4Hr1G009760.1 | TRIUR3_34302 | Seita.9G109400.1 | Sobic.001G106100.3 | Zm00001d033860_T001 |
| orthologID_006734 | PH01002121G0090 | LOC_Os05g41930.2 | Bradi2g20890.4 | HORVU1Hr1G066630.1 | TRIUR3_22557 | Seita.9G008200.1 | Sobic.001G009800.1 | Zm00001d034867_T001 |
| orthologID_009796 | PH01000935G0770 | LOC_Os10g31930.1 | Bradi3g27940.1 | HORVU4Hr1G005960.11 | TRIUR3_19114 | Seita.9G233200.1 | Sobic.001G230300.1 | Zm00001d032805_T001 |
| orthologID_007776 | PH01000750G0280 | LOC_Os03g52670.1 | Bradi1g09420.1 | HORVU4Hr1G011100.2 | TRIUR3_06692 | Seita.9G101000.1 | Sobic.001G098700.1 | Zm00001d013325_T001 |
| orthologID_009769 | PH01002582G0150 | LOC_Os10g34580.1 | Bradi3g29390.1 | HORVU1Hr1G043110.7 | TRIUR3_25677 | Seita.9G213700.1 | Sobic.001G212900.1 | Zm00001d013976_T002 |
| orthologID_007879 | PH01001135G0080 | LOC_Os03g51610.1 | Bradi1g10140.1 | HORVU4Hr1G009540.9 | TRIUR3_05805 | Seita.9G110300.1 | Sobic.001G106900.3 | Zm00001d033853_T002 |
| orthologID_008490 | PH01004875G0030 | LOC_Os03g48030.1 | Bradi1g12280.1 | HORVU4Hr1G004170.2 | TRIUR3_04908 | Seita.9G137500.1 | Sobic.001G134800.1 | Zm00001d013536_T002 |
| orthologID_009758 | PH01000560G0020 | LOC_Os10g32540.1 | Bradi3g28060.1 | HORVU1Hr1G038620.27 | TRIUR3_31505 | Seita.9G231500.1 | Sobic.001G228300.1 | Zm00001d032790_T002 |
| orthologID_008405 | PH01001710G0260 | LOC_Os03g22780.1 | Bradi1g62410.3 | HORVU4Hr1G048620.6 | TRIUR3_02709 | Seita.9G404400.1 | Sobic.001G372600.1 | Zm00001d029150_T001 |
| orthologID_008675 | PH01000017G0720 | LOC_Os03g02730.1 | Bradi1g77270.1 | HORVU4Hr1G085730.2 | TRIUR3_14182 | Seita.9G564700.1 | Sobic.001G529900.2 | Zm00001d027385_T003 |
| orthologID_009714 | PH01000321G0230 | LOC_Os10g37670.1 | Bradi3g31420.1 | HORVU1Hr1G048450.4 | TRIUR3_31227 | Seita.9G350400.1 | Sobic.001G322400.1 | Zm00001d029656_T003 |
| orthologID_009926 | PH01002582G0230 | LOC_Os10g34470.1 | Bradi3g29300.2 | HORVU1Hr1G042820.3 | TRIUR3_34777 | Seita.9G214600.1 | Sobic.001G213400.1 | Zm00001d032652_T001 |
| orthologID_007637 | PH01000430G0320 | LOC_Os03g20460.1 | Bradi1g63967.1 | HORVU4Hr1G052090.1 | TRIUR3_12286 | Seita.9G422300.1 | Sobic.001G389500.1 | Zm00001d028954_T001 |
| orthologID_007834 | PH01003162G0270 | LOC_Os03g32160.1 | Bradi1g03470.1 | HORVU5Hr1G114650.1 | TRIUR3_12153 | Seita.9G030200.1 | Sobic.001G031600.1 | Zm00001d012934_T001 |
| orthologID_008187 | PH01001789G0250 | LOC_Os03g48970.1 | Bradi1g11800.5 | HORVU4Hr1G005670.3 | TRIUR3_09920 | Seita.9G129400.1 | Sobic.001G127800.1 | Zm00001d033602_T002 |
| orthologID_008167 | PH01001012G0570 | LOC_Os03g12660.1 | Bradi1g69040.1 | HORVU4Hr1G065440.16 | TRIUR3_29045 | Seita.9G480100.1 | Sobic.001G445900.1 | Zm00001d028325_T005 |
| orthologID_007797 | PH01000040G1660 | LOC_Os03g61470.1 | Bradi1g03090.1 | HORVU5Hr1G115880.1 | TRIUR3_04648 | Seita.4G003800.1 | Sobic.001G024800.1 | Zm00001d012900_T001 |
| orthologID_008164 | PH01002606G0020 | LOC_Os03g53300.2 | Bradi1g08907.1 | HORVU5Hr1G094400.6 | TRIUR3_21381 | Seita.9G096100.1 | Sobic.001G093500.3 | Zm00001d033985_T001 |
| orthologID_008115 | PH01001260G0050 | LOC_Os03g56790.1 | Bradi1g06821.1 | HORVU5Hr1G100770.1 | TRIUR3_01926 | Seita.9G068800.1 | Sobic.001G070000.2 | Zm00001d013141_T001 |
| orthologID_008469 | PH01001174G0490 | LOC_Os06g06740.1 | Bradi1g48525.2 | HORVU7Hr1G025720.1 | TRIUR3_12421 | Seita.9G054400.1 | Sobic.001G056000.1 | Zm00001d034467_T001 |
| orthologID_008146 | PH01001382G0310 | LOC_Os03g19010.1 | Bradi1g64750.1 | HORVU4Hr1G054560.10 | TRIUR3_19751 | Seita.9G433200.1 | Sobic.001G399100.1 | Zm00001d028834_T001 |
| orthologID_009187 | PH01000362G0600 | LOC_Os07g20150.1 | Bradi3g31397.1 | HORVU1Hr1G048400.1 | TRIUR3_34511 | Seita.9G350900.1 | Sobic.001G322600.1 | Zm00001d029653_T001 |
| orthologID_009859 | PH01000873G0070 | LOC_Os08g14440.1 | Bradi3g18820.8 | HORVU7Hr1G069710.6 | TRIUR3_34685 | Seita.6G090900.1 | Sobic.001G252000.1 | Zm00001d014192_T001 |
| orthologID_007592 | PH01001111G0270 | LOC_Os03g55920.1 | Bradi1g07380.1 | HORVU5Hr1G098320.33 | TRIUR3_09992 | Seita.9G076400.1 | Sobic.001G076600.2 | Zm00001d013176_T002 |
| orthologID_008637 | PH01003832G0070 | LOC_Os03g53530.1 | Bradi1g08810.1 | HORVU5Hr1G094580.1 | TRIUR3_25896 | Seita.9G094800.1 | Sobic.001G091900.1 | Zm00001d013283_T001 |
| orthologID_008088 | PH01000716G0710 | LOC_Os03g55110.1 | Bradi1g08040.1 | HORVU5Hr1G096470.2 | TRIUR3_13990 | Seita.9G085300.1 | Sobic.001G083600.1 | Zm00001d013242_T001 |
| orthologID_009835 | PH01002782G0190 | LOC_Os10g30180.1 | Bradi3g27360.1 | HORVU1Hr1G035950.1 | TRIUR3_01036 | Seita.9G243400.1 | Sobic.001G239200.1 | Zm00001d014146_T001 |
| orthologID_008018 | PH01001215G0280 | LOC_Os03g02410.1 | Bradi1g77547.2 | HORVU4Hr1G086640.4 | TRIUR3_15497 | Seita.9G568900.1 | Sobic.001G533100.1 | Zm00001d027347_T002 |
| orthologID_007734 | PH01002899G0120 | LOC_Os03g55730.1 | Bradi1g07520.3 | HORVU5Hr1G097860.2 | TRIUR3_15042 | Seita.9G078400.1 | Sobic.001G077900.1 | Zm00001d034180_T002 |
| orthologID_008443 | PH01002606G0320 | LOC_Os03g53770.1 | Bradi1g08597.1 | HORVU5Hr1G095000.1 | TRIUR3_13694 | Seita.9G092000.1 | Sobic.001G089500.1 | Zm00001d013271_T003 |
| orthologID_009386 | PH01001233G0040 | LOC_Os10g41940.1 | Bradi3g33890.1 | HORVU1Hr1G054450.2 | TRIUR3_08204 | Seita.9G315400.1 | Sobic.001G291900.1 | Zm00001d030010_T001 |
| orthologID_008489 | PH01002188G0360 | LOC_Os03g45720.1 | Bradi1g13190.1 | HORVU4Hr1G002510.1 | TRIUR3_12275 | Seita.9G149600.1 | Sobic.001G147300.2 | Zm00001d033468_T001 |
| orthologID_008148 | PH01000848G0700 | LOC_Os03g39640.2 | Bradi1g15270.1 | HORVU5Hr1G111010.1 | TRIUR3_17765 | Seita.9G176900.1 | Sobic.001G176700.1 | Zm00001d033155_T001 |
| orthologID_008461 | PH01000294G0480 | LOC_Os03g25130.1 | Bradi1g61610.5 | HORVU3Hr1G012090.11 | TRIUR3_11050 | Seita.9G393600.1 | Sobic.001G361000.1 | Zm00001d047427_T001 |
| orthologID_007565 | PH01000857G0280 | LOC_Os03g17990.1 | Bradi1g65620.1 | HORVU4Hr1G056600.1 | TRIUR3_29907 | Seita.9G441900.1 | Sobic.001G408100.2 | Zm00001d047745_T001 |
| orthologID_006442 | PH01000429G0790 | LOC_Os05g45030.1 | Bradi2g19260.1 | HORVU1Hr1G081980.1 | TRIUR3_11043 | Seita.2G410100.1 | Sobic.001G094200.1 | Zm00001d013295_T001 |
| orthologID_007843 | PH01004662G0080 | LOC_Os03g36650.1 | Bradi1g15297.1 | HORVU4Hr1G085990.1 | TRIUR3_27821 | Seita.9G186000.1 | Sobic.001G188800.1 | Zm00001d032401_T001 |
| orthologID_009487 | PH01000301G1030 | LOC_Os10g30640.1 | Bradi3g27660.1 | HORVU1Hr1G037340.1 | TRIUR3_22281 | Seita.9G239000.1 | Sobic.001G235700.1 | Zm00001d014122_T002 |
| orthologID_006593 | PH01197141G0010 | LOC_Os03g32220.1 | Bradi2g49250.1 | HORVU2Hr1G014870.1 | TRIUR3_09270 | Seita.9G357600.1 | Sobic.001G330400.1 | Zm00001d029584_T001 |
| orthologID_010109 | PH01001109G0380 | LOC_Os03g16950.1 | Bradi1g66450.1 | HORVU4Hr1G059150.6 | TRIUR3_10430 | Seita.9G451300.1 | Sobic.001G418200.1 | Zm00001d028651_T001 |
| orthologID_007576 | PH01000426G0590 | LOC_Os03g49480.1 | Bradi1g11580.1 | HORVU4Hr1G006020.1 | TRIUR3_01048 | Seita.9G126700.1 | Sobic.001G124500.1 | Zm00001d033637_T001 |
| orthologID_009312 | PH01000516G0350 | LOC_Os10g21020.1 | Bradi3g50670.4 | HORVU5Hr1G022110.3 | TRIUR3_09276 | Seita.9G273900.1 | Sobic.001G350000.1 | Zm00001d029396_T001 |
| orthologID_008661 | PH01000157G0540 | LOC_Os03g05260.1 | Bradi1g75360.1 | HORVU4Hr1G079190.1 | TRIUR3_07622 | Seita.9G541600.1 | Sobic.001G505500.1 | Zm00001d027648_T001 |

| | | | | | | | | |
|---|---|---|---|---|---|---|---|---|
| orthologID_008230 | PH01000040G0210 | LOC_Os03g62580.1 | Bradi1g02117.1 | HORVU5Hr1G120090.1 | TRIUR3_19735 | Seita.9G017100.1 | Sobic.001G017800.1 | Zm00001d034796_T005 |
| orthologID_007769 | PH01000371G0120 | LOC_Os03g63400.1 | Bradi1g01600.1 | HORVU5Hr1G122170.4 | TRIUR3_13442 | Seita.9G009500.1 | Sobic.001G011200.1 | Zm00001d034855_T001 |
| orthologID_008166 | PH01000985G0600 | LOC_Os03g61750.1 | Bradi1g02780.1 | HORVU5Hr1G116970.9 | TRIUR3_22091 | Seita.9G022900.1 | Sobic.001G026400.1 | Zm00001d012915_T001 |
| orthologID_007869 | PH01000786G0430 | LOC_Os03g15390.1 | Bradi1g67540.1 | HORVU4Hr1G061880.2 | TRIUR3_11406 | Seita.9G462800.1 | Sobic.001G430800.1 | Zm00001d028493_T002 |
| orthologID_008001 | PH01000365G0320 | LOC_Os03g62610.1 | Bradi1g02080.1 | HORVU5Hr1G120240.2 | TRIUR3_02199 | Seita.9G016800.1 | Sobic.001G017300.1 | Zm00001d034804_T002 |
| orthologID_007613 | PH01001125G0090 | LOC_Os03g52010.1 | Bradi1g09840.1 | HORVU4Hr1G010180.2 | TRIUR3_20748 | Seita.9G107000.1 | Sobic.001G103800.2 | Zm00001d033882_T001 |
| orthologID_008685 | PH01000122G1060 | LOC_Os03g60100.1 | Bradi1g04130.1 | HORVU5Hr1G111640.1 | TRIUR3_12289 | Seita.9G039000.1 | Sobic.001G040000.2 | Zm00001d012998_T001 |
| orthologID_008030 | PH01000370G0290 | LOC_Os03g03400.1 | Bradi1g76850.3 | HORVU4Hr1G083340.2 | TRIUR3_12542 | Seita.9G559000.1 | Sobic.001G524300.1 | Zm00001d027445_T002 |
| orthologID_008395 | PH01003859G0110 | LOC_Os03g32314.1 | Bradi1g15840.1 | HORVU6Hr1G081000.3 | TRIUR3_25644 | Seita.9G357200.1 | Sobic.001G329800.1 | Zm00001d029594_T001 |
| orthologID_006922 | PH01000816G0230 | LOC_Os10g39520.1 | Bradi3g32230.1 | HORVU1Hr1G052110.8 | TRIUR3_13822 | Seita.9G340900.1 | Sobic.001G312600.1 | Zm00001d029768_T001 |
| orthologID_007651 | PH01000370G1160 | LOC_Os03g03710.1 | Bradi1g76480.1 | HORVU4Hr1G082690.4 | TRIUR3_21490 | Seita.9G554800.1 | Sobic.001G519900.1 | Zm00001d027487_T017 |
| orthologID_008283 | PH01002625G0100 | LOC_Os03g20100.1 | Bradi1g64130.2 | HORVU4Hr1G052470.1 | TRIUR3_32878 | Seita.9G425000.1 | Sobic.001G391400.1 | Zm00001d047581_T001 |
| orthologID_008151 | PH01000456G0930 | LOC_Os03g24890.1 | Bradi1g61730.1 | HORVU4Hr1G045510.3 | TRIUR3_14282 | Seita.9G395300.1 | Sobic.001G362300.1 | Zm00001d029256_T002 |
| orthologID_009184 | PH01000560G0780 | LOC_Os10g32860.2 | Bradi3g28257.1 | HORVU1Hr1G039010.6 | TRIUR3_13038 | Seita.9G229100.1 | Sobic.001G226200.1 | Zm00001d014080_T001 |
| orthologID_009109 | PH01000520G0230 | LOC_Os10g35644.1 | Bradi3g30027.2 | HORVU1Hr1G045080.14 | TRIUR3_32579 | Seita.9G204500.1 | Sobic.001G204501.1 | Zm00001d032552_T002 |
| orthologID_008135 | PH01000154G0740 | LOC_Os03g15810.1 | Bradi1g67210.1 | HORVU4Hr1G061060.1 | TRIUR3_20438 | Seita.9G456800.1 | Sobic.001G427400.2 | Zm00001d028538_T003 |
| orthologID_008077 | PH01001477G0270 | LOC_Os03g55280.1 | Bradi1g07910.1 | HORVU5Hr1G097020.5 | TRIUR3_08557 | Seita.9G082600.1 | Sobic.001G081900.1 | Zm00001d013223_T002 |
| orthologID_006569 | PH01000992G0430 | LOC_Os05g24930.1 | Bradi2g31210.2 | HORVU7Hr1G034050.17 | TRIUR3_04300 | Seita.3G289900.1 | Sobic.001G451200.1 | Zm00001d028263_T001 |
| orthologID_009672 | PH01001197G0220 | LOC_Os10g27480.1 | Bradi3g26710.1 | HORVU1Hr1G035080.4 | TRIUR3_34703 | Seita.9G262000.1 | Sobic.001G249100.1 | Zm00001d032962_T003 |
| orthologID_009207 | PH01001078G0280 | LOC_Os10g25230.1 | Bradi3g23190.1 | HORVU7Hr1G041300.1 | TRIUR3_30081 | Seita.9G252100.1 | Sobic.001G259600.1 | Zm00001d033049_T001 |
| orthologID_007894 | PH01000634G0360 | LOC_Os03g51380.1 | Bradi1g10300.1 | HORVU4Hr1G009070.2 | TRIUR3_19712 | Seita.9G112000.1 | Sobic.001G108900.1 | Zm00001d013383_T001 |
| orthologID_008300 | PH01000535G0670 | LOC_Os03g63930.1 | Bradi1g01000.1 | HORVU0Hr1G038610.1 | TRIUR3_04651 | Seita.9G004800.1 | Sobic.001G005700.1 | Zm00001d012818_T001 |
| orthologID_007627 | PH01000953G0180 | LOC_Os03g42710.1 | Bradi1g14460.2 | HORVU4Hr1G012000.11 | TRIUR3_16120 | Seita.9G165400.1 | Sobic.001G163600.1 | Zm00001d033304_T001 |
| orthologID_008020 | PH01004012G0140 | ChrSy.fgenesh.mRNA.86 | Bradi1g78660.3 | HORVU4Hr1G090300.4 | TRIUR3_03034 | Seita.7G331300.1 | Sobic.001G029100.2 | Zm00001d034715_T001 |
| orthologID_008118 | PH01000650G0430 | LOC_Os03g49730.1 | Bradi1g11010.1 | HORVU4Hr1G006720.4 | TRIUR3_08985 | Seita.9G124100.1 | Sobic.001G122200.1 | Zm00001d013471_T001 |
| orthologID_008028 | PH01000287G1140 | LOC_Os03g01200.1 | Bradi1g78440.1 | HORVU4Hr1G088540.8 | TRIUR3_19709 | Seita.9G578200.1 | Sobic.001G543100.2 | Zm00001d048552_T005 |
| orthologID_009834 | PH01001382G0410 | LOC_Os07g22840.1 | Bradi3g57330.1 | HORVU5Hr1G112840.2 | TRIUR3_19790 | Seita.9G432900.1 | Sobic.001G398600.2 | Zm00001d028837_T001 |
| orthologID_009663 | PH01000335G0770 | LOC_Os10g33660.1 | Bradi3g28730.2 | HORVU1Hr1G040930.1 | TRIUR3_10604 | Seita.9G223700.1 | Sobic.001G220400.1 | Zm00001d032710_T003 |
| orthologID_008505 | PH01001135G0160 | LOC_Os03g51650.1 | Bradi1g10110.1 | HORVU4Hr1G009560.2 | TRIUR3_23950 | Seita.9G109900.1 | Sobic.001G106600.1 | Zm00001d033855_T002 |
| orthologID_008435 | PH01001876G0340 | LOC_Os03g58830.1 | Bradi1g05037.1 | HORVU5Hr1G107520.2 | TRIUR3_14778 | Seita.9G049500.1 | Sobic.001G051300.1 | Zm00001d034503_T001 |
| orthologID_007910 | PH01000535G0510 | LOC_Os03g63970.1 | Bradi1g00950.1 | HORVU5Hr1G124120.4 | TRIUR3_34129 | Seita.9G004400.1 | Sobic.001G005300.1 | Zm00001d034898_T001 |
| orthologID_008463 | PH01002278G0120 | LOC_Os03g17610.1 | Bradi1g65870.1 | HORVU4Hr1G057310.1 | TRIUR3_24097 | Seita.9G444800.1 | Sobic.001G410800.1 | Zm00001d028705_T001 |
| orthologID_008317 | PH01004691G0100 | LOC_Os03g46190.1 | Bradi1g12980.3 | HORVU4Hr1G002720.2 | TRIUR3_24510 | Seita.9G147700.1 | Sobic.001G144600.1 | Zm00001d013597_T003 |
| orthologID_008144 | PH01000107G1540 | LOC_Os03g03390.1 | Bradi1g76860.1 | HORVU4Hr1G083360.2 | TRIUR3_30857 | Seita.9G559100.1 | Sobic.001G524400.1 | Zm00001d027444_T003 |
| orthologID_008613 | PH01001219G0530 | LOC_Os03g22230.1 | Bradi1g62880.1 | HORVU7Hr1G032740.1 | TRIUR3_00966 | Seita.9G409900.1 | Sobic.001G377500.1 | Zm00001d029099_T001 |
| orthologID_007861 | PH01000039G0390 | LOC_Os03g43850.1 | Bradi1g13960.1 | HORVU5Hr1G093270.18 | TRIUR3_24505 | Seita.9G159800.1 | Sobic.001G158400.1 | Zm00001d033259_T002 |
| orthologID_008393 | PH01000601G0130 | LOC_Os03g32050.1 | Bradi1g57247.1 | HORVU4Hr1G052060.1 | TRIUR3_28374 | Seita.9G355500.1 | Sobic.001G328100.1 | Zm00001d029604_T001 |
| orthologID_008328 | PH01000535G0700 | LOC_Os03g63910.1 | Bradi1g01021.1 | HORVU5Hr1G124070.4 | TRIUR3_04654 | Seita.9G005100.1 | Sobic.001G006000.1 | Zm00001d034892_T001 |
| orthologID_009502 | PH01000301G0700 | LOC_Os10g30550.1 | Bradi3g27600.2 | HORVU1Hr1G036990.9 | TRIUR3_25582 | Seita.9G239800.1 | Sobic.001G236800.1 | Zm00001d032867_T002 |
| orthologID_008511 | PH01001463G0120 | LOC_Os03g51840.1 | Bradi1g09990.1 | HORVU4Hr1G009900.1 | TRIUR3_01288 | Seita.9G108600.1 | Sobic.001G105400.4 | Zm00001d033869_T001 |
| orthologID_009519 | PH01001054G0170 | LOC_Os10g34350.1 | Bradi3g29210.5 | HORVU1Hr1G042490.5 | TRIUR3_09391 | Seita.9G216200.1 | Sobic.001G215300.2 | Zm00001d013992_T001 |
| orthologID_008576 | PH01001096G0090 | LOC_Os03g45270.1 | Bradi1g13340.1 | HORVU4Hr1G002230.1 | TRIUR3_00736 | Seita.9G151900.1 | Sobic.001G149600.3 | Zm00001d033484_T002 |
| orthologID_008173 | PH01001042G0700 | LOC_Os03g55380.1 | Bradi1g07820.2 | HORVU5Hr1G097220.2 | TRIUR3_19314 | Seita.9G081700.1 | Sobic.001G080800.1 | Zm00001d034124_T001 |
| orthologID_007585 | PH01001292G0350 | LOC_Os03g21060.1 | Bradi1g63600.1 | HORVU4Hr1G051290.1 | TRIUR3_29857 | Seita.9G417600.1 | Sobic.001G385800.1 | Zm00001d028999_T001 |
| orthologID_008312 | PH01001328G0610 | LOC_Os03g07180.1 | Bradi1g73840.1 | HORVU4Hr1G074750.1 | TRIUR3_02049 | Seita.9G526800.1 | Sobic.001G492000.1 | Zm00001d027832_T001 |
| orthologID_007708 | PH01000101G0740 | LOC_Os03g11910.1 | Bradi1g69700.1 | HORVU4Hr1G067430.1 | TRIUR3_12311 | Seita.9G488400.1 | Sobic.001G454000.1 | Zm00001d048073_T002 |
| orthologID_007858 | PH01001042G0520 | LOC_Os03g55340.1 | Bradi1g07850.1 | HORVU5Hr1G097140.8 | TRIUR3_17663 | Seita.9G082000.1 | Sobic.001G081300.1 | Zm00001d034111_T001 |
| orthologID_009912 | PH01001155G0090 | LOC_Os10g40620.1 | Bradi3g33070.1 | HORVU1Hr1G054060.6 | TRIUR3_33724 | Seita.9G327600.1 | Sobic.001G302300.1 | Zm00001d029921_T001 |
| orthologID_008522 | PH01002636G0210 | LOC_Os03g26220.1 | Bradi1g61100.1 | HORVU4Hr1G075970.2 | TRIUR3_05741 | Seita.9G384300.1 | Sobic.001G355500.1 | Zm00001d047184_T001 |
| orthologID_009334 | PH01001428G0330 | LOC_Os10g33700.1 | Bradi3g28747.1 | HORVU1Hr1G041020.17 | TRIUR3_34910 | Seita.9G223400.1 | Sobic.001G220000.2 | Zm00001d032707_T001 |

| | | | | | | | | |
|---|---|---|---|---|---|---|---|---|
| orthologID_009495 | PH01001411G0130 | LOC_Os10g37490.1 | Bradi3g31120.1 | HORVU1Hr1G048250.1 | TRIUR3_22102 | Seita.9G190300.1 | Sobic.001G192900.1 | Zm00001d032439_T002 |
| orthologID_008475 | PH01000101G0140 | LOC_Os03g11660.1 | Bradi1g69840.1 | HORVU4Hr1G067610.1 | TRIUR3_12881 | Seita.9G490000.1 | Sobic.001G455600.1 | Zm00001d028220_T001 |
| orthologID_008250 | PH01000416G0920 | LOC_Os03g22140.1 | Bradi1g62950.1 | HORVU4Hr1G046380.1 | TRIUR3_34576 | Seita.9G410700.1 | Sobic.001G378200.1 | Zm00001d029093_T001 |
| orthologID_008514 | PH01000353G0680 | LOC_Os03g13600.2 | Bradi1g68570.2 | HORVU4Hr1G064150.1 | TRIUR3_15555 | Seita.9G474600.1 | Sobic.001G441500.2 | Zm00001d028371_T001 |
| orthologID_008677 | PH01001537G0080 | LOC_Os03g11260.1 | Bradi1g70360.1 | HORVU4Hr1G068950.1 | TRIUR3_07104 | Seita.9G494800.1 | Sobic.001G460600.1 | Zm00001d048112_T001 |
| orthologID_008550 | PH01000804G0440 | LOC_Os03g61140.1 | Bradi1g03290.1 | HORVU5Hr1G115530.2 | TRIUR3_34397 | Seita.9G027600.1 | Sobic.001G028700.1 | Zm00001d034717_T002 |
| orthologID_008547 | PH01000339G0970 | LOC_Os03g13800.1 | Bradi1g68505.2 | HORVU4Hr1G064020.1 | TRIUR3_11631 | Seita.9G473700.1 | Sobic.001G513500.1 | Zm00001d028384_T006 |
| orthologID_007811 | PH01001259G0240 | LOC_Os03g01110.1 | Bradi1g78610.2 | HORVU4Hr1G090170.17 | TRIUR3_03142 | Seita.9G579800.1 | Sobic.001G544300.1 | Zm00001d027259_T010 |
| orthologID_007862 | PH01000188G0340 | LOC_Os03g07380.1 | Bradi1g73670.2 | HORVU4Hr1G075110.1 | TRIUR3_08657 | Seita.9G524800.1 | Sobic.001G489600.1 | Zm00001d048317_T001 |
| orthologID_009877 | PH01000904G0660 | LOC_Os10g36610.1 | Bradi3g30730.2 | HORVU1Hr1G047570.2 | TRIUR3_26442 | Seita.9G195800.1 | Sobic.001G198000.1 | Zm00001d032481_T001 |
| orthologID_008411 | PH01000122G0090 | LOC_Os03g59540.1 | Bradi1g04550.4 | HORVU5Hr1G110170.2 | TRIUR3_07465 | Seita.9G042700.1 | Sobic.001G043601.1 | Zm00001d013025_T001 |
| orthologID_008262 | PH01001382G0320 | LOC_Os03g19020.1 | Bradi1g64740.1 | HORVU4Hr1G054580.4 | TRIUR3_19750 | Seita.9G433100.1 | Sobic.001G398950.1 | Zm00001d047676_T002 |
| orthologID_008113 | PH01000007G3610 | LOC_Os03g12230.1 | Bradi1g69571.2 | HORVU4Hr1G067020.1 | TRIUR3_33351 | Seita.9G486300.1 | Sobic.001G452100.2 | Zm00001d028252_T002 |
| orthologID_009143 | PH01000170G0510 | LOC_Os10g39810.1 | Bradi3g32510.1 | HORVU1Hr1G052890.4 | TRIUR3_17653 | Seita.9G337500.1 | Sobic.001G309200.1 | Zm00001d029810_T001 |
| orthologID_009631 | PH01000170G0490 | LOC_Os10g39800.1 | Bradi3g32497.1 | HORVU1Hr1G052780.11 | TRIUR3_27920 | Seita.9G337600.1 | Sobic.001G309300.1 | Zm00001d029809_T002 |
| orthologID_009169 | PH01000171G0750 | LOC_Os10g41380.1 | Bradi3g33530.1 | HORVU1Hr1G055530.2 | TRIUR3_10679 | Seita.9G319500.1 | Sobic.001G295500.1 | Zm00001d000035_T001 |
| orthologID_008521 | PH01000601G0550 | LOC_Os11g23170.1 | Bradi1g15960.1 | HORVU4Hr1G078670.1 | TRIUR3_10769 | Seita.9G189100.1 | Sobic.001G191900.1 | Zm00001d032430_T002 |
| orthologID_008612 | PH01000426G0190 | LOC_Os03g49230.1 | Bradi1g11700.1 | HORVU4Hr1G005940.3 | TRIUR3_24976 | Seita.9G128000.1 | Sobic.001G126200.1 | Zm00001d033620_T001 |
| orthologID_009523 | PH01003963G0080 | LOC_Os10g10180.1 | Bradi3g22740.1 | HORVU6Hr1G073180.5 | TRIUR3_15391 | Seita.9G283600.1 | Sobic.001G448400.1 | Zm00001d014284_T002 |
| orthologID_008619 | PH01003619G0120 | LOC_Os03g44520.1 | Bradi1g13700.1 | HORVU5Hr1G092790.1 | TRIUR3_20331 | Seita.9G155800.1 | Sobic.001G154600.1 | Zm00001d033216_T001 |
| orthologID_008132 | PH01000535G0680 | LOC_Os03g63920.1 | Bradi1g01007.1 | HORVU0Hr1G038620.1 | TRIUR3_04652 | Seita.9G004900.1 | Sobic.001G005800.1 | Zm00001d034894_T001 |
| orthologID_009290 | PH01000012G0190 | LOC_Os10g35770.1 | Bradi3g30110.1 | HORVU1Hr1G045220.4 | TRIUR3_02519 | Seita.9G202700.1 | Sobic.001G203200.1 | Zm00001d032536_T001 |
| orthologID_008238 | PH01003849G0120 | LOC_Os03g64350.1 | Bradi1g00530.2 | HORVU5Hr1G124520.3 | TRIUR3_34369 | Seita.9G001700.1 | Sobic.001G001300.1 | Zm00001d034932_T001 |
| orthologID_007980 | PH01000188G0390 | LOC_Os03g07420.1 | Bradi1g73220.6 | HORVU4Hr1G075140.1 | TRIUR3_16126 | Seita.9G524600.1 | Sobic.001G489400.2 | Zm00001d048315_T001 |
| orthologID_007635 | PH01002278G0350 | LOC_Os03g17550.1 | Bradi1g65930.1 | HORVU4Hr1G057520.4 | TRIUR3_12158 | Seita.9G445400.1 | Sobic.001G411600.1 | Zm00001d028701_T001 |
| orthologID_009144 | PH01000012G1620 | LOC_Os10g36530.1 | Bradi3g30757.2 | HORVU1Hr1G047640.18 | TRIUR3_22629 | Seita.9G196100.1 | Sobic.001G198400.2 | Zm00001d013887_T002 |
| orthologID_007808 | PH01000106G1230 | LOC_Os03g11230.1 | Bradi1g70420.1 | HORVU4Hr1G069050.1 | TRIUR3_26055 | Seita.9G495400.1 | Sobic.001G460900.2 | Zm00001d048116_T001 |
| orthologID_008571 | PH01000184G0460 | LOC_Os03g39830.1 | Bradi1g15170.1 | HORVU5Hr1G111330.4 | TRIUR3_04346 | Seita.9G176400.1 | Sobic.001G175700.1 | Zm00001d033150_T001 |
| orthologID_008591 | PH01000113G1750 | LOC_Os03g07980.1 | Bradi1g72875.1 | HORVU4Hr1G076170.1 | TRIUR3_31688 | Seita.9G520300.1 | Sobic.001G484900.1 | Zm00001d027881_T001 |
| orthologID_009352 | PH01000912G0160 | LOC_Os02g41560.1 | Bradi3g49211.1 | HORVU6Hr1G058730.1 | TRIUR3_13731 | Seita.1G239500.1 | Sobic.001G003900.1 | Zm00001d012812_T004 |
| orthologID_008298 | PH01000308G1030 | LOC_Os03g15120.1 | Bradi1g67700.1 | HORVU4Hr1G062310.1 | TRIUR3_21464 | Seita.9G464600.1 | Sobic.001G432300.1 | Zm00001d037287_T001 |
| orthologID_008084 | PH01000113G1270 | LOC_Os03g07820.1 | Bradi1g73000.1 | HORVU4Hr1G075810.7 | TRIUR3_25509 | Seita.9G521600.1 | Sobic.001G486900.1 | Zm00001d048298_T003 |
| orthologID_008291 | PH01000339G0720 | LOC_Os03g14000.1 | Bradi1g68300.1 | HORVU4Hr1G063810.1 | TRIUR3_29906 | Seita.9G472300.1 | Sobic.001G439500.1 | Zm00001d028397_T001 |
| orthologID_009154 | PH01003821G0130 | LOC_Os08g31760.1 | Bradi3g35640.1 | HORVU0Hr1G017530.1 | TRIUR3_16121 | Seita.6G142700.1 | Sobic.001G533000.1 | Zm00001d027348_T001 |
| orthologID_007813 | PH01000308G0850 | LOC_Os03g15040.1 | Bradi1g67740.1 | HORVU4Hr1G062460.32 | TRIUR3_23242 | Seita.9G465600.1 | Sobic.001G433000.1 | Zm00001d028472_T001 |
| orthologID_008455 | PH01000143G1560 | LOC_Os03g02640.1 | Bradi1g77350.3 | HORVU4Hr1G085930.8 | TRIUR3_08174 | Seita.9G565900.1 | Sobic.001G531200.1 | Zm00001d027369_T002 |
| orthologID_008198 | PH01000113G1210 | LOC_Os03g07800.1 | Bradi1g73030.1 | HORVU4Hr1G075700.5 | TRIUR3_15334 | Seita.9G521800.1 | Sobic.001G487100.1 | Zm00001d027869_T002 |
| orthologID_007984 | PH01000468G0840 | LOC_Os03g20570.1 | Bradi1g63890.3 | HORVU4Hr1G051990.7 | TRIUR3_06229 | Seita.9G421500.1 | Sobic.001G388800.1 | Zm00001d033635_T001 |
| orthologID_008728 | PH01000376G0430 | LOC_Os03g56410.1 | Bradi1g07080.1 | HORVU5Hr1G099070.1 | TRIUR3_00987 | Seita.9G072100.1 | Sobic.001G072800.2 | Zm00001d034254_T002 |
| orthologID_007636 | PH01002858G0290 | LOC_Os03g31570.1 | Bradi1g15910.3 | HORVU1Hr1G080920.1 | TRIUR3_26332 | Seita.9G353100.1 | Sobic.001G324900.1 | Zm00001d029627_T002 |
| orthologID_008171 | PH01000733G0140 | LOC_Os03g19680.1 | Bradi1g64320.1 | HORVU4Hr1G053180.1 | TRIUR3_21618 | Seita.9G427500.1 | Sobic.001G393800.1 | Zm00001d028909_T001 |
| orthologID_009643 | PH01000186G1150 | LOC_Os10g34370.1 | Bradi3g29240.1 | HORVU1Hr1G042520.2 | TRIUR3_09389 | Seita.9G216000.1 | Sobic.001G214900.1 | Zm00001d032661_T001 |
| orthologID_008509 | PH01002055G0060 | LOC_Os03g27320.1 | Bradi1g60700.1 | HORVU4Hr1G057650.8 | TRIUR3_31160 | Seita.9G378700.1 | Sobic.001G350400.3 | Zm00001d047217_T001 |
| orthologID_008721 | PH01005362G0060 | LOC_Os03g59440.1 | Bradi1g04470.1 | HORVU5Hr1G110990.1 | TRIUR3_10365 | Seita.9G043200.1 | Sobic.001G044700.1 | Zm00001d034563_T001 |
| orthologID_007710 | PH01000535G0790 | LOC_Os03g63810.1 | Bradi1g01058.2 | HORVU5Hr1G124320.9 | TRIUR3_25300 | Seita.9G005600.1 | Sobic.001G006600.1 | Zm00001d034888_T001 |
| orthologID_007682 | PH01004767G0070 | LOC_Os03g18160.1 | Bradi1g65510.1 | HORVU4Hr1G056100.1 | TRIUR3_28260 | Seita.9G440400.1 | Sobic.001G406100.1 | Zm00001d028753_T003 |
| orthologID_008136 | PH01000992G0570 | LOC_Os03g32580.1 | Bradi1g15850.1 | HORVU7Hr1G098010.4 | TRIUR3_28226 | Seita.9G356500.1 | Sobic.001G329100.1 | Zm00001d029597_T002 |
| orthologID_008124 | PH01000322G1080 | LOC_Os03g18700.1 | Bradi1g65060.2 | HORVU4Hr1G055050.2 | TRIUR3_29514 | Seita.9G436300.1 | Sobic.001G402600.2 | Zm00001d028802_T001 |
| orthologID_007605 | PH01000698G0330 | LOC_Os03g14510.2 | Bradi1g68060.3 | HORVU4Hr1G063020.9 | TRIUR3_34623 | Seita.9G469000.1 | Sobic.001G436400.4 | Zm00001d028424_T002 |
| orthologID_009472 | PH01001507G0330 | LOC_Os10g36030.1 | Bradi3g30270.2 | HORVU1Hr1G046270.2 | TRIUR3_25597 | Seita.9G199900.1 | Sobic.001G201100.1 | Zm00001d032521_T004 |

| | | | | | | | | |
|---|---|---|---|---|---|---|---|---|
| orthologID_010049 | PH01000142G0500 | LOC_Os03g50140.1 | Bradi1g11230.1 | HORVU4Hr1G007190.2 | TRIUR3_05588 | Seita.9G120900.1 | Sobic.001G119200.1 | Zm00001d033709_T001 |
| orthologID_007979 | PH01000913G0280 | LOC_Os03g21870.1 | Bradi1g63120.1 | HORVU4Hr1G049960.1 | TRIUR3_32720 | Seita.9G412200.1 | Sobic.001G380100.2 | Zm00001d029067_T001 |
| orthologID_008266 | PH01000115G0420 | LOC_Os03g08470.1 | Bradi1g72457.1 | HORVU4Hr1G077310.1 | TRIUR3_07780 | Seita.9G516700.1 | Sobic.001G481400.1 | Zm00001d027925_T001 |
| orthologID_007897 | PH01000535G0450 | LOC_Os03g64020.1 | Bradi1g00940.1 | HORVU5Hr1G124210.5 | TRIUR3_31517 | Seita.9G004200.1 | Sobic.001G005200.2 | Zm00001d034900_T001 |
| orthologID_008369 | PH01002072G0370 | LOC_Os03g58940.1 | Bradi1g04930.1 | HORVU5Hr1G109040.1 | TRIUR3_25392 | Seita.9G048200.1 | Sobic.001G050300.1 | Zm00001d034513_T001 |
| orthologID_009313 | PH01001197G0230 | LOC_Os10g27470.1 | Bradi3g26697.3 | HORVU1Hr1G036280.1 | TRIUR3_27934 | Seita.9G261900.1 | Sobic.001G249200.1 | Zm00001d014180_T004 |
| orthologID_008150 | PH01002647G0160 | LOC_Os03g54980.1 | Bradi1g08201.1 | HORVU5Hr1G096200.2 | TRIUR3_04165 | Seita.9G087100.1 | Sobic.001G084900.1 | Zm00001d034064_T001 |
| orthologID_009450 | PH01000912G0640 | LOC_Os02g41470.3 | Bradi3g49150.1 | HORVU6Hr1G058310.3 | TRIUR3_19040 | Seita.9G164300.1 | Sobic.001G022500.1 | Zm00001d034764_T002 |
| orthologID_009188 | PH01001857G0400 | LOC_Os10g42400.1 | Bradi3g34150.1 | HORVU1Hr1G050450.2 | TRIUR3_14773 | Seita.9G311300.1 | Sobic.001G288500.1 | Zm00001d030037_T001 |
| orthologID_008623 | PH01001178G0340 | LOC_Os03g10360.1 | Bradi1g71180.1 | HORVU4Hr1G070400.1 | TRIUR3_07904 | Seita.9G502300.1 | Sobic.001G467100.1 | Zm00001d028113_T001 |
| orthologID_008295 | PH01000365G0420 | LOC_Os05g02810.1 | Bradi1g02010.1 | HORVU5Hr1G120280.1 | TRIUR3_00922 | Seita.9G015800.1 | Sobic.001G016800.1 | Zm00001d007233_T001 |
| orthologID_008210 | PH01001042G0590 | LOC_Os03g55360.1 | Bradi1g07830.1 | HORVU5Hr1G097170.11 | TRIUR3_17665 | Seita.9G081800.1 | Sobic.001G080900.1 | Zm00001d013218_T003 |
| orthologID_008524 | PH01000047G0750 | LOC_Os03g26030.1 | Bradi1g62140.1 | HORVU1Hr1G050340.8 | TRIUR3_14735 | Seita.9G400300.1 | Sobic.001G368100.1 | Zm00001d029206_T002 |
| orthologID_008015 | PH01000193G1120 | LOC_Os03g12890.1 | Bradi1g69160.3 | HORVU0Hr1G008950.4 | TRIUR3_12582 | Seita.9G481900.1 | Sobic.001G447500.2 | Zm00001d028303_T003 |
| orthologID_008068 | PH01001125G0060 | LOC_Os03g52000.1 | Bradi1g09847.1 | HORVU4Hr1G010140.1 | TRIUR3_25194 | Seita.9G107100.1 | Sobic.001G104000.1 | Zm00001d033879_T001 |
| orthologID_007782 | PH01002698G0220 | LOC_Os03g53950.1 | Bradi1g08480.1 | HORVU5Hr1G114720.2 | TRIUR3_05190 | Seita.9G090700.1 | Sobic.001G088200.1 | Zm00001d034032_T001 |
| orthologID_008629 | PH01002850G0100 | LOC_Os03g53250.1 | Bradi1g08977.2 | HORVU5Hr1G094350.1 | TRIUR3_11831 | Seita.9G096600.1 | Sobic.001G094000.1 | Zm00001d033982_T001 |
| orthologID_007685 | PH01003189G0030 | LOC_Os03g55820.1 | Bradi1g07470.2 | HORVU5Hr1G098000.2 | TRIUR3_24927 | Seita.9G077800.1 | Sobic.001G077300.1 | Zm00001d013184_T001 |
| orthologID_008249 | PH01004589G0100 | LOC_Os03g64110.1 | Bradi1g00847.1 | HORVU5Hr1G124850.11 | TRIUR3_11521 | Seita.9G003400.1 | Sobic.001G004100.2 | Zm00001d034915_T001 |
| orthologID_007593 | PH01004461G0140 | LOC_Os03g10320.1 | Bradi1g71210.1 | HORVU4Hr1G070470.4 | TRIUR3_31717 | Seita.9G502900.1 | Sobic.001G467500.1 | Zm00001d028118_T001 |
| orthologID_009428 | PH01002571G0210 | LOC_Os10g41370.1 | Bradi3g33520.1 | HORVU1Hr1G055570.4 | TRIUR3_10676 | Seita.9G321100.1 | Sobic.001G296800.2 | Zm00001d021902_T002 |
| orthologID_008375 | PH01000353G0580 | LOC_Os03g13570.1 | Bradi1g68590.1 | HORVU4Hr1G064220.2 | TRIUR3_22730 | Seita.9G474800.1 | Sobic.001G441700.1 | Zm00001d047970_T001 |
| orthologID_009764 | PH01000215G0080 | LOC_Os10g43075.1 | Bradi3g21588.1 | HORVU1Hr1G051460.14 | TRIUR3_14083 | Seita.9G304500.1 | Sobic.001G280900.2 | Zm00001d030164_T089 |
| orthologID_009640 | PH01001138G0130 | LOC_Os10g33350.1 | Bradi3g28530.2 | HORVU1Hr1G040160.10 | TRIUR3_13227 | Seita.9G225200.1 | Sobic.001G222200.1 | Zm00001d014043_T001 |
| orthologID_009781 | PH01000059G0380 | LOC_Os10g38870.1 | Bradi3g31920.1 | HORVU1Hr1G049420.5 | TRIUR3_34300 | Seita.9G344700.1 | Sobic.001G316400.1 | Zm00001d029716_T002 |
| orthologID_007729 | PH01000033G2130 | LOC_Os03g17560.1 | Bradi1g65920.2 | HORVU4Hr1G057530.2 | TRIUR3_12157 | Seita.9G445300.1 | Sobic.001G411500.1 | Zm00001d045529_T002 |
| orthologID_008385 | PH01003916G0020 | LOC_Os03g58150.1 | Bradi1g05560.1 | HORVU5Hr1G105760.4 | TRIUR3_04678 | Seita.9G056500.1 | Sobic.001G057800.1 | Zm00001d034432_T001 |
| orthologID_007663 | PH01001574G0020 | LOC_Os03g58620.1 | Bradi1g05156.2 | HORVU5Hr1G107030.1 | TRIUR3_31893 | Seita.9G051700.1 | Sobic.001G053000.1 | Zm00001d013061_T002 |
| orthologID_007890 | PH01002250G0270 | LOC_Os03g25610.1 | Bradi1g61390.1 | HORVU4Hr1G078380.1 | TRIUR3_32035 | Seita.9G390600.1 | Sobic.001G358700.1 | Zm00001d029299_T001 |
| orthologID_007966 | PH01003159G0100 | LOC_Os03g43410.1 | Bradi1g14230.1 | HORVU5Hr1G093640.1 | TRIUR3_01368 | Seita.9G162800.1 | Sobic.001G161500.1 | Zm00001d033319_T001 |
| orthologID_009189 | PH01002699G0200 | LOC_Os10g42700.1 | Bradi3g34245.3 | HORVU1Hr1G050970.1 | TRIUR3_22495 | Seita.9G307600.1 | Sobic.001G284200.1 | Zm00001d030109_T002 |
| orthologID_008205 | PH01000531G0510 | LOC_Os03g11500.1 | Bradi1g70040.2 | HORVU4Hr1G068030.12 | TRIUR3_24734 | Seita.9G491700.1 | Sobic.001G457300.1 | Zm00001d048092_T002 |
| orthologID_007926 | PH01000648G0310 | LOC_Os03g18020.1 | Bradi1g65600.1 | HORVU4Hr1G056550.23 | TRIUR3_20630 | Seita.9G441700.1 | Sobic.001G407900.2 | Zm00001d028745_T001 |
| orthologID_008174 | PH01002657G0040 | LOC_Os03g17140.1 | Bradi1g66340.1 | HORVU4Hr1G058760.1 | TRIUR3_24750 | Seita.9G449700.1 | Sobic.001G416300.3 | Zm00001d028665_T001 |
| orthologID_009680 | PH01004280G0050 | LOC_Os08g18044.2 | Bradi3g23140.1 | HORVU1Hr1G034980.3 | TRIUR3_29725 | Seita.9G314900.1 | Sobic.001G291100.1 | Zm00001d030014_T008 |
| orthologID_007913 | PH01000430G0520 | LOC_Os03g20560.1 | Bradi1g63900.1 | HORVU4Hr1G052000.14 | TRIUR3_06230 | Seita.9G421600.1 | Sobic.001G388900.1 | Zm00001d038977_T001 |
| orthologID_008090 | PH01001811G0040 | LOC_Os03g15530.1 | Bradi1g67440.1 | HORVU4Hr1G061680.1 | TRIUR3_10373 | Seita.9G454500.1 | Sobic.001G429700.1 | Zm00001d028510_T001 |
| orthologID_008000 | PH01000648G0050 | LOC_Os03g17860.1 | Bradi1g65710.1 | HORVU4Hr1G056730.1 | TRIUR3_09363 | Seita.9G442800.1 | Sobic.001G408900.1 | Zm00001d028726_T001 |
| orthologID_007780 | PH01001432G0330 | LOC_Os03g60430.2 | Bradi1g03880.1 | HORVU5Hr1G112440.1 | TRIUR3_24919 | Seita.9G035800.1 | Sobic.001G036800.1 | Zm00001d034629_T001 |
| orthologID_007584 | PH01000678G0300 | LOC_Os03g48840.3 | Bradi1g11882.2 | HORVU4Hr1G005500.49 | TRIUR3_08729 | Seita.9G130700.1 | Sobic.001G129300.1 | Zm00001d033590_T001 |
| orthologID_007575 | PH01000430G0510 | LOC_Os03g20550.1 | Bradi1g63910.1 | HORVU4Hr1G052010.2 | TRIUR3_06231 | Seita.9G421700.1 | Sobic.001G389000.1 | Zm00001d028962_T001 |
| orthologID_009204 | PH01002667G0150 | LOC_Os10g42860.1 | Bradi3g34580.1 | HORVU1Hr1G051250.1 | TRIUR3_03770 | Seita.9G305700.1 | Sobic.001G282200.2 | Zm00001d030146_T003 |
| orthologID_007704 | PH01050909G0010 | LOC_Os03g58930.1 | Bradi1g04940.1 | HORVU5Hr1G108970.4 | TRIUR3_10103 | Seita.9G048300.1 | Sobic.001G050400.1 | Zm00001d034512_T002 |
| orthologID_007835 | PH01001119G0310 | LOC_Os03g58350.1 | Bradi1g05410.1 | HORVU5Hr1G106350.2 | TRIUR3_13609 | Seita.9G054500.1 | Sobic.001G056100.1 | Zm00001d013071_T001 |
| orthologID_007826 | PH01002899G0210 | LOC_Os03g55784.1 | Bradi1g07490.1 | HORVU5Hr1G098040.1 | TRIUR3_15746 | Seita.9G078000.1 | Sobic.001G077500.2 | Zm00001d034183_T001 |
| orthologID_008209 | PH01001337G0440 | LOC_Os03g26960.1 | Bradi1g60890.1 | HORVU4Hr1G035890.7 | TRIUR3_19133 | Seita.9G381200.1 | Sobic.001G352900.1 | Zm00001d029375_T002 |
| orthologID_007803 | PH01000113G0120 | LOC_Os03g07260.1 | Bradi1g73790.1 | HORVU4Hr1G074850.2 | TRIUR3_05041 | Seita.9G525900.1 | Sobic.001G491300.1 | Zm00001d027838_T001 |
| orthologID_009520 | PH01003555G0170 | LOC_Os10g11060.1 | Bradi3g22920.3 | HORVU1Hr1G033060.1 | TRIUR3_27065 | Seita.9G284300.1 | Sobic.001G276200.1 | Zm00001d030409_T001 |
| orthologID_008260 | PH01000589G0240 | LOC_Os03g56190.1 | Bradi1g07230.1 | HORVU5Hr1G098690.2 | TRIUR3_01395 | Seita.9G074400.1 | Sobic.001G074700.1 | Zm00001d034218_T001 |
| orthologID_008228 | PH01001727G0210 | LOC_Os03g24830.1 | Bradi1g61771.1 | HORVU6Hr1G080400.2 | TRIUR3_17886 | Seita.9G396200.1 | Sobic.001G363000.1 | Zm00001d029248_T002 |

| | | | | | | | | |
|---|---|---|---|---|---|---|---|---|
| orthologID_008710 | PH01000634G0760 | LOC_Os03g51550.2 | Bradi1g10180.1 | HORVU4Hr1G009500.1 | TRIUR3_15901 | Seita.9G110500.1 | Sobic.001G107300.2 | Zm00001d033849_T002 |
| orthologID_008130 | PH01000339G1090 | LOC_Os03g13750.1 | Bradi1g68520.1 | HORVU4Hr1G064080.1 | TRIUR3_23930 | Seita.9G474000.1 | Sobic.001G441000.1 | Zm00001d028375_T001 |
| orthologID_007857 | PH01001328G0340 | LOC_Os03g07100.1 | Bradi1g73910.1 | HORVU4Hr1G074480.3 | TRIUR3_30825 | Seita.9G527600.1 | Sobic.001G493100.1 | Zm00001d027812_T001 |
| orthologID_008545 | PH01002278G0240 | LOC_Os03g17580.1 | Bradi1g65900.1 | HORVU4Hr1G057450.1 | TRIUR3_33965 | Seita.9G445100.1 | Sobic.001G411300.1 | Zm00001d028702_T001 |
| orthologID_008114 | PH01000040G1640 | LOC_Os03g61530.1 | Bradi1g03056.1 | HORVU5Hr1G115810.1 | TRIUR3_23333 | Seita.9G024800.1 | Sobic.001G024300.1 | Zm00001d034747_T001 |
| orthologID_008700 | PH01001260G0450 | LOC_Os03g57080.1 | Bradi1g06610.1 | HORVU5Hr1G103060.1 | TRIUR3_10892 | Seita.9G065800.1 | Sobic.001G067500.1 | Zm00001d034340_T001 |
| orthologID_009675 | PH01000059G1270 | LOC_Os10g39130.1 | Bradi3g32090.1 | HORVU1Hr1G051660.8 | TRIUR3_32699 | Seita.9G342700.1 | Sobic.001G314400.1 | Zm00001d027425_T001 |
| orthologID_007954 | PH01000804G0610 | LOC_Os03g61280.1 | Bradi1g03207.1 | HORVU5Hr1G115310.4 | TRIUR3_10862 | Seita.9G021900.1 | Sobic.001G027800.2 | Zm00001d034723_T001 |
| orthologID_007935 | PH01001584G0050 | LOC_Os03g47730.1 | Bradi1g12520.2 | HORVU4Hr1G003750.1 | TRIUR3_09679 | Seita.9G139700.1 | Sobic.001G137200.1 | Zm00001d033379_T001 |
| orthologID_009277 | PH01002152G0170 | LOC_Os10g39150.1 | Bradi3g32100.1 | HORVU1Hr1G051700.6 | TRIUR3_32150 | Seita.9G342500.1 | Sobic.001G314200.1 | Zm00001d029745_T003 |
| orthologID_008664 | PH01000033G2260 | LOC_Os03g17510.1 | Bradi1g65990.1 | HORVU4Hr1G057730.2 | TRIUR3_06021 | Seita.9G445800.1 | Sobic.001G412200.1 | Zm00001d028697_T001 |
| orthologID_008588 | PH01000376G0230 | LOC_Os03g56300.1 | Bradi1g07150.1 | HORVU5Hr1G098910.2 | TRIUR3_05436 | Seita.9G072900.1 | Sobic.001G073600.1 | Zm00001d013156_T002 |
| orthologID_008159 | PH01001015G0430 | LOC_Os07g47630.1 | Bradi1g18070.2 | HORVU2Hr1G019410.4 | TRIUR3_07917 | Seita.9G408600.1 | Sobic.001G375700.1 | Zm00001d047498_T010 |
| orthologID_008507 | PH01000129G0320 | LOC_Os03g09180.1 | Bradi1g71730.1 | HORVU4Hr1G071920.1 | TRIUR3_23160 | Seita.9G509300.1 | Sobic.001G473800.1 | Zm00001d028018_T001 |
| orthologID_008306 | PH01005165G0030 | LOC_Os03g05020.1 | Bradi1g75470.3 | HORVU4Hr1G078840.3 | TRIUR3_19094 | Seita.9G543000.1 | Sobic.001G507200.1 | Zm00001d027633_T011 |
| orthologID_008096 | PH01001946G0100 | LOC_Os03g46650.1 | Bradi1g12820.1 | HORVU4Hr1G003110.4 | TRIUR3_10244 | Seita.9G145400.1 | Sobic.001G142100.1 | Zm00001d033422_T001 |
| orthologID_007846 | PH01001096G0120 | LOC_Os03g51920.1 | Bradi1g09950.2 | HORVU4Hr1G009990.2 | TRIUR3_07901 | Seita.9G108100.1 | Sobic.001G104900.1 | Zm00001d033872_T001 |
| orthologID_009569 | PH01000230G0730 | LOC_Os10g05130.1 | Bradi3g21140.1 | HORVU6Hr1G078260.2 | TRIUR3_18793 | Seita.9G290400.1 | Sobic.001G269900.1 | Zm00001d030267_T001 |
| orthologID_007670 | PH01001786G0150 | LOC_Os03g10110.1 | Bradi1g71350.1 | HORVU4Hr1G070970.1 | TRIUR3_16678 | Seita.9G504700.1 | Sobic.001G469300.1 | Zm00001d048176_T010 |
| orthologID_008482 | PH01002996G0020 | LOC_Os03g12236.1 | Bradi1g69564.1 | HORVU4Hr1G066920.5 | TRIUR3_24351 | Seita.9G486200.1 | Sobic.001G452000.2 | Zm00001d028256_T002 |
| orthologID_007778 | PH01002187G0040 | LOC_Os03g55530.1 | Bradi1g07680.4 | HORVU5Hr1G097440.1 | TRIUR3_31724 | Seita.9G080600.1 | Sobic.001G079600.1 | Zm00001d034143_T001 |
| orthologID_008471 | PH01000081G1390 | LOC_Os03g06160.1 | Bradi1g74630.2 | HORVU4Hr1G072590.5 | TRIUR3_02107 | Seita.9G534900.1 | Sobic.001G499000.2 | Zm00001d048420_T001 |
| orthologID_007955 | PH01000143G1600 | LOC_Os03g02590.2 | Bradi1g77380.1 | HORVU4Hr1G085960.2 | TRIUR3_34840 | Seita.9G566700.1 | Sobic.001G531500.1 | Zm00001d027366_T002 |
| orthologID_008352 | PH01002188G0320 | LOC_Os08g39820.1 | Bradi1g13170.2 | HORVU4Hr1G002500.3 | TRIUR3_11840 | Seita.5G427200.1 | Sobic.001G147100.1 | Zm00001d013626_T002 |
| orthologID_008180 | PH01001109G0260 | LOC_Os03g17010.1 | Bradi1g66430.1 | HORVU4Hr1G059090.11 | TRIUR3_31184 | Seita.9G450900.1 | Sobic.001G417900.1 | Zm00001d028655_T002 |
| orthologID_008311 | PH01002074G0190 | LOC_Os03g09870.1 | Bradi1g71570.1 | HORVU4Hr1G079170.3 | TRIUR3_21995 | Seita.9G507000.1 | Sobic.001G471500.2 | Zm00001d048193_T002 |
| orthologID_009620 | PH01180127G0010 | LOC_Os05g33860.1 | Bradi3g39960.1 | HORVU1Hr1G060580.2 | TRIUR3_21980 | Seita.3G245200.1 | Sobic.001G295800.1 | Zm00001d029950_T001 |
| orthologID_008470 | PH01001003G0270 | LOC_Os03g15720.2 | Bradi1g67290.1 | HORVU4Hr1G061190.1 | TRIUR3_22245 | Seita.9G455900.1 | Sobic.001G428200.1 | Zm00001d028531_T001 |
| orthologID_008484 | PH01004682G0110 | LOC_Os03g44820.1 | Bradi1g13490.2 | HORVU5Hr1G092240.2 | TRIUR3_20879 | Seita.9G153300.1 | Sobic.001G151700.1 | Zm00001d013656_T001 |
| orthologID_008483 | PH01014296G0010 | LOC_Os01g13190.1 | Bradi1g17340.1 | HORVU2Hr1G072220.7 | TRIUR3_21528 | Seita.9G065600.1 | Sobic.001G067300.3 | Zm00001d034343_T001 |
| orthologID_008177 | PH01002476G0330 | LOC_Os03g60150.1 | Bradi1g04080.1 | HORVU5Hr1G111670.1 | TRIUR3_21758 | Seita.9G038400.1 | Sobic.001G039500.2 | Zm00001d034608_T004 |
| orthologID_009409 | PH01002340G0130 | LOC_Os10g42620.1 | Bradi3g34280.1 | HORVU1Hr1G050830.3 | TRIUR3_21996 | Seita.9G308700.1 | Sobic.001G285400.1 | Zm00001d030090_T001 |
| orthologID_008497 | PH01000181G0680 | LOC_Os03g19390.1 | Bradi1g64530.2 | HORVU4Hr1G053950.1 | TRIUR3_29340 | Seita.9G430100.1 | Sobic.001G396200.1 | Zm00001d047640_T001 |
| orthologID_008257 | PH01001259G0100 | LOC_Os03g01190.1 | Bradi1g78420.2 | HORVU4Hr1G088610.1 | TRIUR3_09107 | Seita.9G579100.1 | Sobic.001G543300.1 | Zm00001d027276_T001 |
| orthologID_007815 | PH01000908G0440 | LOC_Os03g36439.1 | Bradi1g15750.1 | HORVU2Hr1G096680.2 | TRIUR3_22639 | Seita.9G186700.1 | Sobic.001G189600.1 | Zm00001d013824_T001 |
| orthologID_008617 | PH01001401G0280 | LOC_Os03g23970.1 | Bradi1g62230.1 | HORVU5Hr1G050850.1 | TRIUR3_33770 | Seita.9G400800.1 | Sobic.001G368800.1 | Zm00001d029198_T003 |
| orthologID_007630 | PH01000017G0450 | LOC_Os03g02820.1 | Bradi1g77210.2 | HORVU4Hr1G085310.21 | TRIUR3_01723 | Seita.9G563800.1 | Sobic.001G528900.1 | Zm00001d027402_T004 |
| orthologID_008252 | PH01001098G0450 | LOC_Os03g58340.1 | Bradi1g05420.2 | HORVU5Hr1G106320.1 | TRIUR3_13608 | Seita.9G054600.1 | Sobic.001G056200.1 | Zm00001d013072_T001 |
| orthologID_007915 | PH01001686G0270 | LOC_Os03g26440.1 | Bradi1g61057.2 | HORVU3Hr1G016330.9 | TRIUR3_00522 | Seita.9G384000.1 | Sobic.001G355200.3 | Zm00001d029342_T015 |
| orthologID_008301 | PH01000017G0910 | LOC_Os03g02670.2 | Bradi1g77330.1 | HORVU4Hr1G085820.1 | TRIUR3_33835 | Seita.9G565500.1 | Sobic.001G530700.1 | Zm00001d048499_T001 |
| orthologID_008388 | PH01000971G0560 | LOC_Os03g45400.1 | Bradi1g13226.1 | HORVU4Hr1G002350.1 | TRIUR3_32835 | Seita.9G150400.1 | Sobic.001G148200.1 | Zm00001d033473_T001 |
| orthologID_008495 | PH01000017G0530 | LOC_Os03g02762.1 | Bradi1g77240.1 | HORVU4Hr1G085380.2 | TRIUR3_02097 | Seita.9G564300.1 | Sobic.001G529500.1 | Zm00001d027393_T001 |
| orthologID_008080 | PH01000413G0240 | LOC_Os03g14180.1 | Bradi1g68440.1 | HORVU4Hr1G063350.1 | TRIUR3_21891 | Seita.9G470600.1 | Sobic.001G438000.1 | Zm00001d028408_T001 |
| orthologID_008017 | PH01001727G0310 | LOC_Os03g24880.1 | Bradi1g61740.2 | HORVU4Hr1G045500.6 | TRIUR3_14283 | Seita.9G395400.1 | Sobic.001G362400.2 | Zm00001d029255_T001 |
| orthologID_007831 | PH01001584G0340 | LOC_Os03g47949.1 | Bradi1g12340.2 | HORVU4Hr1G004090.6 | TRIUR3_15387 | Seita.9G138500.1 | Sobic.001G135500.1 | Zm00001d013540_T020 |
| orthologID_008491 | PH01000168G0490 | LOC_Os03g02200.1 | Bradi1g60190.2 | HORVU1Hr1G083370.4 | TRIUR3_23661 | Seita.9G352500.1 | Sobic.001G324400.1 | Zm00001d029632_T002 |
| orthologID_009186 | PH01000903G0430 | LOC_Os01g73514.1 | Bradi3g31390.1 | HORVU6Hr1G076190.1 | TRIUR3_30061 | Seita.9G351000.1 | Sobic.001G322700.1 | Zm00001d029652_T001 |
| orthologID_008269 | PH01000496G0070 | LOC_Os03g46640.1 | Bradi1g12830.2 | HORVU4Hr1G003120.1 | TRIUR3_10245 | Seita.9G145300.1 | Sobic.001G142000.1 | Zm00001d033421_T001 |
| orthologID_008047 | PH01000033G1930 | LOC_Os03g17634.1 | Bradi1g65860.1 | HORVU4Hr1G057330.1 | TRIUR3_24096 | Seita.9G444700.1 | Sobic.001G410700.1 | Zm00001d047759_T003 |
| orthologID_007830 | PH01000184G1040 | LOC_Os03g39000.1 | Bradi1g15350.2 | HORVU4Hr1G034300.6 | TRIUR3_20246 | Seita.9G177900.1 | Sobic.001G178100.1 | Zm00001d033110_T001 |

| | | | | | | | | |
|---|---|---|---|---|---|---|---|---|
| orthologID_008641 | PH01000007G1740 | LOC_Os03g13000.1 | Bradi1g69230.2 | HORVU4Hr1G066010.2 | TRIUR3_26704 | Seita.9G482700.1 | Sobic.001G448600.1 | Zm00001d048007_T003 |
| orthologID_008175 | PH01000804G0270 | LOC_Os03g61100.1 | Bradi1g03350.1 | HORVU5Hr1G114990.1 | TRIUR3_24467 | Seita.9G028200.1 | Sobic.001G029500.1 | Zm00001d034706_T001 |
| orthologID_008706 | PH01003523G0190 | LOC_Os03g41612.1 | Bradi1g14670.1 | HORVU4Hr1G013640.1 | TRIUR3_06487 | Seita.9G168700.1 | Sobic.001G167500.1 | Zm00001d013730_T001 |
| orthologID_006261 | PH01000004G3970 | LOC_Os03g60200.1 | Bradi2g55230.1 | HORVU3Hr1G085150.2 | TRIUR3_15440 | Seita.5G385400.1 | Sobic.001G151300.1 | Zm00001d033495_T001 |
| orthologID_008244 | PH01001003G0150 | LOC_Os03g15650.1 | Bradi1g67360.1 | HORVU4Hr1G061320.1 | TRIUR3_13259 | Seita.9G455300.1 | Sobic.001G428800.1 | Zm00001d047863_T004 |
| orthologID_008279 | PH01000490G1060 | LOC_Os03g64170.1 | Bradi1g00790.2 | HORVU5Hr1G125000.2 | TRIUR3_30090 | Seita.9G003000.1 | Sobic.001G003600.1 | Zm00001d034917_T001 |
| orthologID_008043 | PH01000010G3080 | LOC_Os03g05720.1 | Bradi1g74940.1 | HORVU4Hr1G083410.1 | TRIUR3_01164 | Seita.9G537800.1 | Sobic.001G501400.1 | Zm00001d027706_T004 |
| orthologID_009890 | PH01006518G0030 | LOC_Os08g29590.1 | Bradi3g34880.1 | HORVU7Hr1G063520.1 | TRIUR3_23047 | Seita.6G132500.1 | Sobic.001G113500.1 | Zm00001d033780_T004 |
| orthologID_009689 | PH01001197G0310 | LOC_Os10g27450.1 | Bradi3g26680.2 | HORVU1Hr1G034570.9 | TRIUR3_28717 | Seita.9G261700.1 | Sobic.001G249400.1 | Zm00001d032968_T003 |
| orthologID_008558 | PH01000142G0430 | LOC_Os03g50220.1 | Bradi1g11260.1 | HORVU4Hr1G007340.5 | TRIUR3_05591 | Seita.9G120500.1 | Sobic.001G118800.1 | Zm00001d013447_T001 |
| orthologID_007694 | PH01002189G0190 | LOC_Os03g57090.1 | Bradi1g06590.1 | HORVU5Hr1G103080.2 | TRIUR3_11687 | Seita.9G065700.1 | Sobic.001G067400.1 | Zm00001d034341_T003 |
| orthologID_007865 | PH01001178G0560 | LOC_Os03g10460.1 | Bradi1g71057.1 | HORVU1Hr1G012220.1 | TRIUR3_02495 | Seita.9G501100.1 | Sobic.001G466200.1 | Zm00001d028104_T003 |
| orthologID_008420 | PH01000113G0020 | LOC_Os03g07200.1 | Bradi1g73817.1 | HORVU4Hr1G074780.1 | TRIUR3_19441 | Seita.9G526600.1 | Sobic.001G491800.1 | Zm00001d048333_T001 |
| orthologID_008654 | PH01000154G1020 | LOC_Os03g15900.2 | Bradi1g67140.1 | HORVU4Hr1G060880.1 | TRIUR3_17910 | Seita.9G457500.1 | Sobic.001G426600.1 | Zm00001d028546_T032 |
| orthologID_009790 | PH01000321G0550 | LOC_Os10g37840.2 | Bradi3g31527.1 | HORVU1Hr1G048720.16 | TRIUR3_22053 | Seita.9G348900.1 | Sobic.001G321400.1 | Zm00001d029674_T001 |
| orthologID_009736 | PH01001349G0080 | LOC_Os10g28665.1 | Bradi3g27027.6 | HORVU4Hr1G087570.39 | TRIUR3_16600 | Seita.9G265600.1 | Sobic.001G245000.1 | Zm00001d014162_T001 |
| orthologID_007580 | PH01165045G0010 | LOC_Os03g42480.1 | Bradi1g14486.2 | HORVU4Hr1G012070.13 | TRIUR3_18662 | Seita.9G165900.1 | Sobic.001G164700.1 | Zm00001d033312_T006 |
| orthologID_007570 | PH01000129G0560 | LOC_Os03g09120.1 | Bradi1g71790.1 | HORVU0Hr1G009280.1 | TRIUR3_23859 | Seita.9G509900.1 | Sobic.001G474300.1 | Zm00001d028011_T001 |
| orthologID_008087 | PH01002317G0050 | LOC_Os08g14450.1 | Bradi1g00750.1 | HORVU5Hr1G125110.14 | TRIUR3_06257 | Seita.9G002700.1 | Sobic.001G003200.4 | Zm00001d012810_T003 |
| orthologID_007817 | PH01000829G0320 | LOC_Os03g16580.1 | Bradi1g66750.1 | HORVU4Hr1G060000.1 | TRIUR3_29768 | Seita.9G462000.1 | Sobic.001G422600.2 | Zm00001d028596_T001 |
| orthologID_007838 | PH01000532G0300 | LOC_Os03g14840.1 | Bradi1g67887.1 | HORVU4Hr1G062610.1 | TRIUR3_18097 | Seita.9G467200.1 | Sobic.001G434600.1 | Zm00001d028447_T001 |
| orthologID_008431 | PH01001537G0320 | LOC_Os03g11120.1 | Bradi1g70570.1 | HORVU4Hr1G069130.2 | TRIUR3_10498 | Seita.9G496100.1 | Sobic.001G461700.1 | Zm00001d028038_T008 |
| orthologID_008307 | PH01000921G0270 | LOC_Os03g02040.1 | Bradi1g77840.1 | HORVU4Hr1G089540.2 | TRIUR3_18094 | Seita.9G571000.1 | Sobic.001G535400.1 | Zm00001d027330_T002 |
| orthologID_008050 | PH01003162G0250 | LOC_Os03g60870.1 | Bradi1g03490.1 | HORVU5Hr1G114250.1 | TRIUR3_08443 | Seita.9G030400.1 | Sobic.001G031900.2 | Zm00001d034675_T002 |
| orthologID_007921 | PH01001995G0110 | LOC_Os03g40260.1 | Bradi1g15060.2 | HORVU5Hr1G053180.5 | TRIUR3_21947 | Seita.9G173800.1 | Sobic.001G173400.1 | Zm00001d033172_T001 |
| orthologID_008120 | PH01000353G0210 | LOC_Os03g13450.2 | Bradi1g68720.1 | HORVU4Hr1G064590.2 | TRIUR3_20798 | Seita.9G475800.1 | Sobic.001G442900.4 | Zm00001d028362_T001 |
| orthologID_007976 | PH01000371G0700 | LOC_Os03g63720.1 | Bradi1g01100.1 | HORVU5Hr1G123690.1 | TRIUR3_08678 | Seita.9G006200.1 | Sobic.001G007500.1 | Zm00001d012826_T003 |
| orthologID_009097 | PH01002302G0150 | LOC_Os10g35140.1 | Bradi3g29830.1 | HORVU1Hr1G043730.7 | TRIUR3_22465 | Seita.9G207700.1 | Sobic.001G207800.1 | Zm00001d032604_T001 |
| orthologID_009403 | PH01002569G0160 | LOC_Os10g30290.1 | Bradi3g27407.2 | HORVU1Hr1G036440.1 | TRIUR3_10149 | Seita.9G243000.1 | Sobic.001G238800.1 | Zm00001d032888_T001 |
| orthologID_007853 | PH01002201G0080 | LOC_Os03g21000.1 | Bradi1g63640.1 | HORVU4Hr1G051400.3 | TRIUR3_29500 | Seita.9G418200.1 | Sobic.001G386200.1 | Zm00001d047555_T001 |
| orthologID_008554 | PH01000157G0610 | LOC_Os03g05200.1 | Bradi1g75390.2 | HORVU4Hr1G079100.5 | TRIUR3_23348 | Seita.9G541800.1 | Sobic.001G505600.1 | Zm00001d048392_T006 |
| orthologID_007747 | PH01001010G0200 | LOC_Os03g17790.1 | Bradi1g65780.1 | HORVU4Hr1G057020.2 | TRIUR3_33288 | Seita.9G443500.1 | Sobic.001G409400.1 | Zm00001d028718_T001 |
| orthologID_008616 | PH01003173G0150 | LOC_Os03g44290.1 | Bradi1g13787.4 | HORVU5Hr1G092910.2 | TRIUR3_08652 | Seita.9G157200.1 | Sobic.001G155700.5 | Zm00001d033231_T001 |
| orthologID_009150 | PH01003340G0200 | LOC_Os02g17850.1 | Bradi3g33910.1 | HORVU1Hr1G054340.3 | TRIUR3_09175 | Seita.9G314500.1 | Sobic.001G290766.1 | Zm00001d030018_T001 |
| orthologID_008182 | PH01002606G0090 | LOC_Os03g53360.1 | Bradi1g08880.1 | HORVU5Hr1G094460.3 | TRIUR3_27392 | Seita.9G095700.1 | Sobic.001G093000.1 | Zm00001d033989_T001 |
| orthologID_008321 | PH01001812G0170 | LOC_Os03g57170.1 | Bradi1g06470.1 | HORVU5Hr1G103360.5 | TRIUR3_01382 | Seita.9G064700.1 | Sobic.001G066300.1 | Zm00001d034351_T001 |
| orthologID_008383 | PH01004482G0060 | LOC_Os10g11500.1 | Bradi1g57590.1 | HORVU7Hr1G040730.1 | TRIUR3_07301 | Seita.9G290900.1 | Sobic.001G333400.1 | Zm00001d029558_T001 |
| orthologID_007985 | PH01001831G0240 | LOC_Os03g29730.1 | Bradi1g60100.1 | HORVU6Hr1G009170.26 | TRIUR3_19768 | Seita.9G368000.1 | Sobic.001G340600.1 | Zm00001d029482_T002 |
| orthologID_007814 | PH01000716G0290 | LOC_Os03g54870.1 | Bradi1g08270.4 | HORVU5Hr1G096070.1 | TRIUR3_17672 | Seita.9G087900.1 | Sobic.001G085500.1 | Zm00001d013253_T001 |
| orthologID_009713 | PH01000194G0730 | LOC_Os10g28360.1 | Bradi3g26930.1 | HORVU1Hr1G083380.2 | TRIUR3_25016 | Seita.9G267000.1 | Sobic.001G243200.1 | Zm00001d032925_T001 |
| orthologID_008267 | PH01003966G0020 | LOC_Os03g55470.1 | Bradi1g07760.1 | HORVU5Hr1G097370.4 | TRIUR3_14672 | Seita.9G081000.1 | Sobic.001G079900.1 | Zm00001d034133_T001 |
| orthologID_007624 | PH01000465G0750 | LOC_Os03g18520.1 | Bradi1g65197.1 | HORVU4Hr1G055410.1 | TRIUR3_29694 | Seita.9G438100.1 | Sobic.001G404000.1 | Zm00001d047712_T001 |
| orthologID_007676 | PH01001481G0370 | LOC_Os03g38590.1 | Bradi1g15490.1 | HORVU5Hr1G061580.4 | TRIUR3_05370 | Seita.9G179800.1 | Sobic.001G180900.1 | Zm00001d033090_T002 |
| orthologID_010076 | PH01004495G0130 | LOC_Os03g11650.1 | Bradi1g69870.1 | HORVU4Hr1G067630.7 | TRIUR3_10457 | Seita.9G490100.1 | Sobic.001G455700.1 | Zm00001d028219_T001 |
| orthologID_007855 | PH01001161G0520 | LOC_Os03g53600.1 | Bradi1g08760.5 | HORVU5Hr1G094670.2 | TRIUR3_05232 | Seita.9G094300.1 | Sobic.001G091300.3 | Zm00001d034001_T001 |
| orthologID_007960 | PH01000017G1420 | LOC_Os03g02450.1 | Bradi1g77505.2 | HORVU4Hr1G086630.1 | TRIUR3_08693 | Seita.9G568400.1 | Sobic.001G532600.1 | Zm00001d027352_T001 |
| orthologID_007930 | PH01000299G0820 | LOC_Os06g21530.1 | Bradi1g74030.1 | HORVU4Hr1G074320.3 | TRIUR3_12218 | Seita.9G528500.1 | Sobic.001G493800.1 | Zm00001d048441_T001 |
| orthologID_007727 | PH01000698G0290 | LOC_Os03g14520.1 | Bradi1g68050.1 | HORVU4Hr1G063000.1 | TRIUR3_31720 | Seita.9G468900.1 | Sobic.001G436300.1 | Zm00001d028425_T003 |
| orthologID_008412 | PH01055856G0010 | LOC_Os03g51960.1 | Bradi1g09910.1 | HORVU4Hr1G010120.1 | TRIUR3_34446 | Seita.9G107800.1 | Sobic.001G104600.1 | Zm00001d033874_T002 |
| orthologID_009399 | PH01002702G0180 | LOC_Os06g30460.1 | Bradi3g17280.4 | HORVU7Hr1G072650.1 | TRIUR3_27772 | Seita.9G045700.1 | Sobic.001G047600.1 | Zm00001d034531_T001 |

| | | | | | | | | |
|---|---|---|---|---|---|---|---|---|
| orthologID_008366 | PH01000322G0740 | LOC_Os03g18570.1 | Bradi1g65150.4 | HORVU4Hr1G055230.1 | TRIUR3_17780 | Seita.9G437500.1 | Sobic.001G403500.1 | Zm00001d028786_T003 |
| orthologID_008086 | PH01000921G0090 | LOC_Os03g01970.1 | Bradi1g77887.1 | HORVU4Hr1G089820.7 | TRIUR3_26611 | Seita.9G572100.1 | Sobic.001G536400.1 | Zm00001d027324_T002 |
| orthologID_008094 | PH01000531G0180 | LOC_Os03g11400.1 | Bradi1g70210.2 | HORVU4Hr1G068470.10 | TRIUR3_08563 | Seita.9G493100.1 | Sobic.001G458900.1 | Zm00001d028195_T001 |
| orthologID_008183 | PH01000115G1590 | LOC_Os03g08860.1 | Bradi1g72060.3 | HORVU4Hr1G078130.1 | TRIUR3_06441 | Seita.9G512700.1 | Sobic.001G477300.1 | Zm00001d027976_T002 |
| orthologID_007986 | PH01000113G0130 | LOC_Os03g07300.1 | Bradi1g73770.1 | HORVU4Hr1G074810.5 | TRIUR3_11821 | Seita.9G525700.1 | Sobic.001G491000.1 | Zm00001d027841_T002 |
| orthologID_007730 | PH01000115G0530 | LOC_Os03g08520.1 | Bradi1g72440.1 | HORVU4Hr1G077350.2 | TRIUR3_26835 | Seita.9G516300.1 | Sobic.001G481100.1 | Zm00001d027932_T001 |
| orthologID_008714 | PH01000753G0680 | LOC_Os03g48110.1 | Bradi1g12210.3 | HORVU2Hr1G033130.1 | TRIUR3_13984 | Seita.9G136800.1 | Sobic.001G134200.1 | Zm00001d033531_T001 |
| orthologID_007619 | PH01000113G0530 | LOC_Os03g07440.1 | Bradi1g73200.1 | HORVU4Hr1G075160.1 | TRIUR3_12806 | Seita.9G524300.1 | Sobic.001G489000.1 | Zm00001d048313_T001 |
| orthologID_008549 | PH01002172G0070 | LOC_Os03g53580.1 | Bradi1g08790.1 | HORVU5Hr1G094610.2 | TRIUR3_16984 | Seita.9G094600.1 | Sobic.001G091500.1 | Zm00001d033999_T001 |
| orthologID_008523 | PH01001477G0200 | LOC_Os03g55330.1 | Bradi1g07860.1 | HORVU5Hr1G097100.1 | TRIUR3_10016 | Seita.9G082100.1 | Sobic.001G081400.1 | Zm00001d013219_T001 |
| orthologID_007839 | PH01000010G2650 | LOC_Os03g05470.1 | Bradi1g75140.1 | HORVU4Hr1G080110.9 | TRIUR3_05676 | Seita.9G540100.1 | Sobic.001G503300.1 | Zm00001d027676_T001 |
| orthologID_007739 | PH01000707G0790 | LOC_Os03g63510.1 | Bradi1g01280.1 | HORVU5Hr1G123150.1 | TRIUR3_10042 | Seita.9G007900.1 | Sobic.001G009400.1 | Zm00001d034870_T001 |
| orthologID_009240 | PH01000012G1160 | LOC_Os10g36340.2 | Bradi3g30370.2 | HORVU1Hr1G046610.1 | TRIUR3_08682 | Seita.9G198100.1 | Sobic.001G199900.2 | Zm00001d032508_T001 |
| orthologID_008192 | PH01001995G0240 | LOC_Os03g40310.1 | Bradi1g15040.4 | HORVU6Hr1G087100.1 | TRIUR3_14061 | Seita.9G173600.1 | Sobic.001G172800.1 | Zm00001d013750_T001 |
| orthologID_008577 | PH01000376G0330 | LOC_Os03g56370.1 | Bradi1g07120.1 | HORVU5Hr1G099020.1 | TRIUR3_00982 | Seita.9G072600.1 | Sobic.001G073300.1 | Zm00001d034249_T001 |
| orthologID_008039 | PH01001010G0220 | LOC_Os03g17800.1 | Bradi1g65767.2 | HORVU4Hr1G056950.4 | TRIUR3_32537 | Seita.9G443400.1 | Sobic.001G409300.1 | Zm00001d028720_T001 |
| orthologID_007965 | PH01000081G0810 | LOC_Os03g06370.2 | Bradi1g74500.1 | HORVU4Hr1G073140.4 | TRIUR3_21933 | Seita.9G533400.1 | Sobic.001G497600.1 | Zm00001d027741_T001 |
| orthologID_009901 | PH01000904G0290 | LOC_Os10g36420.1 | Bradi3g30410.1 | HORVU1Hr1G046740.5 | TRIUR3_27479 | Seita.9G197300.1 | Sobic.001G199200.1 | Zm00001d032502_T001 |
| orthologID_009459 | PH01000305G0300 | LOC_Os10g34010.1 | Bradi3g29000.1 | HORVU1Hr1G041960.1 | TRIUR3_31884 | Seita.9G219300.1 | Sobic.001G216800.1 | Zm00001d032672_T001 |
| orthologID_008358 | PH01000157G0500 | LOC_Os03g05280.1 | Bradi1g75340.1 | HORVU4Hr1G079250.1 | TRIUR3_17140 | Seita.9G541400.1 | Sobic.001G505200.1 | Zm00001d048393_T001 |
| orthologID_007633 | PH01000572G0680 | LOC_Os03g37130.1 | Bradi1g15730.1 | HORVU6Hr1G020720.5 | TRIUR3_10925 | Seita.9G183900.1 | Sobic.001G186800.1 | Zm00001d033056_T001 |
| orthologID_007701 | PH01000750G0220 | LOC_Os03g52730.1 | Bradi1g09369.1 | HORVU4Hr1G011320.1 | TRIUR3_13656 | Seita.9G100600.1 | Sobic.001G098200.1 | Zm00001d013317_T001 |
| orthologID_009395 | PH01000321G0540 | LOC_Os10g37830.1 | Bradi3g31520.1 | HORVU1Hr1G048700.4 | TRIUR3_28016 | Seita.9G349000.1 | Sobic.001G321500.1 | Zm00001d029673_T001 |
| orthologID_007661 | PH01000013G2450 | LOC_Os03g19580.1 | Bradi1g64370.1 | HORVU4Hr1G053480.1 | TRIUR3_11959 | Seita.9G428200.1 | Sobic.001G394600.1 | Zm00001d028903_T002 |
| orthologID_008012 | PH01000408G0040 | LOC_Os03g10590.1 | Bradi1g70960.3 | HORVU4Hr1G070180.9 | TRIUR3_02490 | Seita.9G500500.1 | Sobic.001G465300.1 | Zm00001d028097_T001 |
| orthologID_007671 | PH01000013G1830 | LOC_Os03g19720.1 | Bradi1g64290.1 | HORVU4Hr1G053130.1 | TRIUR3_30700 | Seita.9G427100.1 | Sobic.001G393500.2 | Zm00001d047618_T001 |
| orthologID_008325 | PH01000723G0440 | LOC_Os03g13220.1 | Bradi1g68880.1 | HORVU4Hr1G064860.1 | TRIUR3_14676 | Seita.9G477600.1 | Sobic.001G444300.1 | Zm00001d047983_T002 |
| orthologID_007891 | PH01000759G0700 | LOC_Os03g03020.1 | Bradi1g77060.1 | HORVU4Hr1G084830.2 | TRIUR3_18631 | Seita.9G561900.1 | Sobic.001G527100.1 | Zm00001d027421_T001 |
| orthologID_007577 | PH01000047G0020 | LOC_Os03g22680.1 | Bradi1g62490.1 | HORVU4Hr1G010980.12 | TRIUR3_23854 | Seita.9G405000.1 | Sobic.001G373100.1 | Zm00001d029141_T001 |
| orthologID_007749 | PH01001115G0700 | LOC_Os03g15920.1 | Bradi1g67120.1 | HORVU4Hr1G060870.1 | TRIUR3_17911 | Seita.9G457700.1 | Sobic.001G426400.1 | Zm00001d028548_T001 |
| orthologID_007715 | PH01001968G0140 | LOC_Os03g50440.1 | Bradi1g11410.1 | HORVU4Hr1G007560.1 | TRIUR3_10757 | Seita.9G118700.1 | Sobic.001G116700.1 | Zm00001d033744_T001 |
| orthologID_007820 | PH01002828G0120 | LOC_Os03g51350.1 | Bradi1g10310.1 | HORVU4Hr1G008990.1 | TRIUR3_31296 | Seita.9G112100.1 | Sobic.001G109000.1 | Zm00001d013384_T001 |
| orthologID_007849 | PH01000390G0450 | LOC_Os03g04300.3 | Bradi1g76040.1 | HORVU4Hr1G080940.2 | TRIUR3_19773 | Seita.9G548900.1 | Sobic.001G513800.1 | Zm00001d027544_T003 |
| orthologID_009759 | PH01177560G0010 | LOC_Os10g33874.1 | Bradi3g28890.2 | HORVU1Hr1G041650.24 | TRIUR3_18971 | Seita.9G220300.1 | Sobic.001G217900.1 | Zm00001d032689_T005 |
| orthologID_009333 | PH01001683G0170 | LOC_Os10g42130.1 | Bradi3g33957.2 | HORVU1Hr1G049840.9 | TRIUR3_22131 | Seita.9G314000.1 | Sobic.001G290400.1 | Zm00001d030023_T001 |
| orthologID_007917 | PH01001459G0340 | LOC_Os03g21370.1 | Bradi1g63490.1 | HORVU4Hr1G050980.1 | TRIUR3_23917 | Seita.9G415900.1 | Sobic.001G383900.1 | Zm00001d029027_T001 |
| orthologID_009602 | PH01000808G0320 | LOC_Os10g04580.1 | Bradi3g21180.1 | HORVU1Hr1G045310.7 | TRIUR3_12671 | Seita.9G292600.1 | Sobic.001G270800.1 | Zm00001d014330_T014 |
| orthologID_007756 | PH01001698G0420 | LOC_Os03g21810.1 | Bradi1g63160.1 | HORVU4Hr1G050160.2 | TRIUR3_28348 | Seita.9G412600.1 | Sobic.001G380600.1 | Zm00001d047518_T004 |
| orthologID_008643 | PH01000476G0030 | LOC_Os09g07660.2 | Bradi1g02770.1 | HORVU5Hr1G051870.7 | TRIUR3_21765 | Seita.2G153300.1 | Sobic.001G280100.1 | Zm00001d014446_T001 |
| orthologID_007767 | PH01000107G1720 | LOC_Os03g03450.1 | Bradi1g76800.1 | HORVU4Hr1G083210.3 | TRIUR3_06236 | Seita.9G558400.1 | Sobic.001G523800.1 | Zm00001d048461_T001 |
| orthologID_008720 | PH01087563G0010 | LOC_Os03g60910.1 | Bradi1g03450.1 | HORVU5Hr1G114630.1 | TRIUR3_12154 | Seita.9G030000.1 | Sobic.001G031300.1 | Zm00001d022599_T002 |
| orthologID_008578 | PH01000251G0780 | LOC_Os03g10750.1 | Bradi1g70890.1 | HORVU4Hr1G069920.28 | TRIUR3_01460 | Seita.9G499500.1 | Sobic.001G464500.1 | Zm00001d028088_T003 |
| orthologID_008091 | PH01003316G0140 | LOC_Os03g29540.2 | Bradi1g60130.1 | HORVU6Hr1G013240.3 | TRIUR3_30058 | Seita.9G368900.1 | Sobic.001G341300.2 | Zm00001d029475_T001 |
| orthologID_009436 | PH01000520G0320 | LOC_Os10g35670.1 | Bradi3g30040.1 | HORVU1Hr1G045110.3 | TRIUR3_19451 | Seita.9G204200.1 | Sobic.001G204300.1 | Zm00001d032550_T001 |
| orthologID_007305 | PH01000809G0320 | LOC_Os12g17070.1 | Bradi4g39227.2 | HORVU1Hr1G083800.3 | TRIUR3_03667 | Seita.3G298600.1 | Sobic.001G527600.1 | Zm00001d027416_T001 |
| orthologID_008362 | PH01000340G0420 | LOC_Os03g46310.1 | Bradi1g12940.1 | HORVU4Hr1G002890.1 | TRIUR3_11654 | Seita.9G147000.1 | Sobic.001G143900.3 | Zm00001d034956_T001 |
| orthologID_008048 | PH01001917G0130 | LOC_Os03g46660.1 | Bradi1g12810.1 | HORVU4Hr1G003090.2 | TRIUR3_05547 | Seita.9G145500.1 | Sobic.001G142200.1 | Zm00001d013573_T001 |
| orthologID_008465 | PH01000122G1730 | LOC_Os03g60350.1 | Bradi1g03940.1 | HORVU5Hr1G112300.1 | TRIUR3_18542 | Seita.9G036900.1 | Sobic.001G038200.1 | Zm00001d034620_T001 |
| orthologID_008233 | PH01000113G0330 | LOC_Os03g07370.1 | Bradi1g73700.1 | HORVU4Hr1G074980.1 | TRIUR3_08746 | Seita.9G524900.1 | Sobic.001G489800.2 | Zm00001d027847_T001 |
| orthologID_007951 | PH01000187G0610 | LOC_Os03g29850.1 | Bradi1g60110.1 | HORVU6Hr1G018070.12 | TRIUR3_22023 | Seita.9G366600.1 | Sobic.001G339600.1 | Zm00001d029497_T001 |

| | | | | | | | | |
|---|---|---|---|---|---|---|---|---|
| orthologID_007707 | PH01003159G0110 | LOC_Os03g43420.1 | Bradi1g14220.1 | HORVU5Hr1G093650.4 | TRIUR3_01369 | Seita.9G162700.1 | Sobic.001G161400.1 | Zm00001d033318_T002 |
| orthologID_008582 | PH01001979G0270 | LOC_Os03g20010.1 | Bradi1g64160.2 | HORVU4Hr1G052610.2 | TRIUR3_34937 | Seita.9G425600.1 | Sobic.001G391900.1 | Zm00001d028924_T001 |
| orthologID_008102 | PH01000138G1290 | LOC_Os03g16730.1 | Bradi1g66670.2 | HORVU4Hr1G059740.2 | TRIUR3_18953 | Seita.9G453200.1 | Sobic.001G421400.1 | Zm00001d028612_T001 |
| orthologID_008112 | PH01000532G0650 | LOC_Os03g14669.2 | Bradi1g67980.2 | HORVU4Hr1G062890.1 | TRIUR3_09240 | Seita.9G468100.1 | Sobic.001G435500.1 | Zm00001d028432_T001 |
| orthologID_009849 | PH01001507G0510 | LOC_Os10g36190.1 | Bradi3g30330.1 | HORVU1Hr1G046560.2 | TRIUR3_21392 | Seita.9G198700.1 | Sobic.001G200600.1 | Zm00001d032518_T001 |
| orthologID_007728 | PH01000716G0110 | LOC_Os03g54770.1 | Bradi1g08294.1 | HORVU5Hr1G095960.3 | TRIUR3_30915 | Seita.9G088200.1 | Sobic.001G085800.1 | Zm00001d034054_T002 |
| orthologID_007622 | PH01000458G0200 | LOC_Os03g57660.1 | Bradi1g06230.2 | HORVU5Hr1G103900.13 | TRIUR3_10713 | Seita.9G061800.1 | Sobic.001G062900.1 | Zm00001d034383_T003 |
| orthologID_007964 | PH01000134G1270 | LOC_Os03g50070.1 | Bradi1g11150.4 | HORVU4Hr1G007090.1 | TRIUR3_33118 | Seita.9G122000.1 | Sobic.001G120100.1 | Zm00001d013456_T001 |
| orthologID_008693 | PH01000091G0990 | LOC_Os03g26990.1 | Bradi1g60860.1 | HORVU3Hr1G003880.1 | TRIUR3_13239 | Seita.9G381000.1 | Sobic.001G352700.2 | Zm00001d047202_T001 |
| orthologID_006607 | PH01003125G0040 | LOC_Os02g18940.1 | Bradi2g40070.1 | HORVU1Hr1G000110.2 | TRIUR3_00703 | Seita.1G130200.1 | Sobic.001G530900.1 | Zm00001d053290_T001 |
| orthologID_007620 | PH01000376G0870 | LOC_Os03g56610.1 | Bradi1g06930.1 | HORVU5Hr1G099550.1 | TRIUR3_33709 | Seita.9G069500.1 | Sobic.001G070800.1 | Zm00001d034279_T001 |
| orthologID_007698 | PH01000113G1370 | LOC_Os03g07880.1 | Bradi1g72960.2 | HORVU4Hr1G075830.4 | TRIUR3_08409 | Seita.9G521100.1 | Sobic.001G486000.1 | Zm00001d027874_T004 |
| orthologID_007992 | PH01001463G0090 | LOC_Os03g51880.1 | Bradi1g09980.1 | HORVU4Hr1G009950.1 | TRIUR3_01285 | Seita.9G108400.1 | Sobic.001G105300.1 | Zm00001d013359_T002 |
| orthologID_008441 | PH01000707G0390 | LOC_Os03g63360.1 | Bradi1g01660.1 | HORVU5Hr1G121730.1 | TRIUR3_25087 | Seita.9G010100.1 | Sobic.001G011600.1 | Zm00001d034852_T001 |
| orthologID_008370 | PH01000648G0190 | LOC_Os03g17940.1 | Bradi1g65657.1 | HORVU4Hr1G056660.2 | TRIUR3_32905 | Seita.9G442300.1 | Sobic.001G408400.1 | Zm00001d028731_T001 |
| orthologID_009193 | PH01000194G0820 | LOC_Os10g28320.1 | Bradi3g26900.2 | HORVU7Hr1G039930.1 | TRIUR3_28126 | Seita.9G247500.1 | Sobic.001G242800.1 | Zm00001d032921_T001 |
| orthologID_009213 | PH01000397G0760 | LOC_Os10g40650.1 | Bradi3g33090.2 | HORVU1Hr1G054110.3 | TRIUR3_16743 | Seita.9G327400.1 | Sobic.001G302100.1 | Zm00001d029919_T001 |
| orthologID_008396 | PH01000091G1730 | LOC_Os03g27120.1 | Bradi1g60800.1 | HORVU3Hr1G095700.1 | TRIUR3_27784 | Seita.9G379900.1 | Sobic.001G351500.1 | Zm00001d029387_T001 |
| orthologID_008299 | PH01001453G0370 | LOC_Os03g44430.1 | Bradi1g13750.1 | HORVU5Hr1G092880.1 | TRIUR3_20364 | Seita.9G156600.1 | Sobic.001G155200.2 | Zm00001d033221_T001 |
| orthologID_007788 | PH01000157G0460 | LOC_Os03g05290.1 | Bradi1g75290.1 | HORVU4Hr1G079230.1 | TRIUR3_17138 | Seita.9G541300.1 | Sobic.001G505100.1 | Zm00001d027652_T001 |
| orthologID_008386 | PH01001584G0370 | LOC_Os03g47990.1 | Bradi1g12312.1 | HORVU4Hr1G004130.3 | TRIUR3_04904 | Seita.9G138200.1 | Sobic.001G135200.1 | Zm00001d033526_T001 |
| orthologID_009500 | PH01000362G0480 | LOC_Os10g37660.1 | Bradi3g31410.1 | HORVU1Hr1G048430.3 | TRIUR3_33217 | Seita.9G350600.1 | Sobic.001G322500.2 | Zm00001d029654_T003 |
| orthologID_008099 | PH01000007G3680 | LOC_Os03g12160.1 | Bradi1g69590.1 | HORVU4Hr1G067100.2 | TRIUR3_19918 | Seita.9G486700.1 | Sobic.001G452600.1 | Zm00001d028244_T010 |
| orthologID_008256 | PH01002189G0030 | LOC_Os03g57140.1 | Bradi1g06511.1 | HORVU5Hr1G103320.2 | TRIUR3_01390 | Seita.9G065000.1 | Sobic.001G066700.1 | Zm00001d034346_T002 |
| orthologID_009479 | PH01000048G0430 | LOC_Os09g02810.1 | Bradi3g47770.1 | HORVU6Hr1G053890.9 | TRIUR3_22590 | Seita.2G178000.1 | Sobic.001G453400.2 | Zm00001d028238_T001 |
| orthologID_006567 | PH01000804G0030 | LOC_Os03g60939.1 | Bradi2g19800.2 | HORVU5Hr1G081360.11 | TRIUR3_02001 | Seita.9G029800.1 | Sobic.001G031100.1 | Zm00001d034682_T001 |
| orthologID_009158 | PH01000873G0610 | LOC_Os08g14770.1 | Bradi3g18970.2 | HORVU6Hr1G033090.1 | TRIUR3_16844 | Seita.6G093600.1 | Sobic.001G465700.1 | Zm00001d024159_T002 |
| orthologID_008640 | PH01001260G0170 | LOC_Os03g56850.1 | Bradi1g06766.2 | HORVU5Hr1G101000.3 | TRIUR3_05372 | Seita.9G068200.1 | Sobic.001G069300.2 | Zm00001d013136_T001 |
| orthologID_008153 | PH01000115G0020 | LOC_Os03g08320.1 | Bradi1g72600.1 | HORVU4Hr1G076840.5 | TRIUR3_06120 | Seita.9G518000.1 | Sobic.001G482700.1 | Zm00001d027900_T001 |
| orthologID_007993 | PH01005158G0070 | LOC_Os03g53020.1 | Bradi1g09177.1 | HORVU5Hr1G093960.16 | TRIUR3_04618 | Seita.9G098200.1 | Sobic.001G095700.1 | Zm00001d033957_T002 |
| orthologID_008562 | PH01000322G0510 | LOC_Os03g18454.1 | Bradi1g65246.3 | HORVU4Hr1G055720.2 | TRIUR3_04864 | Seita.9G438700.1 | Sobic.001G404500.1 | Zm00001d028775_T001 |
| orthologID_007876 | PH01000716G0750 | LOC_Os03g55140.1 | Bradi1g07991.1 | HORVU5Hr1G096830.6 | TRIUR3_10260 | Seita.9G084900.1 | Sobic.001G083300.5 | Zm00001d034081_T002 |
| orthologID_009107 | PH01000377G0450 | LOC_Os10g39870.1 | Bradi3g32560.1 | HORVU1Hr1G052930.1 | TRIUR3_04219 | Seita.9G336900.1 | Sobic.001G308700.1 | Zm00001d047397_T001 |
| orthologID_009706 | PH01002340G0150 | LOC_Os10g42630.1 | Bradi3g34290.2 | HORVU6Hr1G058250.1 | TRIUR3_21997 | Seita.9G308600.1 | Sobic.001G285300.1 | Zm00001d030091_T001 |
| orthologID_008611 | PH01000786G0600 | LOC_Os03g15440.1 | Bradi1g67500.1 | HORVU4Hr1G061760.1 | TRIUR3_31897 | Seita.9G454200.1 | Sobic.001G430000.1 | Zm00001d028504_T001 |
| orthologID_008081 | PH01000767G0370 | LOC_Os03g57270.1 | Bradi1g06400.1 | HORVU5Hr1G103490.6 | TRIUR3_18516 | Seita.9G063800.1 | Sobic.001G065200.1 | Zm00001d013117_T001 |
| orthologID_007631 | PH01000376G0290 | LOC_Os03g56310.1 | Bradi1g07140.1 | HORVU5Hr1G098980.12 | TRIUR3_31235 | Seita.9G072800.1 | Sobic.001G073500.1 | Zm00001d034245_T004 |
| orthologID_009720 | PH01002057G0210 | LOC_Os10g42170.1 | Bradi3g33967.1 | HORVU1Hr1G049810.23 | TRIUR3_14473 | Seita.9G313900.1 | Sobic.001G290300.2 | Zm00001d047046_T003 |
| orthologID_009351 | PH01004295G0070 | LOC_Os10g42550.1 | Bradi3g34350.1 | HORVU1Hr1G050760.1 | TRIUR3_14767 | Seita.9G309100.1 | Sobic.001G286100.1 | Zm00001d030083_T001 |
| orthologID_007973 | PH01000113G0080 | LOC_Os03g07234.1 | Bradi1g73802.2 | HORVU4Hr1G074830.1 | TRIUR3_05043 | Seita.9G526300.1 | Sobic.001G491500.1 | Zm00001d048332_T001 |
| orthologID_008275 | PH01000370G0180 | LOC_Os03g03360.1 | Bradi1g76880.1 | HORVU4Hr1G084410.1 | TRIUR3_10657 | Seita.9G559300.1 | Sobic.001G524900.1 | Zm00001d027442_T001 |
| orthologID_007999 | PH01001577G0310 | LOC_Os03g01570.1 | Bradi1g78180.1 | HORVU4Hr1G087350.3 | TRIUR3_04506 | Seita.9G576300.1 | Sobic.001G540600.2 | Zm00001d027298_T001 |
| orthologID_007971 | PH01003704G0130 | LOC_Os03g59700.1 | Bradi1g04250.1 | HORVU5Hr1G111410.1 | TRIUR3_04348 | Seita.9G040900.1 | Sobic.001G041200.1 | Zm00001d013010_T002 |
| orthologID_008583 | PH01000365G0930 | LOC_Os03g63020.1 | Bradi1g01840.1 | HORVU5Hr1G120960.1 | TRIUR3_19424 | Seita.9G012900.1 | Sobic.001G014200.5 | Zm00001d034834_T009 |
| orthologID_008565 | PH01000040G1760 | LOC_Os03g61260.1 | Bradi1g03220.1 | HORVU5Hr1G115340.1 | TRIUR3_10860 | Seita.9G022000.1 | Sobic.001G027600.1 | Zm00001d034724_T001 |
| orthologID_007804 | PH01000081G0220 | LOC_Os03g06580.4 | Bradi1g74400.1 | HORVU4Hr1G073580.2 | TRIUR3_31147 | Seita.9G532200.1 | Sobic.001G496500.1 | Zm00001d027755_T002 |
| orthologID_008207 | PH01001681G0160 | LOC_Os03g46390.1 | Bradi1g12890.1 | HORVU4Hr1G002980.2 | TRIUR3_17632 | Seita.9G146600.1 | Sobic.001G143300.1 | Zm00001d013583_T001 |
| orthologID_007787 | PH01002919G0240 | LOC_Os03g51760.1 | Bradi1g10017.1 | HORVU4Hr1G009880.1 | TRIUR3_01291 | Seita.9G109000.1 | Sobic.001G105800.1 | Zm00001d033863_T001 |
| orthologID_007887 | PH01001115G0450 | LOC_Os03g16080.1 | Bradi1g66990.1 | HORVU4Hr1G060540.1 | TRIUR3_20196 | Seita.9G459100.1 | Sobic.001G425150.1 | Zm00001d047838_T001 |
| orthologID_008717 | PH01000113G0190 | LOC_Os03g07340.1 | Bradi1g73720.1 | HORVU4Hr1G075010.10 | TRIUR3_10819 | Seita.9G525300.1 | Sobic.001G490600.1 | Zm00001d027844_T001 |

| | | | | | | | | |
|---|---|---|---|---|---|---|---|---|
| orthologID_007603 | PH01000174G0650 | LOC_Os03g58120.1 | Bradi1g05590.2 | HORVU5Hr1G105140.4 | TRIUR3_15383 | Seita.9G056900.1 | Sobic.001G058200.3 | Zm00001d034428_T001 |
| orthologID_008272 | PH01000312G1160 | LOC_Os03g07220.1 | Bradi1g73810.3 | HORVU4Hr1G074790.8 | TRIUR3_05046 | Seita.9G526500.1 | Sobic.001G491700.1 | Zm00001d027835_T001 |
| orthologID_008105 | PH01000040G0140 | LOC_Os03g62640.2 | Bradi1g02020.1 | HORVU5Hr1G120260.3 | TRIUR3_00923 | Seita.9G016500.1 | Sobic.001G016900.1 | Zm00001d012865_T004 |
| orthologID_008552 | PH01000495G0640 | LOC_Os03g35340.1 | Bradi1g41947.5 | HORVU7Hr1G083160.12 | TRIUR3_23097 | Seita.9G188600.1 | Sobic.001G191700.1 | Zm00001d013829_T001 |
| orthologID_007677 | PH01000376G0170 | LOC_Os03g56260.1 | Bradi1g07190.1 | HORVU5Hr1G098780.4 | TRIUR3_32456 | Seita.9G073300.1 | Sobic.001G074100.1 | Zm00001d034239_T001 |
| orthologID_009850 | PH01002033G0280 | LOC_Os10g38920.1 | Bradi3g31967.3 | HORVU1Hr1G049470.21 | TRIUR3_22865 | Seita.9G344300.1 | Sobic.001G316000.1 | Zm00001d029721_T001 |
| orthologID_008593 | PH01001672G0380 | LOC_Os07g49530.1 | Bradi1g16355.1 | HORVU2Hr1G013090.1 | TRIUR3_12694 | Seita.2G444900.1 | Sobic.001G190000.1 | Zm00001d013827_T001 |
| orthologID_008063 | PH01000040G0280 | LOC_Os03g62490.1 | Bradi1g02170.1 | HORVU5Hr1G119700.1 | TRIUR3_11415 | Seita.9G017700.1 | Sobic.001G018200.1 | Zm00001d034795_T002 |
| orthologID_007690 | PH01000122G1350 | LOC_Os03g60190.1 | Bradi1g04030.1 | HORVU5Hr1G111850.1 | TRIUR3_05774 | Seita.9G037700.1 | Sobic.001G039100.1 | Zm00001d012991_T001 |
| orthologID_008109 | PH01000376G0520 | LOC_Os03g56430.1 | Bradi1g07060.1 | HORVU5Hr1G099180.4 | TRIUR3_00989 | Seita.9G071900.1 | Sobic.001G072600.1 | Zm00001d013154_T001 |
| orthologID_008270 | PH01000122G0570 | LOC_Os03g59650.1 | Bradi1g04292.1 | HORVU5Hr1G111300.1 | TRIUR3_19335 | Seita.9G041500.1 | Sobic.001G041700.3 | Zm00001d034594_T006 |
| orthologID_009284 | PH01000377G0960 | LOC_Os10g40130.1 | Bradi3g32820.3 | HORVU1Hr1G053310.2 | TRIUR3_09767 | Seita.9G333200.1 | Sobic.001G305700.1 | Zm00001d047128_T002 |
| orthologID_009801 | PH01001233G0460 | LOC_Os10g41510.1 | Bradi3g33700.1 | HORVU1Hr1G055300.1 | TRIUR3_14876 | Seita.9G318100.1 | Sobic.001G294300.1 | Zm00001d029976_T006 |
| orthologID_008204 | PH01000371G0270 | LOC_Os03g63450.1 | Bradi1g01500.1 | HORVU2Hr1G016330.1 | TRIUR3_14900 | Seita.9G008700.1 | Sobic.001G010400.1 | Zm00001d034859_T001 |
| orthologID_008061 | PH01000408G0640 | LOC_Os03g10850.1 | Bradi1g70790.1 | HORVU4Hr1G069660.11 | TRIUR3_16682 | Seita.9G498300.1 | Sobic.001G463500.1 | Zm00001d028071_T001 |
| orthologID_008607 | PH01001003G0740 | LOC_Os03g15890.1 | Bradi1g67145.1 | HORVU4Hr1G060940.5 | TRIUR3_23067 | Seita.9G457400.1 | Sobic.001G426800.1 | Zm00001d047847_T003 |
| orthologID_009366 | PH01000171G1240 | LOC_Os10g41180.1 | Bradi3g33377.1 | HORVU1Hr1G056000.4 | TRIUR3_06497 | Seita.9G322600.1 | Sobic.001G043200.1 | Zm00001d029878_T001 |
| orthologID_007902 | PH01000353G0450 | LOC_Os03g13550.1 | Bradi1g68660.2 | HORVU4Hr1G064300.1 | TRIUR3_15315 | Seita.9G475000.1 | Sobic.001G441900.2 | Zm00001d047971_T002 |
| orthologID_007863 | PH01002425G0300 | LOC_Os03g50430.1 | Bradi1g11397.1 | HORVU4Hr1G007570.1 | TRIUR3_10758 | Seita.9G118800.1 | Sobic.001G116800.1 | Zm00001d013432_T001 |
| orthologID_007589 | PH01000797G0560 | LOC_Os03g50730.1 | Bradi1g10710.1 | HORVU4Hr1G008130.10 | TRIUR3_25218 | Seita.9G116200.1 | Sobic.001G114200.1 | Zm00001d013421_T002 |
| orthologID_008268 | PH01000371G0550 | LOC_Os03g63550.1 | Bradi1g01320.1 | HORVU5Hr1G123210.2 | TRIUR3_07633 | Seita.9G007400.1 | Sobic.001G008800.1 | Zm00001d012833_T002 |
| orthologID_007771 | PH01001686G0260 | LOC_Os03g26450.1 | Bradi1g61050.1 | HORVU4Hr1G038670.1 | TRIUR3_30906 | Seita.9G383900.1 | Sobic.001G355100.1 | Zm00001d029343_T002 |
| orthologID_009690 | PH01003938G0210 | LOC_Os06g23160.1 | Bradi3g31760.1 | HORVU7Hr1G081880.11 | TRIUR3_24968 | Seita.4G159900.1 | Sobic.001G168900.1 | Zm00001d033345_T007 |
| orthologID_007924 | PH01002343G0290 | LOC_Os03g27950.1 | Bradi1g60450.3 | HORVU5Hr1G096520.1 | TRIUR3_20742 | Seita.9G374500.1 | Sobic.001G346700.2 | Zm00001d047241_T003 |
| orthologID_008218 | PH01001293G0010 | LOC_Os03g46300.1 | Bradi1g12950.1 | HORVU4Hr1G002850.1 | TRIUR3_11655 | Seita.9G147200.1 | Sobic.001G144100.1 | Zm00001d013593_T001 |
| orthologID_008354 | PH01000038G2120 | LOC_Os03g08960.1 | Bradi1g71930.1 | HORVU4Hr1G078410.2 | TRIUR3_26324 | Seita.9G511500.1 | Sobic.001G476300.1 | Zm00001d027991_T001 |
| orthologID_008258 | PH01008675G0010 | LOC_Os03g22450.1 | Bradi1g62700.1 | HORVU4Hr1G046540.6 | TRIUR3_09242 | Seita.9G407200.1 | Sobic.001G374800.1 | Zm00001d029123_T005 |
| orthologID_007800 | PH01000106G1000 | LOC_Os03g11340.1 | Bradi1g70280.1 | HORVU4Hr1G068770.5 | TRIUR3_03801 | Seita.9G494100.1 | Sobic.001G459900.1 | Zm00001d028189_T001 |
| orthologID_008679 | PH01000497G0620 | LOC_Os03g53070.1 | Bradi1g09150.1 | HORVU0Hr1G013770.1 | TRIUR3_35271 | Seita.9G097800.1 | Sobic.001G095300.1 | Zm00001d033966_T001 |
| orthologID_008737 | PH01001161G0430 | LOC_Os03g53640.1 | Bradi1g08718.1 | HORVU5Hr1G094770.2 | TRIUR3_17919 | Seita.9G093400.1 | Sobic.001G090700.1 | Zm00001d013277_T001 |
| orthologID_009182 | PH01000520G0330 | LOC_Os10g35680.1 | Bradi3g30047.2 | HORVU1Hr1G045100.2 | TRIUR3_19450 | Seita.9G204100.1 | Sobic.001G204200.1 | Zm00001d032556_T001 |
| orthologID_007721 | PH01000882G0420 | LOC_Os03g30830.1 | Bradi1g16230.3 | HORVU6Hr1G019780.2 | TRIUR3_30528 | Seita.9G363400.1 | Sobic.001G334900.1 | Zm00001d047293_T001 |
| orthologID_009616 | PH01001306G0080 | LOC_Os10g30944.2 | Bradi3g27760.1 | HORVU1Hr1G037730.4 | TRIUR3_17761 | Seita.9G236500.1 | Sobic.001G233600.1 | Zm00001d032832_T001 |
| orthologID_009825 | PH01004295G0100 | LOC_Os02g13200.1 | Bradi3g34310.1 | HORVU1Hr1G050810.1 | TRIUR3_34788 | Seita.9G308800.1 | Sobic.001G285800.1 | Zm00001d030087_T001 |
| orthologID_007590 | PH01003838G0050 | LOC_Os03g01800.1 | Bradi1g77990.1 | HORVU4Hr1G090820.2 | TRIUR3_14063 | Seita.9G573900.1 | Sobic.001G538000.1 | Zm00001d027313_T001 |
| orthologID_008567 | PH01001115G0420 | LOC_Os03g16120.1 | Bradi1g66960.1 | HORVU4Hr1G060500.1 | TRIUR3_34296 | Seita.9G459700.1 | Sobic.001G424800.1 | Zm00001d047834_T004 |
| orthologID_009854 | PH01150229G0010 | LOC_Os10g01100.1 | Bradi3g21580.1 | HORVU4Hr1G010030.27 | TRIUR3_07898 | Seita.9G303400.1 | Sobic.001G280300.1 | Zm00001d030171_T001 |
| orthologID_009438 | PH01000186G1110 | LOC_Os10g34390.1 | Bradi3g29260.3 | HORVU1Hr1G042560.3 | TRIUR3_14833 | Seita.9G215800.1 | Sobic.001G214600.1 | Zm00001d013985_T004 |
| orthologID_007658 | PH01001528G0160 | LOC_Os03g60700.1 | Bradi1g03660.1 | HORVU5Hr1G113310.2 | TRIUR3_07273 | Seita.9G032600.1 | Sobic.001G033500.1 | Zm00001d034659_T001 |
| orthologID_009172 | PH01000059G0270 | LOC_Os10g38820.1 | Bradi3g31890.1 | HORVU1Hr1G049310.1 | TRIUR3_15472 | Seita.9G345200.1 | Sobic.001G316900.2 | Zm00001d029711_T001 |
| orthologID_008254 | PH01001597G0370 | LOC_Os03g48930.1 | Bradi1g11830.1 | HORVU0Hr1G040340.1 | TRIUR3_09922 | Seita.9G129900.1 | Sobic.001G128400.1 | Zm00001d033596_T001 |
| orthologID_007655 | PH01000408G0460 | LOC_Os03g10780.1 | Bradi1g70850.4 | HORVU4Hr1G069790.2 | TRIUR3_14056 | Seita.9G499200.1 | Sobic.001G464200.1 | Zm00001d028084_T002 |
| orthologID_009163 | PH01001233G0110 | LOC_Os10g41749.4 | Bradi3g33810.1 | HORVU1Hr1G054610.22 | TRIUR3_11704 | Seita.9G316100.1 | Sobic.001G292700.1 | Zm00001d047063_T003 |
| orthologID_008053 | PH01002002G0040 | LOC_Os03g59250.1 | Bradi1g04710.4 | HORVU5Hr1G109730.4 | TRIUR3_13613 | Seita.9G067900.1 | Sobic.001G047000.1 | Zm00001d034534_T001 |
| orthologID_007644 | PH01002041G0150 | LOC_Os03g59320.1 | Bradi1g04620.1 | HORVU5Hr1G109940.1 | TRIUR3_17719 | Seita.9G044500.1 | Sobic.001G046000.1 | Zm00001d013033_T001 |
| orthologID_008694 | PH01000122G0280 | LOC_Os03g59580.1 | Bradi1g04590.1 | HORVU5Hr1G110120.1 | TRIUR3_10940 | Seita.9G042200.1 | Sobic.001G042800.1 | Zm00001d034586_T001 |
| orthologID_008008 | PH01001260G0210 | LOC_Os03g56869.1 | Bradi1g06750.1 | HORVU5Hr1G101020.12 | TRIUR3_05373 | Seita.9G067800.1 | Sobic.001G069000.1 | Zm00001d013135_T002 |
| orthologID_008364 | PH01002124G0160 | LOC_Os03g49140.1 | Bradi1g11780.1 | HORVU4Hr1G005730.2 | TRIUR3_33699 | Seita.9G129000.1 | Sobic.001G127500.1 | Zm00001d013499_T002 |
| orthologID_007083 | PH01000027G1830 | LOC_Os10g17489.1 | Bradi4g15800.1 | HORVU4Hr1G063660.3 | TRIUR3_32757 | Seita.1G214300.1 | Sobic.001G044400.1 | Zm00001d034569_T001 |
| orthologID_008630 | PH01000353G1080 | LOC_Os03g13840.2 | Bradi1g68210.3 | HORVU4Hr1G063980.6 | TRIUR3_32479 | Seita.9G473100.1 | Sobic.001G440200.1 | Zm00001d047951_T002 |

| | | | | | | | | |
|---|---|---|---|---|---|---|---|---|
| orthologID_008663 | PH01000017G0390 | LOC_Os03g02860.1 | Bradi1g77170.1 | HORVU4Hr1G085090.1 | TRIUR3_09931 | Seita.9G562900.1 | Sobic.001G528300.1 | Zm00001d027405_T001 |
| orthologID_007560 | PH01001119G0270 | LOC_Os03g58390.1 | Bradi1g05390.2 | HORVU5Hr1G106470.4 | TRIUR3_25105 | Seita.9G054100.1 | Sobic.001G055900.1 | Zm00001d034468_T001 |
| orthologID_008610 | PH01000017G1970 | LOC_Os03g02070.1 | Bradi1g77800.1 | HORVU4Hr1G089520.1 | TRIUR3_08669 | Seita.9G570800.1 | Sobic.001G535200.1 | Zm00001d048513_T001 |
| orthologID_007623 | PH01000365G0780 | LOC_Os03g62750.1 | Bradi1g01900.1 | HORVU5Hr1G120570.1 | TRIUR3_02858 | Seita.9G014200.1 | Sobic.001G015500.1 | Zm00001d012861_T001 |
| orthologID_010045 | PH01002850G0170 | LOC_Os03g53200.1 | Bradi1g09042.1 | HORVU5Hr1G094290.9 | TRIUR3_13818 | Seita.9G097000.1 | Sobic.001G094400.1 | Zm00001d033980_T001 |
| orthologID_009173 | PH01000226G0670 | LOC_Os10g35070.1 | Bradi3g29797.1 | HORVU1Hr1G043830.1 | TRIUR3_15853 | Seita.9G208200.1 | Sobic.001G208200.1 | Zm00001d032608_T001 |
| orthologID_008436 | PH01000705G0730 | LOC_Os03g05710.1 | Bradi1g74950.1 | HORVU4Hr1G083470.3 | TRIUR3_01163 | Seita.9G537900.1 | Sobic.001G501600.1 | Zm00001d027703_T001 |
| orthologID_007688 | PH01000537G0750 | LOC_Os03g15230.1 | Bradi1g67640.1 | HORVU4Hr1G062110.1 | TRIUR3_33978 | Seita.9G463700.1 | Sobic.001G431800.1 | Zm00001d028456_T001 |
| orthologID_008179 | PH01000038G0790 | LOC_Os03g08450.1 | Bradi1g72470.1 | HORVU4Hr1G077250.1 | TRIUR3_26202 | Seita.9G516900.1 | Sobic.001G481700.1 | Zm00001d027922_T001 |
| orthologID_008501 | PH01001360G0010 | LOC_Os03g09060.1 | Bradi1g71860.1 | HORVU4Hr1G078510.1 | TRIUR3_30796 | Seita.9G510600.1 | Sobic.001G475600.1 | Zm00001d027999_T001 |
| orthologID_007982 | PH01003495G0070 | LOC_Os03g01720.1 | Bradi1g78063.2 | HORVU4Hr1G086680.2 | TRIUR3_01850 | Seita.9G574600.1 | Sobic.001G538932.1 | Zm00001d027308_T001 |
| orthologID_007888 | PH01004272G0030 | LOC_Os03g04210.1 | Bradi1g76100.1 | HORVU4Hr1G081190.2 | TRIUR3_07248 | Seita.9G550000.1 | Sobic.001G514500.2 | Zm00001d048352_T001 |
| orthologID_009831 | PH01000012G2690 | LOC_Os10g37230.1 | Bradi3g30470.1 | HORVU1Hr1G047000.1 | TRIUR3_19138 | Seita.5G011100.1 | Sobic.001G042500.1 | Zm00001d034588_T001 |
| orthologID_007794 | PH01001584G0170 | LOC_Os03g47610.6 | Bradi1g12550.3 | HORVU4Hr1G003580.22 | TRIUR3_02010 | Seita.9G140100.1 | Sobic.001G138000.1 | Zm00001d033383_T001 |
| orthologID_009090 | PH01000904G0460 | LOC_Os10g36490.1 | Bradi3g30790.2 | HORVU1Hr1G047690.1 | TRIUR3_14952 | Seita.9G197000.1 | Sobic.001G199000.1 | Zm00001d013891_T018 |
| orthologID_008271 | PH01000985G0560 | LOC_Os03g61730.1 | Bradi1g02800.1 | HORVU5Hr1G116930.5 | TRIUR3_21463 | Seita.9G023100.1 | Sobic.001G026200.1 | Zm00001d012913_T005 |
| orthologID_008659 | PH01001631G0530 | LOC_Os03g29960.1 | Bradi1g60040.1 | HORVU0Hr1G010260.4 | TRIUR3_16079 | Seita.9G365600.1 | Sobic.001G338800.1 | Zm00001d029504_T001 |
| orthologID_008234 | PH01001015G0560 | LOC_Os03g22270.1 | Bradi1g62860.1 | HORVU4Hr1G047240.2 | TRIUR3_32286 | Seita.9G409400.1 | Sobic.001G377000.1 | Zm00001d047503_T001 |
| orthologID_008433 | PH01001528G0150 | LOC_Os03g60690.1 | Bradi1g03670.4 | HORVU5Hr1G113330.1 | TRIUR3_07274 | Seita.9G032700.1 | Sobic.001G033600.2 | Zm00001d012961_T001 |
| orthologID_008537 | PH01000750G0690 | LOC_Os03g52450.1 | Bradi1g09550.1 | HORVU4Hr1G010840.2 | TRIUR3_23258 | Seita.9G103000.1 | Sobic.001G100100.1 | Zm00001d013331_T001 |
| orthologID_009318 | PH01000397G0060 | LOC_Os10g40280.1 | Bradi3g32860.2 | HORVU1Hr1G053390.1 | TRIUR3_15340 | Seita.9G333600.1 | Sobic.001G305200.1 | Zm00001d029851_T001 |
| orthologID_008540 | PH01001206G0320 | LOC_Os03g56110.1 | Bradi1g07247.4 | HORVU5Hr1G098570.6 | TRIUR3_30866 | Seita.9G074700.1 | Sobic.001G075101.1 | Zm00001d034212_T001 |
| orthologID_008011 | PH01004181G0030 | LOC_Os03g60120.1 | Bradi1g04110.1 | HORVU5Hr1G111550.1 | TRIUR3_12291 | Seita.9G038700.1 | Sobic.001G039800.1 | Zm00001d034605_T001 |
| orthologID_007746 | PH01000251G0610 | LOC_Os03g10790.1 | Bradi1g70840.1 | HORVU4Hr1G069740.2 | TRIUR3_14055 | Seita.9G499000.1 | Sobic.001G464100.1 | Zm00001d048140_T001 |
| orthologID_009217 | PH01000012G2650 | LOC_Os10g37210.1 | Bradi3g30480.1 | HORVU1Hr1G046990.10 | TRIUR3_19139 | Seita.9G192300.1 | Sobic.001G194900.2 | Zm00001d032455_T002 |
| orthologID_008569 | PH01000370G0900 | LOC_Os03g03630.1 | Bradi1g76610.1 | HORVU4Hr1G082900.2 | TRIUR3_24907 | Seita.9G556300.1 | Sobic.001G521300.1 | Zm00001d027477_T001 |
| orthologID_009699 | PH01002058G0250 | LOC_Os08g14490.1 | Bradi3g18840.2 | HORVU1Hr1G012730.2 | TRIUR3_15067 | Seita.6G091200.1 | Sobic.001G537100.1 | Zm00001d027319_T001 |
| orthologID_008618 | PH01000081G0800 | LOC_Os03g06390.1 | Bradi1g74490.1 | HORVU4Hr1G073230.1 | TRIUR3_34633 | Seita.9G533300.1 | Sobic.001G497500.1 | Zm00001d027742_T001 |
| orthologID_009683 | PH01000097G0400 | LOC_Os10g25430.1 | Bradi3g25370.1 | HORVU6Hr1G031370.7 | TRIUR3_32066 | Seita.9G253500.1 | Sobic.001G258200.1 | Zm00001d033040_T001 |
| orthologID_008178 | PH01001706G0010 | LOC_Os03g44660.1 | Bradi1g13590.1 | HORVU5Hr1G092490.1 | TRIUR3_10478 | Seita.9G154600.1 | Sobic.001G153300.2 | Zm00001d013664_T006 |
| orthologID_008669 | PH01000733G0110 | LOC_Os03g19670.1 | Bradi1g64327.1 | HORVU4Hr1G053190.2 | TRIUR3_29922 | Seita.9G427600.1 | Sobic.001G394000.1 | Zm00001d047620_T002 |
| orthologID_010088 | PH01000560G0790 | LOC_Os10g32870.1 | Bradi3g28270.1 | HORVU1Hr1G039030.1 | TRIUR3_13039 | Seita.9G229000.1 | Sobic.001G226100.2 | Zm00001d014079_T001 |
| orthologID_008292 | PH01000408G0830 | LOC_Os03g10880.1 | Bradi1g70761.1 | HORVU4Hr1G069480.3 | TRIUR3_19882 | Seita.9G497900.1 | Sobic.001G463200.1 | Zm00001d028056_T001 |
| orthologID_008575 | PH01004470G0050 | LOC_Os03g43580.1 | Bradi1g14090.1 | HORVU5Hr1G093430.4 | TRIUR3_32060 | Seita.9G161900.1 | Sobic.001G160100.1 | Zm00001d033280_T001 |
| orthologID_006588 | PH01002747G0220 | LOC_Os08g01120.1 | Bradi2g08130.1 | HORVU7Hr1G078700.1 | TRIUR3_10266 | Seita.9G184500.1 | Sobic.001G187300.1 | Zm00001d033053_T001 |
| orthologID_008123 | PH01004589G0090 | LOC_Os08g39420.1 | Bradi1g00860.1 | HORVU4Hr1G077000.2 | TRIUR3_05029 | Seita.9G003500.1 | Sobic.001G004200.1 | Zm00001d017656_T003 |
| orthologID_007662 | PH01000857G0610 | LOC_Os03g17920.1 | Bradi1g65680.2 | HORVU4Hr1G056670.41 | TRIUR3_11597 | Seita.9G442500.1 | Sobic.001G408600.1 | Zm00001d028728_T001 |
| orthologID_008006 | PH01000707G0040 | LOC_Os03g63290.1 | Bradi1g01780.1 | HORVU5Hr1G121430.1 | TRIUR3_05894 | Seita.9G011400.1 | Sobic.001G012100.1 | Zm00001d012849_T001 |
| orthologID_009557 | PH01000305G0740 | LOC_Os10g33920.1 | Bradi3g28920.2 | HORVU1Hr1G041720.2 | TRIUR3_33221 | Seita.9G220000.1 | Sobic.001G217500.1 | Zm00001d032686_T001 |
| orthologID_008203 | PH01000767G0590 | LOC_Os03g57410.1 | Bradi1g06330.1 | HORVU5Hr1G103740.2 | TRIUR3_03835 | Seita.9G063000.1 | Sobic.001G063900.1 | Zm00001d013110_T001 |
| orthologID_007716 | PH01000159G0760 | LOC_Os03g27930.1 | Bradi1g60460.3 | HORVU1Hr1G093790.4 | TRIUR3_17341 | Seita.9G374700.1 | Sobic.001G346900.1 | Zm00001d029429_T001 |
| orthologID_008019 | PH01004075G0030 | LOC_Os03g57640.1 | Bradi1g06241.1 | HORVU5Hr1G103770.1 | TRIUR3_10712 | Seita.9G062100.1 | Sobic.001G063000.1 | Zm00001d013099_T001 |
| orthologID_007700 | PH01000154G0800 | LOC_Os03g15850.1 | Bradi1g67190.3 | HORVU4Hr1G061050.1 | TRIUR3_23611 | Seita.9G457000.1 | Sobic.001G427200.1 | Zm00001d028540_T001 |
| orthologID_008134 | PH01001268G0390 | LOC_Os03g04545.1 | Bradi1g75850.1 | HORVU4Hr1G083600.1 | TRIUR3_17825 | Seita.9G546700.1 | Sobic.001G511300.3 | Zm00001d027588_T003 |
| orthologID_008624 | PH01002636G0200 | LOC_Os03g26210.1 | Bradi1g61110.1 | HORVU2Hr1G013450.3 | TRIUR3_24765 | Seita.9G384500.1 | Sobic.001G355600.1 | Zm00001d029339_T001 |
| orthologID_008319 | PH01000142G0520 | LOC_Os03g50130.1 | Bradi1g11220.1 | HORVU4Hr1G007180.4 | TRIUR3_05587 | Seita.9G121000.1 | Sobic.001G119400.1 | Zm00001d033707_T001 |
| orthologID_009412 | PH01000012G0490 | LOC_Os10g35870.1 | Bradi3g30180.1 | HORVU1Hr1G045630.2 | TRIUR3_32289 | Seita.9G201200.1 | Sobic.001G202400.1 | Zm00001d032531_T001 |
| orthologID_009802 | PH01000373G0400 | LOC_Os10g05720.1 | Bradi3g21070.1 | HORVU1Hr1G043430.1 | TRIUR3_23075 | Seita.9G287800.1 | Sobic.001G266500.1 | Zm00001d014315_T001 |
| orthologID_007795 | PH01000707G0010 | LOC_Os03g63280.1 | Bradi1g01760.1 | HORVU5Hr1G121600.4 | TRIUR3_17306 | Seita.9G011700.1 | Sobic.001G012700.1 | Zm00001d034844_T001 |
| orthologID_008284 | PH01000113G0180 | LOC_Os03g07330.1 | Bradi1g73730.1 | HORVU4Hr1G075000.4 | TRIUR3_10818 | Seita.9G525500.1 | Sobic.001G490700.1 | Zm00001d027843_T001 |

| | | | | | | | | |
|---|---|---|---|---|---|---|---|---|
| orthologID_009508 | PH01002394G0100 | LOC_Os10g32690.1 | Bradi3g28130.2 | HORVU1Hr1G038800.1 | TRIUR3_22885 | Seita.9G230700.1 | Sobic.001G227700.2 | Zm00001d032777_T001 |
| orthologID_007672 | PH01001056G0700 | LOC_Os03g03640.1 | Bradi1g76600.1 | HORVU4Hr1G082880.1 | TRIUR3_04852 | Seita.9G556200.1 | Sobic.001G521200.1 | Zm00001d027478_T001 |
| orthologID_008722 | PH01001696G0040 | LOC_Os03g55760.1 | Bradi1g07511.1 | HORVU5Hr1G097900.21 | TRIUR3_15041 | Seita.9G078300.1 | Sobic.001G077800.1 | Zm00001d013187_T001 |
| orthologID_008539 | PH01001527G0240 | LOC_Os03g58230.1 | Bradi1g05500.1 | HORVU5Hr1G105950.3 | TRIUR3_33883 | Seita.9G055800.1 | Sobic.001G057100.1 | Zm00001d034444_T002 |
| orthologID_007828 | PH01000108G0120 | LOC_Os03g41330.1 | Bradi1g14780.1 | HORVU4Hr1G013380.2 | TRIUR3_28321 | Seita.9G170000.1 | Sobic.001G169200.1 | Zm00001d033347_T001 |
| orthologID_006827 | PH01000275G0940 | LOC_Os10g41590.1 | Bradi2g33230.2 | HORVU3Hr1G099950.5 | TRIUR3_19205 | Seita.9G367100.1 | Sobic.001G340000.3 | Zm00001d029490_T004 |
| orthologID_008253 | PH01006104G0030 | LOC_Os03g13740.1 | Bradi1g68530.1 | HORVU4Hr1G064070.1 | TRIUR3_23931 | Seita.9G474100.1 | Sobic.001G441100.1 | Zm00001d028374_T001 |
| orthologID_007885 | PH01000322G0310 | LOC_Os03g18420.1 | Bradi1g65310.2 | HORVU4Hr1G055850.1 | TRIUR3_11089 | Seita.9G439000.1 | Sobic.001G404800.1 | Zm00001d028773_T002 |
| orthologID_007674 | PH01001115G0280 | LOC_Os03g16170.1 | Bradi1g66920.1 | HORVU4Hr1G060370.1 | TRIUR3_15727 | Seita.9G460200.1 | Sobic.001G424400.1 | Zm00001d028574_T001 |
| orthologID_007854 | PH01000985G0410 | LOC_Os03g61680.1 | Bradi1g02930.1 | HORVU5Hr1G116650.1 | TRIUR3_05883 | Seita.9G023600.1 | Sobic.001G025600.1 | Zm00001d042829_T001 |
| orthologID_007919 | PH01002146G0070 | LOC_Os03g50280.1 | Bradi1g11280.1 | HORVU4Hr1G007360.1 | TRIUR3_25479 | Seita.9G120000.1 | Sobic.001G118300.1 | Zm00001d033718_T001 |
| orthologID_009122 | PH01000103G1610 | LOC_Os10g26680.1 | Bradi3g24750.1 | HORVU1Hr1G048280.2 | TRIUR3_24386 | Seita.9G259700.1 | Sobic.001G252300.1 | Zm00001d032992_T001 |
| orthologID_006247 | PH01000520G0340 | LOC_Os10g35690.1 | Bradi2g58736.1 | HORVU1Hr1G062620.1 | TRIUR3_15370 | Seita.9G204000.1 | Sobic.001G257700.1 | Zm00001d028672_T001 |
| orthologID_008308 | PH01001096G0100 | LOC_Os03g45260.1 | Bradi1g13350.2 | HORVU4Hr1G002150.1 | TRIUR3_00735 | Seita.9G152000.1 | Sobic.001G149700.1 | Zm00001d033488_T001 |
| orthologID_007841 | PH01001498G0070 | LOC_Os03g09230.1 | Bradi1g71680.1 | HORVU4Hr1G071790.1 | TRIUR3_11802 | Seita.9G508700.1 | Sobic.001G473200.1 | Zm00001d048202_T002 |
| orthologID_008541 | PH01001363G0420 | LOC_Os03g14280.1 | Bradi1g68160.1 | HORVU4Hr1G063150.3 | TRIUR3_28932 | Seita.9G469900.1 | Sobic.001G437200.1 | Zm00001d028414_T002 |
| orthologID_008863 | PH01000338G0890 | LOC_Os04g52710.1 | Bradi5g21900.1 | HORVU2Hr1G104280.1 | TRIUR3_13128 | Seita.7G238800.1 | Sobic.006G217000.1 | Zm00001d002296_T001 |
| orthologID_008792 | PH01001214G0300 | LOC_Os04g52920.1 | Bradi5g22090.2 | HORVU2Hr1G103570.5 | TRIUR3_05455 | Seita.7G241200.1 | Sobic.006G219500.1 | Zm00001d002274_T001 |
| orthologID_008857 | PH01000583G0420 | LOC_Os04g35180.2 | Bradi5g10250.1 | HORVU2Hr1G075280.2 | TRIUR3_12244 | Seita.7G099800.1 | Sobic.006G082500.1 | Zm00001d003663_T001 |
| orthologID_008864 | PH01003491G0120 | LOC_Os04g54790.1 | Bradi5g23450.1 | HORVU2Hr1G115690.4 | TRIUR3_04048 | Seita.7G257500.1 | Sobic.006G235400.3 | Zm00001d002129_T001 |
| orthologID_008912 | PH01000087G0040 | LOC_Os04g58330.1 | Bradi5g26350.3 | HORVU2Hr1G118680.7 | TRIUR3_15619 | Seita.3G013800.1 | Sobic.006G269700.1 | Zm00001d001811_T001 |
| orthologID_008827 | PH01000388G0640 | LOC_Os05g22970.1 | Bradi5g03830.1 | HORVU2Hr1G010100.2 | TRIUR3_16591 | Seita.7G030600.1 | Sobic.006G029800.1 | Zm00001d004123_T001 |
| orthologID_008844 | PH01006750G0030 | LOC_Os04g28060.1 | Bradi5g05700.1 | HORVU2Hr1G067430.9 | TRIUR3_16826 | Seita.7G057000.1 | Sobic.006G040000.1 | Zm00001d004090_T004 |
| orthologID_008886 | PH01000810G0630 | LOC_Os08g03870.1 | Bradi5g25480.1 | HORVU2Hr1G118410.1 | TRIUR3_07651 | Seita.3G031000.1 | Sobic.006G258300.3 | Zm00001d026556_T001 |
| orthologID_008800 | PH01001287G0410 | LOC_Os04g43550.1 | Bradi5g15580.1 | HORVU2Hr1G089130.1 | TRIUR3_32230 | Seita.7G166200.1 | Sobic.006G147200.1 | Zm00001d025859_T001 |
| orthologID_007823 | PH01000101G0280 | LOC_Os03g11690.2 | Bradi1g69810.2 | HORVU4Hr1G067550.5 | TRIUR3_15634 | Seita.9G489600.1 | Sobic.006G067000.2 | Zm00001d028225_T001 |
| orthologID_008956 | PH01002995G0220 | LOC_Os04g33830.1 | Bradi5g09530.1 | HORVU2Hr1G073370.3 | TRIUR3_34121 | Seita.7G092700.1 | Sobic.006G073500.1 | Zm00001d003767_T001 |
| orthologID_008890 | PH01001326G0560 | LOC_Os04g30910.1 | Bradi5g07350.1 | HORVU2Hr1G068530.2 | TRIUR3_31401 | Seita.7G066000.1 | Sobic.006G049700.1 | Zm00001d004023_T002 |
| orthologID_008934 | PH01000968G0250 | LOC_Os04g37740.1 | Bradi5g11880.1 | HORVU2Hr1G079170.1 | TRIUR3_34655 | Seita.7G118000.1 | Sobic.006G100200.1 | Zm00001d003497_T001 |
| orthologID_008941 | PH01001058G0600 | LOC_Os04g55590.1 | Bradi5g24080.1 | HORVU2Hr1G113820.5 | TRIUR3_10705 | Seita.7G263200.1 | Sobic.006G241000.1 | Zm00001d026453_T001 |
| orthologID_009510 | PH01001817G0130 | LOC_Os08g23730.1 | Bradi3g20590.1 | HORVU6Hr1G009430.2 | TRIUR3_04203 | Seita.6G113200.1 | Sobic.006G278500.2 | Zm00001d026685_T002 |
| orthologID_008952 | PH01000000G2440 | LOC_Os04g48490.1 | Bradi5g18950.1 | HORVU2Hr1G094510.1 | TRIUR3_19371 | Seita.7G206900.1 | Sobic.006G186300.2 | Zm00001d002607_T001 |
| orthologID_008789 | PH01002075G0050 | LOC_Os04g38910.1 | Bradi5g12540.1 | HORVU2Hr1G080770.4 | TRIUR3_29321 | Seita.7G126400.1 | Sobic.006G108700.1 | Zm00001d003399_T001 |
| orthologID_006983 | PH01001556G0340 | LOC_Os05g05460.1 | Bradi2g36187.3 | HORVU1Hr1G019660.1 | TRIUR3_09281 | Seita.3G039700.1 | Sobic.006G010800.1 | Zm00001d004480_T001 |
| orthologID_006617 | PH01000443G0680 | LOC_Os01g12470.1 | Bradi2g07390.1 | HORVU3Hr1G030340.10 | TRIUR3_17523 | Seita.5G015400.1 | Sobic.006G275100.6 | Zm00001d026670_T001 |
| orthologID_008831 | PH01005560G0070 | LOC_Os04g55100.1 | Bradi5g23717.1 | HORVU2Hr1G114900.13 | TRIUR3_31947 | Seita.7G260000.1 | Sobic.006G237600.1 | Zm00001d026432_T001 |
| orthologID_008995 | PH01000152G0320 | LOC_Os04g51166.1 | Bradi5g20570.1 | HORVU2Hr1G101940.1 | TRIUR3_34372 | Seita.7G224200.1 | Sobic.006G203100.1 | Zm00001d002433_T002 |
| orthologID_009033 | PH01000028G1180 | LOC_Os04g44890.1 | Bradi5g16580.1 | HORVU2Hr1G091370.3 | TRIUR3_06662 | Seita.7G178400.1 | Sobic.006G158100.1 | Zm00001d025919_T001 |
| orthologID_008839 | PH01001899G0110 | LOC_Os04g57670.1 | Bradi5g25817.1 | HORVU2Hr1G120630.5 | TRIUR3_11004 | Seita.3G019500.1 | Sobic.006G262800.1 | Zm00001d026592_T001 |
| orthologID_008981 | PH01000000G2050 | LOC_Os04g48880.1 | Bradi5g19130.1 | HORVU2Hr1G094160.2 | TRIUR3_22453 | Seita.7G209000.1 | Sobic.006G188200.2 | Zm00001d002589_T001 |
| orthologID_008943 | PH01000042G0410 | LOC_Os04g46050.1 | Bradi5g17390.2 | HORVU2Hr1G093330.17 | TRIUR3_22537 | Seita.2G157900.1 | Sobic.006G166200.1 | Zm00001d002791_T002 |
| orthologID_008983 | PH01000087G0120 | LOC_Os04g58240.1 | Bradi5g26280.2 | HORVU2Hr1G118930.1 | TRIUR3_21790 | Seita.3G014300.1 | Sobic.006G268900.1 | Zm00001d026630_T001 |
| orthologID_006467 | PH01000076G1900 | LOC_Os01g52170.1 | Bradi2g48367.2 | HORVU3Hr1G068300.36 | TRIUR3_24327 | Seita.5G299100.1 | Sobic.006G010900.1 | Zm00001d004478_T001 |
| orthologID_008959 | PH01002905G0090 | LOC_Os04g59580.1 | Bradi5g27580.2 | HORVU2Hr1G127190.12 | TRIUR3_24137 | Seita.3G001800.1 | Sobic.006G280900.2 | Zm00001d026701_T001 |
| orthologID_008812 | PH01001918G0180 | LOC_Os04g33880.1 | Bradi5g09570.1 | HORVU2Hr1G073410.4 | TRIUR3_33017 | Seita.7G093100.1 | Sobic.006G073900.1 | Zm00001d025362_T002 |
| orthologID_009017 | PH01001636G0120 | LOC_Os04g33660.2 | Bradi5g09340.1 | HORVU2Hr1G073030.6 | TRIUR3_33715 | Seita.7G090500.1 | Sobic.006G069300.1 | Zm00001d003781_T002 |
| orthologID_008992 | PH01000561G0570 | LOC_Os04g35900.1 | Bradi5g10800.1 | HORVU2Hr1G076790.4 | TRIUR3_26307 | Seita.7G106100.1 | Sobic.006G088000.1 | Zm00001d025453_T006 |
| orthologID_009039 | PH01001668G0080 | LOC_Os04g42520.1 | Bradi5g14910.1 | HORVU2Hr1G087090.5 | TRIUR3_04270 | Seita.7G157900.1 | Sobic.006G138800.1 | Zm00001d003108_T007 |
| orthologID_009024 | PH01000020G0110 | LOC_Os04g40130.1 | Bradi5g13435.1 | HORVU2Hr1G082620.1 | TRIUR3_34538 | Seita.7G137300.1 | Sobic.006G120200.2 | Zm00001d003294_T001 |
| orthologID_008779 | PH01000657G0060 | LOC_Os04g51300.1 | Bradi5g20670.1 | HORVU2Hr1G101730.3 | TRIUR3_15946 | Seita.7G225300.1 | Sobic.006G204000.2 | Zm00001d002426_T003 |

| | | | | | | | | |
|---|---|---|---|---|---|---|---|---|
| orthologID_008866 | PH01000160G1200 | LOC_Os04g49260.1 | Bradi5g19270.2 | HORVU2Hr1G093870.4 | TRIUR3_24418 | Seita.7G211400.1 | Sobic.006G190600.2 | Zm00001d002558_T001 |
| orthologID_009069 | PH01001688G0260 | LOC_Os04g02140.1 | Bradi5g01920.2 | HORVU2Hr1G005110.4 | TRIUR3_05250 | Seita.7G011900.1 | Sobic.006G008800.1 | Zm00001d004512_T001 |
| orthologID_008881 | PH01000020G1790 | LOC_Os04g41140.1 | Bradi5g14060.1 | HORVU2Hr1G084600.11 | TRIUR3_13774 | Seita.7G146100.1 | Sobic.006G128400.1 | Zm00001d003227_T002 |
| orthologID_008823 | PH01002036G0170 | LOC_Os04g01510.1 | Bradi5g02270.2 | HORVU2Hr1G005680.1 | TRIUR3_02863 | Seita.7G006400.1 | Sobic.006G003100.1 | Zm00001d024925_T001 |
| orthologID_009054 | PH01000041G1500 | LOC_Os04g41150.1 | Bradi5g14070.2 | HORVU2Hr1G084590.1 | TRIUR3_13773 | Seita.7G145800.1 | Sobic.006G128300.4 | Zm00001d003224_T002 |
| orthologID_008808 | PH01000630G0250 | LOC_Os04g35430.1 | Bradi5g10440.4 | HORVU2Hr1G075850.13 | TRIUR3_33277 | Seita.7G101700.1 | Sobic.006G083900.1 | Zm00001d003648_T002 |
| orthologID_008764 | PH01001910G0450 | LOC_Os04g56320.1 | Bradi5g24590.1 | HORVU0Hr1G005570.3 | TRIUR3_03343 | Seita.3G023400.1 | Sobic.006G248700.2 | Zm00001d001995_T003 |
| orthologID_008788 | PH01000016G1050 | LOC_Os04g32590.1 | Bradi5g08600.2 | HORVU2Hr1G071100.8 | TRIUR3_13384 | Seita.7G079900.1 | Sobic.006G058400.1 | Zm00001d025275_T001 |
| orthologID_008783 | PH01003129G0210 | LOC_Os04g54820.1 | Bradi5g23500.1 | HORVU2Hr1G115630.5 | TRIUR3_32184 | Seita.7G257900.1 | Sobic.006G235700.1 | Zm00001d002127_T001 |
| orthologID_008904 | PH01000615G0020 | LOC_Os04g15840.1 | Bradi5g04120.1 | HORVU2Hr1G010800.1 | TRIUR3_14684 | Seita.7G029300.1 | Sobic.006G031900.1 | Zm00001d025018_T001 |
| orthologID_008930 | PH01000378G0230 | LOC_Os04g33370.1 | Bradi5g09120.1 | HORVU2Hr1G072400.1 | TRIUR3_26817 | Seita.7G088100.1 | Sobic.006G066400.1 | Zm00001d003823_T001 |
| orthologID_008794 | PH01001918G0130 | LOC_Os04g33940.1 | Bradi5g09630.1 | HORVU2Hr1G073740.1 | TRIUR3_07003 | Seita.7G093700.1 | Sobic.006G074500.1 | Zm00001d003756_T001 |
| orthologID_008982 | PH01000442G0880 | LOC_Os04g44340.1 | Bradi5g16200.1 | HORVU2Hr1G090400.3 | TRIUR3_11764 | Seita.7G173300.1 | Sobic.006G153000.2 | Zm00001d002971_T001 |
| orthologID_008810 | PH01000001G1780 | LOC_Os04g45200.1 | Bradi5g16850.1 | HORVU2Hr1G091850.2 | TRIUR3_02526 | Seita.7G180900.1 | Sobic.006G160300.2 | Zm00001d025940_T001 |
| orthologID_009066 | PH01001053G0410 | LOC_Os04g55410.1 | Bradi5g23940.1 | HORVU2Hr1G114260.1 | TRIUR3_12521 | Seita.7G262100.1 | Sobic.006G239800.1 | Zm00001d002085_T001 |
| orthologID_009000 | PH01000000G0480 | LOC_Os04g49610.1 | Bradi5g19590.1 | HORVU2Hr1G104940.1 | TRIUR3_27923 | Seita.7G214400.1 | Sobic.006G193500.1 | Zm00001d026025_T002 |
| orthologID_009073 | PH01000020G1500 | LOC_Os04g40940.1 | Bradi5g13910.1 | HORVU2Hr1G084050.1 | TRIUR3_08259 | Seita.7G144300.1 | Sobic.006G126700.1 | Zm00001d025721_T001 |
| orthologID_008951 | PH01005398G0070 | LOC_Os04g43420.1 | Bradi5g15550.2 | HORVU2Hr1G089030.8 | TRIUR3_13397 | Seita.7G164700.1 | Sobic.006G146300.1 | Zm00001d025848_T001 |
| orthologID_008049 | PH01002685G0190 | LOC_Os04g43922.1 | Bradi1g04570.1 | HORVU7Hr1G038290.5 | TRIUR3_28456 | Seita.7G169800.1 | Sobic.006G251600.1 | Zm00001d001973_T001 |
| orthologID_008909 | PH01000030G0550 | LOC_Os04g50110.1 | Bradi5g19970.1 | HORVU2Hr1G105930.2 | TRIUR3_25488 | Seita.7G218700.1 | Sobic.006G197700.1 | Zm00001d002490_T001 |
| orthologID_008813 | PH01000810G0690 | LOC_Os04g57310.1 | Bradi5g25506.1 | HORVU2Hr1G118360.1 | TRIUR3_12911 | Seita.3G031300.1 | Sobic.006G258600.2 | Zm00001d026560_T001 |
| orthologID_008961 | PH01000625G0220 | LOC_Os04g38420.1 | Bradi5g12180.1 | HORVU2Hr1G079930.1 | TRIUR3_29462 | Seita.7G121900.1 | Sobic.006G104100.1 | Zm00001d003448_T002 |
| orthologID_009040 | PH01000561G0140 | LOC_Os04g35770.1 | Bradi5g10700.1 | HORVU2Hr1G076520.2 | TRIUR3_05261 | Seita.7G104600.1 | Sobic.006G086900.1 | Zm00001d003617_T001 |
| orthologID_008802 | PH01000044G1910 | LOC_Os04g58580.1 | Bradi5g26500.1 | HORVU2Hr1G122950.1 | TRIUR3_19116 | Seita.3G011800.1 | Sobic.006G271500.1 | Zm00001d001789_T001 |
| orthologID_008806 | PH01000224G0300 | LOC_Os04g51880.1 | Bradi5g21040.1 | HORVU2Hr1G100380.3 | TRIUR3_20846 | Seita.7G230800.1 | Sobic.006G208200.1 | Zm00001d002377_T001 |
| orthologID_008905 | PH01005530G0080 | LOC_Os04g43220.1 | Bradi5g15450.1 | HORVU2Hr1G088780.1 | TRIUR3_23722 | Seita.7G163200.1 | Sobic.006G144000.2 | Zm00001d025837_T001 |
| orthologID_008907 | PH01000825G0440 | LOC_Os04g35530.1 | Bradi5g10500.1 | HORVU2Hr1G076010.6 | TRIUR3_26992 | Seita.7G102300.1 | Sobic.006G084500.1 | Zm00001d025435_T002 |
| orthologID_008825 | PH01002660G0040 | LOC_Os04g48360.1 | Bradi5g18853.1 | HORVU2Hr1G094760.8 | TRIUR3_26658 | Seita.7G205300.1 | Sobic.006G185000.1 | Zm00001d002616_T001 |
| orthologID_008840 | PH01000318G0370 | LOC_Os04g20990.1 | Bradi5g03740.1 | HORVU2Hr1G065180.24 | TRIUR3_31883 | Seita.7G028100.1 | Sobic.006G038500.1 | Zm00001d004196_T002 |
| orthologID_008911 | PH01005619G0070 | LOC_Os04g55150.2 | Bradi5g23760.1 | HORVU2Hr1G114660.3 | TRIUR3_06846 | Seita.7G260300.1 | Sobic.006G237900.1 | Zm00001d002095_T001 |
| orthologID_008928 | PH01000200G0930 | LOC_Os04g48060.1 | Bradi5g18710.1 | HORVU2Hr1G095220.3 | TRIUR3_25906 | Seita.7G203200.1 | Sobic.006G183100.1 | Zm00001d026089_T001 |
| orthologID_009018 | PH01001075G0390 | LOC_Os04g32620.1 | Bradi5g08380.1 | HORVU2Hr1G071270.4 | TRIUR3_27406 | Seita.7G080300.1 | Sobic.006G058732.1 | Zm00001d003884_T001 |
| orthologID_008854 | PH01000561G0150 | LOC_Os04g35780.1 | Bradi5g10730.1 | HORVU0Hr1G004790.1 | TRIUR3_35101 | Seita.7G104800.1 | Sobic.006G087400.1 | Zm00001d025452_T001 |
| orthologID_008816 | PH01000028G0330 | LOC_Os04g45280.1 | Bradi5g16890.1 | HORVU2Hr1G091890.10 | TRIUR3_18629 | Seita.7G181200.1 | Sobic.006G160600.1 | Zm00001d025941_T001 |
| orthologID_009071 | PH01000300G0610 | LOC_Os04g52780.1 | Bradi5g21960.1 | HORVU2Hr1G104030.4 | TRIUR3_18210 | Seita.7G239900.1 | Sobic.006G217900.1 | Zm00001d002287_T001 |
| orthologID_008985 | PH01000801G0160 | LOC_Os04g43150.2 | Bradi5g15390.1 | HORVU2Hr1G088550.12 | TRIUR3_26083 | Seita.7G162600.1 | Sobic.006G143300.1 | Zm00001d003033_T005 |
| orthologID_008837 | PH01000300G0400 | LOC_Os04g52700.1 | Bradi5g21895.1 | HORVU2Hr1G104270.1 | TRIUR3_13129 | Seita.7G238700.1 | Sobic.006G216900.2 | Zm00001d002301_T001 |
| orthologID_009063 | PH01000044G1890 | LOC_Os04g58600.2 | Bradi5g26520.1 | HORVU2Hr1G123070.30 | TRIUR3_01484 | Seita.3G011500.1 | Sobic.006G271700.1 | Zm00001d026643_T002 |
| orthologID_008885 | PH01000087G0440 | LOC_Os04g58080.1 | Bradi5g26150.1 | HORVU2Hr1G119550.1 | TRIUR3_09473 | Seita.3G015900.1 | Sobic.006G266900.1 | Zm00001d026617_T001 |
| orthologID_008752 | PH01001151G0360 | LOC_Os04g55240.1 | Bradi5g23893.1 | HORVU2Hr1G114570.1 | TRIUR3_16712 | Seita.7G261300.1 | Sobic.006G239100.2 | Zm00001d002088_T001 |
| orthologID_008819 | PH01000028G2440 | LOC_Os02g54000.1 | Bradi5g16280.2 | HORVU2Hr1G090680.26 | TRIUR3_27488 | Seita.7G174100.1 | Sobic.006G154100.1 | Zm00001d002961_T002 |
| orthologID_008929 | PH01000598G0080 | LOC_Os04g42870.2 | Bradi5g15167.4 | HORVU2Hr1G087840.3 | TRIUR3_23512 | Seita.7G160700.1 | Sobic.006G140900.2 | Zm00001d025819_T001 |
| orthologID_006283 | PH01005398G0080 | LOC_Os04g43430.2 | Bradi2g37360.1 | HORVU3Hr1G015010.23 | TRIUR3_16759 | Seita.7G164600.1 | Sobic.006G146400.4 | Zm00001d003047_T001 |
| orthologID_009070 | PH01001918G0370 | LOC_Os04g33840.1 | Bradi5g09540.2 | HORVU2Hr1G073360.13 | TRIUR3_30518 | Seita.7G092800.1 | Sobic.006G073600.2 | Zm00001d003765_T001 |
| orthologID_008879 | PH01000328G0720 | LOC_Os04g36760.1 | Bradi5g11100.1 | HORVU2Hr1G077760.1 | TRIUR3_31726 | Seita.7G112200.1 | Sobic.006G093900.1 | Zm00001d025509_T001 |
| orthologID_008787 | PH01003796G0120 | LOC_Os04g56620.1 | Bradi5g24930.1 | HORVU2Hr1G110680.1 | TRIUR3_16099 | Seita.3G025900.1 | Sobic.006G251800.1 | Zm00001d026515_T003 |
| orthologID_008938 | PH01000000G0720 | LOC_Os04g49520.1 | Bradi5g19440.1 | HORVU2Hr1G104660.3 | TRIUR3_01838 | Seita.7G213300.1 | Sobic.006G192600.1 | Zm00001d002539_T001 |
| orthologID_008908 | PH01001636G0330 | LOC_Os04g33700.1 | Bradi5g09380.1 | HORVU2Hr1G073100.11 | TRIUR3_18786 | Seita.7G090900.1 | Sobic.006G071362.1 | Zm00001d003778_T001 |
| orthologID_008968 | PH01001058G0330 | LOC_Os04g55700.1 | Bradi5g24250.1 | HORVU2Hr1G113320.9 | TRIUR3_01781 | Seita.7G264700.1 | Sobic.006G243100.2 | Zm00001d002053_T001 |
| orthologID_008170 | PH01000630G0170 | LOC_Os04g35480.1 | Bradi1g54570.1 | HORVU2Hr1G075910.13 | TRIUR3_20740 | Seita.7G101800.1 | Sobic.006G084000.1 | Zm00001d025433_T003 |

| | | | | | | | | |
|---|---|---|---|---|---|---|---|---|
| orthologID_008853 | PH01000041G1890 | LOC_Os04g38850.1 | Bradi5g12490.1 | HORVU2Hr1G080580.11 | TRIUR3_19321 | Seita.7G125800.1 | Sobic.006G108200.2 | Zm00001d003404_T001 |
| orthologID_008828 | PH01001023G0020 | LOC_Os04g57950.1 | Bradi5g26080.1 | HORVU2Hr1G119720.1 | TRIUR3_13184 | Seita.3G016500.1 | Sobic.006G266200.2 | Zm00001d026610_T001 |
| orthologID_009072 | PH01000087G1080 | LOC_Os04g57780.1 | Bradi5g25910.2 | HORVU2Hr1G120390.31 | TRIUR3_00810 | Seita.3G018500.1 | Sobic.006G263700.4 | Zm00001d001861_T001 |
| orthologID_008925 | PH01000087G1240 | LOC_Os04g57690.1 | Bradi5g25840.3 | HORVU2Hr1G120590.2 | TRIUR3_25797 | Seita.3G019200.1 | Sobic.006G263100.1 | Zm00001d001868_T001 |
| orthologID_008918 | PH01002245G0210 | LOC_Os04g53690.1 | Bradi5g22710.1 | HORVU2Hr1G107800.3 | TRIUR3_17741 | Seita.7G247800.1 | Sobic.006G225300.2 | Zm00001d026361_T001 |
| orthologID_008948 | PH01001828G0240 | LOC_Os04g56654.1 | Bradi5g24967.2 | HORVU2Hr1G110600.1 | TRIUR3_01830 | Seita.3G026300.1 | Sobic.006G252300.1 | Zm00001d026518_T001 |
| orthologID_008856 | PH01001270G0050 | LOC_Os04g27850.1 | Bradi5g05630.1 | HORVU2Hr1G066950.2 | TRIUR3_33837 | Seita.7G053400.1 | Sobic.006G035500.1 | Zm00001d004120_T001 |
| orthologID_009048 | PH01001613G0110 | LOC_Os04g36750.1 | Bradi5g11110.1 | HORVU2Hr1G077710.1 | TRIUR3_31593 | Seita.7G111700.1 | Sobic.006G093500.1 | Zm00001d003554_T001 |
| orthologID_008903 | PH01000679G0350 | LOC_Os04g54190.1 | Bradi5g23120.1 | HORVU2Hr1G116960.2 | TRIUR3_10287 | Seita.7G253200.1 | Sobic.006G230300.1 | Zm00001d002167_T001 |
| orthologID_008996 | PH01000386G0490 | LOC_Os04g50720.1 | Bradi5g20070.1 | HORVU2Hr1G106250.3 | TRIUR3_15994 | Seita.7G219900.1 | Sobic.006G199200.1 | Zm00001d002478_T001 |
| orthologID_008745 | PH01002905G0130 | LOC_Os04g59560.1 | Bradi5g27560.1 | HORVU2Hr1G127090.4 | TRIUR3_03893 | Seita.3G002000.1 | Sobic.006G280600.1 | Zm00001d002942_T002 |
| orthologID_006895 | PH01000432G0880 | LOC_Os12g19090.1 | Bradi2g38290.1 | HORVU2Hr1G083530.1 | TRIUR3_08422 | Seita.7G141600.1 | Sobic.006G124300.2 | Zm00001d003268_T001 |
| orthologID_008421 | PH01002979G0200 | LOC_Os07g39090.1 | Bradi1g20737.1 | HORVU2Hr1G029200.4 | TRIUR3_19832 | Seita.7G165500.1 | Sobic.006G146900.1 | Zm00001d003050_T001 |
| orthologID_009043 | PH01001603G0190 | LOC_Os04g37630.1 | Bradi5g11757.5 | HORVU2Hr1G078840.14 | TRIUR3_07454 | Seita.7G116500.1 | Sobic.006G097700.1 | Zm00001d003511_T001 |
| orthologID_008799 | PH01000810G0430 | LOC_Os04g57150.1 | Bradi5g25400.1 | HORVU2Hr1G108810.3 | TRIUR3_21007 | Seita.3G029900.1 | Sobic.006G257300.1 | Zm00001d026549_T006 |
| orthologID_008793 | PH01000467G0350 | LOC_Os04g42784.1 | Bradi5g15120.1 | HORVU2Hr1G087660.41 | TRIUR3_21148 | Seita.7G160000.1 | Sobic.006G140400.1 | Zm00001d025816_T002 |
| orthologID_008774 | PH01000030G0050 | LOC_Os04g50770.1 | Bradi5g20130.1 | HORVU2Hr1G106400.3 | TRIUR3_18702 | Seita.7G220500.1 | Sobic.006G199800.1 | Zm00001d002476_T001 |
| orthologID_008761 | PH01007831G0010 | LOC_Os04g59630.1 | Bradi5g27617.1 | HORVU2Hr1G124850.1 | TRIUR3_03594 | Seita.3G001200.1 | Sobic.006G281500.1 | Zm00001d026702_T002 |
| orthologID_009001 | PH01000028G2140 | LOC_Os04g44540.2 | Bradi5g16330.1 | HORVU2Hr1G090800.1 | TRIUR3_11622 | Seita.7G175300.1 | Sobic.006G154900.1 | Zm00001d025904_T001 |
| orthologID_008940 | PH01000384G0860 | LOC_Os04g35650.1 | Bradi5g10630.3 | HORVU2Hr1G076380.1 | TRIUR3_18454 | Seita.7G103700.1 | Sobic.006G085900.1 | Zm00001d025463_T001 |
| orthologID_008990 | PH01000001G1770 | LOC_Os04g45210.1 | Bradi5g16860.1 | HORVU2Hr1G091870.2 | TRIUR3_02525 | Seita.7G181000.1 | Sobic.006G160400.1 | Zm00001d002834_T001 |
| orthologID_009020 | PH01001688G0350 | LOC_Os04g02000.1 | Bradi5g01960.1 | HORVU2Hr1G004930.1 | TRIUR3_05246 | Seita.7G011500.1 | Sobic.006G008600.2 | Zm00001d004517_T005 |
| orthologID_008776 | PH01000488G0370 | LOC_Os04g19684.1 | Bradi5g05225.2 | HORVU2Hr1G012840.4 | TRIUR3_20569 | Seita.7G028500.1 | Sobic.006G034100.1 | Zm00001d025128_T004 |
| orthologID_008978 | PH01001667G0430 | LOC_Os04g34984.3 | Bradi5g10117.2 | HORVU2Hr1G074950.11 | TRIUR3_31224 | Seita.7G098300.1 | Sobic.006G080400.1 | Zm00001d003678_T003 |
| orthologID_008947 | PH01000225G1060 | LOC_Os04g51180.1 | Bradi5g20597.1 | HORVU2Hr1G101760.2 | TRIUR3_24949 | Seita.7G224400.1 | Sobic.006G203300.1 | Zm00001d026239_T004 |
| orthologID_008936 | PH01000118G0230 | LOC_Os04g42580.1 | Bradi5g14970.1 | HORVU2Hr1G087400.1 | TRIUR3_16548 | Seita.7G158400.1 | Sobic.006G139000.1 | Zm00001d025893_T001 |
| orthologID_009034 | PH01000001G2410 | LOC_Os04g44870.1 | Bradi5g16570.1 | HORVU2Hr1G091320.5 | TRIUR3_30883 | Seita.7G178300.1 | Sobic.006G157900.1 | Zm00001d002850_T001 |
| orthologID_008875 | PH01000224G0970 | LOC_Os04g51580.2 | Bradi5g20880.2 | HORVU2Hr1G101090.6 | TRIUR3_18618 | Seita.7G228100.1 | Sobic.006G206700.1 | Zm00001d002402_T001 |
| orthologID_008760 | PH01003848G0010 | LOC_Os04g36890.1 | Bradi5g10980.1 | HORVU2Hr1G078350.3 | TRIUR3_20435 | Seita.7G113400.1 | Sobic.006G095400.1 | Zm00001d025522_T001 |
| orthologID_008804 | PH01000152G0680 | LOC_Os04g51310.1 | Bradi5g20690.1 | HORVU2Hr1G101610.1 | TRIUR3_11078 | Seita.7G225400.1 | Sobic.006G204200.1 | Zm00001d026244_T001 |
| orthologID_008429 | PH01002503G0110 | LOC_Os04g32970.1 | Bradi1g72850.1 | HORVU2Hr1G071830.18 | TRIUR3_29562 | Seita.7G083700.1 | Sobic.006G062800.1 | Zm00001d025307_T004 |
| orthologID_008922 | PH01000615G0340 | LOC_Os04g15690.1 | Bradi5g04030.1 | HORVU2Hr1G010560.1 | TRIUR3_00604 | Seita.7G029500.1 | Sobic.006G031200.1 | Zm00001d025016_T003 |
| orthologID_008932 | PH01000041G1770 | LOC_Os04g38920.1 | Bradi5g12547.1 | HORVU2Hr1G080820.2 | TRIUR3_30247 | Seita.7G126500.1 | Sobic.006G108832.1 | Zm00001d003398_T001 |
| orthologID_008987 | PH01000432G0250 | LOC_Os04g40990.1 | Bradi5g13940.1 | HORVU2Hr1G084130.1 | TRIUR3_14305 | Seita.7G144600.1 | Sobic.006G127100.1 | Zm00001d003247_T002 |
| orthologID_008974 | PH01000901G0070 | LOC_Os04g58570.1 | Bradi5g26490.1 | HORVU2Hr1G123030.2 | TRIUR3_24257 | Seita.3G011900.1 | Sobic.006G271300.1 | Zm00001d026642_T001 |
| orthologID_008858 | PH01001666G0060 | LOC_Os04g21340.1 | Bradi5g05120.1 | HORVU2Hr1G065390.1 | TRIUR3_29245 | Seita.7G026000.1 | Sobic.006G037100.1 | Zm00001d004209_T001 |
| orthologID_008851 | PH01001058G0610 | LOC_Os04g55570.1 | Bradi5g24090.1 | HORVU2Hr1G113830.11 | TRIUR3_10706 | Seita.7G263100.1 | Sobic.006G240900.1 | Zm00001d026452_T001 |
| orthologID_009256 | PH01000952G0060 | LOC_Os02g21880.1 | Bradi3g10690.1 | HORVU1Hr1G008620.1 | TRIUR3_30742 | Seita.1G145900.1 | Sobic.006G228500.1 | Zm00001d026381_T001 |
| orthologID_008859 | PH01001030G0610 | LOC_Os04g30430.1 | Bradi5g07090.1 | HORVU2Hr1G042280.3 | TRIUR3_18851 | Seita.7G063800.1 | Sobic.006G041100.2 | Zm00001d004070_T003 |
| orthologID_006257 | PH01002344G0070 | LOC_Os01g43360.1 | Bradi2g43870.1 | HORVU3Hr1G055630.1 | TRIUR3_29691 | Seita.5G230700.1 | Sobic.006G240800.1 | Zm00001d026451_T001 |
| orthologID_009042 | PH01001668G0040 | LOC_Os04g42540.1 | Bradi5g14920.1 | HORVU2Hr1G087070.3 | TRIUR3_04271 | Seita.7G158100.1 | Sobic.006G138900.2 | Zm00001d003107_T001 |
| orthologID_006201 | PH01004000G0140 | LOC_Os05g24880.1 | Bradi2g31190.1 | HORVU6Hr1G063770.2 | TRIUR3_10423 | Seita.3G290100.1 | Sobic.006G222100.1 | Zm00001d002246_T001 |
| orthologID_008821 | PH01000583G0580 | LOC_Os04g35100.1 | Bradi5g10177.2 | HORVU2Hr1G075110.5 | TRIUR3_02641 | Seita.7G099100.1 | Sobic.006G081200.1 | Zm00001d003671_T001 |
| orthologID_009410 | PH01002027G0070 | LOC_Os02g29380.1 | Bradi3g43720.1 | HORVU3Hr1G003060.16 | TRIUR3_01116 | Seita.1G164500.1 | Sobic.006G224100.1 | Zm00001d016521_T001 |
| orthologID_008979 | PH01000016G1090 | LOC_Os04g32600.1 | Bradi5g08591.1 | HORVU2Hr1G071170.4 | TRIUR3_35088 | Seita.7G080000.1 | Sobic.006G058500.1 | Zm00001d003888_T002 |
| orthologID_008756 | PH01001222G0560 | LOC_Os04g34940.1 | Bradi5g10100.1 | HORVU2Hr1G074850.7 | TRIUR3_21259 | Seita.7G098000.1 | Sobic.006G080000.1 | Zm00001d003684_T001 |
| orthologID_008778 | PH01000087G0500 | LOC_Os04g58060.1 | Bradi5g26120.1 | HORVU2Hr1G119480.1 | TRIUR3_09322 | Seita.3G016100.1 | Sobic.006G266700.1 | Zm00001d026614_T001 |
| orthologID_008797 | PH01001543G0540 | LOC_Os04g36830.1 | Bradi5g11020.2 | HORVU2Hr1G078270.7 | TRIUR3_18838 | Seita.7G112900.1 | Sobic.006G094900.1 | Zm00001d003544_T003 |
| orthologID_010125 | PH01001250G0050 | LOC_Os04g50100.1 | Bradi5g19960.1 | HORVU4Hr1G004390.2 | TRIUR3_07473 | Seita.7G218600.1 | Sobic.006G197600.1 | Zm00001d002491_T001 |
| orthologID_009021 | PH01000000G5760 | LOC_Os04g47310.1 | Bradi5g18260.5 | HORVU5Hr1G048810.2 | TRIUR3_23125 | Seita.7G197800.1 | Sobic.006G177600.1 | Zm00001d026137_T002 |

| | | | | | | | | |
|---|---|---|---|---|---|---|---|---|
| orthologID_008949 | PH01000000G3550 | LOC_Os04g48130.1 | Bradi5g18750.1 | HORVU2Hr1G095170.3 | TRIUR3_20089 | Seita.7G203700.1 | Sobic.006G183400.1 | Zm00001d002632_T001 |
| orthologID_008901 | PH01000192G0380 | LOC_Os04g41300.1 | Bradi5g14160.1 | HORVU2Hr1G084680.6 | TRIUR3_16659 | Seita.7G147700.1 | Sobic.006G129900.1 | Zm00001d025745_T001 |
| orthologID_008809 | PH01002245G0230 | LOC_Os04g53670.1 | Bradi5g22700.1 | HORVU2Hr1G107580.20 | TRIUR3_02182 | Seita.7G247600.1 | Sobic.006G224800.1 | Zm00001d026359_T003 |
| orthologID_008939 | PH01000000G0350 | LOC_Os04g49660.1 | Bradi5g19610.1 | HORVU2Hr1G104960.1 | TRIUR3_33801 | Seita.7G214800.1 | Sobic.006G193900.1 | Zm00001d002523_T001 |
| orthologID_008818 | PH01001276G0430 | LOC_Os04g08410.1 | Bradi5g00630.1 | HORVU2Hr1G000130.2 | TRIUR3_06075 | Seita.7G021200.1 | Sobic.006G017800.1 | Zm00001d024873_T001 |
| orthologID_010126 | PH01000272G1270 | LOC_Os04g47690.1 | Bradi5g18460.1 | HORVU2Hr1G095890.1 | TRIUR3_17303 | Seita.7G200000.1 | Sobic.006G179800.1 | Zm00001d002659_T001 |
| orthologID_008876 | PH01000300G0920 | LOC_Os04g52870.1 | Bradi5g22050.4 | HORVU2Hr1G103700.3 | TRIUR3_16031 | Seita.4G120600.1 | Sobic.006G219000.1 | Zm00001d026326_T001 |
| orthologID_009028 | PH01000020G2630 | LOC_Os04g41460.1 | Bradi5g14367.1 | HORVU2Hr1G085160.2 | TRIUR3_03272 | Seita.7G148600.1 | Sobic.006G130800.1 | Zm00001d025749_T001 |
| orthologID_009499 | PH01001522G0130 | LOC_Os10g30870.1 | Bradi3g27730.1 | HORVU1Hr1G037650.5 | TRIUR3_26439 | Seita.9G237700.1 | Sobic.006G269400.1 | Zm00001d001812_T001 |
| orthologID_009005 | PH01000304G1020 | LOC_Os04g36650.1 | Bradi5g11180.1 | HORVU2Hr1G077570.1 | TRIUR3_34498 | Seita.7G110700.1 | Sobic.006G092300.1 | Zm00001d025485_T001 |
| orthologID_009075 | PH01003804G0130 | LOC_Os04g51786.1 | Bradi5g20960.2 | HORVU2Hr1G100600.2 | TRIUR3_09828 | Seita.7G229700.1 | Sobic.006G207700.1 | Zm00001d002385_T002 |
| orthologID_009633 | PH01002694G0120 | LOC_Os10g12354.1 | Bradi3g23020.1 | HORVU0Hr1G038290.1 | TRIUR3_17388 | Seita.9G420000.1 | Sobic.006G265300.1 | Zm00001d001847_T001 |
| orthologID_008973 | PH01005340G0060 | LOC_Os04g30740.1 | Bradi5g07180.1 | HORVU2Hr1G068170.7 | TRIUR3_14174 | Seita.7G064800.1 | Sobic.006G041800.1 | Zm00001d004051_T001 |
| orthologID_008917 | PH01003491G0140 | LOC_Os04g54640.1 | Bradi5g23437.1 | HORVU2Hr1G115710.3 | TRIUR3_08218 | Seita.2G021300.1 | Sobic.006G235350.1 | Zm00001d002130_T002 |
| orthologID_008945 | PH01000225G1240 | LOC_Os04g51280.1 | Bradi5g20660.1 | HORVU2Hr1G101720.2 | TRIUR3_15945 | Seita.7G225200.1 | Sobic.006G204100.1 | Zm00001d026243_T002 |
| orthologID_008795 | PH01003239G0190 | LOC_Os04g31390.1 | Bradi5g07790.1 | HORVU2Hr1G069390.1 | TRIUR3_33706 | Seita.2G399300.1 | Sobic.006G046700.1 | Zm00001d025210_T001 |
| orthologID_008873 | PH01001688G0150 | LOC_Os04g02500.1 | Bradi5g01880.1 | HORVU2Hr1G000590.6 | TRIUR3_16274 | Seita.7G012400.1 | Sobic.006G009400.2 | Zm00001d004498_T001 |
| orthologID_008791 | PH01000152G0440 | LOC_Os04g51190.1 | Bradi5g20607.1 | HORVU2Hr1G101770.7 | TRIUR3_11171 | Seita.7G224500.1 | Sobic.006G203400.1 | Zm00001d026240_T006 |
| orthologID_009003 | PH01000265G0330 | LOC_Os04g33560.1 | Bradi5g09260.1 | HORVU0Hr1G004630.1 | TRIUR3_23343 | Seita.7G089700.1 | Sobic.006G068200.1 | Zm00001d003806_T001 |
| orthologID_008926 | PH01001603G0430 | LOC_Os04g37520.1 | Bradi5g11650.1 | HORVU2Hr1G078640.1 | TRIUR3_23467 | Seita.7G115600.1 | Sobic.006G096700.1 | Zm00001d003520_T001 |
| orthologID_008751 | PH01000087G1630 | LOC_Os04g57510.1 | Bradi5g25660.1 | HORVU2Hr1G121240.5 | TRIUR3_03019 | Seita.3G033300.1 | Sobic.006G260900.1 | Zm00001d026580_T002 |
| orthologID_009011 | PH01000152G0760 | LOC_Os04g51350.1 | Bradi5g20810.1 | HORVU2Hr1G101470.2 | TRIUR3_15349 | Seita.7G226000.1 | Sobic.006G204600.1 | Zm00001d002421_T001 |
| orthologID_008931 | PH01000023G0550 | LOC_Os04g46600.1 | Bradi5g17730.1 | HORVU2Hr1G097640.1 | TRIUR3_29209 | Seita.7G190100.1 | Sobic.006G171050.1 | Zm00001d026173_T001 |
| orthologID_008868 | PH01000023G1110 | LOC_Os04g46370.1 | Bradi5g17580.1 | HORVU2Hr1G097890.16 | TRIUR3_07325 | Seita.7G188100.1 | Sobic.006G169500.1 | Zm00001d002751_T001 |
| orthologID_009045 | PH01005239G0070 | LOC_Os04g55560.2 | Bradi5g24100.1 | HORVU2Hr1G113880.23 | TRIUR3_10707 | Seita.7G263000.1 | Sobic.006G240700.3 | Zm00001d002075_T001 |
| orthologID_010136 | PH01001692G0270 | LOC_Os04g46730.1 | Bradi5g17840.1 | HORVU0Hr1G018480.1 | TRIUR3_05265 | Seita.7G192000.1 | Sobic.006G171900.1 | Zm00001d002727_T002 |
| orthologID_008894 | PH01000020G1560 | LOC_Os04g40970.1 | Bradi5g13920.3 | HORVU2Hr1G084110.2 | TRIUR3_14303 | Seita.7G144400.1 | Sobic.006G126800.1 | Zm00001d025722_T002 |
| orthologID_009067 | PH01000023G1200 | LOC_Os04g46330.1 | Bradi5g17560.3 | HORVU2Hr1G097980.1 | TRIUR3_22485 | Seita.7G187600.1 | Sobic.006G168900.1 | Zm00001d002756_T001 |
| orthologID_008753 | PH01001959G0020 | LOC_Os04g15790.1 | Bradi5g04050.1 | HORVU2Hr1G010520.1 | TRIUR3_00605 | Seita.7G029400.1 | Sobic.006G031300.1 | Zm00001d004132_T001 |
| orthologID_008750 | PH01000020G1370 | LOC_Os04g40870.1 | Bradi5g13850.1 | HORVU7Hr1G090670.1 | TRIUR3_33812 | Seita.7G143700.1 | Sobic.006G126200.1 | Zm00001d025715_T001 |
| orthologID_008755 | PH01000630G0150 | LOC_Os04g35500.1 | Bradi5g10470.1 | HORVU2Hr1G075950.1 | TRIUR3_32843 | Seita.7G102000.1 | Sobic.006G084201.1 | Zm00001d003645_T001 |
| orthologID_009009 | PH01000154G0330 | LOC_Os06g20354.1 | Bradi5g13440.1 | HORVU1Hr1G010220.3 | TRIUR3_05731 | Seita.J024900.1 | Sobic.006G198900.1 | Zm00001d026199_T001 |
| orthologID_008865 | PH01000055G2050 | LOC_Os04g55090.1 | Bradi5g23710.1 | HORVU2Hr1G114930.2 | TRIUR3_01914 | Seita.7G259900.1 | Sobic.006G237500.1 | Zm00001d002098_T001 |
| orthologID_008993 | PH01000001G4480 | LOC_Os04g44210.1 | Bradi5g16060.1 | HORVU2Hr1G090090.1 | TRIUR3_14011 | Seita.7G171500.1 | Sobic.006G151200.1 | Zm00001d002992_T002 |
| orthologID_008777 | PH01000084G1440 | LOC_Os04g52164.1 | Bradi5g21350.1 | HORVU2Hr1G099550.2 | TRIUR3_02819 | Seita.7G233100.1 | Sobic.006G211000.1 | Zm00001d002353_T001 |
| orthologID_008891 | PH01000432G0630 | LOC_Os04g40740.1 | Bradi5g13770.1 | HORVU2Hr1G083780.2 | TRIUR3_10973 | Seita.7G143100.1 | Sobic.006G125700.1 | Zm00001d003259_T006 |
| orthologID_008884 | PH01219833G0010 | LOC_Os04g58120.1 | Bradi5g26180.1 | HORVU2Hr1G119470.1 | TRIUR3_17036 | Seita.3G015400.1 | Sobic.006G267400.1 | Zm00001d026620_T001 |
| orthologID_009041 | PH01000571G0010 | LOC_Os04g36850.1 | Bradi5g11000.1 | HORVU2Hr1G078280.1 | TRIUR3_25588 | Seita.7G113100.1 | Sobic.006G095100.1 | Zm00001d003540_T003 |
| orthologID_009022 | PH01000028G1770 | LOC_Os04g44620.2 | Bradi5g16410.2 | HORVU2Hr1G090950.3 | TRIUR3_22647 | Seita.7G176100.1 | Sobic.006G155800.1 | Zm00001d025907_T001 |
| orthologID_008946 | PH01001870G0190 | LOC_Os04g41820.1 | Bradi5g14497.1 | HORVU2Hr1G085560.4 | TRIUR3_23538 | Seita.7G151800.1 | Sobic.006G133700.1 | Zm00001d003180_T001 |
| orthologID_009053 | PH01004538G0060 | LOC_Os04g53220.1 | Bradi5g22280.2 | HORVU2Hr1G103100.1 | TRIUR3_28987 | Seita.7G243000.1 | Sobic.006G220700.2 | Zm00001d002259_T002 |
| orthologID_008902 | PH01003491G0150 | LOC_Os04g54630.1 | Bradi5g23430.1 | HORVU2Hr1G115730.1 | TRIUR3_08219 | Seita.7G257400.1 | Sobic.006G235300.1 | Zm00001d002131_T001 |
| orthologID_009893 | PH01000736G0030 | LOC_Os08g19830.1 | Bradi3g19947.1 | HORVU3Hr1G014250.5 | TRIUR3_29504 | Seita.2G043600.1 | Sobic.006G252900.1 | Zm00001d001962_T014 |
| orthologID_008766 | PH01000000G2820 | LOC_Os03g46480.1 | Bradi5g18857.2 | HORVU2Hr1G119540.2 | TRIUR3_09472 | Seita.7G205400.1 | Sobic.006G185100.1 | Zm00001d002615_T003 |
| orthologID_008849 | PH01001250G0020 | LOC_Os04g50120.1 | Bradi5g19980.2 | HORVU2Hr1G106020.2 | TRIUR3_25489 | Seita.7G218800.1 | Sobic.006G197800.1 | Zm00001d002489_T001 |
| orthologID_009051 | PH01000044G0910 | LOC_Os04g59060.1 | Bradi5g27050.2 | HORVU2Hr1G124560.2 | TRIUR3_34147 | Seita.3G005600.1 | Sobic.006G277000.1 | Zm00001d002891_T001 |
| orthologID_008897 | PH01000041G0930 | LOC_Os04g39240.1 | Bradi5g12830.2 | HORVU2Hr1G081400.8 | TRIUR3_30012 | Seita.7G129600.1 | Sobic.006G112100.1 | Zm00001d003369_T004 |
| orthologID_008913 | PH01000087G0360 | LOC_Os04g58090.1 | Bradi5g26160.1 | HORVU2Hr1G119500.1 | TRIUR3_27489 | Seita.3G015800.1 | Sobic.006G267000.1 | Zm00001d001832_T001 |
| orthologID_008765 | PH01000968G0320 | LOC_Os04g37750.1 | Bradi5g11887.1 | HORVU2Hr1G079160.5 | TRIUR3_20461 | Seita.7G118100.1 | Sobic.006G100266.1 | Zm00001d003495_T008 |
| orthologID_009006 | PH01000968G0230 | LOC_Os04g37720.1 | Bradi5g11860.1 | HORVU2Hr1G079060.5 | TRIUR3_26852 | Seita.7G117800.1 | Sobic.006G100000.1 | Zm00001d003500_T001 |

| | | | | | | | | |
|---|---|---|---|---|---|---|---|---|
| orthologID_008927 | PH01000438G0770 | LOC_Os04g58710.1 | Bradi5g26740.2 | HORVU2Hr1G123770.1 | TRIUR3_34181 | Seita.3G010200.1 | Sobic.006G272600.1 | Zm00001d026649_T001 |
| orthologID_008861 | PH01000001G0900 | LOC_Os04g45600.1 | Bradi5g17027.1 | HORVU2Hr1G092280.2 | TRIUR3_10110 | Seita.7G183200.1 | Sobic.006G162200.1 | Zm00001d002815_T001 |
| orthologID_009015 | PH01000152G1130 | LOC_Os04g51510.1 | Bradi5g20734.1 | HORVU2Hr1G101160.1 | TRIUR3_22756 | Seita.7G227500.1 | Sobic.006G205700.1 | Zm00001d026251_T001 |
| orthologID_008997 | PH01000037G0430 | LOC_Os04g55860.1 | Bradi5g24420.2 | HORVU2Hr1G112600.2 | TRIUR3_07915 | Seita.3G034700.1 | Sobic.006G244600.1 | Zm00001d026475_T001 |
| orthologID_008801 | PH01000349G0410 | LOC_Os04g38330.1 | Bradi5g12100.1 | HORVU2Hr1G079820.1 | TRIUR3_32639 | Seita.7G121200.1 | Sobic.006G103100.1 | Zm00001d003462_T001 |
| orthologID_009531 | PH01000030G0070 | LOC_Os04g50710.1 | Bradi3g48400.1 | HORVU2Hr1G106130.1 | TRIUR3_15992 | Seita.8G085500.1 | Sobic.006G199000.1 | Zm00001d026200_T001 |
| orthologID_008910 | PH01001279G0250 | LOC_Os04g44190.1 | Bradi5g16050.1 | HORVU2Hr1G090070.8 | TRIUR3_14009 | Seita.7G171300.1 | Sobic.006G151001.1 | Zm00001d025874_T001 |
| orthologID_009031 | PH01000304G0290 | LOC_Os04g36600.1 | Bradi5g11320.1 | HORVU2Hr1G077110.1 | TRIUR3_11056 | Seita.7G109000.1 | Sobic.006G090800.2 | Zm00001d003593_T001 |
| orthologID_008878 | PH01000020G1170 | LOC_Os04g40670.1 | Bradi5g13710.1 | HORVU2Hr1G083620.1 | TRIUR3_06971 | Seita.7G142000.1 | Sobic.006G125000.1 | Zm00001d025711_T001 |
| orthologID_006439 | PH01001223G0360 | LOC_Os01g33800.1 | Bradi2g13840.1 | HORVU2Hr1G110940.2 | TRIUR3_11510 | Seita.5G183500.1 | Sobic.006G167600.1 | Zm00001d026000_T004 |
| orthologID_009038 | PH01003280G0040 | LOC_Os04g08350.1 | Bradi5g00610.1 | HORVU2Hr1G000280.3 | TRIUR3_23795 | Seita.7G020900.1 | Sobic.006G016900.1 | Zm00001d004413_T002 |
| orthologID_008895 | PH01001692G0240 | LOC_Os04g46740.1 | Bradi5g17850.1 | HORVU2Hr1G097380.3 | TRIUR3_05264 | Seita.7G192100.1 | Sobic.006G172000.1 | Zm00001d002726_T002 |
| orthologID_008833 | PH01000023G1450 | LOC_Os04g46170.1 | Bradi5g17440.3 | HORVU2Hr1G093490.1 | TRIUR3_13877 | Seita.2G156600.1 | Sobic.006G167200.1 | Zm00001d025996_T001 |
| orthologID_008980 | PH01000265G1050 | LOC_Os04g33300.1 | Bradi5g08980.1 | HORVU2Hr1G072140.5 | TRIUR3_22240 | Seita.7G087200.1 | Sobic.006G065800.1 | Zm00001d003836_T005 |
| orthologID_008991 | PH01000087G1070 | LOC_Os04g57790.1 | Bradi5g25920.2 | HORVU2Hr1G120400.1 | TRIUR3_00809 | Seita.3G018400.1 | Sobic.006G263900.1 | Zm00001d026598_T001 |
| orthologID_009046 | PH01000561G0130 | LOC_Os04g35760.1 | Bradi5g10690.2 | HORVU2Hr1G076530.5 | TRIUR3_05260 | Seita.7G104500.1 | Sobic.006G086800.1 | Zm00001d003618_T001 |
| orthologID_008796 | PH01000593G0040 | LOC_Os04g31270.1 | Bradi5g07710.1 | HORVU2Hr1G068980.1 | TRIUR3_27084 | Seita.7G069500.1 | Sobic.006G047300.1 | Zm00001d025203_T001 |
| orthologID_008838 | PH01001910G0160 | LOC_Os04g56180.1 | Bradi5g24650.1 | HORVU2Hr1G111610.3 | TRIUR3_19383 | Seita.3G022600.1 | Sobic.006G247800.1 | Zm00001d002004_T003 |
| orthologID_009007 | PH01000505G0350 | LOC_Os04g48410.1 | Bradi5g18900.1 | HORVU2Hr1G094650.1 | TRIUR3_10120 | Seita.7G206200.1 | Sobic.006G185700.2 | Zm00001d002611_T002 |
| orthologID_009025 | PH01000384G0870 | LOC_Os04g35610.1 | Bradi5g10620.1 | HORVU2Hr1G076370.2 | TRIUR3_18453 | Seita.7G103600.1 | Sobic.006G085800.1 | Zm00001d025446_T002 |
| orthologID_008970 | PH01001942G0260 | LOC_Os04g14680.1 | Bradi5g03640.1 | HORVU2Hr1G009940.3 | TRIUR3_26505 | Seita.7G023900.1 | Sobic.006G021100.1 | Zm00001d025047_T001 |
| orthologID_008950 | PH01001151G0350 | LOC_Os04g55250.1 | Bradi5g23900.1 | HORVU2Hr1G114460.2 | TRIUR3_30074 | Seita.7G261500.1 | Sobic.006G239200.1 | Zm00001d026441_T001 |
| orthologID_008742 | PH01000875G0280 | LOC_Os04g04030.1 | Bradi5g01270.1 | HORVU2Hr1G001830.1 | TRIUR3_00562 | Seita.7G016500.1 | Sobic.006G012700.1 | Zm00001d004465_T001 |
| orthologID_010128 | PH01000597G0530 | LOC_Os04g55970.2 | Bradi5g24360.2 | HORVU2Hr1G112280.5 | TRIUR3_26191 | Seita.3G033600.1 | Sobic.006G245500.1 | Zm00001d002025_T001 |
| orthologID_008814 | PH01001172G0260 | LOC_Os04g13190.1 | Bradi5g03455.1 | HORVU2Hr1G012380.2 | TRIUR3_33928 | Seita.7G039500.1 | Sobic.006G022301.1 | Zm00001d025040_T001 |
| orthologID_008994 | PH01000041G1250 | LOC_Os04g39110.1 | Bradi5g12714.1 | HORVU2Hr1G081190.1 | TRIUR3_31020 | Seita.7G128300.1 | Sobic.006G110850.1 | Zm00001d003380_T001 |
| orthologID_008834 | PH01000598G0070 | LOC_Os04g42880.1 | Bradi5g15180.5 | HORVU2Hr1G087860.4 | TRIUR3_14774 | Seita.7G160800.1 | Sobic.006G141000.3 | Zm00001d025820_T001 |
| orthologID_009010 | PH01000603G0050 | LOC_Os04g31340.1 | Bradi5g07750.1 | HORVU2Hr1G069060.2 | TRIUR3_28926 | Seita.7G070300.1 | Sobic.006G047000.1 | Zm00001d004006_T001 |
| orthologID_009012 | PH01000239G1090 | LOC_Os04g34930.1 | Bradi5g10090.1 | HORVU2Hr1G074780.1 | TRIUR3_21258 | Seita.7G097800.1 | Sobic.006G079900.2 | Zm00001d003685_T001 |
| orthologID_009044 | PH01000300G0590 | LOC_Os04g52770.1 | Bradi5g21950.1 | HORVU2Hr1G104040.6 | TRIUR3_18209 | Seita.7G239800.1 | Sobic.006G217800.1 | Zm00001d002288_T002 |
| orthologID_008832 | PH01001566G0140 | LOC_Os04g48260.1 | Bradi5g18810.1 | HORVU2Hr1G094980.10 | TRIUR3_27083 | Seita.7G204700.1 | Sobic.006G184200.1 | Zm00001d002624_T002 |
| orthologID_008771 | PH01001543G0150 | LOC_Os04g36720.1 | Bradi5g11147.1 | HORVU2Hr1G077650.1 | TRIUR3_21493 | Seita.7G111300.1 | Sobic.006G093100.1 | Zm00001d003559_T001 |
| orthologID_008805 | PH01000118G0450 | LOC_Os04g42490.1 | Bradi5g14887.2 | HORVU2Hr1G087040.2 | TRIUR3_04268 | Seita.7G157600.1 | Sobic.006G138600.1 | Zm00001d003109_T002 |
| orthologID_009047 | PH01000886G0120 | LOC_Os04g32270.1 | Bradi5g08370.1 | HORVU2Hr1G070670.2 | TRIUR3_29441 | Seita.7G077100.1 | Sobic.006G054500.1 | Zm00001d025252_T001 |
| orthologID_008780 | PH01000467G0640 | LOC_Os04g42900.1 | Bradi5g15190.1 | HORVU2Hr1G087890.3 | TRIUR3_15228 | Seita.7G160900.1 | Sobic.006G141400.1 | Zm00001d025821_T001 |
| orthologID_009032 | PH01000042G1900 | LOC_Os04g45610.1 | Bradi5g17050.7 | HORVU2Hr1G092340.3 | TRIUR3_16404 | Seita.7G183400.1 | Sobic.006G162400.1 | Zm00001d002812_T002 |
| orthologID_008935 | PH01001023G0060 | LOC_Os04g58020.1 | Bradi5g26110.2 | HORVU2Hr1G119610.3 | TRIUR3_23521 | Seita.3G016200.1 | Sobic.006G266600.1 | Zm00001d001837_T001 |
| orthologID_008767 | PH01000505G0760 | LOC_Os04g18090.1 | Bradi5g19000.1 | HORVU2Hr1G094400.5 | TRIUR3_18738 | Seita.7G207300.1 | Sobic.006G186700.1 | Zm00001d002601_T001 |
| orthologID_008953 | PH01000087G1640 | LOC_Os04g57500.1 | Bradi5g25667.1 | HORVU2Hr1G121250.11 | TRIUR3_03020 | Seita.3G033200.1 | Sobic.006G260800.2 | Zm00001d001896_T002 |
| orthologID_009008 | PH01000030G0890 | LOC_Os04g49990.1 | Bradi5g19900.4 | HORVU2Hr1G105800.13 | TRIUR3_07266 | Seita.7G217900.1 | Sobic.006G196900.2 | Zm00001d026036_T001 |
| orthologID_008957 | PH01000044G2040 | LOC_Os04g58420.1 | Bradi5g26435.1 | HORVU2Hr1G118780.6 | TRIUR3_06368 | Seita.3G012900.1 | Sobic.006G270300.2 | Zm00001d001804_T005 |
| orthologID_008775 | PH01000000G2070 | LOC_Os04g48870.1 | Bradi5g19120.1 | HORVU2Hr1G094220.1 | TRIUR3_22452 | Seita.7G208900.1 | Sobic.006G188100.1 | Zm00001d002590_T001 |
| orthologID_008746 | PH01000825G0330 | LOC_Os04g35490.1 | Bradi5g10460.2 | HORVU2Hr1G075930.2 | TRIUR3_20741 | Seita.7G101900.1 | Sobic.006G084100.1 | Zm00001d025434_T001 |
| orthologID_008798 | PH01000067G1350 | LOC_Os04g18790.1 | Bradi5g04730.2 | HORVU2Hr1G011810.1 | TRIUR3_18324 | Seita.7G041000.1 | Sobic.006G033500.1 | Zm00001d004165_T001 |
| orthologID_008850 | PH01000467G0080 | LOC_Os04g42650.1 | Bradi5g15040.1 | HORVU2Hr1G087510.1 | TRIUR3_08820 | Seita.7G159100.1 | Sobic.006G139600.1 | Zm00001d025807_T001 |
| orthologID_008988 | PH01000222G0280 | LOC_Os02g18810.1 | Bradi5g10954.1 | HORVU2Hr1G076970.5 | TRIUR3_06830 | Seita.7G108300.1 | Sobic.006G089900.1 | Zm00001d025468_T001 |
| orthologID_008867 | PH01000300G0160 | LOC_Os04g52580.1 | Bradi5g21850.10 | HORVU2Hr1G098550.6 | TRIUR3_07059 | Seita.7G237200.1 | Sobic.006G215300.1 | Zm00001d002314_T001 |
| orthologID_008855 | PH01001914G0290 | LOC_Os04g53360.1 | Bradi5g22343.1 | HORVU2Hr1G106480.7 | TRIUR3_15512 | Seita.7G244400.1 | Sobic.006G221400.1 | Zm00001d026345_T001 |
| orthologID_008807 | PH01000224G0980 | LOC_Os04g51570.1 | Bradi5g20857.1 | HORVU2Hr1G101070.1 | TRIUR3_18619 | Seita.7G228000.1 | Sobic.006G206600.3 | Zm00001d002403_T001 |
| orthologID_008965 | PH01000890G0540 | LOC_Os04g52180.1 | Bradi5g21360.1 | HORVU2Hr1G099610.3 | TRIUR3_13006 | Seita.7G233200.1 | Sobic.006G211200.1 | Zm00001d002352_T001 |

| | | | | | | | | |
|---|---|---|---|---|---|---|---|---|
| orthologID_008874 | PH01001914G0170 | LOC_Os04g53490.1 | Bradi5g22510.1 | HORVU2Hr1G106880.1 | TRIUR3_10099 | Seita.7G245800.1 | Sobic.006G222700.1 | Zm00001d002240_T001 |
| orthologID_008919 | PH01000378G0910 | LOC_Os04g33060.1 | Bradi5g08820.1 | HORVU2Hr1G071860.2 | TRIUR3_13643 | Seita.7G084700.1 | Sobic.006G063900.1 | Zm00001d000123_T003 |
| orthologID_006240 | PH01000016G0600 | LOC_Os04g32350.1 | Bradi2g34200.2 | HORVU1Hr1G022720.1 | TRIUR3_08966 | Seita.7G078000.1 | Sobic.006G055300.1 | Zm00001d003917_T007 |
| orthologID_008882 | PH01002245G0110 | LOC_Os04g53740.1 | Bradi5g22770.1 | HORVU2Hr1G107990.1 | TRIUR3_12014 | Seita.7G248600.1 | Sobic.006G226100.2 | Zm00001d026368_T001 |
| orthologID_008762 | PH01000265G0640 | LOC_Os04g33450.1 | Bradi5g09130.1 | HORVU2Hr1G072420.11 | TRIUR3_32840 | Seita.7G088200.1 | Sobic.006G066500.1 | Zm00001d025338_T001 |
| orthologID_008977 | PH01000030G0270 | LOC_Os04g50208.1 | Bradi5g20020.2 | HORVU2Hr1G106090.14 | TRIUR3_15767 | Seita.7G219400.1 | Sobic.006G198400.2 | Zm00001d026192_T006 |
| orthologID_008871 | PH01000041G1920 | LOC_Os04g38810.1 | Bradi5g12470.1 | HORVU2Hr1G080560.3 | TRIUR3_28173 | Seita.7G125600.1 | Sobic.006G108000.1 | Zm00001d003406_T002 |
| orthologID_006874 | PH01004707G0070 | LOC_Os01g18880.2 | Bradi2g11270.4 | HORVU2Hr1G086870.3 | TRIUR3_09998 | Seita.5G030400.1 | Sobic.006G263000.1 | Zm00001d001869_T001 |
| orthologID_007137 | PH01000023G0560 | LOC_Os04g46570.1 | Bradi5g17710.2 | HORVU2Hr1G097570.20 | TRIUR3_18244 | Seita.7G189900.1 | Sobic.006G170900.1 | Zm00001d002742_T002 |
| orthologID_008826 | PH01001603G0100 | LOC_Os04g37670.1 | Bradi5g11790.2 | HORVU2Hr1G078890.16 | TRIUR3_07452 | Seita.7G117100.1 | Sobic.006G098200.1 | Zm00001d003508_T001 |
| orthologID_009029 | PH01001142G0170 | LOC_Os04g39670.1 | Bradi5g13212.2 | HORVU2Hr1G082000.1 | TRIUR3_18993 | Seita.7G134400.1 | Sobic.006G116500.2 | Zm00001d003322_T007 |
| orthologID_009059 | PH01002245G0200 | LOC_Os04g53700.1 | Bradi5g22730.1 | HORVU2Hr1G107860.15 | TRIUR3_08639 | Seita.7G247900.1 | Sobic.006G225400.1 | Zm00001d002185_T003 |
| orthologID_008998 | PH01000044G1350 | LOC_Os04g58860.1 | Bradi5g26600.1 | HORVU2Hr1G123370.1 | TRIUR3_09712 | Seita.3G008200.1 | Sobic.006G274400.1 | Zm00001d026664_T001 |
| orthologID_009065 | PH01003264G0020 | LOC_Os04g57480.1 | Bradi5g25650.1 | HORVU2Hr1G122000.17 | TRIUR3_05718 | Seita.3G032900.1 | Sobic.006G260600.1 | Zm00001d001897_T001 |
| orthologID_007617 | PH01000023G1630 | LOC_Os04g46120.1 | Bradi1g68150.1 | HORVU2Hr1G093430.2 | TRIUR3_13879 | Seita.2G157400.1 | Sobic.006G166700.1 | Zm00001d002773_T001 |
| orthologID_009058 | PH01001315G0360 | LOC_Os04g46930.1 | Bradi5g17980.1 | HORVU3Hr1G068730.1 | TRIUR3_18936 | Seita.7G193800.1 | Sobic.006G173600.2 | Zm00001d002711_T001 |
| orthologID_008758 | PH01003624G0170 | LOC_Os04g31690.1 | Bradi5g07870.2 | HORVU2Hr1G069610.4 | TRIUR3_24216 | Seita.7G072600.1 | Sobic.006G050400.2 | Zm00001d003975_T001 |
| orthologID_009394 | PH01000104G0650 | LOC_Os08g37940.1 | Bradi3g38777.2 | HORVU2Hr1G085300.10 | TRIUR3_06872 | Seita.6G190000.1 | Sobic.006G263800.1 | Zm00001d001860_T001 |
| orthologID_009035 | PH01000558G0430 | LOC_Os04g53550.1 | Bradi5g22580.1 | HORVU2Hr1G106990.2 | TRIUR3_00420 | Seita.5G165700.1 | Sobic.006G223500.1 | Zm00001d026352_T001 |
| orthologID_008899 | PH01000041G2180 | LOC_Os04g38700.2 | Bradi5g12400.2 | HORVU2Hr1G080450.1 | TRIUR3_09914 | Seita.7G124700.1 | Sobic.006G107200.1 | Zm00001d003415_T002 |
| orthologID_008924 | PH01000875G0180 | LOC_Os04g03990.1 | Bradi5g01340.1 | HORVU2Hr1G001690.1 | TRIUR3_00558 | Seita.7G015900.1 | Sobic.006G012300.1 | Zm00001d004468_T004 |
| orthologID_008883 | PH01000016G0110 | LOC_Os04g32150.1 | Bradi5g08360.1 | HORVU3Hr1G094270.1 | TRIUR3_11859 | Seita.7G076900.1 | Sobic.006G054300.1 | Zm00001d025249_T001 |
| orthologID_008972 | PH01000432G0200 | LOC_Os04g41040.1 | Bradi5g13967.1 | HORVU2Hr1G084300.6 | TRIUR3_06084 | Seita.7G144900.1 | Sobic.006G127500.1 | Zm00001d003242_T001 |
| orthologID_008841 | PH01000087G1550 | LOC_Os04g57540.1 | Bradi5g25720.4 | HORVU2Hr1G121070.1 | TRIUR3_11270 | Seita.3G020700.1 | Sobic.006G261300.4 | Zm00001d026584_T001 |
| orthologID_008782 | PH01000825G0150 | LOC_Os04g35380.1 | Bradi5g10400.1 | HORVU2Hr1G075710.17 | TRIUR3_31830 | Seita.7G101400.1 | Sobic.006G083600.1 | Zm00001d003652_T004 |
| orthologID_008955 | PH01000850G0530 | LOC_Os04g32930.1 | Bradi5g08550.10 | HORVU2Hr1G071670.10 | TRIUR3_14532 | Seita.7G082900.1 | Sobic.006G062200.4 | Zm00001d025304_T002 |
| orthologID_008678 | PH01000532G0700 | LOC_Os03g14040.1 | Bradi1g67990.5 | HORVU4Hr1G062920.2 | TRIUR3_07107 | Seita.1G118300.1 | Sobic.006G103400.1 | Zm00001d003459_T001 |
| orthologID_009056 | PH01000696G0320 | LOC_Os04g34600.1 | Bradi5g10027.1 | HORVU2Hr1G074770.3 | TRIUR3_26722 | Seita.7G097200.1 | Sobic.006G078500.1 | Zm00001d003712_T001 |
| orthologID_008803 | PH01000001G0140 | LOC_Os04g45821.1 | Bradi5g17187.1 | HORVU2Hr1G092730.1 | TRIUR3_29405 | Seita.7G184700.1 | Sobic.006G164000.1 | Zm00001d025966_T001 |
| orthologID_008954 | PH01000875G0370 | LOC_Os04g04320.1 | Bradi5g01220.2 | HORVU2Hr1G002050.1 | TRIUR3_11170 | Seita.7G016900.1 | Sobic.006G013100.1 | Zm00001d004460_T004 |
| orthologID_008923 | PH01000000G4930 | LOC_Os04g47680.1 | Bradi5g18450.1 | HORVU2Hr1G095910.4 | TRIUR3_29162 | Seita.7G199900.1 | Sobic.006G179700.1 | Zm00001d026124_T001 |
| orthologID_009019 | PH01000016G1570 | LOC_Os04g32660.1 | Bradi5g08390.1 | HORVU2Hr1G071350.1 | TRIUR3_30560 | Seita.7G080500.1 | Sobic.006G058900.1 | Zm00001d025286_T001 |
| orthologID_009004 | PH01001619G0270 | LOC_Os04g39950.1 | Bradi5g13290.1 | HORVU2Hr1G082220.1 | TRIUR3_32080 | Seita.7G135500.1 | Sobic.006G118000.1 | Zm00001d003314_T001 |
| orthologID_008141 | PH01000064G1610 | LOC_Os07g43540.1 | Bradi1g20550.1 | HORVU2Hr1G028500.1 | TRIUR3_07116 | Seita.2G400500.1 | Sobic.002G385400.1 | Zm00001d022226_T002 |
| orthologID_007844 | PH01000288G0350 | LOC_Os07g07974.1 | Bradi1g55710.1 | HORVU4Hr1G004280.3 | TRIUR3_11462 | Seita.2G050700.1 | Sobic.002G050700.2 | Zm00001d018930_T003 |
| orthologID_007190 | PH01000894G0480 | LOC_Os09g36030.1 | Bradi4g35820.1 | HORVU5Hr1G080980.4 | TRIUR3_11771 | Seita.2G283100.1 | Sobic.002G271800.1 | Zm00001d006182_T001 |
| orthologID_008389 | PH01003073G0070 | LOC_Os07g31720.1 | Bradi1g26850.1 | HORVU4Hr1G090370.2 | TRIUR3_35182 | Seita.2G319800.1 | Sobic.002G307600.1 | Zm00001d021529_T002 |
| orthologID_008695 | PH01002295G0240 | LOC_Os07g34598.1 | Bradi1g25830.1 | HORVU2Hr1G045220.1 | TRIUR3_21572 | Seita.7G294600.1 | Sobic.002G324300.1 | Zm00001d021669_T008 |
| orthologID_007545 | PH01004610G0130 | LOC_Os09g38510.1 | Bradi4g37670.1 | HORVU5Hr1G088530.7 | TRIUR3_05994 | Seita.2G304800.1 | Sobic.002G294000.2 | Zm00001d006327_T001 |
| orthologID_007798 | PH01002743G0130 | LOC_Os07g41340.1 | Bradi1g22010.1 | HORVU2Hr1G033090.1 | TRIUR3_24112 | Seita.2G382800.1 | Sobic.002G368500.2 | Zm00001d022084_T001 |
| orthologID_006191 | PH01000005G4060 | LOC_Os09g07460.1 | Bradi2g38270.1 | HORVU5Hr1G051310.1 | TRIUR3_20300 | Seita.2G154400.1 | Sobic.002G082000.1 | Zm00001d019221_T001 |
| orthologID_008631 | PH01003401G0090 | LOC_Os07g11110.1 | Bradi1g54030.1 | HORVU6Hr1G004860.10 | TRIUR3_21293 | Seita.2G082400.1 | Sobic.002G077000.1 | Zm00001d019177_T001 |
| orthologID_006549 | PH01001166G0090 | LOC_Os01g37510.1 | Bradi2g41027.1 | HORVU3Hr1G047870.5 | TRIUR3_30388 | Seita.5G195300.1 | Sobic.002G396200.1 | Zm00001d022337_T002 |
| orthologID_007311 | PH01003503G0210 | LOC_Os05g24560.1 | Bradi4g43270.1 | HORVU5Hr1G043560.3 | TRIUR3_24889 | Seita.7G335300.1 | Sobic.002G315700.3 | Zm00001d021614_T001 |
| orthologID_008219 | PH01000490G0650 | LOC_Os07g40770.1 | Bradi1g22566.3 | HORVU2Hr1G034100.7 | TRIUR3_14270 | Seita.2G378200.1 | Sobic.002G364000.1 | Zm00001d006763_T001 |
| orthologID_008339 | PH01000605G0380 | LOC_Os07g32480.1 | Bradi1g26580.1 | HORVU0Hr1G039990.2 | TRIUR3_33854 | Seita.2G326500.1 | Sobic.002G313700.1 | Zm00001d021588_T002 |
| orthologID_008165 | PH01001650G0130 | LOC_Os07g41650.1 | Bradi1g21810.1 | HORVU2Hr1G032220.4 | TRIUR3_22430 | Seita.2G384400.1 | Sobic.002G370300.1 | Zm00001d022104_T001 |
| orthologID_007760 | PH01001750G0080 | LOC_Os07g10890.1 | Bradi1g54180.1 | HORVU2Hr1G055160.1 | TRIUR3_31142 | Seita.2G081300.1 | Sobic.002G076100.2 | Zm00001d005244_T002 |
| orthologID_008460 | PH01000591G0710 | LOC_Os07g45300.1 | Bradi1g19550.2 | HORVU2Hr1G024840.7 | TRIUR3_10116 | Seita.2G413400.1 | Sobic.002G398700.2 | Zm00001d022352_T001 |
| orthologID_010108 | PH01001437G0440 | LOC_Os07g30970.1 | Bradi1g27120.1 | HORVU4Hr1G044040.16 | TRIUR3_21918 | Seita.2G319600.1 | Sobic.002G306900.1 | Zm00001d021526_T001 |

| | | | | | | | | |
|---|---|---|---|---|---|---|---|---|
| orthologID_007085 | PH01001123G0180 | LOC_Os09g20420.1 | Bradi4g29050.1 | HORVU5Hr1G058410.2 | TRIUR3_22325 | Seita.2G188400.1 | Sobic.002G184800.1 | Zm00001d005661_T001 |
| orthologID_008348 | PH01001114G0220 | LOC_Os07g28840.1 | Bradi1g28267.2 | HORVU2Hr1G104570.6 | TRIUR3_03471 | Seita.2G310400.1 | Sobic.002G299100.1 | Zm00001d021346_T002 |
| orthologID_007653 | PH01003181G0090 | LOC_Os07g37310.1 | Bradi1g24830.1 | HORVU2Hr1G041110.1 | TRIUR3_32389 | Seita.2G352000.1 | Sobic.002G338400.1 | Zm00001d021774_T001 |
| orthologID_008138 | PH01000471G0590 | LOC_Os07g13600.1 | Bradi1g53567.1 | HORVU0Hr1G003010.1 | TRIUR3_22188 | Seita.2G092300.1 | Sobic.002G084900.1 | Zm00001d005177_T001 |
| orthologID_008580 | PH01002295G0130 | LOC_Os07g34390.1 | Bradi1g25900.2 | HORVU2Hr1G045370.3 | TRIUR3_17320 | Seita.2G336700.1 | Sobic.002G323300.1 | Zm00001d021662_T002 |
| orthologID_008200 | PH01004068G0170 | LOC_Os07g49300.1 | Bradi1g16590.2 | HORVU2Hr1G013900.17 | TRIUR3_02765 | Seita.2G443200.1 | Sobic.002G430000.2 | Zm00001d022584_T001 |
| orthologID_008313 | PH01000620G0190 | LOC_Os07g28470.1 | Bradi1g28353.1 | HORVU2Hr1G016660.20 | TRIUR3_01434 | Seita.2G308800.1 | Sobic.002G297500.1 | Zm00001d021365_T001 |
| orthologID_007691 | PH01001912G0330 | LOC_Os07g06600.1 | Bradi1g57120.1 | HORVU2Hr1G059940.1 | TRIUR3_27143 | Seita.2G037600.1 | Sobic.002G039100.1 | Zm00001d018827_T001 |
| orthologID_007104 | PH01000883G0280 | LOC_Os09g25860.1 | Bradi4g30760.1 | HORVU5Hr1G063810.2 | TRIUR3_22195 | Seita.2G211700.1 | Sobic.002G208000.1 | Zm00001d020558_T001 |
| orthologID_007187 | PH01000976G0450 | LOC_Os09g19980.1 | Bradi4g28910.1 | HORVU5Hr1G057760.1 | TRIUR3_09649 | Seita.2G187100.1 | Sobic.002G183300.1 | Zm00001d020256_T001 |
| orthologID_008869 | PH01000541G0170 | LOC_Os07g11290.1 | Bradi5g25284.2 | HORVU7Hr1G094990.2 | TRIUR3_12746 | Seita.2G083000.1 | Sobic.002G347700.2 | Zm00001d021873_T001 |
| orthologID_007770 | PH01000811G0110 | LOC_Os07g48720.1 | Bradi1g17280.1 | HORVU2Hr1G016800.2 | TRIUR3_14501 | Seita.2G439400.1 | Sobic.002G424900.1 | Zm00001d007197_T001 |
| orthologID_007365 | PH01003299G0050 | LOC_Os09g25310.1 | Bradi4g30450.1 | HORVU5Hr1G065220.4 | TRIUR3_04550 | Seita.2G207300.1 | Sobic.002G204000.1 | Zm00001d020506_T001 |
| orthologID_007783 | PH01000363G0360 | LOC_Os07g37830.1 | Bradi1g24500.2 | HORVU2Hr1G039880.8 | TRIUR3_21334 | Seita.2G355000.1 | Sobic.002G341500.1 | Zm00001d006610_T001 |
| orthologID_007163 | PH01000770G0620 | LOC_Os09g31380.1 | Bradi4g33727.2 | HORVU5Hr1G073330.1 | TRIUR3_35209 | Seita.2G253700.1 | Sobic.002G247200.1 | Zm00001d006026_T001 |
| orthologID_008683 | PH01000591G0700 | LOC_Os07g45310.1 | Bradi1g19540.1 | HORVU2Hr1G024900.5 | TRIUR3_25773 | Seita.2G413500.1 | Sobic.002G398800.1 | Zm00001d022353_T001 |
| orthologID_007284 | PH01001573G0270 | LOC_Os09g31482.1 | Bradi4g33870.2 | HORVU5Hr1G073680.3 | TRIUR3_00890 | Seita.2G255600.1 | Sobic.002G249500.2 | Zm00001d020954_T001 |
| orthologID_007431 | PH01000978G0270 | LOC_Os09g39400.1 | Bradi4g38450.1 | HORVU4Hr1G001720.1 | TRIUR3_12780 | Seita.2G132700.1 | Sobic.002G134001.1 | Zm00001d005344_T001 |
| orthologID_008414 | PH01000451G0650 | LOC_Os07g30170.1 | Bradi1g27460.1 | HORVU5Hr1G076080.1 | TRIUR3_28431 | Seita.2G316000.1 | Sobic.002G304000.1 | Zm00001d021506_T001 |
| orthologID_007254 | PH01000063G0560 | LOC_Os09g33490.1 | Bradi4g34800.1 | HORVU5Hr1G077010.4 | TRIUR3_21221 | Seita.2G268500.1 | Sobic.002G259600.1 | Zm00001d006106_T001 |
| orthologID_007738 | PH01000482G0720 | LOC_Os07g24230.1 | Bradi1g29040.1 | HORVU3Hr1G013630.1 | TRIUR3_17908 | Seita.2G116100.1 | Sobic.002G119100.1 | Zm00001d019510_T001 |
| orthologID_007742 | PH01001527G0030 | LOC_Os07g08210.1 | Bradi1g55520.1 | HORVU6Hr1G060690.1 | TRIUR3_02967 | Seita.2G054600.1 | Sobic.002G052900.2 | Zm00001d018946_T001 |
| orthologID_007209 | PH01000533G0590 | LOC_Os09g26870.1 | Bradi4g31300.1 | HORVU5Hr1G062110.4 | TRIUR3_16582 | Seita.2G218300.1 | Sobic.002G215600.1 | Zm00001d020622_T002 |
| orthologID_007103 | PH01000788G0390 | LOC_Os09g39690.1 | Bradi4g38750.1 | HORVU4Hr1G000090.6 | TRIUR3_00427 | Seita.2G136300.1 | Sobic.002G137400.1 | Zm00001d019765_T001 |
| orthologID_007379 | PH01000319G0220 | LOC_Os09g29060.1 | Bradi4g32520.1 | HORVU5Hr1G069860.1 | TRIUR3_17547 | Seita.2G235800.1 | Sobic.002G230500.1 | Zm00001d005924_T002 |
| orthologID_007173 | PH01002168G0010 | LOC_Os09g11230.1 | Bradi4g27240.1 | HORVU5Hr1G053560.4 | TRIUR3_27787 | Seita.2G161900.1 | Sobic.002G157300.1 | Zm00001d005110_T001 |
| orthologID_008671 | PH01000751G0390 | LOC_Os07g12640.1 | Bradi1g53747.1 | HORVU4Hr1G075040.19 | TRIUR3_08658 | Seita.2G088200.1 | Sobic.002G081500.1 | Zm00001d019218_T001 |
| orthologID_008255 | PH01003216G0040 | LOC_Os07g08880.1 | Bradi1g55090.1 | HORVU2Hr1G054100.3 | TRIUR3_19030 | Seita.2G062000.1 | Sobic.002G058300.3 | Zm00001d019039_T002 |
| orthologID_007864 | PH01000068G1050 | LOC_Os07g44060.1 | Bradi1g20280.2 | HORVU2Hr1G027680.1 | TRIUR3_10946 | Seita.2G403900.1 | Sobic.002G388400.1 | Zm00001d022254_T002 |
| orthologID_007338 | PH01004612G0090 | LOC_Os09g24710.2 | Bradi4g30210.1 | HORVU5Hr1G065890.1 | TRIUR3_28763 | Seita.2G204200.1 | Sobic.002G200900.1 | Zm00001d005762_T002 |
| orthologID_007210 | PH01000885G0520 | LOC_Os09g38755.1 | Bradi4g37960.1 | HORVU5Hr1G089510.1 | TRIUR3_10602 | Seita.2G307700.1 | Sobic.002G296400.1 | Zm00001d021372_T001 |
| orthologID_007983 | PH01001437G0340 | LOC_Os07g30980.1 | Bradi1g27110.1 | HORVU2Hr1G051300.27 | TRIUR3_20998 | Seita.2G322900.1 | Sobic.002G310300.1 | Zm00001d006443_T001 |
| orthologID_008555 | PH01000553G0200 | LOC_Os07g31260.1 | Bradi1g27080.1 | HORVU2Hr1G051560.3 | TRIUR3_34480 | Seita.2G322800.1 | Sobic.002G310200.1 | Zm00001d000436_T001 |
| orthologID_008152 | PH01000205G0180 | ChrSy.fgenesh.mRNA.49 | Bradi1g03550.1 | HORVU5Hr1G114090.4 | TRIUR3_27202 | Seita.4G130600.1 | Sobic.002G009300.1 | Zm00001d007945_T004 |
| orthologID_008635 | PH01000068G0720 | LOC_Os07g44310.1 | Bradi1g20140.1 | HORVU2Hr1G027160.15 | TRIUR3_09162 | Seita.2G405700.1 | Sobic.002G390300.1 | Zm00001d006942_T007 |
| orthologID_008548 | PH01005139G0110 | LOC_Os07g01850.1 | Bradi1g59260.1 | HORVU2Hr1G063840.2 | TRIUR3_24558 | Seita.2G142300.1 | Sobic.002G142600.1 | Zm00001d019699_T002 |
| orthologID_008609 | PH01000068G1300 | LOC_Os07g43870.3 | Bradi1g20410.2 | HORVU2Hr1G028260.5 | TRIUR3_16919 | Seita.2G401800.1 | Sobic.002G386600.1 | Zm00001d007009_T001 |
| orthologID_008449 | PH01000364G0120 | LOC_Os07g47300.1 | Bradi1g18570.1 | HORVU2Hr1G020240.1 | TRIUR3_12172 | Seita.2G427500.1 | Sobic.002G409900.1 | Zm00001d007117_T001 |
| orthologID_007239 | PH01000085G1190 | LOC_Os07g23720.1 | Bradi4g27310.2 | HORVU7Hr1G038140.4 | TRIUR3_12790 | Seita.2G113900.1 | Sobic.002G117200.2 | Zm00001d019492_T001 |
| orthologID_007054 | PH01000063G0270 | LOC_Os09g32988.1 | Bradi4g34770.1 | HORVU5Hr1G076760.1 | TRIUR3_00757 | Seita.2G267600.1 | Sobic.002G259200.1 | Zm00001d021063_T001 |
| orthologID_009496 | PH01005166G0070 | LOC_Os07g01410.1 | Bradi3g55850.1 | HORVU7Hr1G018530.3 | TRIUR3_04133 | Seita.J003100.1 | Sobic.002G003500.1 | Zm00001d018619_T001 |
| orthologID_007132 | PH01001386G0150 | LOC_Os09g27240.2 | Bradi4g31560.2 | HORVU5Hr1G066630.3 | TRIUR3_26455 | Seita.2G222200.1 | Sobic.002G218700.1 | Zm00001d005834_T001 |
| orthologID_008274 | PH01002108G0260 | LOC_Os07g17120.1 | Bradi1g53000.2 | HORVU3Hr1G016340.3 | TRIUR3_00523 | Seita.2G102400.1 | Sobic.002G103900.1 | Zm00001d019345_T001 |
| orthologID_009175 | PH01002479G0170 | LOC_Os07g18750.1 | Bradi3g58060.1 | HORVU4Hr1G082600.1 | TRIUR3_09733 | Seita.2G106500.1 | Sobic.002G109100.1 | Zm00001d005146_T001 |
| orthologID_007928 | PH01000064G2290 | LOC_Os07g44020.1 | Bradi1g20310.1 | HORVU2Hr1G027750.5 | TRIUR3_22879 | Seita.2G403400.1 | Sobic.002G388000.3 | Zm00001d022250_T001 |
| orthologID_007521 | PH01002194G0200 | LOC_Os12g21930.1 | Bradi4g08090.5 | HORVU2Hr1G106290.7 | TRIUR3_16619 | Seita.2G322600.1 | Sobic.002G310000.3 | Zm00001d000435_T001 |
| orthologID_007081 | PH01000070G1540 | LOC_Os09g32280.1 | Bradi4g34180.1 | HORVU5Hr1G074940.3 | TRIUR3_08399 | Seita.2G260400.1 | Sobic.002G253200.1 | Zm00001d006054_T001 |
| orthologID_008646 | PH01001776G0230 | LOC_Os07g25500.1 | Bradi1g28980.3 | HORVU1Hr1G020190.3 | TRIUR3_23409 | Seita.2G117400.1 | Sobic.002G120200.1 | Zm00001d019525_T001 |
| orthologID_007526 | PH01004069G0020 | LOC_Os09g17680.1 | Bradi4g28380.1 | HORVU5Hr1G056620.2 | TRIUR3_24713 | Seita.2G182300.1 | Sobic.002G176800.1 | Zm00001d005632_T001 |
| orthologID_007639 | PH01000068G1510 | LOC_Os07g43530.2 | Bradi1g20560.2 | HORVU2Hr1G028530.3 | TRIUR3_07115 | Seita.2G400400.1 | Sobic.002G385300.1 | Zm00001d022225_T001 |

| | | | | | | | | |
|---|---|---|---|---|---|---|---|---|
| orthologID_008560 | PH01000056G0470 | LOC_Os07g46330.1 | Bradi1g19250.1 | HORVU2Hr1G023480.3 | TRIUR3_33338 | Seita.2G415600.1 | Sobic.002G401000.1 | Zm00001d022371_T001 |
| orthologID_008092 | PH01000204G0690 | LOC_Os07g14700.1 | Bradi1g53290.1 | HORVU2Hr1G108350.1 | TRIUR3_19784 | Seita.2G098100.1 | Sobic.002G092700.1 | Zm00001d019304_T001 |
| orthologID_007180 | PH01000073G0710 | LOC_Os07g39910.1 | Bradi1g22990.1 | HORVU2Hr1G035300.1 | TRIUR3_22696 | Seita.2G373100.1 | Sobic.002G358900.3 | Zm00001d021980_T001 |
| orthologID_007552 | PH01000314G0770 | LOC_Os09g35970.1 | Bradi4g35867.2 | HORVU5Hr1G081010.1 | TRIUR3_19186 | Seita.2G283600.1 | Sobic.002G272500.1 | Zm00001d006185_T001 |
| orthologID_008121 | PH01001314G0350 | LOC_Os07g32800.2 | Bradi1g26400.4 | HORVU2Hr1G046860.2 | TRIUR3_34617 | Seita.2G329800.1 | Sobic.002G315900.1 | Zm00001d006474_T002 |
| orthologID_008574 | PH01001895G0240 | LOC_Os07g38830.1 | Bradi1g23675.6 | HORVU2Hr1G037020.1 | TRIUR3_15101 | Seita.2G365600.1 | Sobic.002G351200.2 | Zm00001d021899_T002 |
| orthologID_008538 | PH01003175G0220 | LOC_Os07g38490.1 | Bradi1g23980.1 | HORVU2Hr1G038030.1 | TRIUR3_21474 | Seita.2G362400.1 | Sobic.002G347900.1 | Zm00001d006640_T001 |
| orthologID_007880 | PH01000553G0540 | LOC_Os07g31460.1 | Bradi1g26930.7 | HORVU2Hr1G048680.2 | TRIUR3_29272 | Seita.2G320700.1 | Sobic.002G308500.1 | Zm00001d006422_T006 |
| orthologID_007501 | PH01001159G0360 | LOC_Os09g29760.1 | Bradi4g32880.2 | HORVU5Hr1G070460.1 | TRIUR3_24324 | Seita.2G240900.1 | Sobic.002G234700.1 | Zm00001d005959_T001 |
| orthologID_007799 | PH01000064G0020 | LOC_Os07g42820.1 | Bradi1g21210.6 | HORVU1Hr1G014610.3 | TRIUR3_29925 | Seita.2G393300.1 | Sobic.002G378300.1 | Zm00001d022172_T001 |
| orthologID_007953 | PH01001807G0490 | LOC_Os07g27180.1 | Bradi1g28660.1 | HORVU4Hr1G052640.5 | TRIUR3_16990 | Seita.2G124700.1 | Sobic.002G126800.1 | Zm00001d005403_T001 |
| orthologID_007686 | PH01000666G0640 | LOC_Os07g41730.2 | Bradi1g21755.2 | HORVU2Hr1G032060.4 | TRIUR3_25869 | Seita.2G385400.1 | Sobic.002G370900.1 | Zm00001d006836_T002 |
| orthologID_008604 | PH01001260G0290 | LOC_Os03g56960.1 | Bradi1g06650.2 | HORVU5Hr1G102230.1 | TRIUR3_20577 | Seita.9G237800.1 | Sobic.002G336800.1 | Zm00001d004983_T001 |
| orthologID_008302 | PH01000096G0970 | LOC_Os07g19494.1 | Bradi1g52600.1 | HORVU2Hr1G053240.2 | TRIUR3_28053 | Seita.2G108200.1 | Sobic.002G110500.1 | Zm00001d019423_T002 |
| orthologID_007996 | PH01001273G0220 | LOC_Os07g29420.4 | Bradi1g28100.8 | HORVU5Hr1G099240.1 | TRIUR3_35111 | Seita.2G312200.1 | Sobic.002G300500.1 | Zm00001d021335_T003 |
| orthologID_007208 | PH01002328G0120 | LOC_Os09g21710.1 | Bradi4g29530.1 | HORVU5Hr1G059900.1 | TRIUR3_25782 | Seita.2G195200.1 | Sobic.002G192000.2 | Zm00001d005698_T001 |
| orthologID_007130 | PH01000332G0920 | LOC_Os09g36976.1 | Bradi4g36650.1 | HORVU5Hr1G084390.4 | TRIUR3_31812 | Seita.2G291900.1 | Sobic.002G281300.1 | Zm00001d022604_T001 |
| orthologID_009279 | PH01000732G0750 | LOC_Os02g55570.1 | Bradi3g54300.6 | HORVU6Hr1G086600.5 | TRIUR3_32010 | Seita.2G080300.1 | Sobic.002G074500.12 | Zm00001d019148_T002 |
| orthologID_008450 | PH01002992G0200 | LOC_Os07g47820.1 | Bradi1g17950.2 | HORVU2Hr1G018720.7 | TRIUR3_02387 | Seita.2G430500.1 | Sobic.002G415900.1 | Zm00001d007158_T014 |
| orthologID_007048 | PH01002176G0070 | LOC_Os09g20970.1 | Bradi4g29330.1 | HORVU5Hr1G059170.1 | TRIUR3_33687 | Seita.2G192400.1 | Sobic.002G188000.1 | Zm00001d020322_T001 |
| orthologID_007155 | PH01000191G1290 | LOC_Os09g34190.1 | Bradi4g35336.1 | HORVU5Hr1G079690.2 | TRIUR3_18585 | Seita.2G275400.1 | Sobic.002G265300.1 | Zm00001d006133_T001 |
| orthologID_007269 | PH01001244G0440 | LOC_Os09g12710.1 | Bradi4g27580.2 | HORVU5Hr1G054250.4 | TRIUR3_24922 | Seita.2G168100.1 | Sobic.002G160700.1 | Zm00001d020011_T001 |
| orthologID_007874 | PH01000742G0250 | LOC_Os07g33660.1 | Bradi1g26070.2 | HORVU2Hr1G045620.2 | TRIUR3_10983 | Seita.2G334500.1 | Sobic.002G320700.1 | Zm00001d021645_T001 |
| orthologID_008533 | PH01000363G0900 | LOC_Os07g38060.1 | Bradi1g24297.1 | HORVU2Hr1G039170.2 | TRIUR3_20044 | Seita.2G357000.1 | Sobic.002G343000.3 | Zm00001d021828_T004 |
| orthologID_007629 | PH01000360G0200 | LOC_Os07g42760.1 | Bradi1g21102.3 | HORVU2Hr1G030550.20 | TRIUR3_22403 | Seita.2G392600.1 | Sobic.002G377200.5 | Zm00001d006884_T001 |
| orthologID_007962 | PH01003175G0160 | LOC_Os07g38460.1 | Bradi1g24000.3 | HORVU2Hr1G038060.2 | TRIUR3_16464 | Seita.2G361900.1 | Sobic.002G347400.2 | Zm00001d021869_T001 |
| orthologID_007851 | PH01001014G0410 | LOC_Os07g36250.1 | Bradi1g25310.1 | HORVU2Hr1G043310.10 | TRIUR3_06326 | Seita.2G344600.1 | Sobic.002G331100.1 | Zm00001d021715_T001 |
| orthologID_008403 | PH01000085G0310 | LOC_Os07g22950.1 | Bradi1g19480.1 | HORVU6Hr1G013200.1 | TRIUR3_09324 | Seita.2G112700.1 | Sobic.002G116200.2 | Zm00001d019481_T001 |
| orthologID_008009 | PH01002211G0290 | LOC_Os07g10540.1 | Bradi1g54327.2 | HORVU1Hr1G042060.1 | TRIUR3_22577 | Seita.2G078900.1 | Sobic.002G073200.1 | Zm00001d019130_T005 |
| orthologID_007411 | PH01000070G2000 | LOC_Os09g32430.1 | Bradi4g34260.3 | HORVU5Hr1G075260.1 | TRIUR3_22266 | Seita.2G261700.1 | Sobic.002G254500.1 | Zm00001d006061_T001 |
| orthologID_008214 | PH01002475G0080 | LOC_Os07g25430.1 | Bradi1g29000.1 | HORVU5Hr1G113900.2 | TRIUR3_03113 | Seita.2G117000.1 | Sobic.002G119500.3 | Zm00001d005446_T001 |
| orthologID_007073 | PH01001334G0240 | LOC_Os09g04050.1 | Bradi4g08647.1 | HORVU5Hr1G050940.1 | TRIUR3_34211 | Seita.2G147600.1 | Sobic.002G146000.1 | Zm00001d019669_T001 |
| orthologID_007872 | PH01000361G0850 | LOC_Os07g06440.1 | Bradi1g57200.1 | HORVU2Hr1G059360.2 | TRIUR3_07354 | Seita.2G036800.1 | Sobic.002G038200.1 | Zm00001d007881_T003 |
| orthologID_007266 | PH01000507G0160 | LOC_Os09g29420.1 | Bradi4g32720.1 | HORVU5Hr1G070240.1 | TRIUR3_32966 | Seita.2G210000.1 | Sobic.002G206500.1 | Zm00001d020541_T003 |
| orthologID_007937 | PH01003181G0010 | LOC_Os07g37250.1 | Bradi1g24860.1 | HORVU2Hr1G041260.1 | TRIUR3_21998 | Seita.2G351700.1 | Sobic.002G338100.1 | Zm00001d006588_T001 |
| orthologID_008687 | PH01000826G0010 | LOC_Os07g07060.1 | Bradi1g56640.1 | HORVU2Hr1G057880.3 | TRIUR3_20508 | Seita.2G043100.1 | Sobic.002G043600.1 | Zm00001d018870_T001 |
| orthologID_007229 | PH01002461G0070 | LOC_Os09g33800.1 | Bradi4g34980.1 | HORVU5Hr1G078000.8 | TRIUR3_11749 | Seita.2G270800.1 | Sobic.002G261900.1 | Zm00001d021123_T002 |
| orthologID_007553 | PH01001206G0420 | LOC_Os09g39790.1 | Bradi4g38890.1 | HORVU4Hr1G000940.2 | TRIUR3_22799 | Seita.2G137900.1 | Sobic.002G139100.1 | Zm00001d005313_T001 |
| orthologID_007017 | PH01163365G0010 | LOC_Os09g19570.1 | Bradi4g28580.2 | HORVU5Hr1G056820.4 | TRIUR3_09094 | Seita.2G184100.1 | Sobic.002G179100.1 | Zm00001d005641_T001 |
| orthologID_010010 | PH01004184G0110 | LOC_Os09g10940.1 | Bradi4g27130.1 | HORVU5Hr1G053370.3 | TRIUR3_06062 | Seita.2G162900.1 | Sobic.002G153500.1 | Zm00001d005504_T001 |
| orthologID_007030 | PH01000941G0160 | LOC_Os09g29970.1 | Bradi4g33010.1 | HORVU5Hr1G070930.3 | TRIUR3_19993 | Seita.2G242500.1 | Sobic.002G236700.2 | Zm00001d020838_T001 |
| orthologID_007249 | PH01001036G0570 | LOC_Os09g27340.1 | Bradi4g31610.1 | HORVU5Hr1G066780.1 | TRIUR3_17543 | Seita.2G223100.1 | Sobic.002G219500.1 | Zm00001d020656_T001 |
| orthologID_008005 | PH01000064G0590 | LOC_Os07g43040.1 | Bradi1g21380.1 | HORVU2Hr1G029720.1 | TRIUR3_13507 | Seita.2G395500.1 | Sobic.002G380400.2 | Zm00001d022188_T003 |
| orthologID_007382 | PH01000945G0580 | LOC_Os01g32730.1 | Bradi4g27210.1 | HORVU5Hr1G053480.1 | TRIUR3_23685 | Seita.5G184500.1 | Sobic.002G158800.1 | Zm00001d019956_T002 |
| orthologID_007556 | PH01000653G0330 | LOC_Os09g37920.1 | Bradi4g37260.2 | HORVU5Hr1G087670.18 | TRIUR3_06867 | Seita.2G300600.1 | Sobic.002G290000.2 | Zm00001d021430_T013 |
| orthologID_007217 | PH01001622G0070 | LOC_Os09g01590.1 | Bradi4g08200.1 | HORVU5Hr1G049730.1 | TRIUR3_18245 | Seita.2G140000.1 | Sobic.002G140700.1 | Zm00001d019721_T001 |
| orthologID_007504 | PH01000131G1400 | LOC_Os09g35670.1 | Bradi4g35690.1 | HORVU5Hr1G080550.1 | TRIUR3_24270 | Seita.2G281500.1 | Sobic.002G270100.1 | Zm00001d006175_T001 |
| orthologID_007434 | PH01000651G0690 | LOC_Os09g12530.1 | Bradi4g27470.1 | HORVU5Hr1G054010.2 | TRIUR3_15756 | Seita.2G166000.1 | Sobic.002G154800.1 | Zm00001d019993_T001 |
| orthologID_007200 | PH01000540G0640 | LOC_Os09g14680.1 | Bradi4g27920.1 | HORVU5Hr1G055380.2 | TRIUR3_28411 | Seita.2G174000.1 | Sobic.002G167500.1 | Zm00001d020063_T001 |
| orthologID_007152 | PH01002495G0210 | LOC_Os09g39180.1 | Bradi4g38260.2 | HORVU5Hr1G091810.2 | TRIUR3_25560 | Seita.2G130500.1 | Sobic.002G131700.1 | Zm00001d019648_T001 |

| | | | | | | | | |
|---|---|---|---|---|---|---|---|---|
| orthologID_007289 | PH01001055G0590 | LOC_Os09g20590.1 | Bradi4g29190.1 | HORVU5Hr1G058830.1 | TRIUR3_15046 | Seita.6G237300.1 | Sobic.002G186000.3 | Zm00001d020283_T001 |
| orthologID_010050 | PH01002478G0040 | LOC_Os07g32560.1 | Bradi1g26540.1 | HORVU2Hr1G047480.2 | TRIUR3_25279 | Seita.2G327400.1 | Sobic.002G314200.1 | Zm00001d006466_T001 |
| orthologID_008290 | PH01000204G1220 | LOC_Os07g14530.3 | Bradi1g53347.1 | HORVU3Hr1G036190.1 | TRIUR3_34131 | Seita.2G096900.1 | Sobic.002G092050.1 | Zm00001d019297_T001 |
| orthologID_007873 | PH01147644G0010 | LOC_Os07g34520.1 | Bradi1g25870.1 | HORVU2Hr1G045340.1 | TRIUR3_17318 | Seita.2G337700.1 | Sobic.002G324000.1 | Zm00001d021666_T001 |
| orthologID_008652 | PH01000826G0720 | LOC_Os12g17830.3 | Bradi1g29220.1 | HORVU7Hr1G105860.2 | TRIUR3_31680 | Seita.2G267200.1 | Sobic.002G258600.1 | Zm00001d021060_T001 |
| orthologID_007466 | PH01000970G0380 | LOC_Os09g32095.1 | Bradi4g34056.1 | HORVU5Hr1G074360.8 | TRIUR3_24569 | Seita.2G258900.1 | Sobic.002G252200.1 | Zm00001d020976_T001 |
| orthologID_007281 | PH01002891G0030 | LOC_Os09g12770.1 | Bradi4g27620.1 | HORVU5Hr1G054450.1 | TRIUR3_29870 | Seita.2G168900.1 | Sobic.002G161800.1 | Zm00001d020020_T003 |
| orthologID_007294 | PH01000060G0860 | LOC_Os09g36240.1 | Bradi4g36220.2 | HORVU5Hr1G081750.14 | TRIUR3_26853 | Seita.2G286300.1 | Sobic.002G275300.1 | Zm00001d021295_T007 |
| orthologID_007538 | PH01000486G0650 | LOC_Os09g26210.1 | Bradi4g30920.1 | HORVU5Hr1G063360.2 | TRIUR3_28597 | Seita.2G213700.1 | Sobic.002G209900.1 | Zm00001d020577_T001 |
| orthologID_007268 | PH01001036G0580 | LOC_Os09g27330.1 | Bradi4g31600.1 | HORVU5Hr1G066760.1 | TRIUR3_24345 | Seita.2G223000.1 | Sobic.002G219400.1 | Zm00001d020655_T001 |
| orthologID_007043 | PH01001001G0500 | LOC_Os09g21250.1 | Bradi4g29430.1 | HORVU5Hr1G059530.2 | TRIUR3_13588 | Seita.2G193700.1 | Sobic.002G189600.1 | Zm00001d020344_T001 |
| orthologID_007640 | PH01000306G0540 | LOC_Os07g41160.1 | Bradi1g22180.1 | HORVU2Hr1G032690.1 | TRIUR3_28445 | Seita.2G381600.1 | Sobic.002G367100.1 | Zm00001d022069_T001 |
| orthologID_007117 | PH01001075G0530 | LOC_Os09g34090.1 | Bradi4g35257.1 | HORVU5Hr1G079180.2 | TRIUR3_11324 | Seita.2G274300.1 | Sobic.002G264300.1 | Zm00001d006128_T002 |
| orthologID_009551 | PH01002571G0180 | LOC_Os10g41730.1 | Bradi3g33790.1 | HORVU1Hr1G054590.4 | TRIUR3_11706 | Seita.9G316300.1 | Sobic.002G381900.2 | Zm00001d022194_T001 |
| orthologID_007140 | PH01000319G0250 | LOC_Os09g29100.1 | Bradi4g32556.1 | HORVU7Hr1G033350.3 | TRIUR3_31847 | Seita.2G236100.1 | Sobic.002G230900.1 | Zm00001d005928_T002 |
| orthologID_007154 | PH01001676G0170 | LOC_Os09g23070.1 | Bradi4g29630.1 | HORVU5Hr1G060110.16 | TRIUR3_31557 | Seita.2G196300.1 | Sobic.002G193500.1 | Zm00001d005711_T005 |
| orthologID_007950 | PH01000442G0280 | LOC_Os07g46410.1 | Bradi1g19140.1 | HORVU2Hr1G023170.1 | TRIUR3_27644 | Seita.2G417000.1 | Sobic.002G401800.1 | Zm00001d022381_T016 |
| orthologID_007564 | PH01000451G0320 | LOC_Os07g30110.1 | Bradi1g27595.1 | HORVU6Hr1G040270.9 | TRIUR3_30197 | Seita.2G315800.1 | Sobic.002G303800.1 | Zm00001d021505_T001 |
| orthologID_007078 | PH01001228G0180 | LOC_Os09g28160.1 | Bradi4g32020.2 | HORVU5Hr1G068060.1 | TRIUR3_16873 | Seita.2G228300.1 | Sobic.002G224100.1 | Zm00001d020702_T001 |
| orthologID_007507 | PH01001882G0330 | LOC_Os09g11250.1 | Bradi4g27270.1 | HORVU5Hr1G053680.3 | TRIUR3_22645 | Seita.2G161600.1 | Sobic.002G157100.1 | Zm00001d005513_T001 |
| orthologID_008098 | PH01000086G1010 | LOC_Os07g38970.1 | Bradi1g23600.1 | HORVU2Hr1G036950.1 | TRIUR3_30087 | Seita.2G366700.1 | Sobic.002G352200.1 | Zm00001d021908_T005 |
| orthologID_006784 | PH01001032G0490 | LOC_Os01g29430.1 | Bradi2g13310.1 | HORVU5Hr1G095110.1 | TRIUR3_24819 | Seita.9G530700.1 | Sobic.002G251800.2 | Zm00001d020972_T010 |
| orthologID_008712 | PH01000736G0070 | LOC_Os07g27480.1 | Bradi1g28540.1 | HORVU7Hr1G021530.12 | TRIUR3_20141 | Seita.2G125900.1 | Sobic.002G127900.2 | Zm00001d019591_T006 |
| orthologID_008037 | PH01000365G0310 | LOC_Os03g62600.1 | Bradi1g02100.1 | HORVU5Hr1G120210.4 | TRIUR3_02197 | Seita.9G387600.1 | Sobic.002G103500.1 | Zm00001d005495_T001 |
| orthologID_007107 | PH01002940G0220 | LOC_Os12g16690.1 | Bradi4g39250.1 | HORVU6Hr1G075300.1 | TRIUR3_28559 | Seita.3G298300.1 | Sobic.002G426800.1 | Zm00001d022562_T002 |
| orthologID_010040 | PH01000436G0570 | LOC_Os07g44690.1 | Bradi1g19900.1 | HORVU2Hr1G026020.1 | TRIUR3_10874 | Seita.2G407800.1 | Sobic.002G392500.1 | Zm00001d022295_T001 |
| orthologID_008195 | PH01001763G0210 | LOC_Os07g01100.1 | Bradi1g59820.1 | HORVU2Hr1G065090.1 | TRIUR3_33196 | Seita.J005100.1 | Sobic.002G001100.3 | Zm00001d018599_T001 |
| orthologID_007040 | PH01000050G1100 | LOC_Os09g30466.1 | Bradi4g33440.1 | HORVU5Hr1G072470.2 | TRIUR3_05958 | Seita.2G249600.1 | Sobic.002G243650.1 | Zm00001d020900_T001 |
| orthologID_009449 | PH01006829G0040 | LOC_Os02g24080.4 | Bradi3g11220.5 | HORVU7Hr1G008760.1 | TRIUR3_01355 | Seita.1G147700.1 | Sobic.002G118600.1 | Zm00001d019505_T001 |
| orthologID_008684 | PH01003449G0120 | LOC_Os07g39930.1 | Bradi1g22970.2 | HORVU2Hr1G035200.1 | TRIUR3_17669 | Seita.2G373400.1 | Sobic.002G359100.1 | Zm00001d006732_T001 |
| orthologID_008032 | PH01004483G0030 | LOC_Os07g13280.1 | Bradi1g53620.2 | HORVU2Hr1G056980.13 | TRIUR3_30079 | Seita.2G091500.1 | Sobic.002G083900.1 | Zm00001d019234_T003 |
| orthologID_007306 | PH01001123G0170 | LOC_Os09g20430.1 | Bradi4g29060.8 | HORVU5Hr1G058400.9 | TRIUR3_22324 | Seita.2G188500.1 | Sobic.002G184900.1 | Zm00001d005662_T001 |
| orthologID_007946 | PH01000068G0620 | LOC_Os07g44360.1 | Bradi1g20067.2 | HORVU2Hr1G026860.2 | TRIUR3_25091 | Seita.2G406200.1 | Sobic.002G390700.2 | Zm00001d006938_T001 |
| orthologID_008373 | PH01001059G0440 | LOC_Os07g44170.1 | Bradi1g20230.1 | HORVU2Hr1G027440.18 | TRIUR3_15064 | Seita.2G404600.1 | Sobic.002G389200.2 | Zm00001d022263_T001 |
| orthologID_007277 | PH01001838G0110 | LOC_Os08g28190.1 | Bradi4g18730.1 | HORVU4Hr1G007240.3 | TRIUR3_03422 | Seita.4G100400.1 | Sobic.002G426300.1 | Zm00001d022558_T001 |
| orthologID_008520 | PH01000591G0660 | LOC_Os07g45350.1 | Bradi1g19510.1 | HORVU2Hr1G024550.4 | TRIUR3_07655 | Seita.2G413800.1 | Sobic.002G399100.1 | Zm00001d007060_T003 |
| orthologID_008486 | PH01000855G0400 | LOC_Os07g06570.1 | Bradi1g57100.1 | HORVU2Hr1G059520.2 | TRIUR3_26931 | Seita.2G037400.1 | Sobic.002G038866.1 | Zm00001d018825_T001 |
| orthologID_007185 | PH01000009G1200 | LOC_Os09g36040.1 | Bradi4g35810.1 | HORVU1Hr1G045190.2 | TRIUR3_11770 | Seita.2G283000.1 | Sobic.002G271700.2 | Zm00001d021245_T001 |
| orthologID_008265 | PH01001572G0270 | LOC_Os07g10300.3 | Bradi1g54480.2 | HORVU1Hr1G042020.1 | TRIUR3_18664 | Seita.2G076600.1 | Sobic.002G071100.1 | Zm00001d007754_T002 |
| orthologID_008650 | PH01000736G0450 | LOC_Os07g27810.1 | Bradi1g28490.1 | HORVU3Hr1G092420.1 | TRIUR3_01235 | Seita.2G127400.1 | Sobic.002G128800.1 | Zm00001d005394_T001 |
| orthologID_007421 | PH01002328G0330 | LOC_Os08g27860.1 | Bradi4g29580.1 | HORVU5Hr1G060000.1 | TRIUR3_30694 | Seita.2G195800.1 | Sobic.002G193000.1 | Zm00001d020364_T001 |
| orthologID_008530 | PH01001195G0510 | LOC_Os07g28900.1 | Bradi1g28220.1 | HORVU7Hr1G091700.1 | TRIUR3_22331 | Seita.2G311000.1 | Sobic.002G299400.1 | Zm00001d021343_T001 |
| orthologID_007714 | PH01000442G0510 | LOC_Os07g46500.1 | Bradi1g19030.1 | HORVU2Hr1G022900.2 | TRIUR3_16022 | Seita.2G418600.1 | Sobic.002G403200.1 | Zm00001d022393_T001 |
| orthologID_007893 | PH01002574G0010 | LOC_Os07g40120.1 | Bradi1g22890.2 | HORVU2Hr1G035060.2 | TRIUR3_29044 | Seita.2G374500.1 | Sobic.002G360200.1 | Zm00001d022003_T001 |
| orthologID_007551 | PH01004197G0100 | LOC_Os09g15810.1 | Bradi4g28150.1 | HORVU5Hr1G056400.1 | TRIUR3_17732 | Seita.2G178100.1 | Sobic.002G172600.1 | Zm00001d005614_T001 |
| orthologID_007121 | PH01157768G0010 | LOC_Os09g19900.1 | Bradi4g28830.2 | HORVU5Hr1G057500.1 | TRIUR3_07245 | Seita.2G186300.1 | Sobic.002G181400.1 | Zm00001d020234_T001 |
| orthologID_007112 | PH01000149G0040 | LOC_Os09g28600.1 | Bradi4g32270.1 | HORVU0Hr1G013120.2 | TRIUR3_34376 | Seita.2G232100.1 | Sobic.002G227300.1 | Zm00001d005901_T001 |
| orthologID_007193 | PH01000511G0190 | LOC_Os09g37006.1 | Bradi4g36710.1 | HORVU5Hr1G084510.3 | TRIUR3_02686 | Seita.2G292300.1 | Sobic.002G281600.1 | Zm00001d006250_T001 |
| orthologID_007066 | PH01000405G0510 | LOC_Os09g25060.1 | Bradi4g30360.1 | HORVU5Hr1G065420.1 | TRIUR3_16803 | Seita.2G206000.1 | Sobic.002G202700.1 | Zm00001d020492_T001 |
| orthologID_008701 | PH01001113G0300 | LOC_Os07g38220.1 | Bradi1g24170.2 | HORVU2Hr1G038800.2 | TRIUR3_14167 | Seita.2G358300.1 | Sobic.002G345000.3 | Zm00001d021840_T001 |

| | | | | | | | | |
|---|---|---|---|---|---|---|---|---|
| orthologID_007020 | PH01213953G0010 | LOC_Os09g19750.1 | Bradi4g28680.1 | HORVU5Hr1G057100.1 | TRIUR3_19762 | Seita.2G185500.1 | Sobic.002G180600.1 | Zm00001d020224_T001 |
| orthologID_008572 | PH01168297G0010 | LOC_Os07g40700.2 | Bradi1g22607.1 | HORVU2Hr1G034350.4 | TRIUR3_23295 | Seita.2G377900.1 | Sobic.002G363500.2 | Zm00001d022045_T001 |
| orthologID_009341 | PH01004568G0130 | LOC_Os08g20400.1 | Bradi3g20010.1 | HORVU7Hr1G022210.2 | TRIUR3_29346 | Seita.8G006200.1 | Sobic.002G422400.1 | Zm00001d022531_T001 |
| orthologID_008129 | PH01000064G0640 | LOC_Os07g43020.1 | Bradi1g21366.1 | HORVU2Hr1G029760.2 | TRIUR3_23793 | Seita.2G395200.1 | Sobic.002G380000.1 | Zm00001d006904_T001 |
| orthologID_007684 | PH01003453G0220 | LOC_Os07g01720.1 | Bradi1g59350.1 | HORVU2Hr1G064070.2 | TRIUR3_33701 | Seita.2G001200.1 | Sobic.002G006300.1 | Zm00001d018636_T001 |
| orthologID_007860 | PH01001479G0370 | LOC_Os07g47670.2 | Bradi1g18030.1 | HORVU2Hr1G019250.1 | TRIUR3_04143 | Seita.2G429300.1 | Sobic.002G414600.1 | Zm00001d007153_T001 |
| orthologID_007182 | PH01000379G0100 | LOC_Os09g25950.1 | Bradi4g30797.1 | HORVU0Hr1G013040.2 | TRIUR3_10358 | Seita.2G212000.1 | Sobic.002G208400.2 | Zm00001d020561_T002 |
| orthologID_007933 | PH01001195G0300 | LOC_Os07g28940.1 | Bradi1g28167.2 | HORVU2Hr1G059170.16 | TRIUR3_22943 | Seita.2G311400.1 | Sobic.002G299700.1 | Zm00001d006357_T001 |
| orthologID_007972 | PH01002880G0120 | LOC_Os07g39730.2 | Bradi1g23140.2 | HORVU2Hr1G035780.2 | TRIUR3_23479 | Seita.2G370800.1 | Sobic.002G356600.1 | Zm00001d021960_T001 |
| orthologID_007397 | PH01002230G0270 | LOC_Os09g03610.1 | Bradi4g08727.2 | HORVU5Hr1G050820.6 | TRIUR3_35136 | Seita.2G146900.1 | Sobic.002G145800.1 | Zm00001d005281_T004 |
| orthologID_008466 | PH01000306G0610 | LOC_Os07g41370.1 | Bradi1g21980.2 | HORVU0Hr1G003020.3 | TRIUR3_20924 | Seita.2G383000.1 | Sobic.002G368700.1 | Zm00001d022088_T004 |
| orthologID_008427 | PH01001472G0030 | LOC_Os07g49480.2 | Bradi1g16470.1 | HORVU2Hr1G013370.2 | TRIUR3_21509 | Seita.2G444600.1 | Sobic.002G421100.1 | Zm00001d007241_T001 |
| orthologID_008356 | PH01001843G0240 | LOC_Os07g12330.1 | Bradi1g53780.2 | HORVU4Hr1G068690.4 | TRIUR3_03805 | Seita.2G086200.1 | Sobic.002G079900.1 | Zm00001d019206_T002 |
| orthologID_006562 | PH01000751G0100 | LOC_Os07g12530.1 | Bradi2g36950.1 | HORVU7Hr1G054230.1 | TRIUR3_32439 | Seita.2G088100.1 | Sobic.002G081400.1 | Zm00001d019217_T002 |
| orthologID_008410 | PH01000364G1150 | LOC_Os07g47790.1 | Bradi1g17961.1 | HORVU2Hr1G018830.1 | TRIUR3_02386 | Seita.2G430400.1 | Sobic.002G415600.1 | Zm00001d022461_T001 |
| orthologID_008243 | PH01000056G0250 | LOC_Os07g46170.1 | Bradi1g19380.1 | HORVU0Hr1G002680.2 | TRIUR3_10061 | Seita.2G414700.1 | Sobic.002G399900.1 | Zm00001d007067_T003 |
| orthologID_008672 | PH01001113G0360 | LOC_Os07g38240.1 | Bradi1g24140.1 | HORVU2Hr1G038760.3 | TRIUR3_28482 | Seita.2G358600.1 | Sobic.002G345300.2 | Zm00001d021842_T001 |
| orthologID_006343 | PH01003397G0070 | LOC_Os05g23610.1 | Bradi2g31120.2 | HORVU4Hr1G011360.1 | TRIUR3_26720 | Seita.6G097300.1 | Sobic.002G087900.1 | Zm00001d019263_T002 |
| orthologID_007165 | PH01004196G0130 | LOC_Os09g35880.1 | Bradi4g35950.1 | HORVU5Hr1G081190.1 | TRIUR3_22144 | Seita.2G284200.1 | Sobic.002G273300.1 | Zm00001d021278_T001 |
| orthologID_008046 | PH01000361G0700 | LOC_Os07g06080.1 | Bradi1g57360.1 | HORVU2Hr1G059320.1 | TRIUR3_07352 | Seita.2G035900.1 | Sobic.002G037800.1 | Zm00001d018820_T004 |
| orthologID_007325 | PH01000950G0020 | LOC_Os09g27260.1 | Bradi4g31582.1 | HORVU5Hr1G066720.2 | TRIUR3_29138 | Seita.2G222800.1 | Sobic.002G219200.1 | Zm00001d020652_T001 |
| orthologID_007542 | PH01000250G0780 | LOC_Os09g09470.1 | Bradi4g26970.1 | HORVU5Hr1G052660.2 | TRIUR3_30521 | Seita.1G161000.1 | Sobic.002G150800.1 | Zm00001d005269_T002 |
| orthologID_008536 | PH01000064G0570 | LOC_Os07g43060.1 | Bradi1g21400.1 | HORVU2Hr1G029620.1 | TRIUR3_13505 | Seita.2G395700.1 | Sobic.002G380600.1 | Zm00001d006907_T002 |
| orthologID_007218 | PH01003334G0070 | LOC_Os09g32526.1 | Bradi4g34370.1 | HORVU5Hr1G075490.3 | TRIUR3_19464 | Seita.2G263100.1 | Sobic.002G255400.1 | Zm00001d021021_T001 |
| orthologID_007243 | PH01003825G0020 | LOC_Os09g20684.1 | Bradi4g29243.1 | HORVU5Hr1G058910.2 | TRIUR3_31354 | Seita.2G190600.1 | Sobic.002G186600.1 | Zm00001d020306_T001 |
| orthologID_007035 | PH01000314G0440 | LOC_Os09g35780.1 | Bradi4g35770.2 | HORVU5Hr1G080820.1 | TRIUR3_27491 | Seita.2G282300.1 | Sobic.002G271000.1 | Zm00001d021266_T001 |
| orthologID_007093 | PH01001034G0110 | LOC_Os09g38768.2 | Bradi4g37980.1 | HORVU5Hr1G094000.6 | TRIUR3_02690 | Seita.2G307900.1 | Sobic.002G296600.1 | Zm00001d021371_T001 |
| orthologID_009883 | PH01001879G0150 | LOC_Os08g23360.1 | Bradi3g07210.1 | HORVU5Hr1G042050.1 | TRIUR3_27314 | Seita.8G114100.1 | Sobic.002G371700.1 | Zm00001d022115_T002 |
| orthologID_007938 | PH01003652G0040 | LOC_Os07g09050.1 | Bradi1g54950.1 | HORVU5Hr1G009150.5 | TRIUR3_10466 | Seita.2G066800.1 | Sobic.002G062100.1 | Zm00001d019053_T001 |
| orthologID_007387 | PH01000096G1860 | LOC_Os07g19030.1 | Bradi4g27410.1 | HORVU3Hr1G071890.1 | TRIUR3_29059 | Seita.2G107200.1 | Sobic.002G109600.1 | Zm00001d019404_T001 |
| orthologID_007359 | PH01000444G0130 | LOC_Os09g24520.2 | Bradi4g30017.1 | HORVU5Hr1G066470.6 | TRIUR3_00974 | Seita.2G202400.1 | Sobic.002G198801.1 | Zm00001d020435_T001 |
| orthologID_007181 | PH01000646G0290 | LOC_Os09g19640.1 | Bradi4g28590.1 | HORVU5Hr1G056860.2 | TRIUR3_09092 | Seita.2G184400.1 | Sobic.002G179500.1 | Zm00001d005645_T001 |
| orthologID_007049 | PH01005316G0070 | LOC_Os09g16280.1 | Bradi4g28260.1 | HORVU5Hr1G056270.1 | TRIUR3_17713 | Seita.2G179500.1 | Sobic.002G173800.1 | Zm00001d020133_T001 |
| orthologID_008119 | PH01000306G0830 | LOC_Os07g41280.3 | Bradi1g22050.1 | HORVU2Hr1G032970.4 | TRIUR3_01707 | Seita.2G382400.1 | Sobic.002G368000.2 | Zm00001d022077_T003 |
| orthologID_007337 | PH01002271G0140 | LOC_Os09g26580.1 | Bradi4g31070.1 | HORVU5Hr1G062740.3 | TRIUR3_12448 | Seita.2G216200.1 | Sobic.002G213400.1 | Zm00001d020603_T004 |
| orthologID_008029 | PH01000442G0350 | LOC_Os07g46440.1 | Bradi1g19110.2 | HORVU2Hr1G023120.1 | TRIUR3_17096 | Seita.2G417700.1 | Sobic.002G402500.1 | Zm00001d022386_T001 |
| orthologID_007416 | PH01002633G0060 | LOC_Os11g19220.1 | Bradi4g20250.3 | HORVU4Hr1G012340.1 | TRIUR3_25433 | Seita.8G107400.1 | Sobic.002G377900.1 | Zm00001d022164_T002 |
| orthologID_008331 | PH01000073G0750 | LOC_Os07g39920.1 | Bradi1g22980.1 | HORVU2Hr1G035230.1 | TRIUR3_12045 | Seita.2G373300.1 | Sobic.002G359000.1 | Zm00001d021985_T001 |
| orthologID_007221 | PH01001149G0270 | LOC_Os09g02284.1 | Bradi4g08350.1 | HORVU5Hr1G050250.10 | TRIUR3_10978 | Seita.2G142700.1 | Sobic.002G145000.1 | Zm00001d005295_T001 |
| orthologID_008459 | PH01000086G0640 | LOC_Os07g39300.1 | Bradi1g23480.1 | HORVU2Hr1G036750.4 | TRIUR3_12803 | Seita.2G368400.1 | Sobic.002G353600.1 | Zm00001d006681_T004 |
| orthologID_007487 | PH01000063G1010 | LOC_Os09g33580.2 | Bradi4g34870.3 | HORVU5Hr1G077390.1 | TRIUR3_15679 | Seita.2G269400.1 | Sobic.002G260600.1 | Zm00001d021086_T001 |
| orthologID_007348 | PH01001386G0540 | LOC_Os09g27140.1 | Bradi4g31500.1 | HORVU5Hr1G061510.1 | TRIUR3_13890 | Seita.2G221600.1 | Sobic.002G218000.1 | Zm00001d020644_T001 |
| orthologID_008342 | PH01000541G0110 | LOC_Os07g11280.2 | Bradi1g54007.2 | HORVU2Hr1G056820.1 | TRIUR3_32701 | Seita.2G082900.1 | Sobic.002G347600.1 | Zm00001d021871_T002 |
| orthologID_007141 | PH01000332G1000 | LOC_Os09g37012.1 | Bradi4g36720.1 | HORVU5Hr1G084490.1 | TRIUR3_02687 | Seita.2G292400.1 | Sobic.002G281700.1 | Zm00001d022600_T001 |
| orthologID_008601 | PH01000364G1030 | LOC_Os07g47720.1 | Bradi1g17997.3 | HORVU2Hr1G019150.1 | TRIUR3_16010 | Seita.2G429600.1 | Sobic.002G414800.1 | Zm00001d007155_T001 |
| orthologID_008036 | PH01003658G0140 | LOC_Os07g49410.1 | Bradi1g16500.1 | HORVU2Hr1G013480.1 | TRIUR3_16102 | Seita.2G444100.1 | Sobic.002G431300.1 | Zm00001d022588_T001 |
| orthologID_007852 | PH01003453G0070 | LOC_Os07g01780.1 | Bradi1g59232.1 | HORVU2Hr1G063790.1 | TRIUR3_33612 | Seita.2G002200.1 | Sobic.002G010200.4 | Zm00001d018668_T002 |
| orthologID_007517 | PH01000507G0180 | LOC_Os09g29404.1 | Bradi4g32707.1 | HORVU5Hr1G070160.2 | TRIUR3_24745 | Seita.2G237100.1 | Sobic.002G233600.1 | Zm00001d020799_T003 |
| orthologID_007400 | PH01002111G0240 | LOC_Os09g20930.1 | Bradi4g29320.2 | HORVU5Hr1G059140.9 | TRIUR3_31610 | Seita.2G192100.1 | Sobic.002G187700.2 | Zm00001d005684_T001 |
| orthologID_008100 | PH01000553G0030 | LOC_Os07g30820.1 | Bradi1g27210.1 | HORVU2Hr1G050290.5 | TRIUR3_31910 | Seita.2G319000.1 | Sobic.002G306400.1 | Zm00001d006409_T002 |

| | | | | | | | | |
|---|---|---|---|---|---|---|---|---|
| orthologID_007216 | PH01002098G0290 | LOC_Os09g23360.1 | Bradi4g41440.2 | HORVU5Hr1G060450.5 | TRIUR3_16029 | Seita.2G197900.1 | Sobic.002G194800.2 | Zm00001d020399_T007 |
| orthologID_008365 | PH01002322G0260 | LOC_Os07g38000.1 | Bradi1g24320.1 | HORVU2Hr1G039510.2 | TRIUR3_31827 | Seita.2G356500.1 | Sobic.002G342600.1 | Zm00001d021825_T001 |
| orthologID_007450 | PH01028363G0010 | LOC_Os09g37280.1 | Bradi4g36920.8 | HORVU5Hr1G085520.1 | TRIUR3_22683 | Seita.2G294600.1 | Sobic.002G284200.1 | Zm00001d006270_T001 |
| orthologID_008247 | PH01001858G0220 | LOC_Os07g08070.1 | Bradi1g55650.1 | HORVU3Hr1G105460.2 | TRIUR3_34972 | Seita.2G051800.1 | Sobic.002G051700.1 | Zm00001d018938_T001 |
| orthologID_007340 | PH01001832G0330 | LOC_Os09g37540.1 | Bradi4g37070.1 | HORVU5Hr1G086520.6 | TRIUR3_26011 | Seita.2G297500.1 | Sobic.002G286400.1 | Zm00001d021450_T001 |
| orthologID_008127 | PH01000068G0540 | LOC_Os07g44410.1 | Bradi1g20050.1 | HORVU2Hr1G026840.1 | TRIUR3_14409 | Seita.2G406400.1 | Sobic.002G390900.1 | Zm00001d022277_T001 |
| orthologID_008585 | PH01000056G1590 | LOC_Os07g46750.1 | Bradi1g18710.1 | HORVU2Hr1G022090.4 | TRIUR3_03789 | Seita.2G421400.1 | Sobic.002G406500.1 | Zm00001d022513_T001 |
| orthologID_008407 | PH01001014G0230 | LOC_Os07g36390.1 | Bradi1g28870.2 | HORVU2Hr1G043170.1 | TRIUR3_06322 | Seita.2G345300.1 | Sobic.002G331800.1 | Zm00001d021716_T001 |
| orthologID_007602 | PH01002546G0390 | LOC_Os07g29320.1 | Bradi1g04477.2 | HORVU4Hr1G078580.2 | TRIUR3_18474 | Seita.2G311900.1 | Sobic.002G300300.2 | Zm00001d015662_T001 |
| orthologID_007354 | PH01000894G0310 | LOC_Os09g35800.1 | Bradi4g35790.1 | HORVU5Hr1G080860.9 | TRIUR3_25331 | Seita.2G282600.1 | Sobic.002G271200.1 | Zm00001d021249_T003 |
| orthologID_007781 | PH01002295G0230 | LOC_Os07g34589.2 | Bradi1g25837.1 | HORVU2Hr1G045250.4 | TRIUR3_30345 | Seita.2G338000.1 | Sobic.002G324200.1 | Zm00001d021668_T002 |
| orthologID_008125 | PH01006391G0040 | LOC_Os05g19390.1 | Bradi1g15820.3 | HORVU6Hr1G014060.1 | TRIUR3_32871 | Seita.3G295200.1 | Sobic.002G380100.1 | Zm00001d022184_T008 |
| orthologID_007386 | PH01002135G0120 | LOC_Os09g19650.1 | Bradi4g28597.1 | HORVU5Hr1G056870.5 | TRIUR3_13978 | Seita.2G184500.1 | Sobic.002G179700.1 | Zm00001d020206_T001 |
| orthologID_008657 | PH01003073G0020 | LOC_Os07g31650.1 | Bradi1g26857.4 | HORVU2Hr1G048520.8 | TRIUR3_31100 | Seita.2G320000.1 | Sobic.002G307800.1 | Zm00001d021532_T001 |
| orthologID_007350 | PH01001001G0330 | LOC_Os09g21380.1 | Bradi4g29450.1 | HORVU5Hr1G059690.7 | TRIUR3_10468 | Seita.2G194000.1 | Sobic.002G189900.1 | Zm00001d020348_T001 |
| orthologID_007634 | PH01000553G0290 | LOC_Os07g31370.1 | Bradi1g27010.1 | HORVU2Hr1G050270.1 | TRIUR3_28923 | Seita.2G322100.1 | Sobic.002G309500.1 | Zm00001d006433_T001 |
| orthologID_008705 | PH01001245G0310 | LOC_Os07g39430.1 | Bradi1g23400.2 | HORVU2Hr1G036490.11 | TRIUR3_21656 | Seita.2G369200.1 | Sobic.002G354600.1 | Zm00001d006700_T003 |
| orthologID_007932 | PH01000656G0620 | LOC_Os07g36090.3 | Bradi1g25420.3 | HORVU2Hr1G043970.1 | TRIUR3_13700 | Seita.2G343100.1 | Sobic.002G329700.1 | Zm00001d021704_T001 |
| orthologID_007144 | PH01000978G0590 | LOC_Os09g39490.1 | Bradi4g38527.1 | HORVU4Hr1G001140.1 | TRIUR3_05688 | Seita.2G133600.1 | Sobic.002G135100.1 | Zm00001d019865_T001 |
| orthologID_007273 | PH01001229G0490 | LOC_Os09g39980.1 | Bradi4g31350.1 | HORVU5Hr1G062010.4 | TRIUR3_22838 | Seita.2G143500.1 | Sobic.002G143000.1 | Zm00001d019671_T002 |
| orthologID_007412 | PH01000191G1450 | LOC_Os09g34300.1 | Bradi4g35440.1 | HORVU5Hr1G079750.36 | TRIUR3_03571 | Seita.2G276900.1 | Sobic.002G266300.1 | Zm00001d021170_T004 |
| orthologID_007744 | PH01000752G0660 | LOC_Os07g27350.1 | Bradi1g28600.1 | HORVU7Hr1G078870.10 | TRIUR3_17836 | Seita.2G125300.1 | Sobic.002G127550.1 | Zm00001d019587_T007 |
| orthologID_007513 | PH01000486G0360 | LOC_Os09g26190.1 | Bradi4g30897.1 | HORVU5Hr1G063380.19 | TRIUR3_32398 | Seita.2G213400.1 | Sobic.002G209600.3 | Zm00001d005785_T006 |
| orthologID_007457 | PH01000063G1970 | LOC_Os09g34000.1 | Bradi4g35170.1 | HORVU5Hr1G078830.3 | TRIUR3_04065 | Seita.2G272700.1 | Sobic.002G264000.1 | Zm00001d006126_T001 |
| orthologID_007069 | PH01000050G1230 | LOC_Os09g30411.1 | Bradi4g33395.1 | HORVU5Hr1G072140.4 | TRIUR3_07862 | Seita.2G248700.1 | Sobic.002G242800.1 | Zm00001d020882_T001 |
| orthologID_008510 | PH01000436G0090 | LOC_Os07g45060.1 | Bradi1g19660.1 | HORVU2Hr1G025150.1 | TRIUR3_23618 | Seita.2G412000.1 | Sobic.002G396400.1 | Zm00001d007045_T001 |
| orthologID_007293 | PH01002454G0130 | LOC_Os09g09370.2 | Bradi4g08917.6 | HORVU0Hr1G012710.2 | TRIUR3_17683 | Seita.2G149300.1 | Sobic.002G145500.1 | Zm00001d019944_T004 |
| orthologID_008645 | PH01000096G0870 | LOC_Os07g19530.1 | Bradi1g52690.1 | HORVU2Hr1G051970.9 | TRIUR3_15904 | Seita.2G108500.1 | Sobic.002G111000.1 | Zm00001d019432_T002 |
| orthologID_008059 | PH01000490G0700 | LOC_Os06g12876.1 | Bradi1g22590.1 | HORVU2Hr1G034130.5 | TRIUR3_24760 | Seita.3G402200.1 | Sobic.002G363800.3 | Zm00001d006762_T004 |
| orthologID_007319 | PH01000379G0560 | LOC_Os09g25770.1 | Bradi4g30720.1 | HORVU5Hr1G064010.18 | TRIUR3_10721 | Seita.2G211200.1 | Sobic.002G207400.1 | Zm00001d020555_T001 |
| orthologID_007211 | PH01003389G0190 | LOC_Os09g31270.1 | Bradi4g33660.1 | HORVU3Hr1G088950.1 | TRIUR3_09892 | Seita.2G252700.1 | Sobic.002G246400.1 | Zm00001d020931_T001 |
| orthologID_007968 | PH01000064G1820 | LOC_Os07g43670.1 | Bradi1g20520.1 | HORVU2Hr1G028430.1 | TRIUR3_20260 | Seita.2G400700.1 | Sobic.002G385500.1 | Zm00001d022228_T001 |
| orthologID_008133 | PH01002112G0230 | LOC_Os07g11060.1 | Bradi1g54050.1 | HORVU2Hr1G052170.1 | TRIUR3_13349 | Seita.2G082200.1 | Sobic.002G076800.1 | Zm00001d019172_T001 |
| orthologID_007638 | PH01000068G1660 | LOC_Os07g43470.1 | Bradi1g20620.1 | HORVU2Hr1G028600.1 | TRIUR3_07542 | Seita.2G399400.1 | Sobic.002G383800.1 | Zm00001d022209_T001 |
| orthologID_008603 | PH01001623G0370 | LOC_Os07g26610.1 | Bradi1g28790.1 | HORVU5Hr1G027250.12 | TRIUR3_12987 | Seita.2G123400.1 | Sobic.002G125800.1 | Zm00001d019568_T001 |
| orthologID_007476 | PH01002891G0050 | LOC_Os09g12780.1 | Bradi4g27627.1 | HORVU5Hr1G054480.1 | TRIUR3_26531 | Seita.3G047200.1 | Sobic.002G162300.2 | Zm00001d020021_T001 |
| orthologID_007725 | PH01000640G0190 | LOC_Os07g37010.1 | Bradi1g24990.2 | HORVU0Hr1G003280.14 | TRIUR3_27243 | Seita.2G349600.1 | Sobic.002G335900.1 | Zm00001d021745_T001 |
| orthologID_007906 | PH01001424G0190 | LOC_Os07g13540.1 | Bradi1g53600.1 | HORVU2Hr1G023100.1 | TRIUR3_17097 | Seita.2G091900.1 | Sobic.002G084400.1 | Zm00001d019240_T001 |
| orthologID_007940 | PH01000977G0200 | LOC_Os07g01550.1 | Bradi1g59430.1 | HORVU2Hr1G064170.2 | TRIUR3_18304 | Seita.2G000300.1 | Sobic.002G004800.1 | Zm00001d018626_T001 |
| orthologID_008334 | PH01001835G0300 | LOC_Os07g07480.1 | Bradi1g56137.1 | HORVU7Hr1G027910.1 | TRIUR3_09757 | Seita.2G046100.1 | Sobic.002G046900.1 | Zm00001d018895_T001 |
| orthologID_007026 | PH01000431G0610 | LOC_Os09g20910.1 | Bradi4g29310.2 | HORVU5Hr1G059040.4 | TRIUR3_04909 | Seita.2G192000.1 | Sobic.002G187600.1 | Zm00001d020312_T017 |
| orthologID_007057 | PH01000379G0980 | LOC_Os09g25580.1 | Bradi4g30600.1 | HORVU5Hr1G064090.1 | TRIUR3_11907 | Seita.2G209700.1 | Sobic.002G206200.1 | Zm00001d020537_T001 |
| orthologID_007923 | PH01001639G0210 | LOC_Os07g47090.2 | Bradi1g18347.7 | HORVU2Hr1G021090.1 | TRIUR3_30807 | Seita.2G422600.1 | Sobic.002G408000.3 | Zm00001d007105_T002 |
| orthologID_007736 | PH01005295G0040 | LOC_Os07g16130.1 | Bradi1g53050.1 | HORVU5Hr1G039960.2 | TRIUR3_30433 | Seita.9G386700.1 | Sobic.002G102200.1 | Zm00001d019343_T001 |
| orthologID_007345 | PH01000050G1020 | LOC_Os09g30486.1 | Bradi4g33490.3 | HORVU5Hr1G072620.4 | TRIUR3_32885 | Seita.2G250200.1 | Sobic.002G244300.1 | Zm00001d020903_T001 |
| orthologID_008343 | PH01000073G1210 | LOC_Os07g40240.1 | Bradi1g22863.1 | HORVU2Hr1G034860.1 | TRIUR3_20452 | Seita.2G374900.1 | Sobic.002G360900.1 | Zm00001d006750_T001 |
| orthologID_010207 | PH01000534G0230 | LOC_Os09g25070.1 | Bradi4g30372.1 | HORVU5Hr1G065380.2 | TRIUR3_16804 | Seita.2G206100.1 | Sobic.002G202800.1 | Zm00001d020495_T001 |
| orthologID_007316 | PH01001868G0360 | LOC_Os09g25314.1 | Bradi4g30460.1 | HORVU5Hr1G065230.2 | TRIUR3_04551 | Seita.2G207400.1 | Sobic.002G204100.3 | Zm00001d020510_T003 |
| orthologID_008374 | PH01001827G0280 | LOC_Os07g47210.1 | Bradi1g18470.1 | HORVU2Hr1G020430.2 | TRIUR3_20913 | Seita.2G424400.1 | Sobic.002G409000.1 | Zm00001d022496_T001 |
| orthologID_007977 | PH01001650G0340 | LOC_Os07g41750.1 | Bradi1g21730.2 | HORVU2Hr1G031940.9 | TRIUR3_33092 | Seita.2G385700.1 | Sobic.002G371100.1 | Zm00001d022111_T002 |

| | | | | | | | | |
|---|---|---|---|---|---|---|---|---|
| orthologID_008432 | PH01000083G0680 | LOC_Os09g24240.1 | Bradi1g51870.1 | HORVU2Hr1G075670.1 | TRIUR3_02941 | Seita.2G173900.1 | Sobic.002G167400.1 | Zm00001d020062_T001 |
| orthologID_007884 | PH01003394G0190 | LOC_Os07g37400.1 | Bradi1g24790.1 | HORVU2Hr1G040830.5 | TRIUR3_27000 | Seita.2G352600.1 | Sobic.002G339000.1 | Zm00001d006591_T001 |
| orthologID_008137 | PH01000961G0160 | LOC_Os09g01680.1 | Bradi1g27880.1 | HORVU7Hr1G105200.27 | TRIUR3_16296 | Seita.5G033300.1 | Sobic.002G395700.2 | Zm00001d022332_T001 |
| orthologID_009568 | PH01001068G0250 | LOC_Os08g07060.1 | Bradi3g16010.3 | HORVU6Hr1G009510.1 | TRIUR3_09163 | Seita.6G049100.1 | Sobic.002G198000.1 | Zm00001d049490_T001 |
| orthologID_008341 | PH01001842G0020 | LOC_Os07g33330.1 | Bradi1g26210.1 | HORVU2Hr1G046380.3 | TRIUR3_29253 | Seita.2G332400.1 | Sobic.002G318600.1 | Zm00001d021633_T006 |
| orthologID_008399 | PH01005525G0020 | LOC_Os07g12160.1 | Bradi1g53910.1 | HORVU4Hr1G086340.3 | TRIUR3_22490 | Seita.2G083800.1 | Sobic.002G078000.1 | Zm00001d019184_T001 |
| orthologID_007342 | PH01000544G0310 | LOC_Os09g29120.1 | Bradi4g32570.2 | HORVU5Hr1G069750.7 | TRIUR3_26481 | Seita.2G236200.1 | Sobic.002G231000.1 | Zm00001d020773_T002 |
| orthologID_008440 | PH01003497G0070 | LOC_Os07g37760.1 | Bradi1g24590.1 | HORVU2Hr1G040000.8 | TRIUR3_26204 | Seita.2G354600.1 | Sobic.002G341000.1 | Zm00001d006606_T001 |
| orthologID_007847 | PH01000363G0710 | LOC_Os07g37960.1 | Bradi1g24360.1 | HORVU2Hr1G039700.1 | TRIUR3_20417 | Seita.2G356200.1 | Sobic.002G342300.1 | Zm00001d021821_T001 |
| orthologID_007334 | PH01006625G0050 | LOC_Os09g34870.1 | Bradi4g35376.2 | HORVU5Hr1G080030.2 | TRIUR3_09034 | Seita.2G278000.1 | Sobic.002G267600.1 | Zm00001d006154_T001 |
| orthologID_007025 | PH01000050G1280 | LOC_Os09g30410.2 | Bradi4g33377.4 | HORVU5Hr1G072120.4 | TRIUR3_09460 | Seita.2G248600.1 | Sobic.002G242600.1 | Zm00001d006002_T001 |
| orthologID_008699 | PH01250366G0010 | LOC_Os07g43370.1 | Bradi1g20780.1 | HORVU2Hr1G028980.1 | TRIUR3_34230 | Seita.2G398500.1 | Sobic.002G383400.1 | Zm00001d022203_T001 |
| orthologID_007168 | PH01005948G0040 | LOC_Os09g19910.1 | Bradi4g28850.1 | HORVU5Hr1G057560.1 | TRIUR3_07146 | Seita.2G186400.1 | Sobic.002G181600.2 | Zm00001d020236_T001 |
| orthologID_008739 | PH01000086G1500 | LOC_Os07g38620.1 | Bradi1g23900.1 | HORVU2Hr1G037680.2 | TRIUR3_33761 | Seita.2G363800.1 | Sobic.002G349100.1 | Zm00001d021881_T001 |
| orthologID_007458 | PH01000052G0290 | LOC_Os09g26670.1 | Bradi4g31140.1 | HORVU5Hr1G062450.1 | TRIUR3_26077 | Seita.2G216900.1 | Sobic.002G214300.1 | Zm00001d020610_T001 |
| orthologID_008584 | PH01002923G0150 | LOC_Os05g19270.1 | Bradi1g15260.1 | HORVU1Hr1G030930.12 | TRIUR3_27336 | Seita.3G295400.1 | Sobic.002G362200.1 | Zm00001d022028_T005 |
| orthologID_007440 | PH01000458G0440 | LOC_Os09g38784.1 | Bradi4g37996.1 | HORVU5Hr1G089770.2 | TRIUR3_07450 | Seita.2G308400.1 | Sobic.002G297100.1 | Zm00001d006343_T001 |
| orthologID_008004 | PH01004353G0020 | LOC_Os07g09970.1 | Bradi1g54553.1 | HORVU0Hr1G016430.1 | TRIUR3_32819 | Seita.2G075700.1 | Sobic.002G070200.1 | Zm00001d019100_T001 |
| orthologID_007443 | PH01001080G0510 | LOC_Os09g26260.1 | Bradi4g30930.1 | HORVU5Hr1G063340.3 | TRIUR3_30814 | Seita.2G213800.1 | Sobic.002G210100.1 | Zm00001d020578_T001 |
| orthologID_007650 | PH01001650G0080 | LOC_Os07g41600.1 | Bradi1g21880.1 | HORVU2Hr1G032280.1 | TRIUR3_35043 | Seita.2G384700.1 | Sobic.002G370100.1 | Zm00001d022102_T001 |
| orthologID_008735 | PH01000436G0430 | LOC_Os07g44744.1 | Bradi1g19850.1 | HORVU0Hr1G002730.5 | TRIUR3_03926 | Seita.2G409200.1 | Sobic.002G393500.1 | Zm00001d007032_T001 |
| orthologID_007413 | PH01002495G0170 | LOC_Os03g46750.1 | Bradi4g38200.1 | HORVU5Hr1G091740.8 | TRIUR3_17180 | Seita.2G130200.1 | Sobic.002G131400.1 | Zm00001d005370_T004 |
| orthologID_007437 | PH01001228G0110 | LOC_Os09g28100.1 | Bradi4g31990.1 | HORVU5Hr1G067910.3 | TRIUR3_05820 | Seita.2G227700.1 | Sobic.002G223700.1 | Zm00001d005848_T003 |
| orthologID_008535 | PH01000056G0870 | LOC_Os07g46480.1 | Bradi1g19070.1 | HORVU2Hr1G022820.1 | TRIUR3_16024 | Seita.2G418400.1 | Sobic.002G403000.1 | Zm00001d022390_T001 |
| orthologID_008034 | PH01000579G0090 | LOC_Os07g06980.1 | Bradi1g56740.4 | HORVU5Hr1G017550.9 | TRIUR3_17245 | Seita.2G042600.1 | Sobic.002G043100.3 | Zm00001d018864_T002 |
| orthologID_008131 | PH01000643G0410 | LOC_Os07g40730.1 | Bradi1g22600.1 | HORVU2Hr1G034200.5 | TRIUR3_16314 | Seita.2G378100.1 | Sobic.002G363700.1 | Zm00001d022046_T012 |
| orthologID_007468 | PH01000945G0120 | LOC_Os09g11240.1 | Bradi4g27260.1 | HORVU5Hr1G053580.1 | TRIUR3_26671 | Seita.2G161800.1 | Sobic.002G157200.1 | Zm00001d005503_T001 |
| orthologID_008126 | PH01000361G0860 | LOC_Os07g06450.1 | Bradi1g57190.1 | HORVU2Hr1G059380.2 | TRIUR3_07355 | Seita.2G036900.1 | Sobic.002G038300.1 | Zm00001d018821_T001 |
| orthologID_008345 | PH01001298G0380 | LOC_Os07g08950.1 | Bradi1g55030.1 | HORVU2Hr1G104450.6 | TRIUR3_03476 | Seita.2G062200.1 | Sobic.002G058500.1 | Zm00001d019040_T001 |
| orthologID_008579 | PH01005694G0030 | LOC_Os07g02710.1 | Bradi1g58920.1 | HORVU2Hr1G062850.3 | TRIUR3_31136 | Seita.2G011500.1 | Sobic.002G014800.2 | Zm00001d007961_T001 |
| orthologID_007550 | PH01001845G0140 | LOC_Os09g38060.2 | Bradi4g37380.2 | HORVU5Hr1G087990.2 | TRIUR3_26799 | Seita.2G301800.1 | Sobic.002G291400.1 | Zm00001d021419_T001 |
| orthologID_007136 | PH01003773G0100 | LOC_Os09g32940.1 | Bradi4g34656.1 | HORVU5Hr1G076330.27 | TRIUR3_25108 | Seita.2G266400.1 | Sobic.002G257750.1 | Zm00001d006090_T002 |
| orthologID_007463 | PH01001080G0330 | LOC_Os09g26480.1 | Bradi4g31050.1 | HORVU5Hr1G062850.1 | TRIUR3_28093 | Seita.2G216000.1 | Sobic.002G213200.1 | Zm00001d005799_T001 |
| orthologID_007614 | PH01002527G0220 | LOC_Os07g48920.1 | Bradi1g17080.3 | HORVU2Hr1G016040.2 | TRIUR3_26321 | Seita.2G440100.1 | Sobic.002G426100.1 | Zm00001d022554_T003 |
| orthologID_008330 | PH01000515G0140 | LOC_Os07g06880.1 | Bradi1g56870.2 | HORVU2Hr1G079270.1 | TRIUR3_04050 | Seita.2G041700.1 | Sobic.002G042300.1 | Zm00001d018857_T001 |
| orthologID_007087 | PH01003200G0210 | LOC_Os09g36760.1 | Bradi4g36477.3 | HORVU5Hr1G084030.4 | TRIUR3_15371 | Seita.2G290100.1 | Sobic.002G279400.1 | Zm00001d022626_T001 |
| orthologID_007135 | PH01001617G0380 | LOC_Os09g24200.1 | Bradi4g29870.1 | HORVU5Hr1G060920.5 | TRIUR3_14094 | Seita.2G201100.1 | Sobic.002G197100.1 | Zm00001d005732_T001 |
| orthologID_008446 | PH01001184G0160 | LOC_Os07g48570.1 | Bradi1g17410.6 | HORVU2Hr1G017290.1 | TRIUR3_32855 | Seita.2G436800.1 | Sobic.002G421900.1 | Zm00001d022525_T001 |
| orthologID_007260 | PH01002541G0040 | LOC_Os09g12500.1 | Bradi4g27460.4 | HORVU5Hr1G053980.5 | TRIUR3_15755 | Seita.2G165900.1 | Sobic.002G155000.1 | Zm00001d019989_T001 |
| orthologID_007298 | PH01000978G0420 | LOC_Os09g39474.1 | Bradi4g38520.2 | HORVU4Hr1G001130.1 | TRIUR3_05689 | Seita.2G133500.1 | Sobic.002G135000.1 | Zm00001d019866_T001 |
| orthologID_008397 | PH01000068G1270 | LOC_Os07g43900.1 | Bradi1g20390.1 | HORVU2Hr1G028230.14 | TRIUR3_16921 | Seita.2G402000.1 | Sobic.002G386800.1 | Zm00001d022239_T001 |
| orthologID_007970 | PH01001827G0140 | LOC_Os07g47150.2 | Bradi1g18420.1 | HORVU2Hr1G020730.4 | TRIUR3_34529 | Seita.2G423400.1 | Sobic.002G408600.1 | Zm00001d007108_T001 |
| orthologID_009682 | PH01000620G0360 | LOC_Os10g32300.1 | Bradi3g03210.1 | HORVU3Hr1G001960.10 | TRIUR3_00638 | Seita.2G308700.1 | Sobic.002G297400.1 | Zm00001d021368_T001 |
| orthologID_007146 | PH01003811G0100 | LOC_Os09g13940.2 | Bradi4g27850.2 | HORVU5Hr1G055190.3 | TRIUR3_17496 | Seita.2G172500.1 | Sobic.002G165700.1 | Zm00001d020043_T002 |
| orthologID_007500 | PH01000228G0990 | LOC_Os01g32130.1 | Bradi4g08840.1 | HORVU1Hr1G051410.1 | TRIUR3_26823 | Seita.6G156000.1 | Sobic.002G046300.1 | Zm00001d018894_T001 |
| orthologID_007287 | PH01000379G0690 | LOC_Os09g25650.1 | Bradi4g30665.1 | HORVU5Hr1G063950.3 | TRIUR3_04287 | Seita.2G210700.1 | Sobic.002G206900.3 | Zm00001d005781_T002 |
| orthologID_007754 | PH01000500G0090 | LOC_Os07g05380.1 | Bradi1g58297.1 | HORVU2Hr1G038850.7 | TRIUR3_28963 | Seita.2G030600.1 | Sobic.002G033200.1 | Zm00001d018793_T001 |
| orthologID_008731 | PH01000158G0110 | LOC_Os07g42300.1 | Bradi1g21550.2 | HORVU2Hr1G031450.6 | TRIUR3_30764 | Seita.2G386600.1 | Sobic.002G373800.1 | Zm00001d022134_T002 |
| orthologID_008402 | PH01000331G0190 | LOC_Os07g05480.1 | Bradi1g58350.1 | HORVU2Hr1G060480.1 | TRIUR3_21954 | Seita.2G031300.1 | Sobic.002G033900.1 | Zm00001d018797_T001 |
| orthologID_008544 | PH01001153G0030 | LOC_Os07g48610.1 | Bradi1g17380.1 | HORVU2Hr1G017080.2 | TRIUR3_22926 | Seita.2G437100.1 | Sobic.002G422200.2 | Zm00001d022529_T001 |

| | | | | | | | | |
|---|---|---|---|---|---|---|---|---|
| orthologID_007403 | PH01000177G0360 | LOC_Os09g24620.1 | Bradi4g30130.1 | HORVU5Hr1G066130.1 | TRIUR3_31710 | Seita.2G203100.1 | Sobic.002G199900.1 | Zm00001d020443_T001 |
| orthologID_007045 | PH01000453G0130 | LOC_Os09g23220.1 | Bradi4g29707.1 | HORVU5Hr1G060340.1 | TRIUR3_31267 | Seita.2G197200.1 | Sobic.002G194500.1 | Zm00001d020392_T002 |
| orthologID_007886 | PH01002479G0140 | LOC_Os07g18720.1 | Bradi1g52707.5 | HORVU2Hr1G105830.5 | TRIUR3_25337 | Seita.2G106400.1 | Sobic.002G109000.1 | Zm00001d019397_T006 |
| orthologID_007995 | PH01000436G0710 | LOC_Os07g44640.1 | Bradi1g19940.1 | HORVU2Hr1G026200.10 | TRIUR3_10876 | Seita.2G142600.1 | Sobic.002G145300.1 | Zm00001d019697_T001 |
| orthologID_008208 | PH01001546G0100 | LOC_Os07g08820.1 | Bradi1g55151.1 | HORVU2Hr1G112650.4 | TRIUR3_05767 | Seita.2G061400.1 | Sobic.002G057500.1 | Zm00001d018988_T002 |
| orthologID_007731 | PH01002968G0190 | LOC_Os07g40940.1 | Bradi1g22290.1 | HORVU2Hr1G033470.1 | TRIUR3_18941 | Seita.2G380300.1 | Sobic.002G366000.1 | Zm00001d022059_T003 |
| orthologID_008408 | PH01000360G0390 | LOC_Os07g42640.1 | Bradi1g21060.1 | HORVU2Hr1G030670.4 | TRIUR3_09861 | Seita.2G391400.1 | Sobic.002G376300.1 | Zm00001d022159_T001 |
| orthologID_007842 | PH01000936G0390 | LOC_Os07g46780.1 | Bradi1g18660.1 | HORVU2Hr1G021870.23 | TRIUR3_07123 | Seita.2G421700.1 | Sobic.002G406700.1 | Zm00001d022512_T001 |
| orthologID_007376 | PH01002616G0140 | LOC_Os09g38110.1 | Bradi4g37430.1 | HORVU5Hr1G087860.1 | TRIUR3_14883 | Seita.2G302400.1 | Sobic.002G291900.1 | Zm00001d021414_T001 |
| orthologID_007014 | PH01002328G0180 | LOC_Os09g21760.1 | Bradi4g29547.1 | HORVU5Hr1G059930.6 | TRIUR3_27164 | Seita.2G195400.1 | Sobic.002G192200.1 | Zm00001d020359_T003 |
| orthologID_007196 | PH01000314G0220 | LOC_Os09g35680.1 | Bradi4g35700.1 | HORVU5Hr1G080530.1 | TRIUR3_23573 | Seita.2G281600.1 | Sobic.002G270200.1 | Zm00001d021224_T001 |
| orthologID_007929 | PH01002295G0010 | LOC_Os07g34070.1 | Bradi1g25937.1 | HORVU2Hr1G046050.9 | TRIUR3_20291 | Seita.2G336300.1 | Sobic.002G322800.2 | Zm00001d021655_T001 |
| orthologID_007456 | PH01003477G0090 | LOC_Os09g29950.3 | Bradi4g32987.1 | HORVU5Hr1G070880.3 | TRIUR3_24225 | Seita.2G242300.1 | Sobic.002G236500.1 | Zm00001d020835_T027 |
| orthologID_007460 | PH01003361G0120 | LOC_Os09g10840.2 | Bradi4g27100.1 | HORVU5Hr1G053330.2 | TRIUR3_06066 | Seita.2G163200.1 | Sobic.002G153100.1 | Zm00001d005143_T001 |
| orthologID_007138 | PH01001027G0150 | LOC_Os09g28770.1 | Bradi4g32370.1 | HORVU5Hr1G069130.5 | TRIUR3_16341 | Seita.2G233900.1 | Sobic.002G228700.1 | Zm00001d005911_T001 |
| orthologID_009223 | PH01000591G0690 | LOC_Os02g06650.1 | Bradi3g04754.1 | HORVU6Hr1G026320.1 | TRIUR3_31113 | Seita.2G413600.1 | Sobic.002G398901.1 | Zm00001d022354_T001 |
| orthologID_007275 | PH01001127G0180 | LOC_Os09g27960.1 | Bradi4g31900.3 | HORVU5Hr1G067740.1 | TRIUR3_04010 | Seita.2G226500.1 | Sobic.002G222900.1 | Zm00001d005858_T001 |
| orthologID_007075 | PH01000618G0440 | LOC_Os03g31490.1 | Bradi4g38350.1 | HORVU5Hr1G092170.2 | TRIUR3_03209 | Seita.2G131000.1 | Sobic.002G132400.1 | Zm00001d005359_T001 |
| orthologID_007172 | PH01000897G0270 | LOC_Os03g19650.3 | Bradi4g26900.1 | HORVU4Hr1G024360.2 | TRIUR3_18942 | Seita.2G009700.1 | Sobic.002G013700.1 | Zm00001d018693_T001 |
| orthologID_007555 | PH01000070G1130 | LOC_Os09g32180.1 | Bradi4g34090.2 | HORVU5Hr1G074520.10 | TRIUR3_28052 | Seita.2G259400.1 | Sobic.002G252400.1 | Zm00001d020978_T005 |
| orthologID_007441 | PH01000379G0610 | LOC_Os09g25760.1 | Bradi4g30710.1 | HORVU5Hr1G064020.3 | TRIUR3_10720 | Seita.2G211100.1 | Sobic.002G207300.2 | Zm00001d020552_T001 |
| orthologID_007505 | PH01000177G0530 | LOC_Os09g24680.1 | Bradi4g30190.1 | HORVU5Hr1G066010.3 | TRIUR3_01902 | Seita.2G203900.1 | Sobic.002G200500.3 | Zm00001d020448_T001 |
| orthologID_007149 | PH01000177G0450 | LOC_Os09g24660.1 | Bradi4g30160.1 | HORVU5Hr1G066030.1 | TRIUR3_01899 | Seita.2G203500.1 | Sobic.002G200100.1 | Zm00001d020444_T003 |
| orthologID_008525 | PH01000620G0750 | LOC_Os07g28790.1 | Bradi1g28310.1 | HORVU2Hr1G101440.4 | TRIUR3_15467 | Seita.2G310100.1 | Sobic.002G298800.1 | Zm00001d021351_T003 |
| orthologID_007250 | PH01000379G1000 | LOC_Os09g25560.1 | Bradi4g30578.1 | HORVU5Hr1G064070.3 | TRIUR3_11905 | Seita.2G209500.1 | Sobic.002G206000.1 | Zm00001d020535_T001 |
| orthologID_007303 | PH01000476G0800 | LOC_Os09g10750.1 | Bradi4g27040.2 | HORVU5Hr1G053270.2 | TRIUR3_23461 | Seita.2G163700.1 | Sobic.002G152700.1 | Zm00001d019899_T001 |
| orthologID_007153 | PH01004105G0040 | LOC_Os09g20550.1 | Bradi4g29170.1 | HORVU5Hr1G058760.1 | TRIUR3_34825 | Seita.2G189500.1 | Sobic.002G185900.1 | Zm00001d020292_T001 |
| orthologID_007899 | PH01000656G0500 | LOC_Os07g35860.1 | Bradi1g25480.1 | HORVU2Hr1G044220.1 | TRIUR3_06305 | Seita.2G342500.1 | Sobic.002G329200.1 | Zm00001d021700_T001 |
| orthologID_007543 | PH01004434G0050 | LOC_Os09g13930.1 | Bradi4g27840.2 | HORVU5Hr1G055180.4 | TRIUR3_17497 | Seita.2G172300.1 | Sobic.002G165500.2 | Zm00001d005581_T001 |
| orthologID_006383 | PH01000085G0730 | LOC_Os07g23494.1 | Bradi2g27286.1 | HORVU7Hr1G021720.12 | TRIUR3_06171 | Seita.2G113400.1 | Sobic.002G116800.1 | Zm00001d005169_T001 |
| orthologID_007068 | PH01000084G0230 | LOC_Os09g32300.1 | Bradi4g34220.3 | HORVU5Hr1G075110.3 | TRIUR3_13060 | Seita.2G260600.1 | Sobic.002G253600.2 | Zm00001d006055_T002 |
| orthologID_007407 | PH01001001G0210 | LOC_Os09g21450.1 | Bradi4g29467.4 | HORVU5Hr1G059770.3 | TRIUR3_24490 | Seita.2G194200.1 | Sobic.002G190700.2 | Zm00001d020353_T001 |
| orthologID_007801 | PH01004948G0090 | LOC_Os07g30670.1 | Bradi1g27280.2 | HORVU7Hr1G023140.1 | TRIUR3_25345 | Seita.2G318000.1 | Sobic.002G305700.1 | Zm00001d021515_T001 |
| orthologID_007034 | PH01000646G0440 | LOC_Os09g19560.1 | Bradi4g28570.1 | HORVU5Hr1G056790.1 | TRIUR3_09095 | Seita.2G184000.1 | Sobic.002G179000.1 | Zm00001d020188_T001 |
| orthologID_008713 | PH01000549G0330 | LOC_Os07g02200.1 | Bradi1g59120.2 | HORVU2Hr1G063510.2 | TRIUR3_14278 | Seita.2G002000.1 | Sobic.002G007200.1 | Zm00001d018641_T001 |
| orthologID_007088 | PH01000770G0710 | LOC_Os09g31350.1 | Bradi4g33700.1 | HORVU5Hr1G073120.2 | TRIUR3_13762 | Seita.2G253400.1 | Sobic.002G246900.1 | Zm00001d006024_T001 |
| orthologID_007331 | PH01000131G0760 | LOC_Os09g34850.3 | Bradi4g35390.1 | HORVU5Hr1G080050.2 | TRIUR3_23595 | Seita.2G277800.1 | Sobic.002G267400.1 | Zm00001d021162_T001 |
| orthologID_007027 | PH01002445G0060 | LOC_Os05g22730.1 | Bradi4g07440.2 | HORVU6Hr1G093240.3 | TRIUR3_17213 | Seita.9G423600.1 | Sobic.002G033800.1 | Zm00001d018796_T001 |
| orthologID_007206 | PH01001005G0580 | LOC_Os09g36530.1 | Bradi4g36380.1 | HORVU5Hr1G083000.3 | TRIUR3_10729 | Seita.2G288800.1 | Sobic.002G278000.4 | Zm00001d021318_T003 |
| orthologID_007276 | PH01002495G0180 | LOC_Os03g46740.1 | Bradi4g38210.1 | HORVU5Hr1G091750.1 | TRIUR3_17181 | Seita.2G130300.1 | Sobic.002G131500.1 | Zm00001d019644_T003 |
| orthologID_007455 | PH01000191G0930 | LOC_Os09g34070.1 | Bradi4g35250.17 | HORVU5Hr1G079190.2 | TRIUR3_33068 | Seita.2G274200.1 | Sobic.002G264200.2 | Zm00001d021188_T006 |
| orthologID_007234 | PH01002627G0320 | LOC_Os09g32450.1 | Bradi4g34290.6 | HORVU5Hr1G075340.15 | TRIUR3_18494 | Seita.2G262400.1 | Sobic.002G254800.1 | Zm00001d021015_T002 |
| orthologID_008229 | PH01002586G0010 | LOC_Os07g34030.1 | Bradi1g25977.2 | HORVU2Hr1G046000.3 | TRIUR3_34438 | Seita.2G336000.1 | Sobic.002G322400.1 | Zm00001d006512_T002 |
| orthologID_007439 | PH01001005G0480 | LOC_Os09g36480.1 | Bradi4g36330.2 | HORVU5Hr1G082770.2 | TRIUR3_08891 | Seita.2G288400.1 | Sobic.002G277600.1 | Zm00001d006227_T003 |
| orthologID_007237 | PH01002593G0160 | LOC_Os09g26370.1 | Bradi4g30990.1 | HORVU5Hr1G063320.4 | TRIUR3_17606 | Seita.2G214700.1 | Sobic.002G211000.1 | Zm00001d020589_T001 |
| orthologID_008082 | PH01005214G0090 | LOC_Os07g32900.1 | Bradi1g26330.1 | HORVU2Hr1G046710.2 | TRIUR3_24254 | Seita.2G330900.1 | Sobic.002G317100.2 | Zm00001d006478_T001 |
| orthologID_007070 | PH01000052G0510 | LOC_Os09g26760.1 | Bradi4g31227.1 | HORVU5Hr1G062320.2 | TRIUR3_18690 | Seita.2G217400.1 | Sobic.002G214700.1 | Zm00001d005812_T002 |
| orthologID_008515 | PH01000811G0370 | LOC_Os07g48810.1 | Bradi1g17220.1 | HORVU2Hr1G016510.5 | TRIUR3_19936 | Seita.2G438800.1 | Sobic.002G424100.1 | Zm00001d022544_T002 |
| orthologID_007699 | PH01000541G0700 | LOC_Os07g12150.1 | Bradi1g53880.1 | HORVU7Hr1G037330.1 | TRIUR3_21337 | Seita.2G084000.1 | Sobic.002G078300.1 | Zm00001d019186_T001 |
| orthologID_007339 | PH01003361G0070 | LOC_Os09g10850.1 | Bradi4g27110.3 | HORVU5Hr1G053350.1 | TRIUR3_06065 | Seita.2G163100.1 | Sobic.002G153400.1 | Zm00001d005140_T003 |

| | | | | | | | | |
|---|---|---|---|---|---|---|---|---|
| orthologID_007449 | PH01000507G0280 | LOC_Os09g29380.1 | Bradi4g32670.2 | HORVU5Hr1G070010.1 | TRIUR3_15337 | Seita.2G237800.1 | Sobic.002G233100.1 | Zm00001d005944_T001 |
| orthologID_008190 | PH01001327G0100 | LOC_Os07g33850.2 | Bradi1g26020.1 | HORVU2Hr1G045810.4 | TRIUR3_25082 | Seita.2G335500.1 | Sobic.002G321500.1 | Zm00001d006509_T001 |
| orthologID_008468 | PH01000064G1490 | LOC_Os07g43460.1 | Bradi1g20610.1 | HORVU2Hr1G028610.1 | TRIUR3_07543 | Seita.2G399500.1 | Sobic.002G383900.1 | Zm00001d006920_T001 |
| orthologID_007871 | PH01000056G0980 | LOC_Os07g46555.1 | Bradi1g18980.1 | HORVU2Hr1G022660.2 | TRIUR3_14698 | Seita.2G419300.1 | Sobic.002G404200.1 | Zm00001d022400_T003 |
| orthologID_006703 | PH01001117G0100 | LOC_Os01g14610.2 | Bradi2g62300.1 | HORVU1Hr1G084860.2 | TRIUR3_15078 | Seita.2G443300.1 | Sobic.002G430200.2 | Zm00001d022585_T001 |
| orthologID_008305 | PH01000751G0750 | LOC_Os07g12890.1 | Bradi1g53680.1 | HORVU2Hr1G025400.1 | TRIUR3_14343 | Seita.2G090600.1 | Sobic.002G083200.2 | Zm00001d019228_T001 |
| orthologID_007371 | PH01000653G0110 | LOC_Os09g38010.2 | Bradi4g37327.1 | HORVU5Hr1G087850.1 | TRIUR3_13806 | Seita.2G301300.1 | Sobic.002G290900.1 | Zm00001d021423_T001 |
| orthologID_008655 | PH01001767G0340 | LOC_Os07g07620.1 | Bradi1g56040.1 | HORVU2Hr1G053030.5 | TRIUR3_31171 | Seita.2G047600.1 | Sobic.002G048200.1 | Zm00001d007838_T001 |
| orthologID_008240 | PH01001703G0350 | LOC_Os07g09370.1 | Bradi1g54840.1 | HORVU5Hr1G113990.21 | TRIUR3_15565 | Seita.2G068700.1 | Sobic.002G065200.1 | Zm00001d007774_T001 |
| orthologID_008013 | PH01000068G0730 | LOC_Os07g44299.1 | Bradi1g20150.1 | HORVU2Hr1G027150.1 | TRIUR3_10442 | Seita.2G405600.1 | Sobic.002G390200.2 | Zm00001d006943_T001 |
| orthologID_007116 | PH01000149G0690 | LOC_Os09g28510.1 | Bradi4g32190.1 | HORVU5Hr1G068520.1 | TRIUR3_31811 | Seita.2G231100.1 | Sobic.002G226400.1 | Zm00001d005895_T001 |
| orthologID_007868 | PH01000553G0220 | LOC_Os07g31270.1 | Bradi1g27060.1 | HORVU2Hr1G050450.3 | TRIUR3_31170 | Seita.2G322500.1 | Sobic.002G309900.2 | Zm00001d021558_T001 |
| orthologID_007753 | PH01002764G0030 | LOC_Os07g08180.2 | Bradi1g55540.3 | HORVU2Hr1G057760.1 | TRIUR3_30573 | Seita.2G054100.1 | Sobic.002G052400.1 | Zm00001d018943_T003 |
| orthologID_007406 | PH01001159G0380 | LOC_Os09g29750.1 | Bradi4g32870.1 | HORVU5Hr1G070450.1 | TRIUR3_24325 | Seita.2G240800.1 | Sobic.002G234600.1 | Zm00001d005958_T002 |
| orthologID_007094 | PH01000228G0310 | LOC_Os01g32280.1 | Bradi4g08120.1 | HORVU5Hr1G029490.2 | TRIUR3_26477 | Seita.3G233700.1 | Sobic.002G333100.1 | Zm00001d006567_T002 |
| orthologID_007515 | PH01000060G1580 | LOC_Os09g36100.2 | Bradi4g35990.5 | HORVU5Hr1G081380.3 | TRIUR3_32945 | Seita.2G284600.1 | Sobic.002G273900.1 | Zm00001d021282_T001 |
| orthologID_007018 | PH01000778G0140 | LOC_Os09g19920.1 | Bradi4g28860.1 | HORVU5Hr1G057570.2 | TRIUR3_07145 | Seita.2G186500.1 | Sobic.002G181700.2 | Zm00001d020237_T001 |
| orthologID_007998 | PH01001153G0250 | LOC_Os07g48450.1 | Bradi1g17480.1 | HORVU2Hr1G017470.1 | TRIUR3_32458 | Seita.2G436000.1 | Sobic.002G420700.1 | Zm00001d022424_T001 |
| orthologID_008042 | PH01001810G0250 | LOC_Os07g06800.1 | Bradi1g56930.3 | HORVU1Hr1G013970.1 | TRIUR3_11683 | Seita.2G039300.1 | Sobic.002G041000.1 | Zm00001d018847_T001 |
| orthologID_007604 | PH01001298G0080 | LOC_Os07g08810.1 | Bradi1g55160.1 | HORVU2Hr1G057400.17 | TRIUR3_08380 | Seita.2G061000.1 | Sobic.002G057400.1 | Zm00001d018987_T001 |
| orthologID_008103 | PH01000068G1090 | LOC_Os07g44040.1 | Bradi1g20300.1 | HORVU2Hr1G027660.3 | TRIUR3_10944 | Seita.2G403500.1 | Sobic.002G388100.1 | Zm00001d022251_T002 |
| orthologID_007286 | PH01000070G1810 | LOC_Os09g32330.1 | Bradi4g34240.1 | HORVU5Hr1G075200.2 | TRIUR3_14133 | Seita.2G261500.1 | Sobic.002G254200.1 | Zm00001d006059_T001 |
| orthologID_007850 | PH01000364G0880 | LOC_Os07g47650.1 | Bradi1g18040.4 | HORVU2Hr1G019280.4 | TRIUR3_04142 | Seita.2G429100.1 | Sobic.002G414500.1 | Zm00001d007152_T001 |
| orthologID_008426 | PH01005776G0060 | LOC_Os07g34020.1 | Bradi1g25990.1 | HORVU2Hr1G045970.3 | TRIUR3_32218 | Seita.2G335900.1 | Sobic.002G322300.1 | Zm00001d021654_T001 |
| orthologID_008169 | PH01003569G0100 | LOC_Os07g14100.1 | Bradi1g53482.1 | HORVU5Hr1G117070.1 | TRIUR3_26955 | Seita.2G095100.1 | Sobic.002G088800.1 | Zm00001d019269_T001 |
| orthologID_008500 | PH01003073G0030 | LOC_Os07g31660.1 | Bradi1g26853.1 | HORVU2Hr1G050580.1 | TRIUR3_28767 | Seita.2G319900.1 | Sobic.002G307700.1 | Zm00001d006416_T001 |
| orthologID_008581 | PH01000361G0620 | LOC_Os07g05870.1 | Bradi1g58500.1 | HORVU2Hr1G111150.1 | TRIUR3_10317 | Seita.2G034600.1 | Sobic.002G036300.1 | Zm00001d018812_T001 |
| orthologID_007530 | PH01005345G0020 | LOC_Os09g29460.1 | Bradi4g32737.1 | HORVU5Hr1G070260.1 | TRIUR3_31014 | Seita.2G236800.1 | Sobic.002G233800.1 | Zm00001d020804_T001 |
| orthologID_007192 | PH01002111G0170 | LOC_Os09g20900.1 | Bradi4g29300.1 | HORVU5Hr1G059050.1 | TRIUR3_04910 | Seita.2G191900.1 | Sobic.002G187500.1 | Zm00001d020311_T001 |
| orthologID_007706 | PH01000068G1390 | LOC_Os07g43720.1 | Bradi1g20490.1 | HORVU2Hr1G028370.1 | TRIUR3_23717 | Seita.2G401200.1 | Sobic.002G385900.1 | Zm00001d022231_T001 |
| orthologID_007819 | PH01002701G0030 | LOC_Os07g28400.1 | Bradi1g28410.1 | HORVU5Hr1G078960.1 | TRIUR3_04068 | Seita.2G309200.1 | Sobic.002G297900.1 | Zm00001d021360_T001 |
| orthologID_007127 | PH01000405G0880 | LOC_Os12g14080.1 | Bradi4g39460.2 | HORVU6Hr1G020920.1 | TRIUR3_32140 | Seita.3G086300.1 | Sobic.002G387300.1 | Zm00001d022244_T002 |
| orthologID_007473 | PH01001849G0230 | LOC_Os12g21890.1 | Bradi4g08080.1 | HORVU5Hr1G031300.2 | TRIUR3_04058 | Seita.2G322700.1 | Sobic.002G008800.1 | Zm00001d007942_T002 |
| orthologID_008079 | PH01000538G0870 | LOC_Os07g39720.1 | Bradi1g23150.1 | HORVU2Hr1G035800.1 | TRIUR3_23478 | Seita.2G370900.1 | Sobic.002G356500.1 | Zm00001d021959_T001 |
